# Point-Process Modelling of Cell-Type Interaction Structure in CosMx Glioma Data for Hypothesis-Driven Tumor Inference from scRNA-seq

**DOI:** 10.64898/2026.10.02.756019

**Authors:** Felipe Pimenta Carcanholo, Murilo Henrique Anzolini Cassiano, Enzo Marques Panepucci, Márcio Poletti Laurini, Tathiane Maistro Malta Pereira

## Abstract

Glioblastoma (GBM) exhibits marked spatial heterogeneity that is lost after tissue dissociation for single-cell RNA sequencing. Here, we developed a spatial-statistical framework to characterize tumor–microenvironment organization and derive molecularly predictable spatial phenotypes from CosMx data. Eight GBM specimens comprising 2,427,362 cells, 15 cell populations, and four malignant states were analyzed, revealing substantial inter-patient differences in cellular composition and density. Fine finite-element meshes best preserved continuous spatial intensity and supported whole-tissue Log-Gaussian Cox Process modelling. Immune populations, including macrophages, microglia, monocytes, cDCs, neutrophils, and T cells, showed spatial attraction toward malignant states. These models generated cell-level measures of conditional intensity and tumor-proximity probability. Transcript abundances and metabolic pathway scores served as the sole predictive features to model these spatial phenotypes. Transcriptomic representations outperformed metabolic pathway scores. Leave-one-patient-out validation reduced predictive performance, and patient-wise harmonization failed to restore cross-patient transportability. Spatial outcome distributions varied between specimens, indicating that molecular–spatial relationships are strongly conditioned by patient-specific tumor architecture. Predicted interaction scores stratified transcriptional subclusters within cell types in external non-spatial single-cell datasets. This framework links spatial point-process phenotypes to molecular states and provides a basis for inferring spatial context from non-spatial single-cell data. Generalizing these models requires larger and more diverse spatial cohorts.

## Introduction

Gliomas are brain tumors with histological characteristics of glial cells, known for their diverse features and generally poor therapeutic response. They are classified based on their molecular characteristics and histological degree of malignancy. Malignant gliomas are the most frequent and lethal cancers originating in the central nervous system, with glioblastoma (GBM) being the most aggressive subtype ^1^.

The spatial organisation of tumour and immune populations is increasingly recognised as a determinant of anti-tumour immunity and is strongly associated with patient outcome ^2^. In glioblastoma, tumour complexity resides not only in cellular diversity but also in its spatial organization ^3^. The GBM tumor microenvironment (TME) is dynamic and structured into distinct niches, such as perivascular, invasive, and hypoxic sub-regions, where different cell types adapt and interact ^3^. One of the primary drivers of this spatial organization is tissue hypoxia ^4^. Hypoxic niches in GBM are associated with extensive metabolic reprogramming, rapid tumor evolution, and worse patient prognosis, being preferentially populated by mesenchymal-like (MES-like) tumor cells ^4^. Interactions between immune and tumor cells have been shown to be necessary for the formation of hypoxic zones, and their effects extend beyond visible necrotic areas, acting as a global organizing factor of the tumor ^4,5^.

The dynamic interactions between immune and tumor cells within these specialized niches are associated with tumor progression. However, the precise physical context of these interactions, including how proximity to the tumor influences cellular metabolic adaptation and immune evasion, cannot be preserved when tissues are dissociated for single-cell profiling.

Single-cell RNA sequencing (scRNA-seq) measures gene expression at the individual-cell level and has been increasingly used because of reduced costs, technological advances in next generation sequencing, and advances in computational methods ^6^. By profiling cells individually, scRNA-seq enables detailed characterisation of cellular heterogeneity and the measurement of thousands of transcripts across millions of cells in parallel, despite limitations such as data sparsity and low detection efficiency ^6^. The rapid evolution of scRNA-seq methods has led to manifold discoveries over a short time, including the identification of novel cell types and subtypes ^7–9^, the description of rare cell populations, revisions to established differentiation hierarchies, and the characterization of cell-state changes during development and in response to external stimuli ^6^. Therefore, the combination of molecular resolution and scalability makes scRNA-seq an efficient technique for the characterization of the cellular state and transcriptional program in complex tissues. Nevertheless, the molecular resolution comes at the cost of loss of spatial context as the tissues have to be disaggregated into a suspension of single cells or single nuclei, which hinders the identification of the location of the cells within the tissue.

Novel spatial transcriptomic methods address this limitation by retaining the relationship between spatial organisation and cellular states. Spatial “omics” constitute a range of technologies that allow molecular measurements with spatial resolution. Each technology involves a trade-off between the number of genes, spatial resolution, and detection sensitivity. These platforms can generically be divided into sequencing-based and imaging-based approaches. The former typically provide broader molecular coverage, including whole-transcriptome measurements, but at the expense of lower spatial resolution. The latter, on the other hand, provide higher spatial resolution, including single-molecule resolution, but typically target fewer genes ^10^.

One such example of spatial imaging technology is the CosMx Spatial Molecular Imager (NanoString). The technology uses multiplexed fluorescence in situ hybridization (FISH) and provides single-molecule level resolution ^11^. Currently, 6,000 simultaneous RNA targets can be quantified by CosMx technology in their spatial location along with negative probes and system controls for background noise and readout errors. Although the technology provides high sensitivity and cellular resolution, the process of image segmentation takes place independently in each field of view (FOV). This leads to technical artifacts like fractured or duplicated cells in the edges of the FOVs ^11^.

Despite the structural advantages of spatial transcriptomics, single-cell experiments provide a larger volume of publicly available data than spatial transcriptomics ^12^. This disparity creates an opportunity to use spatially resolved datasets to identify molecular characteristics associated with spatial phenotypes and to investigate whether these relationships can be used to infer spatial localisation from molecular profiles in the much larger body of existing non-spatial single-cell data.

Spatial proximity alone does not necessarily imply a specific cellular interaction. Two populations co-occur when they occupy the same anatomical or microenvironmental region. In these cases, an observed enrichment of one population around another can reflect shared spatial clustering rather than a direct association between cell types. In glioblastoma (GBM), tumor cells and immune populations organize into spatially heterogeneous niches. Consequently, analyses based on coarse cell counts, neighbourhood overlaps, or ad-hoc spatial strategies may identify spatial co-occurrence without distinguishing whether the observed association reflects interaction between cell populations or their shared localisation within the tissue ^2,13^.

Statistical approaches must account for the underlying spatial organization of the tissue. Spatial point-process methods represent individual cells as points in space, incorporate cell identities as marks, and test cellular relationships against explicit spatial null models ^14–16^. Spatial statistics have been applied to spatially resolved tissue data to characterise tissue architecture, cellular neighbourhoods, immune infiltration, and spatial relationships between cellular populations ^13,14,17^. Applying these methods in GBM maps the distinct microenvironmental niches organizing malignant and non-malignant populations ^5^.

The biological relevance of such spatial interactions is closely connected to cellular metabolism. The reprogramming of energy metabolism is a hallmark of cancer, driven by the high energy demands of tumor growth, which require significant adjustments to support uncontrolled proliferation ^18^. Tumor cells remodel multiple pathways, including the metabolism of amino acids, nucleotides, the urea cycle, lipids, iron, and central carbon metabolism (glycolysis, tricarboxylic acid (TCA) cycle, pentose phosphate pathway (PPP), and oxidative phosphorylation) ^19^. This metabolic flexibility allows cells to adapt to a constantly changing environment characterized by fluctuations in nutrient and oxygen availability.

This metabolic complexity is also observed in gliomas. Under physiological conditions, the brain is protected by the blood-brain barrier (BBB). This barrier prevents the entry of peripheral immune cells, protecting the cerebral tissue from inflammation ^20^. However, gliomas originating from glial cells responsible for BBB maintenance are capable of disrupting this structure, allowing the infiltration of immune cells and the formation of a unique tumor microenvironment (TME) ^21^. Immune cells are crucial components of the glioma TME; nevertheless, glioma cells and glioma stem-like cells (GSCs, brain tumor-initiating cells) interact with adaptive immune cells and recruited immunotolerant innate immune cells, inducing them to inhibit or evade anti-tumor responses ^22^. These interactions occur within an environment characterised by strong spatial and metabolic heterogeneity, in which local availability of oxygen and nutrients can influence the state and function of both tumour and immune populations.

Because these hypoxic niches are associated with metabolic reprogramming, the physical distance of a cell from the tumor core can be associated with changes in its phenotype, making the inference of spatial proximity biologically relevant. Thus, metabolic state represents a potential molecular correlate of the spatial context in which individual cells reside. If spatially defined phenotypes are associated with reproducible metabolic programs, these programs may provide information about spatial localisation even when the original spatial coordinates are unavailable. This raises the possibility that molecular profiles measured in spatially resolved datasets could be used to identify metabolic characteristics associated with specific spatial contexts and subsequently investigate whether these relationships are detectable in non-spatial single-cell datasets.

Here, we aim to quantify cell-type-specific spatial interactions while accounting for the underlying spatial organisation of the tissue, and to determine whether these spatial relationships are associated with cellular metabolic states. We further investigate whether spatial phenotypes related to tumour proximity can be predicted from metabolic profiles and whether these relationships can be extended to large, non-spatial single-cell datasets. By connecting spatial organisation with cellular metabolic state, this framework seeks to distinguish specific cell-type interactions from shared spatial organisation and to provide a basis for investigating spatially associated phenotypes in existing single-cell cohorts.

## Methods

### Reference spatial dataset

We used a publicly available CosMx 6k dataset of GBM samples from Migliozzi et al., 2025 ^23^. It comprises 8 specimens, 2,663 fields of view (FOVs), and a total of 2,750,202 cells (Table S1). The dataset had already undergone image segmentation and cell-centroid identification, and cell-type annotations were provided. In addition to the broad cell-type annotations, tumour cells were classified into four cellular states: GPM, MTC, NEU and, PPR ^23^. These cellular states represent biological traits distributed along two primary axes of tumor evolution, neurodevelopmental and metabolic. Along the neurodevelopmental axis, PPR stands for the Proliferative/Progenitor state, while NEU represents the Neuronal state. Along the metabolic axis, MTC denotes the Mitochondrial state, which is characterized by a reliance on oxidative phosphorylation for energy production. In contrast, GPM stands for the Glycolytic/Plurimetabolic state, a phenotype sustained by aerobic glycolysis combined with amino acid and lipid metabolism ^23,24^.

### Single Cell dataset

To assess the algorithm on single-cell data, we used an extensive pre-annotated glioblastoma single-cell RNA sequencing (scRNA-seq) dataset (GBmap). This harmonised collection comprises more than 1.1 million cells from 240 patients across 26 different datasets, with cells annotated into 18 major cell types ^25,26^. To obtain the same cellular populations across both assays, we standardised the annotations by transferring cell-type labels using a classification model built with the CellTypist package ^27,28^. The scRNA-seq expression matrix was also restricted to the ∼6,000 genes targeted by the CosMx spatial panel, so that the same set of features was available for the spatial predictive model when applied to the single-cell data.

### Metabolic score

To calculate the baseline metabolic activation score for each cell and pathway, we used single-sample Gene Set Enrichment Analysis (ssGSEA) ^29^. As a rank-based method, ssGSEA evaluates the expression profile of individual cells and measures the relative enrichment of genes belonging to a specific biological pathway ^29,30^. Gene sets were obtained from the KEGG database, comprising 85 metabolic pathways ^31,32^. Those pathways were collected using R packages KEGGREST and clusterProfiler to systematically query and retrieve reference pathways from the KEGG database that were stored on this package. To ensure a comprehensive and standardized representation of cellular metabolism without introducing manual selection bias, we filtered the retrieved data using KEGG internal class annotation, isolating pathways strictly categorized under Metabolism.

The CosMx SMI 6k platform measures approximately 6,000 targeted genes, compared with approximately 20,000 genes in whole-transcriptome spatial transcriptomic assays such as Visium Spatial Transcriptomics. CosMx data also contain a high proportion of zero-count values and provide single-molecule detection ^33^. We therefore used an alternative metabolic score based on gene detection. For each cell and pathway, the score was calculated as the percentage of genes in the pathway with at least one detected transcript. The pathway gene sets were restricted to genes present in the CosMx 6k panel before calculating the score.

### Space discretization

The eight complete CosMx specimens provided millions of individually resolved cells for spatial analysis. Direct specimen-scale point-process modeling demands heavy computation. Each cell centroid represents an observed point ui = (xi, yi) with cell-type annotation as its mark. For a pattern containing N cells, there are N(N -1) / 2 unique unordered point pairs potentially contributing to pairwise interaction statistics. Even trimming the pair count with spatial indexing and finite interaction ranges, the re-evaluated interaction terms at each Gibbs fitting step and absorbed high compute cost at CosMx scale.

We constructed a triangular finite-element mesh for each of the eight CosMx specimens using the fmesher package to provide a tractable numerical representation of the specimen geometry ^34^. The mesh differs from the observed point pattern, it represents the continuous spatial domain numerically, and the original cell centroids remain the observed point pattern used for subsequent inference. Mesh vertices act as computational locations connected by triangular elements and lack a one-to-one correspondence to biological cells. We incorporated the mesh into the specimen-scale spatial workflow and evaluated alternative point-process formulations containing a latent spatial field.

Constructing the mesh using every observed cell coordinate as an input location creates excess compute cost. We used a reduced set of spatial locations as mesh-construction seeds. For specimens containing more than 5,000 cells, we sampled 3,500 observed cell coordinates at random and obtained additional locations from a 25×25 spatial grid by calculating the median x- and y-coordinates of the cells within each occupied grid cell. Occupied grid cells alone contributed additional locations, keeping the realized number of seeds specimen-dependent and below the nominal target of 5,000. We used seeds to guide mesh construction, retaining the observed cells for subsequent analyses. This hybrid strategy combined direct sampling from the observed cellular distribution with broader spatial coverage of the specimen extent.

We calculated Gaussian kernel density estimation (KDE) surfaces from the complete set of cell coordinates and from the corresponding seed locations for each specimen to evaluate the spatial representation of the seeds. We estimated both KDE surfaces using a fixed bandwidth of σ = 800 coordinate units, enabled edge correction, and applied a 256 × 256 evaluation grid. The seed set contained fewer points than the complete cellular pattern. We multiplied the seed-derived KDE by the ratio between the number of observed cells and the number of seeds to scale both surfaces. Pearson correlation quantified the spatial similarity between the complete-cell and seed-derived KDE surfaces. This analysis assessed the retention of the broad spatial density structure of the complete cellular point pattern at the evaluated smoothing scale.

A non-convex hull represents the spatial boundary of each specimen, allowing the mesh to follow irregular geometry. We scaled mesh parameters according to the spatial diameter D of each specimen, defining D from the diagonal extent of its spatial coordinates. We defined a reference geometric scale as C = max(600, 0.006D). We set the non-convex hull parameters to C for the convex component and 1.6C for the concavity component. We generated three candidate mesh resolutions: coarse, medium, and fine. The coarse mesh used max.edge = (2.5C, 5C), cutoff = 0.30C, and offset = (0.60C, 2C); the medium mesh used max.edge = (1.4C, 3.5C), cutoff = 0.25C, and offset = (0.50C, 1.6C); and the fine mesh used max.edge = (0.9C, 2.2C), cutoff = 0.18C, and offset = (0.40C, 1.2C). We imposed a minimum triangle angle of 24 degrees for all resolutions to reduce the formation of narrow triangular elements. The max.edge parameter controls the allowed size of triangular elements, cutoff prevents a dense concentration of coincident vertices, and offset controls the numerical extension of the mesh beyond the main specimen boundary.

The extension beyond the observed boundary acts as a numerical buffer. Representing a continuous spatial field requires this buffer. Vertices located at the edge of a finite domain have support only on the observed side of the boundary. Extending the triangulation beyond the observed tissue reduces numerical boundary effects. These outer triangles constitute computational support and lack interpretation as observed tissue.

We used the same triangular representation to evaluate models incorporating a latent Gaussian spatial field through the stochastic partial differential equation (SPDE) approach. This formulation approximates a continuous Gaussian field ⍵(s) with a Matérn covariance structure using a Gaussian Markov random field defined at the mesh vertices. Piecewise-linear basis functions interpolate the latent field between vertices, representing a continuous spatial effect with a sparse precision matrix. This reduces computational requirements. We evaluated SPDE-based models as an alternative formulation to model residual spatial autocorrelation.

### Mesh fidelity assessment

The triangular mesh approximates a continuous domain. Selecting a mesh required testing coarse, medium, and fine resolutions through two evaluation checks. The evaluation assessed numerical behavior under Matérn fields at varying scales and measured retention of observed cell density via Kernel Density Estimation (KDE). The KDE analysis first processed all cells across the three resolutions, then evaluated individual cell populations for the chosen resolution. The Matérn fields operated with a unit marginal standard deviation (σ=1) and smoothness *α*=2. The spatial range ρ dictates the distance over which the latent field remains spatially correlated. Short ranges capture fine local variation. Broad ranges represent larger spatial trends. Diagnostic ranges of 416, 832, 1664, 2080, 2496, and 3328 coordinate units provided evaluation scales. These fixed diagnostic values remain separate from prior range values derived during model estimation.

Each combination of specimen, mesh resolution, and diagnostic range generated a Matérn precision matrix *Q*. Inverting *Q* yielded the covariance matrix *S* = *Q*^−1^ among vertex coefficients. A common evaluation lattice overlaid the tissue domain at 212.5-unit spacing. Matrix *A* defined the finite-element projection from vertices to lattice points. The diagonal of *AS A^T^*provided the standard deviation of the projected field. Departures from the target unit marginal standard deviation indicate numerical distortion from the finite-element representation.

The 95th percentile (P95) summarized each diagnostic, capturing the upper tail of error to resist dominance by extreme outliers. Local discretization error, P95(|sd.dev-1|), tracks local irregularity against the expected standard deviation to reflect finite-element discretization. Interpolation loss, |sd.bound-sd|/sd.bound, isolates discrepancy inside triangles. The sd.bound metric originates from linear interpolation of vertex-level standard deviations. The sd metric utilizes the full projected covariance. Boundary and vertex deviation, P95(|sd.bound-1|), measures deviation at the vertex level and reflects finite mesh geometry with boundary distortion. Total standard deviation deviation, P95(|sd-1|), tracks the overall departure from the nominal standard deviation.

Kernel Density Estimation (KDE) transforms discrete cell locations into a continuous density surface. Centering a Gaussian kernel on each observed cell and summing the spatial contributions built this surface. The bandwidth σ determines the smoothing scale. Narrow bandwidths preserve local variation. Broad bandwidths generate a smoother surface. This evaluation fixed the bandwidth at σ=800 coordinate units. The original coordinates of all observed cells generated a KDE surface for each specimen. Evaluating this surface at the vertices of coarse, medium, and fine meshes enabled surface reconstruction over the tissue via piecewise-linear interpolation through mesh triangles. The independent 212.5-unit lattice served as the comparison ground for direct and reconstructed surfaces. Keeping original coordinates, KDE parameters, and evaluation points uniform across resolutions ensures that fidelity differences reflect the mesh resolution. Pearson correlation scored the agreement.

The fine mesh proceeded to an identical evaluation across 15 annotated cell populations. All available cells within a population formed the KDE for each type and specimen, avoiding subsampling. Evaluating the KDE at every fine-mesh vertex allowed reconstruction via piecewise-linear interpolation. Linear concordance loss quantified the discrepancy between direct and reconstructed surfaces on the same independent lattice. Calculating KDE surfaces over a rectangular window padded 4σ beyond the mesh extent, with edge parameters disabled, isolated mesh-interpolation error from KDE boundary effects. This padding maintained defined KDE values at outer vertices and restricted all comparisons to evaluation points inside the observed tissue. The analysis measures numerical reconstruction of the point pattern.

### Log-Gaussian Cox Process Modelling

The spatstat package converted cellular subpopulations into spatial point patterns ^16^. The fmesher package discretized the tissue space into a finite triangular mesh bounded by the spatial polygon of the patient sample ^34^. The fmesher::fm_int function pre-calculated spatial integration points to prevent memory allocation redundancies. For Complete Spatial Randomness and Independence (CSRI) testing, Z-score standardization applied a zero mean and unit variance to the target tumor density raster. The INLA and inlabru packages executed approximate Bayesian inference ^35^. An Empirical Bayes integration strategy (int.strategy = “eb”) optimized computational memory over the pre-calculated integration points. For cellular proximity probability extraction, Z-score standardization applied a zero mean and unit standard deviation to the synthesized Niche Density raster matrix to ensure INLA optimization engine convergence. The Log-Gaussian Cox Process models utilized the established INLA and inlabru Empirical Bayes configuration.

### Molecular representations of cells and the matched-comparison design

Each cell that entered the interaction analysis carries two molecular descriptions derived from the same CosMx imaging run: (i) metabolic-pathway activity scores over 84 curated pathway feature columns, and (ii) transcript abundances over the 6,175-gene panel. The prediction question was posed as a matched comparison: the outcome rows, folds, model classes and metrics are held fixed while only the feature representation changes (metabolic scores vs transcript abundances). This design attributes any performance difference to the representation rather than to data or validation idiosyncrasies, which is the rationale for pooling all model comparisons into one evaluation harness The four interaction-target classes are approximately balanced across the pair rows (NEU 29.2%, MTC 28.1%, GPM 19.5%, PPR 23.1%), so no class-weighting correction was applied. The pathway feature matrix is dense, with zero missing entries across all 84 columns; transcript abundances are structurally sparse (about 13.0% of matrix entries non-zero), and undetected transcripts were kept as zeros rather than imputed, because the same zero semantics apply to all models.

### Outcomes: spatial-interaction measures as prediction targets

The upstream spatial-statistics stage assigns, for each cell and each interaction-target class in tumor = {GPM, PPR, MTC, NEU}, a probability of finding a partner within a 30-unit interaction radius (’probabilidade_raio30’) and a conditional interaction intensity (’intensidade_condicional’). These two quantities are the prediction outcomes; the model’s task is to infer them from molecular profile alone, i.e. without any spatial coordinate information. Probabilities were logit-transformed after clipping to [1e-3, 1-1e-3] (target ‘prob_logit’) so that a bounded variable is modelled on an unbounded scale suitable for squared-error regression. Intensity was modelled as log10 after restricting to finite values in (0, 1] (target ‘intensity_log10’); non-finite or non-positive entries are upstream sentinels and were excluded from intensity fits and evaluation rather than imputed.

### Pooled interaction model with interaction-identity encoding

Rather than separately fitting a model for each interaction target (four targets x 15 observed cell types), one model per target variable was fit on all cell-to-target rows jointly, appended by a one-hot encoding of the cell-type-by-target combination (56 observed combinations) to the feature block. This pooling lets statistical strength be borrowed across interaction classes, and allows a single model to rank interactions within a cell, which is the intended downstream use; the interaction identity is an input, so predictions remain interaction-specific.

### Algorithms

Candidate algorithms span three families to avoid favouring one model class: (i) gradient-boosted trees (XGBoost): 600 trees, learning rate 0.05, max depth 8, min child weight 10, row and column subsampling 0.8, histogram method, CUDA device, with training rows subsampled to 350,000 (fixed seed 7) for the transcriptomic arm; the metabolic arm used 900 trees with native categorical handling and no row subsampling; (ii) sparse linear baselines: ElasticNet with cross-validated penalty; for the transcriptomic arm l1-ratio 0.5, alpha grid log-scaled with 3 points between 1e-3 and 10, 3-fold internal CV, maximum 300 iterations (fit on 120,000 subsampled rows); for the metabolic arm l1-ratio in {0.1, 0.5, 0.9}, 10-point alpha grid, standardized features, 600,000 subsampled rows; (iii) supporting nonlinear learners: a random forest (300,000-row subsample) and a two-hidden-layer MLP (512-256 units, ReLU, dropout 0.1, CUDA), and a ridge regression on the combination one-hot alone as a pure-interaction-identity baseline. All regressions minimize squared error on the transformed targets.

### Within-cohort generalization and cross-patient transportability

We validated using two grouped cross-validation schemes in parallel, as they ask different questions about generalization:(1) ‘cellgroup5’: GroupKFold with five splits grouped by cell identity, so that any held-out fold has no cells present during training, but patients can be seen more than once; this tests generalization to novel cells from the same patients, avoiding same-cell leakage across the pair rows of a cell.

Two grouped cross-validation schemes were run in parallel because they answer two different deployment questions. (1) ‘cellgroup5’: GroupKFold with five splits grouped by cell identity, so every held-out fold contains only cells unseen during training while patients may recur; this asks whether models generalize to new cells of the same patients and prevents same-cell leakage across the pair rows of a cell. (2) ‘losò: leave-one-subject-out GroupKFold with eight splits grouped by sample identity (’um08’-’um15’), asking the stronger question of transportability to patients unseen at all. The analysis includes 7,572,841 cell-by-target rows from eight patients, four interaction-target classes and 15 cell types. Fold assignment is deterministic within a group, and all metrics are computed out-of-fold.

### Per-patient rank harmonization and centered-target arms

To test transportability, we built a harmonized transcriptomic matrix where, within each patient and gene, the detected log1p abundances have been replaced with their within-patient rank (0, 1], and undetected entries (i.e. zero abundance) remain at the lowest rank to preserve the sparsity pattern. Two additional loso arms were built using this harmonized matrix and the transcriptomic XGBoost configuration: H+T, trained on the raw ‘prob_logit’ target, and H+C, trained on targets with each patient’s mean subtracted to predict within-patient deviations instead of absolute levels.

### Cross-platform annotation transfer and scRNA-seq deployment

Cell-type labels were transferred from the CosMx reference to the GBmap scRNA-seq object (338,564 cells) using scArches plus SCANVI, aligning the two platforms on the shared 5,982 genes ^25,26,36,37^. To avoid biologically implausible mappings, non-neoplastic query cells were masked to the 11 microenvironment types (malignant-state posteriors zeroed), while B, plasma and mast cells (not found in the CosMx 15) were forced to unassigned; finally, any cell with a maximum posterior probability below 0.5 was also moved to unassigned. The deployment booster is the H+C XGBoost, retrained on all eight patients (500,000 rows, within-patient centered targets). Of the 193 genes not present in scRNA-seq, the deployment booster replaces their values in each cell with zeros (bottom rank). Features were rank-harmonized within each donor, and cells scored against all four target classes, with the four homotypic malignant combinations (never seen in training) encoded as zero vectors.

### Type-only reclustering and all-target subcluster analysis

The GBmap cells were first subset using the transferred type labels before being reclustered in the SCANVI latent space using 15 nearest neighbors and Leiden community detection with resolution 0.8 for the large types (GPM, Macrophage, PPR, Microglia and Tcells), 0.6 for Monocyte, Neutrophil and NEU, 0.5 for vasculature and 0.4 for cDC; random seed 0 followed by UMAP with seed 0. The fixed reclusters shown below are then used to score all four target classes, so that subcluster partners are always compared within the same partition.

Per-recluster means to use donor-clustered standard errors: the square root of the summed squared donor-level residual sum of squares divided by cell count. Single-donor reclusters are plotted without a generalizable interval and few-donor reclusters are flagged, along with a within-donor permutation null from 20 shuffles of the recluster labels within donor and the between donor variance share reported alongside eta-squared.

### Metrics and their interpretation

For each fold, coefficient of determination (R²) was computed on the transformed scale, and Spearman rank correlation was computed on pooled out-of-fold predictions per target. For the cross-validated mean of per-fold R², it can be strongly influenced by a single easy or hard fold, hence per-fold minima and maxima were retained with means, and are reported separately alongside the pooled R² (all out-of-fold rows scored jointly); the two aggregation methods answer slightly different questions (“typical fold” vs “typical held-out row”) and are not directly comparable. Additionally, a centered R² (’r2_centered’) was computed, with the mean of observed and of predicted values removed within each patient; this isolates within-patient pattern consistency, by removing any systematic offset that might arise due to calibration error. For all pooled metrics, sampling uncertainty was assessed with a 200-replicate percentile bootstrap. Under ‘cellgroup5’, rows were resampled directly, but with each replicate limited to 600,000 rows, thereby approximating the distribution of a statistic estimated on a subsampled dataset; this is conservative, since the actual uncertainty would be lower for a full-size resample. Under ‘losò, the resampling was done at the patient level (all rows of one patient placed into the sample together), with the concatenated resample then limited to 600,000 rows; because one very large patient could fill the cap, many replicates ended up containing only one or two patients, inflating the uncertainty beyond what an 8-patient cluster bootstrap would report.

### Sample size, prespecification and exploratory patient-level diagnostics

The prediction component is exploratory and was not prespecified; the patient cohort was fixed at data generation and no a priori sample-size calculation was done, hence patient-count limitations are addressed by patient-cluster bootstrap intervals rather than by power arguments. Post hoc, and explicitly exploratory, marginal distributions of both outcomes and of assay depth (detected genes and total counts per cell, read directly from the source expression matrix) were summarized per patient to test whether cross-patient failure was technical (depth/panel) or distributional (outcome base rates and composition).

### Software and reproducibility

Spatial-statistical analyses of the CosMx data were performed in R 4.4.3 within the Mamba environment named “r-spatstat” ^38,39^. Point-pattern representation, kernel-density estimation, spatial summary functions, and spatial randomization and simulation procedures used spatstat 3.6-0 and its component packages spatstat.geom 3.8-1, spatstat.explore 3.8-0, spatstat.model 3.7-0, and spatstat.random 3.4-5 ^16,40^. Data manipulation and tabular workflows used tidyverse 2.0.0 and data.table 1.18.4 ^41,42^. Vector spatial geometries were handled with sf 1.1-2, whereas raster-based spatial covariates and continuous spatial surfaces were handled with terra 1.9-34 ^43–45^. Finite-element mesh construction, non-convex tissue-boundary representation, spatial projection, and numerical mesh operations used fmesher 0.8.0 ^34^. Log-Gaussian Cox Process inference, latent Gaussian spatial fields, and stochastic partial differential equation (SPDE) formulations were implemented with INLA 26.8.7 and inlabru 2.15.0, with sparse-matrix operations supported by Matrix 1.7-5 ^35,46,47^. Spatial integration points for specimen-scale LGCP fitting were generated with fmesher::fm_int, and approximate Bayesian inference used the Empirical Bayes integration strategy implemented through INLA and inlabru. Figures generated within the spatial R workflow used ggplot2 4.0.3, patchwork 1.3.2, and RColorBrewer 1.1-3 ^48–50^.

**Table 1.** Spatial representativeness of mesh-construction seed locations across CosMx specimens. KDE surfaces from all observed cells and from the corresponding seed locations were calculated using σ = 800 coordinate units (edge = TRUE, dimyx = 256). Seed-derived KDEs were scaled by the cell-to-seed ratio and compared with the complete-cell KDEs using Pearson correlation. Higher values indicate greater similarity in spatial density patterns. n_cells and n_seeds denote the numbers of observed cells and mesh-construction seeds, respectively.

| Specimen | number of cells | number of seeds | pearson |
| --- | --- | --- | --- |
| um08 | 507809 | 3919 | 0.85 |
| um09 | 540864 | 3964 | 0.83 |
| um10 | 668041 | 3894 | 0.84 |
| um11 | 325967 | 3878 | 0.88 |
| um12 | 111244 | 3718 | 0.95 |
| um13 | 183838 | 3917 | 0.90 |
| um14 | 22045 | 3679 | 0.99 |
| um15 | 67554 | 3661 | 0.97 |

Metabolic-pathway analyses were performed separately in an R/Bioconductor environment. KEGG pathway definitions were queried and retrieved using KEGGREST 1.48.1 and clusterProfiler 4.16.0, after which pathways annotated under the KEGG Metabolism class were retained ^51–55^. Single-sample Gene Set Enrichment Analysis (ssGSEA) was computed using GSVA 2.2.0 ^29,56^. The same metabolic-analysis workflow was used for the Visium benchmarking experiment, in which ssGSEA scores derived from the complete transcriptome and from the CosMx-restricted gene set were compared across the 85 metabolic pathways. The alternative gene-detection score was calculated as the proportion of pathway genes with at least one detected transcript after restriction to the relevant assay gene set.

Machine-learning analyses were performed in Python 3.12 within the mamba environment pppred ^39,57^. Data handling, numerical computation, and intermediate columnar-data storage used pandas 3.0.5, NumPy 2.5.3, SciPy 1.18.1, and PyArrow 25.0.1 ^58–61^. Grouped cross-validation, including cell-grouped five-fold validation and leave-one-subject-out validation, as well as ElasticNet and ridge regression, random forests, preprocessing procedures, and regression metrics, were implemented with scikit-learn 1.9.1^62^. Gradient-boosted tree models, including the raw, rank-harmonized, and centered-target modelling arms, were implemented with XGBoost 3.4.1 ^63^. The multilayer perceptron model was implemented with PyTorch 2.11.0 ^64^. GPU-accelerated analyses were run on a single NVIDIA GeForce RTX 5060 Ti using NVIDIA driver 595.84 and a PyTorch CUDA 12.8 build.

Single-cell analyses of the GBmap dataset and the CosMx-to-scRNA-seq annotation-transfer workflow were performed in Python. Annotated expression matrices were represented with anndata 0.13.3, and neighborhood-graph construction, dimensionality reduction, UMAP visualization, and downstream single-cell analyses were performed with Scanpy 1.12.4 ^65,66^. Query-to-reference label transfer between CosMx and GBmap using the scArches/SCANVI framework was implemented with scvi-tools 1.5.1 using the 5,982 genes shared between the two platforms ^67^. Graph-based community detection of the transferred cell populations used leidenalg 0.12.0, with cell-type-specific reclustering performed in the SCANVI latent representation using the neighborhood and resolution parameters described above ^68^.

Random seeds were fixed explicitly for stochastic procedures where applicable, including mesh-seed sampling, machine-learning row subsampling, clustering, and dimensionality-reduction steps. Cross-validation assignments were deterministic within grouping units to prevent leakage between repeated cell-by-target rows. Complete software specifications for the r-spatstat environment, the separate R/Bioconductor environment used for metabolic-pathway analyses, and the pppred environment, together with R session information, Python package inventories, and analysis scripts, will accompany the analysis code to document the computational dependencies used to generate the reported results, tables, and figures.

## Results

### Spatial Heterogeneity in the Glioma Dataset

The analysed dataset comprised 8 GBM specimens, spanning 2,463 FOVs and 2,427,362 cells in total. The number of FOVs varied across specimens, with a median of 342 FOVs per specimen and a range of 61–548 ^23^. Cell numbers also varied substantially, with a median of 254,902 cells per specimen and a range of 22,045–668,041 cells (Table S1). Whole-tissue spatial maps showing the cellular composition of each specimen are shown in Figure 1A, while 1B shows the relative proportions of the 15 annotated cell populations across specimens. All 15 populations differed significantly in their relative representation between specimens (χ² test for homogeneity of proportions, cell type vs. all other cells, df = 7; BH-adjusted P < 1 × 10⁻¹⁶ for all populations; global χ² = 667,971, df = 98, P < 0.001; Table S2). Tumour cells were further characterised according to the four cellular states defined in the original dataset (PPR, NEU, MTC, and GPM) which represent neurodevelopmental and metabolic axes of tumour evolution ^23^. Figure 1C shows the relative proportions of these states within the tumour-cell compartment across specimens. All four tumour states also differed significantly in their relative representation between specimens (χ² test for homogeneity of proportions, state vs. all other tumour cells, df = 7; BH-adjusted P < 1 × 10⁻¹⁶; χ² = 70,791–204,621; Table S2).

**Figure 1.**
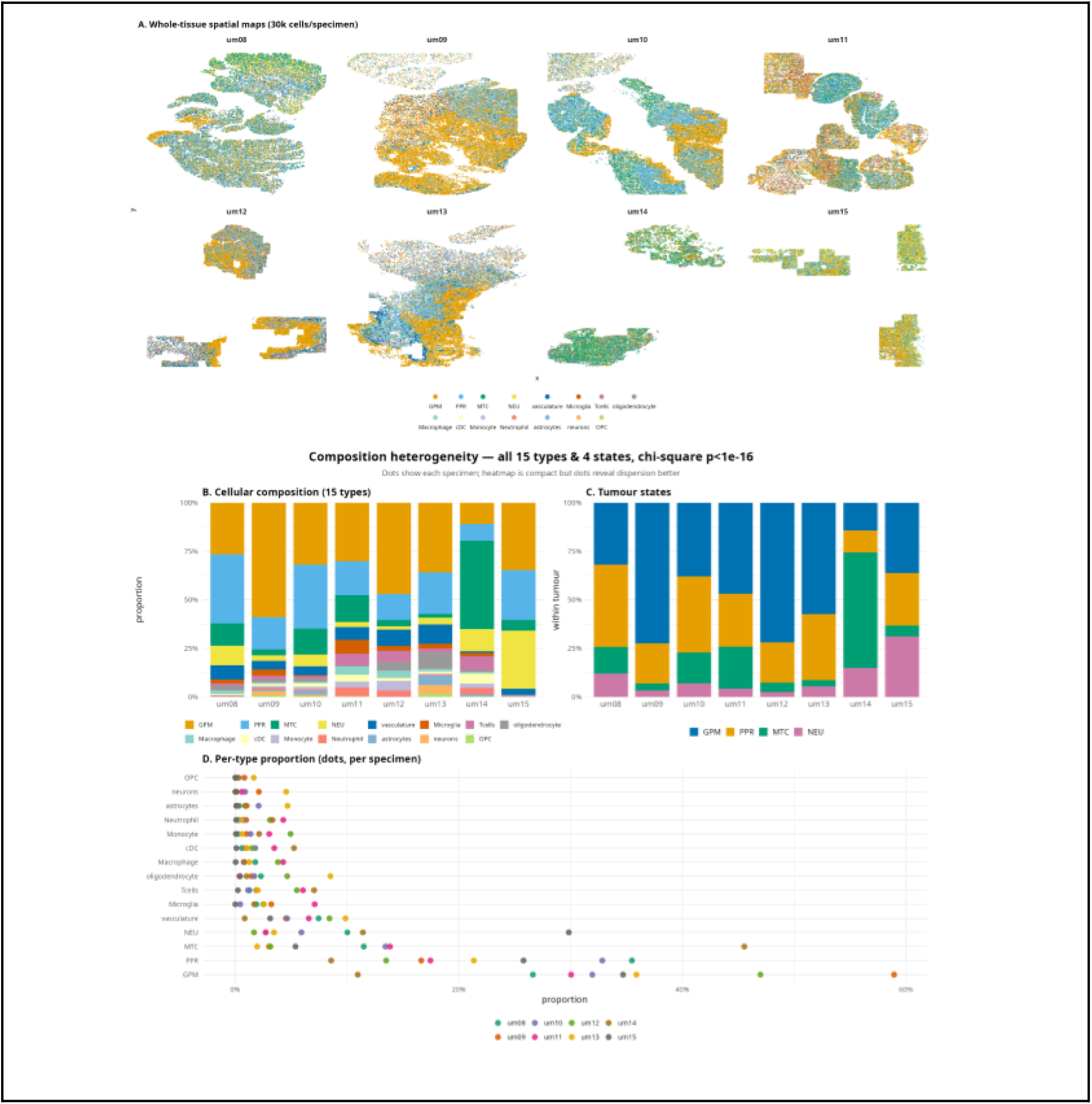
Dataset overview and cellular composition across GBM specimens. **(A)** Whole-tissue spatial maps of 8 GBM specimens (um08–um15), comprising 2,463 FOVs and 2,427,362 cells. Each point represents a cell coloured by cell type; 30,000 cells per specimen were randomly sampled for visualisation. **(B)** Relative proportions of the 15 annotated cell populations across specimens. **(C)** Relative proportions of the four tumour states (GPM, PPR, MTC, and NEU) within the tumour-cell compartment (1,904,720 cells). **(D)** Cell-type proportions across specimens. Each point represents one specimen. Differences were assessed using χ² tests with Benjamini–Hochberg correction; all 15 cell types and 4 tumour states showed significant differences (adjusted P < 1 × 10⁻¹⁶; Table S2).

We next examined cell density across the spatially profiled specimens. Overall cell density showed marked variation between specimens (Figure 2A), which was also observed when density was examined separately for each annotated cell population (Figure 2B). Differences between specimens were significant both overall (Kruskal–Wallis, H = 629, df = 7, P = 1.6 × 10⁻¹³¹) and for each of the 15 cell populations (H = 367–1,083, df = 7, BH-adjusted P < 1 × 10⁻⁷⁴ for all populations; Table S1, Table S3).

**Figure 2.**
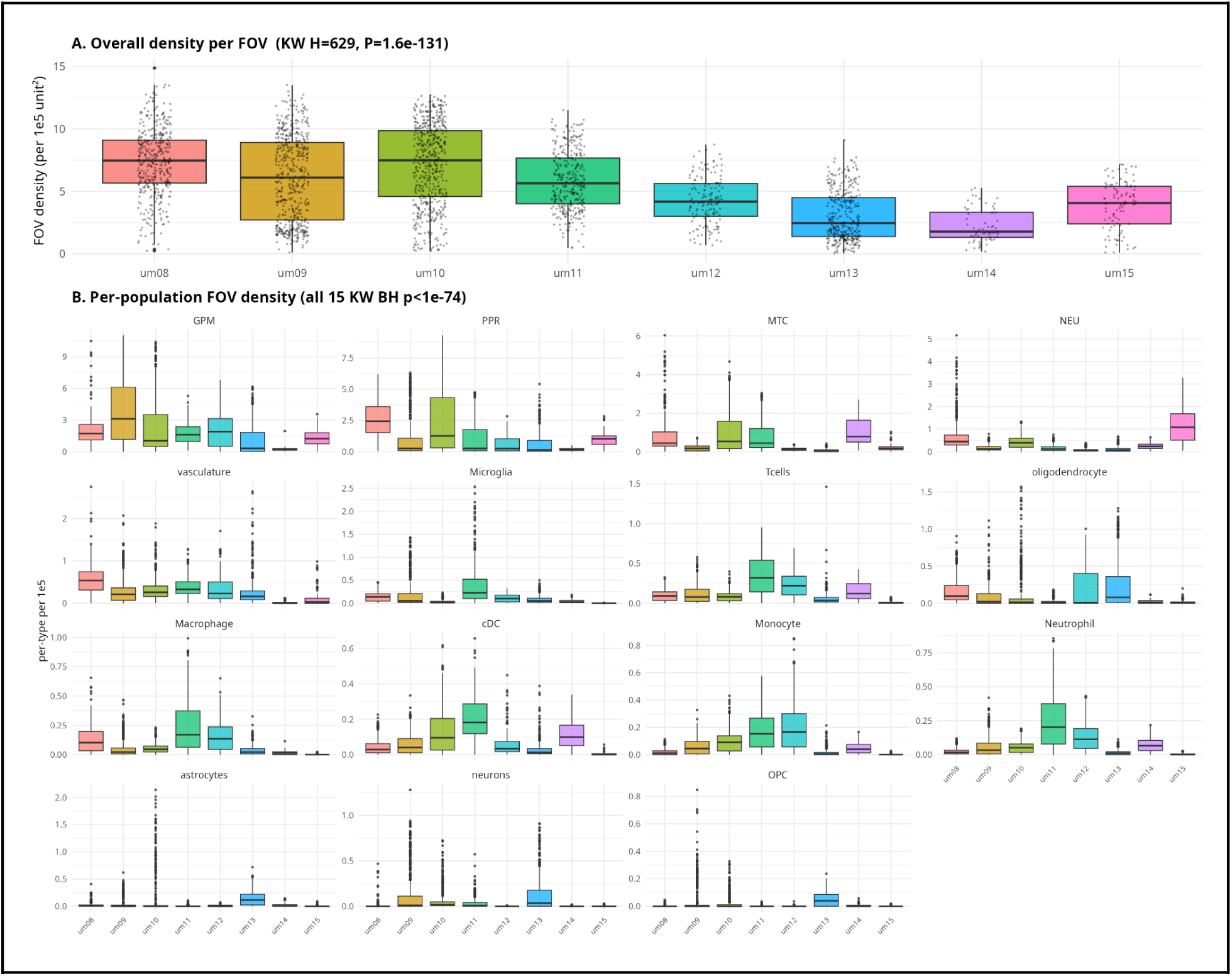
Cell density heterogeneity across FOVs. **(A)** Overall cell density per FOV, expressed per 10^5 unit². Four zero-area FOVs were excluded. Median FOV density ranged from 1.77 (um14) to 7.48 (um10) per 10^5 unit² (Kruskal–Wallis, H = 629, df = 7, P < 1 × 10−16; Tables S1, S3). **(B)** Density per FOV for each of the 15 cell populations, including zero-density FOVs. All populations differed significantly between specimens (Kruskal–Wallis, H = 367–1,083, df = 7, BH-adjusted P < 1 × 10−74; Table 3). FOV density reflects both cellular distribution and tissue occupancy (0.36–0.71); hull-corrected densities are shown in Table S1.

Together, these results demonstrate substantial heterogeneity in cellular density between specimens, highlighting the need to account for spatial background structure when evaluating cell–cell interactions.

### Fine Mesh Resolution Maximizes Numerical Fidelity for Spatial Field Reconstruction

We first evaluated if spatial locations selected as mesh-construction seeds represented the cellular density patterns across the CosMx specimens (Table 1). Each specimen yielded two KDE surfaces, one from all observed cell coordinates and one from the mesh-construction seed locations. Scaling the seed-derived KDE by the ratio of observed cells to seeds enabled comparison. Pearson correlation quantified spatial similarity between the surfaces. This analysis measured the fidelity of the reduced seed locations in reproducing the broad spatial distribution of the complete cellular pattern.

The mesh-construction seeds showed high correspondence with the complete cellular distributions across all eight specimens (Table 1). Pearson correlations ranged from 0.83 to 0.99, with a median of 0.89. Specimens um09 and um10 produced the lowest correlations (0.83 and 0.84). Correlations for um12, um14, and um15 exceeded 0.95. Specimen sizes ranged from 22,045 to 668,041 observed cells. The reduced seed sets maintained 3,700 to 4,000 locations and retained the spatial density structure of the complete patterns. These results confirmed the hybrid seed-selection strategy for subsequent mesh construction.

The three candidate meshes increased numerical resolution across all eight specimens (Figure 3A). Median mesh size grew from 7,625 vertices and 15,079 triangular elements in the coarse representation to 11,450 vertices and 22,669 triangles in the medium representation and 23,980 vertices and 47,563 triangles in the fine representation. Triangle-edge length decreased alongside this refinement. Median edge length dropped from 1,084.8 coordinate units in the coarse meshes to 901.6 in the medium and 623.0 in the fine meshes, a 42.6% reduction. The median 95th-percentile edge length decreased from 2,953.7 to 1,575.7 coordinate units. The three candidate meshes provided finer spatial discretizations alongside higher element counts.

**Figure 3.**
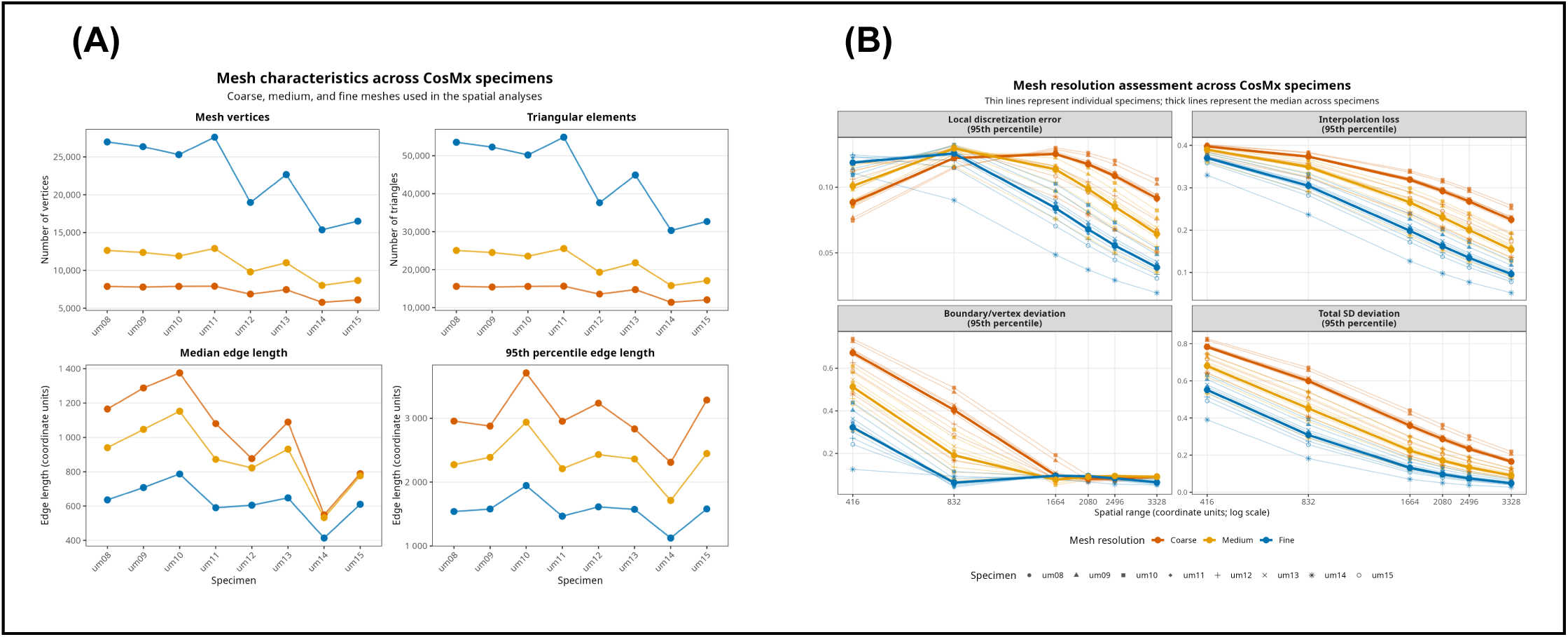
Geometric characterization and numerical assessment of candidate mesh resolutions. **(A)** Geometric properties (number of vertices, triangular elements, median, and 95th-percentile edge lengths) for coarse, medium, and fine triangular meshes across all eight CosMx specimens. **(B)** Numerical fidelity of the three mesh resolutions evaluated across six diagnostic spatial ranges (416 to 3328 coordinate units) using Matérn spatial fields. Panels display four 95th-percentile error diagnostics: local discretization error, interpolation loss, boundary/vertex deviation, and total standard-deviation deviation. Thin lines represent individual specimens, and thick lines indicate the median, with lower values signifying smaller numerical deviation and higher spatial fidelity.

Further analysis tested if geometric resolution changes improved the numerical representation of continuous spatial fields (Figure 3B). The fine mesh produced the lowest interpolation loss and total standard-deviation deviation across all eight specimens and diagnostic spatial scales. At the 416-unit range, for the coarse, medium, and fine meshes median interpolation loss measured 0.398, 0.390, and 0.370 and median total SD deviation measured 0.783, 0.681, and 0.552, respectively. At the 3328-unit range, interpolation loss dropped to 0.224, 0.154, and 0.096 and total SD deviation fell to 0.165, 0.091, and 0.049, respectively.

Local discretization error showed a stronger dependence on spatial scale. At the 416-unit and 832-unit ranges, increasing mesh resolution did not reduce this diagnostic. From 1664 units onward, the fine mesh produced lower local error. At 1664 units, median values decreased from 0.125 (coarse) to 0.114 (medium) and 0.084 (fine). At 3328 units, values dropped from 0.092 to 0.064 and 0.039.

Boundary/vertex deviation followed a different pattern. Mesh refinement reduced this component at short spatial ranges. At 832 units, the median deviation decreased from 0.404 in the coarse mesh to 0.191 in the medium mesh and 0.062 in the fine mesh. At spatial ranges of 1664 units and above, values converged across resolutions. The relative ordering varied, indicating this component stabilized without further mesh refinement.

Mesh refinement improved interpolation and total field reconstruction. The fine mesh provided the highest numerical fidelity across the evaluated range, even though the local-error diagnostic did not favor it at the shortest spatial scales and boundary-related differences diminished at broader scales.

#### Fine Mesh Reconstructions Preserve Spatial Intensity Across Cell Populations

KDE-based reconstruction analysis distinguished the three candidate mesh resolutions (Figure 4A). Estimating KDE surfaces from all observed cells revealed decreasing reconstruction loss as mesh resolution increased. The fine mesh produced the lowest 1−Pearson values across specimens. The medium mesh yielded intermediate reconstruction loss. The coarse mesh generated the largest deviations from the estimated KDE surface. These results established the fine mesh as the standard for subsequent spatial analyses.

**Figure 4.**
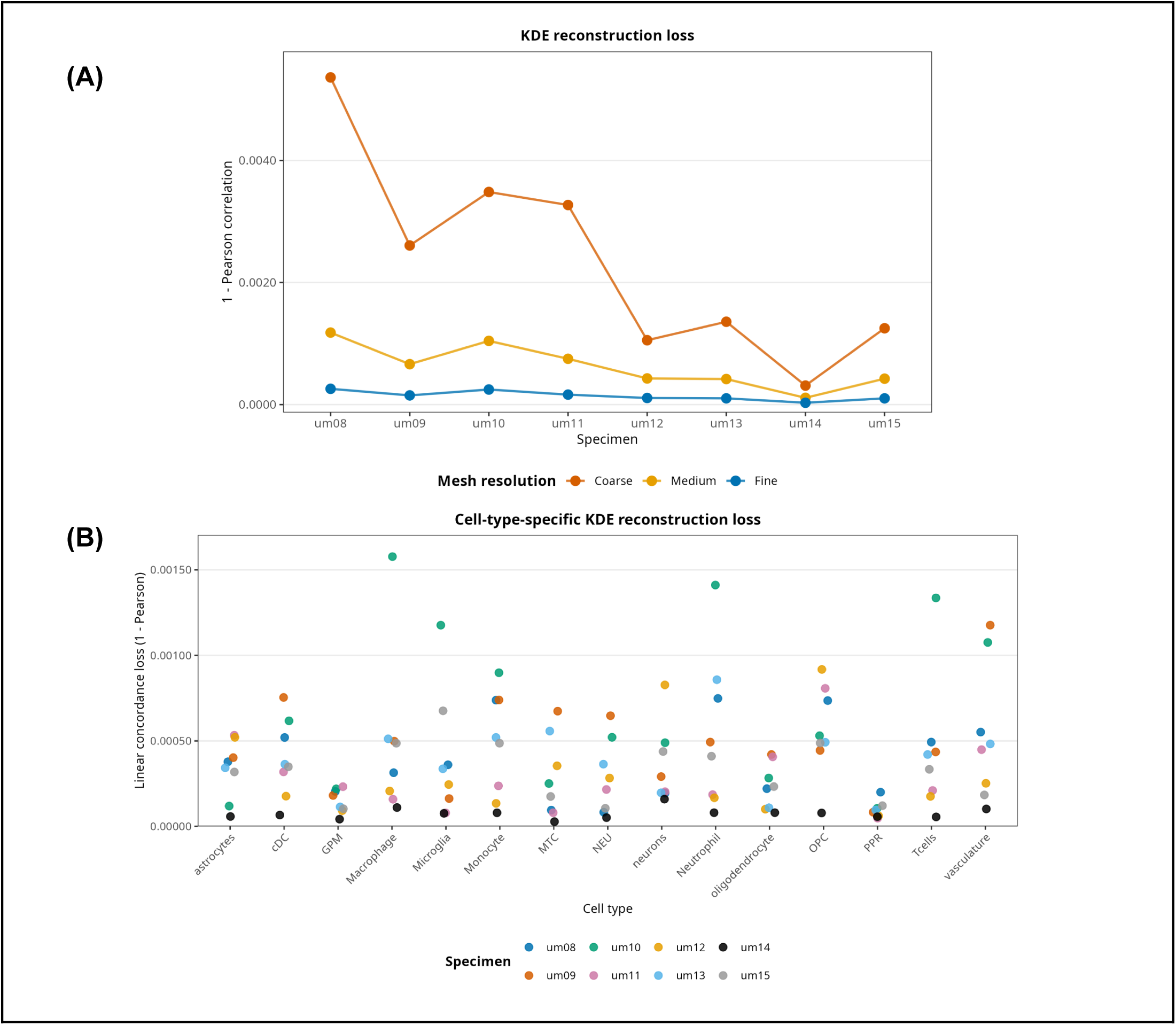
KDE-based assessment of spatial signal retention across mesh resolutions and cell populations. **(A)** Reconstruction loss (1−Pearson linear concordance loss) for Gaussian kernel density estimation (KDE) surfaces calculated from all observed cells and reconstructed using coarse, medium, and fine meshes. Lower values indicate greater agreement between the directly estimated and mesh-reconstructed surfaces. **(B)** Cell-type-specific reconstruction loss for the 15 annotated populations using the selected fine mesh. Points represent individual specimens, with lower 1−Pearson values indicating greater preservation of the original cell-type-specific spatial intensity pattern.

Analysis of the 15 annotated cell populations measured reconstruction fidelity using the fine mesh (Figure 4B). Cell-type-specific KDE surfaces produced low linear concordance loss across specimens, indicating agreement between the estimated spatial intensity surfaces and the finite-mesh reconstructions. Sparse populations exhibited higher variation in reconstruction loss compared to the all-cell analysis. Agreement remained high across the evaluated cell types.

Geometric, Matérn-based, and KDE assessments confirm the fine mesh provides the highest fidelity among the candidate resolutions. The fine mesh retains the spatial intensity structure of the complete cellular pattern and individual cell populations.

### Null Models for Multi-Scale Spatial Point Pattern Analysis

Modeling the relative intensity of cellular proximity to the tumor requires testing the null hypothesis of Complete Spatial Randomness and Independence (CSRI). Establishing CSRI as a baseline identifies biological interactions, such as structural clustering or cellular exclusion, within the spatial distribution of cells ^15^.

Computational simulations evaluated the spatial point patterns against two null models, random marks and random locations. Testing for random labeling or spatial independence requires comparing distinct cellular populations. The random labeling model fixed the spatial coordinates of all cells and reassigned their biological identities (cell-type labels or “marks”). This step determines the dependence of spatial relationships between neighboring points on specific cellular identities. The random spatial shifting model translated one cellular pattern relative to the other across the tissue area and preserved its internal spatial configuration. This evaluation compares the observed spatial association between two populations against expected spatial independence ^15^.

#### Local Spatial Organization of the Tumor Microenvironment

Three complementary approaches analyzed the spatial organization of the tumor compartment and its distinct malignant states (GPM, MTC, NEU, and PPR) relative to the microenvironment on a Field of View (FOV) basis. The analysis required a minimum of 500 cells per FOV to establish biological representation. The J-function evaluated the global spatial distribution of tumor cells against all other combined microenvironmental cells. Ripley’s cross-K function, paired with random labeling, assessed the bivariate spatial dependence between tumor cells and specific immune and stromal cell types. The K-cross function, combined with random spatial shifting, tested the structural independence of these pairwise interactions ^15^. Results represent the percentage of evaluable FOVs achieving statistical significance (p < 0.05) (Figures 5A–C; detailed FOV-level distributions are shown in Figure S1).

**Figure 5.**
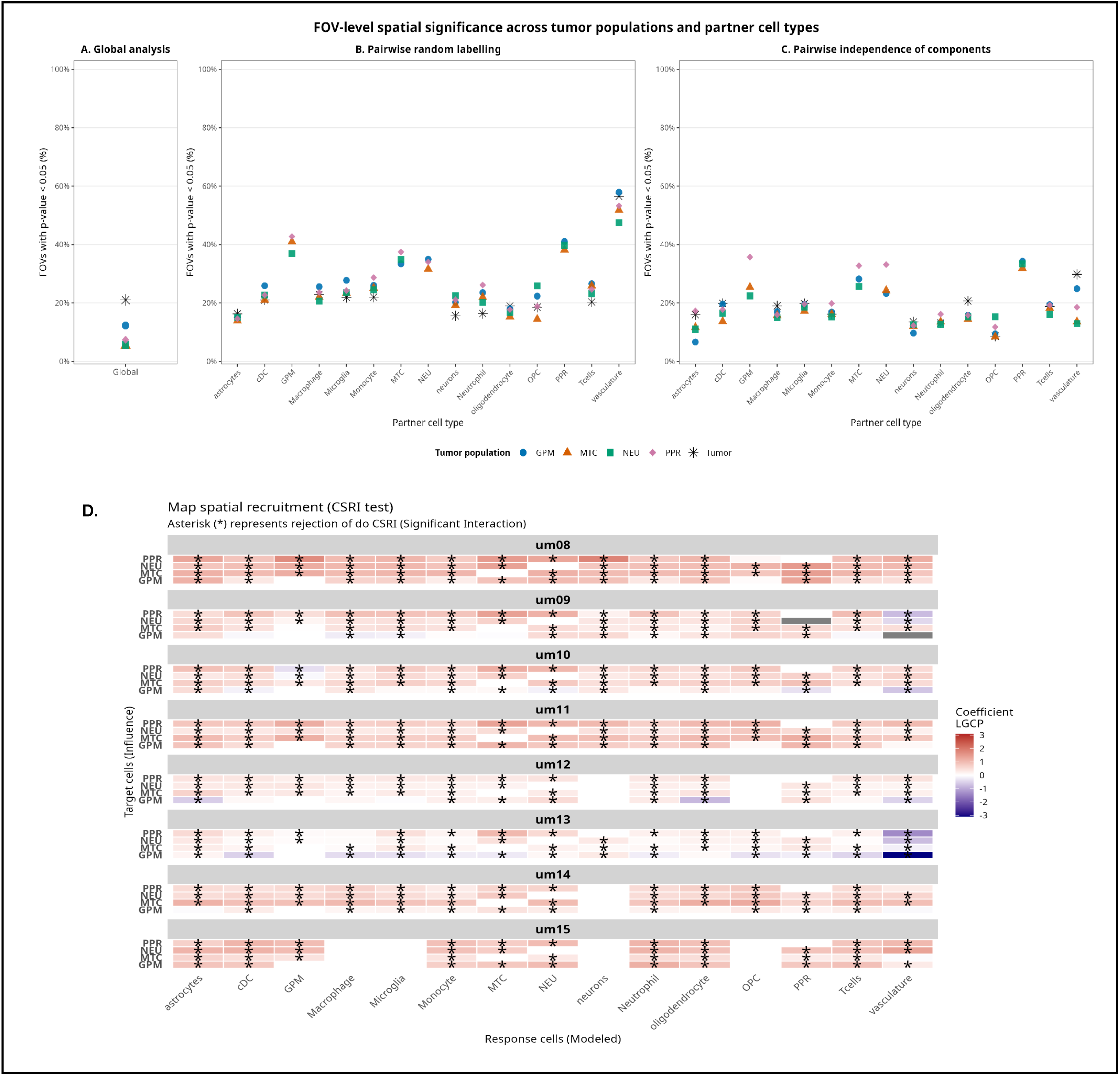
Spatial significance and interaction mapping of tumor populations within the microenvironment. **(A)** Global FOV-level spatial analysis for individual (GPM, MTC, NEU, PPR) and pooled tumor states, showing the percentage of evaluable FOVs with significant spatial dependence ($p < 0.05$). **(B-C)** Pairwise spatial analyses between focal tumor populations and annotated partner cell types evaluated against **(B)** random-labeling and **(C)** independence-of-components (random spatial shifting) null models. The y-axis displays the percentage of significant FOVs ($p < 0.05$) for each combination. Non-evaluable FOVs were excluded from the denominators in **A-C**. **(D)** Spatial recruitment heatmap displaying Log-Gaussian Cox Process (LGCP) coefficients across the patient cohort. Red indicates spatial attraction (positive coefficient) and blue represents spatial repulsion (negative coefficient) between tumor states (y-axis) and partner cell types (x-axis). Asterisks (*) denote significant rejection of the Complete Spatial Randomness and Independence (CSRI) null hypothesis; grey tiles indicate pairs with insufficient cell counts.

The global analysis revealed distinct spatial behaviors among tumor populations (Figure 5A). The pooled tumor compartment yielded the highest proportion of significant FOVs (21.0%, 373 of 1,773). Among individual malignant states, GPM produced the highest frequency of significant FOVs (12.3%), followed by PPR (7.4%), NEU (5.7%), and MTC (5.4%). The pooled tumor compartment exhibited departures from the global null expectation more often than individual, separated tumor states.

Pairwise random-labeling analysis demonstrated spatial heterogeneity across partner cell types (Figure 5B). Vasculature produced the highest proportions of significant FOVs among non-tumor populations, ranging from 47.5% to 57.9% across all tumor states. Immune populations exhibited recurrent departures from random labeling. Microglia and Monocyte comparisons yielded significance in 23% to 29% of FOVs, while cDC, Macrophage, and T-cell comparisons achieved significance in 20% to 25% of evaluable FOVs. Pairwise comparisons among malignant states produced high rejection frequencies. Comparisons involving GPM or PPR as partner populations reached significance in 36.9% to 42.7% of FOVs. Comparisons involving MTC and NEU exceeded 30%. These results indicate that the spatial allocation of malignant-state labels within the observed point configurations was frequently inconsistent with random labelling.

The independence-of-components analysis generated lower proportions of significant FOVs than the random-labeling analysis while maintaining recurrent spatial patterns (Figure 5C). Vasculature maintained prominent spatial dependence, with significant FOV proportions ranging from 12.9% to 29.8%. Microglia maintained significance in 17% to 20% of FOVs, and other immune populations maintained dependence in 14% to 20% of evaluable FOVs. Spatial dependence among malignant states persisted under the independence-of-components null model. Comparisons involving PPR as a partner population achieved significance in 31.9% to 34.3% of FOVs. GPM-related pairings ranged from 22% to 36%, and comparisons involving MTC or NEU exceeded 20% of evaluable FOVs.

FOV-level tests operate on a local scale. Local abundance and spatial configuration dictate the detection of departures from the null model. Sparse populations or locally homogeneous spatial configurations restrict evidence for rejecting the null hypothesis within evaluable FOVs. The global analyses capture recurrent significant deviations across numerous cell-type combinations despite this local variability. The variation in significance frequencies among GPM, MTC, NEU, PPR, and the pooled Tumor compartment highlights the spatial heterogeneity of the malignant compartment, also emphasizing the limitations of treating individual FOVs as isolated spatial units. Evaluating spatial relationships across the continuous tissue domain requires whole-specimen modeling to capture both local and broader-scale spatial variation.

#### Malignant States Attract Immune Populations Across the Continuous Tissue Domain

Classical point process analyses calculate exact pairwise distances but become computationally intractable on multiplexed, whole-slide transcriptomics datasets. Dividing tissue into artificial FOV-by-FOV tiles introduces edge effects that truncate continuous biological gradients and mask macro-architectural recruitment patterns. Discretizing the whole-tissue space into a finite triangular mesh establishes the computational topology required to evaluate the unbroken tissue domain. Log-Gaussian Cox Processes (LGCP) modeled via Stochastic Partial Differential Equations (SPDE) operate on this mesh to capture microscopic cellular niches and long-range spatial gradients. The LGCP represents a “response” cell type as a Poisson point process. The spatial density of the “target” tumor subpopulation, calculated via Gaussian kernel density estimation, serves as the sole environmental covariate. Utilizing only the independent target density as a spatial predictor avoids structural endogeneity.

The model defines the log-intensity of the response cells, λ(s), as λ(s) = exp(β_0_ + β_1_X_target_(s) + ⍵(s)) ^35^. The parameter β_0_ represents the baseline cellular intensity. The fixed effect coefficient β_1_ measures the influence of the target tumor density, (X_target_). The Matérn spatial random field ⍵(s) acts as a spatial confounding filter, absorbing background tissue heterogeneity, unobserved spatial clustering, and stochastic noise. The random field accounts for baseline spatial autocorrelation and isolates the log-linear spatial dependence β_1_ between the response cells and the target tumor cells.

The LGCP framework tests Complete Spatial Randomness (CSR) against spatial attraction or repulsion. A baseline scenario of spatial independence yields a β_1_ coefficient equal to zero. The 95% Credible Interval (CI) of the β_1_ posterior distribution classifies the spatial interaction. A CI lower bound greater than zero rejects the CSR null hypothesis and indicates spatial attraction. A CI upper bound less than zero rejects the CSR null hypothesis and indicates spatial repulsion. A CI crossing zero indicates spatial independence.

The LGCP models identified spatial attraction and repulsion patterns between the malignant tumor states (GPM, MTC, NEU, and PPR) and the surrounding microenvironment (Figure 5D; patient-specific whole-specimen heatmaps are shown in Figure S2). Rejection of the CSR null hypothesis confirms biological cellular aggregation. Innate and adaptive immune compartments, including macrophages, microglia, monocytes, cDCs, neutrophils, and T cells, exhibited attraction to all tumor states across the patient cohort. Vasculature displayed inter-patient heterogeneity, showed structural attraction to the tumor states in specimens um08, um10, um11, and um15, and high spatial repulsion in specimen um13 against the GPM state.

### Quantifying the Probability of Cellular Proximity to Tumors

The selected fine mesh discretizes the continuous spatial domain and provides the topological infrastructure for solving Stochastic Partial Differential Equations (SPDE). The analysis converts interrelated cellular subpopulations into paired spatial point patterns. Each pair consists of a target population (e.g., GPM, PPR, NEU, MTC) and a modeled response population (e.g., macrophages, microglia). Gaussian kernel density estimation generates continuous surfaces of cellular abundance for both populations in the pair. Cell-type-specific density maps across the complete cohort are shown in Figure S3. The harmonic mean of these densities synthesizes the Niche Density covariate: 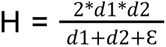.This formulation penalizes tissue regions dominated by a single population and emphasizes regions in which both populations are simultaneously abundant. The resulting Niche Density surface was standardized as a Z-score, with zero mean and unit standard deviation. Whole-specimen Niche Density maps for all tumor-state–partner-cell combinations are shown in Figure S4.

Including the response cell density (d2) in the predictor introduces structural endogeneity. This condition violates classic causal inference assumptions, but the objective of this step prioritizes extracting continuous predictive features. It anchors the maximum conditional intensity at direct interaction boundaries to isolate microenvironments of probable metabolic exchange.

The Log-Gaussian Cox Process (LGCP) specifies the conditional intensity rate λ(s) of the response population through a log-linked linear predictor: λ(s) = exp(β_0_ + β_1_X_niche_ + ⍵(s)). The parameter β_0_represents the global intercept. The fixed effect coefficient β_1_ measures the predictive niche covariate X_niche_. The Matérn spatial random field ⍵(s)completes the Bayesian formula. During training, the spatial random field ⍵(s)absorbs latent spatial autocorrelations, independent clustering, and structural tissue artifacts. A topologically dissociated prediction step extracts the final conditional intensity. Limiting the predictive function to the fixed components, λ_pred_ = exp(β_0_ + β_1_X_niche_), excludes the spatial field. This exclusion ensures the predicted per capita cellular intensity rate reflects the pure influence of the modeled interaction niche.

The cumulative distribution function of the Poisson distribution, P(N ≥ 1) = 1 - exp(-λ_pred_. A), transforms the extracted conditional intensity λ_pred_ into a probability metric of biological encounter within a pre-defined spatial area A. This equation translates the abstract magnitude of the Poisson intensity into a standardized metric of cellular connectivity bounded between 0 and 1. This continuous probability value provides the weighting required for downstream spatial-dependent metabolic activation and reprogramming algorithms. Whole-specimen encounter-probability maps for all tumor-state–partner-cell combinations are shown in Figure S5.

Visual dissection of interaction topologies in patient um08 validates the spatial framework (Figure 6). Comparing independent spatial density maps against model-derived encounter probabilities (30 µm radius) demonstrates the isolation of co-localized niches. GPM cells concentrate in the inferior tissue region (Figure 6A). Macrophages distribute across the tissue (Figure 6B). Astrocytes cluster in the upper-right (Figure 6C), and classical dendritic cells (cDCs) peak in the bottom-left (Figure 6D). The calculated encounter probabilities filter these broad distributions. High interaction probabilities for GPM and macrophages map to their overlapping inferior niche (Figure 6E). The probability scores for astrocytes and cDCs isolate specific focal points of mutual concentration and bypass non-overlapping dense regions (Figure 6F, 6G).

**Figure 6.**
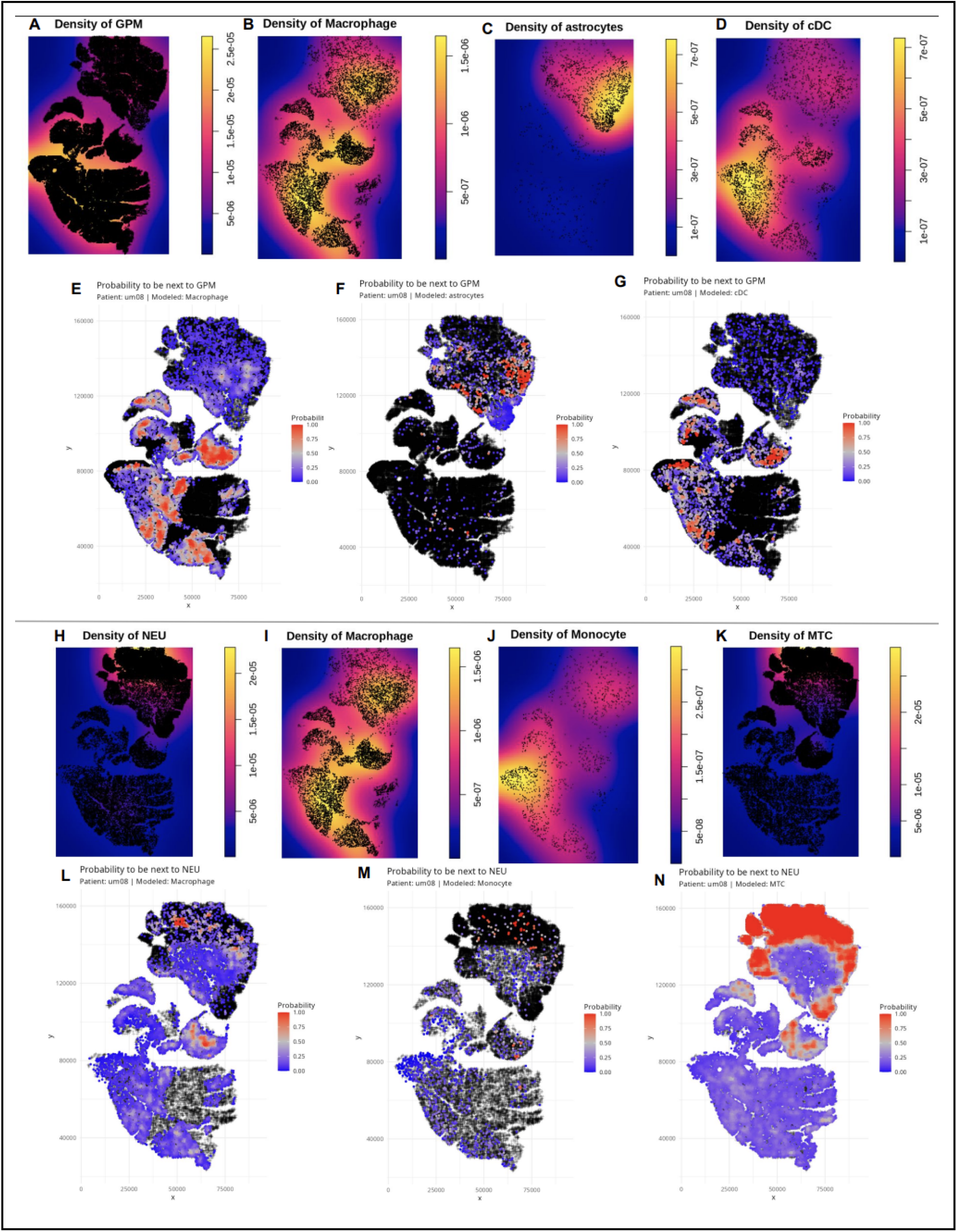
Spatial validation of LGCP-derived cellular encounter probabilities in patient um08. **A–D)** Continuous Gaussian kernel density estimation maps illustrating the spatial distribution of the target GPM population (A) and the modeled response populations: macrophages (B), astrocytes (C), and classical dendritic cells (cDCs) (D). **(E–G)** Resulting spatial probability maps projecting the 0 to 1 likelihood of encountering a GPM cell within a 30 μm radius for macrophages (E), astrocytes (F), and cDCs (G). The probability metric successfully filters broad density distributions to isolate specific niches of symmetric co-localization, such as the inferior overlapping niche of GPM and macrophages. **(H–K)** Independent spatial density maps for a distinct target population, NEU (H), alongside modeled response populations: macrophages (I), monocytes (J), and MTC cells (K). **(L–N)** Corresponding spatial probability maps displaying the likelihood of encountering a NEU cell within a 30 μm radius for macrophages (L), monocytes (M), and MTC cells (N).

Designating NEU as the target population shifts the interaction probabilities to match its dense superior niche (Figure 6H). Macrophage density remains widespread across the tissue (Figure 6I). Their NEU-encounter probability maps to the upper region (Figure 6L). Monocytes form a dense cluster in the center-left (Figure 6J). The model bypasses this isolated cluster and assigns high encounter probabilities to sparse monocyte populations overlapping with the superior NEU niche (Figure 6M). MTC cells exhibit a high encounter probability matching their co-localization at the top of the tissue (Figure 6K, 6N).

These visualizations confirm the mathematical framework requires overlapping target and response densities to generate high spatial encounter probabilities. Having established the reliability of these quantitative spatial probabilities, we will apply these scores as the features for our downstream metabolic/transcriptome prediction models.

### Transcriptomic profiles outperform metabolic pathway scores within the cohort

Under cell-grouped validation (cellgroup5), the transcriptomic representation achieved pooled R² of 0.649 and Spearman 0.817 for prob_logit, and 0.679 and 0.824 for intensity_log10 (XGBoost), while the metabolic representation reached 0.384/0.619 and 0.410/0.625 respectively (XGBoost; the supporting learners peak at 0.397/0.629 on prob_logit, still far below transcripts; Table S4). The ordering was not an artefact of model family: within each representation, XGBoost exceeded ElasticNet (transcriptomic ElasticNet: 0.453/0.692 and 0.476/0.695; metabolic ElasticNet: 0.322/0.557 and 0.347/0.566), and the supporting learners agreed that transcripts dominate (Table 1). A ridge model using only interaction-identity encoding achieved 0.305/0.520 for prob_logit (Table S5), indicating that the molecular representations provided additional predictive information beyond cell-type identity. Under loso, performance was much lower for both representations: ElasticNet achieved 0.003/0.339 and 0.047/0.377 for transcripts, and 0.077/0.296 and 0.081/0.308 for metabolism. Thus, neither representation generalized well across patients.

### Uncertainty increases under leave-one-patient-out validation

The patient-cluster bootstrap shows substantially greater uncertainty under loso than under cellgroup5. Under cellgroup5, transcriptomic XGBoost achieved R² 0.648–0.651 and Spearman 0.816–0.818 for prob_logit, compared with 0.382–0.386 and 0.618–0.621 for metabolism, respectively. Under loso, intervals were much wider: transcriptomic XGBoost prob_logit R² ranged from −2.856 to 0.352 and Spearman from 0.098 to 0.623, compared with −4.103 to 0.474 and 0.128 to 0.675 for metabolism. These intervals do not provide clear evidence of a difference between representations at the eight-patient sample size. The ElasticNet results showed similarly wide intervals (prob_logit: transcripts R² −3.224 to 0.349, Spearman 0.247–0.597; metabolism R² −4.076 to 0.452, Spearman 0.131–0.624), indicating that uncertainty is driven primarily by the number of independent patients rather than the number of rows.

### Cross-patient generalization is limited

Under leave-one-patient-out validation, absolute predictive performance was poor for both representations. XGBoost had mean per-fold R² of −0.742 for transcriptomic features (range −2.617 to 0.264) and −1.263 for metabolic features (range −4.345 to 0.231) on prob_logit. Pooled out-of-fold R² remained positive (0.051 and 0.059, respectively; Table 2), reflecting the different weighting of between-patient and within-patient prediction in the pooled and per-fold summaries. Spearman correlations were more stable, with mean values of 0.369 for transcripts and 0.334 for metabolism. ElasticNet showed the same pattern, with mean per-fold R² of −0.826 and −1.289 and mean Spearman values of 0.322 and 0.319 for transcripts and metabolism, respectively. The supporting learners showed similarly poor cross-patient performance, with mean per-fold R² of −1.30 for random forest and −1.29 for MLP and mean Spearman values of 0.327 and 0.331. The identity-only ridge model reached a pooled Spearman of 0.106 for prob_logit (Tables S4–S5).

**Table 2.**
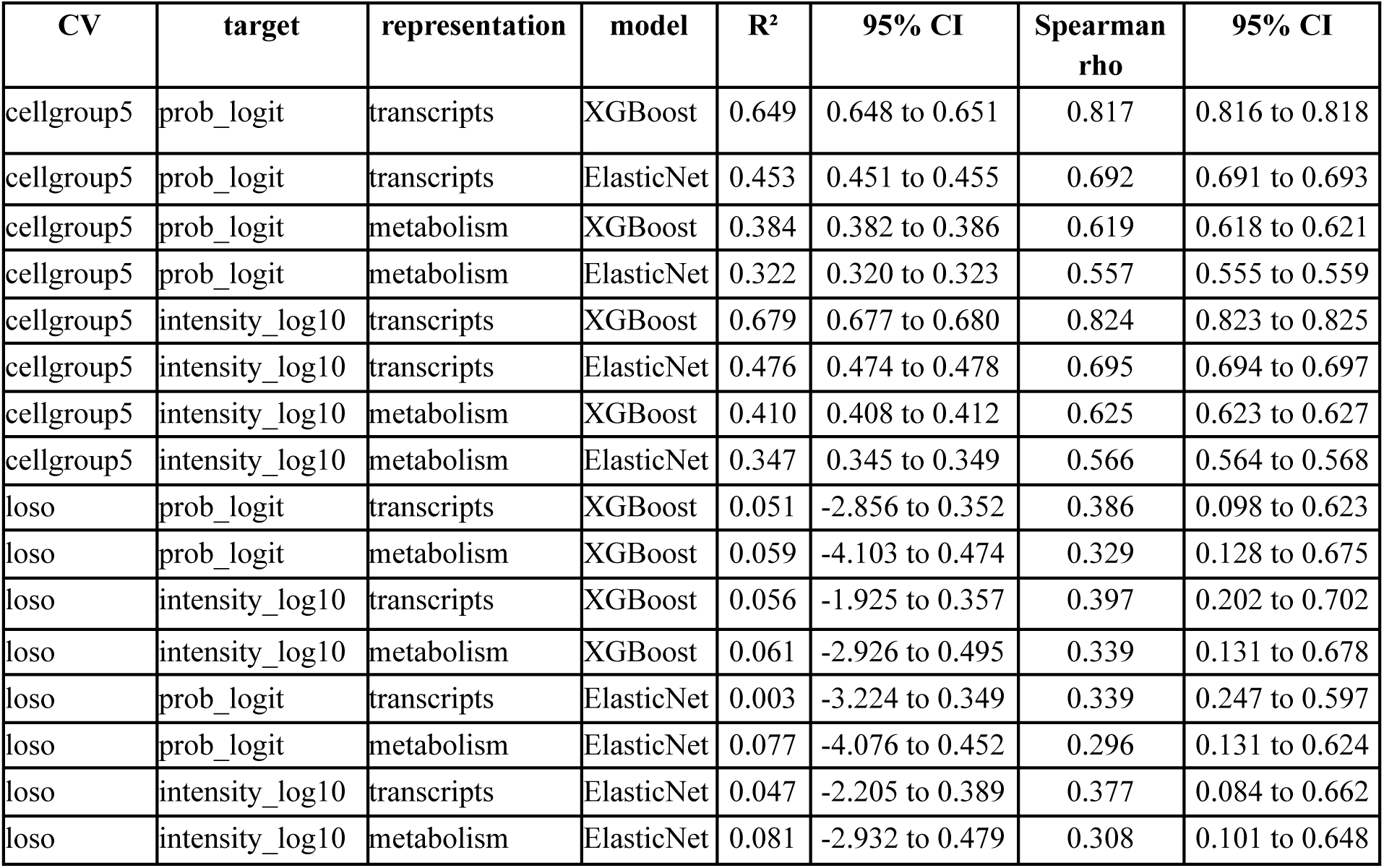
Headline performance: pooled out-of-fold R² and Spearman ρ on the transformed scale (logit probabilities and log10 intensities), 95% bootstrap interval (200 replicates). cellgroup5 holds out cells from patients represented in the training data, whereas loso holds out entire patients. Under cellgroup5, the transcriptomic representation outperforms metabolism across all learners. Under loso, differences between representations and learners become small, with the supporting random forest, MLP, and identity-only ridge showing the same pattern (Tables S4-S5).

### Harmonization improves within-patient performance but does not restore cross-patient generalization

Per-patient feature harmonization produced a modest improvement in transcriptomic LOSO performance on the raw scale: mean per-fold R² improved from −0.742 to −0.710, while mean within-patient R² improved from −0.136 to −0.093 (H+T; Table 3). Mean Spearman remained similar (0.369 vs. 0.365). Centering the target in addition to feature harmonization (H+C) further improved mean within-patient R² to −0.067, with positive within-patient R² for um10, um09, and um11 (0.216–0.265) but negative values for um13 and um15. Spearman performance remained similar across harmonization strategies (0.365–0.369), indicating that harmonization improved within-patient fit without improving rank performance or restoring cross-patient generalization.

**Table 3.**
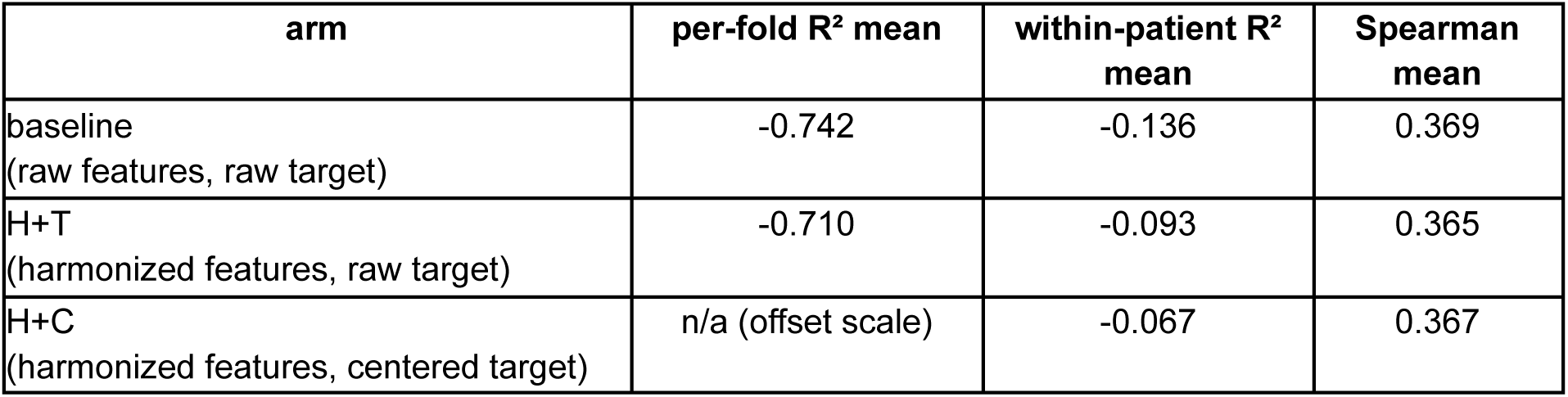
Harmonization experiment, loso, prob_logit, transcriptomic XGBoost: per-fold means over eight folds.

| arm | per-fold $R^2$ mean | within-patient $R^2$ mean | Spearman mean |
| --- | --- | --- | --- |
| baseline<br>(raw features, raw target) | -0.742 | -0.136 | 0.369 |
| H+T<br>(harmonized features, raw target) | -0.710 | -0.093 | 0.365 |
| H+C<br>(harmonized features, centered target) | n/a (offset scale) | -0.067 | 0.367 |

### Patient-level heterogeneity contributes to transportability loss

Per-patient analysis of the transcriptomic XGBoost arm (Table 4) showed moderate-to-good rank correlations in seven of eight held-out patients (Spearman 0.257–0.547), despite substantial variation in R². The largest negative R² values occurred in patients with lower outcome base rates. Patient-level outcome distributions also varied markedly, with median per-cell interaction probabilities ranging from 0.047 to 0.961. In contrast, assay depth was more similar across patients, with median detected genes per cell ranging from 481 to 929 and median counts from 559 to 1412. Removing patient-level offsets reduced the R² penalty for the most extreme case, um12, from −2.617 to −0.196 while preserving a Spearman correlation of 0.409. The metabolic representation showed a similar pattern, with transcriptomic XGBoost achieving equal or higher Spearman correlation in six of eight patients, including um13 (0.481 vs. 0.369). For um15, however, Spearman correlation was low for both representations (−0.015 metabolic, 0.039 transcriptomic). Together, these results indicate that patient-level differences in outcome scale contribute to the loss of cross-patient predictive performance.

**Table 4.**
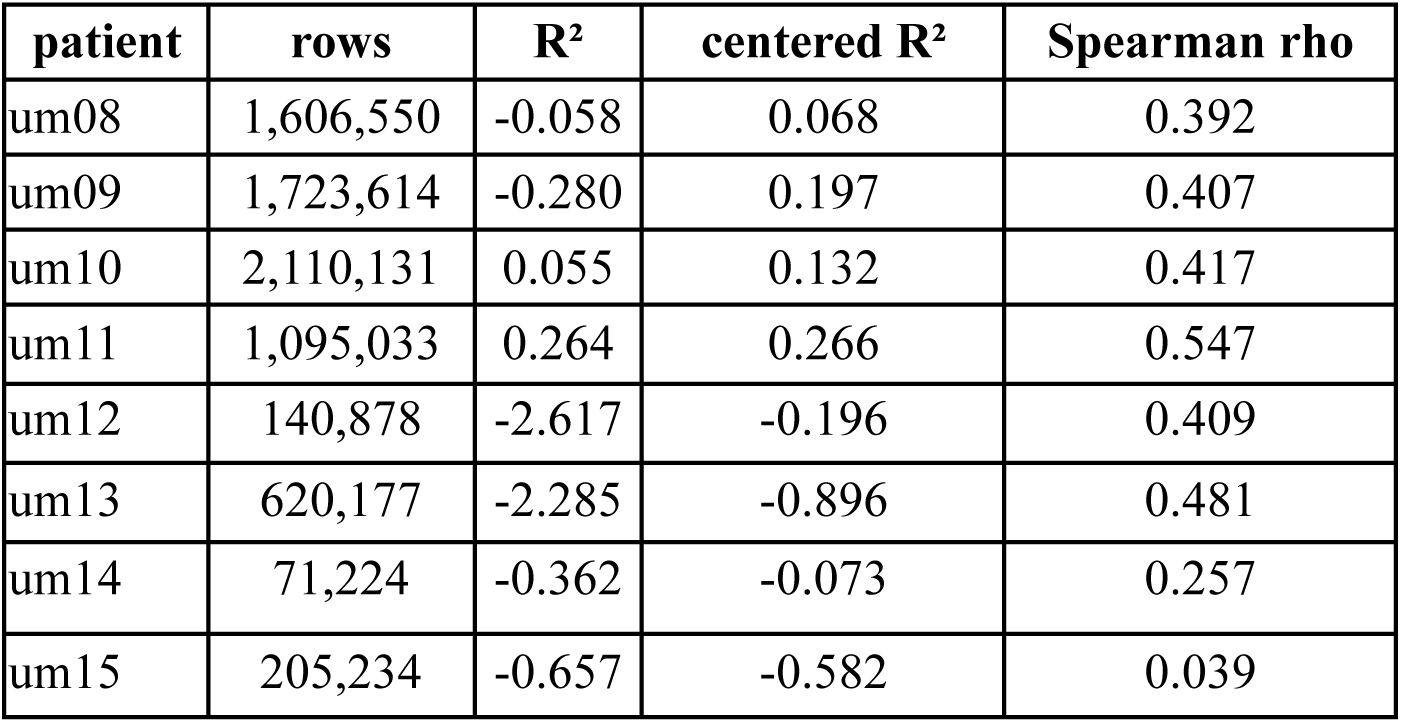
Leave-one-patient-out validation for prob_logit using transcriptomic XGBoost, shown by held-out patient.

| patient | rows | R<sup>2</sup> | centered R<sup>2</sup> | Spearman rho |
| --- | --- | --- | --- | --- |
| um08 | 1,606,550 | -0.058 | 0.068 | 0.392 |
| um09 | 1,723,614 | -0.280 | 0.197 | 0.407 |
| um10 | 2,110,131 | 0.055 | 0.132 | 0.417 |
| um11 | 1,095,033 | 0.264 | 0.266 | 0.547 |
| um12 | 140,878 | -2.617 | -0.196 | 0.409 |
| um13 | 620,177 | -2.285 | -0.896 | 0.481 |
| um14 | 71,224 | -0.362 | -0.073 | 0.257 |
| um15 | 205,234 | -0.657 | -0.582 | 0.039 |

### Gene overlap and label transfer support single-cell deployment

Of the 6,175 genes in the CosMx panel, 5,982 (96.9%) were present in the GBmap query and were used for transfer, while the remaining 193 (3.1%) were zero-filled at deployment. The GBmap query contained 27,625 unique feature names, including 21,643 that were not represented in the CosMx panel (Table S6).

Label transfer assigned 95.6% of GBmap cells to a CosMx cell type, leaving 4.4% unassigned (14,960 of 338,564). Malignant cells were assigned to the four CosMx states at 96.1% (122,542 of 127,521; GPM 58,213; PPR 51,201; NEU 12,173; MTC 955). Mature T cells mapped to Tcells at 89.6% (48,595 of 54,257), microglial cells to Microglia at 67.1%, and dendritic cells to cDC at 60.0%. B, plasma, and mast cells remained unassigned by construction (Table 5). Residual myeloid ambiguity, including GBmap macrophages mapping to Macrophage at 34.1% and microglial cells mapping to Macrophage or Neutrophil, was retained as posterior uncertainty rather than treated as hard misclassification. Deployment inference covered 1,294,416 cell-by-target pairs across 323,604 cells from 110 donors and was evaluated strictly using within-donor ranks.

**Table 5.** Label-transfer outcomes for GBmap cell types using masked scANVI with a posterior threshold of 0.5. cells = number of GBmap cells of each type; top assignment = most frequent CosMx label, with n and % denoting the number and proportion of cells assigned to that label; unassigned = cells forced to unassigned because no CosMx counterpart was available (B, plasma, and mast cells) or because the maximum posterior probability was <0.5. Rows sum to the TOTAL of 338,564 cells, of which 4.4% were unassigned. Non-top assignments are retained in the table to show the distribution of label-transfer outcomes across cell types.

| GBmap type | cells | top assignment | n | % of type | unassigned n (%) |
| --- | --- | --- | --- | --- | --- |
| malignant cell | 127,521 | states<br>(GPM/PPR/NEU/MTC) | 122,542 | 96.1% | 2,047 (1.6%) |
| macrophage | 61,134 | Macrophage | 20,857 | 34.1% | 4,540 (7.4%) |
| microglial cell | 54,722 | Microglia | 36,721 | 67.1% | 2,383 (4.4%) |
| mature T cell | 54,257 | Tcells | 48,595 | 89.6% | 1,591 (2.9%) |
| monocyte | 14,215 | Monocyte | 9,144 | 64.3% | 232 (1.6%) |
| oligodendrocyte | 12,481 | oligodendrocyte | 11,561 | 92.6% | 118 (0.9%) |
| dendritic cell | 3,961 | cDC | 2,377 | 60.0% | 139 (3.5%) |
| radial glial cell | 2,807 | unassigned | 1,598 | 56.9% | 1,598 (56.9%) |
| natural killer cell | 2,489 | Tcells | 2,287 | 91.9% | 53 (2.1%) |
| mural cell | 1,418 | vasculature | 1,351 | 95.3% | 25 (1.8%) |
| B cell | 1,250 | unassigned (forced) | 1,250 | 100% | 1,250 (100%) |
| endothelial cell | 673 | vasculature | 638 | 94.8% | 23 (3.4%) |
| plasma cell | 572 | unassigned (forced) | 572 | 100% | 572 (100%) |
| oligodendrocyte precursor cell | 496 | OPC | 483 | 97.4% | 7 (1.4%) |
| mast cell | 373 | unassigned (forced) | 373 | 100% | 373 (100%) |
| astrocyte | 173 | astrocytes | 153 | 88.4% | 9 (5.2%) |
| neuron | 22 | neurons | 19 | 86.4% | 0 (0%) |
| TOTAL | 338,564 | — | — | — | 14,960 (4.4%) |

### Predicted interaction scores stratify transcriptional subclusters within type

We next asked whether the spatial interaction phenotype was sufficiently encoded in transcriptional variation to enable its application to scRNA-seq data. Type-only reclustering revealed substantial variation in predicted interaction scores across transcriptional subclusters, with the strongest associations concentrated in specific type–partner pairs (Figure 7; Table 6). Across all 40 type–target combinations, GPM→PPR showed the largest departure from the within-donor shuffle null (η² = 0.379; null = 0.083; gap = 0.296; between-donor share = 0.257; 58,213 cells from 90 donors in 20 reclusters), followed by Monocyte→MTC (η² = 0.269; null = 0.082; gap = 0.187; between-donor share = 0.393; 16,974 cells from 91 donors in 13 reclusters). Among the best-partner pairs for each cell type, Macrophage→MTC ranked third (η² = 0.238; null = 0.125; gap = 0.113; between-donor share = 0.444; 34,178 cells from 86 donors in 19 reclusters), followed by Vasculature→GPM (η² = 0.416; null = 0.317; gap = 0.099; between-donor share = 0.386; 13,908 cells from 96 donors in 16 reclusters). The corresponding recluster means spanned −0.61 to 2.45, −2.29 to −0.23, −2.76 to −0.50, and −0.77 to 2.16, respectively.

**Figure 7.**
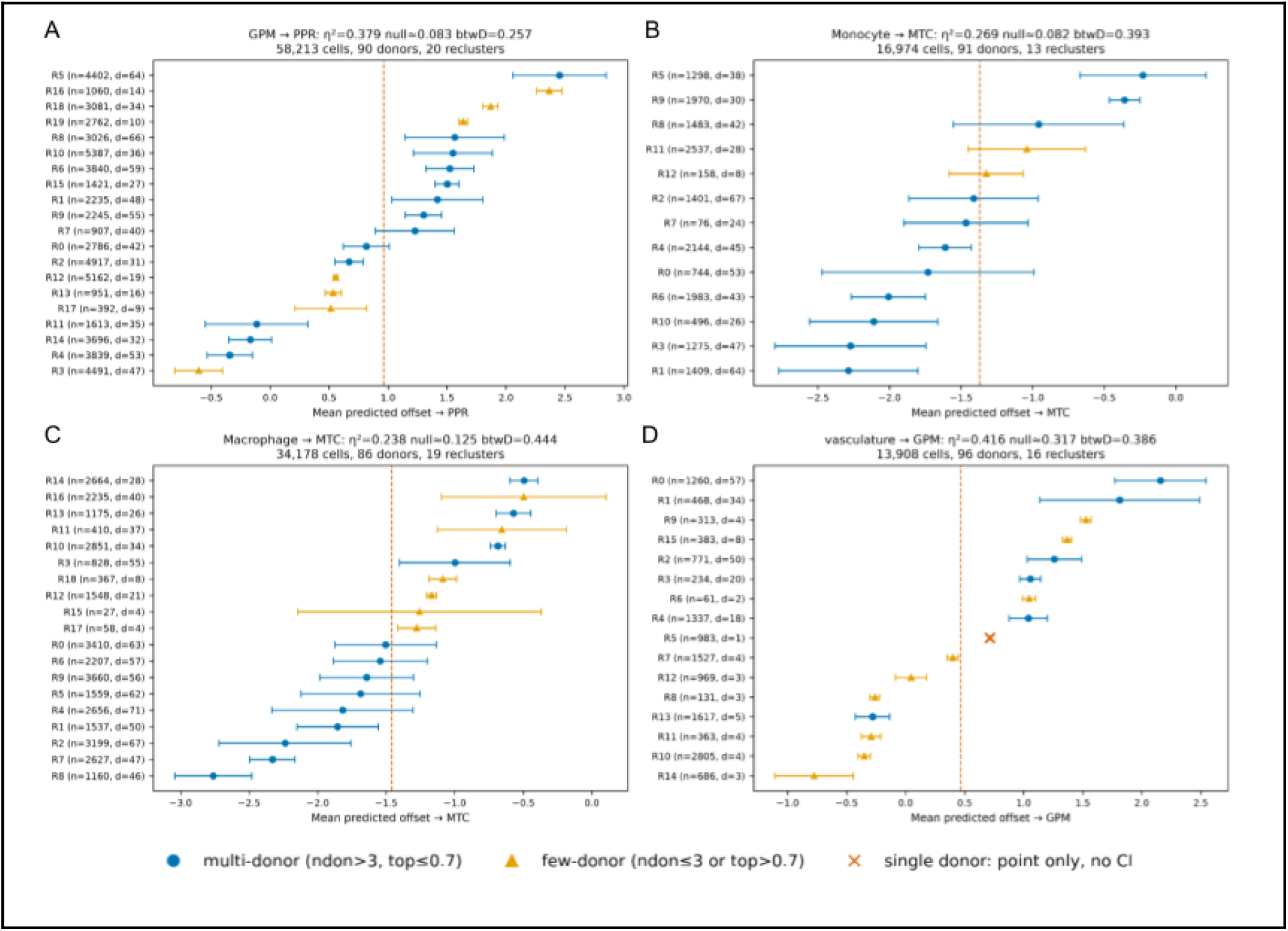
Subcluster forest plots for the four strongest partner-specific signals. **A)** GPM toward PPR, **B)** monocytes toward MTC, **C)** macrophages toward MTC, and **D)** vasculature toward GPM. Points show mean predicted within-donor offsets for each recluster; bars indicate 95% donor-clustered intervals (±1.96 × SE). Circles denote multi-donor reclusters, triangles reclusters with limited donor representation (≤3 donors or >70% of cells from a single donor), and crosses single-donor reclusters, for which no generalizable interval is shown. The dashed line marks the corresponding type mean. Panel titles report η², the within-donor shuffle null, between-donor variance share, and cell, donor, and recluster counts.

**Table 6.** Type-only reclustering, best partner per type: subcluster association (eta-squared) with within-donor shuffle null and between-donor share.

| type | best partner | cells | donors | reclusters | eta2 | null | btw-donor | range |
| --- | --- | --- | --- | --- | --- | --- | --- | --- |
| GPM | PPR | 58,213 | 90 | 20 | 0.379 | 0.083 | 0.257 | -0.61 to 2.45 |
| vasculature | GPM | 13,908 | 96 | 16 | 0.416 | 0.317 | 0.386 | -0.77 to 2.16 |
| Macrophage | MTC | 34,178 | 86 | 19 | 0.238 | 0.125 | 0.444 | -2.76 to -0.50 |
| PPR | GPM | 51,201 | 91 | 24 | 0.139 | 0.102 | 0.282 | 0.91 to 2.73 |
| Microglia | MTC | 49,580 | 87 | 15 | 0.196 | 0.186 | 0.600 | -1.03 to 0.63 |
| Tcells | GPM | 51,519 | 87 | 14 | 0.067 | 0.037 | 0.125 | -0.23 to 0.54 |
| Monocyte | MTC | 16,974 | 91 | 13 | 0.269 | 0.082 | 0.393 | -2.29 to -0.23 |
| Neutrophil | MTC | 15,920 | 88 | 13 | 0.159 | 0.109 | 0.585 | -2.89 to -1.19 |
| NEU | GPM | 12,173 | 69 | 14 | 0.234 | 0.209 | 0.432 | -2.13 to 0.13 |
| cDC | MTC | 5,310 | 79 | 13 | 0.203 | 0.179 | 0.480 | -2.15 to -0.29 |

The strongest gradients were represented across multiple donors. GPM R4 and R5 comprised 3,839 cells from 53 donors and 4,402 cells from 64 donors, respectively, with the largest donor contributing 13% of R5. Monocyte contrasts involved reclusters represented by 30–64 donors, and macrophage contrasts by 26–71 donors. The lower end of the Vasculature→GPM gradient included clusters represented by five or fewer donors, including one single-donor recluster containing 983 cells. These differences in donor representation provide context for the interpretation of the observed gradients.

These results show that the predicted spatial interaction phenotype is structured by transcriptional variation within cell type, providing a rationale for exploring its potential application to scRNA-seq data.

### State dependence is partner-specific

The same transcriptional subclusters were evaluated across all four interaction partners, allowing partner-specific variation to be assessed within a fixed cellular partition. GPM subclusters showed a pronounced gradient toward PPR, whereas scores toward MTC remained high across reclusters (η² = 0.193; range = 1.09–2.59). In monocytes, MTC showed the largest departure from the shuffle null (gap = 0.187), compared with ≤0.113 for the other partners. Vasculature showed departures from the null for GPM, NEU (η² = 0.331; null = 0.243), and PPR (η² = 0.232; null = 0.138), whereas MTC showed a smaller departure (η² = 0.132; null = 0.037). Macrophage subclusters also varied across partners, with between-donor shares ranging from 0.44 to 0.59.

The remaining best-partner combinations showed smaller departures from their within-donor nulls: PPR→GPM (η² = 0.139 vs. 0.102), Microglia→MTC (0.196 vs. 0.186), NEU→GPM (0.234 vs. 0.209), cDC→MTC (0.203 vs. 0.179), Neutrophil→MTC (0.159 vs. 0.109), and T cells→GPM (0.067 vs. 0.037). The complete set of 40 type–target combinations is reported in Table S7, with the remaining best-partner profiles shown in Figure S6.

Because clustering and score calculation were derived from the same expression matrix, η² may be optimistic; the within-donor shuffle therefore provides the reference for evaluating the excess association. Type–partner pairs were selected post hoc by η², so these patterns represent candidate interaction-associated transcriptional subclusters rather than validated biological states.

Thus, transcriptionally encoded variation in the predicted spatial phenotype was partner-specific rather than a uniform property of cellular subclustering.

## Conclusions

A whole-specimen spatial framework integrated finite-element meshes, point-process statistics, and LGCP/SPDE modeling. This approach extracted continuous cell-level spatial phenotypes, including conditional cellular intensity and tumor-proximity probability. Molecular profiles successfully predicted these spatial metrics without relying on spatial coordinates. Within the training cohort, transcriptomic representations outperformed metabolic pathway scores in predicting spatial proximity. Leave-one-patient-out validation revealed limited cross-patient generalization. Patient-level feature harmonization improved within-patient fit but failed to restore cross-patient transportability. Patient-specific tumor architecture strongly conditions these molecular-spatial relationships. The small sample of independent spatial specimens restricts broad generalization across new patients.

Deploying the spatial framework on external single-cell RNA sequencing data successfully stratified transcriptional subclusters within cell types using predicted interaction scores. The transcriptomes encode partner-specific spatial variation. Generalizing these findings requires larger, more diverse spatial cohorts. Future validation must test cross-modality transfer using datasets with matched spatial transcriptomics and single-cell RNA sequencing from the exact same patients.

## Acknowledgements

This study was financed, in part, by the São Paulo Research Foundation (FAPESP), Brasil. Process Numbers: #2025/15993-2 ; #2025/26980-9 ; #2025/17477-1 ; #2018/00583-0. The opinions, hypotheses, and conclusions or recommendations expressed in this material are the responsibility of the authors and do not necessarily reflect the views of FAPESP.

## Supplementary Data

**Table S1**. **Specimen-level sampling and cellular density**. Number of FOVs, total cell counts, and overall cell density for each GBM specimen. Density is reported for the complete cellular compartment and summarised across FOVs within each specimen. Values include the number of FOVs and cells contributing to each specimen-level estimate.

**Table S2. Cell-type and tumour-state composition across GBM specimens.** Cell counts and relative proportions for the 15 annotated cell populations and the four tumour-cell states (PPR, NEU, MTC, and GPM) in each specimen. P values from χ² tests for homogeneity assess differences in the representation of each cell population or tumour state between specimens. P values were adjusted for multiple testing using the Benjamini-Hochberg (BH) procedure.

**Table S3. Cell-type-specific spatial density across GBM specimens.** Density distributions for each of the 15 annotated cell populations across FOVs within each specimen. Differences in density between specimens were assessed using Kruskal-Wallis tests, with BH adjustment for multiple comparisons. The table reports the corresponding test statistics and adjusted P values for each cell population.

**Table S4. Supporting learners**: random forest and MLP results for the metabolic representation, reported as pooled out-of-fold R² and Spearman ρ on the transformed scale.

**Table S5. Identity-only baseline**: ridge regression using one-hot encoding of the cell-type-by-target combination, without molecular features.

**Table S6. CosMx–GBmap gene overlap**: Three membership lists over model-space gene symbols: both_5982 (panel genes found in the query; carried transfer and applied), cosmx_only_193 (panel genes absent from GBmap; zero-filled at deployment), gbmap_only_21643 (query genes outside the panel; unused).

**Table S7. Full subcluster association results:** all 40 type-by-target combinations (10 reclustered types by 4 targets) with cell, donor and recluster counts, eta-squared, within-donor shuffle null (20 shuffles), gap (eta minus null) and between-donor share, plus the observed range of recluster means.

**Figure S1. FOV-level CSRI analyses across tumor populations and cellular partners.** Violin and dot plots show FOV-specific results for GPM, MTC, NEU, PPR, and the pooled Tumor compartment across global spatial-randomness, random-labelling, and component-independence tests. Each point represents an evaluable FOV and is colored according to significance at p < 0.05.

**Figure S2. Whole-specimen LGCP interaction profiles across the CosMx cohort.** Panels A–H correspond to specimens um08–um15. Heatmaps show LGCP coefficients between tumor states (rows) and partner cell types (columns). Positive coefficients indicate spatial recruitment and negative coefficients indicate spatial avoidance. Asterisks denote significant effects, whereas grey tiles indicate pairs with insufficient cell counts.

**Figure S3. Cell-type-specific density distributions across CosMx specimens.** Spatial density distributions of the 15 annotated cell populations are shown across FOVs for specimens um08–um15, illustrating inter-specimen heterogeneity in the abundance and spatial representation of individual cell types.

**Figure S4. Whole-specimen relative-density maps across the CosMx cohort.** Panels A–H correspond to specimens um08–um15. For each tumor-state–partner-cell pair, Gaussian KDE surfaces, for target and modeled cell types, were calculated separately from the observed cellular coordinates, used to calculate a harmonic mean and standardized as Z-scores. These maps represent the relative spatial enrichment of co-occurrence of both cell types population across the tissue, with higher values indicating regions of high-density of both cell types. The maps are derived directly from the observed cellular density surfaces and do not include the LGCP/SPDE latent spatial field, thus representing descriptive spatial organization rather than model-adjusted interaction effects.

**Figure S5. Whole-specimen probability maps across the CosMx cohort.** Panels A–H correspond to specimens um08–um15. For each tumor-state–partner-cell pair, maps show the model-derived probability of encountering a target cell within the predefined interaction radius, based on the LGCP-estimated conditional intensity. Probability values range from 0 to 1, with higher values indicating a greater predicted likelihood of local cellular proximity. Unlike the descriptive density maps shown in Figures S3 and S4, these maps represent the probabilistic output of the spatial model.

**Figure S6. Subcluster forest plots for the six remaining best-partner pairs** (PPR toward GPM, Microglia toward MTC, T cells toward GPM, Neutrophil toward MTC, NEU toward GPM, cDC toward MTC), using the same encoding as Figure 7. None shows a substantial departure from its within-donor shuffle null; the PPR toward GPM top recluster (R23, 30 cells, 2 donors) and the cDC toward MTC bottom recluster (R9, 20 cells, 1 donor) illustrate why few- and single-donor extremes provide limited evidence for generalizable state contrasts.

## References

1. Parsons, D. W. et al. An Integrated Genomic Analysis of Human Glioblastoma Multiforme. Science 321, 1807–1812 (2008).

2. Schürch, C. M. et al. Coordinated Cellular Neighborhoods Orchestrate Antitumoral Immunity at the Colorectal Cancer Invasive Front. Cell 182, 1341–1359.e19 (2020).

3. Hambardzumyan, D. & Bergers, G. Glioblastoma: Defining Tumor Niches. Trends Cancer 1, 252–265 (2015).

4. Whittle, J. R. et al. Spatial omics shed light on the tumour organisation of glioblastoma. Semin. Cell Dev. Biol. 167, 1–9 (2025).

5. Greenwald, A. C. et al. Integrative spatial analysis reveals a multi-layered organization of glioblastoma. Cell 187, 2485–2501.e26 (2024).

6. Piwecka, M., Rajewsky, N. & Rybak-Wolf, A. Single-cell and spatial transcriptomics: deciphering brain complexity in health and disease. Nat. Rev. Neurol. 19, 346–362 (2023).

7. Tasic, B. et al. Adult mouse cortical cell taxonomy revealed by single cell transcriptomics. Nat. Neurosci. 19, 335–346 (2016).

8. Zeisel, A. et al. Cell types in the mouse cortex and hippocampus revealed by single-cell RNA-seq. Science 347, 1138–1142 (2015).

9. Eze, U. C., Bhaduri, A., Haeussler, M., Nowakowski, T. J. & Kriegstein, A. R. Single-cell atlas of early human brain development highlights heterogeneity of human neuroepithelial cells and early radial glia. Nat. Neurosci. 24, 584–594 (2021).

10. Cheng, M. et al. Spatially resolved transcriptomics: a comprehensive review of their technological advances, applications, and challenges. J. Genet. Genomics 50, 625–640 (2023).

11. He, S. et al. High-plex imaging of RNA and proteins at subcellular resolution in fixed tissue by spatial molecular imaging. Nat. Biotechnol. 40, 1794–1806 (2022).

12. Li, X. et al. STellaris: a web server for accurate spatial mapping of single cells based on spatial transcriptomics data. Nucleic Acids Res. 51, W560–W568 (2023).

13. Lopez Janeiro, A., et al. Spatially resolved tissue imaging to analyze the tumor immune microenvironment: beyond cell-type densities. J. Immunother. Cancer 12, e008589 (2024).

14. Vu, T. et al. SPF: A spatial and functional data analytic approach to cell imaging data. PLOS Comput. Biol. 18, e1009486 (2022).

15. Baddeley, A., Rubak, E. & Turner, R. Spatial Point Patterns: Methodology and Applications with R. (Chapman and Hall/CRC Press, London, 2015).

16. Baddeley, A. & Turner, R. spatstat: An R Package for Analyzing Spatial Point Patterns. J. Stat. Softw. 12, 1–42 (2005).

17. Ding, D. Y. et al. Quantitative characterization of tissue states using multiomics and ecological spatial analysis. Nat. Genet. 57, 910–921 (2025).

18. Hanahan, D. & Weinberg, R. A. Hallmarks of Cancer: The Next Generation. Cell 144, 646–674 (2011).

19. Mao, Y., Xia, Z., Xia, W. & Jiang, P. Metabolic reprogramming, sensing, and cancer therapy. Cell Rep. 43, 115064 (2024).

20. Engelhardt, B., Vajkoczy, P. & Weller, R. O. The movers and shapers in immune privilege of the CNS. Nat. Immunol. 18, 123–131 (2017).

21. Qiu, R., Zhong, Y., Li, Q., Li, Y. & Fan, H. Metabolic Remodeling in Glioma Immune Microenvironment: Intercellular Interactions Distinct From Peripheral Tumors. Front. Cell Dev. Biol. 9, (2021).

22. Broekman, M. L. et al. Multidimensional communication in the microenvirons of glioblastoma. Nat. Rev. Neurol. 14, 482–495 (2018).

23. Migliozzi, S. et al. Restraint of cancer cell plasticity by spatial homotypic clustering. Cancer Cell 43, 2206–2223.e10 (2025).

24. Nomura, M. et al. The multilayered transcriptional architecture of glioblastoma ecosystems. Nat. Genet. 57, 1155–1167 (2025).

25. Program, C. S.-C. B. et al. CZ CELL×GENE Discover: A single-cell data platform for scalable exploration, analysis and modeling of aggregated data. 2023.10.30.563174 Preprint at 10.1101/2023.10.30.563174 (2023).

26. Ruiz-Moreno, C. et al. Harmonized single-cell landscape, intercellular crosstalk and tumor architecture of glioblastoma. 2022.08.27.505439 Preprint at 10.1101/2022.08.27.505439 (2022).

27. Domínguez Conde, C., et al. Cross-tissue immune cell analysis reveals tissue-specific features in humans. Science 376, eabl5197 (2022).

28. Xu, C. et al. Automatic cell-type harmonization and integration across Human Cell Atlas datasets. Cell 186, 5876–5891.e20 (2023).

29. Barbie, D. A. et al. Systematic RNA interference reveals that oncogenic KRAS-driven cancers require TBK1. Nature 462, 108–112 (2009).

30. Subramanian, A. et al. Gene set enrichment analysis: A knowledge-based approach for interpreting genome-wide expression profiles. Proc. Natl. Acad. Sci. 102, 15545–15550 (2005).

31. Kanehisa, M., Furumichi, M., Sato, Y., Kawashima, M. & Ishiguro-Watanabe, M. KEGG for taxonomy-based analysis of pathways and genomes. Nucleic Acids Res. 51, D587–D592 (2023).

32. Kanehisa, M. & Goto, S. KEGG: Kyoto Encyclopedia of Genes and Genomes. Nucleic Acids Res. 28, 27–30 (2000).

33. He, S. et al. High-plex imaging of RNA and proteins at subcellular resolution in fixed tissue by spatial molecular imaging. Nat. Biotechnol. 40, 1794–1806 (2022).

34. Lindgren, F. Fmesher: Triangle Meshes and Related Geometry Tools. (2026).

35. Bachl, F. E., Lindgren, F., Borchers, D. L. & Illian, J. B. inlabru: an R package for Bayesian spatial modelling from ecological survey data. Methods Ecol. Evol. 10, 760–766 (2019).

36. Xu, C. et al. Probabilistic harmonization and annotation of single-cell transcriptomics data with deep generative models. Mol. Syst. Biol. 17, MSB20209620 (2021).

37. Lotfollahi, M. et al. Mapping single-cell data to reference atlases by transfer learning. Nat. Biotechnol. 40, 121–130 (2022).

38. R Core Team. R: A Language and Environment for Statistical Computing. (R Foundation for Statistical Computing, Vienna, Austria, 2025).

39. QuantStack and Mamba Contributors. mamba. (2019).

40. Baddeley, A., Turner, R., Mateu, J. & Bevan, A. Hybrids of Gibbs Point Process Models and Their Implementation. J. Stat. Softw. 55, 1–43 (2013).

41. Wickham, H. et al. Welcome to the tidyverse. J. Open Source Softw. 4, 1686 (2019).

42. Barrett, T. et al. Data.Table: Extension of ‘data.Framè. (2026).

43. Pebesma, E. & Bivand, R. Spatial Data Science: With Applications in R. (Chapman and Hall/CRC, New York, 2023). doi:10.1201/9780429459016.

44. Pebesma, E. The R Journal: Simple Features for R: Standardized Support for Spatial Vector Data. R J. 10, 439–446 (2018).

45. Hijmans, R. J., Brown, A. & Barbosa, M. Terra: Spatial Data Analysis. (2026).

46. Lindgren, F. & Rue, H. Bayesian Spatial Modelling with R-INLA. J. Stat. Softw. 63, 1–25 (2015).

47. Bates, D., Maechler, M. & Jagan, M. Matrix: Sparse and Dense Matrix Classes and Methods. (2026).

48. Wickham, H. Data Analysis. in ggplot2: Elegant Graphics for Data Analysis (ed. Wickham, H.) 189–201 (Springer International Publishing, Cham, 2016). doi:10.1007/978-3-319-24277-4_9.

49. Pedersen, T. L. Patchwork: The Composer of Plots. (2025).

50. Neuwirth, E. RColorBrewer: ColorBrewer Palettes. (2022).

51. Yu, G. Thirteen years of clusterProfiler. The Innovation 5, 100722 (2024).

52. Xu, S. et al. Using clusterProfiler to characterize multiomics data. Nat. Protoc. 19, 3292–3320 (2024).

53. Wu, T. et al. clusterProfiler 4.0: A universal enrichment tool for interpreting omics data. The Innovation 2, 100141 (2021).

54. Yu, G., Wang, L.-G., Han, Y. & He, Q.-Y. clusterProfiler: an R package for comparing biological themes among gene clusters. OMICS J. Integr. Biol. 16, 284–287 (2012).

55. Tenenbaum, D. & Maintainer, B. P. KEGGREST: Client-Side REST Access to the Kyoto Encyclopedia of Genes and Genomes (KEGG). (2025). doi:10.18129/B9.bioc.KEGGREST.

56. Hänzelmann, S., Castelo, R. & Guinney, J. GSVA: gene set variation analysis for microarray and RNA-Seq data. BMC Bioinformatics 14, 7 (2013).

57. van Rossum, G. & Drake, F. L. Python Reference Manual.

58. team, T. pandas development. pandas-dev/pandas: Pandas. Zenodo 10.5281/zenodo.3509134 (2020).

59. Harris, C. R. et al. Array programming with NumPy. Nature 585, 357–362 (2020).

60. Virtanen, P. et al. SciPy 1.0: Fundamental Algorithms for Scientific Computing in Python. Nat. Methods 17, 261–272 (2020).

61. Richardson, N., et al. Arrow: Integration to ‘Apache’ ‘Arrow’. (2026).

62. Pedregosa, F. et al. Scikit-learn: Machine Learning in Python. J Mach Learn Res 12, 2825–2830 (2011).

63. Chen, T. & Guestrin, C. XGBoost: A Scalable Tree Boosting System. in Proceedings of the 22nd ACM SIGKDD International Conference on Knowledge Discovery and Data Mining 785–794 (Association for Computing Machinery, New York, NY, USA, 2016). doi:10.1145/2939672.2939785.

64. Ansel, J. et al. PyTorch 2: Faster Machine Learning Through Dynamic Python Bytecode Transformation and Graph Compilation. in Proceedings of the 29th ACM International Conference on Architectural Support for Programming Languages and Operating Systems, Volume 2 929–947 (Association for Computing Machinery, New York, NY, USA, 2024). doi:10.1145/3620665.3640366.

65. Virshup, I., Rybakov, S., Theis, F. J., Angerer, P. & Wolf, F. A. anndata: Access and store annotated data matrices. J. Open Source Softw. 9, 4371 (2024).

66. Wolf, F. A., Angerer, P. & Theis, F. J. SCANPY: large-scale single-cell gene expression data analysis. Genome Biol. 19, 15 (2018).

67. Gayoso, A. et al. A Python library for probabilistic analysis of single-cell omics data. Nat. Biotechnol. 40, 163–166 (2022).

68. Traag, V. A., Waltman, L. & van Eck, N. J. From Louvain to Leiden: guaranteeing well-connected communities. Sci. Rep. 9, 5233 (2019).

