## Supplementary Figures 1 for "Point-Process Modelling of Cell-Type Interaction Structure in CosMx Glioma Data for Hypothesis-Driven Tumor Inference from scRNA-seq"

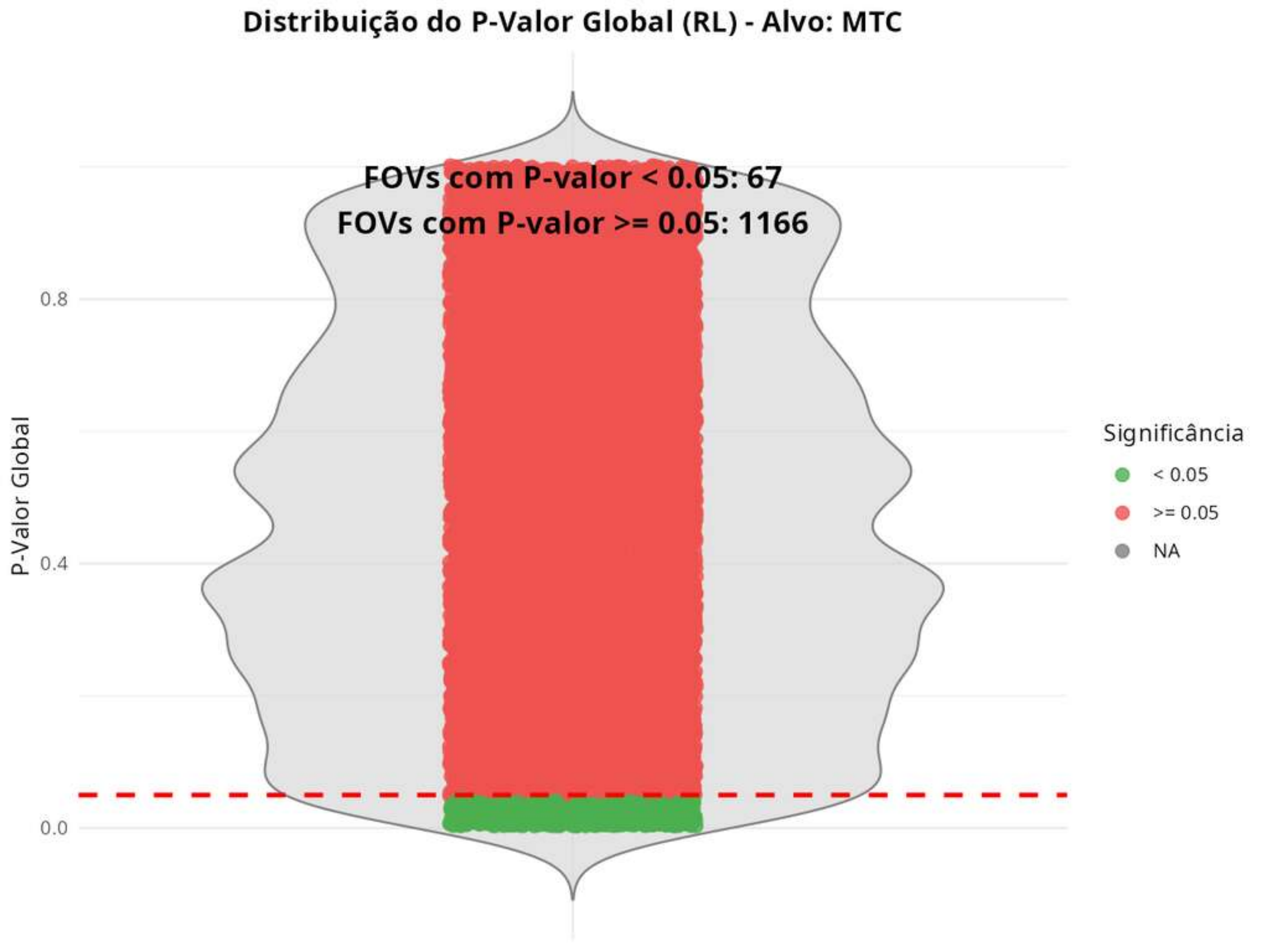
Distribuição do P-Valor Global (RL) - Alvo: MTC
FOVs com P-valor < 0.05: 67
FOVs com P-valor >= 0.05: 1166
P-Valor Global
0.8
0.4
0.0
Significância
< 0.05
>= 0.05
NA

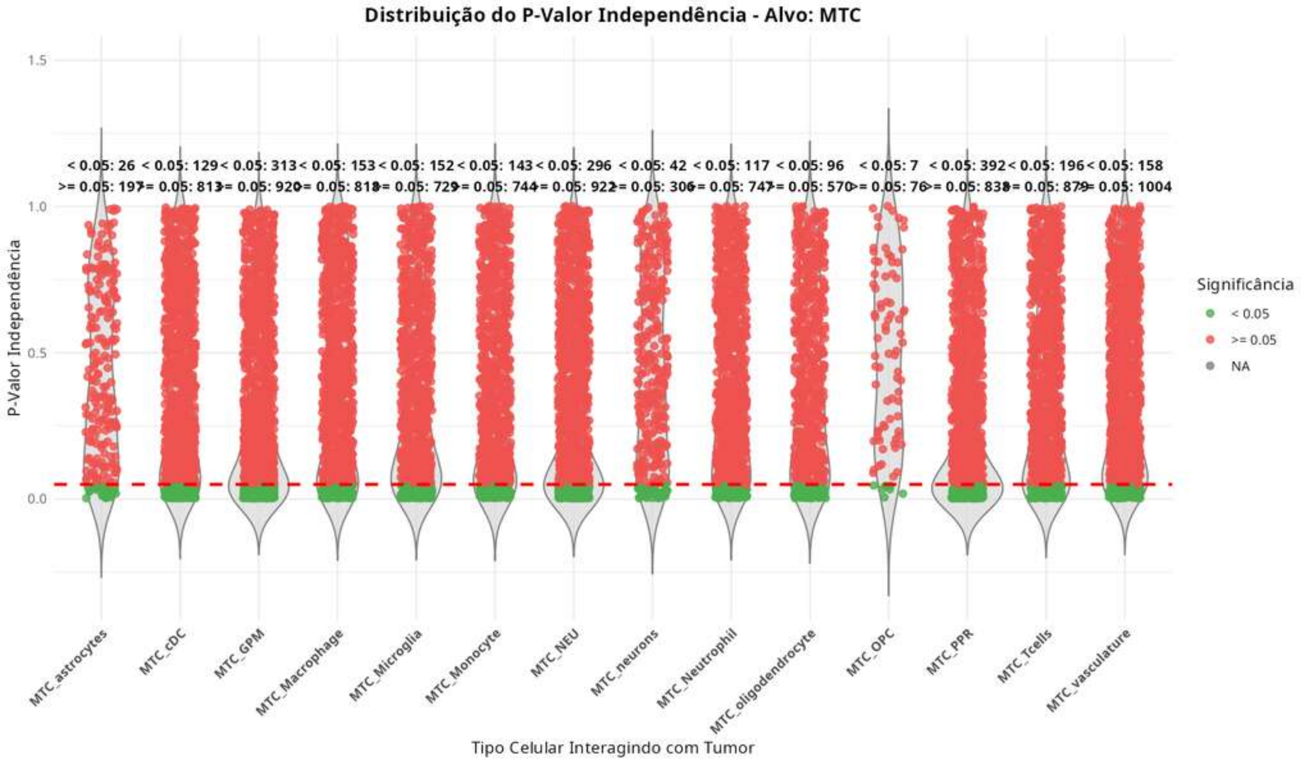

Distribuição do P-Valor Independência - Alvo: MTC
< 0.05: 26
>= 0.05: 197
< 0.05: 129
>= 0.05: 813
< 0.05: 313
>= 0.05: 920
< 0.05: 153
>= 0.05: 818
< 0.05: 152
>= 0.05: 729
< 0.05: 143
>= 0.05: 744
< 0.05: 296
>= 0.05: 922
< 0.05: 42
>= 0.05: 306
< 0.05: 117
>= 0.05: 747
< 0.05: 96
>= 0.05: 570
< 0.05: 7
>= 0.05: 76
< 0.05: 392
>= 0.05: 838
< 0.05: 196
>= 0.05: 879
< 0.05: 158
>= 0.05: 1004
P-Valor Independência
1.5
1.0
0.5
0.0
MTC_astrocytes
MTC_cDC
MTC_GPM
MTC_Macrophage
MTC_Microglia
MTC_Monocyte
MTC_NEU
MTC_neurons
MTC_Neutrophil
MTC_oligodendrocyte
MTC_OPC
MTC_PPR
MTC_Tcells
MTC_vasculature
Tipo Celular Interagindo com Tumor
Significância
< 0.05
>= 0.05
NA

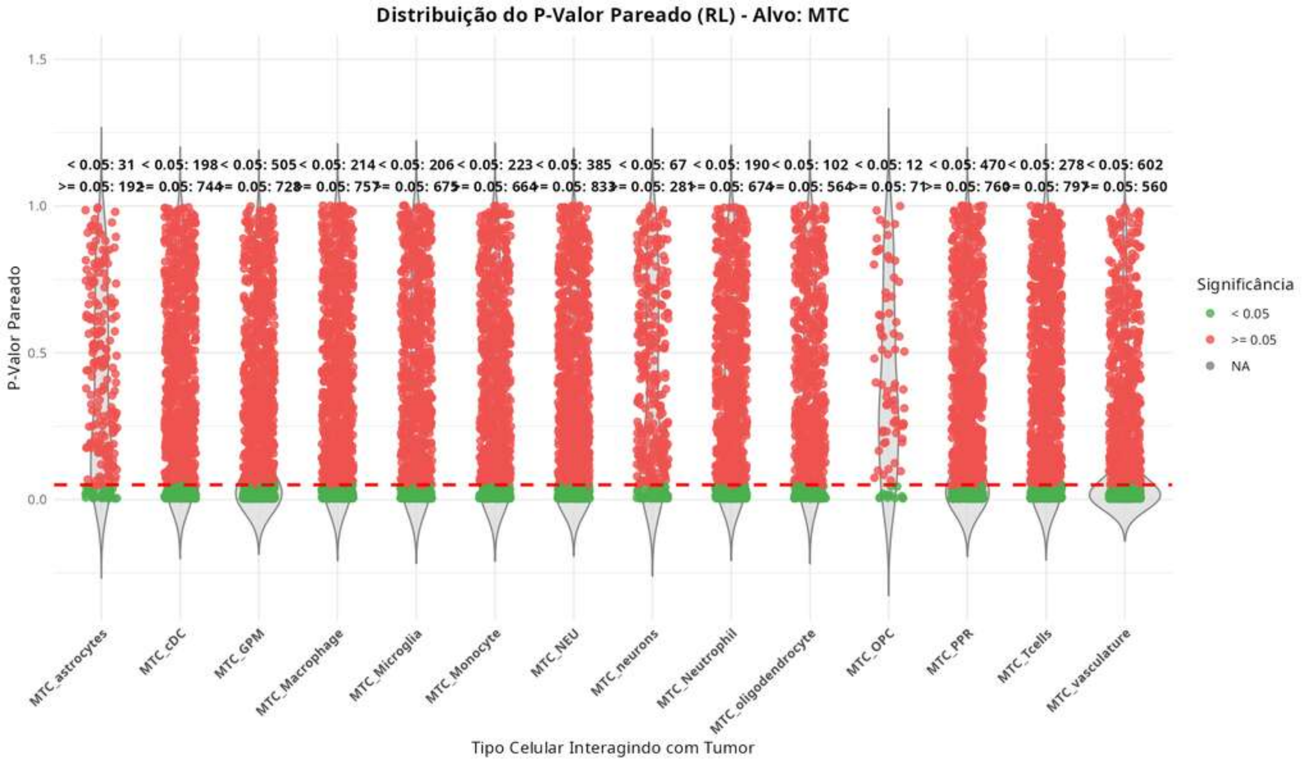

Distribuição do P-Valor Pareado (RL) - Alvo: MTC
P-Valor Pareado
1.5
1.0
0.5
0.0
< 0.05: 31
>= 0.05: 192
< 0.05: 198
>= 0.05: 744
< 0.05: 505
>= 0.05: 728
< 0.05: 214
>= 0.05: 757
< 0.05: 206
>= 0.05: 675
< 0.05: 223
>= 0.05: 664
< 0.05: 385
>= 0.05: 833
< 0.05: 67
>= 0.05: 281
< 0.05: 190
>= 0.05: 674
< 0.05: 102
>= 0.05: 564
< 0.05: 12
>= 0.05: 71
< 0.05: 470
>= 0.05: 760
< 0.05: 278
>= 0.05: 797
< 0.05: 602
>= 0.05: 560
MTC_astrocytes
MTC_cDC
MTC_GPM
MTC_Macrophage
MTC_Microglia
MTC_Monocyte
MTC_NEU
MTC_neurons
MTC_Neutrophil
MTC_oligodendrocyte
MTC_OPC
MTC_PPR
MTC_Tcells
MTC_vasculature
Tipo Celular Interagindo com Tumor
Significância
< 0.05
>= 0.05
NA

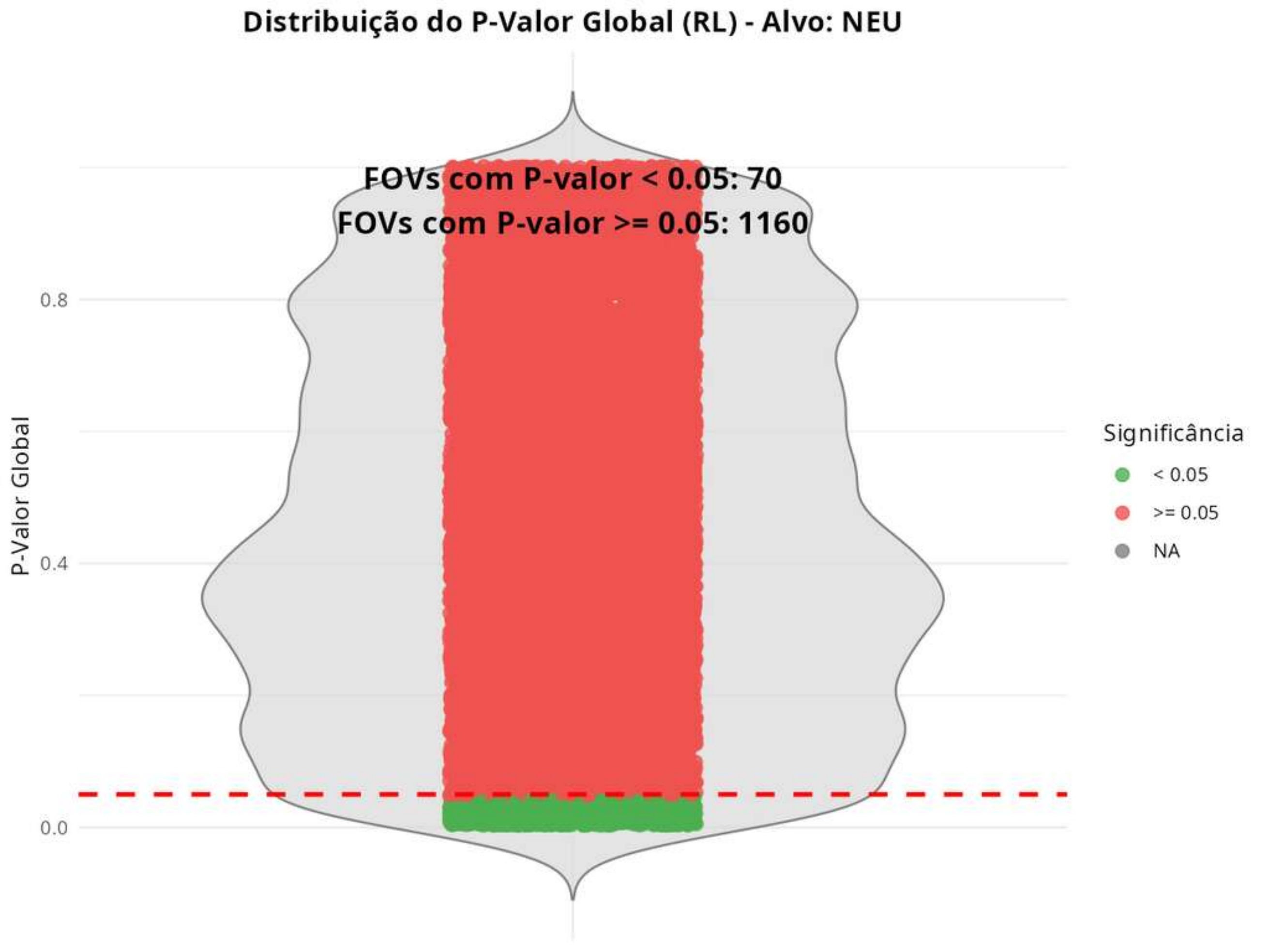
Distribuição do P-Valor Global (RL) - Alvo: NEU
FOVs com P-valor < 0.05: 70
FOVs com P-valor >= 0.05: 1160
P-Valor Global
0.8
0.4
0.0
Significância
< 0.05
>= 0.05
NA

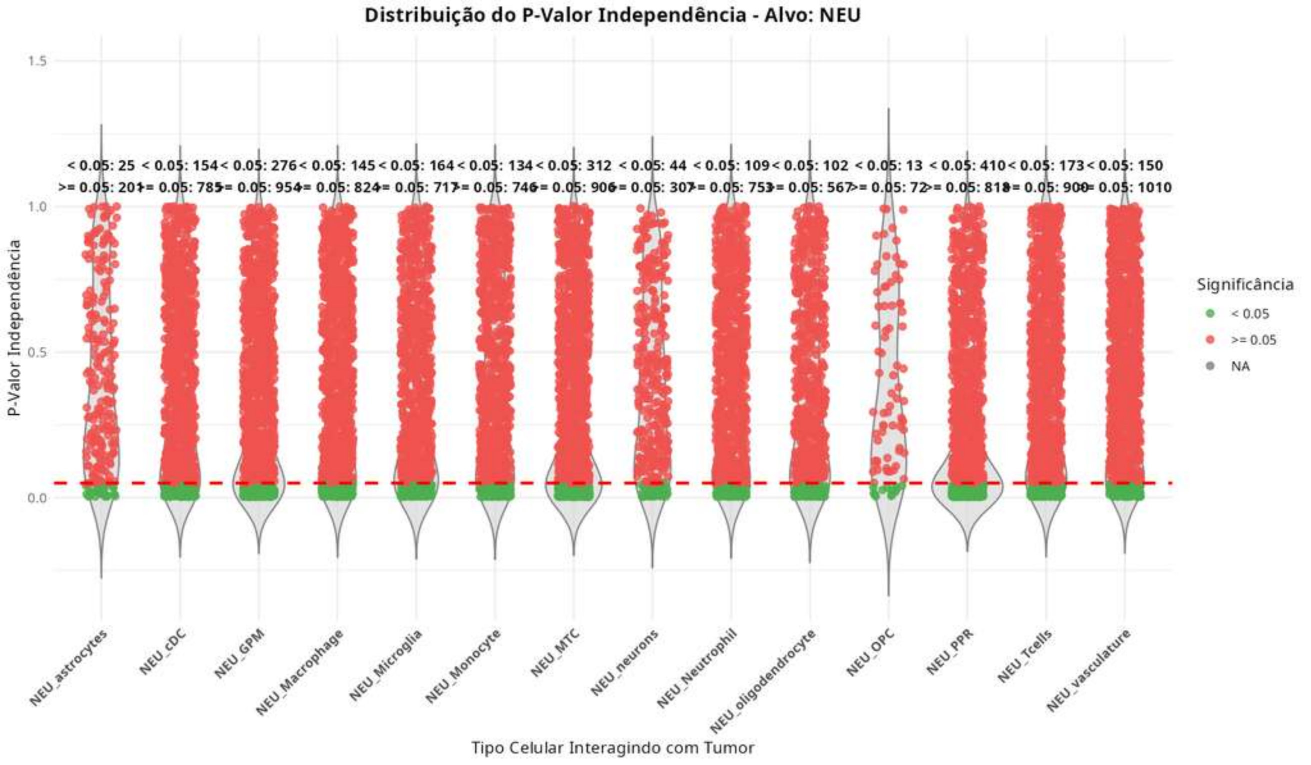
Distribuição do P-Valor Independência - Alvo: NEU
P-Valor Independência
1.5
1.0
0.5
0.0
< 0.05: 25
>= 0.05: 201
< 0.05: 154
>= 0.05: 785
< 0.05: 276
>= 0.05: 954
< 0.05: 145
>= 0.05: 824
< 0.05: 164
>= 0.05: 717
< 0.05: 134
>= 0.05: 746
< 0.05: 312
>= 0.05: 906
< 0.05: 44
>= 0.05: 307
< 0.05: 109
>= 0.05: 753
< 0.05: 102
>= 0.05: 567
< 0.05: 13
>= 0.05: 72
< 0.05: 410
>= 0.05: 818
< 0.05: 173
>= 0.05: 900
< 0.05: 150
>= 0.05: 1010
NEU_astrocytes
NEU_cDC
NEU_GPM
NEU_Macrophage
NEU_Microglia
NEU_Monocyte
NEU_MTC
NEU_neurons
NEU_Neutrophil
NEU_oligodendrocyte
NEU_OPC
NEU_PPR
NEU_Tcells
NEU_vasculature
Tipo Celular Interagindo com Tumor
Significância
< 0.05
>= 0.05
NA

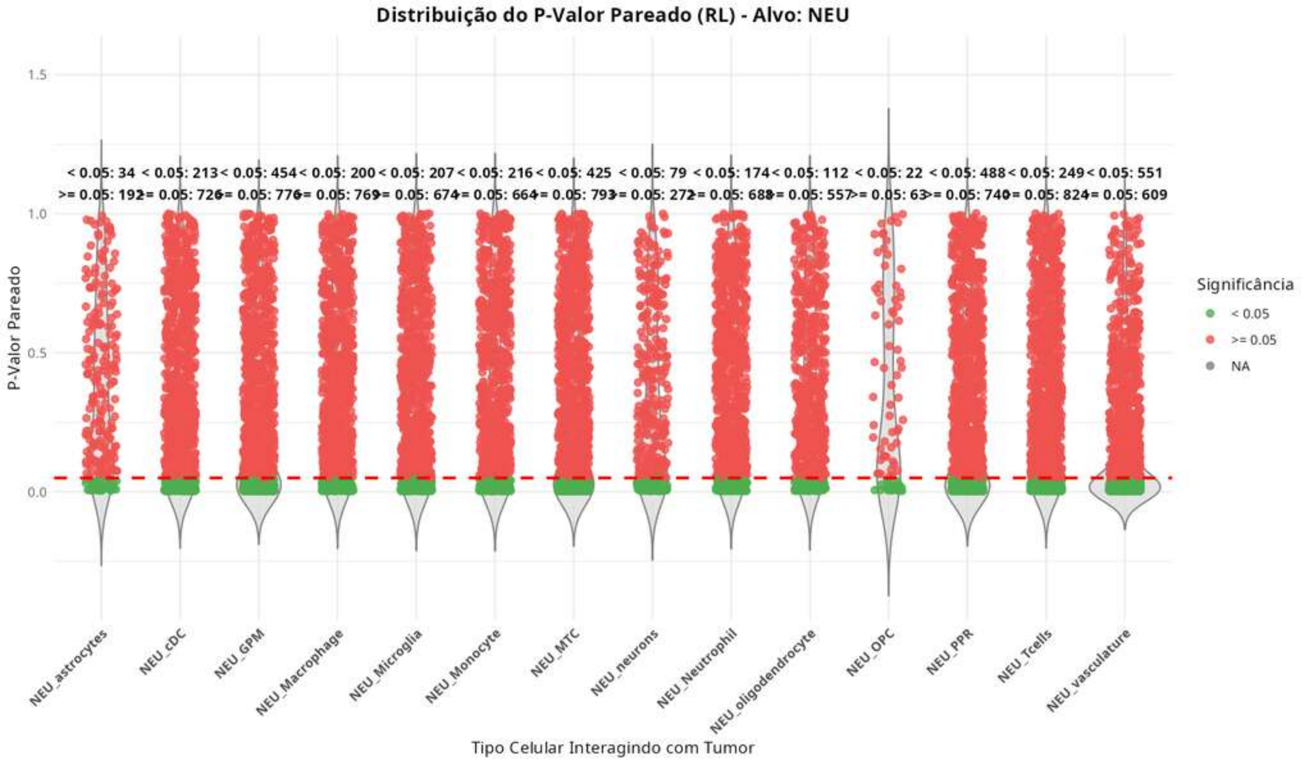

Distribuição do P-Valor Pareado (RL) - Alvo: NEU
P-Valor Pareado
1.5
1.0
0.5
0.0
< 0.05: 34
>= 0.05: 192
< 0.05: 213
>= 0.05: 726
< 0.05: 454
>= 0.05: 776
< 0.05: 200
>= 0.05: 769
< 0.05: 207
>= 0.05: 674
< 0.05: 216
>= 0.05: 664
< 0.05: 425
>= 0.05: 793
< 0.05: 79
>= 0.05: 272
< 0.05: 174
>= 0.05: 688
< 0.05: 112
>= 0.05: 557
< 0.05: 22
>= 0.05: 63
< 0.05: 488
>= 0.05: 740
< 0.05: 249
>= 0.05: 824
< 0.05: 551
>= 0.05: 609
NEU_astrocytes
NEU_cDC
NEU_GPM
NEU_Macrophage
NEU_Microglia
NEU_Monocyte
NEU_MTC
NEU_neurons
NEU_Neutrophil
NEU_oligodendrocyte
NEU_OPC
NEU_PPR
NEU_Tcells
NEU_vasculature
Tipo Celular Interagindo com Tumor
Significância
< 0.05
>= 0.05
NA

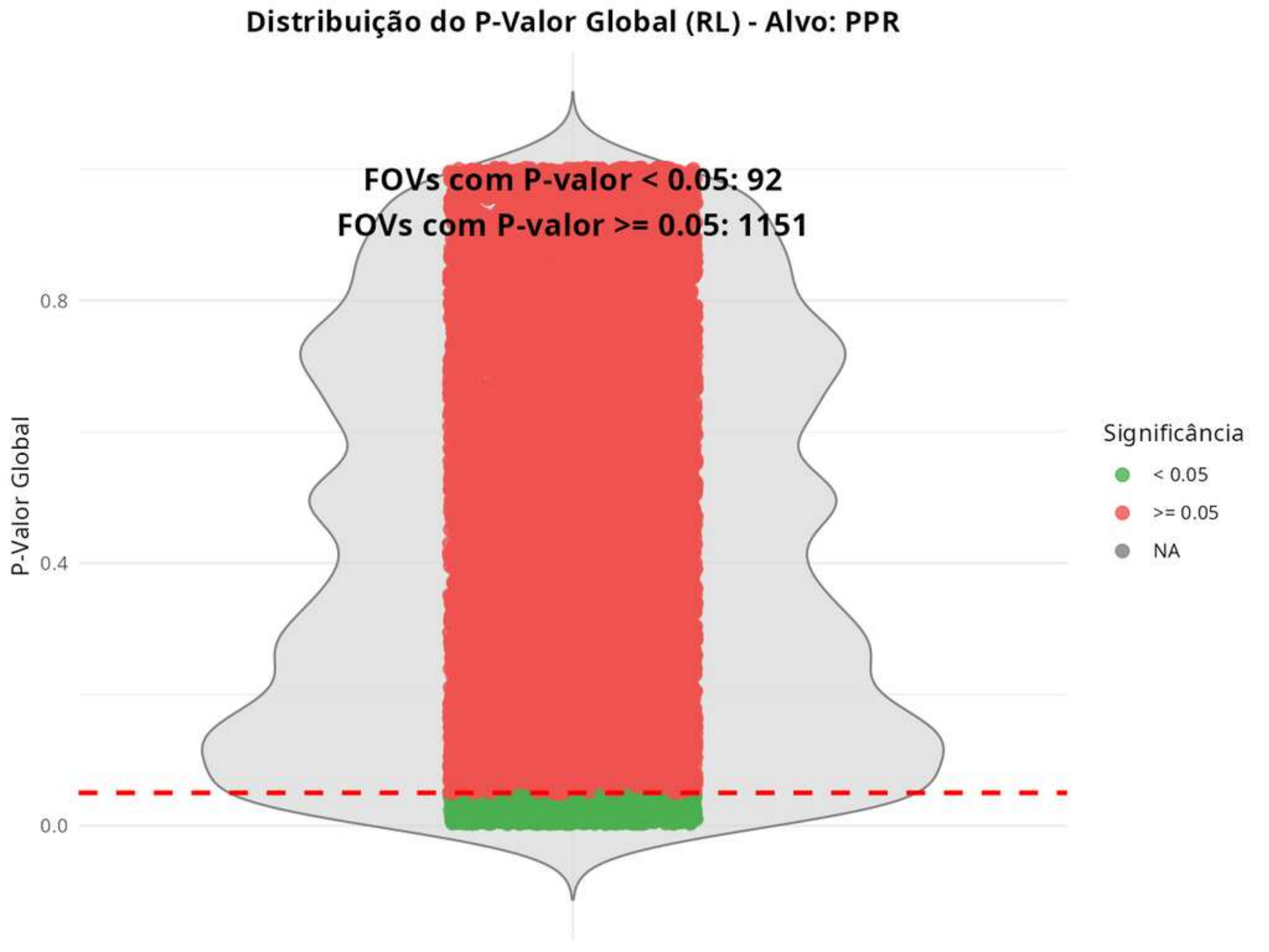
Distribuição do P-Valor Global (RL) - Alvo: PPR
FOVs com P-valor < 0.05: 92
FOVs com P-valor >= 0.05: 1151
P-Valor Global
0.8
0.4
0.0
Significância
< 0.05
>= 0.05
NA

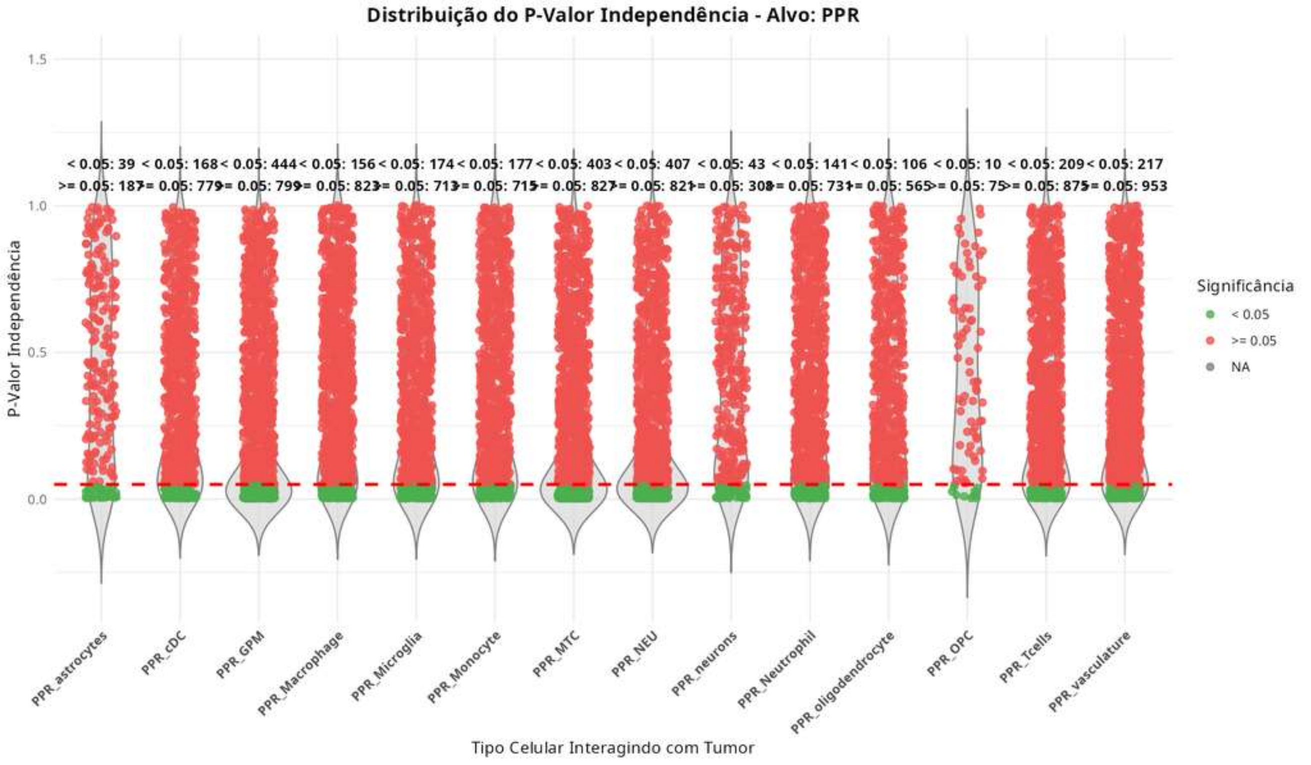
Distribuição do P-Valor Independência - Alvo: PPR
P-Valor Independência
1.5
1.0
0.5
0.0
< 0.05: 39
>= 0.05: 187
< 0.05: 168
>= 0.05: 779
< 0.05: 444
>= 0.05: 799
< 0.05: 156
>= 0.05: 823
< 0.05: 174
>= 0.05: 713
< 0.05: 177
>= 0.05: 715
< 0.05: 403
>= 0.05: 827
< 0.05: 407
>= 0.05: 821
< 0.05: 43
>= 0.05: 308
< 0.05: 141
>= 0.05: 731
< 0.05: 106
>= 0.05: 565
< 0.05: 10
>= 0.05: 75
< 0.05: 209
>= 0.05: 875
< 0.05: 217
>= 0.05: 953
PPR_astrocytes
PPR_cDC
PPR_GPM
PPR_Macrophage
PPR_Microglia
PPR_Monocyte
PPR_MTC
PPR_NEU
PPR_neurons
PPR_Neutrophil
PPR_oligodendrocyte
PPR_OPC
PPR_Tcells
PPR_vasculature
Tipo Celular Interagindo com Tumor
Significância
< 0.05
>= 0.05
NA

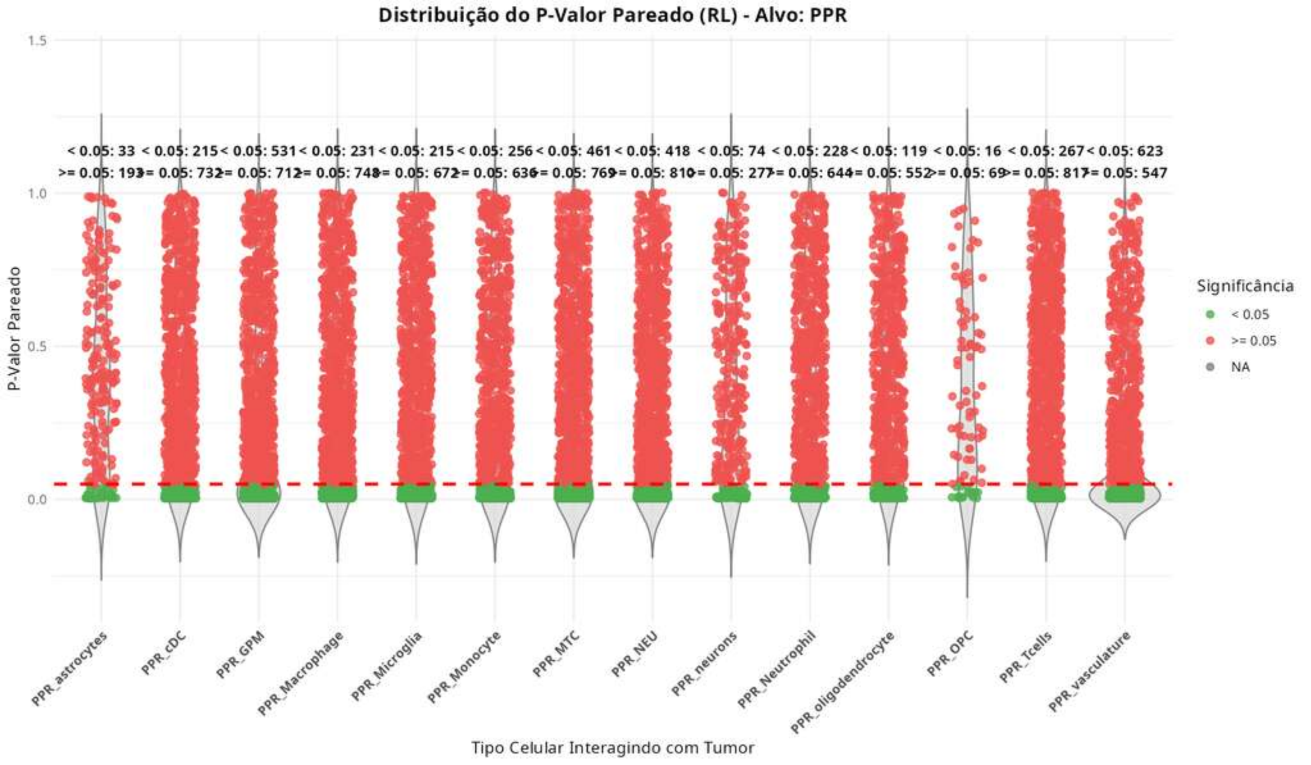

Distribuição do P-Valor Pareado (RL) - Alvo: PPR
P-Valor Pareado
1.5
1.0
0.5
0.0
< 0.05: 33
>= 0.05: 193
< 0.05: 215
>= 0.05: 732
< 0.05: 531
>= 0.05: 712
< 0.05: 231
>= 0.05: 748
< 0.05: 215
>= 0.05: 672
< 0.05: 256
>= 0.05: 636
< 0.05: 461
>= 0.05: 769
< 0.05: 418
>= 0.05: 810
< 0.05: 74
>= 0.05: 277
< 0.05: 228
>= 0.05: 644
< 0.05: 119
>= 0.05: 552
< 0.05: 16
>= 0.05: 69
< 0.05: 267
>= 0.05: 817
< 0.05: 623
>= 0.05: 547
PPR_astrocytes
PPR_cDC
PPR_GPM
PPR_Macrophage
PPR_Microglia
PPR_Monocyte
PPR_MTC
PPR_NEU
PPR_neurons
PPR_Neutrophil
PPR_oligodendrocyte
PPR_OPC
PPR_Tcells
PPR_vasculature
Tipo Celular Interagindo com Tumor
Significância
< 0.05
>= 0.05
NA

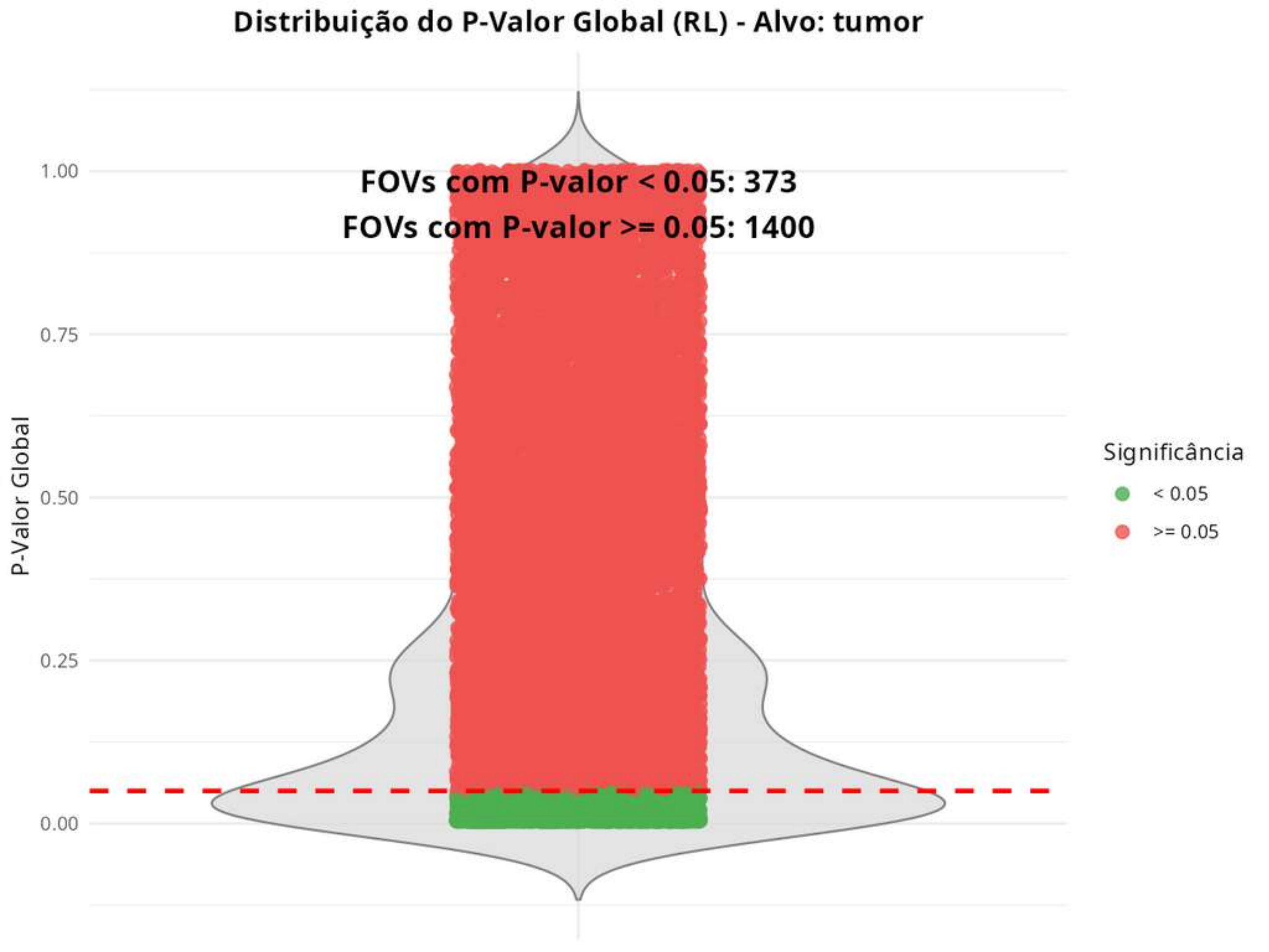
Distribuição do P-Valor Global (RL) - Alvo: tumor
FOVs com P-valor < 0.05: 373
FOVs com P-valor >= 0.05: 1400
P-Valor Global
1.00
0.75
0.50
0.25
0.00
Significância
< 0.05
>= 0.05

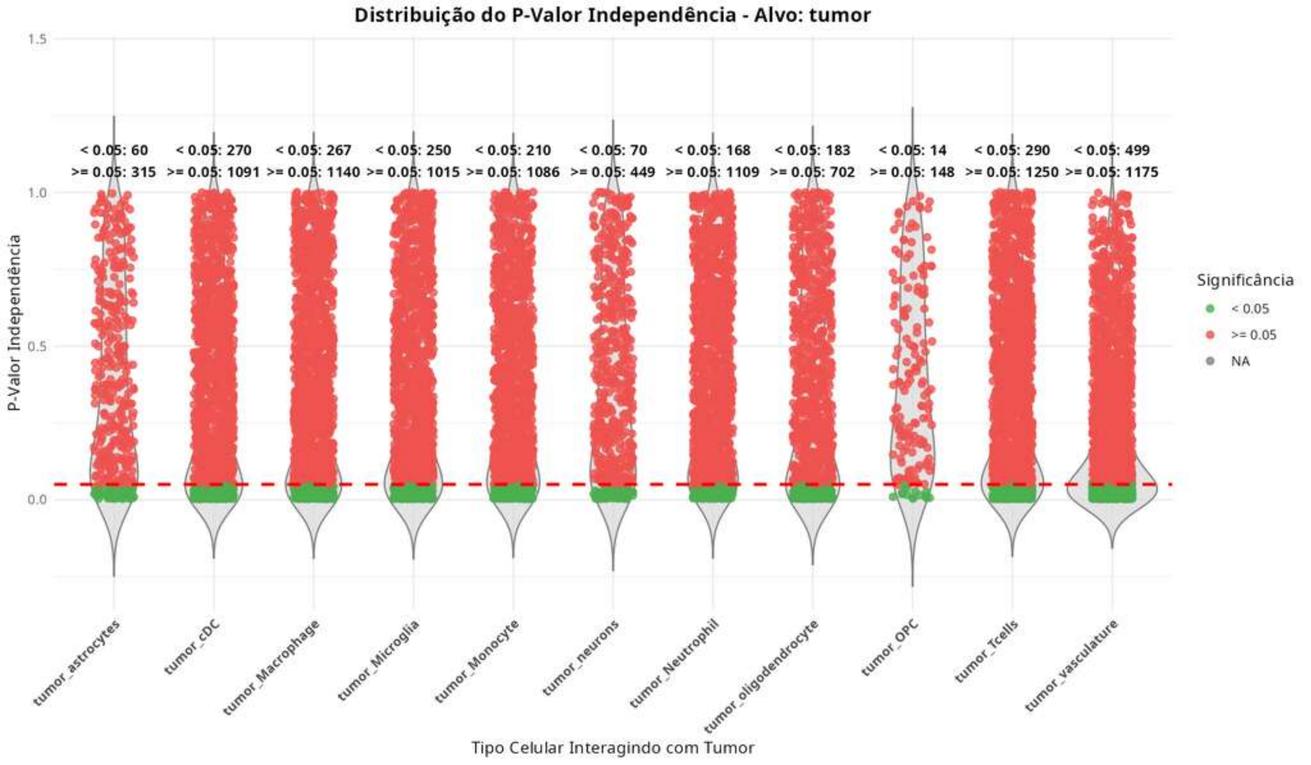
Distribuição do P-Valor Independência - Alvo: tumor
< 0.05: 60
>= 0.05: 315
< 0.05: 270
>= 0.05: 1091
< 0.05: 267
>= 0.05: 1140
< 0.05: 250
>= 0.05: 1015
< 0.05: 210
>= 0.05: 1086
< 0.05: 70
>= 0.05: 449
< 0.05: 168
>= 0.05: 1109
< 0.05: 183
>= 0.05: 702
< 0.05: 14
>= 0.05: 148
< 0.05: 290
>= 0.05: 1250
< 0.05: 499
>= 0.05: 1175
P-Valor Independência
1.5
1.0
0.5
0.0
Significância
< 0.05
>= 0.05
NA
tumor_astrocytes
tumor_cDC
tumor_Macrophage
tumor_Microglia
tumor_Monocyte
tumor_neurons
tumor_Neutrophil
tumor_oligodendrocyte
tumor_OPC
tumor_Tcells
tumor_vasculature
Tipo Celular Interagindo com Tumor

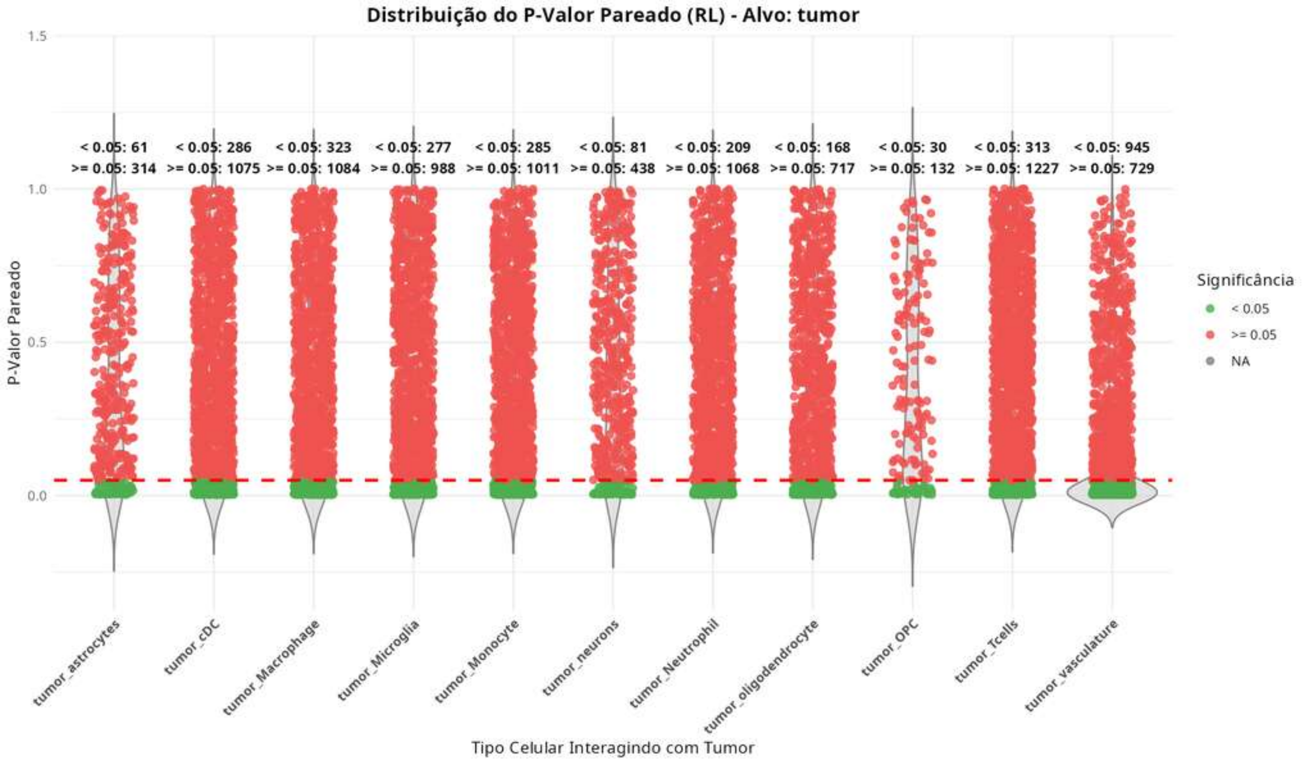
Distribuição do P-Valor Pareado (RL) - Alvo: tumor
P-Valor Pareado
1.5
1.0
0.5
0.0
< 0.05: 61
>= 0.05: 314
< 0.05: 286
>= 0.05: 1075
< 0.05: 323
>= 0.05: 1084
< 0.05: 277
>= 0.05: 988
< 0.05: 285
>= 0.05: 1011
< 0.05: 81
>= 0.05: 438
< 0.05: 209
>= 0.05: 1068
< 0.05: 168
>= 0.05: 717
< 0.05: 30
>= 0.05: 132
< 0.05: 313
>= 0.05: 1227
< 0.05: 945
>= 0.05: 729
tumor_astrocytes
tumor_cDC
tumor_Macrophage
tumor_Microglia
tumor_Monocyte
tumor_neurons
tumor_Neutrophil
tumor_oligodendrocyte
tumor_OPC
tumor_Tcells
tumor_vasculature
Tipo Celular Interagindo com Tumor
Significância
< 0.05
>= 0.05
NA
