## Supplementary figures and images for "Point-Process Modelling of Cell-Type Interaction Structure in CosMx Glioma Data for Hypothesis-Driven Tumor Inference from scRNA-seq"

### Supplementary Figures 2

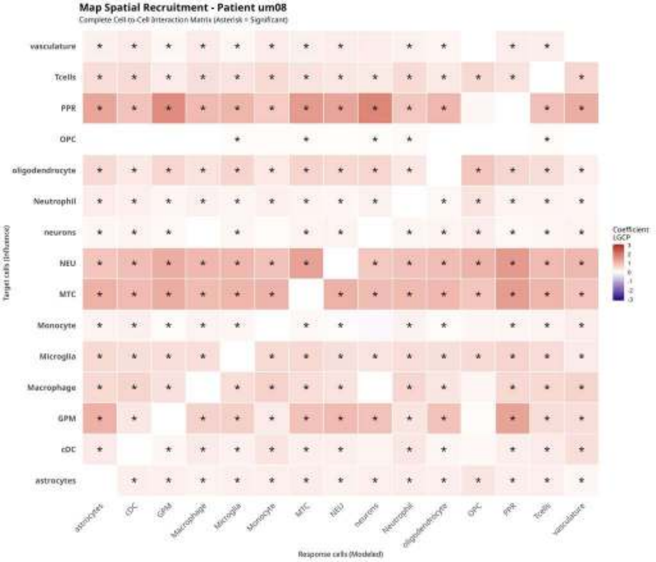

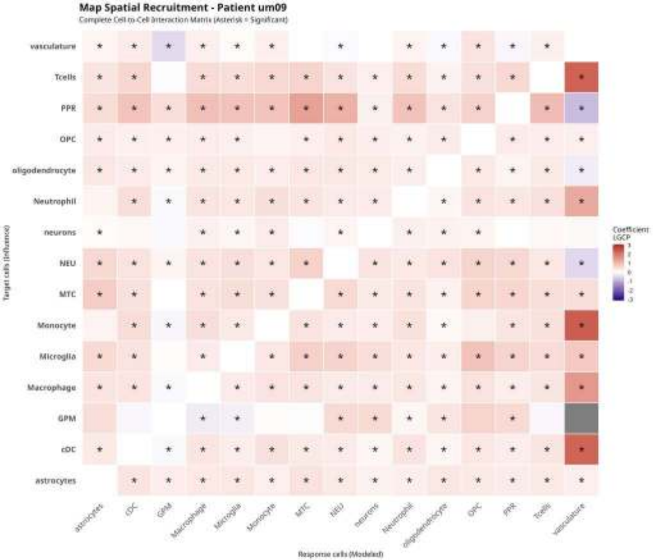

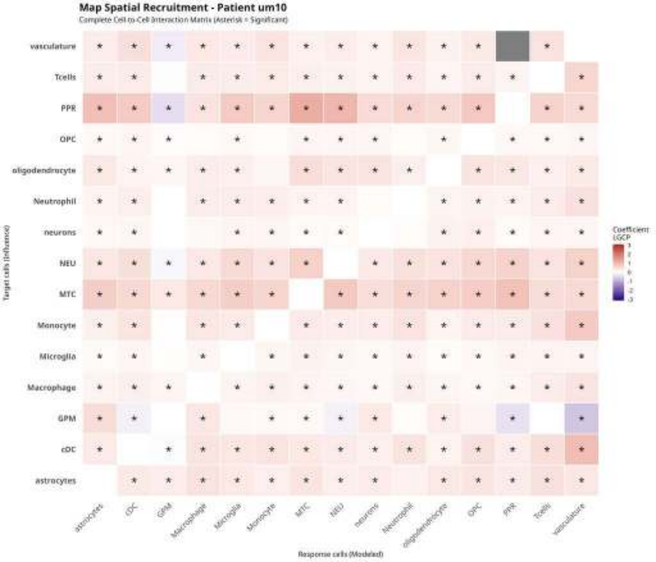

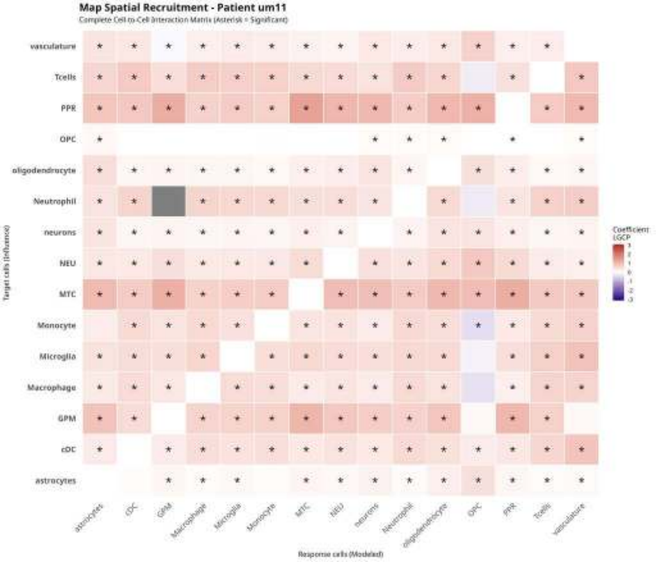

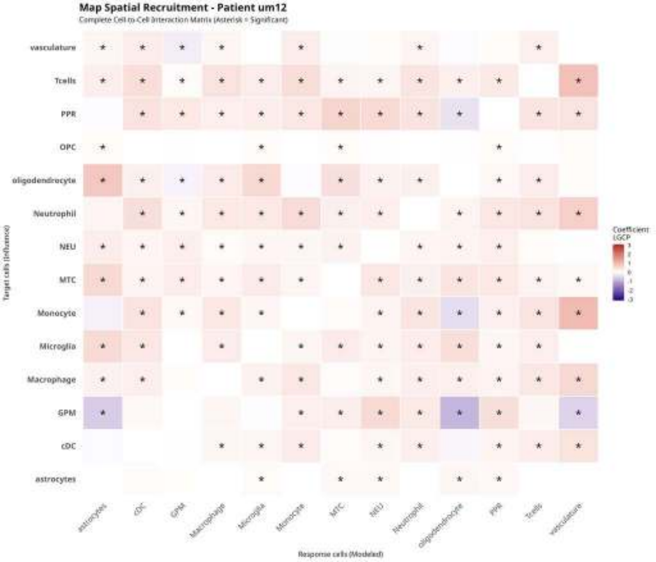

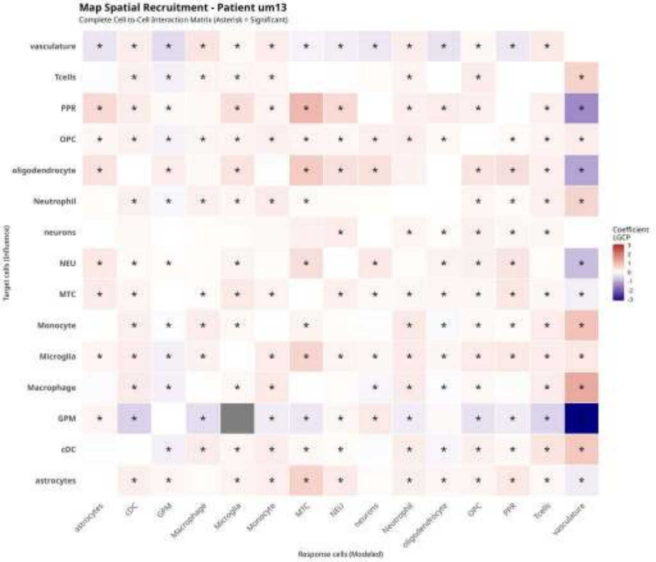

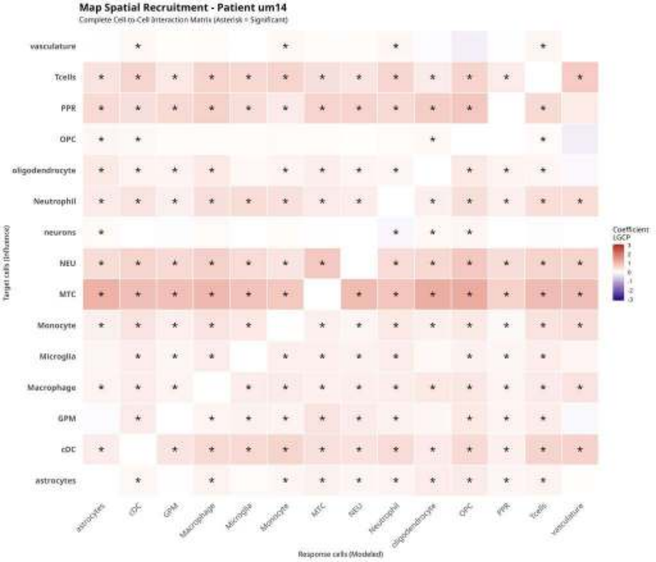

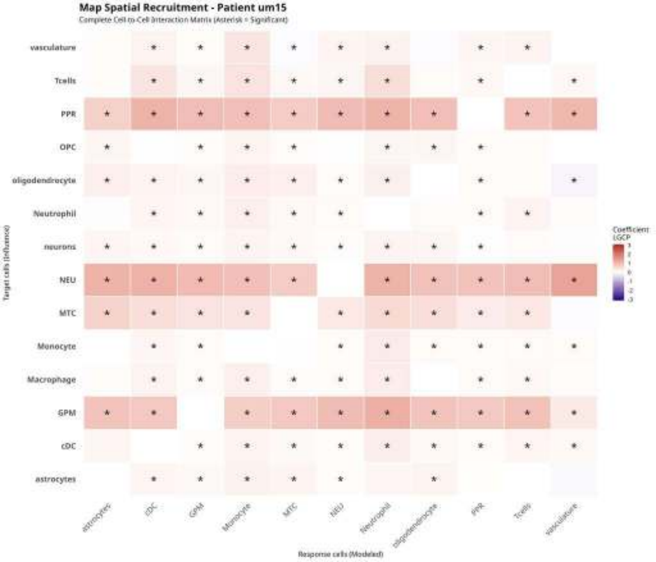
