## Supplementary Figures 3 for "Point-Process Modelling of Cell-Type Interaction Structure in CosMx Glioma Data for Hypothesis-Driven Tumor Inference from scRNA-seq"

#### Density of Monocyte on um08

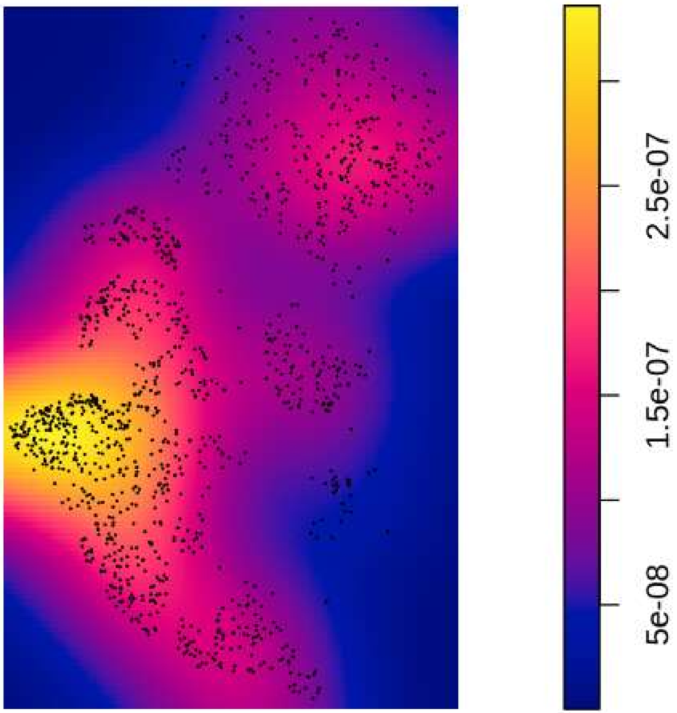

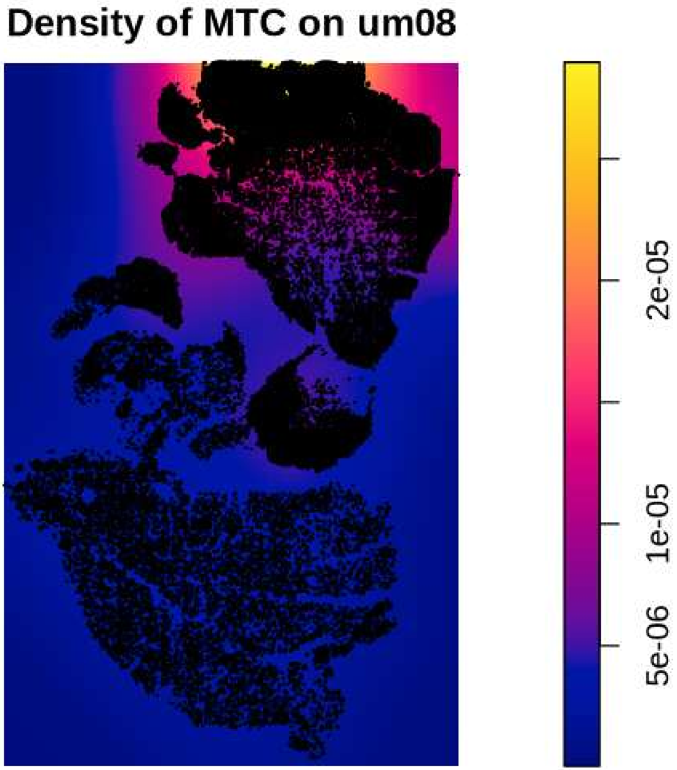
Density of MTC on um08
2e-05
1e-05
5e-06

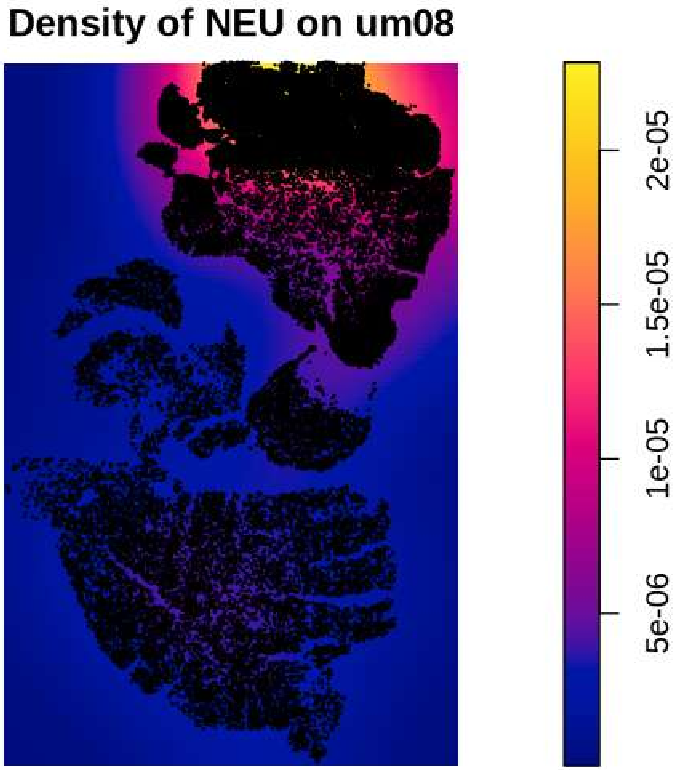
Density of NEU on um08
2e-05
1.5e-05
1e-05
5e-06

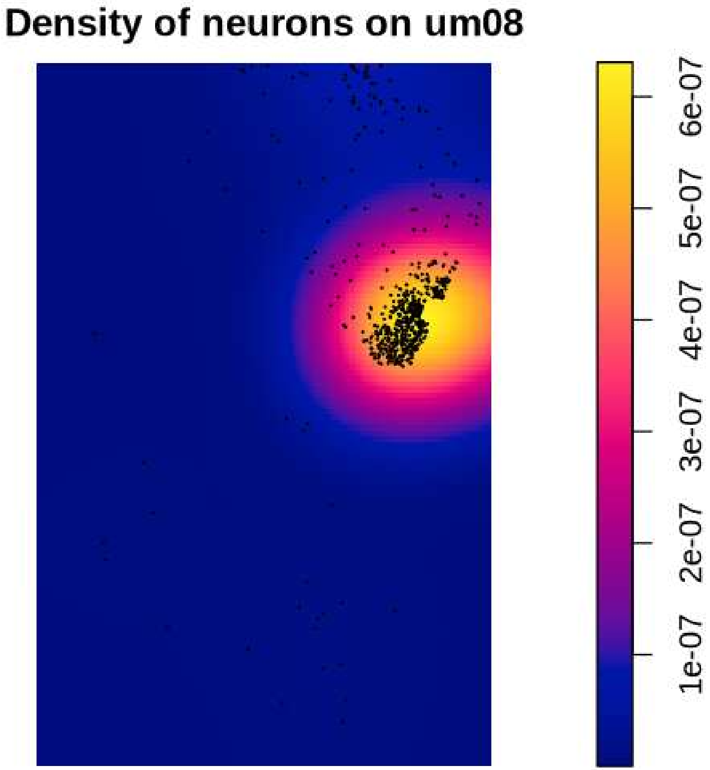
Density of neurons on um08
1e-07
2e-07
3e-07
4e-07
5e-07
6e-07

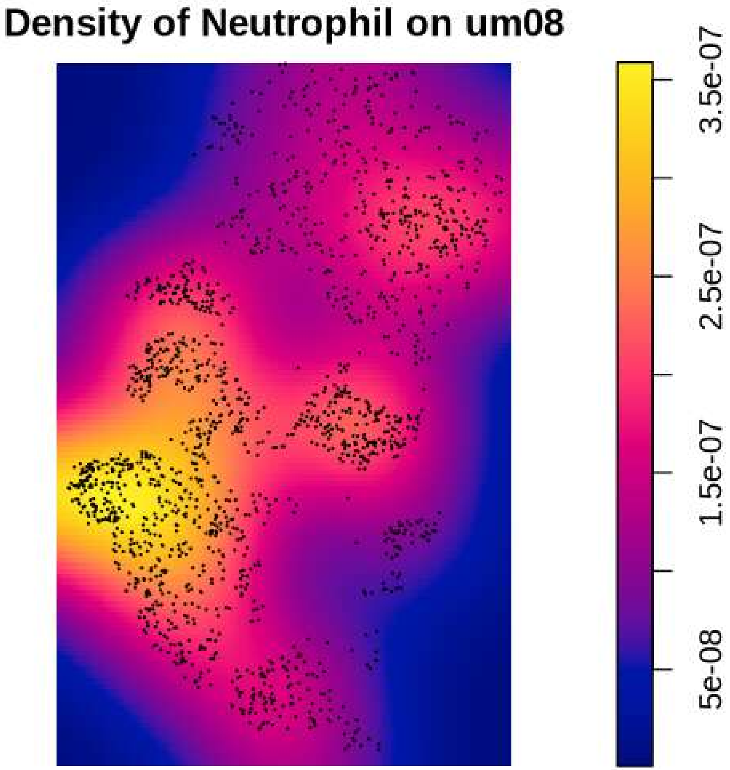
Density of Neutrophil on um08
3.5e-07
2.5e-07
1.5e-07
5e-08

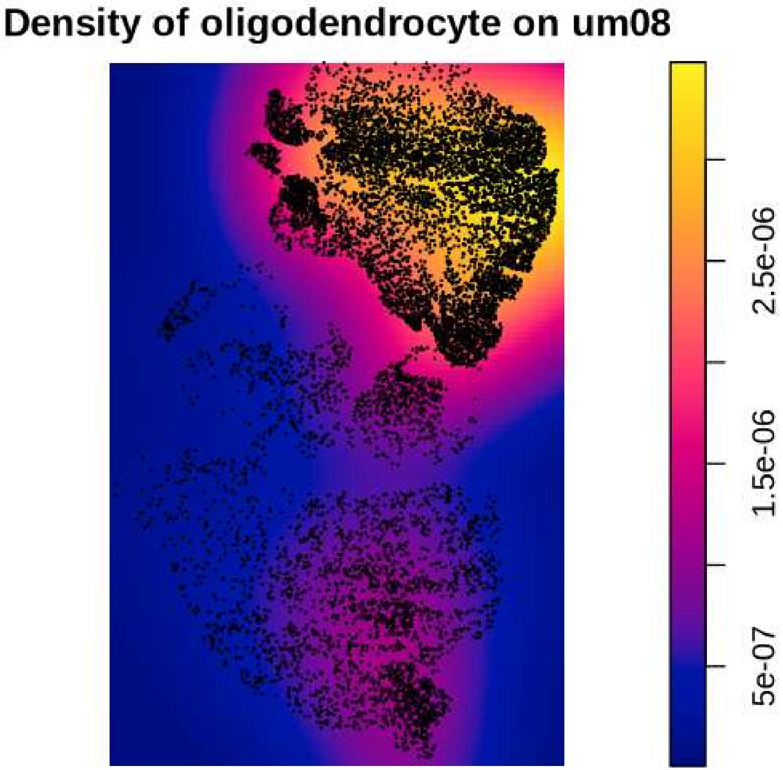
Density of oligodendrocyte on um08
2.5e-06
1.5e-06
5e-07

#### Density of OPC on um08

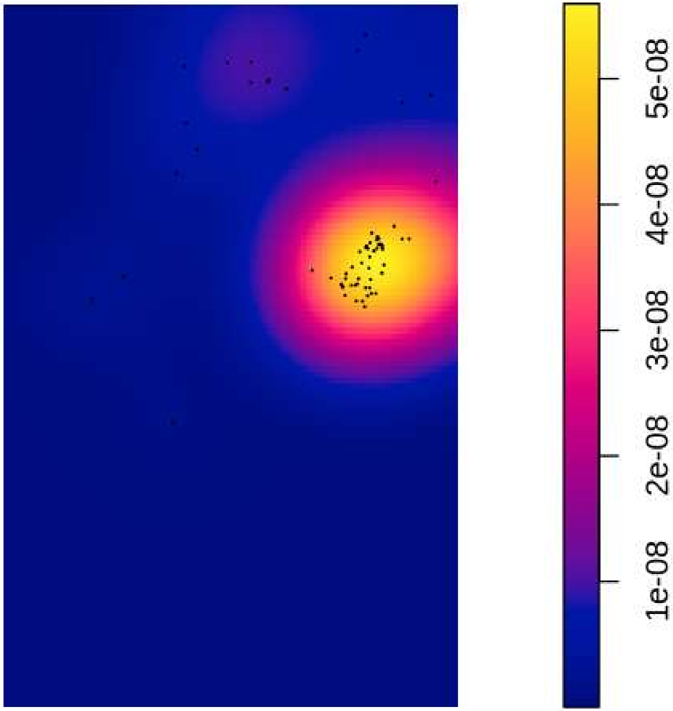

#### Density of PPR on um08

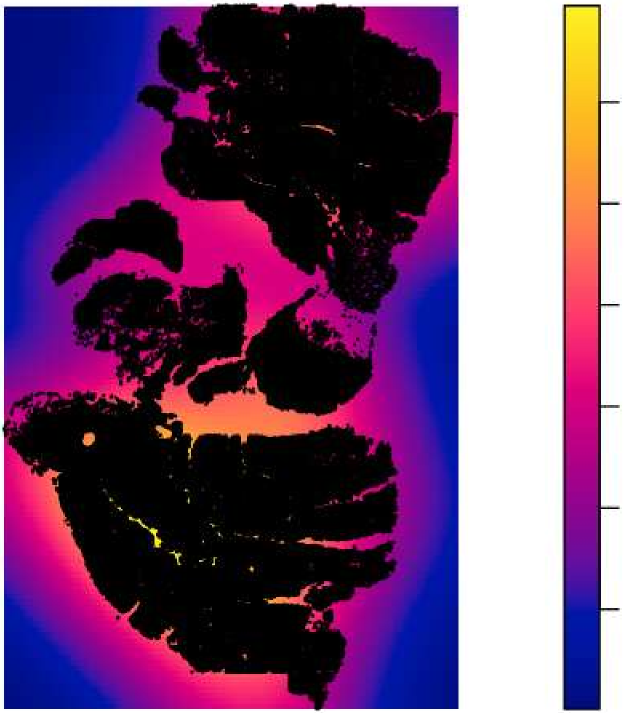

#### Density of Tcells on um08

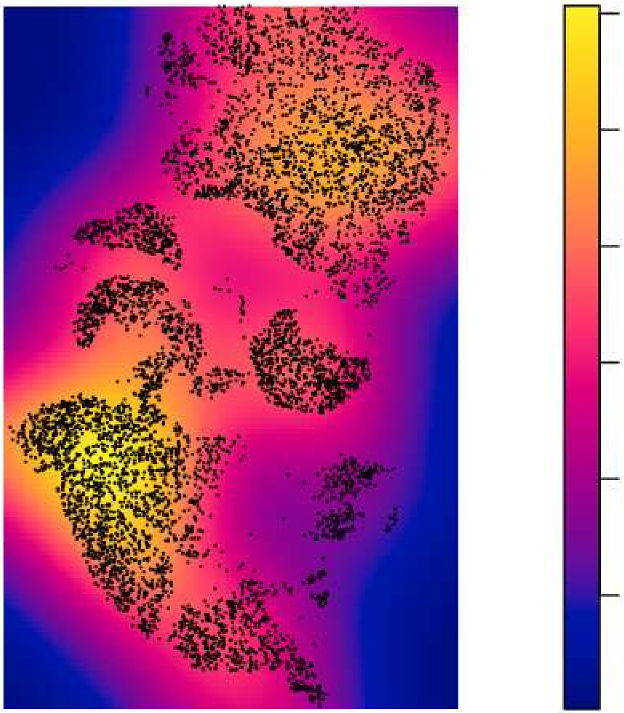

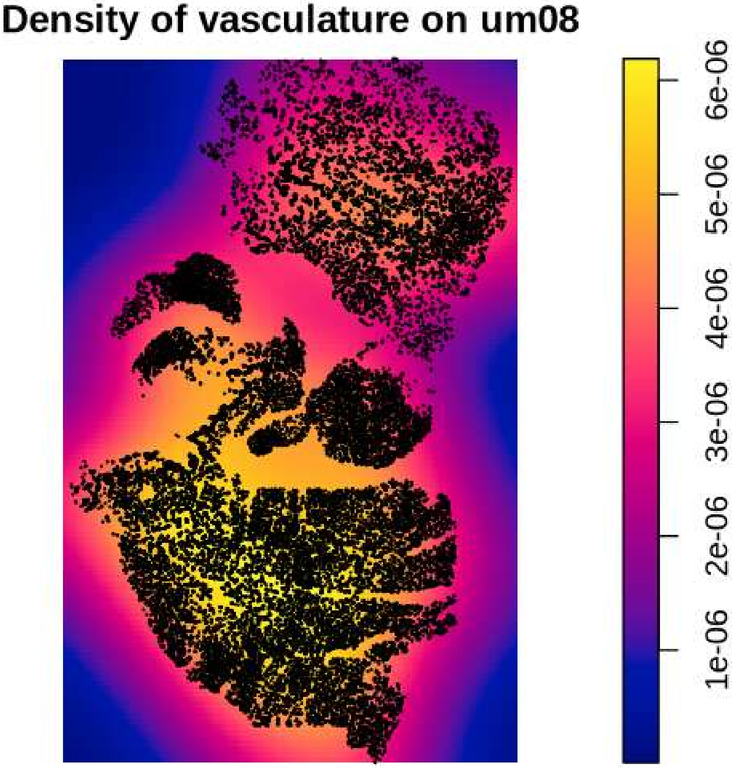

Density of vasculature on um08
6e-06
5e-06
4e-06
3e-06
2e-06
1e-06

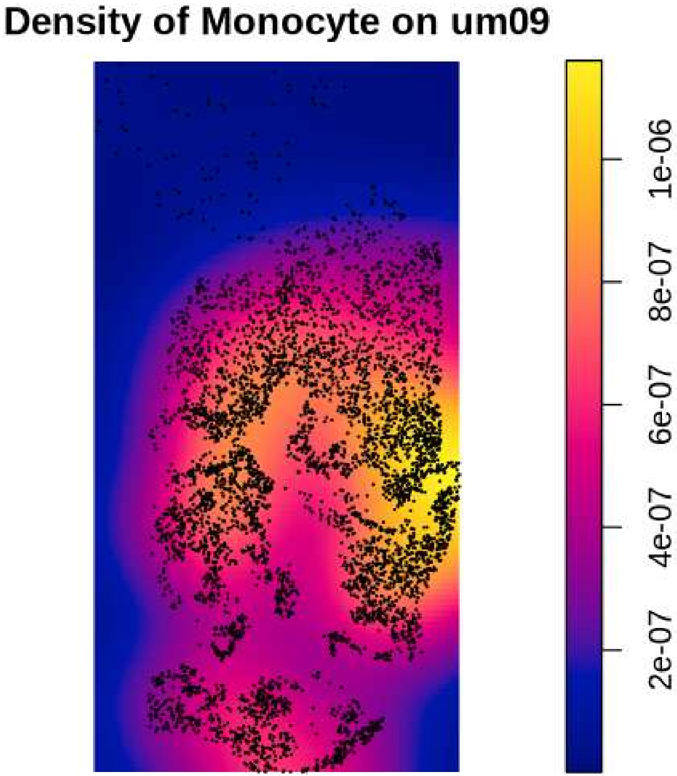

Density of Monocyte on um09
1e-06
8e-07
6e-07
4e-07
2e-07

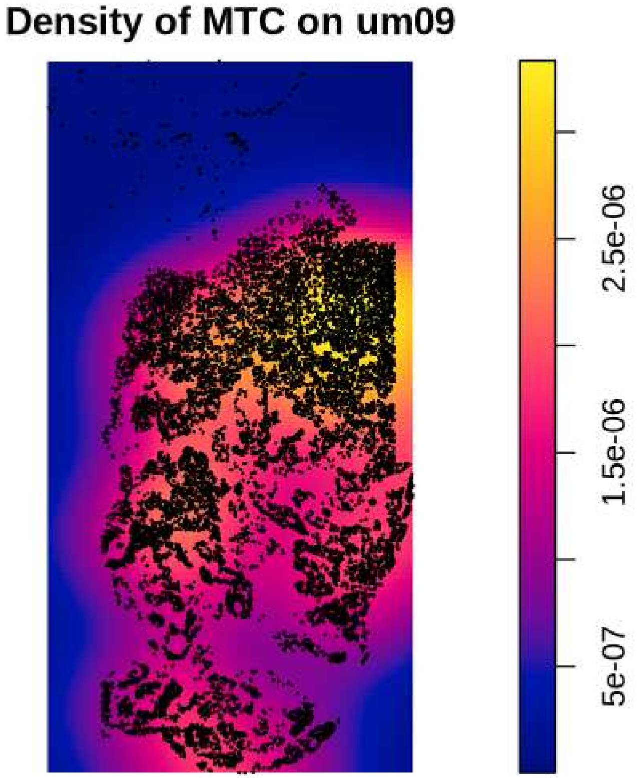

Density of MTC on um09
2.5e-06
1.5e-06
5e-07

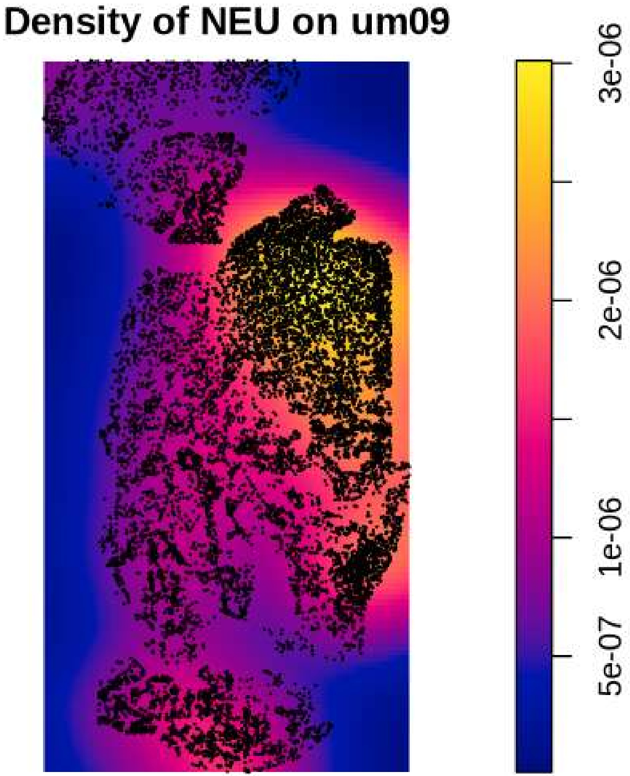
Density of NEU on um09
3e-06
2e-06
1e-06
5e-07

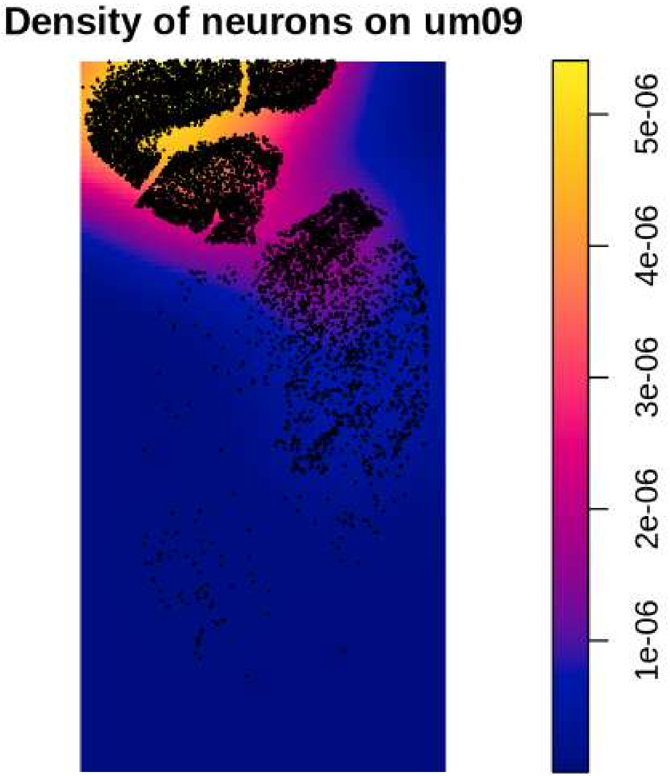

Density of neurons on um09
5e-06
4e-06
3e-06
2e-06
1e-06

Density of Neutrophil on um09

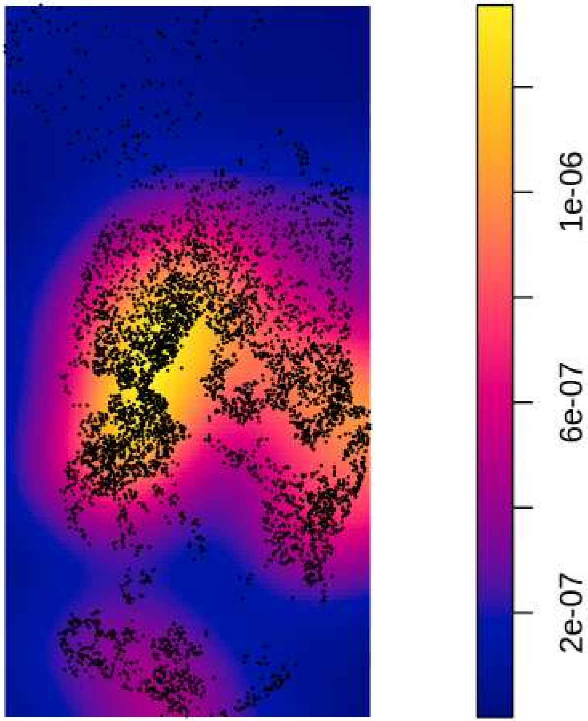

### Density of oligodendrocyte on um09

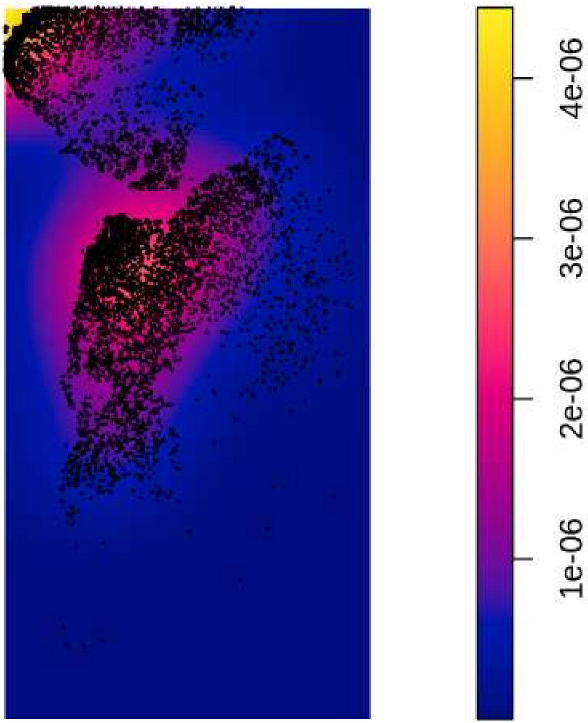

#### Density of OPC on um09

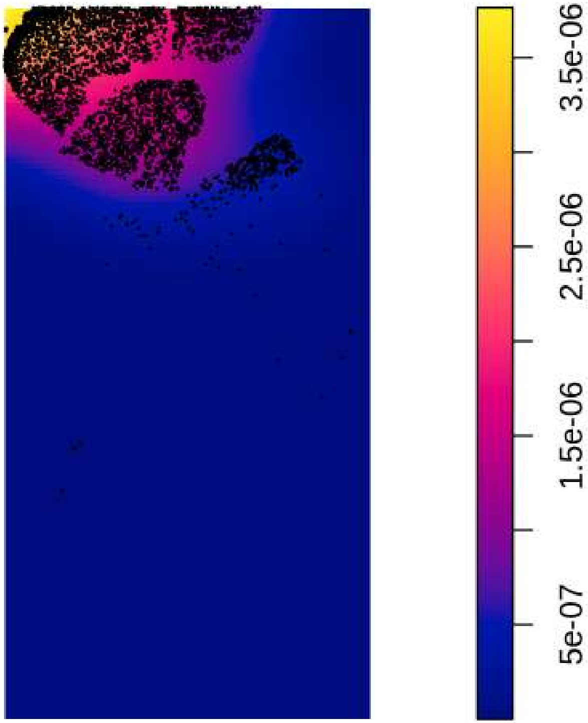

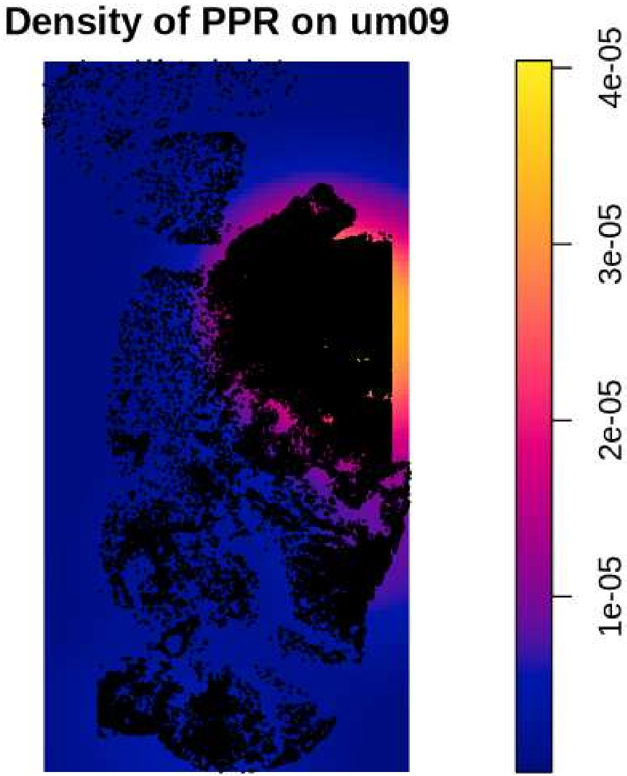

Density of PPR on um09
4e-05
3e-05
2e-05
1e-05

#### Density of Tcells on um09

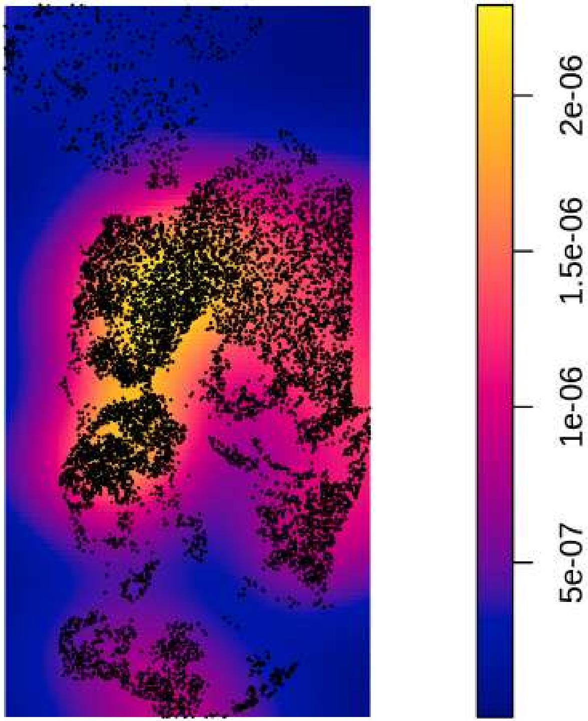

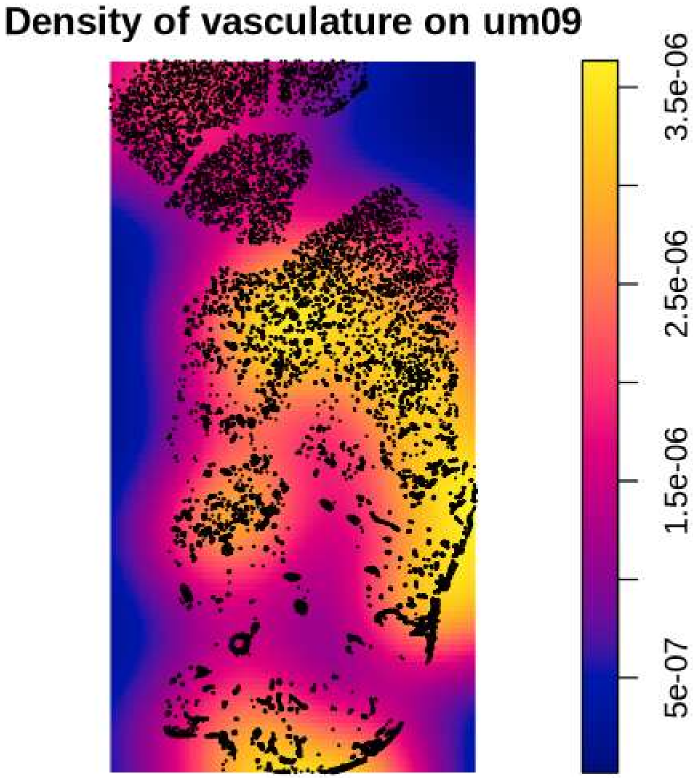

Density of vasculature on um09
3.5e-06
2.5e-06
1.5e-06
5e-07

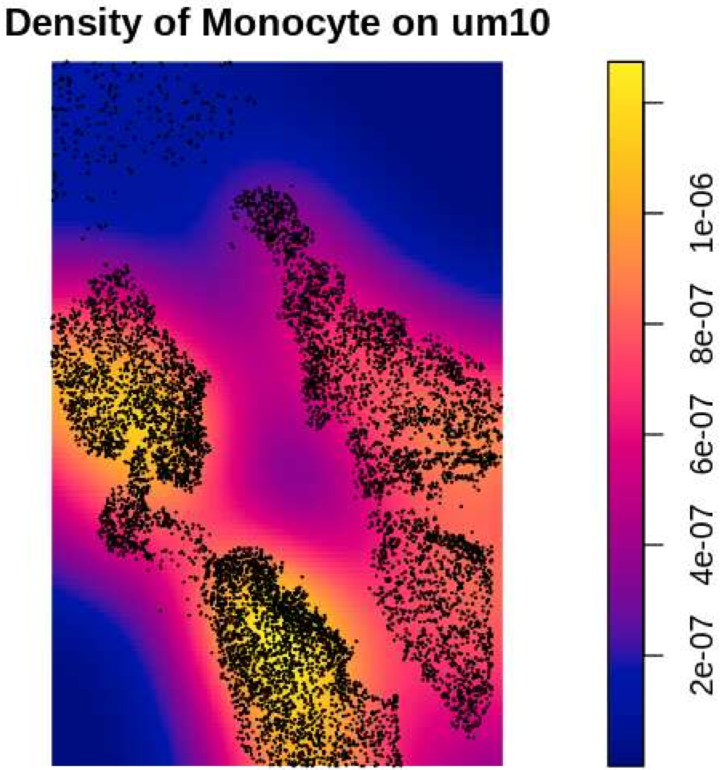
Density of Monocyte on um10
2e-07
4e-07
6e-07
8e-07
1e-06

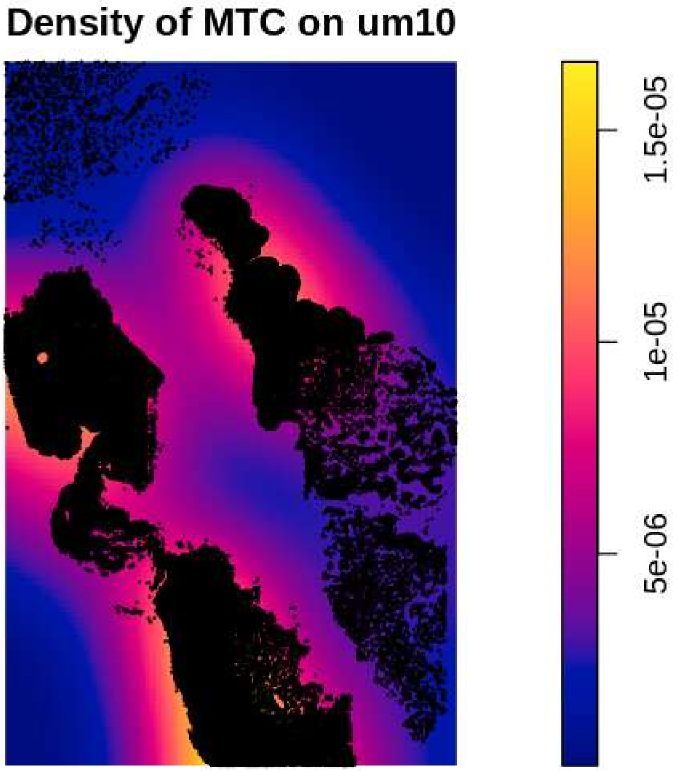

Density of MTC on um10
1.5e-05
1e-05
5e-06

Density of NEU on um10

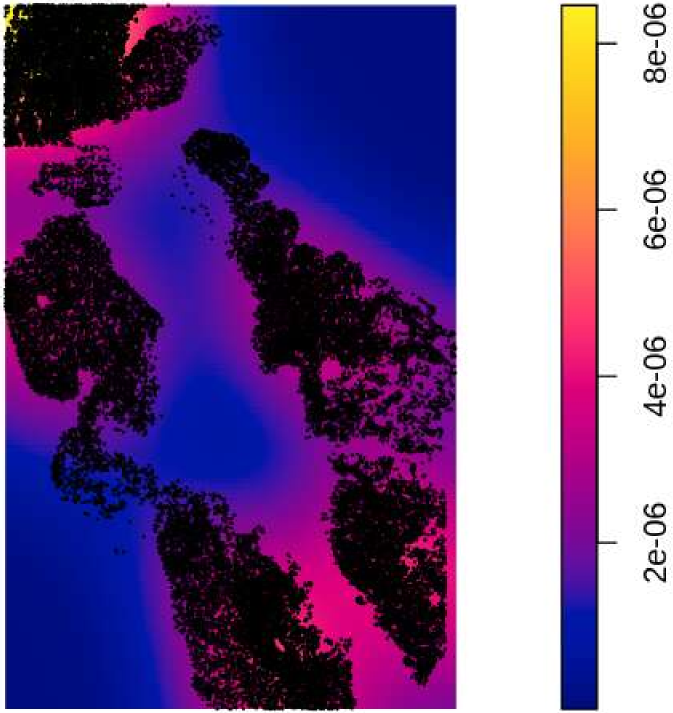

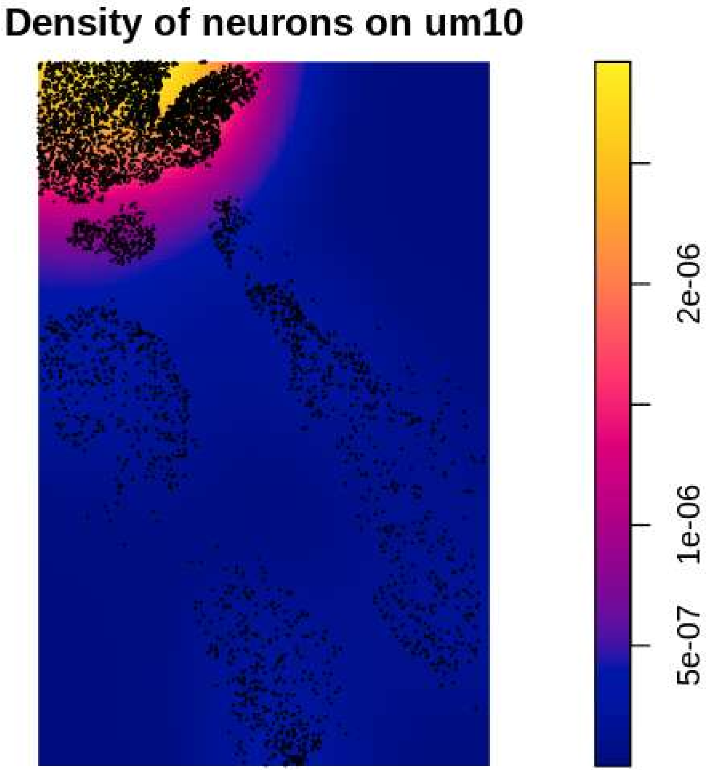
Density of neurons on um10
5e-07
1e-06
2e-06

### Density of Neutrophil on um10

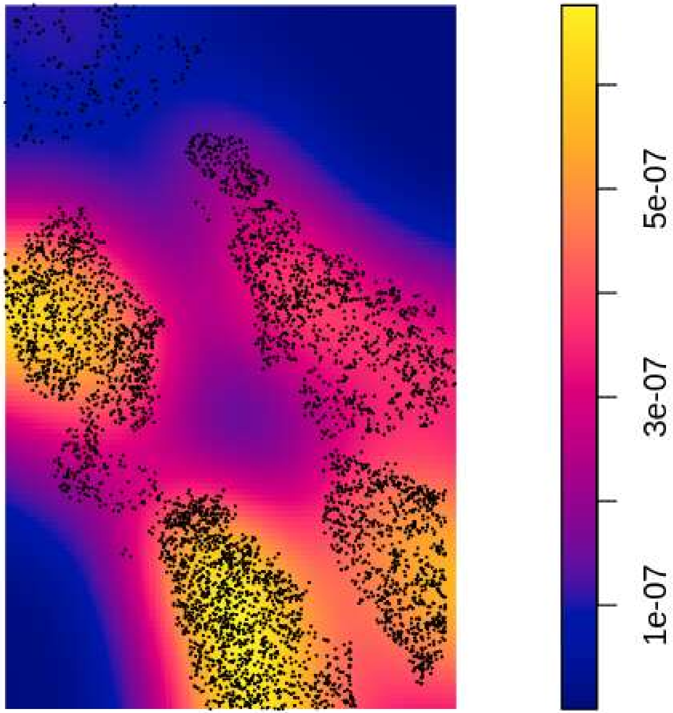

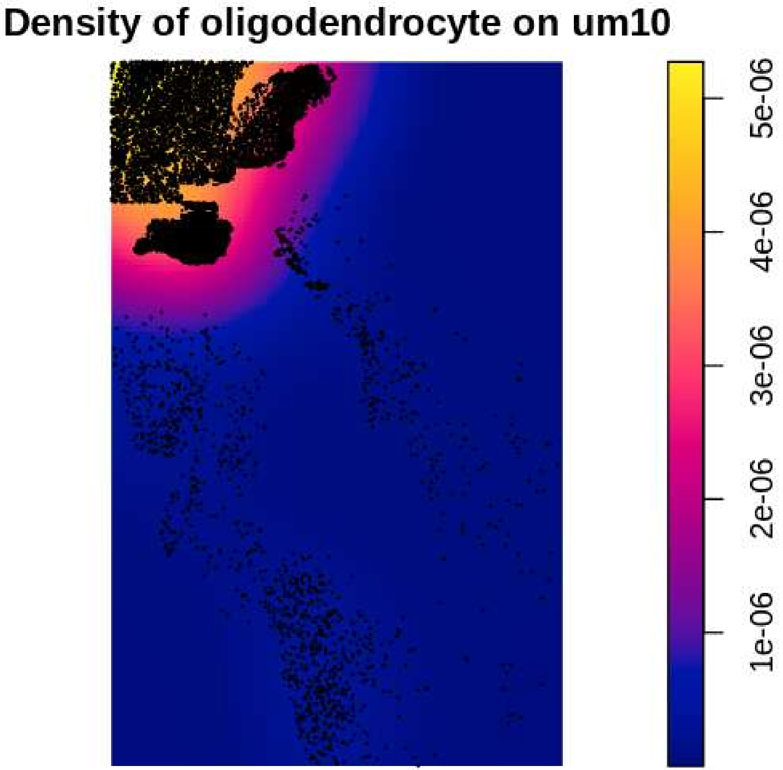

Density of oligodendrocyte on um10
5e-06
4e-06
3e-06
2e-06
1e-06

#### Density of OPC on um10

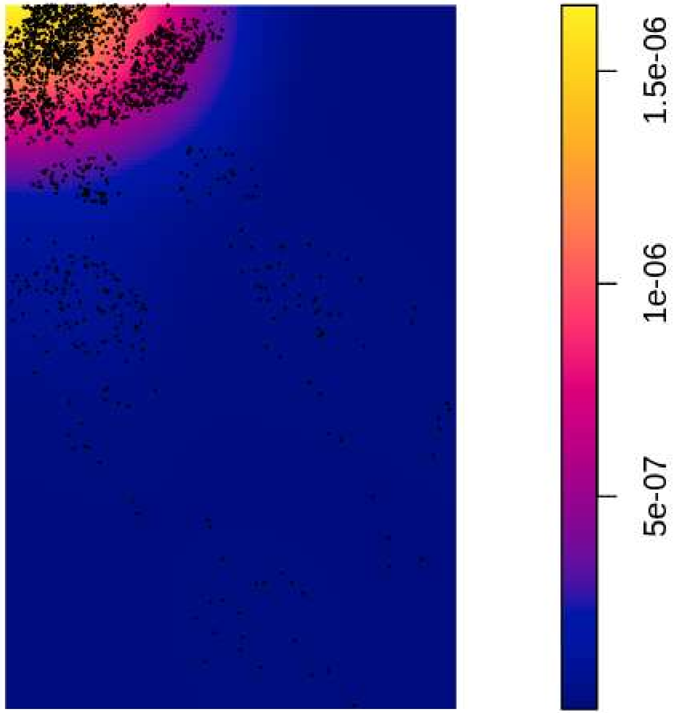

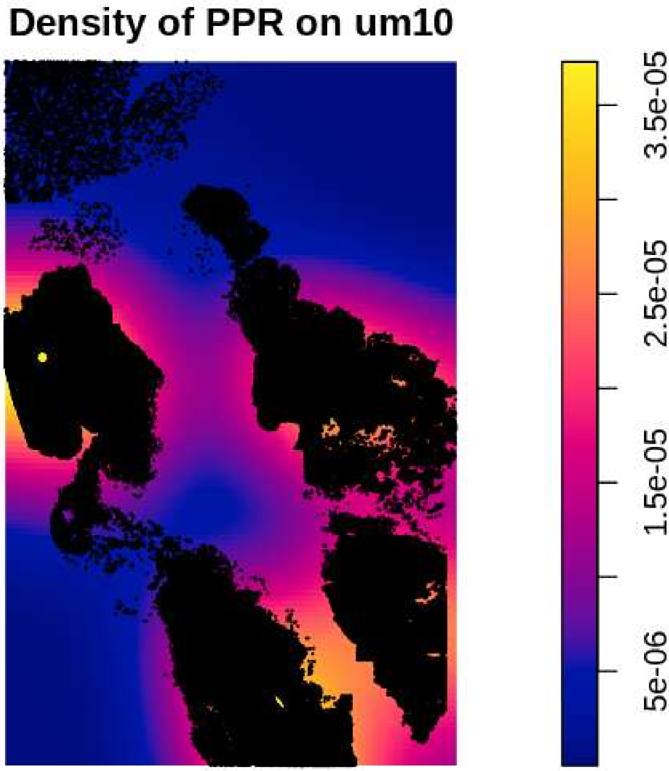
Density of PPR on um10
3.5e-05
2.5e-05
1.5e-05
5e-06

#### Density of Tcells on um10

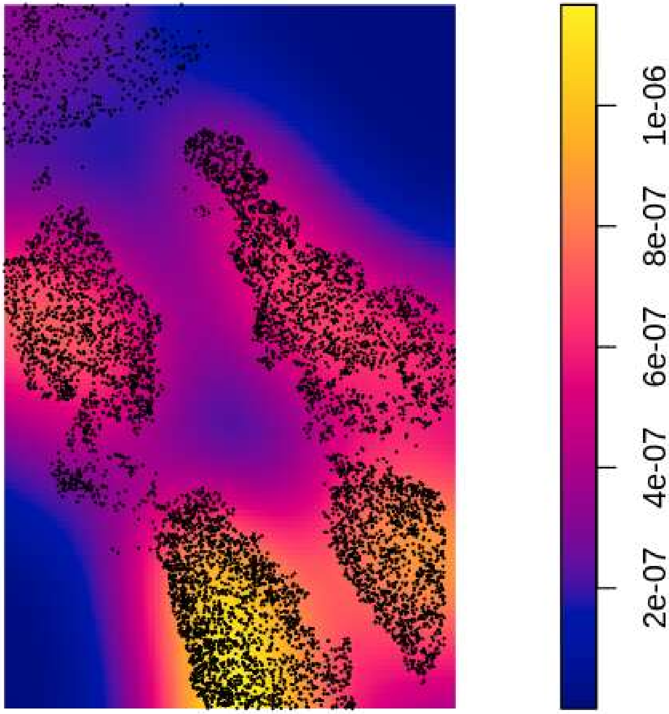

#### Density of vasculature on um10

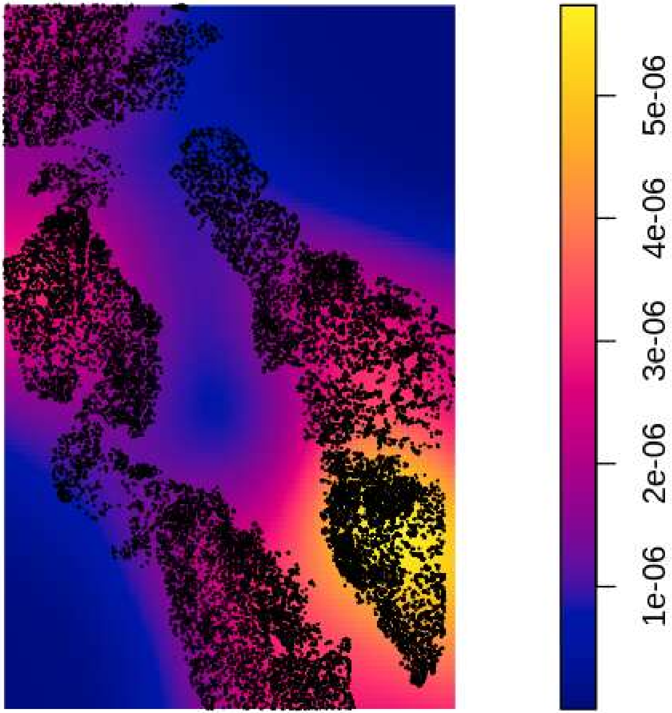

#### Density of Monocyte on um11

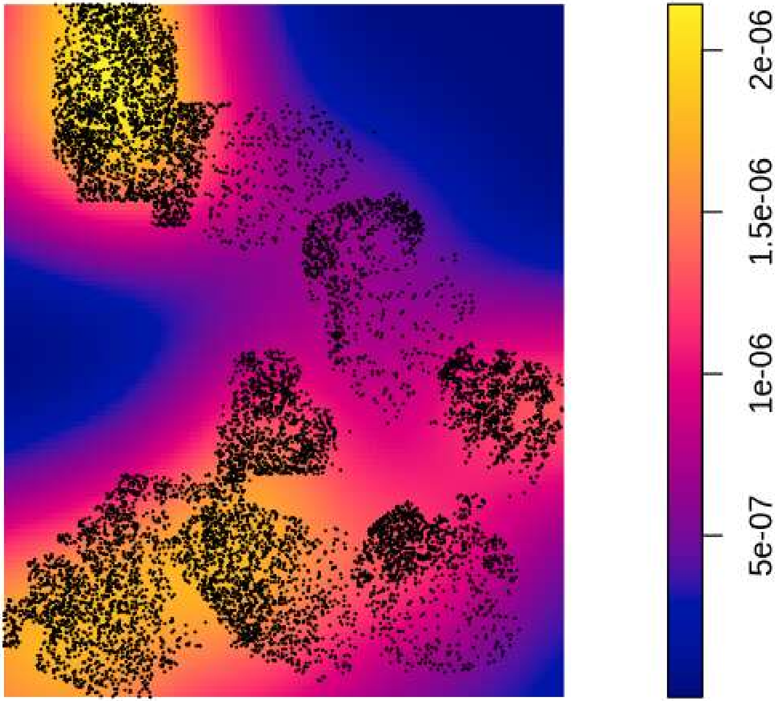

#### Density of MTC on um11

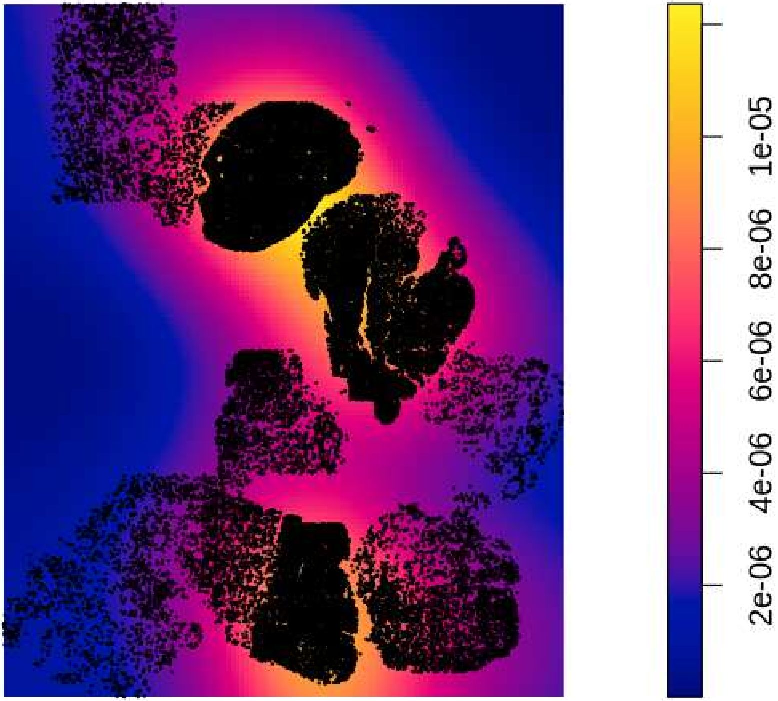

Density of NEU on um11

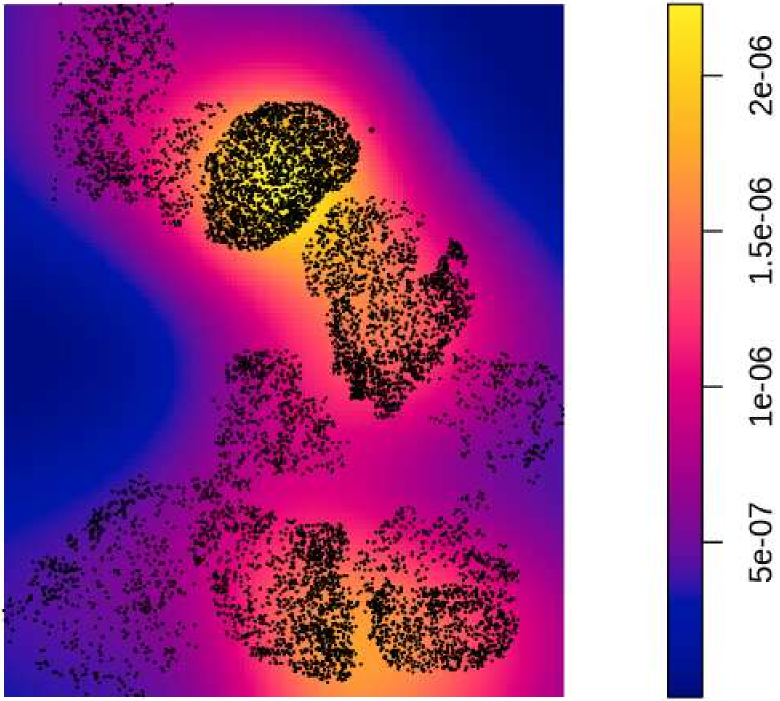

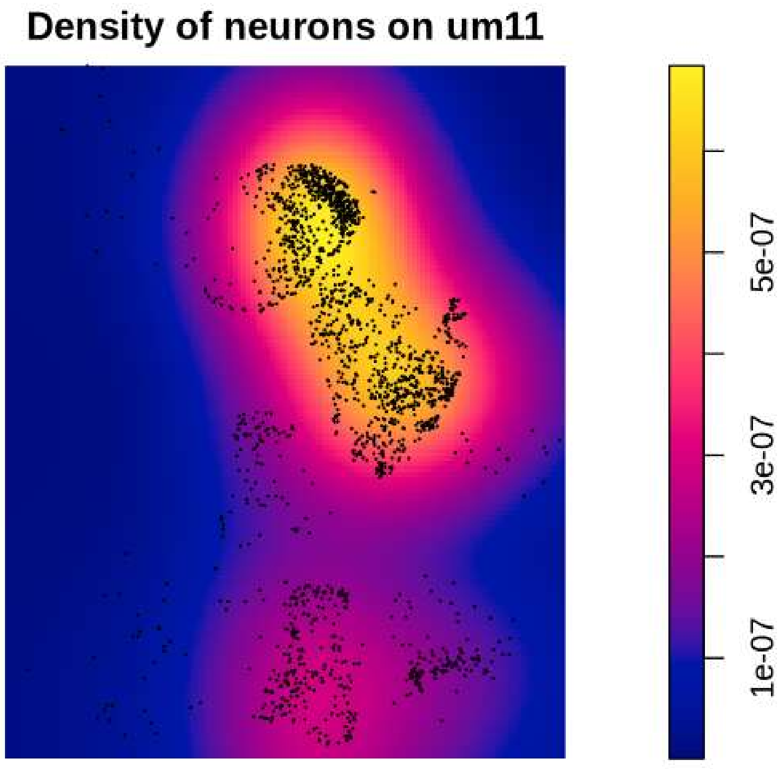
Density of neurons on um11
5e-07
3e-07
1e-07

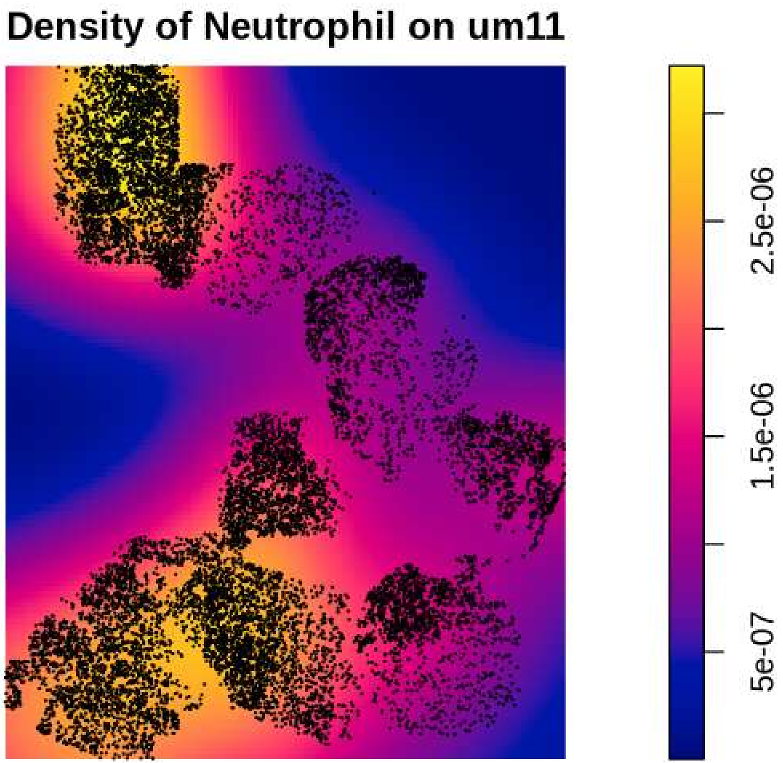
Density of Neutrophil on um11
2.5e-06
1.5e-06
5e-07

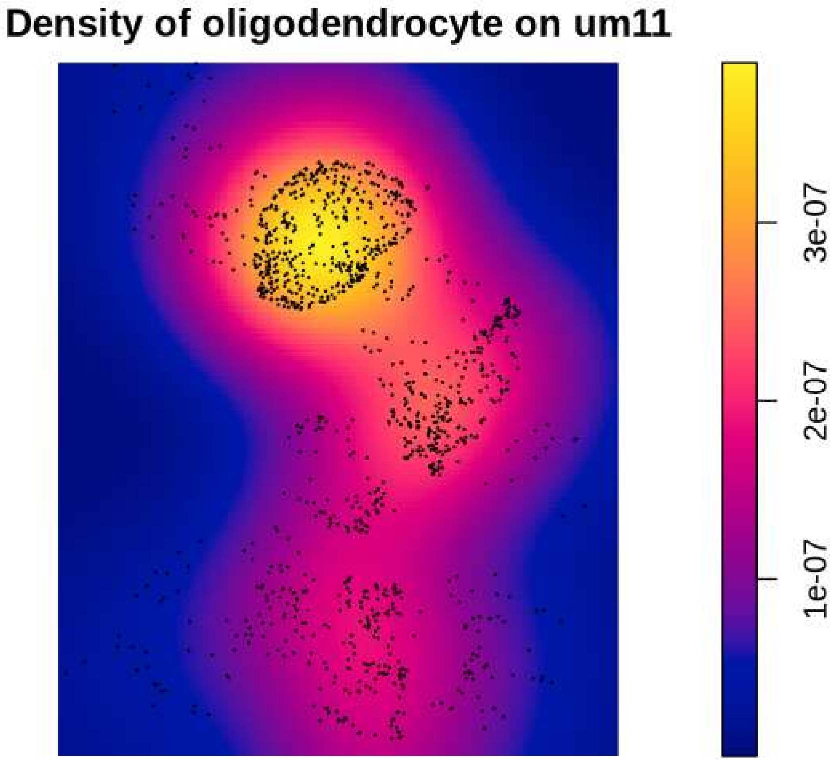

Density of oligodendrocyte on um11
3e-07
2e-07
1e-07

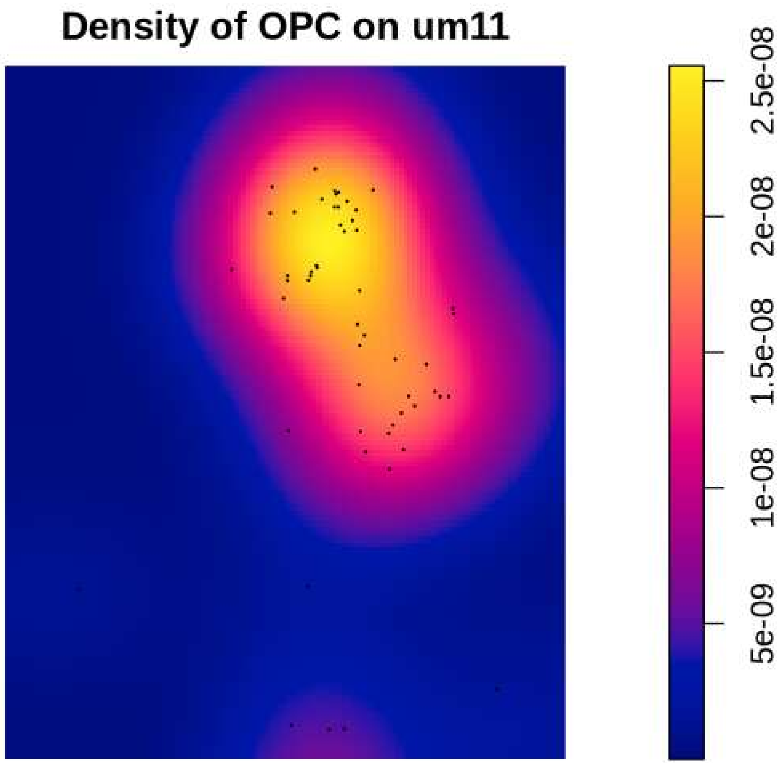
Density of OPC on um11
2.5e-08
2e-08
1.5e-08
1e-08
5e-09

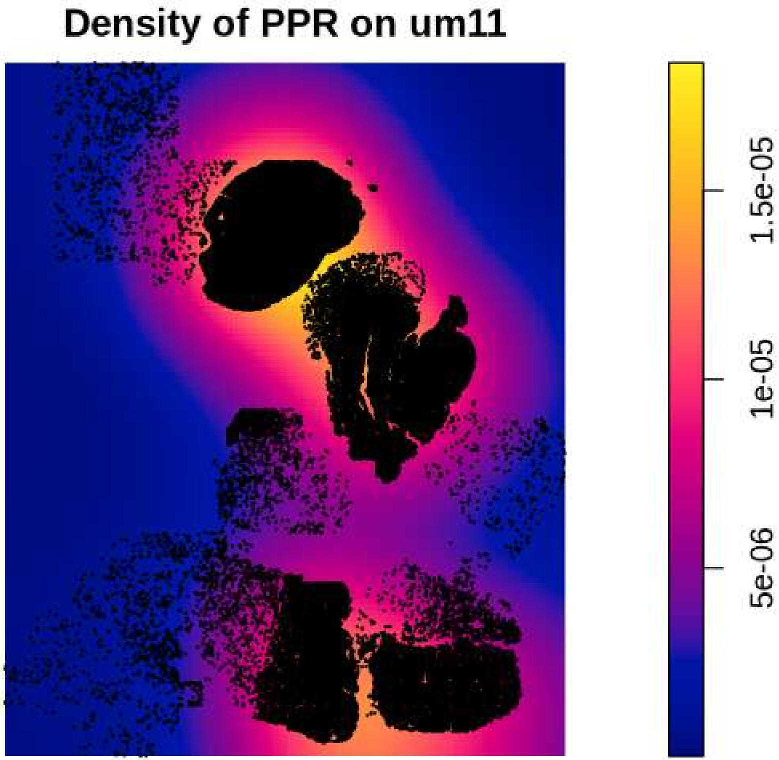
Density of PPR on um11
1.5e-05
1e-05
5e-06

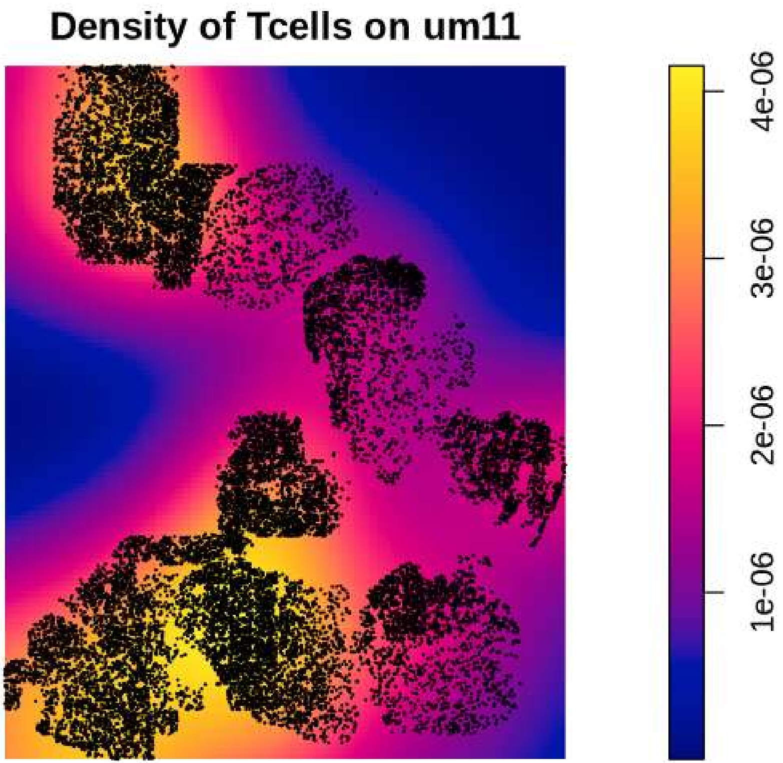
Density of Tcells on um11
4e-06
3e-06
2e-06
1e-06

**Density of vasculature on um11**

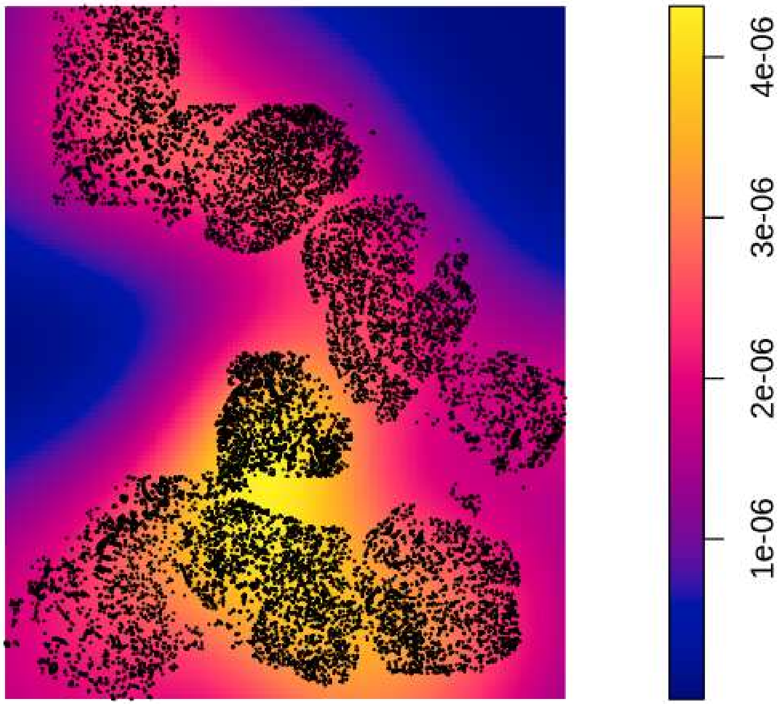

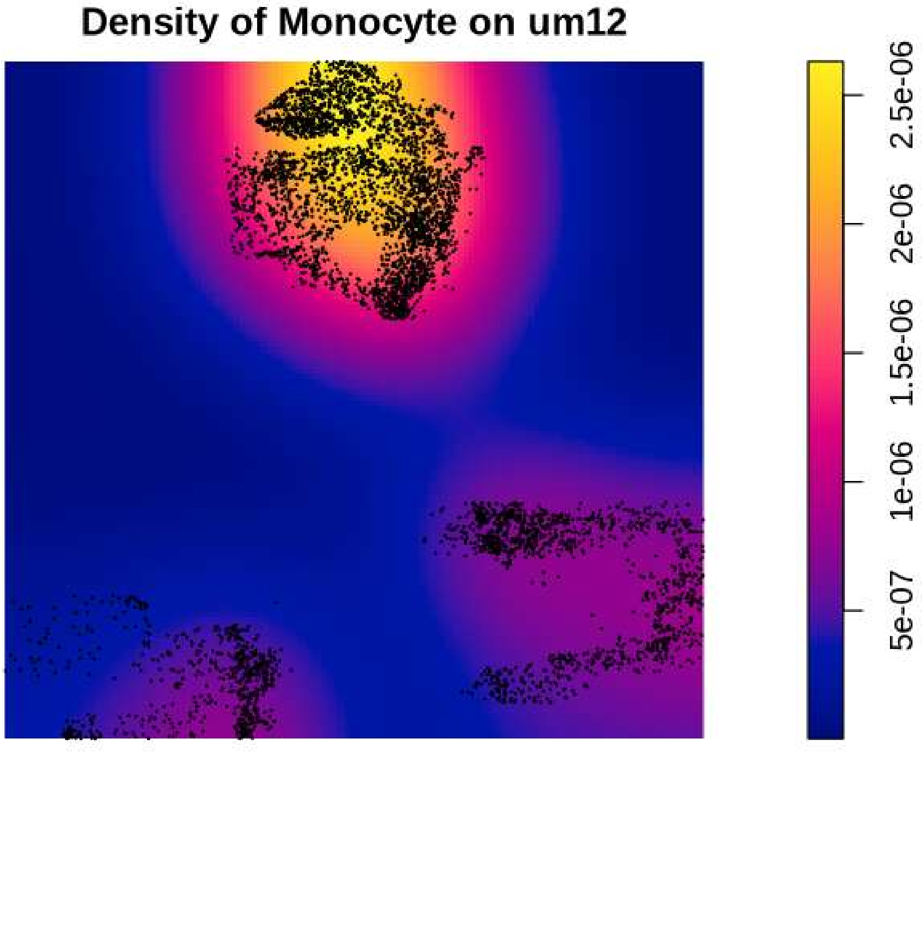
Density of Monocyte on um12
2.5e-06
2e-06
1.5e-06
1e-06
5e-07

#### Density of MTC on um12

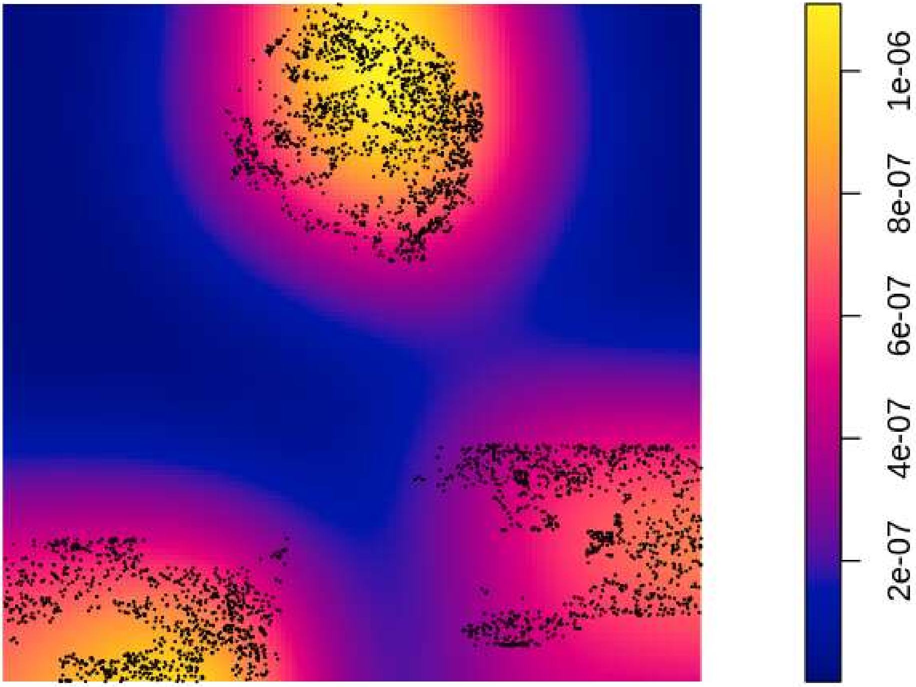

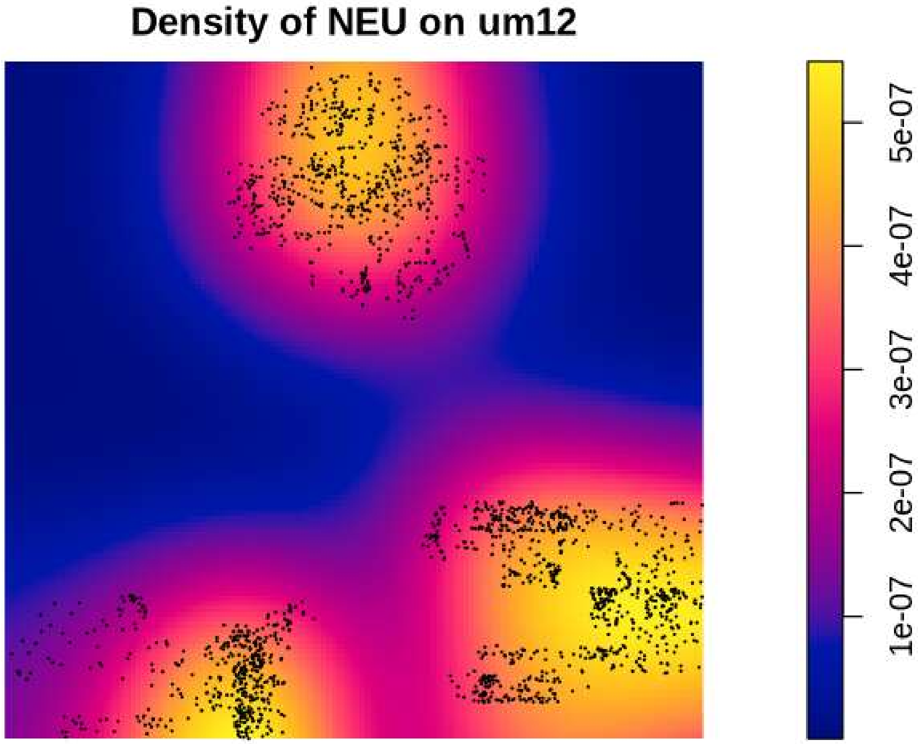

Density of NEU on um12
5e-07
4e-07
3e-07
2e-07
1e-07

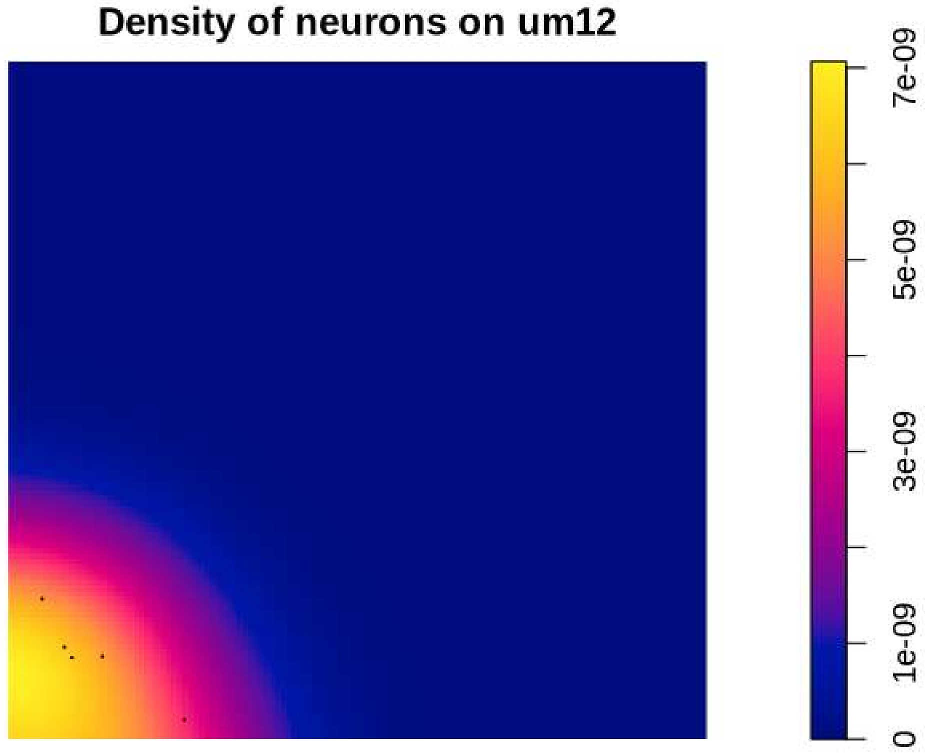
Density of neurons on um12
7e-09
5e-09
3e-09
1e-09
0

**Density of Neutrophil on um12**

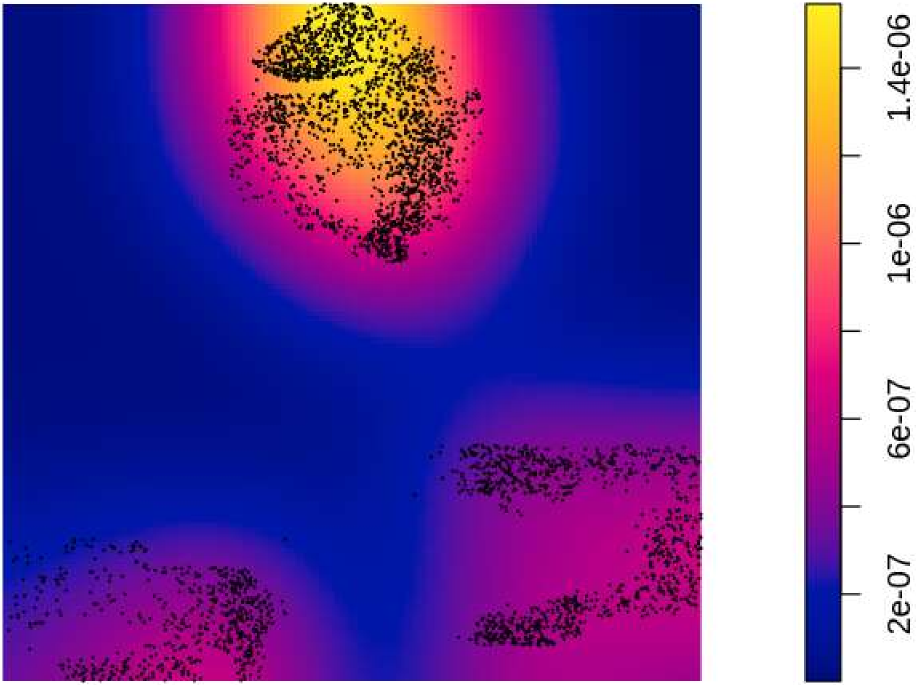

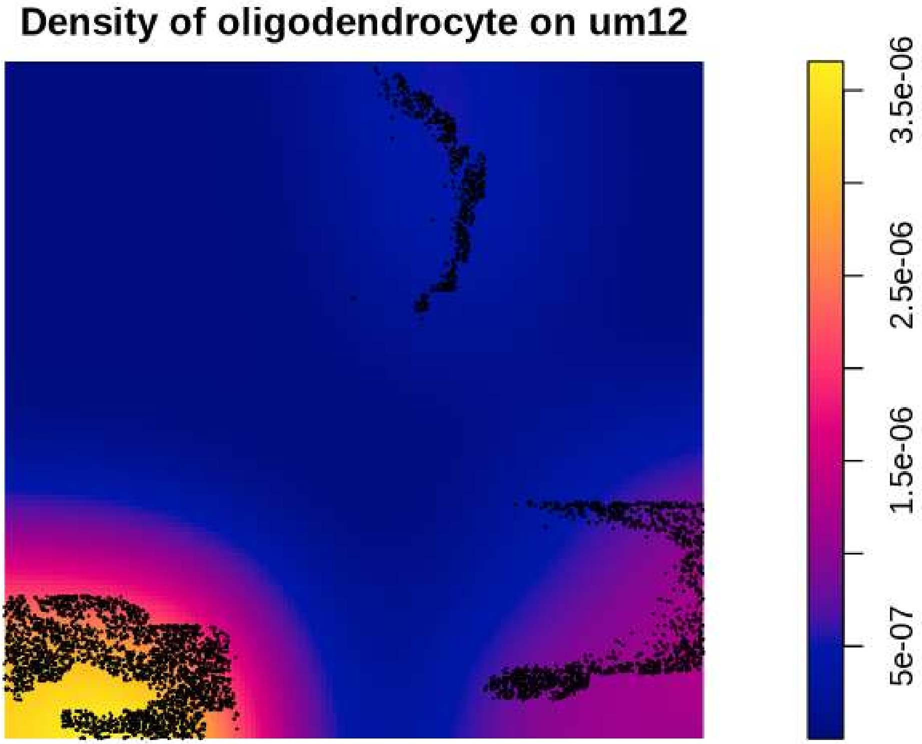
Density of oligodendrocyte on um12
3.5e-06
2.5e-06
1.5e-06
5e-07

#### Density of OPC on um12

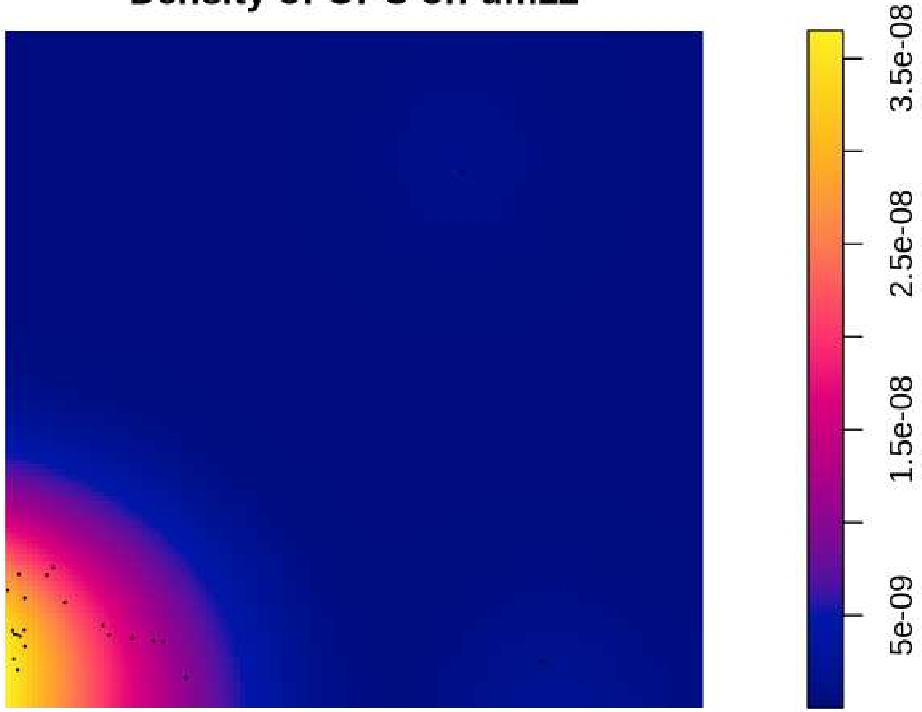

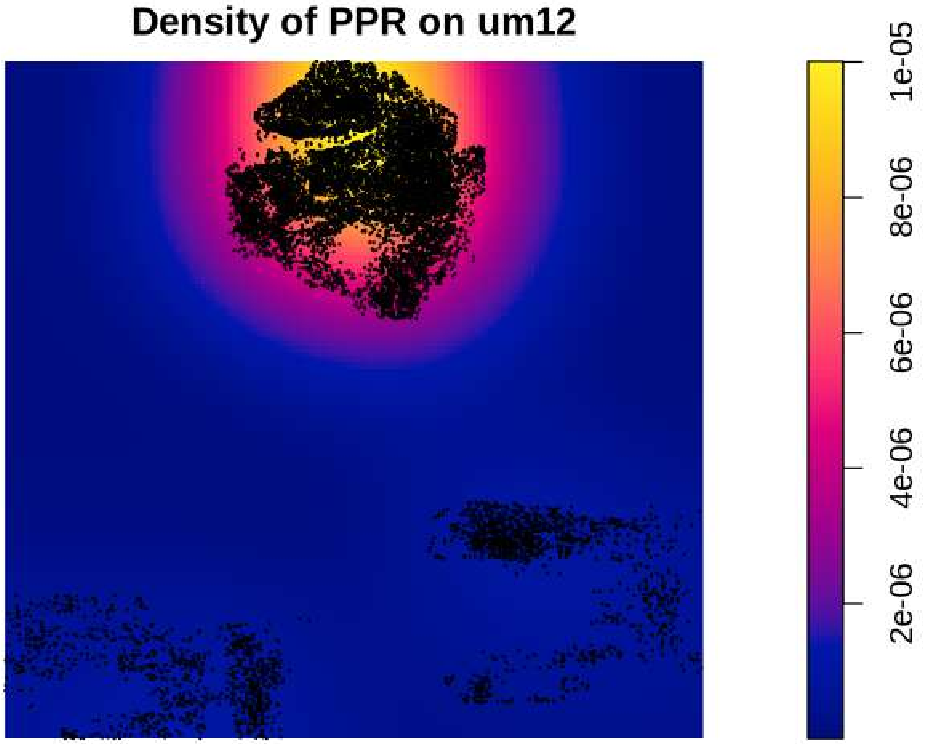

Density of PPR on um12
1e-05
8e-06
6e-06
4e-06
2e-06

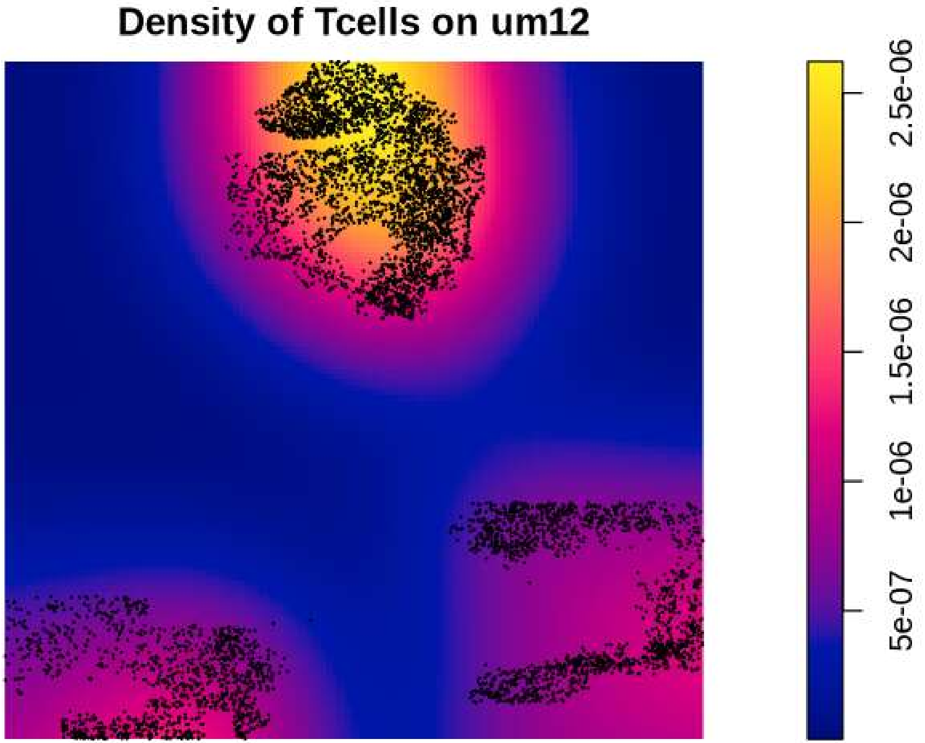

Density of Tcells on um12
2.5e-06
2e-06
1.5e-06
1e-06
5e-07

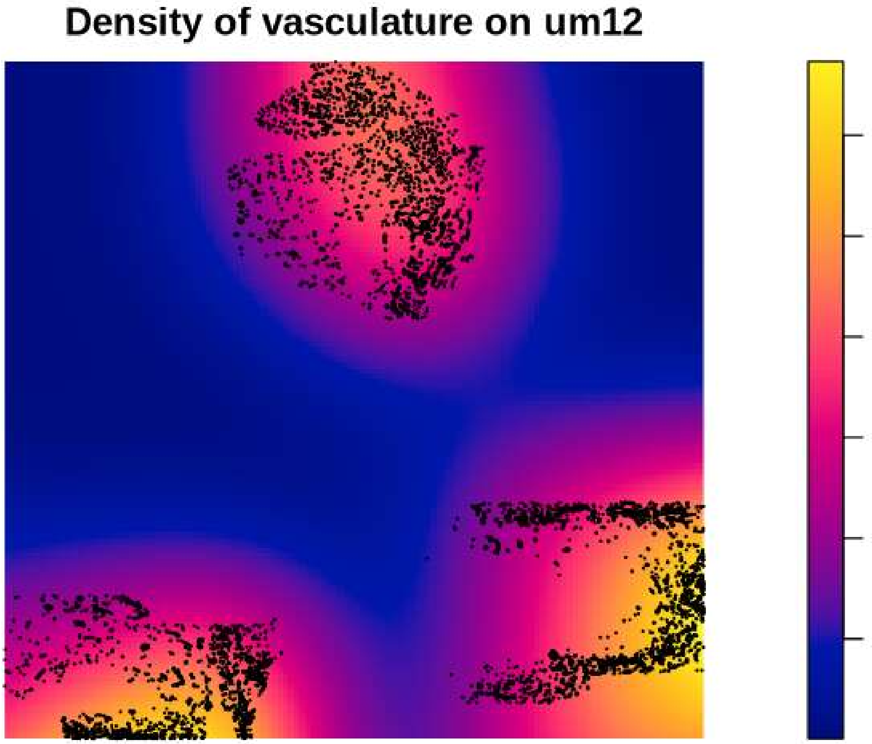

Density of vasculature on um12
5e-07
1.5e-06
2.5e-06

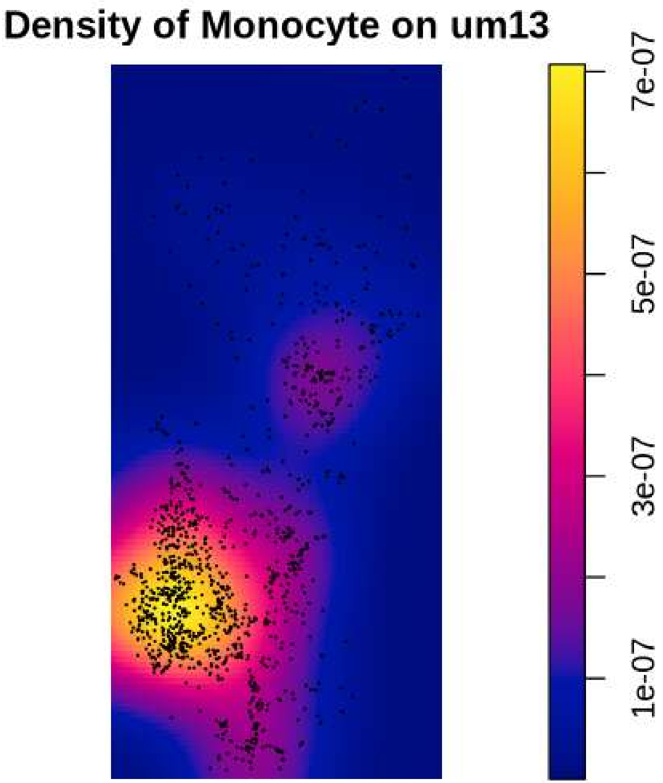
Density of Monocyte on um13
7e-07
5e-07
3e-07
1e-07

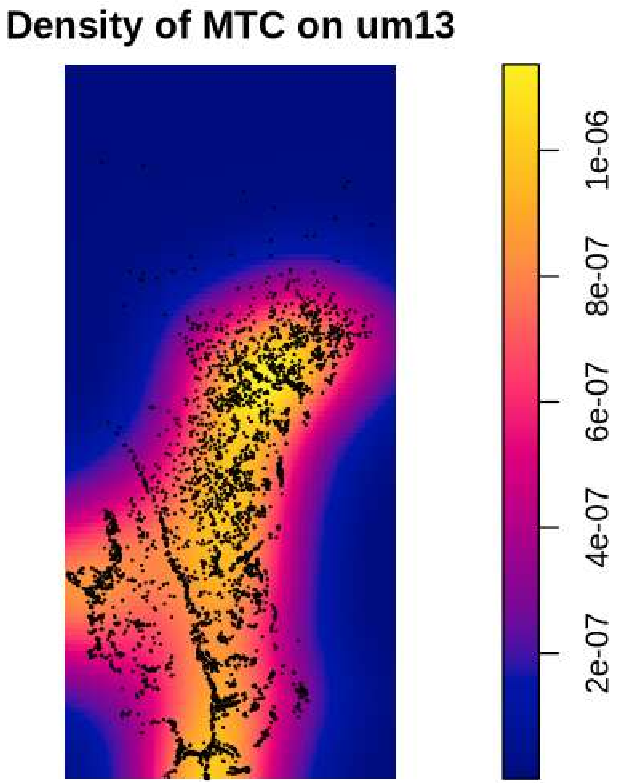
Density of MTC on um13
1e-06
8e-07
6e-07
4e-07
2e-07

Density of NEU on um13

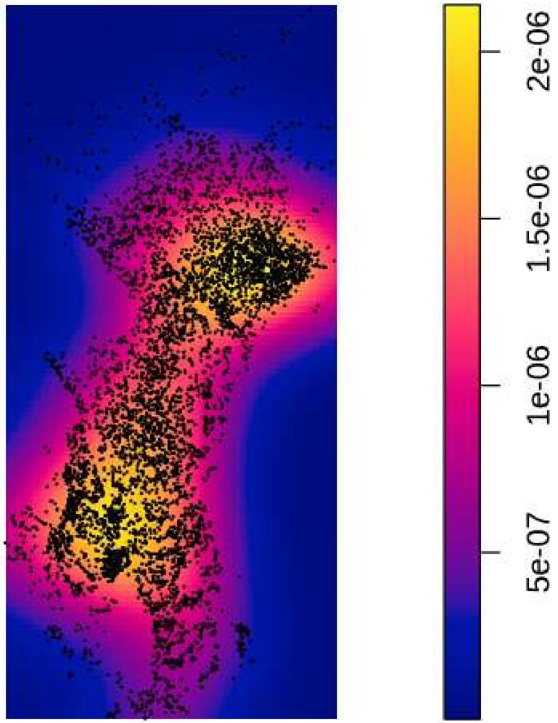

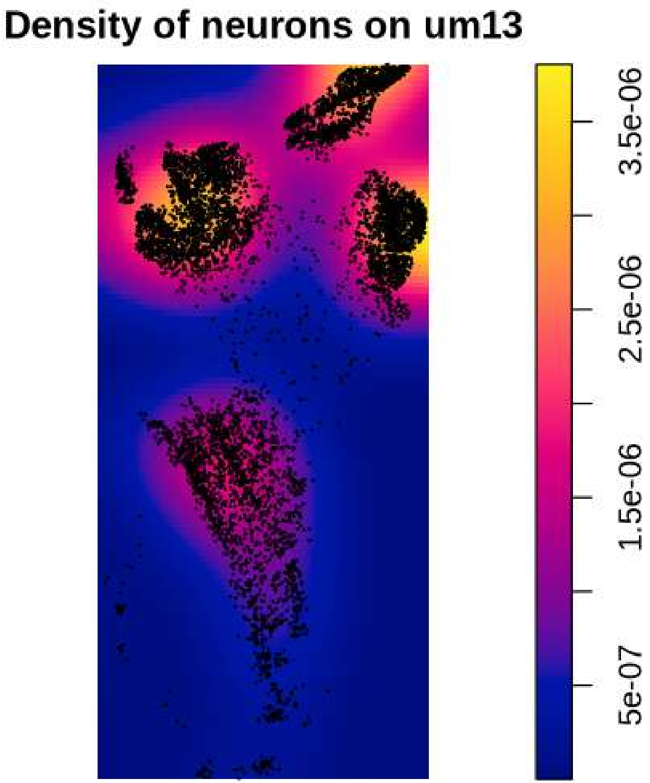
Density of neurons on um13
3.5e-06
2.5e-06
1.5e-06
5e-07

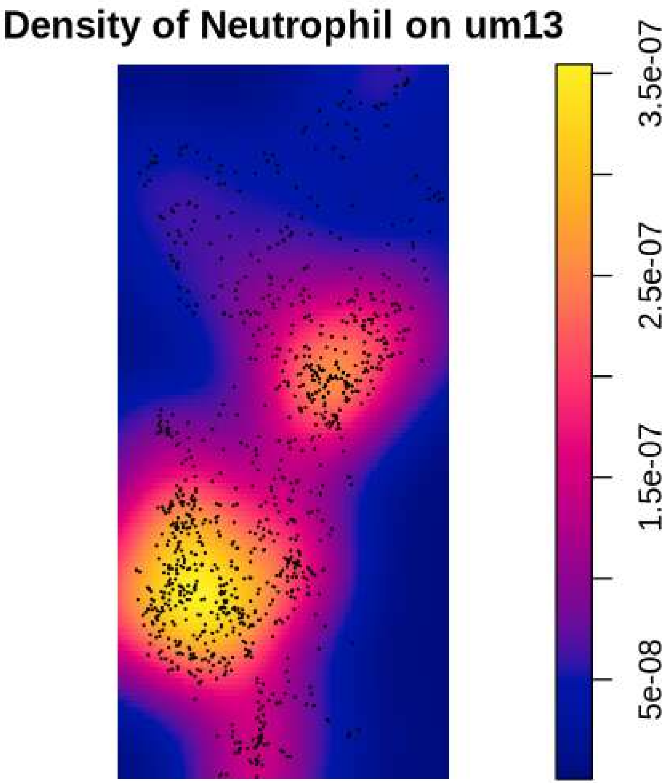

Density of Neutrophil on um13
3.5e-07
2.5e-07
1.5e-07
5e-08

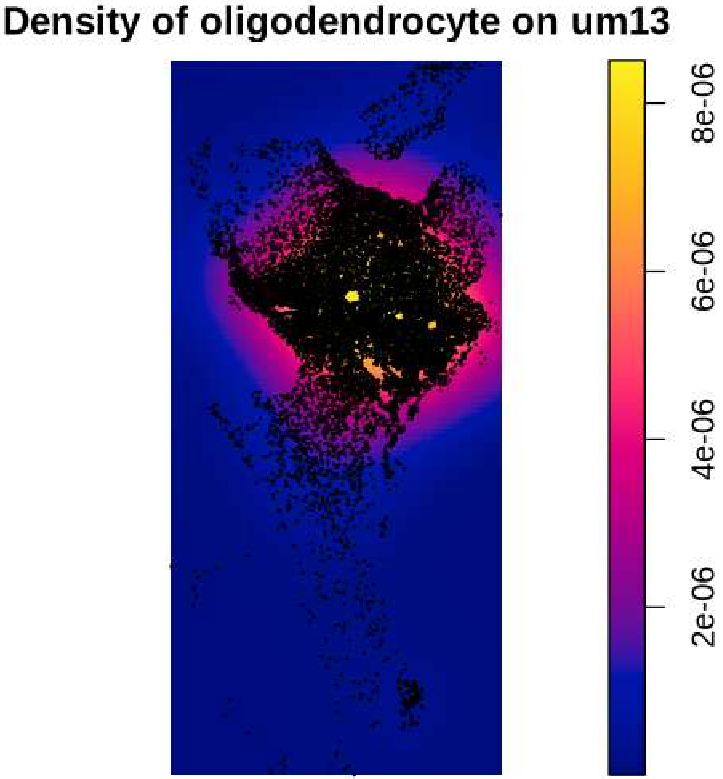
Density of oligodendrocyte on um13
8e-06
6e-06
4e-06
2e-06

#### Density of OPC on um13

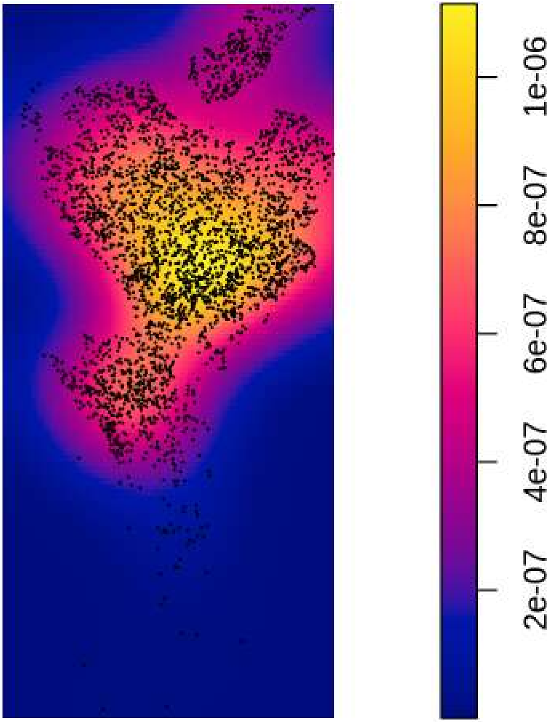

### Density of PPR on um13

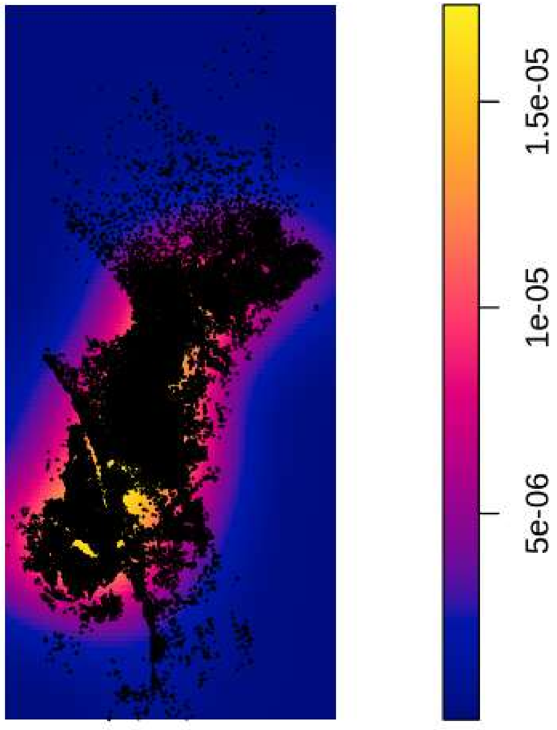

Density of Tcells on um13

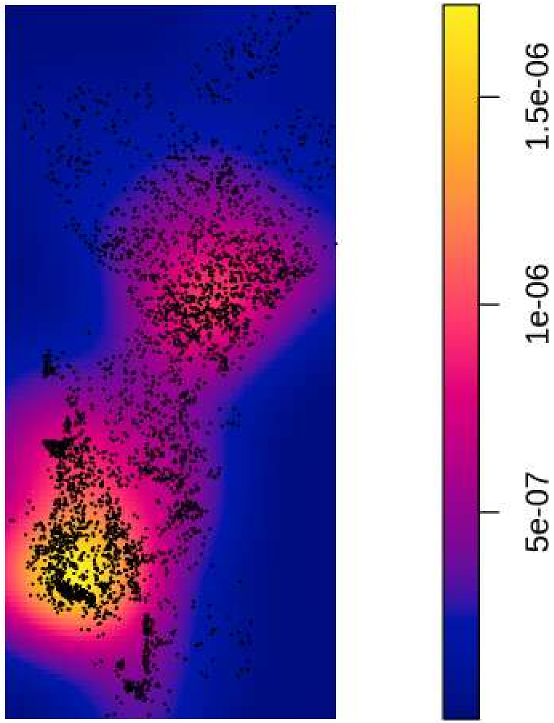

#### Density of vasculature on um13

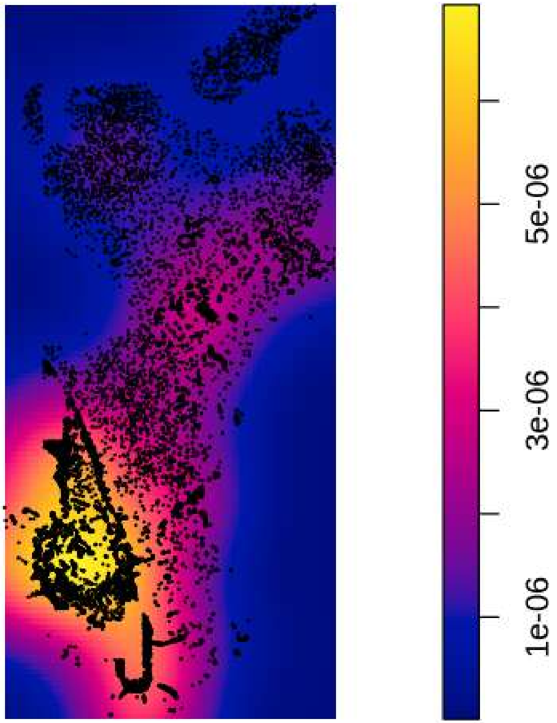

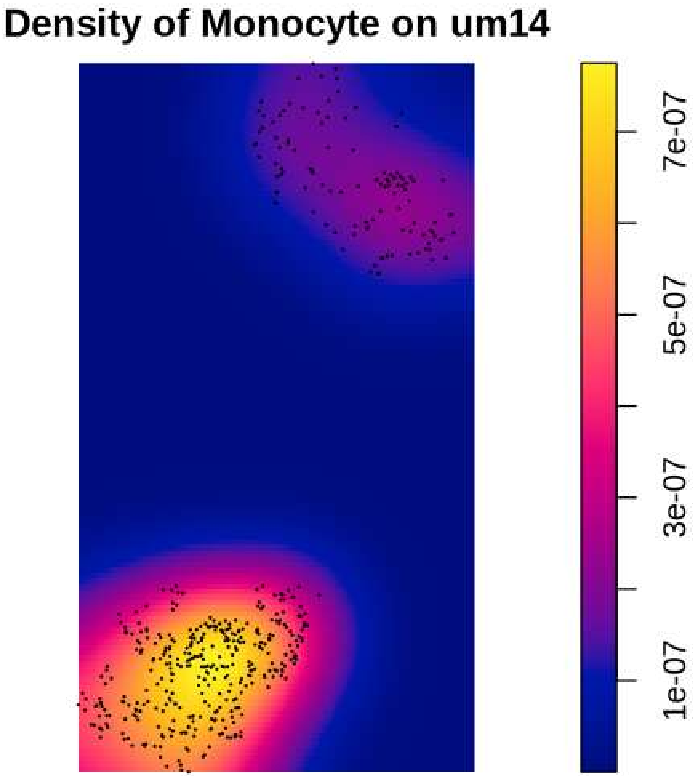
Density of Monocyte on um14
7e-07
5e-07
3e-07
1e-07

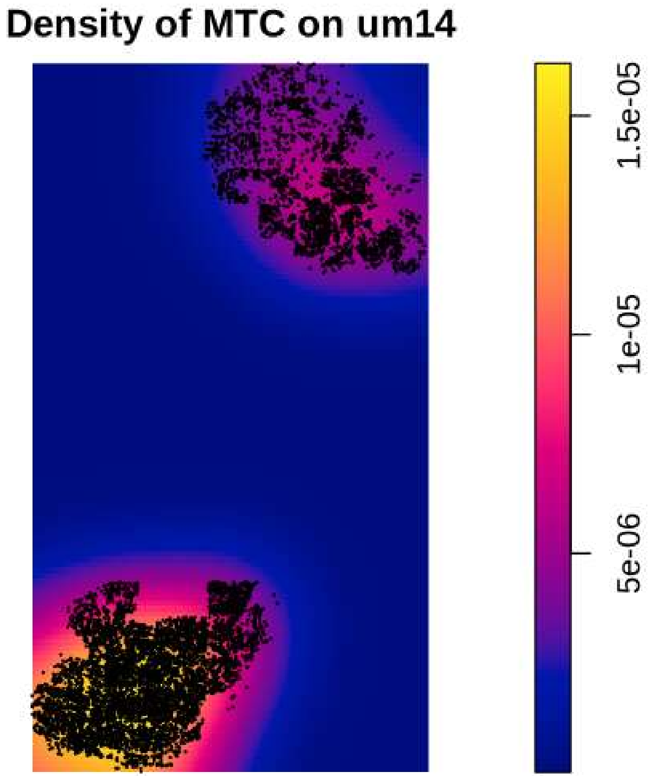
Density of MTC on um14
1.5e-05
1e-05
5e-06

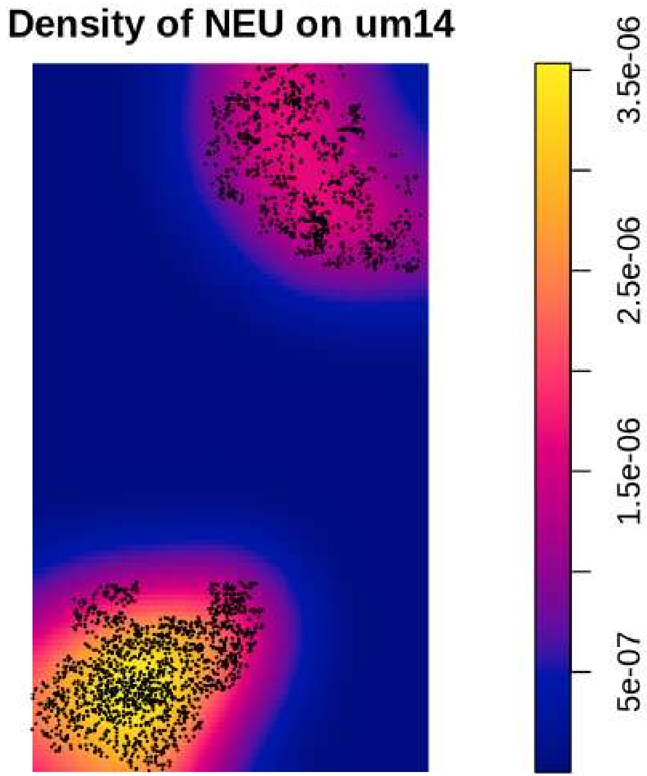

Density of NEU on um14
3.5e-06
2.5e-06
1.5e-06
5e-07

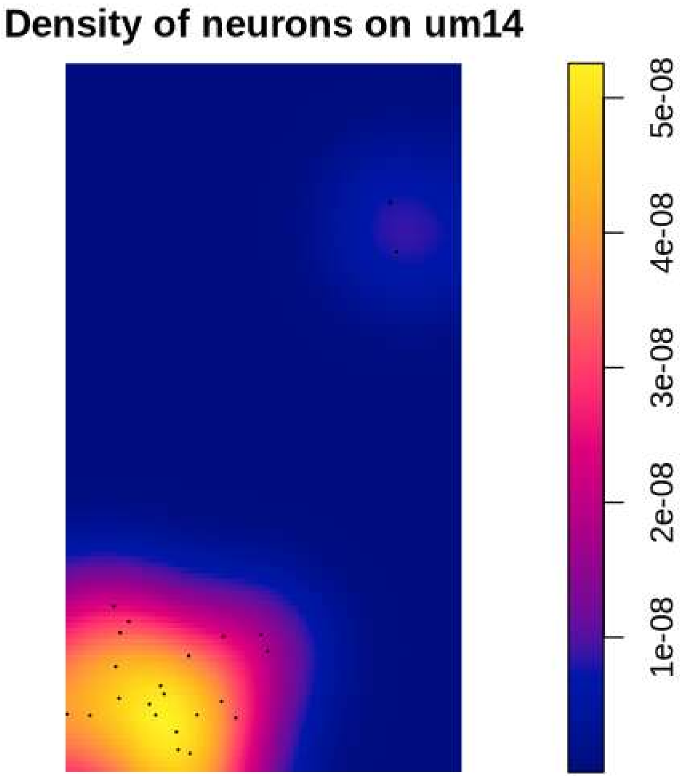
Density of neurons on um14
5e-08
4e-08
3e-08
2e-08
1e-08

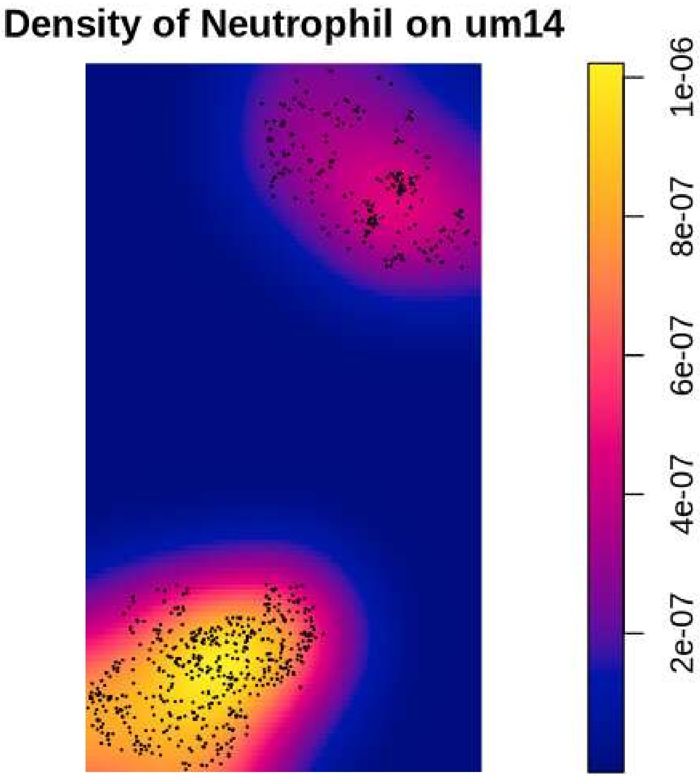

Density of Neutrophil on um14
1e-06
8e-07
6e-07
4e-07
2e-07

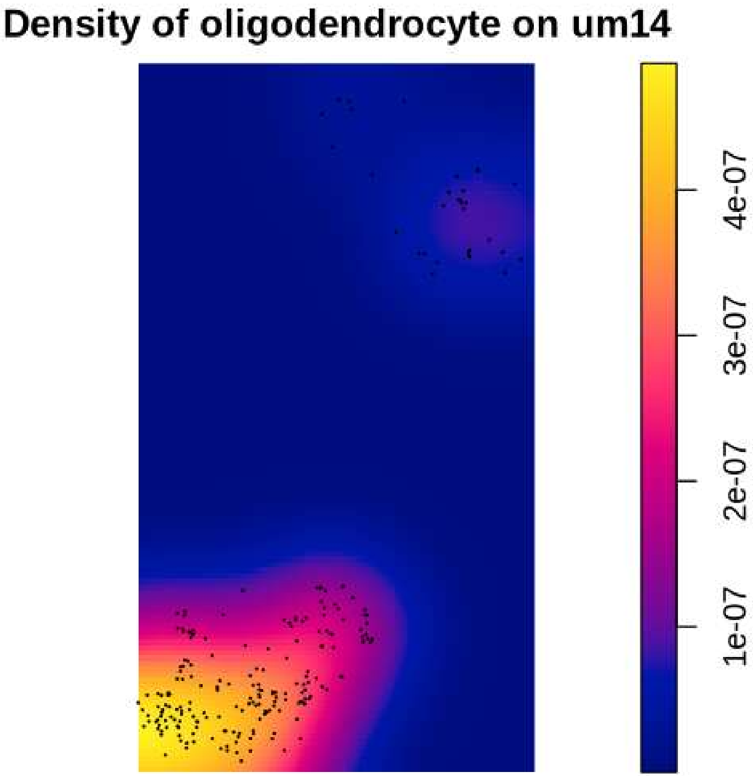
Density of oligodendrocyte on um14
4e-07
3e-07
2e-07
1e-07

#### Density of OPC on um14

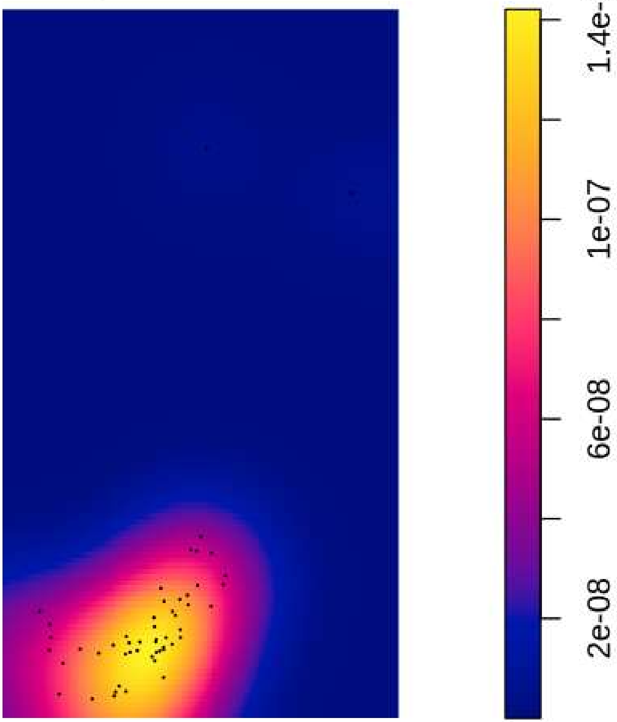

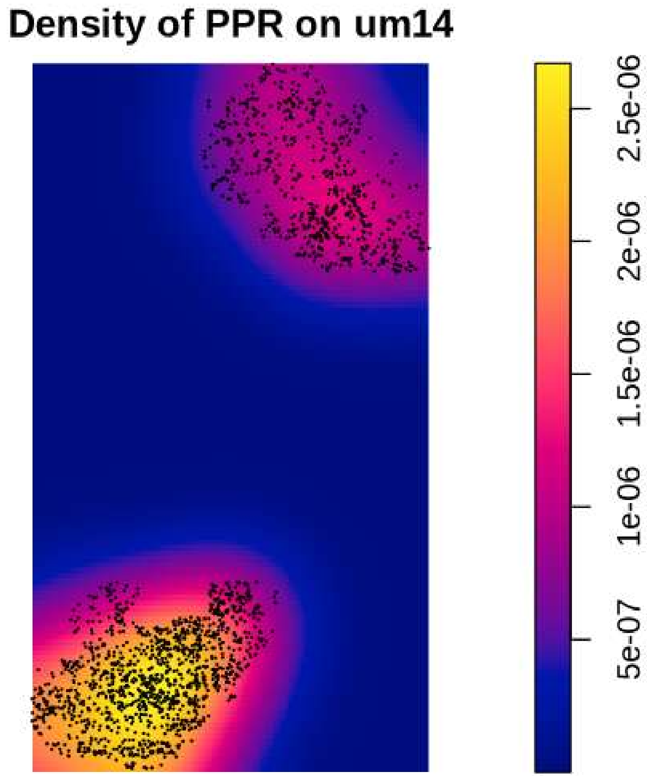
Density of PPR on um14
2.5e-06
2e-06
1.5e-06
1e-06
5e-07

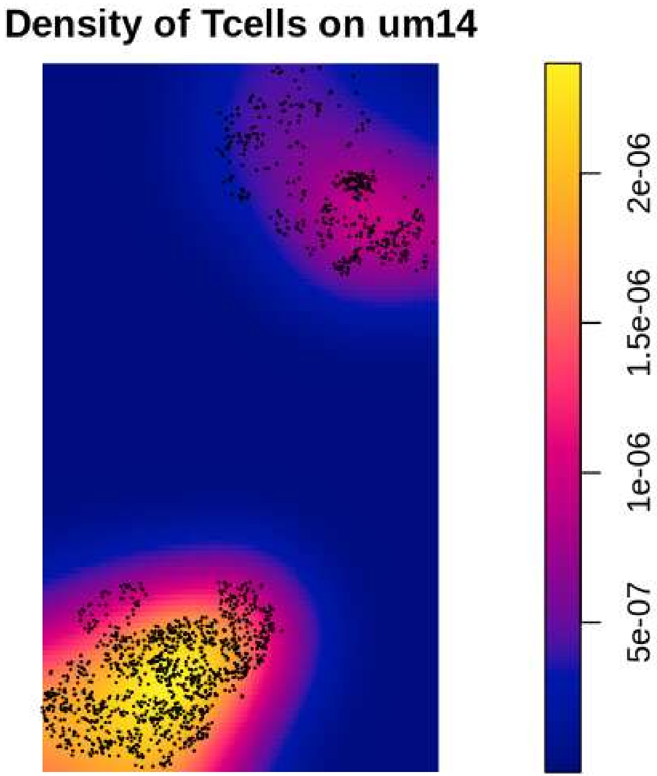
Density of Tcells on um14
2e-06
1.5e-06
1e-06
5e-07

Density of vasculature on um14

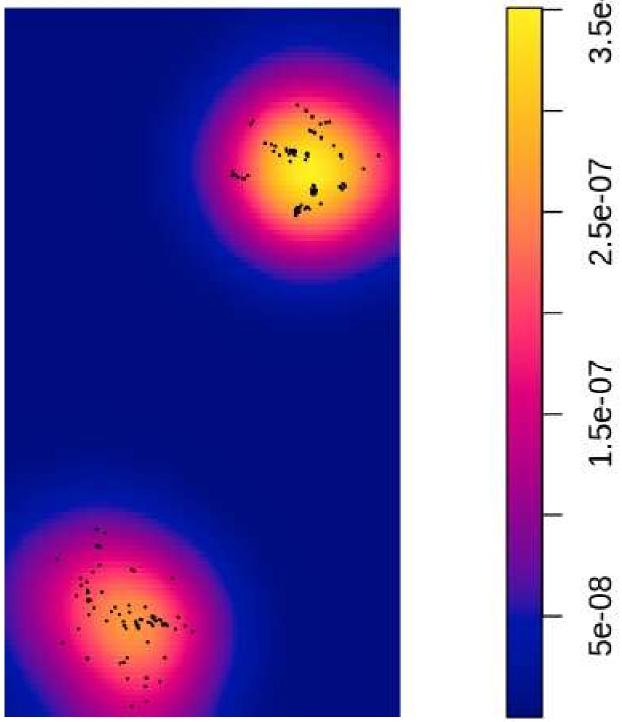

#### Density of Macrophage on um15

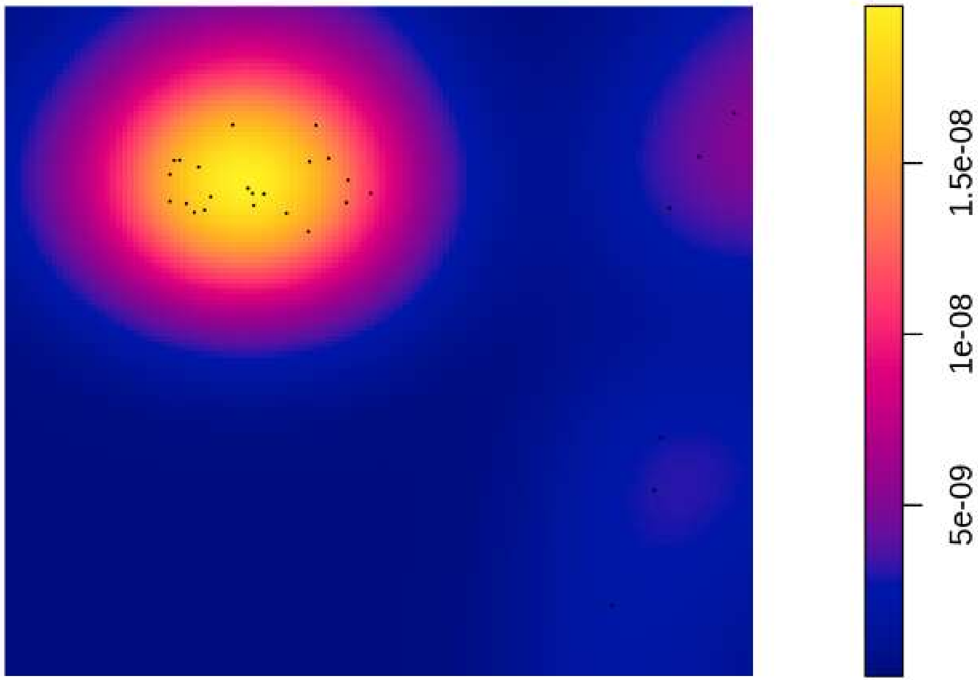

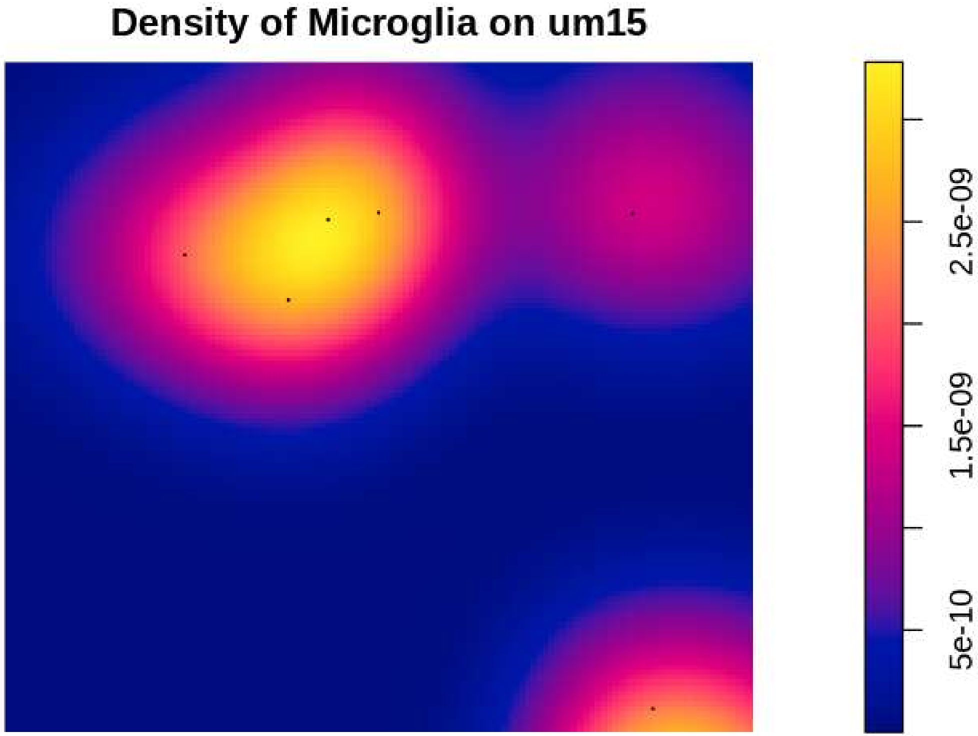
Density of Microglia on um15
5e-10
1.5e-09
2.5e-09

#### Density of Monocyte on um15

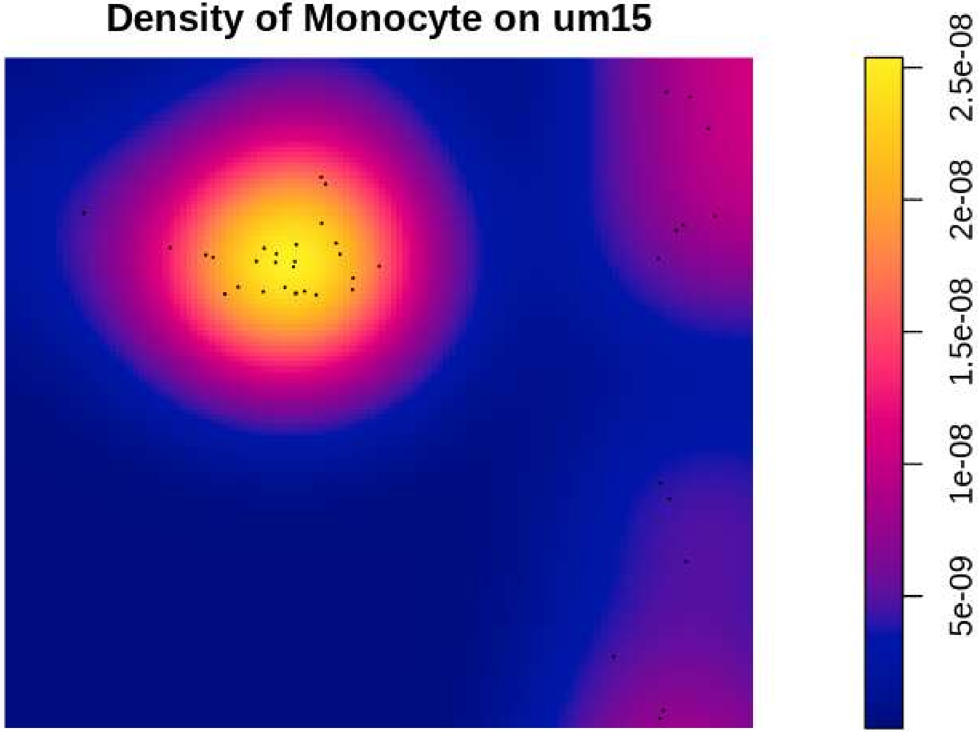

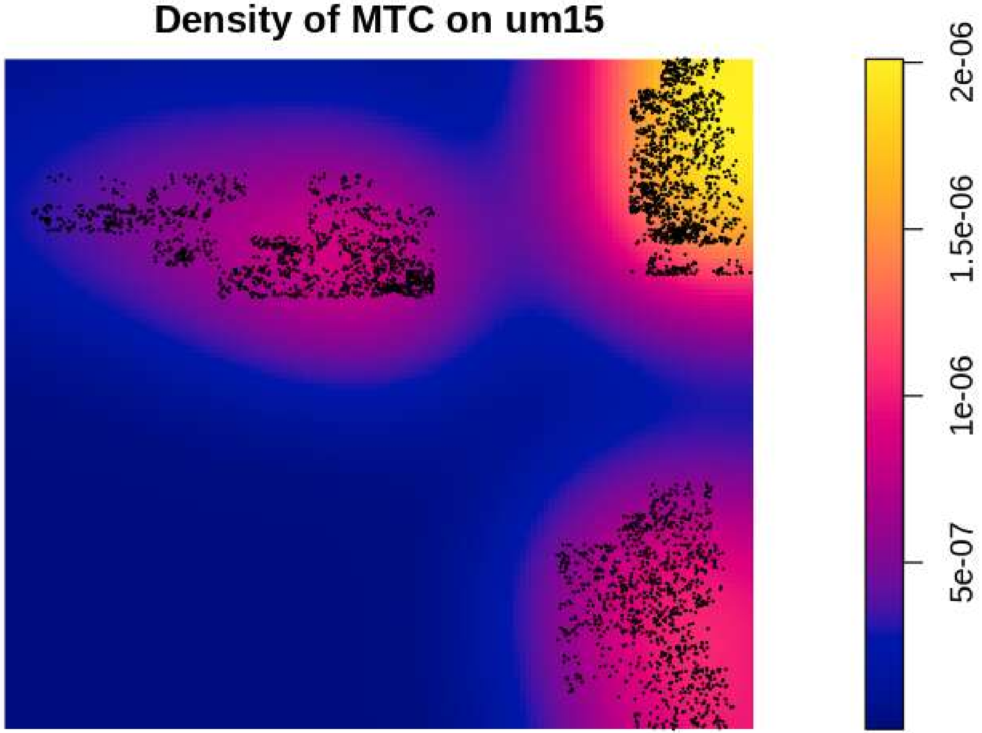
Density of MTC on um15
2e-06
1.5e-06
1e-06
5e-07

#### Density of NEU on um15

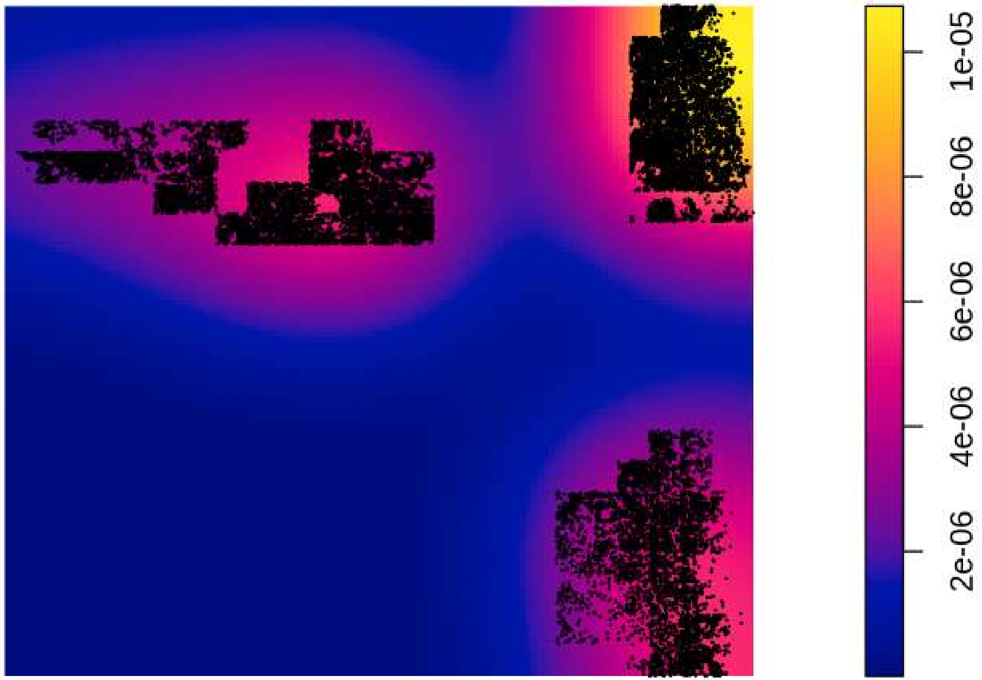

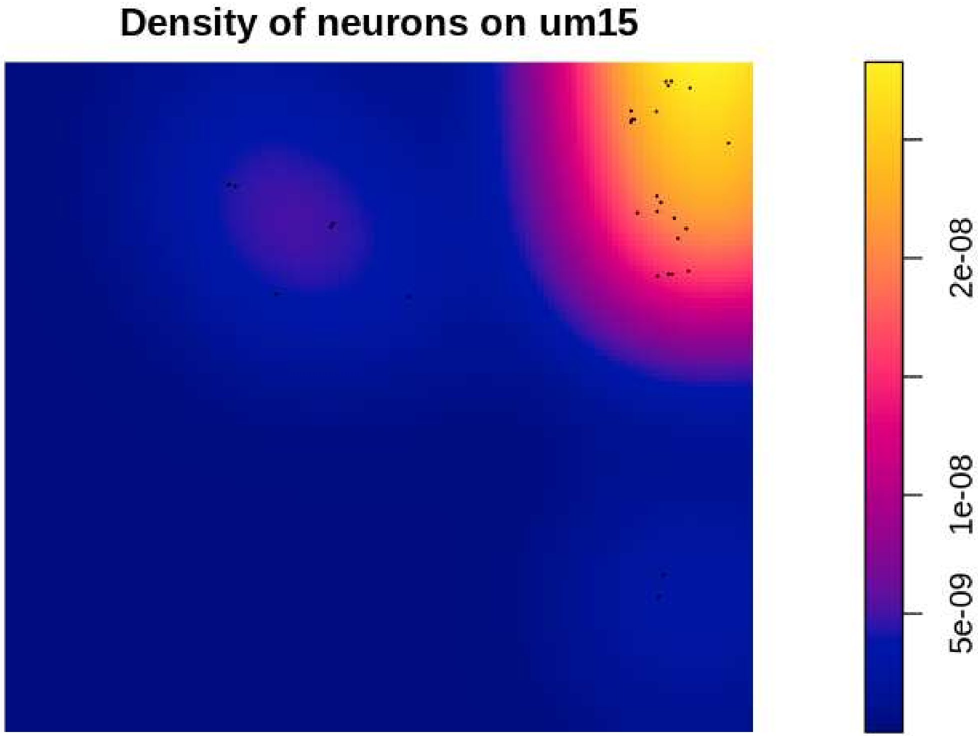
Density of neurons on um15
5e-09
1e-08
2e-08

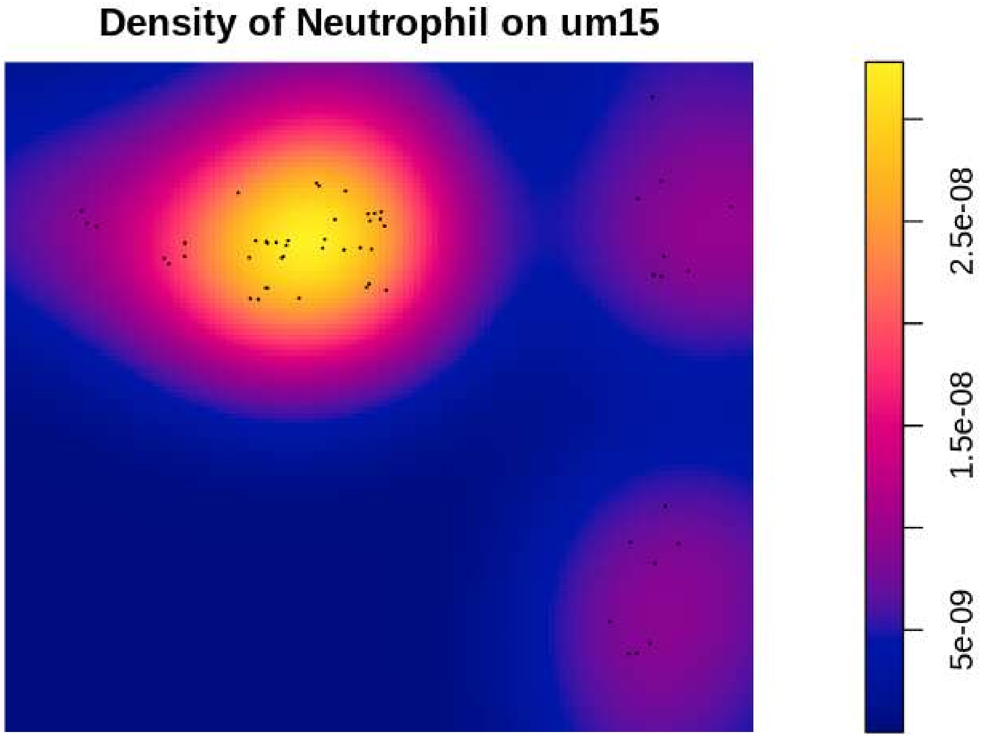
Density of Neutrophil on um15
5e-09
1.5e-08
2.5e-08

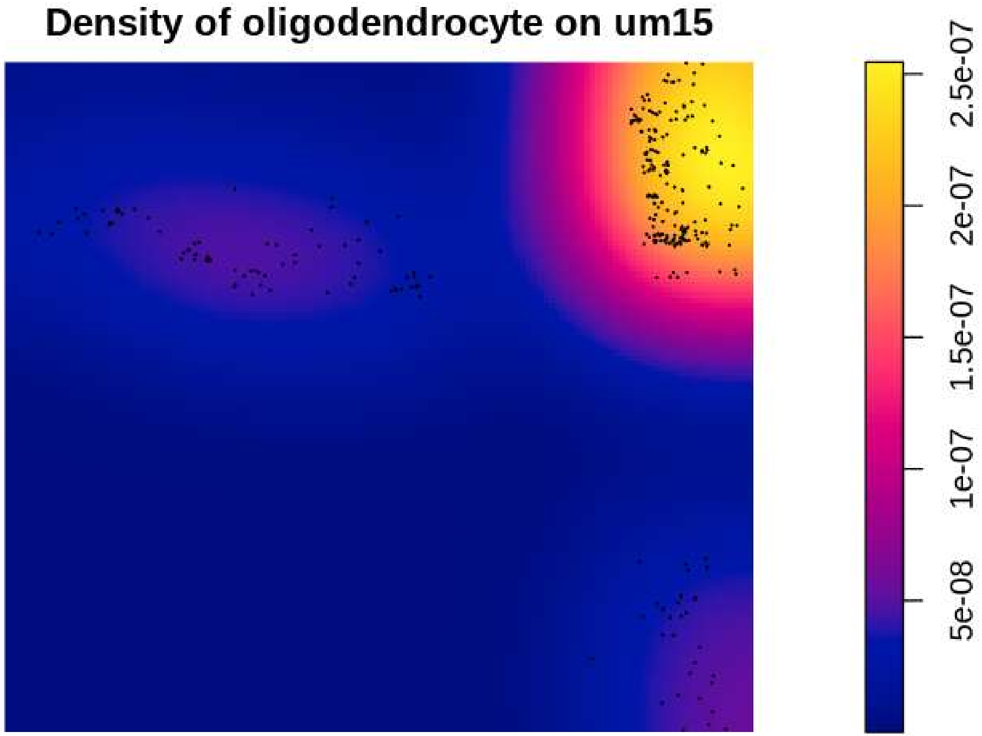
Density of oligodendrocyte on um15
2.5e-07
2e-07
1.5e-07
1e-07
5e-08

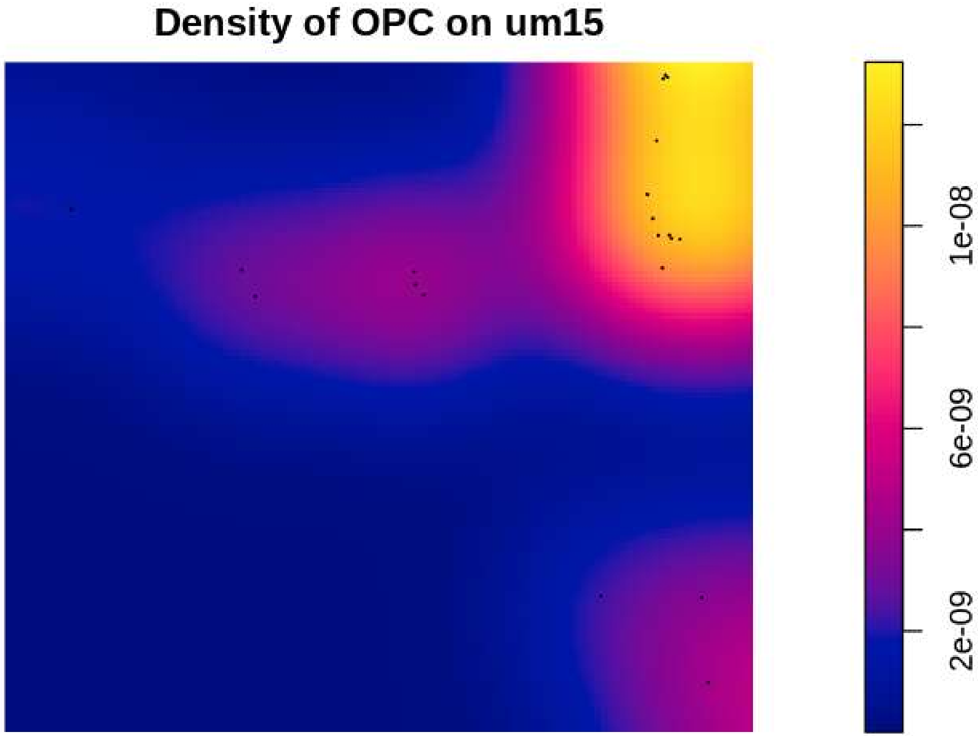

Density of OPC on um15
1e-08
6e-09
2e-09

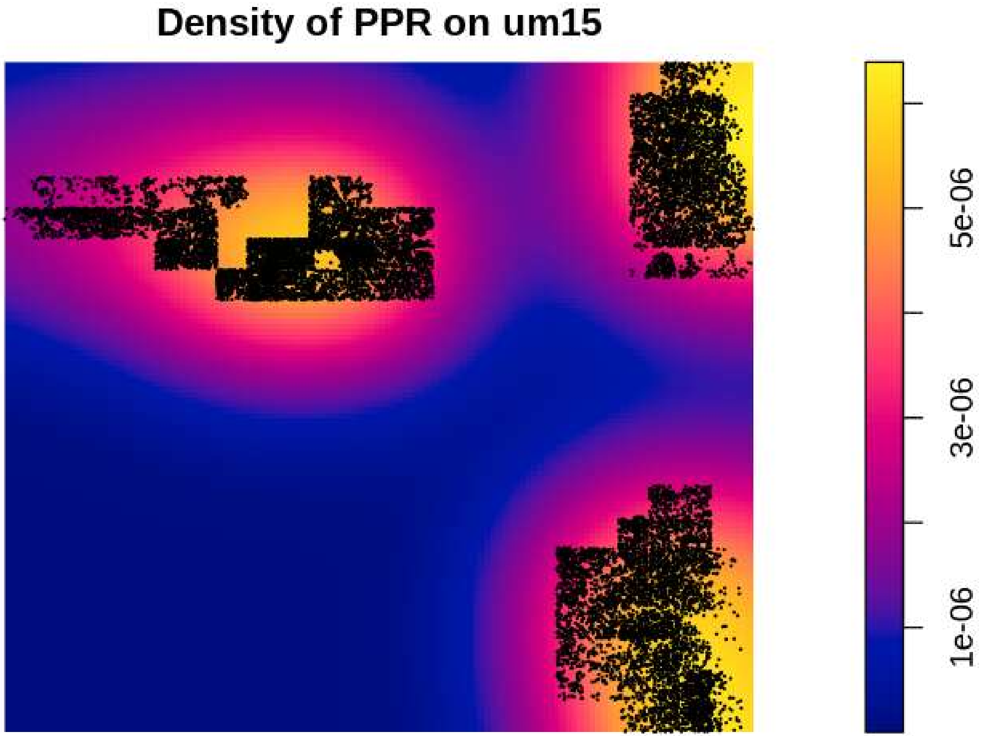

Density of PPR on um15
1e-06
3e-06
5e-06

#### Density of Tcells on um15

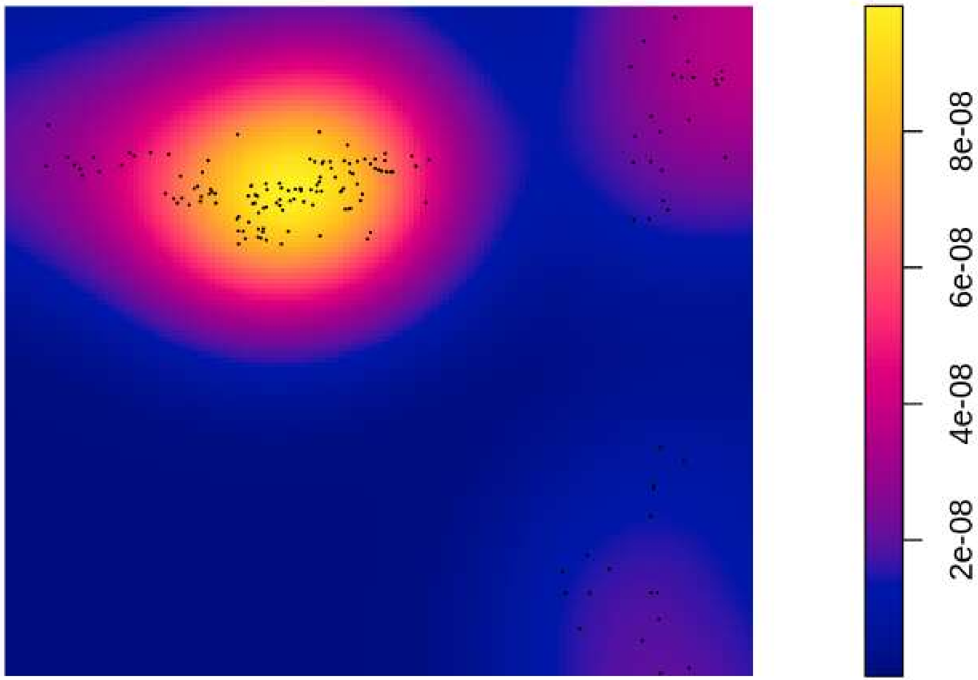

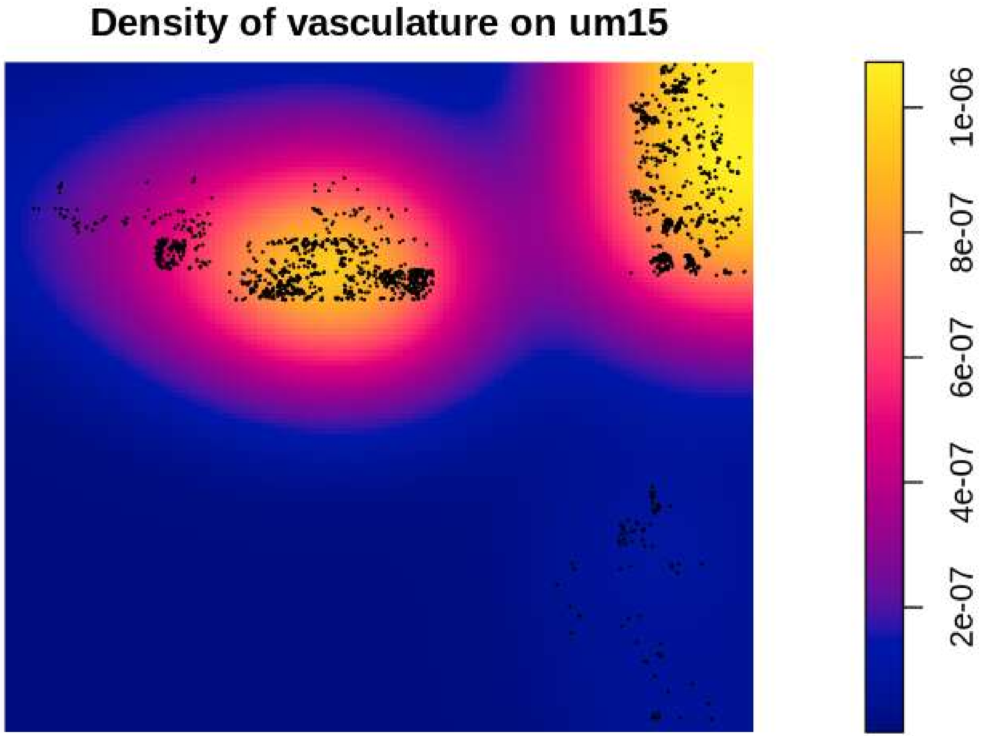
Density of vasculature on um15
1e-06
8e-07
6e-07
4e-07
2e-07
