## Supplementary Figures 4 for "Point-Process Modelling of Cell-Type Interaction Structure in CosMx Glioma Data for Hypothesis-Driven Tumor Inference from scRNA-seq"

Relative Density - Patient: um08 | Target: GPM | Modeled: astrocytes

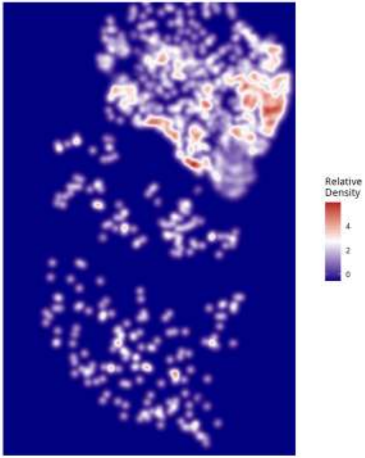

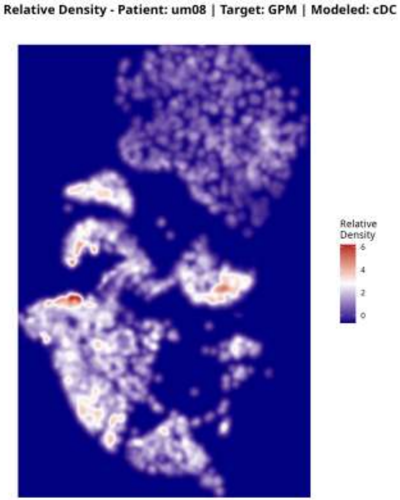
Relative Density - Patient: um08 | Target: GPM | Modeled: cDC
Relative
Density
6
4
2
0

Relative Density - Patient: um08 | Target: GPM | Modeled: GPM

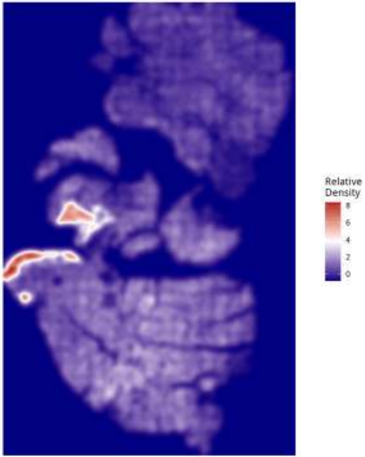

Relative Density - Patient: um08 | Target: GPM | Modeled: Macrophage

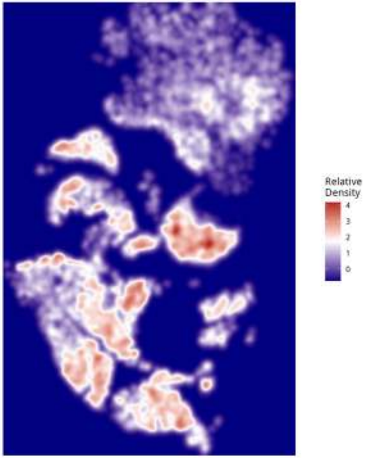

**Relative Density - Patient: um08 | Target: GPM | Modeled: Microglia**

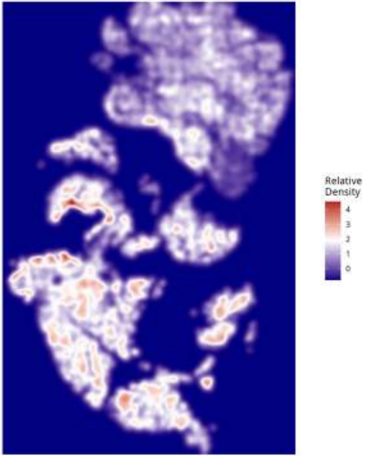

**Relative Density - Patient: um08 | Target: GPM | Modeled: Monocyte**

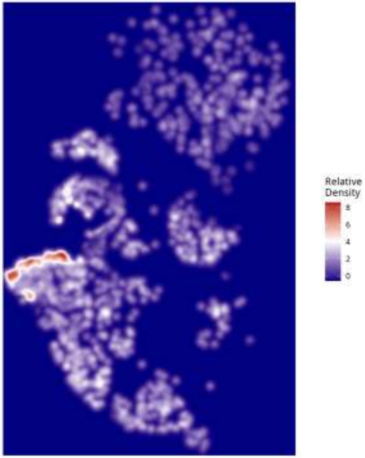

**Relative Density - Patient: um08 | Target: GPM | Modeled: MTC**

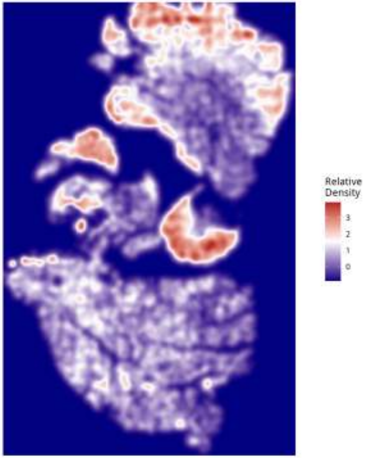

Relative Density - Patient: um08 | Target: GPM | Modeled: NEU

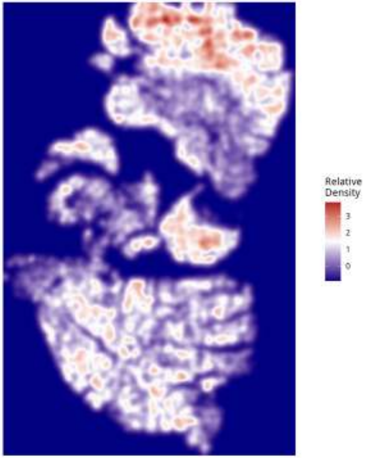

Relative Density - Patient: um08 | Target: GPM | Modeled: neurons

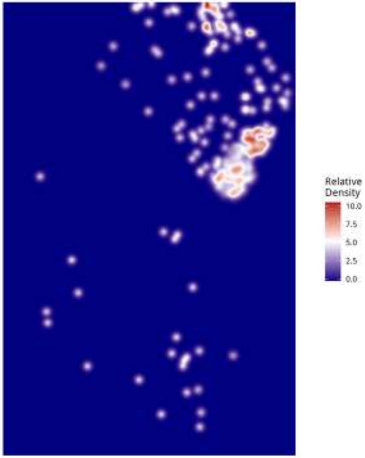

Relative Density - Patient: um08 | Target: GPM | Modeled: Neutrophil

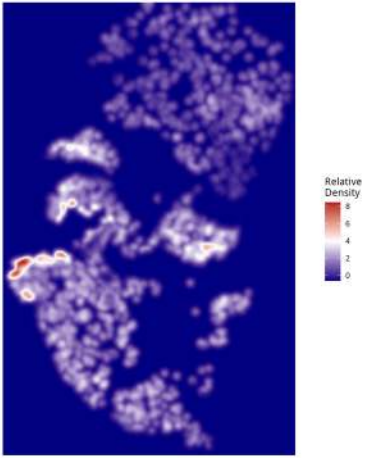

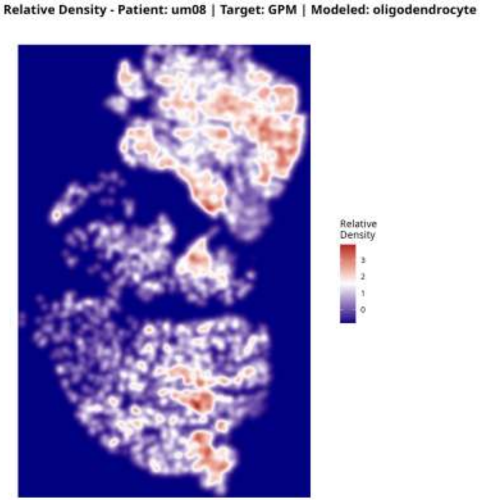

Relative Density - Patient: um08 | Target: GPM | Modeled: oligodendrocyte
Relative Density
3
2
1
0

Relative Density - Patient: um08 | Target: GPM | Modeled: OPC

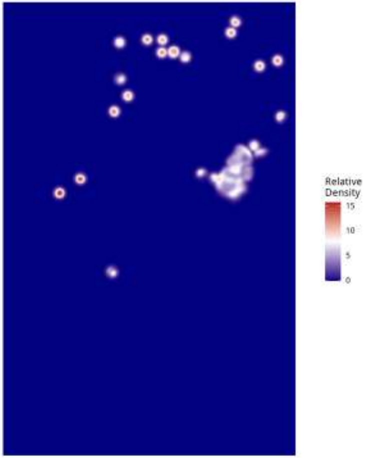

Relative Density - Patient: um08 | Target: GPM | Modeled: PPR

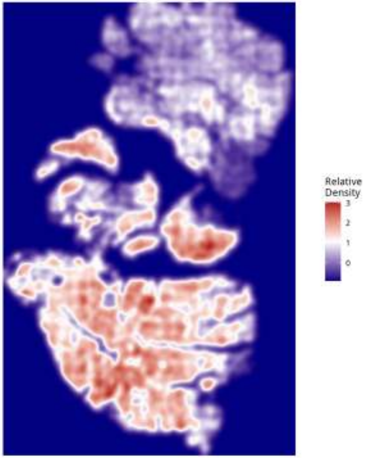

Relative Density - Patient: um08 | Target: GPM | Modeled: Tcells

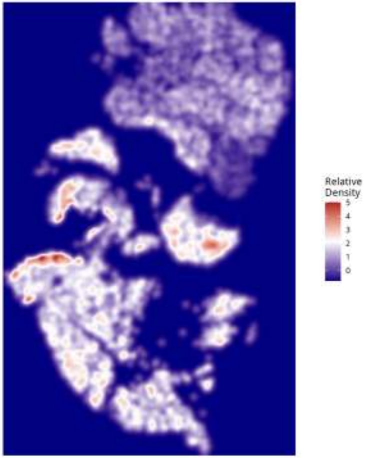

**Relative Density - Patient: um08 | Target: GPM | Modeled: vasculature**

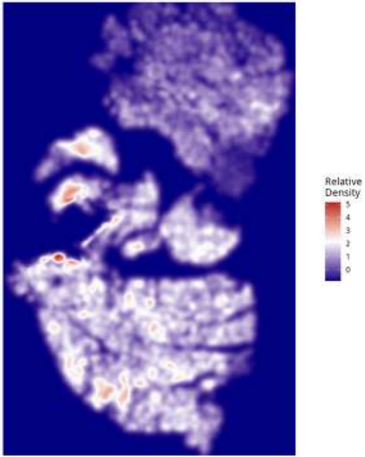

**Relative Density - Patient: um08 | Target: MTC | Modeled: astrocytes**

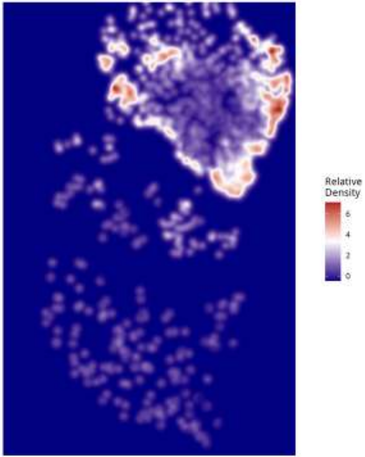

Relative Density - Patient: um08 | Target: MTC | Modeled: cDC

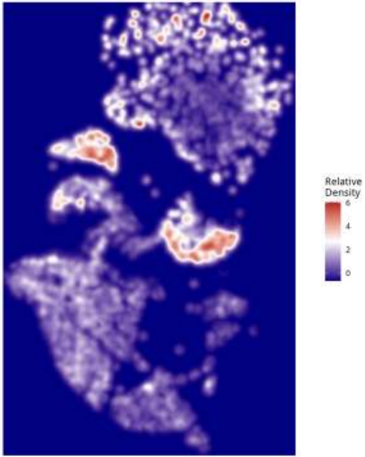

Relative Density - Patient: um08 | Target: MTC | Modeled: GPM

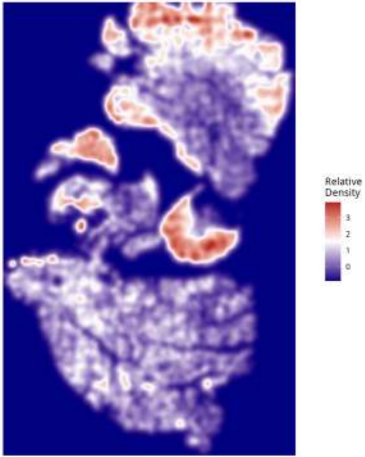

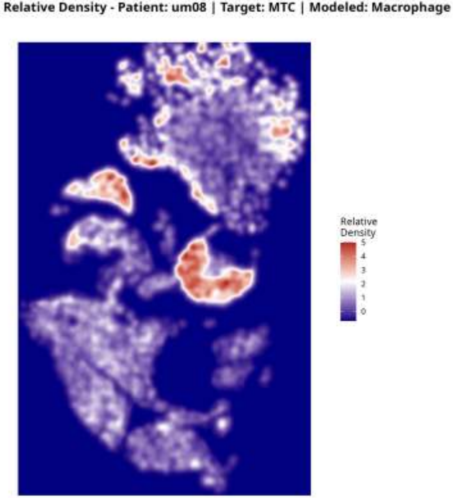

Relative Density - Patient: um08 | Target: MTC | Modeled: Macrophage
Relative Density
5
4
3
2
1
0

Relative Density - Patient: um08 | Target: MTC | Modeled: Microglia

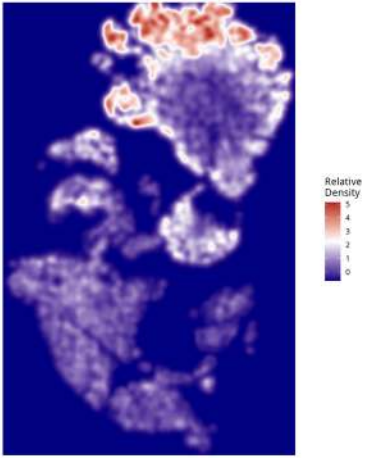

Relative Density - Patient: um08 | Target: MTC | Modeled: Monocyte

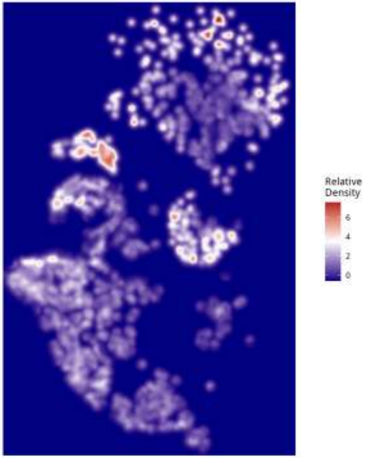

**Relative Density - Patient: um08 | Target: MTC | Modeled: MTC**

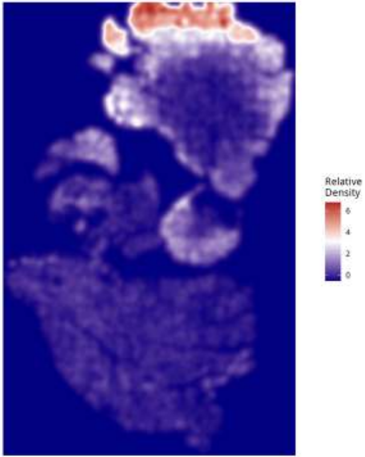

Relative Density - Patient: um08 | Target: MTC | Modeled: NEU

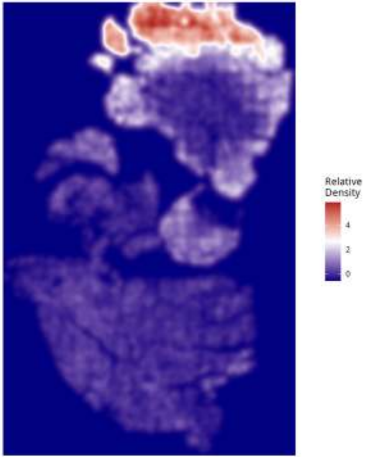

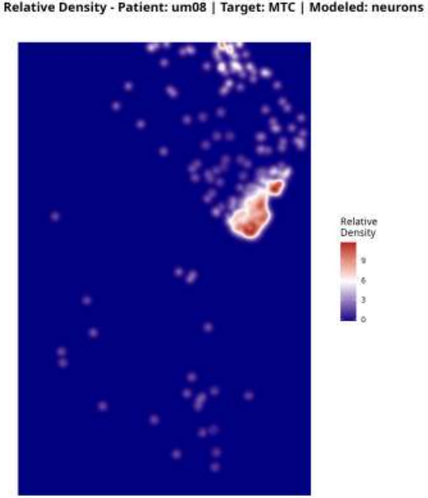
Relative Density - Patient: um08 | Target: MTC | Modeled: neurons
Relative
Density
9
6
3
0

**Relative Density - Patient: um08 | Target: MTC | Modeled: Neutrophil**

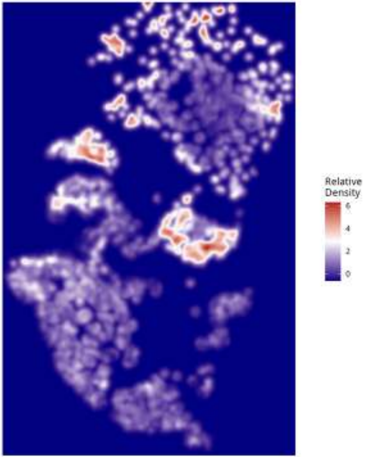

Relative Density - Patient: um08 | Target: MTC | Modeled: oligodendrocyte

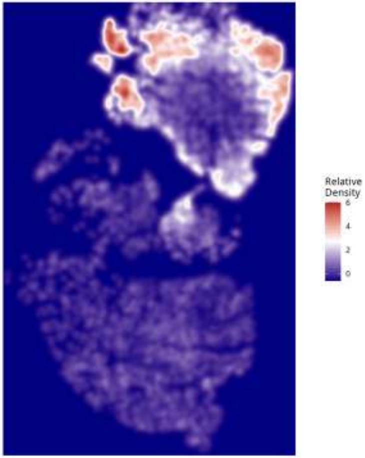

Relative Density - Patient: um08 | Target: MTC | Modeled: OPC

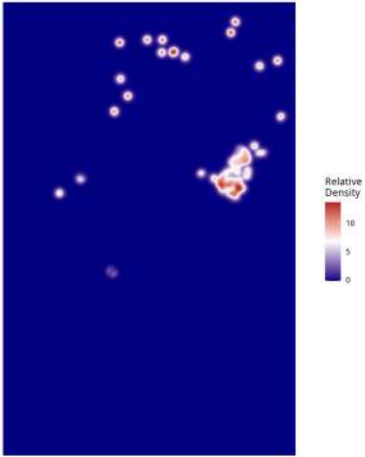

Relative Density - Patient: um08 | Target: MTC | Modeled: PPR

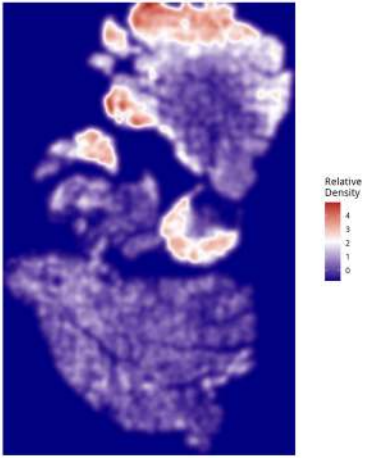

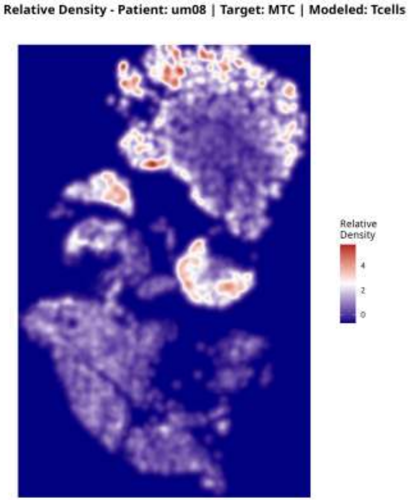
Relative Density - Patient: um08 | Target: MTC | Modeled: Tcells
Relative Density
4
2
0

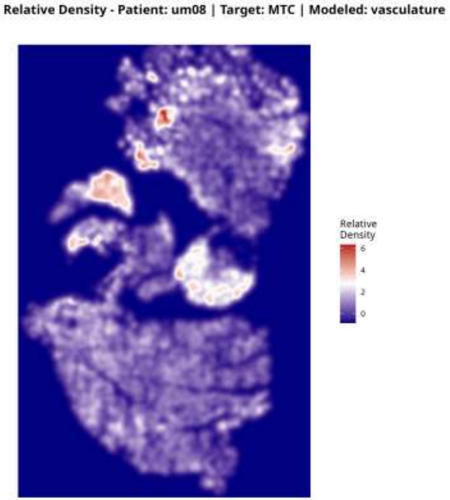
Relative Density - Patient: um08 | Target: MTC | Modeled: vasculature
Relative Density
6
4
2
0

### Relative Density - Patient: um08 | Target: NEU | Modeled: astrocytes

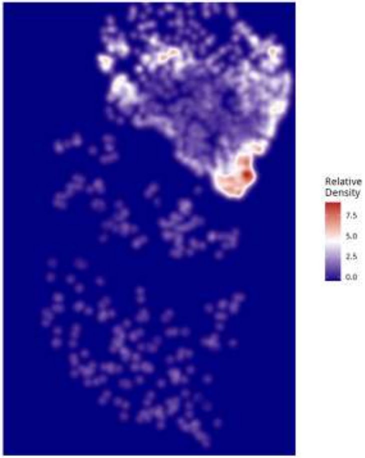

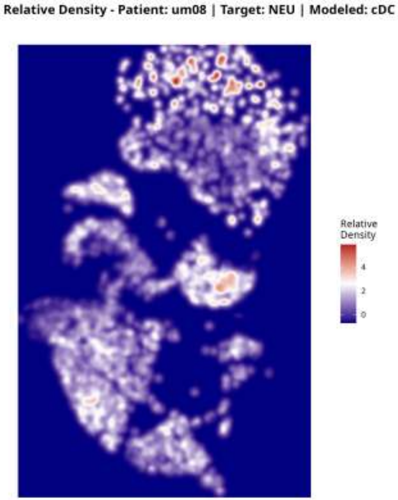

Relative Density - Patient: um08 | Target: NEU | Modeled: cDC
Relative Density
4
2
0

**Relative Density - Patient: um08 | Target: NEU | Modeled: GPM**

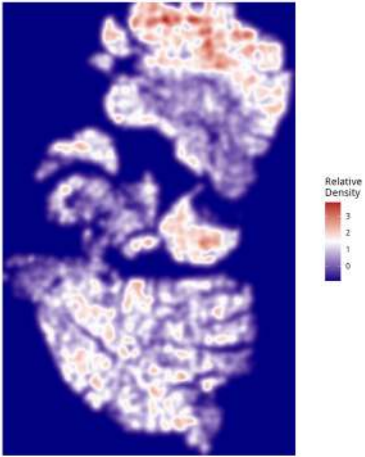

Relative Density - Patient: um08 | Target: NEU | Modeled: Macrophage

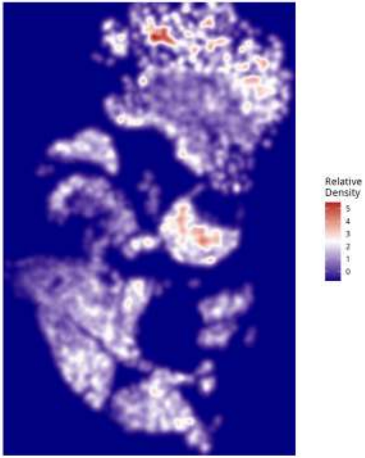

**Relative Density - Patient: um08 | Target: NEU | Modeled: Microglia**

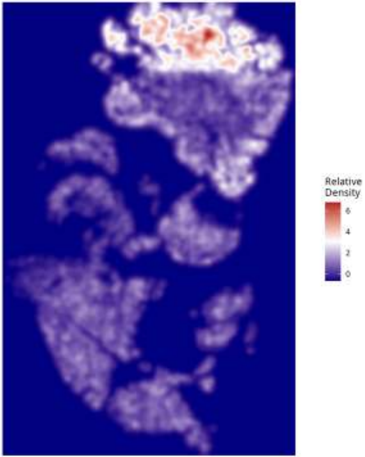

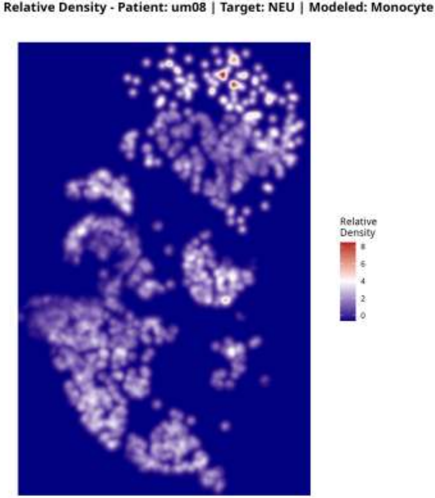

Relative Density - Patient: um08 | Target: NEU | Modeled: Monocyte
Relative Density
8
6
4
2
0

Relative Density - Patient: um08 | Target: NEU | Modeled: MTC

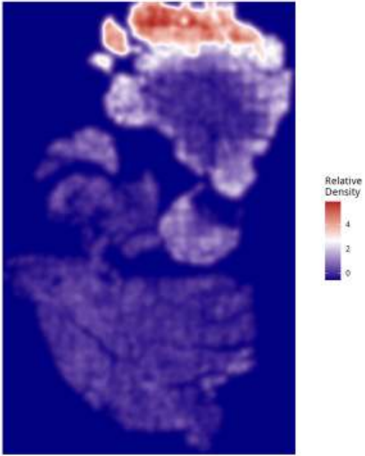

Relative Density - Patient: um08 | Target: NEU | Modeled: NEU

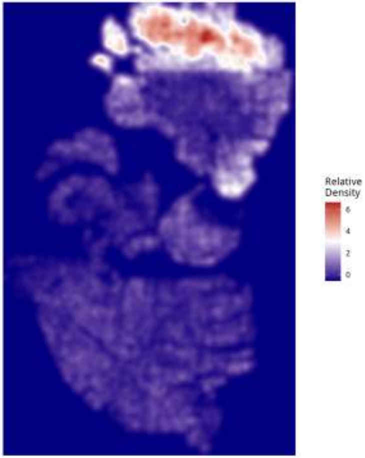

**Relative Density - Patient: um08 | Target: NEU | Modeled: neurons**

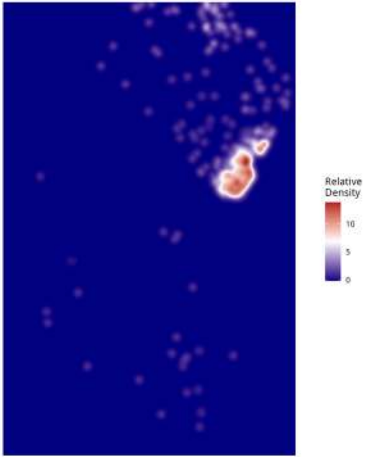

**Relative Density - Patient: um08 | Target: NEU | Modeled: Neutrophil**

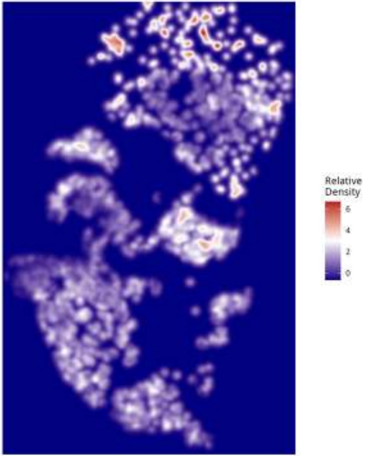

Relative Density - Patient: um08 | Target: NEU | Modeled: oligodendrocyte

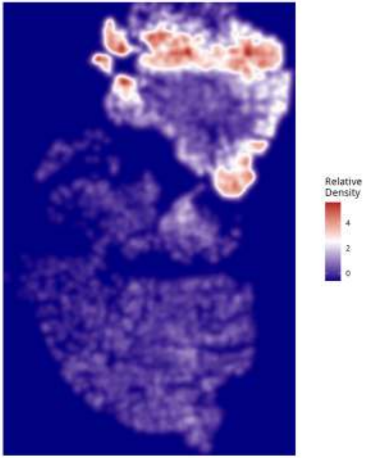

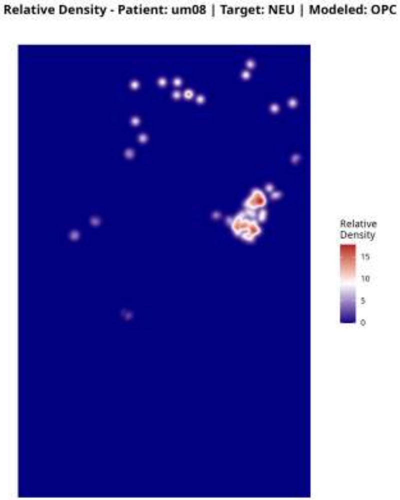

Relative Density - Patient: um08 | Target: NEU | Modeled: OPC
Relative Density
15
10
5
0

Relative Density - Patient: um08 | Target: NEU | Modeled: PPR

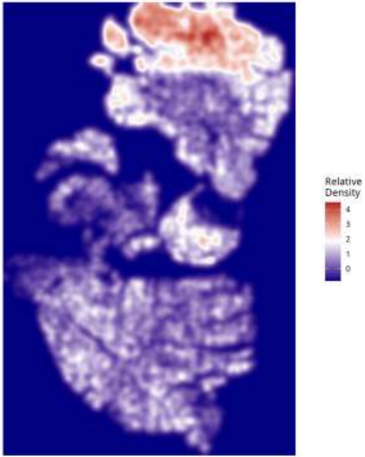

Relative Density - Patient: um08 | Target: NEU | Modeled: Tcells

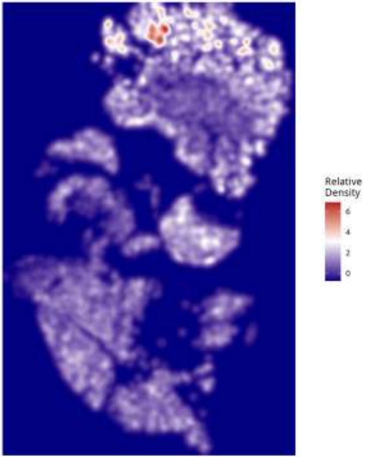

Relative Density - Patient: um08 | Target: NEU | Modeled: vasculature

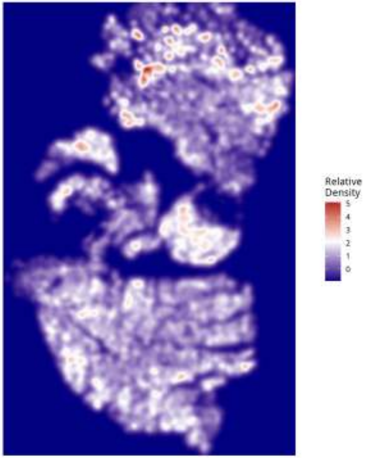

Relative Density - Patient: um08 | Target: PPR | Modeled: astrocytes

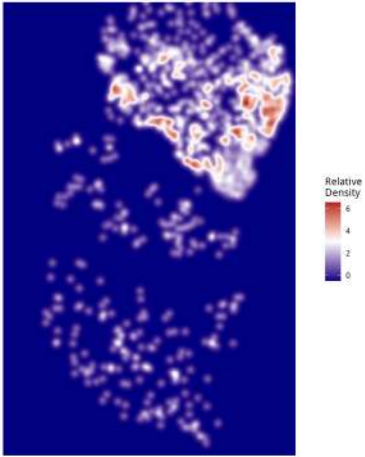

Relative Density - Patient: um08 | Target: PPR | Modeled: cDC

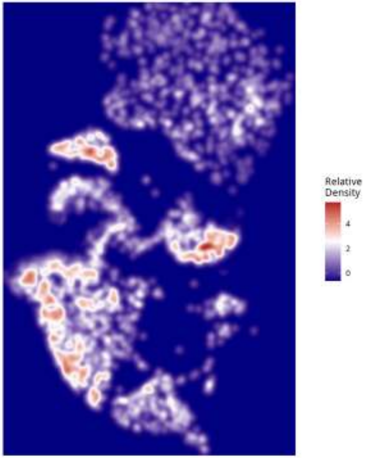

**Relative Density - Patient: um08 | Target: PPR | Modeled: GPM**

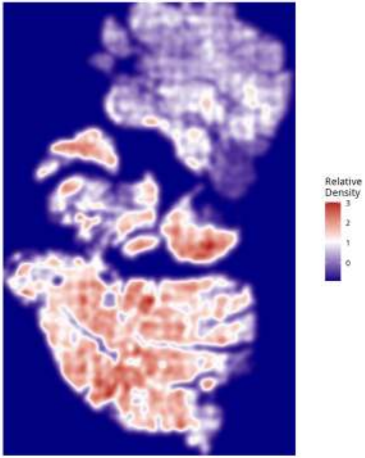

Relative Density - Patient: um08 | Target: PPR | Modeled: Macrophage

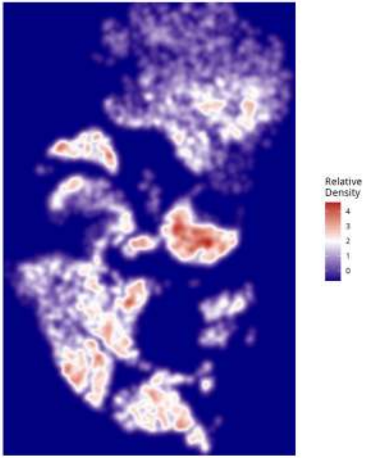

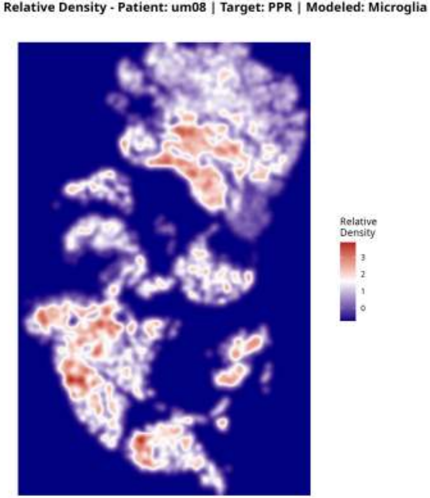
Relative Density - Patient: um08 | Target: PPR | Modeled: Microglia
Relative Density
3
2
1
0

**Relative Density - Patient: um08 | Target: PPR | Modeled: Monocyte**

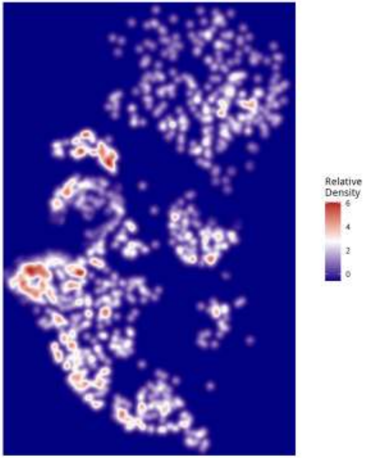

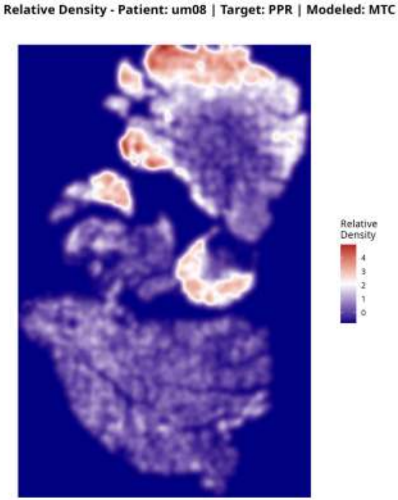
Relative Density - Patient: um08 | Target: PPR | Modeled: MTC
Relative Density
4
3
2
1
0

Relative Density - Patient: um08 | Target: PPR | Modeled: NEU

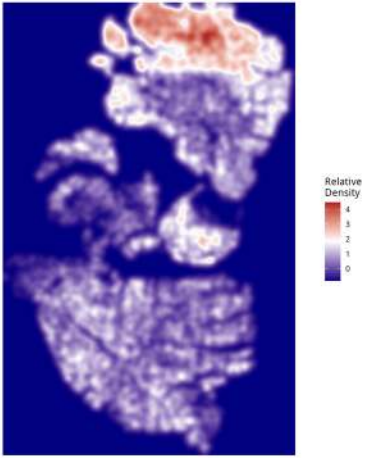

Relative Density - Patient: um08 | Target: PPR | Modeled: neurons

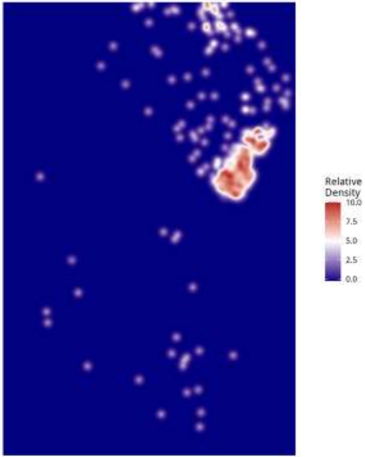

**Relative Density - Patient: um08 | Target: PPR | Modeled: Neutrophil**

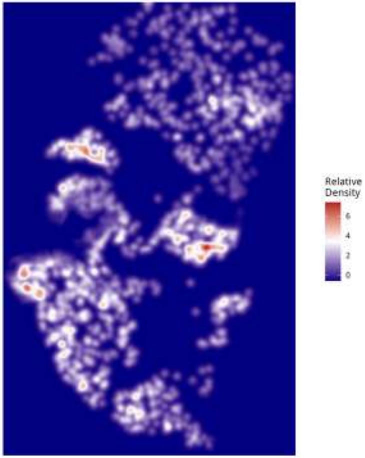

**Relative Density - Patient: um08 | Target: PPR | Modeled: oligodendrocyte**

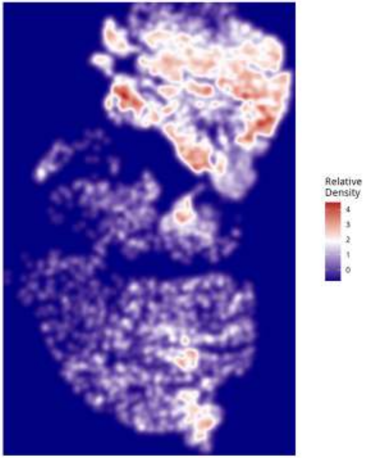

Relative Density - Patient: um08 | Target: PPR | Modeled: OPC

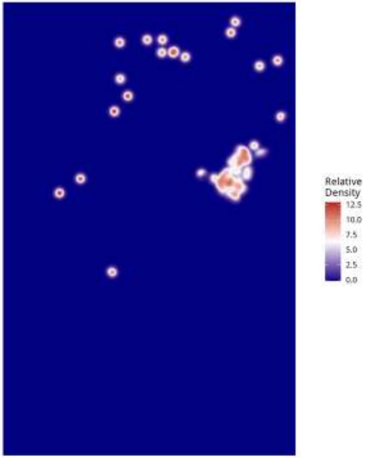

Relative Density - Patient: um08 | Target: PPR | Modeled: PPR

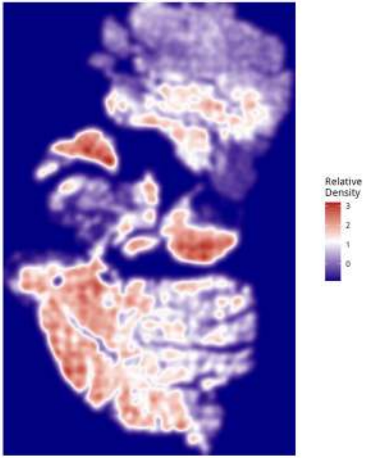

**Relative Density - Patient: um08 | Target: PPR | Modeled: Tcells**

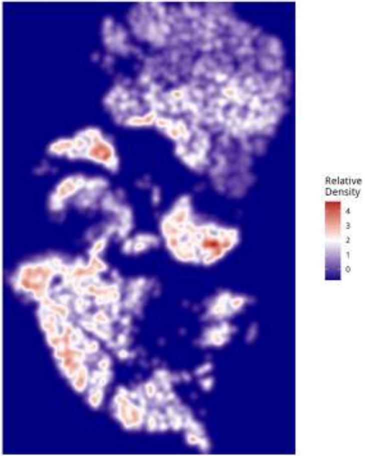

Relative Density - Patient: um08 | Target: PPR | Modeled: vasculature

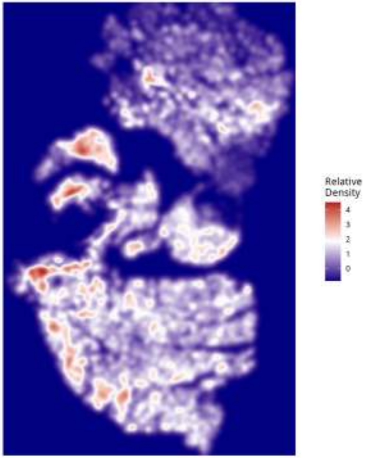

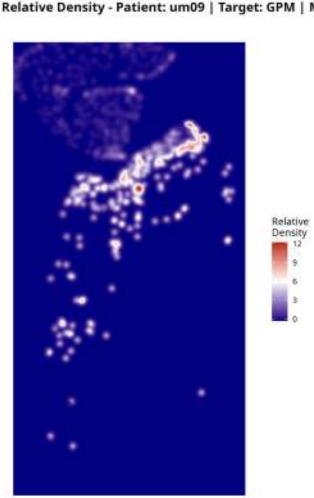
Relative Density - Patient: um09 | Target: GPM | Modeled: astrocytes
Relative Density
12
9
6
3
0

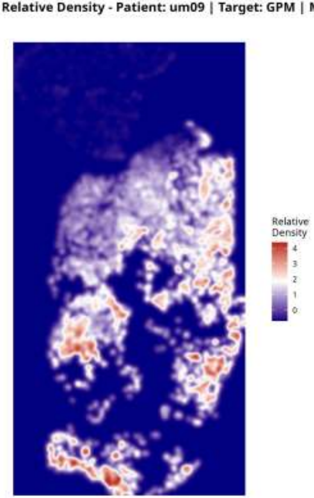
Relative Density - Patient: um09 | Target: GPM | Modeled: cDC
Relative Density
4
3
2
1
0

Relative Density - Patient: um09 | Target: GPM | Modeled: GPM

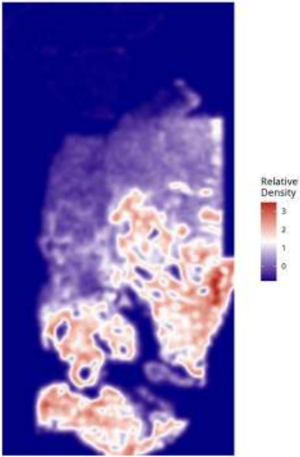

Relative Density - Patient: um09 | Target: GPM | Modeled: Macrophage

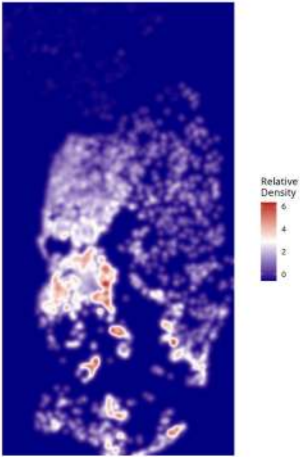

Relative Density - Patient: um09 | Target: GPM | Modeled: Microglia

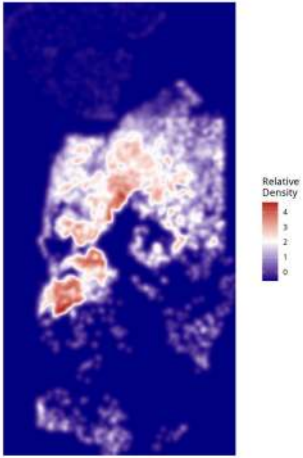

**Relative Density - Patient: um09 | Target: GPM | Modeled: Monocyte**

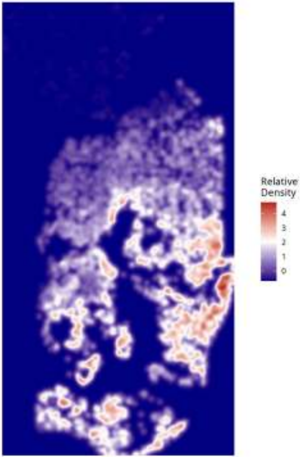

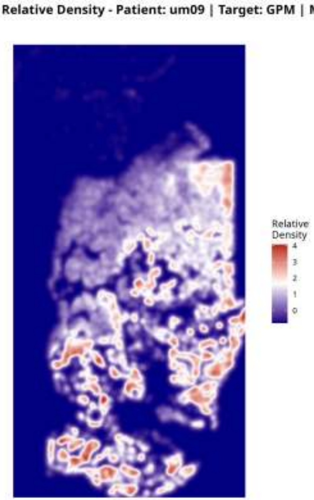
Relative Density - Patient: um09 | Target: GPM | Modeled: MTC
Relative
Density
4
3
2
1
0

Relative Density - Patient: um09 | Target: GPM | Modeled: NEU

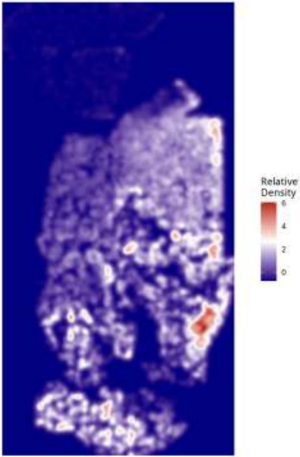

**Relative Density - Patient: um09 | Target: GPM | Modeled: neurons**

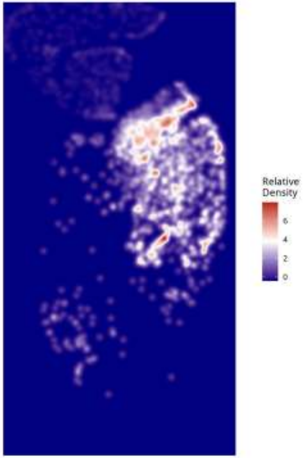

Relative Density - Patient: um09 | Target: GPM | Modeled: Neutrophil

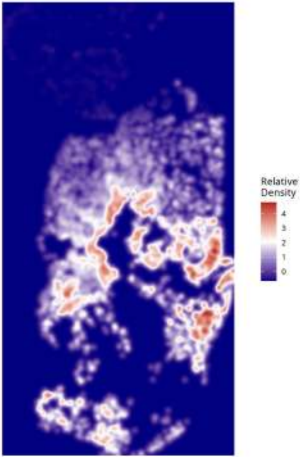

Relative Density - Patient: um09 | Target: GPM | Modeled: oligodendrocyte

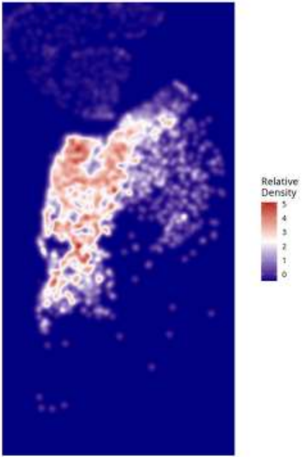

#### Relative Density - Patient: um09 | Target: GPM | Modeled: OPC

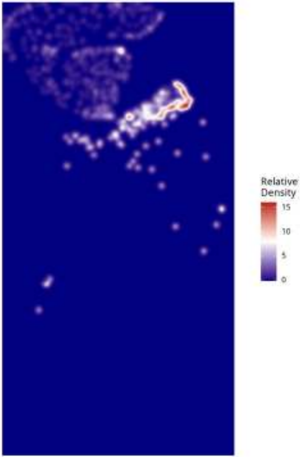

**Relative Density - Patient: um09 | Target: GPM | Modeled: PPR**

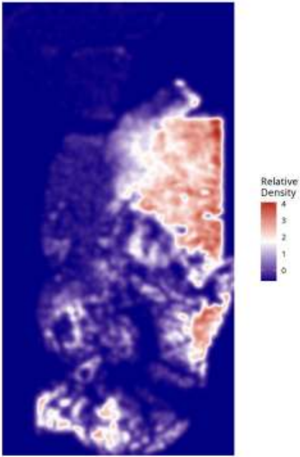

Relative Density - Patient: um09 | Target: GPM | Modeled: Tcells

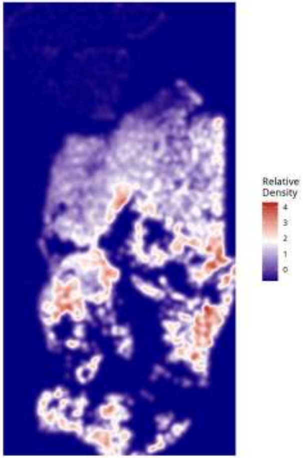

#### Relative Density - Patient: um09 | Target: GPM | Modeled: vasculature

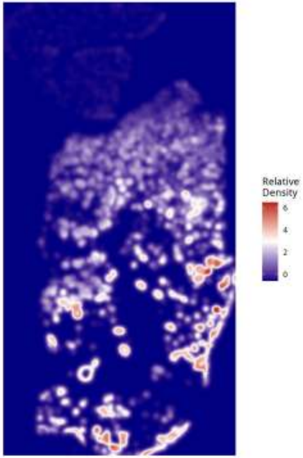

**Relative Density - Patient: um09 | Target: MTC | Modeled: astrocytes**

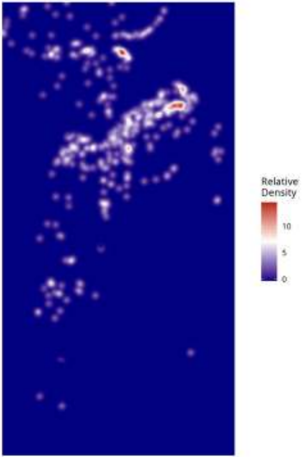

Relative Density - Patient: um09 | Target: MTC | Modeled: cDC

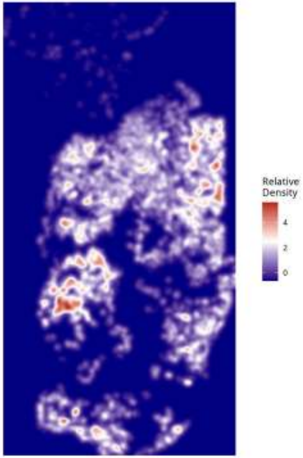

Relative Density - Patient: um09 | Target: MTC | Modeled: GPM

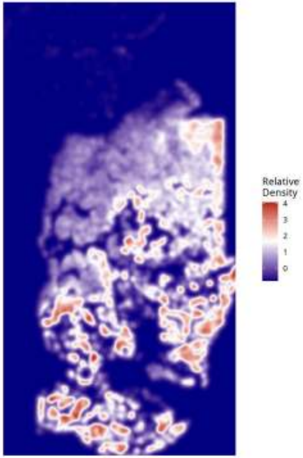

Relative Density - Patient: um09 | Target: MTC | Modeled: Macrophage

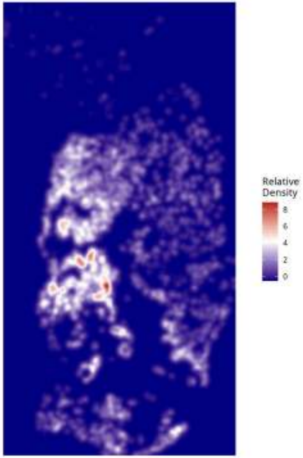

Relative Density - Patient: um09 | Target: MTC | Modeled: Microglia

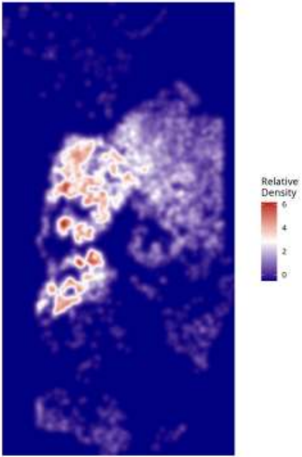

Relative Density - Patient: um09 | Target: MTC | Modeled: Monocyte

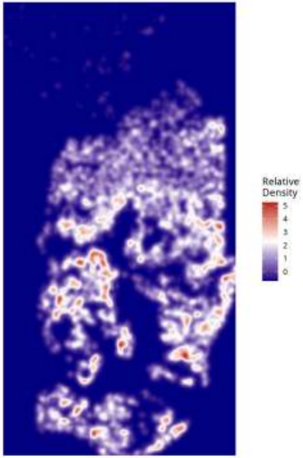

Relative Density - Patient: um09 | Target: MTC | Modeled: MTC

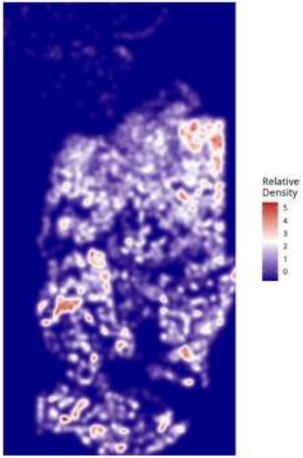

Relative Density - Patient: um09 | Target: MTC | Modeled: NEU

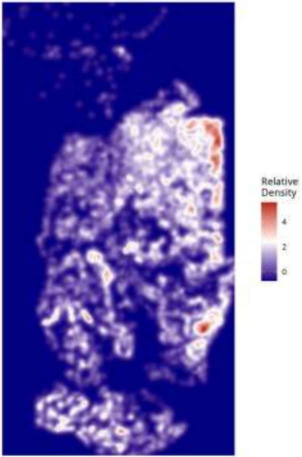

Relative Density - Patient: um09 | Target: MTC | Modeled: neurons

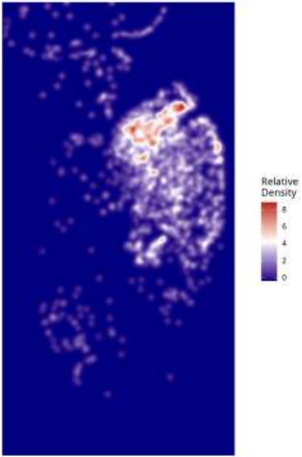

Relative Density - Patient: um09 | Target: MTC | Modeled: Neutrophil

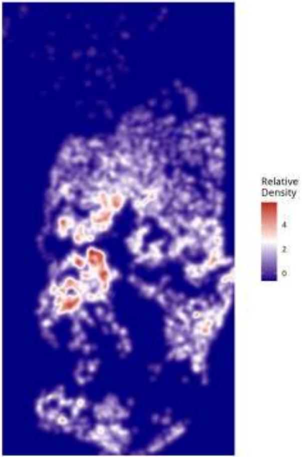

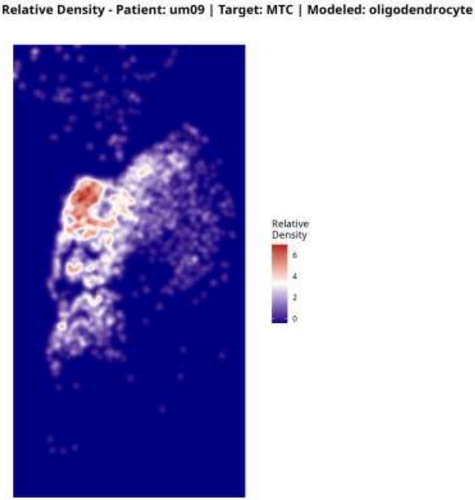

Relative Density - Patient: um09 | Target: MTC | Modeled: oligodendrocyte
Relative Density
6
4
2
0

Relative Density - Patient: um09 | Target: MTC | Modeled: OPC

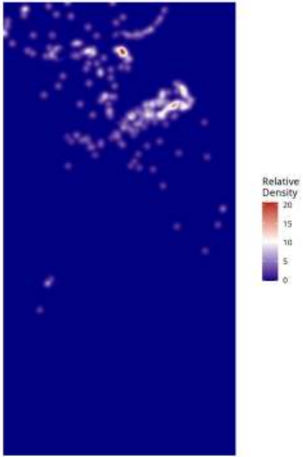

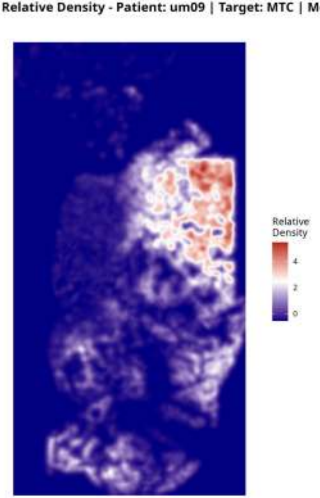
Relative Density - Patient: um09 | Target: MTC | Modeled: PPR
Relative Density
4
2
0

Relative Density - Patient: um09 | Target: MTC | Modeled: Tcells

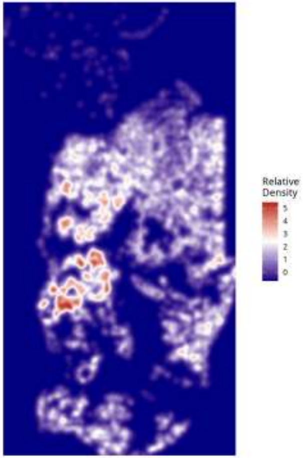

Relative Density - Patient: um09 | Target: MTC | Modeled: vasculature

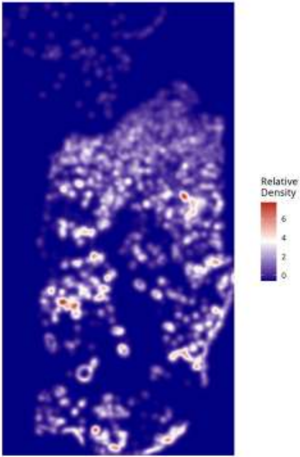

Relative Density - Patient: um09 | Target: NEU | Modeled: astrocytes

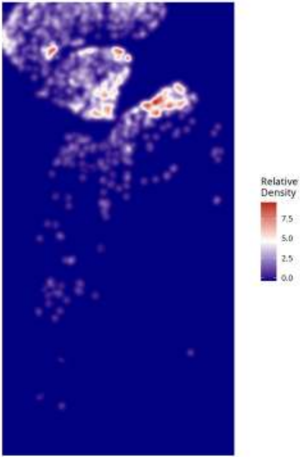

Relative Density - Patient: um09 | Target: NEU | Modeled: cDC

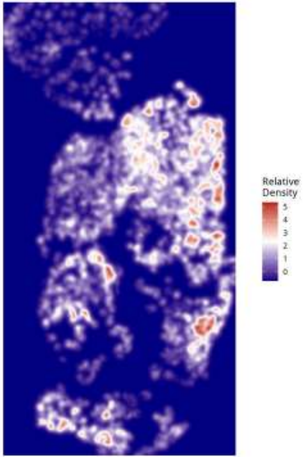

#### Relative Density - Patient: um09 | Target: NEU | Modeled: GPM

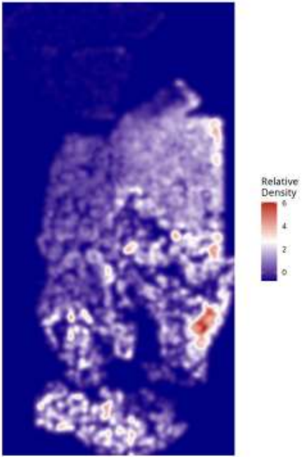

Relative Density - Patient: um09 | Target: NEU | Modeled: Macrophage

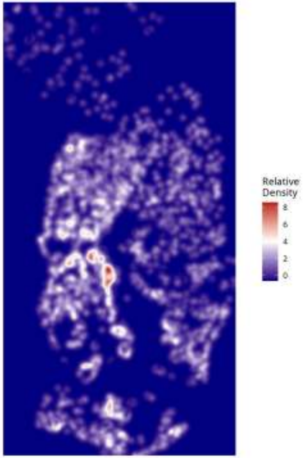

**Relative Density - Patient: um09 | Target: NEU | Modeled: Microglia**

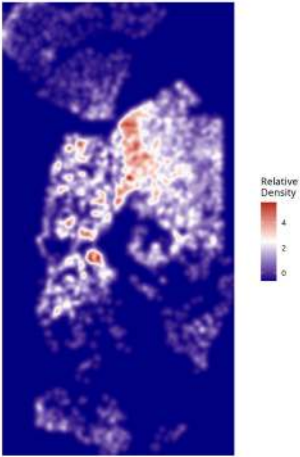

#### Relative Density - Patient: um09 | Target: NEU | Modeled: Monocyte

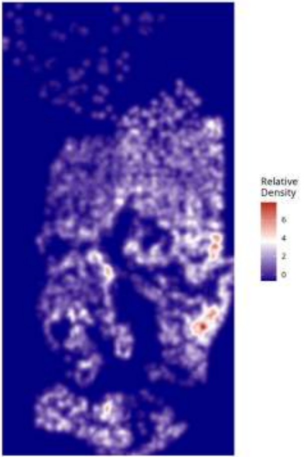

Relative Density - Patient: um09 | Target: NEU | Modeled: MTC

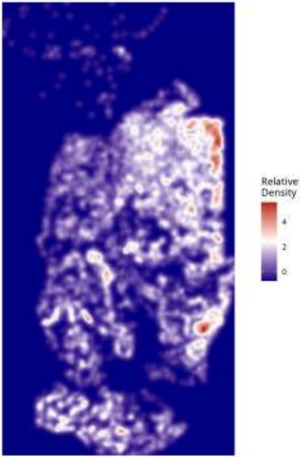

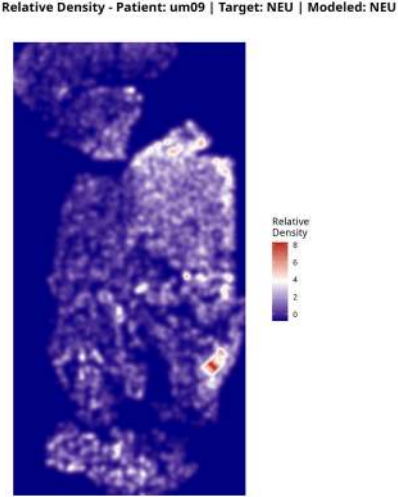
Relative Density - Patient: um09 | Target: NEU | Modeled: NEU
Relative Density
8
6
4
2
0

Relative Density - Patient: um09 | Target: NEU | Modeled: neurons

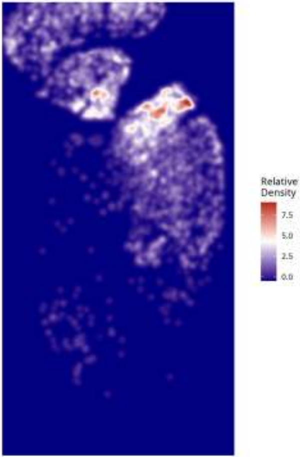

Relative Density - Patient: um09 | Target: NEU | Modeled: Neutrophil

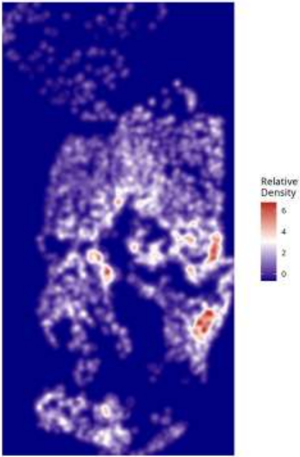

Relative Density - Patient: um09 | Target: NEU | Modeled: oligodendrocyte

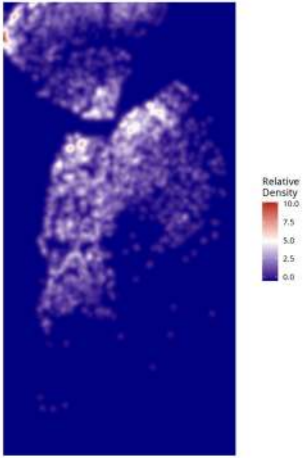

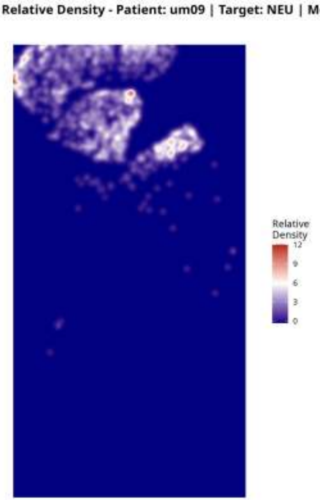
Relative Density - Patient: um09 | Target: NEU | Modeled: OPC
Relative
Density
12
9
6
3
0

Relative Density - Patient: um09 | Target: NEU | Modeled: PPR

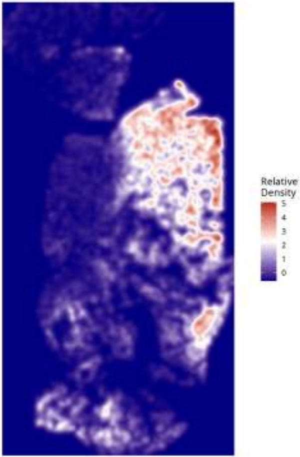

Relative Density - Patient: um09 | Target: NEU | Modeled: Tcells

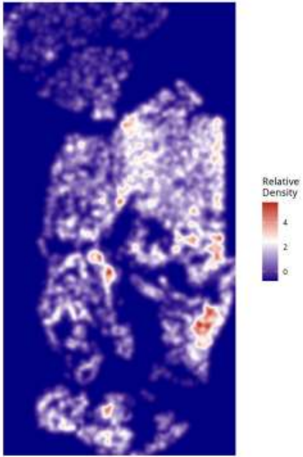

Relative Density - Patient: um09 | Target: NEU | Modeled: vasculature

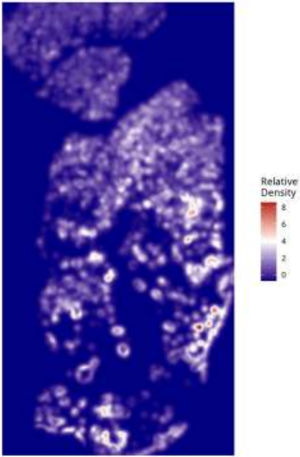

Relative Density - Patient: um09 | Target: PPR | Modeled: astrocytes

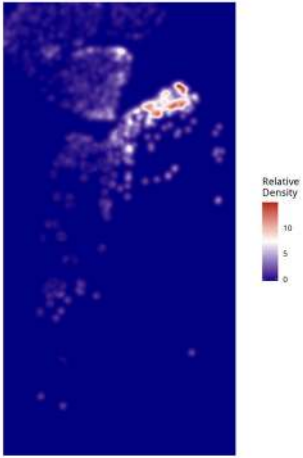

Relative Density - Patient: um09 | Target: PPR | Modeled: cDC

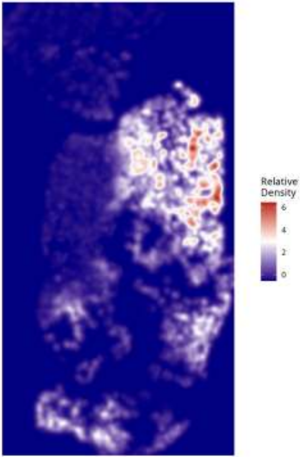

Relative Density - Patient: um09 | Target: PPR | Modeled: GPM

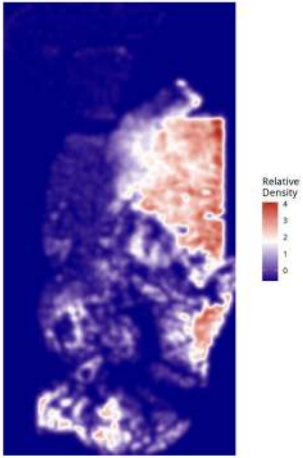

Relative Density - Patient: um09 | Target: PPR | Modeled: Macrophage

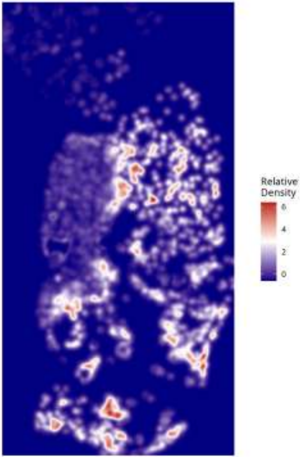

Relative Density - Patient: um09 | Target: PPR | Modeled: Microglia

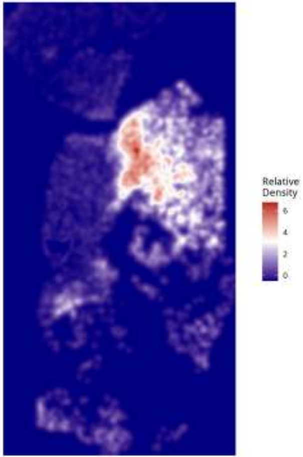

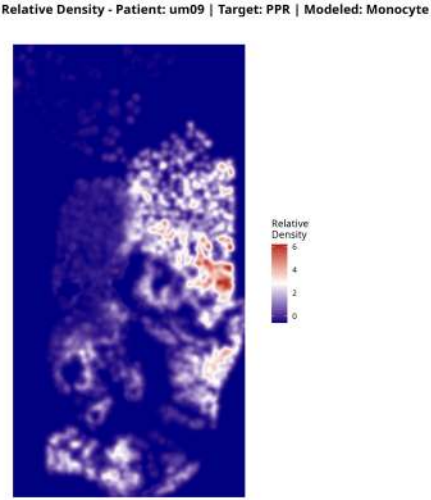
Relative Density - Patient: um09 | Target: PPR | Modeled: Monocyte
Relative Density
6
4
2
0

Relative Density - Patient: um09 | Target: PPR | Modeled: MTC

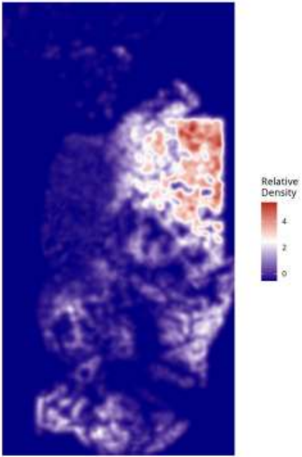

Relative Density - Patient: um09 | Target: PPR | Modeled: NEU

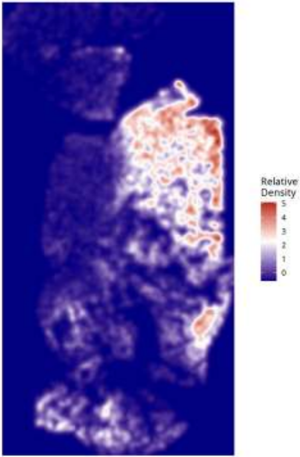

**Relative Density - Patient: um09 | Target: PPR | Modeled: neurons**

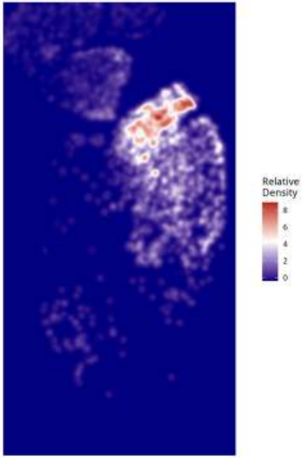

Relative Density - Patient: um09 | Target: PPR | Modeled: Neutrophil

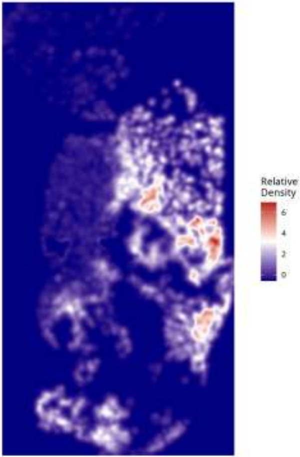

Relative Density - Patient: um09 | Target: PPR | Modeled: oligodendrocyte

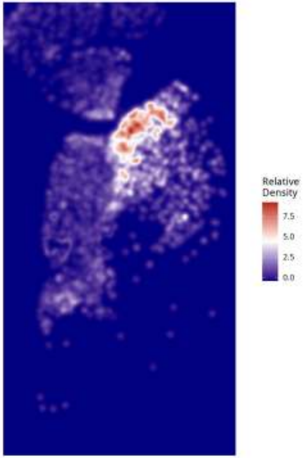

**Relative Density - Patient: um09 | Target: PPR | Modeled: OPC**

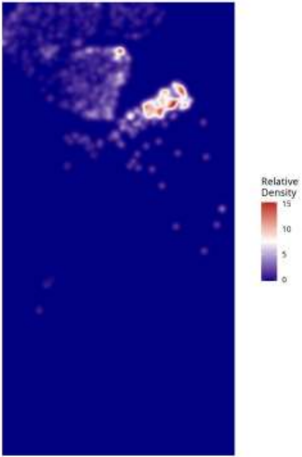

Relative Density - Patient: um09 | Target: PPR | Modeled: PPR

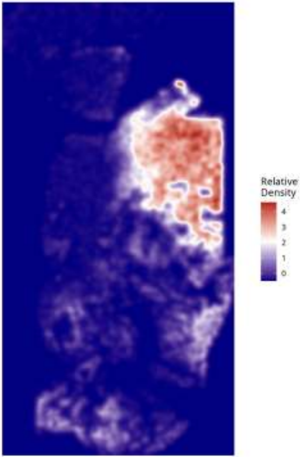

Relative Density - Patient: um09 | Target: PPR | Modeled: Tcells

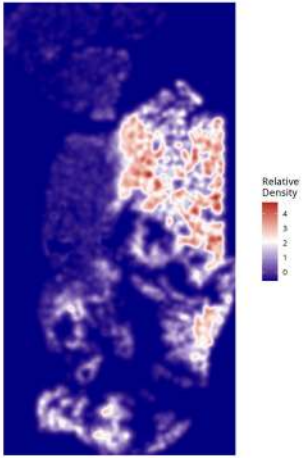

Relative Density - Patient: um09 | Target: PPR | Modeled: vasculature

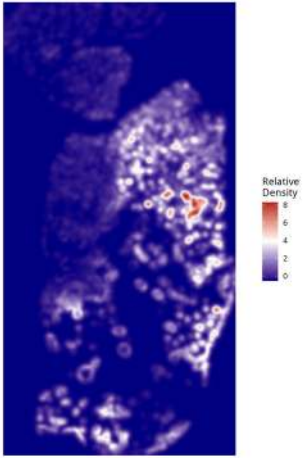

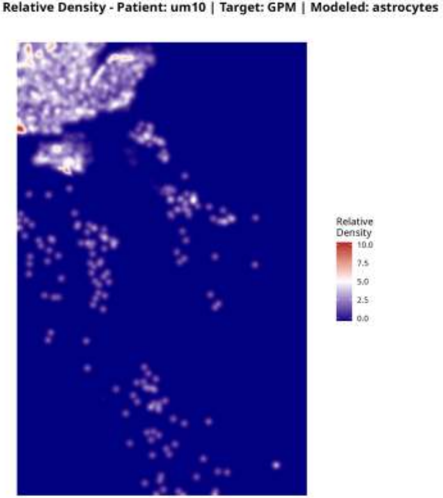
Relative Density - Patient: um10 | Target: GPM | Modeled: astrocytes
Relative Density
10.0
7.5
5.0
2.5
0.0

Relative Density - Patient: um10 | Target: GPM | Modeled: cDC

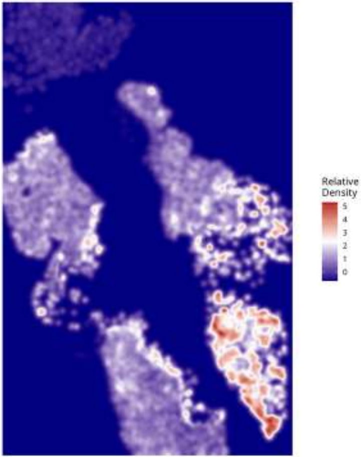

Relative Density - Patient: um10 | Target: GPM | Modeled: GPM

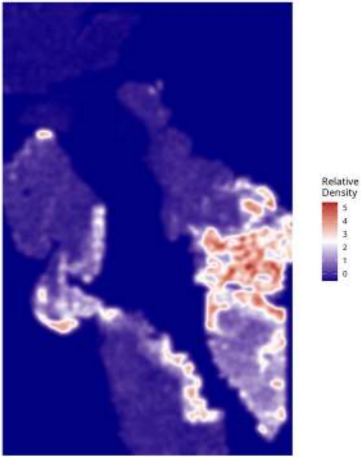

Relative Density - Patient: um10 | Target: GPM | Modeled: Macrophage

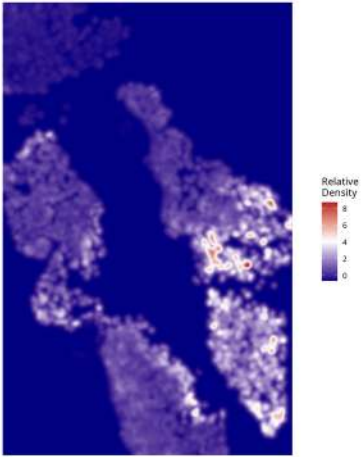

Relative Density - Patient: um10 | Target: GPM | Modeled: Microglia

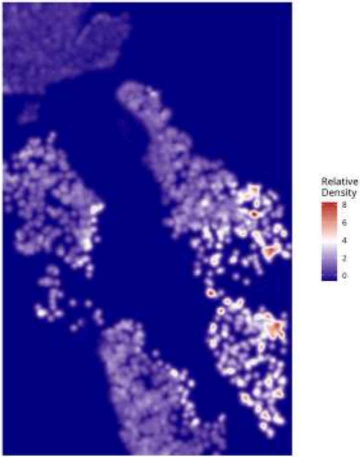

Relative Density - Patient: um10 | Target: GPM | Modeled: Monocyte

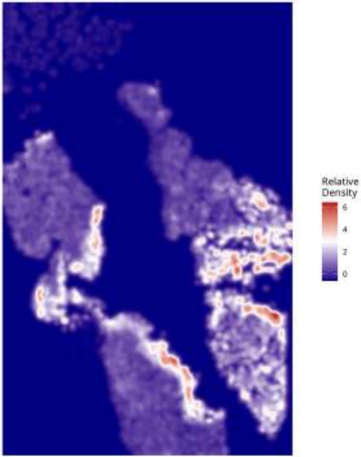

**Relative Density - Patient: um10 | Target: GPM | Modeled: MTC**

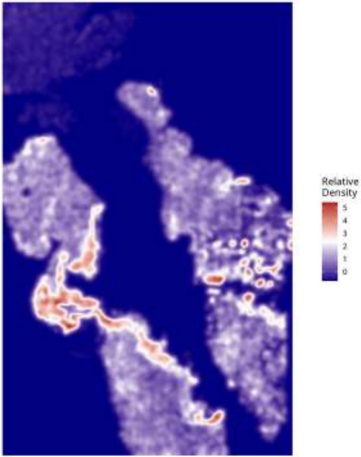

Relative Density - Patient: um10 | Target: GPM | Modeled: NEU

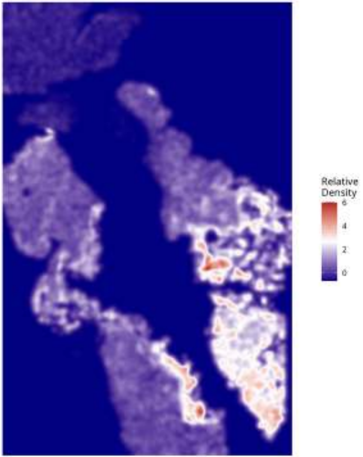

**Relative Density - Patient: um10 | Target: GPM | Modeled: neurons**

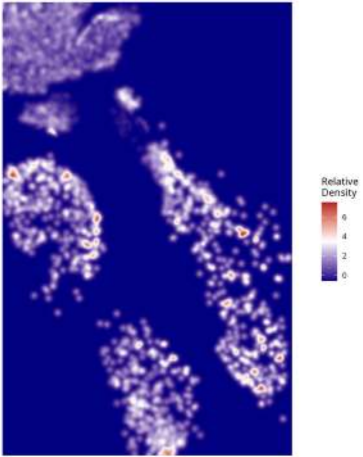

**Relative Density - Patient: um10 | Target: GPM | Modeled: Neutrophil**

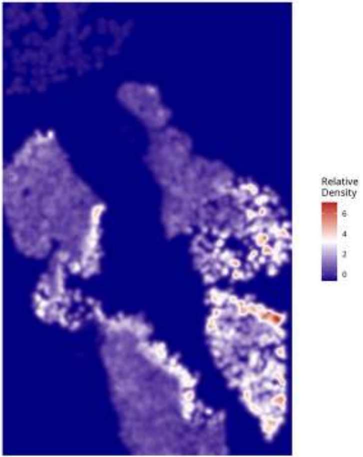

Relative Density - Patient: um10 | Target: GPM | Modeled: oligodendrocyte

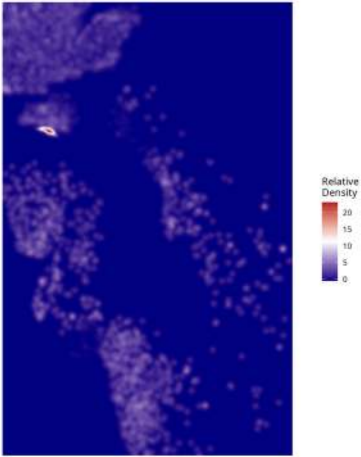

Relative Density - Patient: um10 | Target: GPM | Modeled: OPC

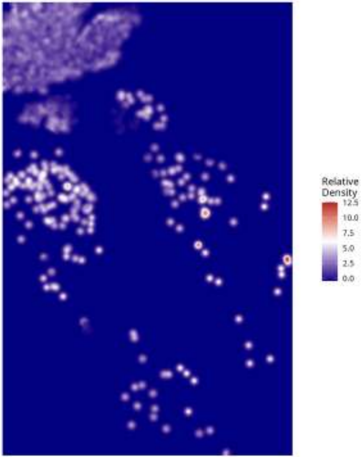

Relative Density - Patient: um10 | Target: GPM | Modeled: PPR

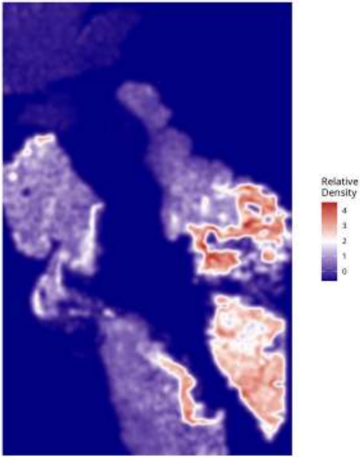

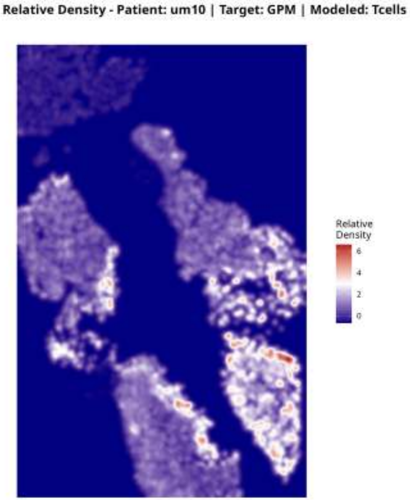
Relative Density - Patient: um10 | Target: GPM | Modeled: Tcells
Relative Density
6
4
2
0

Relative Density - Patient: um10 | Target: GPM | Modeled: vasculature

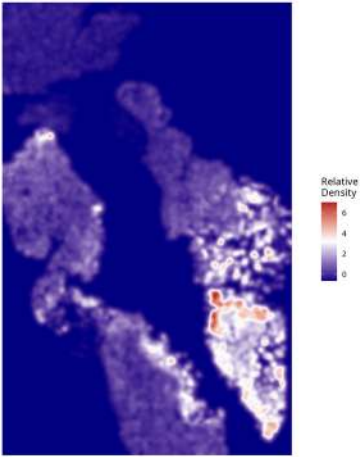

Relative Density - Patient: um10 | Target: MTC | Modeled: astrocytes

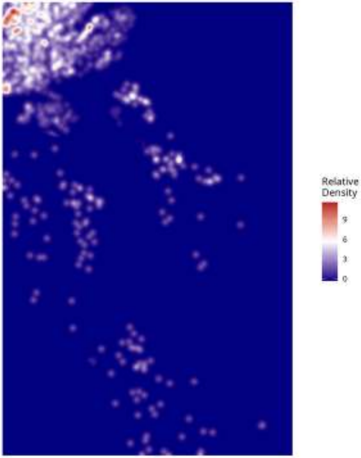

#### Relative Density - Patient: um10 | Target: MTC | Modeled: cDC

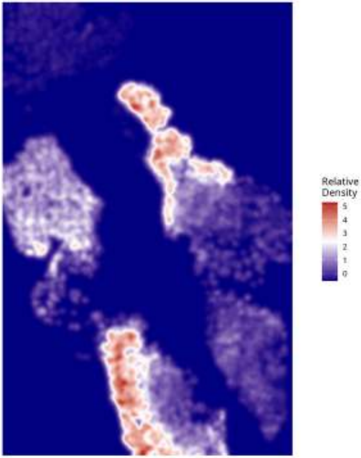

**Relative Density - Patient: um10 | Target: MTC | Modeled: GPM**

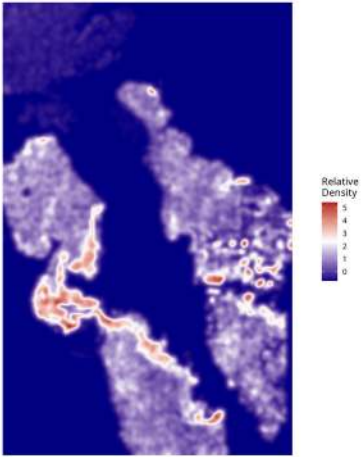

**Relative Density - Patient: um10 | Target: MTC | Modeled: Macrophage**

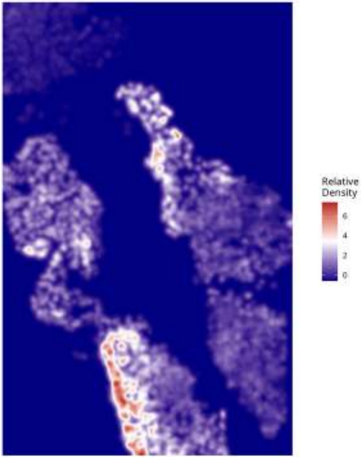

Relative Density - Patient: um10 | Target: MTC | Modeled: Microglia

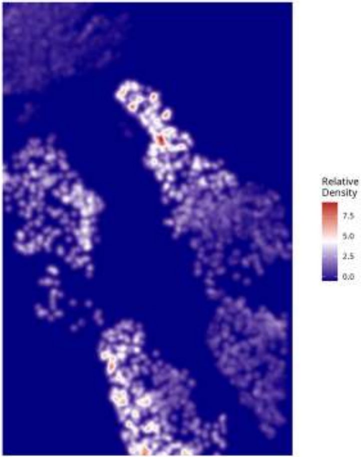

#### Relative Density - Patient: um10 | Target: MTC | Modeled: Monocyte

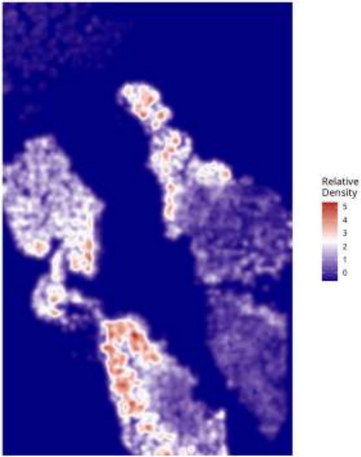

**Relative Density - Patient: um10 | Target: MTC | Modeled: MTC**

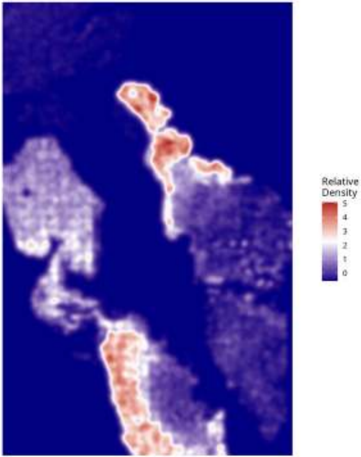

Relative Density - Patient: um10 | Target: MTC | Modeled: NEU

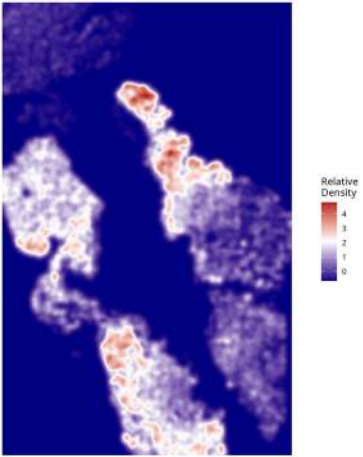

**Relative Density - Patient: um10 | Target: MTC | Modeled: neurons**

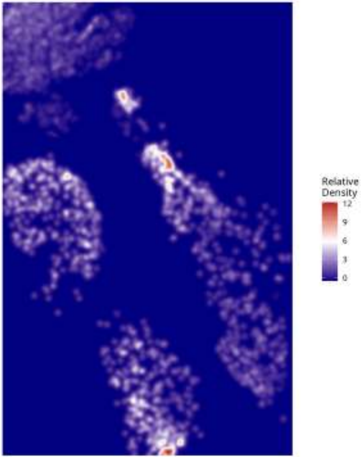

Relative Density - Patient: um10 | Target: MTC | Modeled: Neutrophil

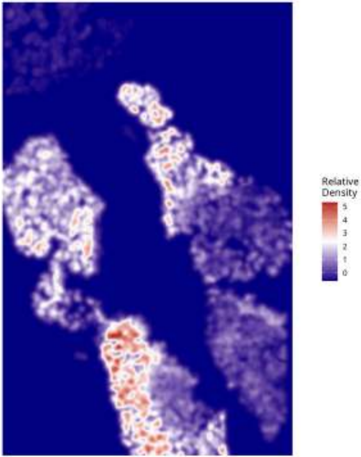

**Relative Density - Patient: um10 | Target: MTC | Modeled: oligodendrocyte**

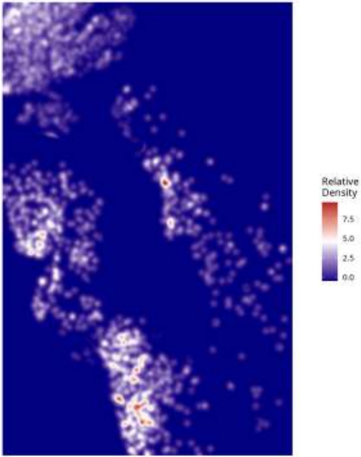

Relative Density - Patient: um10 | Target: MTC | Modeled: OPC

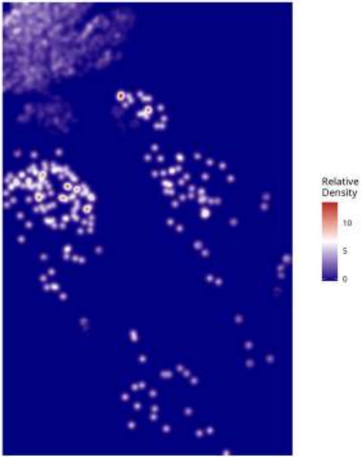

**Relative Density - Patient: um10 | Target: MTC | Modeled: PPR**

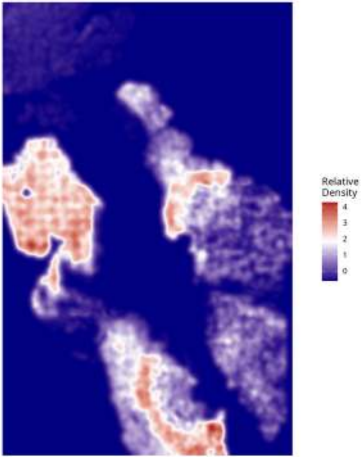

**Relative Density - Patient: um10 | Target: MTC | Modeled: Tcells**

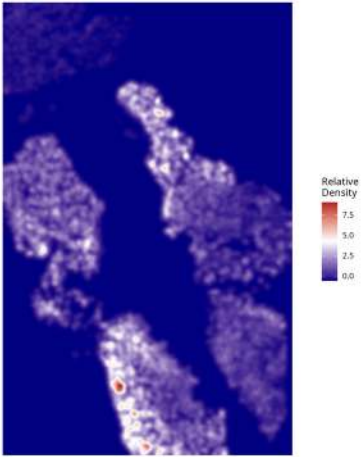

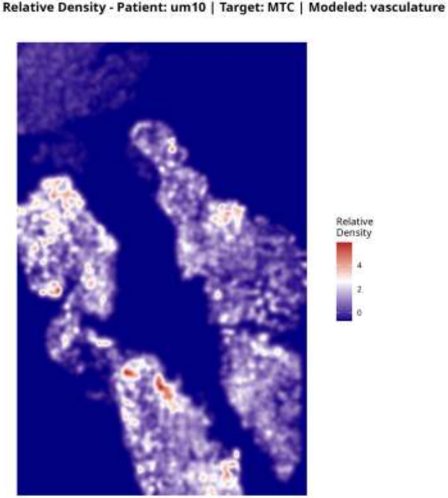
Relative Density - Patient: um10 | Target: MTC | Modeled: vasculature
Relative
Density
4
2
0

Relative Density - Patient: um10 | Target: NEU | Modeled: astrocytes

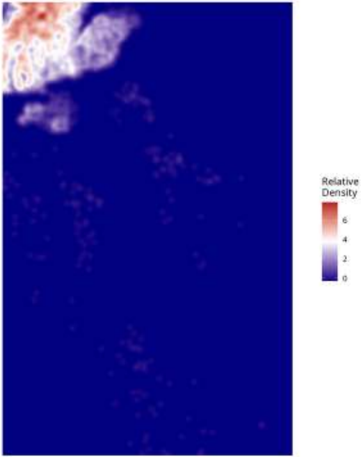

Relative Density - Patient: um10 | Target: NEU | Modeled: cDC

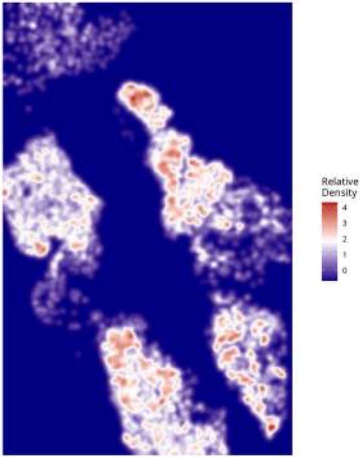

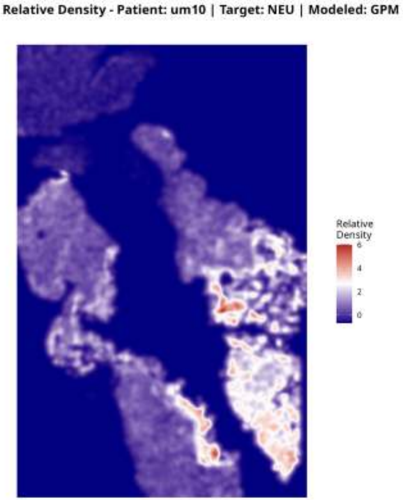
Relative Density - Patient: um10 | Target: NEU | Modeled: GPM
Relative Density
6
4
2
0

Relative Density - Patient: um10 | Target: NEU | Modeled: Macrophage

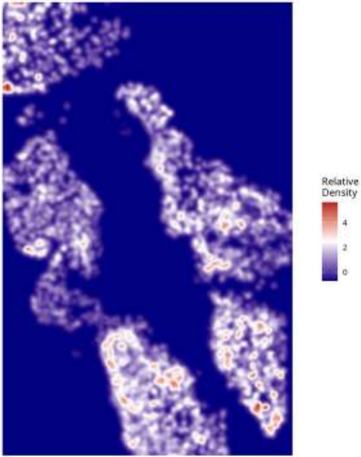

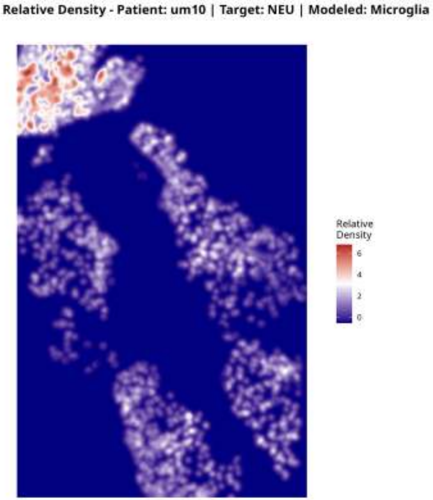
Relative Density - Patient: um10 | Target: NEU | Modeled: Microglia
Relative Density
6
4
2
0

Relative Density - Patient: um10 | Target: NEU | Modeled: Monocyte

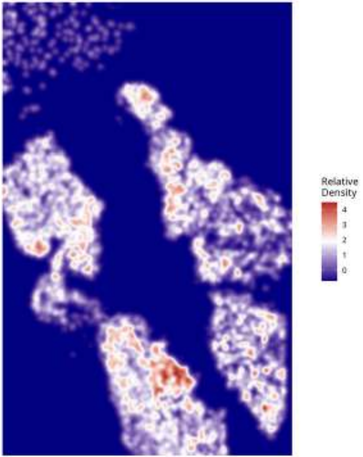

**Relative Density - Patient: um10 | Target: NEU | Modeled: MTC**

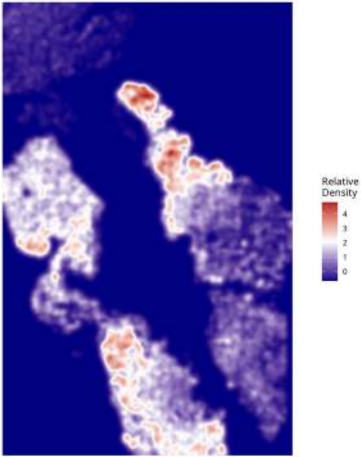

Relative Density - Patient: um10 | Target: NEU | Modeled: NEU

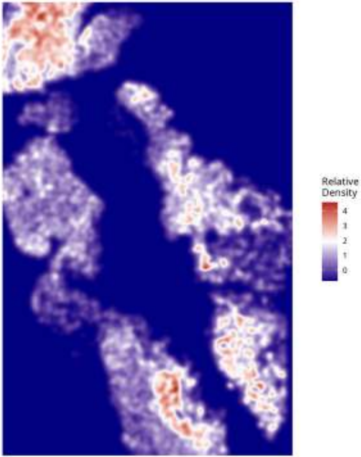

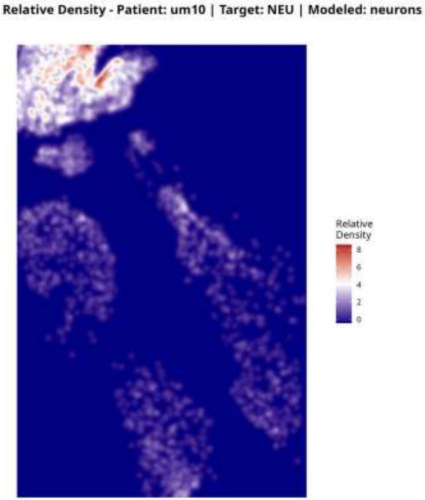
Relative Density - Patient: um10 | Target: NEU | Modeled: neurons
Relative Density
8
6
4
2
0

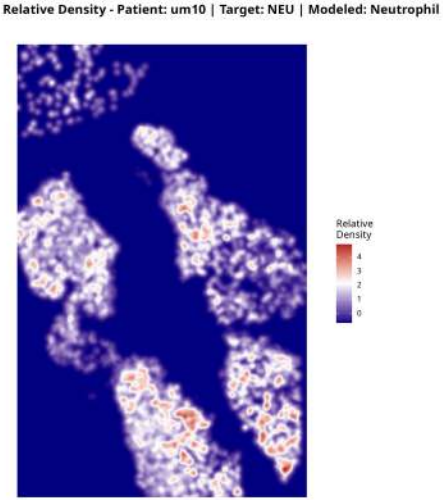
Relative Density - Patient: um10 | Target: NEU | Modeled: Neutrophil
Relative
Density
4
3
2
1
0

Relative Density - Patient: um10 | Target: NEU | Modeled: oligodendrocyte

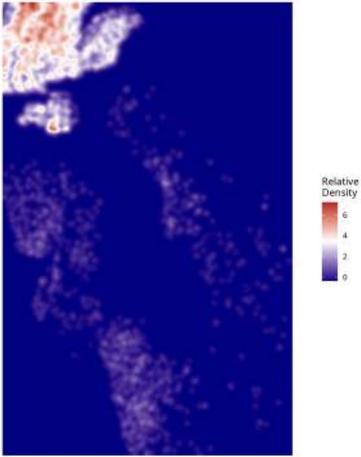

Relative Density - Patient: um10 | Target: NEU | Modeled: OPC

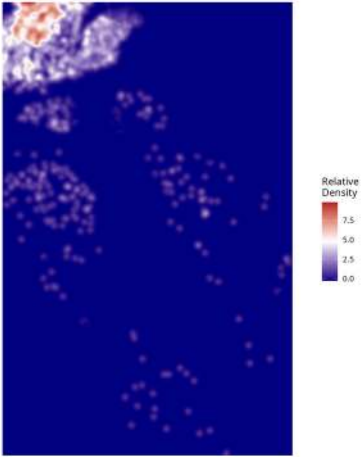

**Relative Density - Patient: um10 | Target: NEU | Modeled: PPR**

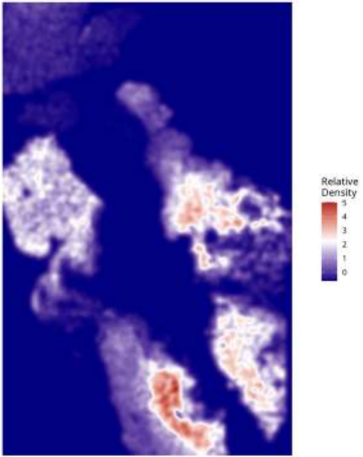

Relative Density - Patient: um10 | Target: NEU | Modeled: Tcells

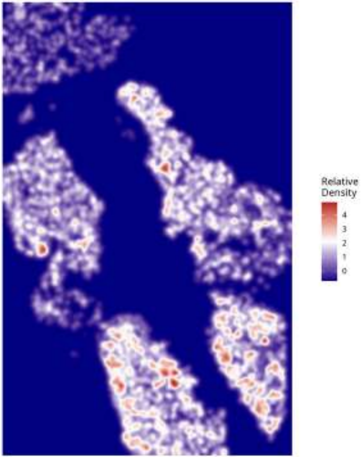

#### Relative Density - Patient: um10 | Target: NEU | Modeled: vasculature

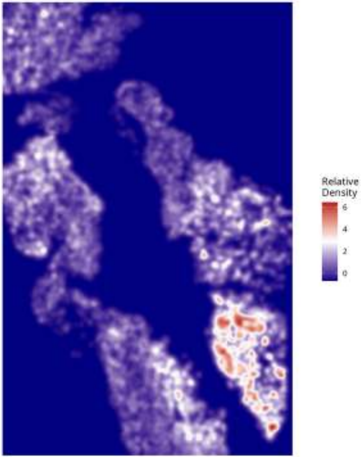

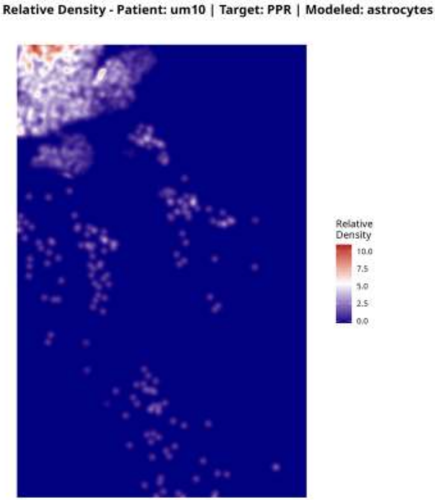

Relative Density - Patient: um10 | Target: PPR | Modeled: astrocytes
Relative Density
10.0
7.5
5.0
2.5
0.0

**Relative Density - Patient: um10 | Target: PPR | Modeled: cDC**

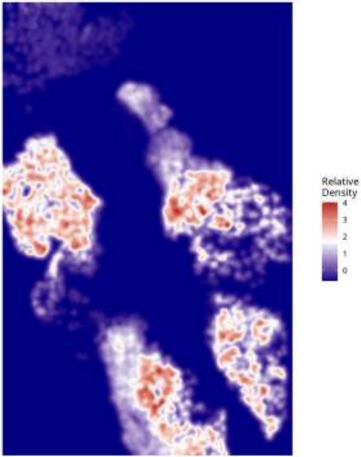

Relative Density - Patient: um10 | Target: PPR | Modeled: GPM

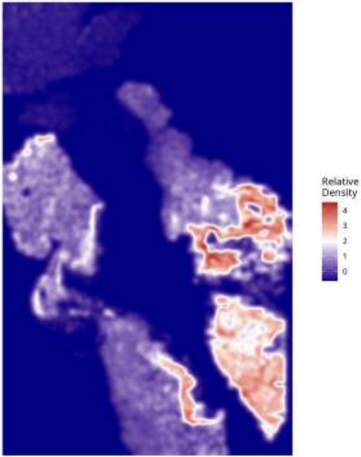

Relative Density - Patient: um10 | Target: PPR | Modeled: Macrophage

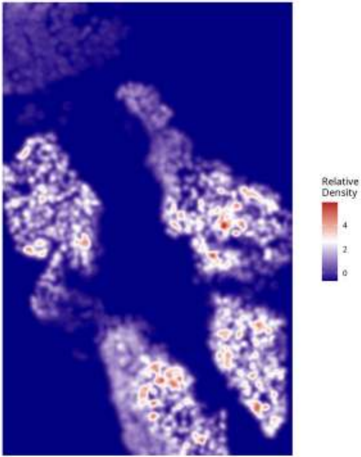

#### Relative Density - Patient: um10 | Target: PPR | Modeled: Microglia

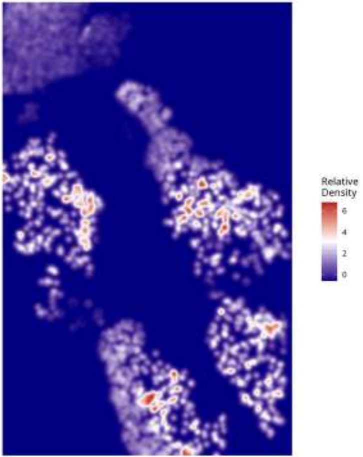

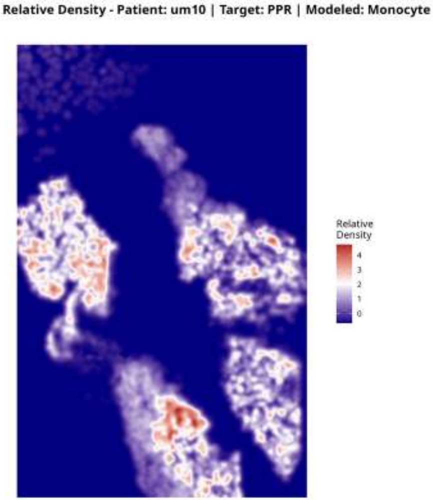
Relative Density - Patient: um10 | Target: PPR | Modeled: Monocyte
Relative Density
4
3
2
1
0

Relative Density - Patient: um10 | Target: PPR | Modeled: MTC

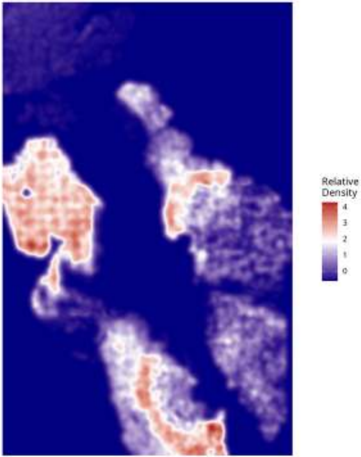

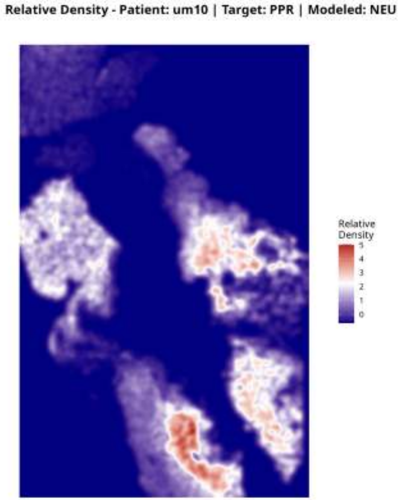
Relative Density - Patient: um10 | Target: PPR | Modeled: NEU
Relative Density
5
4
3
2
1
0

**Relative Density - Patient: um10 | Target: PPR | Modeled: neurons**

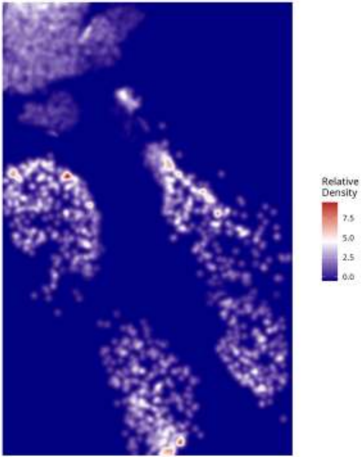

**Relative Density - Patient: um10 | Target: PPR | Modeled: Neutrophil**

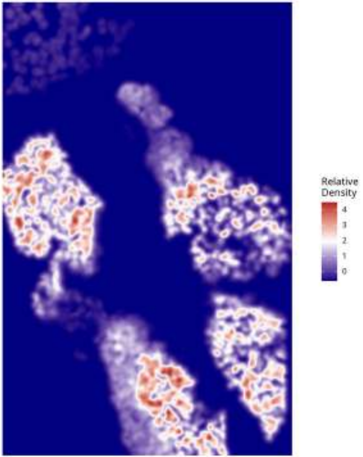

#### Relative Density - Patient: um10 | Target: PPR | Modeled: oligodendrocyte

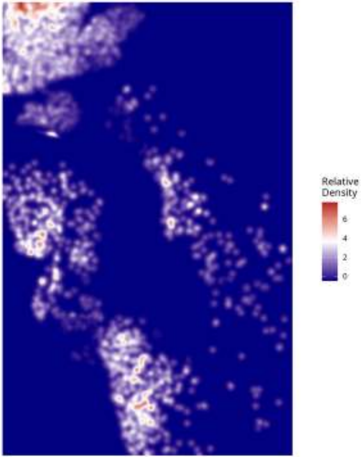

Relative Density - Patient: um10 | Target: PPR | Modeled: OPC

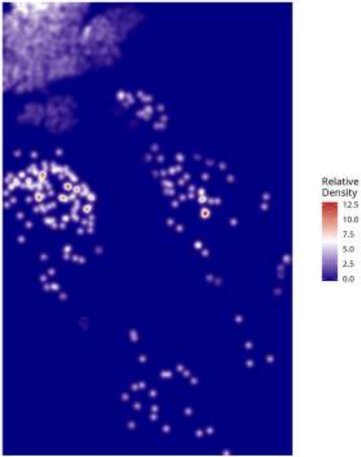

Relative Density - Patient: um10 | Target: PPR | Modeled: PPR

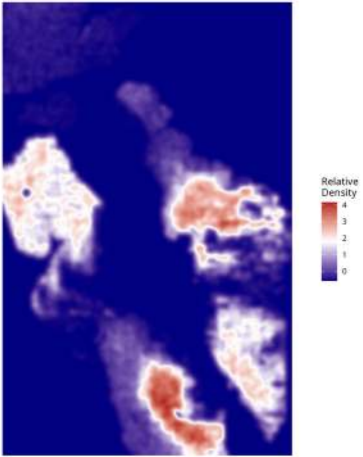

Relative Density - Patient: um10 | Target: PPR | Modeled: Tcells

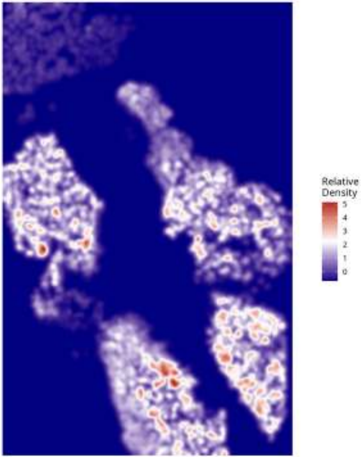

Relative Density - Patient: um10 | Target: PPR | Modeled: vasculature

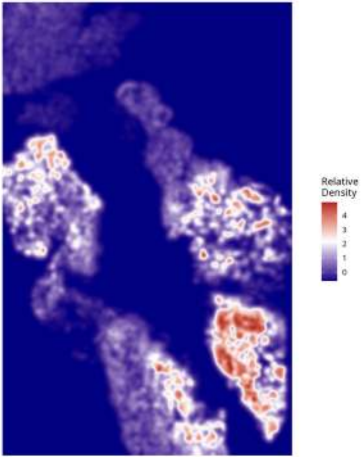

Relative Density - Patient: um11 | Target: GPM | Modeled: astrocytes

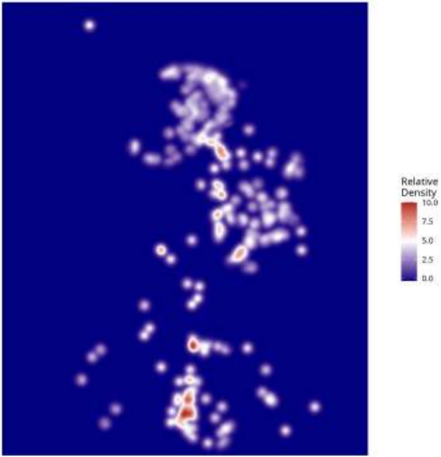

**Relative Density - Patient: um11 | Target: GPM | Modeled: cDC**

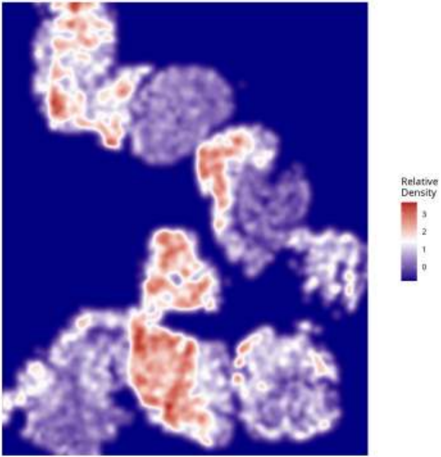

**Relative Density - Patient: um11 | Target: GPM | Modeled: GPM**

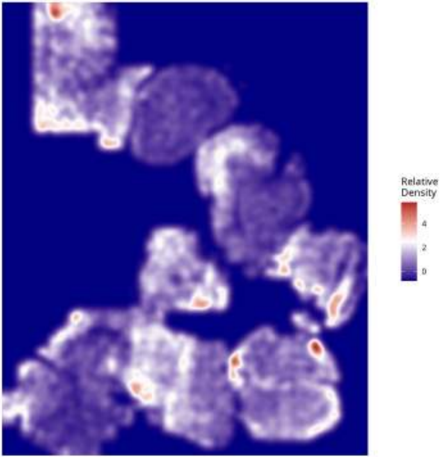

Relative Density - Patient: um11 | Target: GPM | Modeled: Macrophage

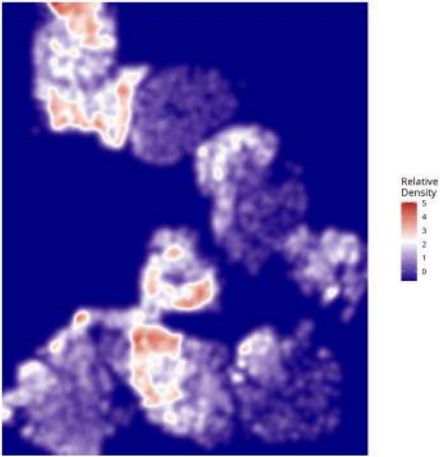

Relative Density - Patient: um11 | Target: GPM | Modeled: Microglia

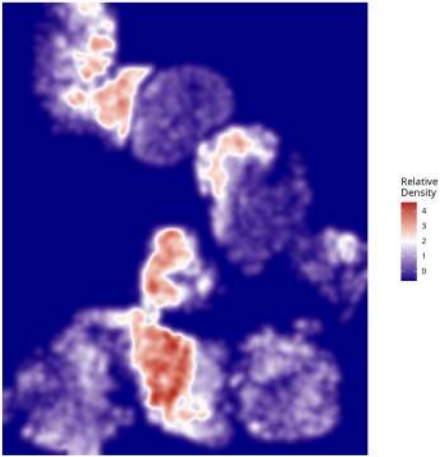

Relative Density - Patient: um11 | Target: GPM | Modeled: Monocyte

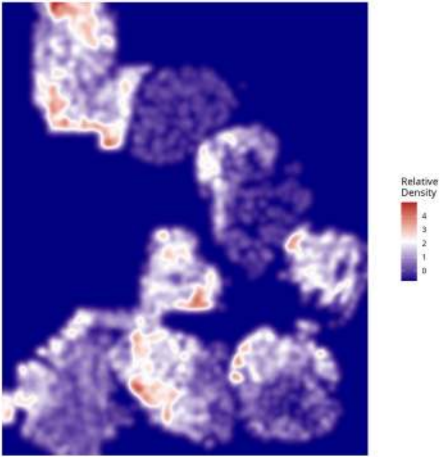

Relative Density - Patient: um11 | Target: GPM | Modeled: MTC

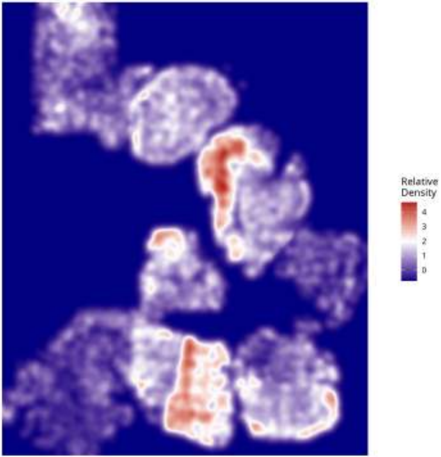

Relative Density - Patient: um11 | Target: GPM | Modeled: NEU

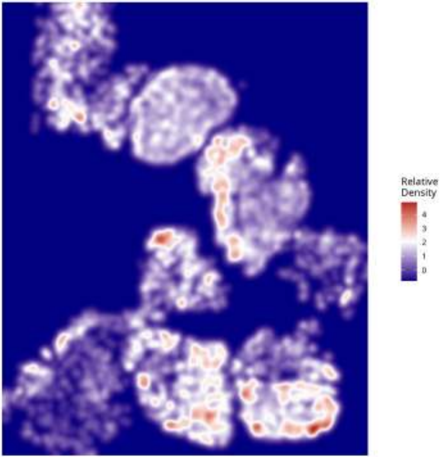

Relative Density - Patient: um11 | Target: GPM | Modeled: neurons

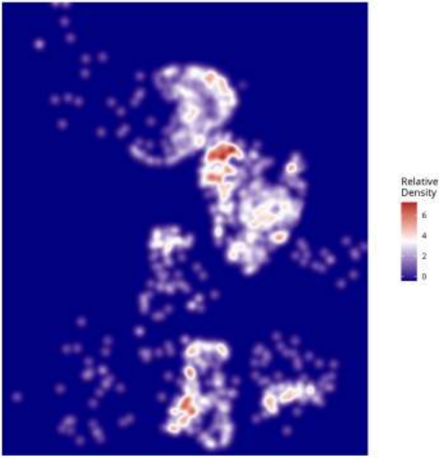

Relative Density - Patient: um11 | Target: GPM | Modeled: Neutrophil

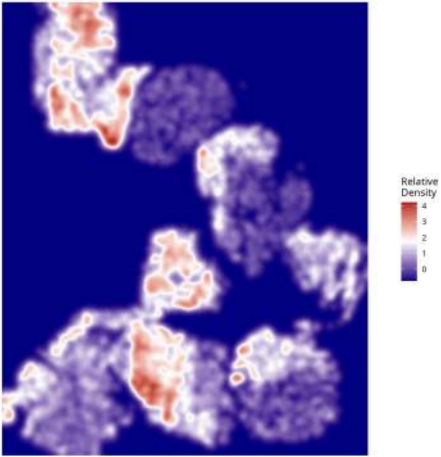

**Relative Density - Patient: um11 | Target: GPM | Modeled: oligodendrocyte**

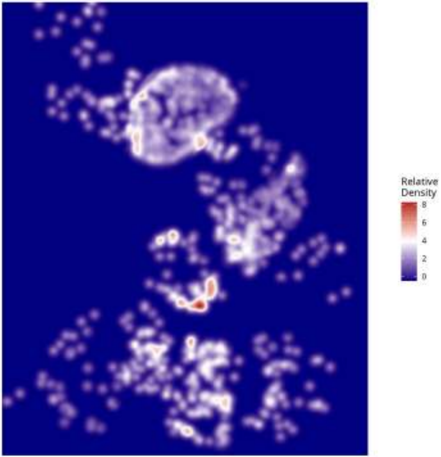

Relative Density - Patient: um11 | Target: GPM | Modeled: OPC

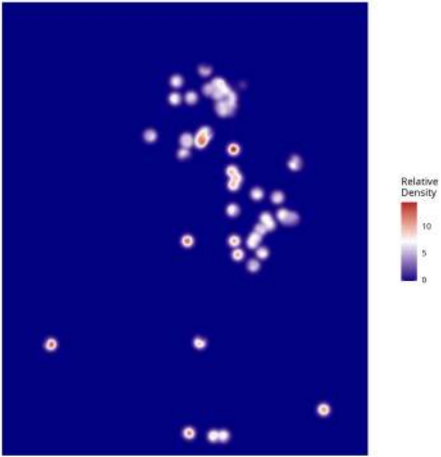

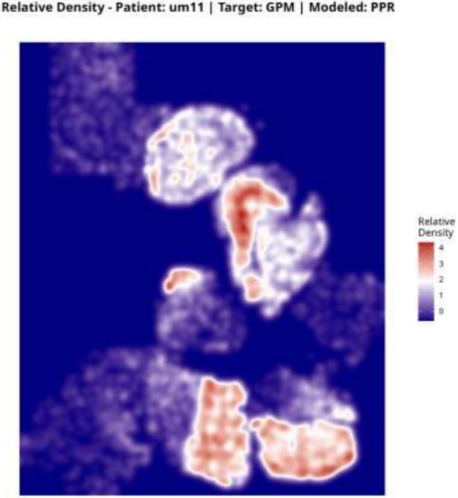

Relative Density - Patient: um11 | Target: GPM | Modeled: PPR
Relative Density
4
3
2
1
0

Relative Density - Patient: um11 | Target: GPM | Modeled: Tcells

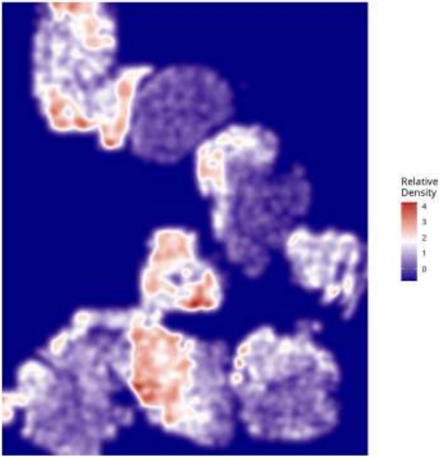

Relative Density - Patient: um11 | Target: GPM | Modeled: vasculature

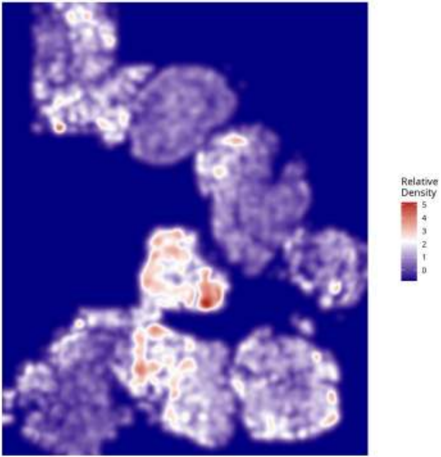

**Relative Density - Patient: um11 | Target: MTC | Modeled: astrocytes**

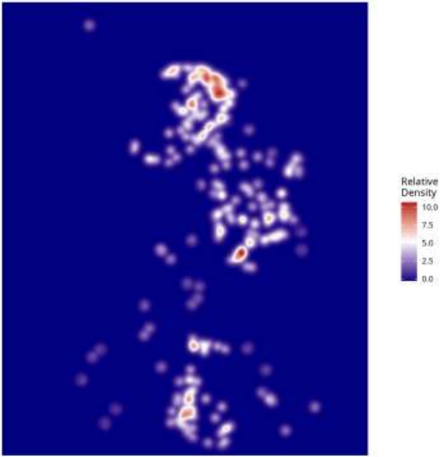

**Relative Density - Patient: um11 | Target: MTC | Modeled: cDC**

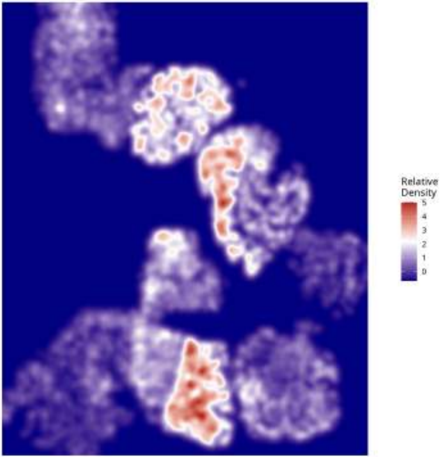

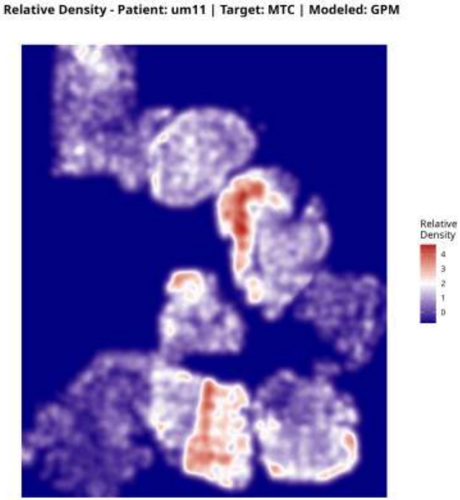
Relative Density - Patient: um11 | Target: MTC | Modeled: GPM
Relative
Density
4
3
2
1
0

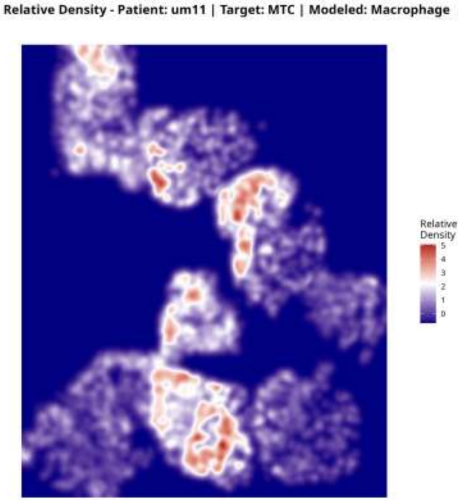
Relative Density - Patient: um11 | Target: MTC | Modeled: Macrophage
Relative Density
5
4
3
2
1
0

**Relative Density - Patient: um11 | Target: MTC | Modeled: Microglia**

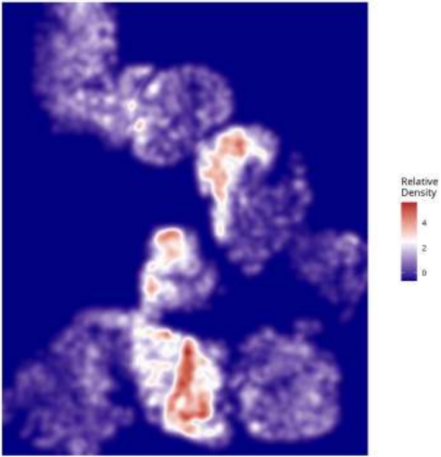

Relative Density - Patient: um11 | Target: MTC | Modeled: Monocyte

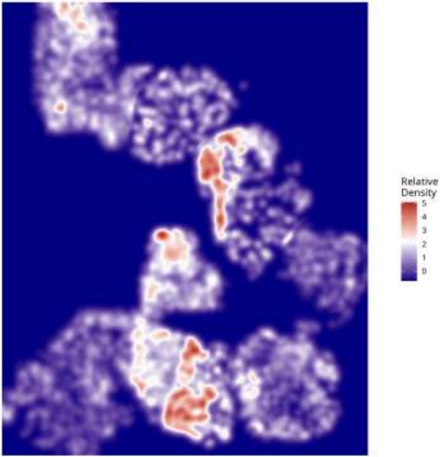

Relative Density - Patient: um11 | Target: MTC | Modeled: MTC

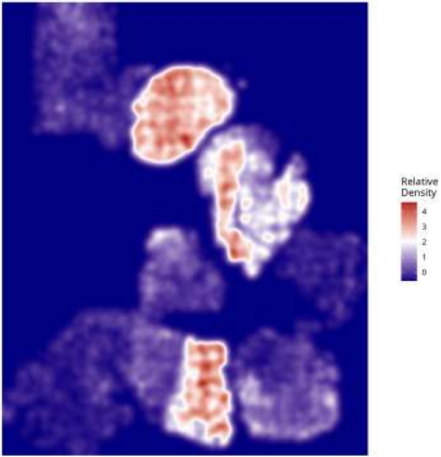

**Relative Density - Patient: um11 | Target: MTC | Modeled: NEU**

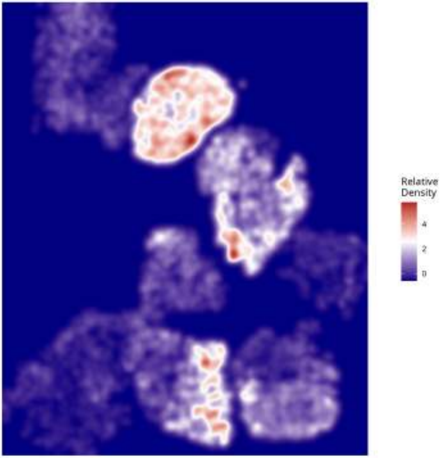

Relative Density - Patient: um11 | Target: MTC | Modeled: neurons

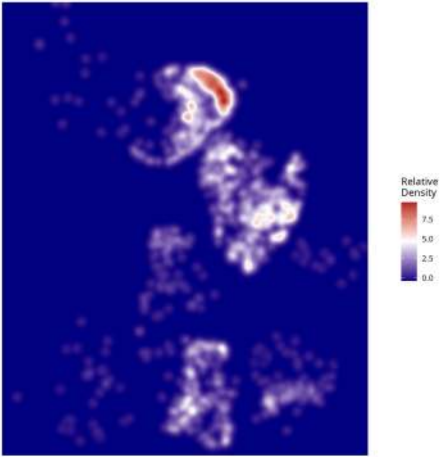

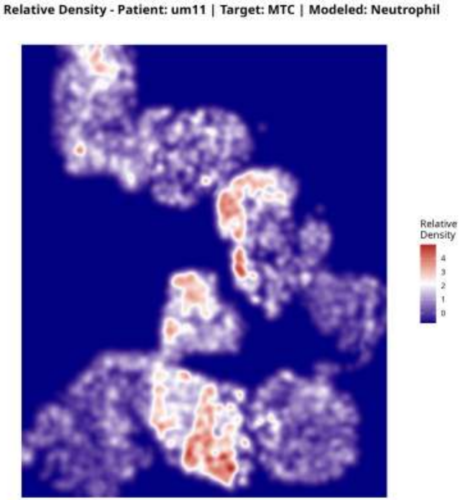
Relative Density - Patient: um11 | Target: MTC | Modeled: Neutrophil
Relative
Density
4
3
2
1
0

**Relative Density - Patient: um11 | Target: MTC | Modeled: oligodendrocyte**

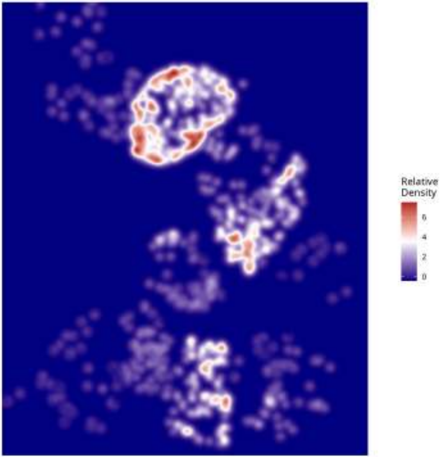

Relative Density - Patient: um11 | Target: MTC | Modeled: OPC

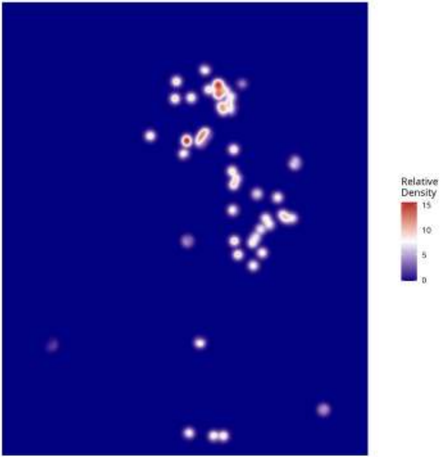

Relative Density - Patient: um11 | Target: MTC | Modeled: PPR

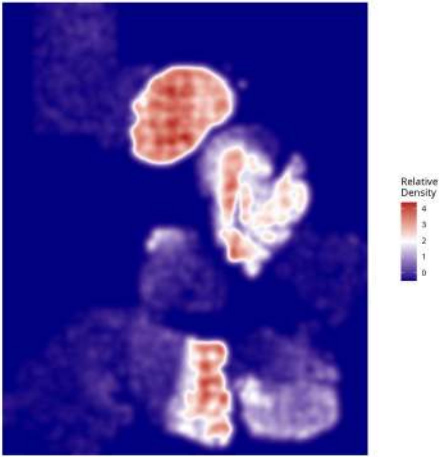

**Relative Density - Patient: um11 | Target: MTC | Modeled: Tcells**

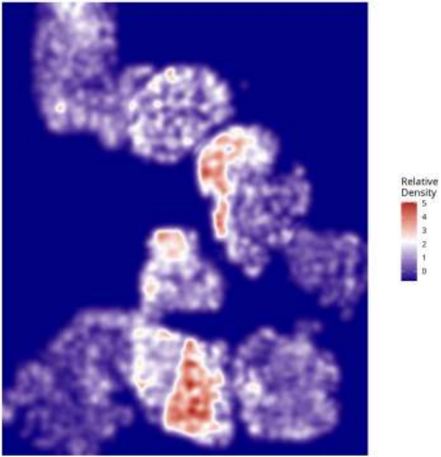

Relative Density - Patient: um11 | Target: MTC | Modeled: vasculature

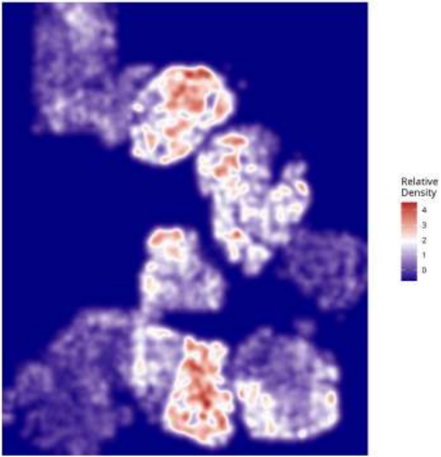

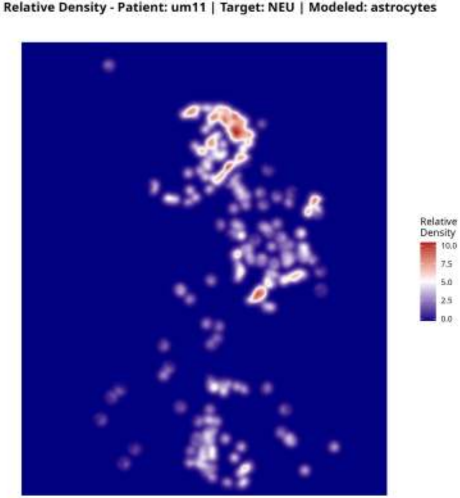
Relative Density - Patient: um11 | Target: NEU | Modeled: astrocytes
Relative Density
10.0
7.5
5.0
2.5
0.0

Relative Density - Patient: um11 | Target: NEU | Modeled: cDC

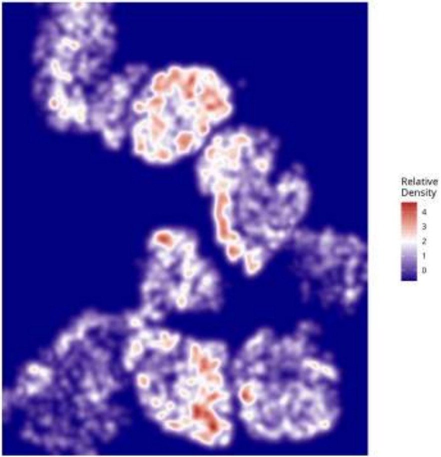

**Relative Density - Patient: um11 | Target: NEU | Modeled: GPM**

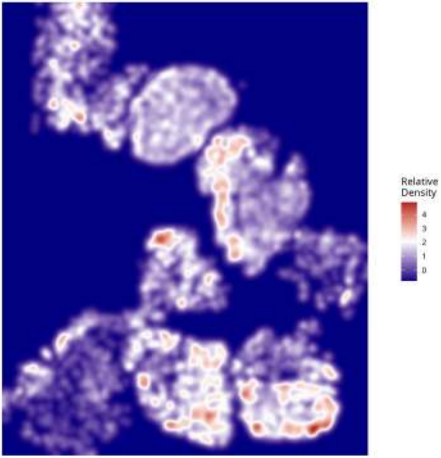

**Relative Density - Patient: um11 | Target: NEU | Modeled: Macrophage**

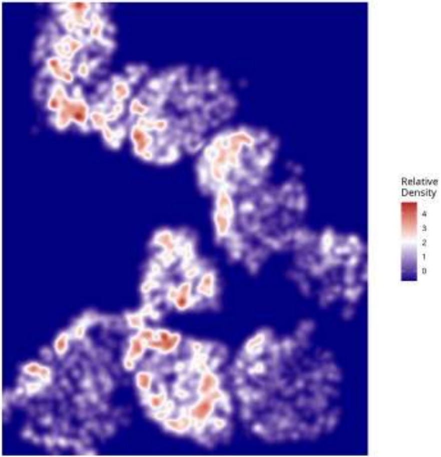

Relative Density - Patient: um11 | Target: NEU | Modeled: Microglia

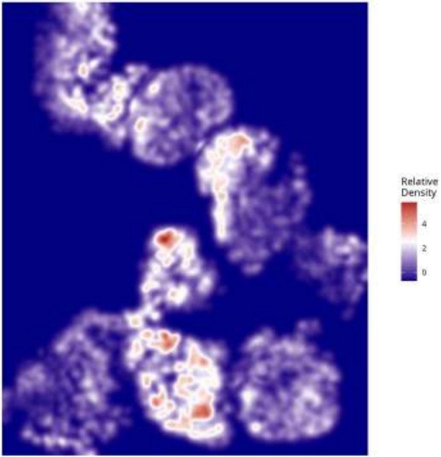

Relative Density - Patient: um11 | Target: NEU | Modeled: Monocyte

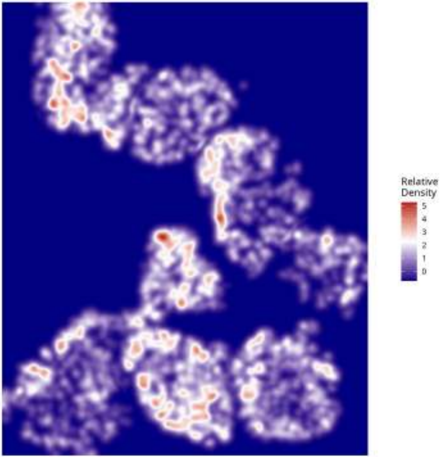

Relative Density - Patient: um11 | Target: NEU | Modeled: MTC

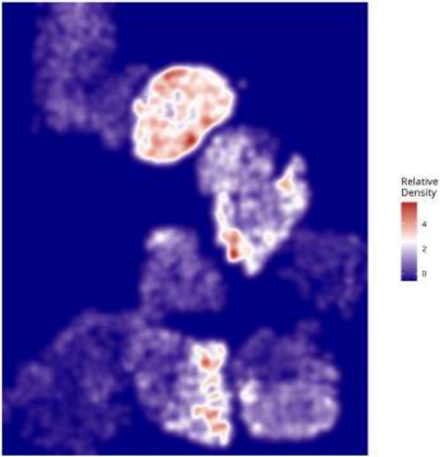

**Relative Density - Patient: um11 | Target: NEU | Modeled: NEU**

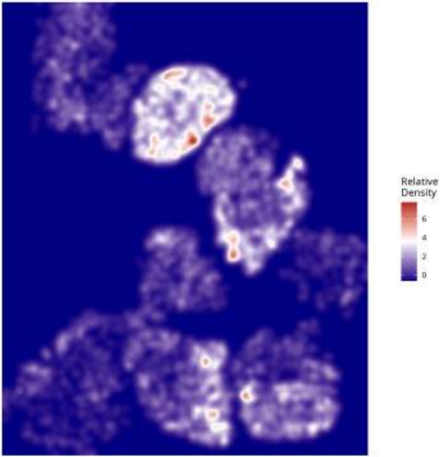

Relative Density - Patient: um11 | Target: NEU | Modeled: neurons

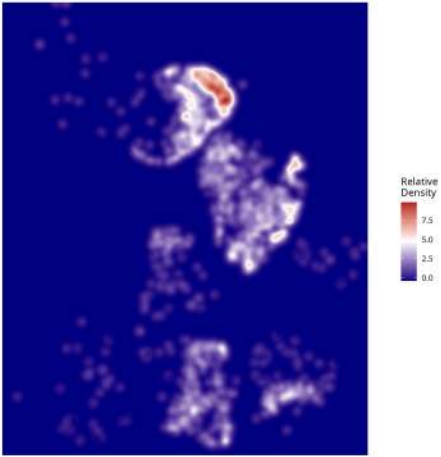

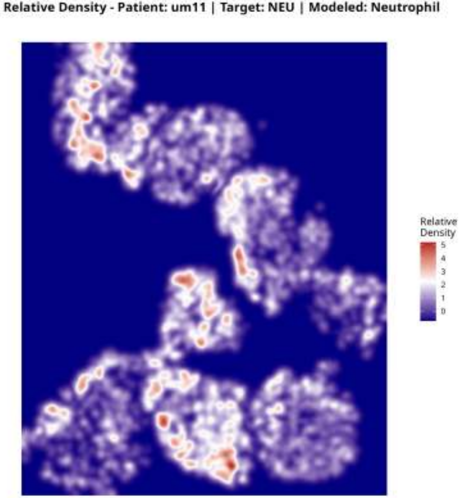
Relative Density - Patient: um11 | Target: NEU | Modeled: Neutrophil
Relative Density
5
4
3
2
1
0

Relative Density - Patient: um11 | Target: NEU | Modeled: oligodendrocyte

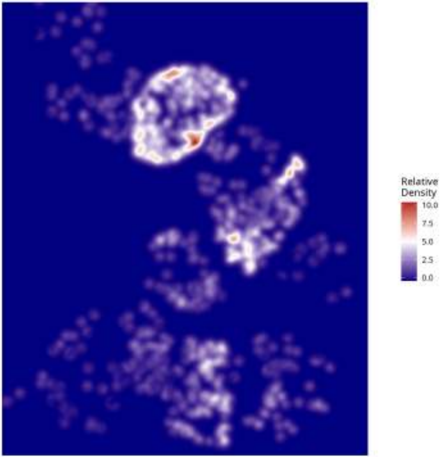

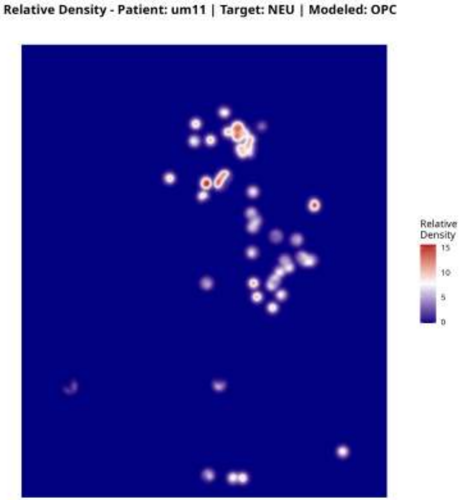
Relative Density - Patient: um11 | Target: NEU | Modeled: OPC
Relative Density
15
10
5
0

Relative Density - Patient: um11 | Target: NEU | Modeled: PPR

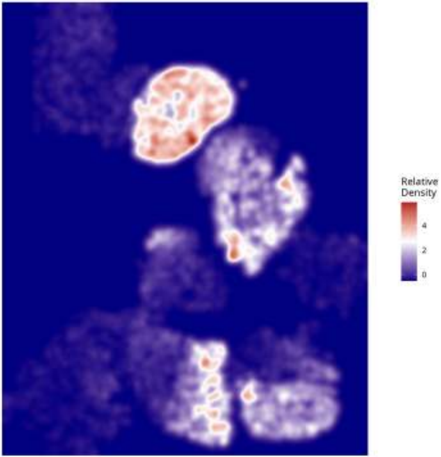

Relative Density - Patient: um11 | Target: NEU | Modeled: Tcells

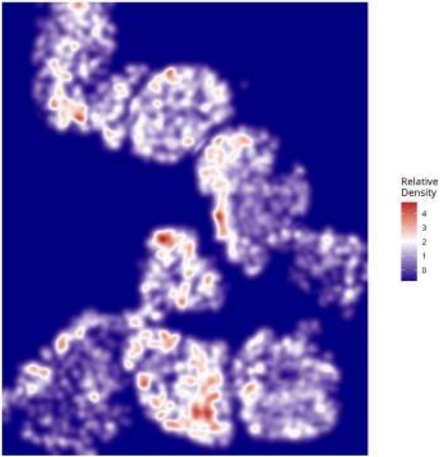

Relative Density - Patient: um11 | Target: NEU | Modeled: vasculature

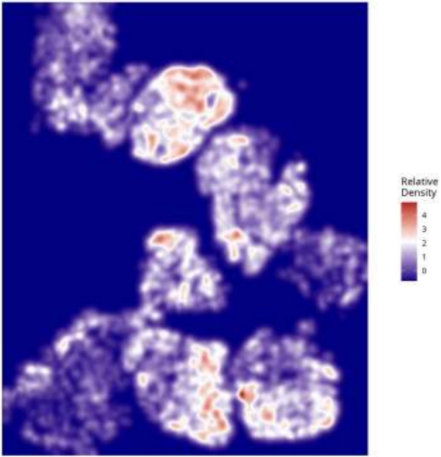

**Relative Density - Patient: um11 | Target: PPR | Modeled: astrocytes**

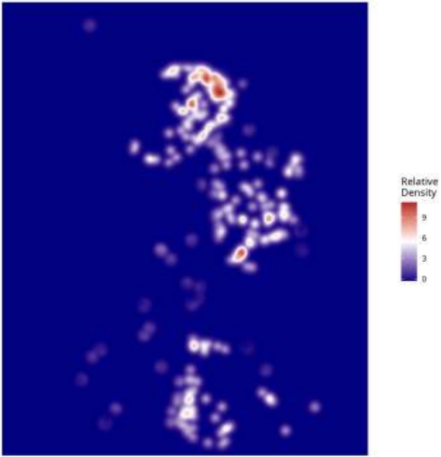

**Relative Density - Patient: um11 | Target: PPR | Modeled: cDC**

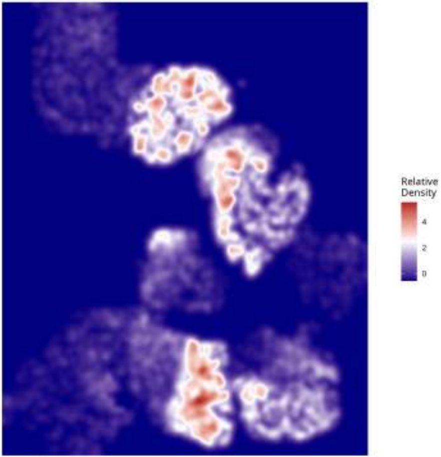

**Relative Density - Patient: um11 | Target: PPR | Modeled: GPM**

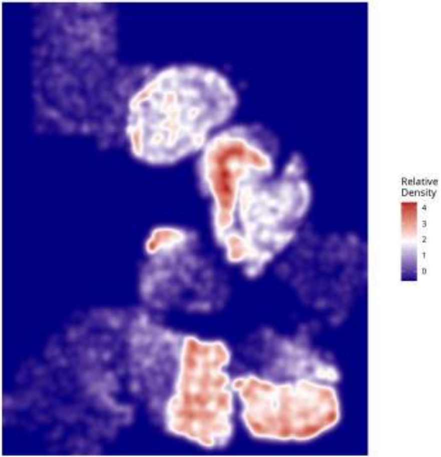

**Relative Density - Patient: um11 | Target: PPR | Modeled: Macrophage**

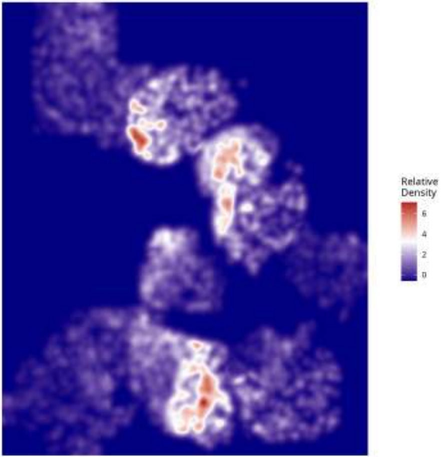

**Relative Density - Patient: um11 | Target: PPR | Modeled: Microglia**

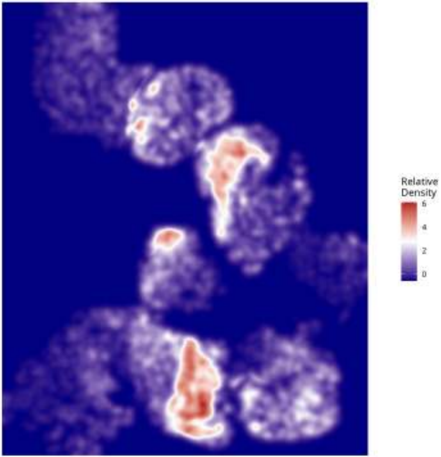

**Relative Density - Patient: um11 | Target: PPR | Modeled: Monocyte**

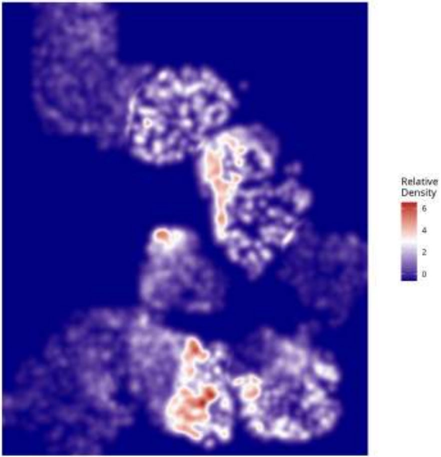

**Relative Density - Patient: um11 | Target: PPR | Modeled: MTC**

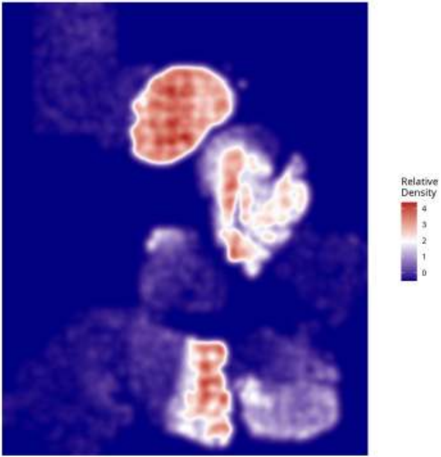

Relative Density - Patient: um11 | Target: PPR | Modeled: NEU

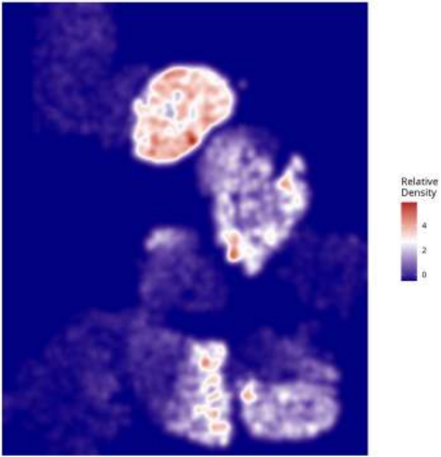

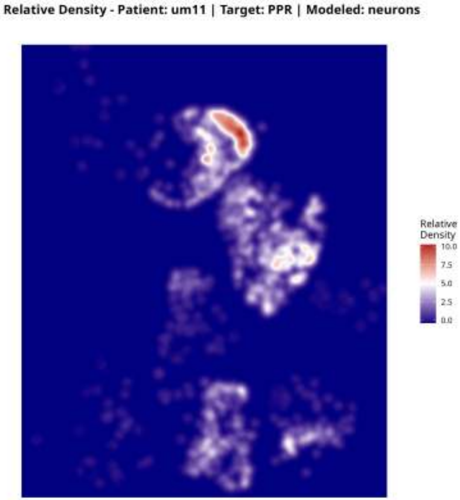

Relative Density - Patient: um11 | Target: PPR | Modeled: neurons
Relative Density
10.0
7.5
5.0
2.5
0.0

Relative Density - Patient: um11 | Target: PPR | Modeled: Neutrophil

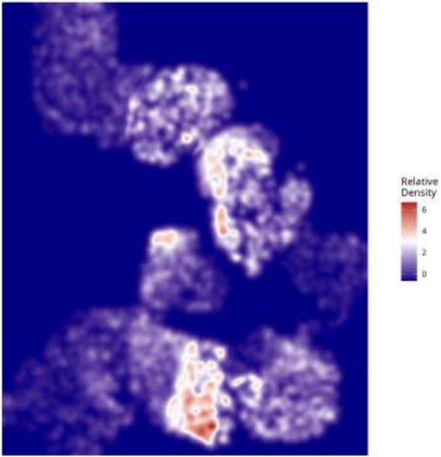

Relative Density - Patient: um11 | Target: PPR | Modeled: oligodendrocyte

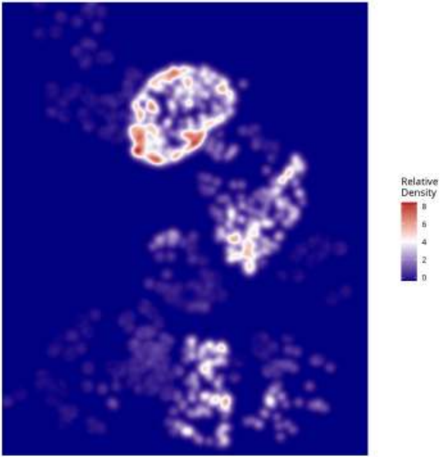

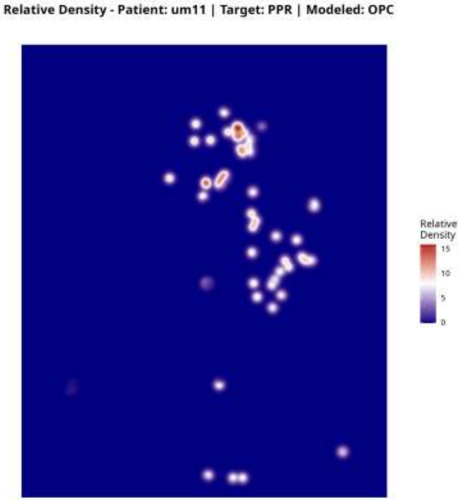

Relative Density - Patient: um11 | Target: PPR | Modeled: OPC
Relative Density
15
10
5
0

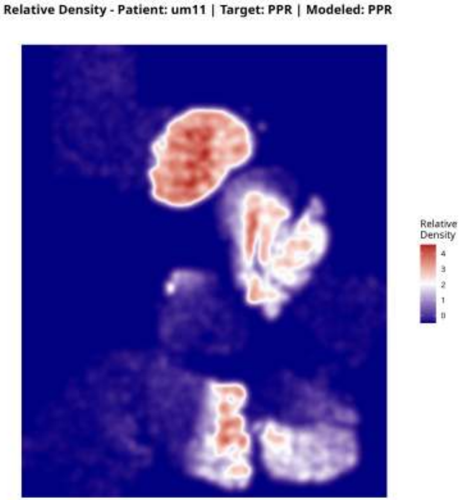
Relative Density - Patient: um11 | Target: PPR | Modeled: PPR
Relative Density
4
3
2
1
0

Relative Density - Patient: um11 | Target: PPR | Modeled: Tcells

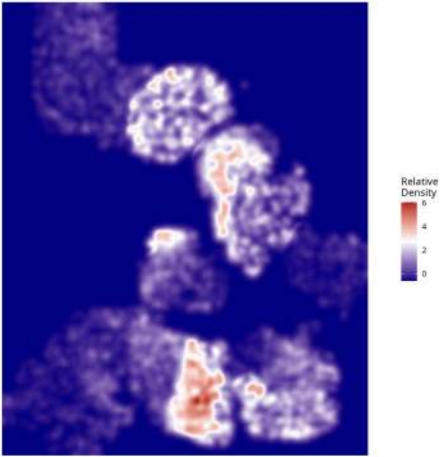

**Relative Density - Patient: um11 | Target: PPR | Modeled: vasculature**

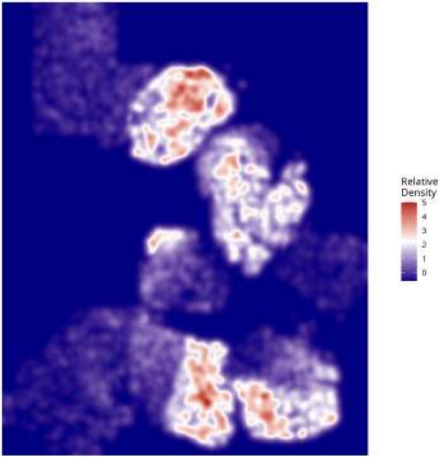

**Relative Density - Patient: um12 | Target: GPM | Modeled: astrocytes**

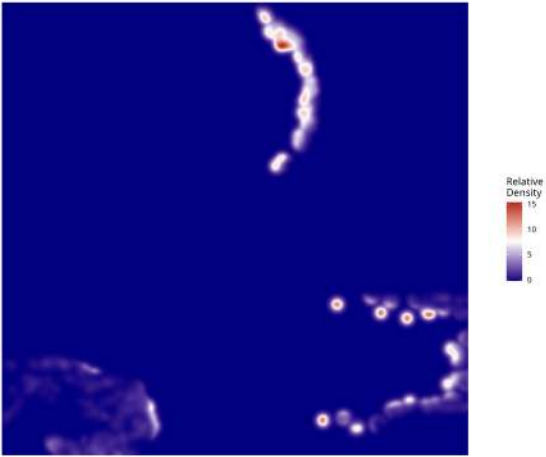

Relative Density - Patient: um12 | Target: GPM | Modeled: cDC

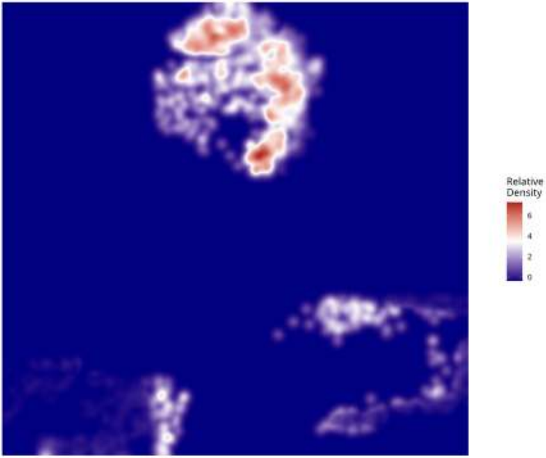

**Relative Density - Patient: um12 | Target: GPM | Modeled: GPM**

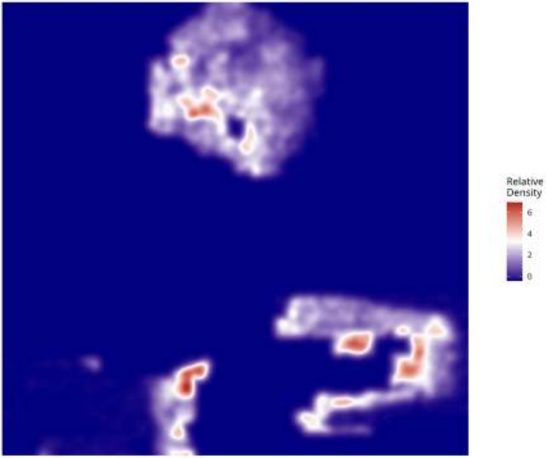

Relative Density - Patient: um12 | Target: GPM | Modeled: Macrophage

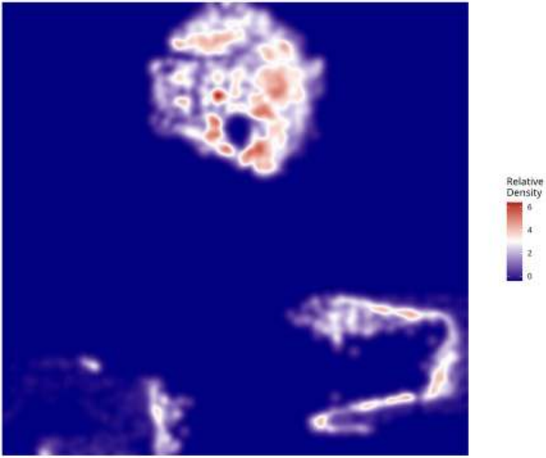

**Relative Density - Patient: um12 | Target: GPM | Modeled: Microglia**

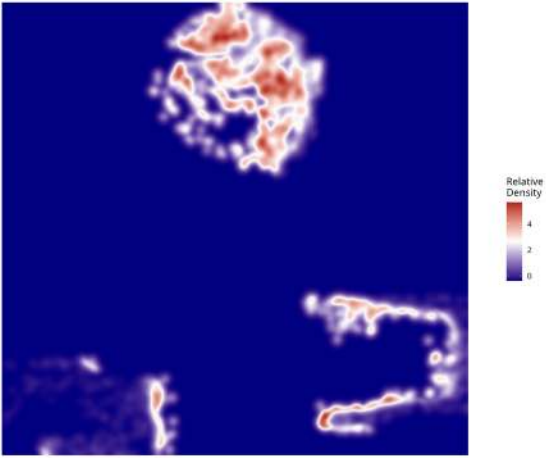

**Relative Density - Patient: um12 | Target: GPM | Modeled: Monocyte**

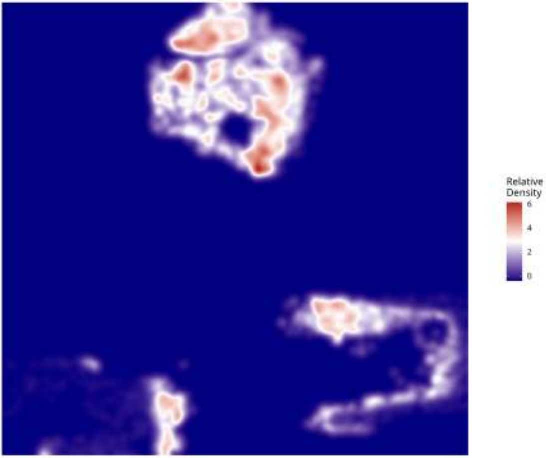

Relative Density - Patient: um12 | Target: GPM | Modeled: MTC

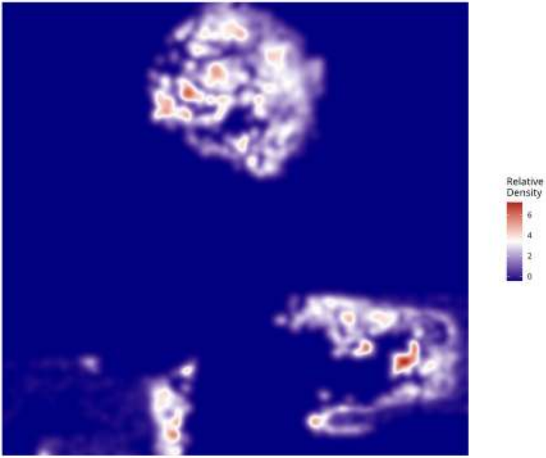

**Relative Density - Patient: um12 | Target: GPM | Modeled: NEU**

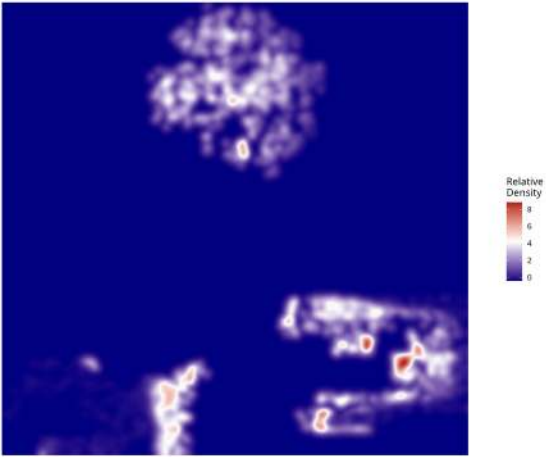

**Relative Density - Patient: um12 | Target: GPM | Modeled: Neutrophil**

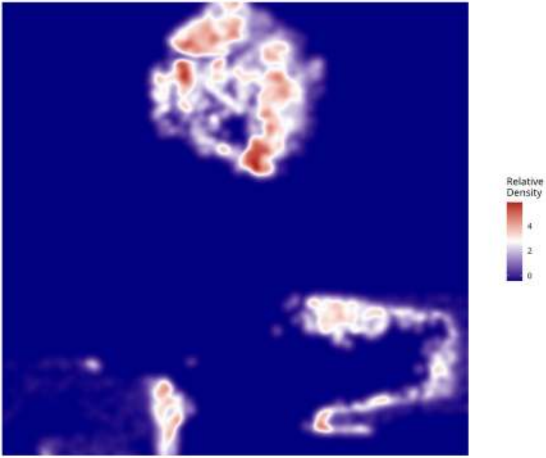

**Relative Density - Patient: um12 | Target: GPM | Modeled: oligodendrocyte**

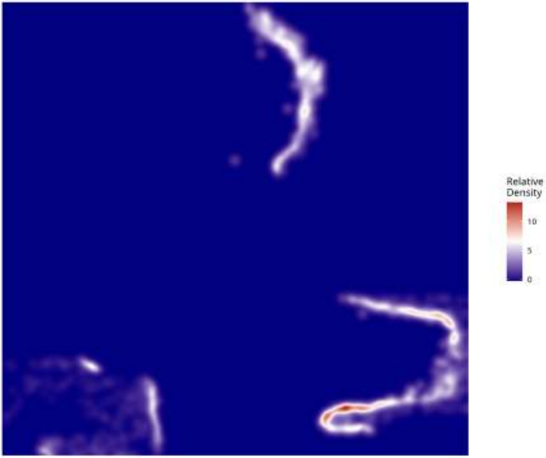

**Relative Density - Patient: um12 | Target: GPM | Modeled: OPC**

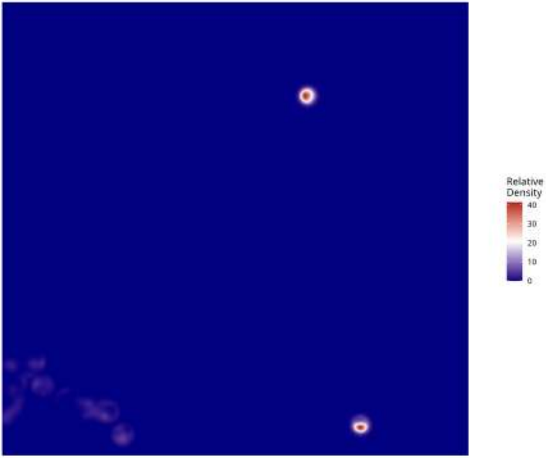

**Relative Density - Patient: um12 | Target: GPM | Modeled: PPR**

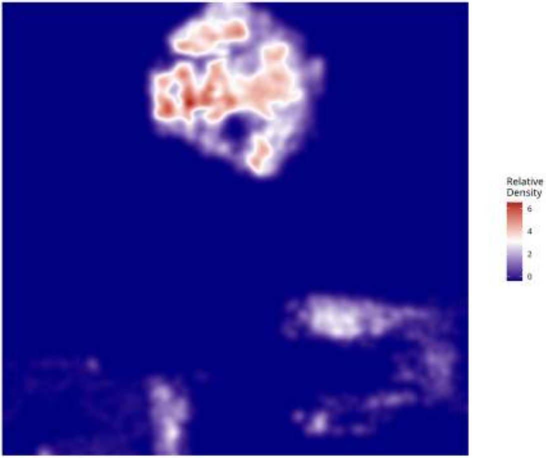

Relative Density - Patient: um12 | Target: GPM | Modeled: Tcells

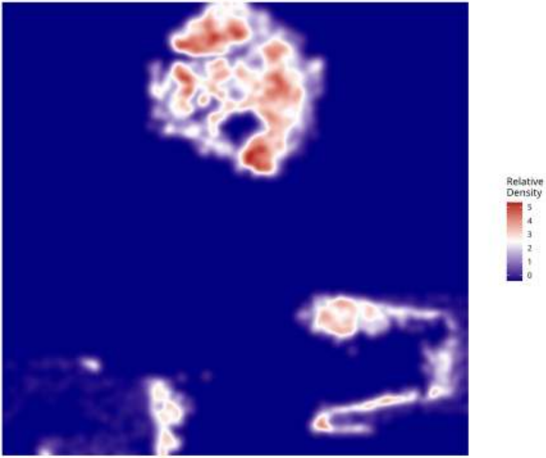

**Relative Density - Patient: um12 | Target: GPM | Modeled: vasculature**

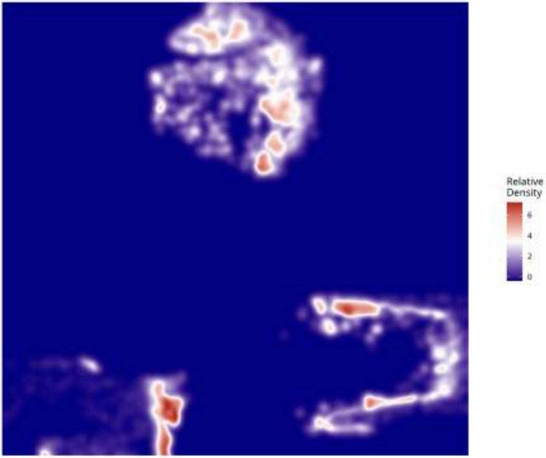

**Relative Density - Patient: um12 | Target: MTC | Modeled: astrocytes**

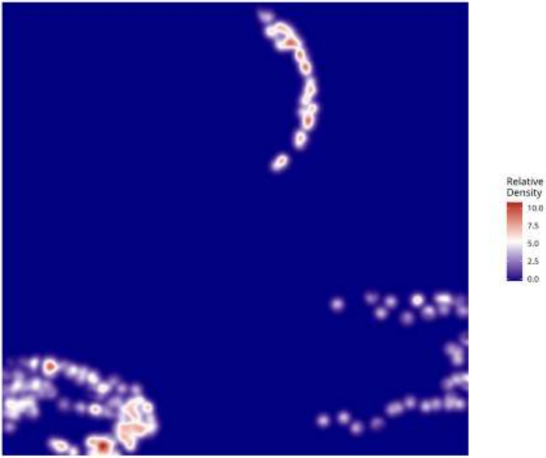

Relative Density - Patient: um12 | Target: MTC | Modeled: cDC

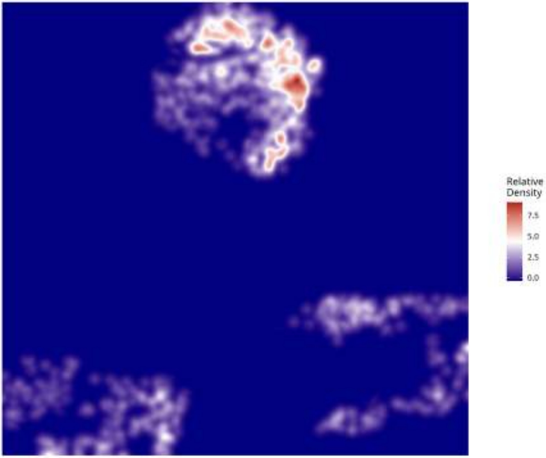

**Relative Density - Patient: um12 | Target: MTC | Modeled: GPM**

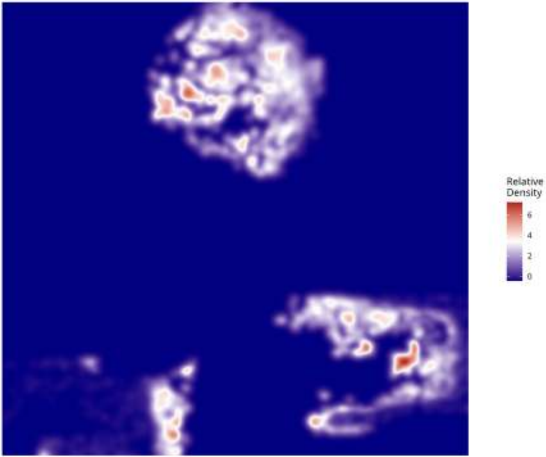

**Relative Density - Patient: um12 | Target: MTC | Modeled: Macrophage**

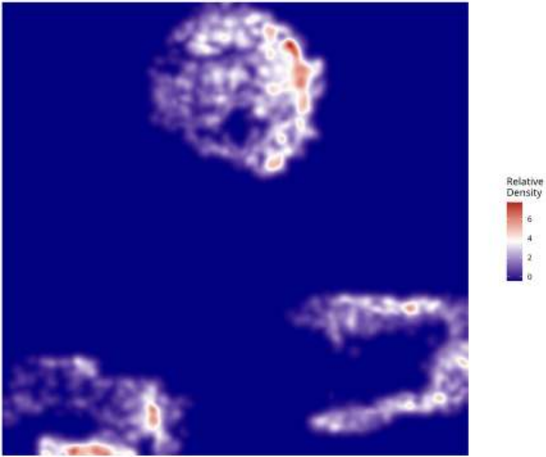

**Relative Density - Patient: um12 | Target: MTC | Modeled: Microglia**

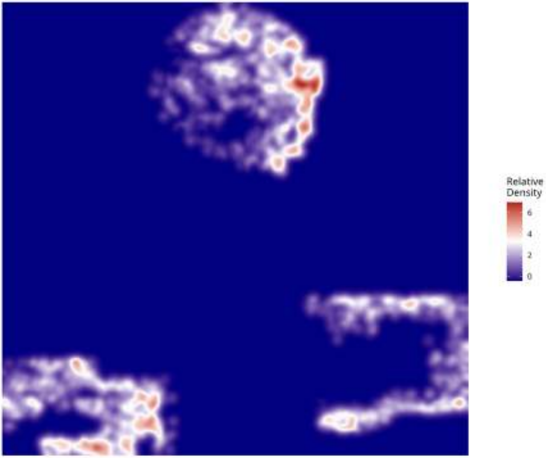

**Relative Density - Patient: um12 | Target: MTC | Modeled: Monocyte**

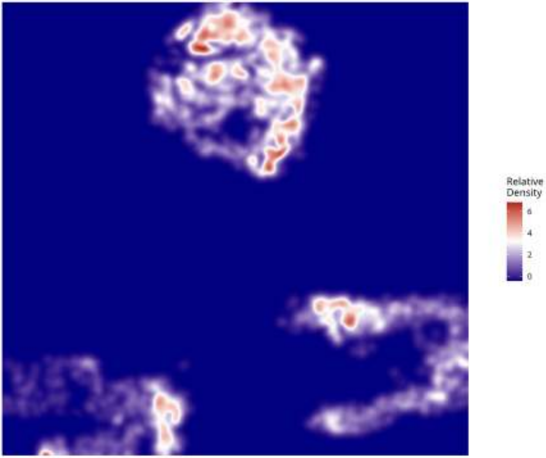

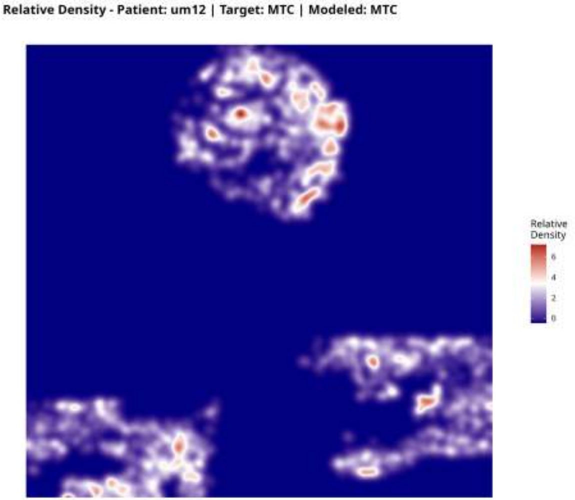
Relative Density - Patient: um12 | Target: MTC | Modeled: MTC
Relative Density
6
4
2
0

**Relative Density - Patient: um12 | Target: MTC | Modeled: NEU**

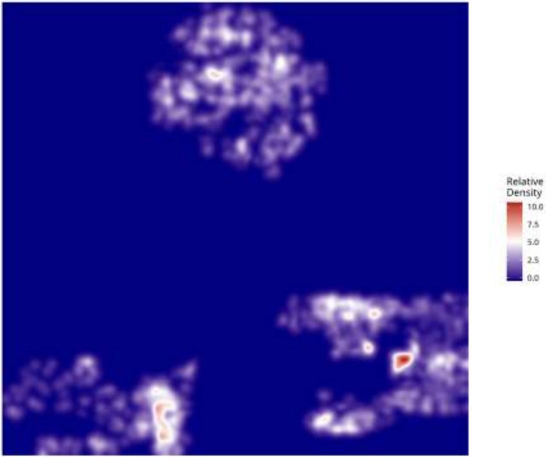

**Relative Density - Patient: um12 | Target: MTC | Modeled: Neutrophil**

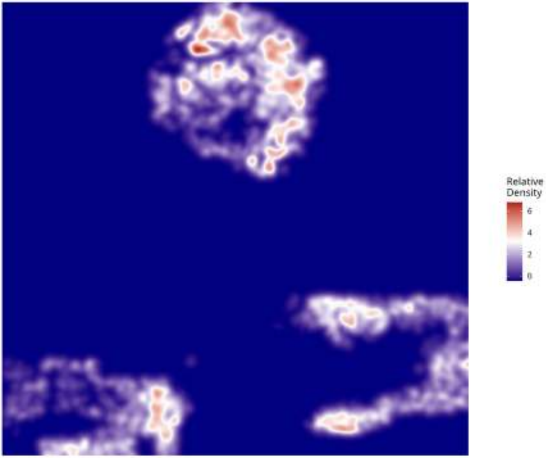

**Relative Density - Patient: um12 | Target: MTC | Modeled: oligodendrocyte**

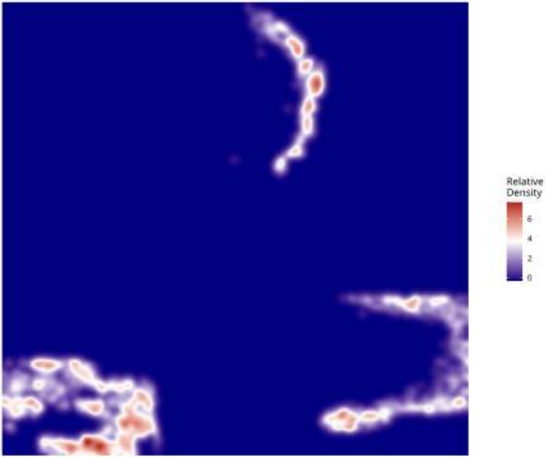

**Relative Density - Patient: um12 | Target: MTC | Modeled: OPC**

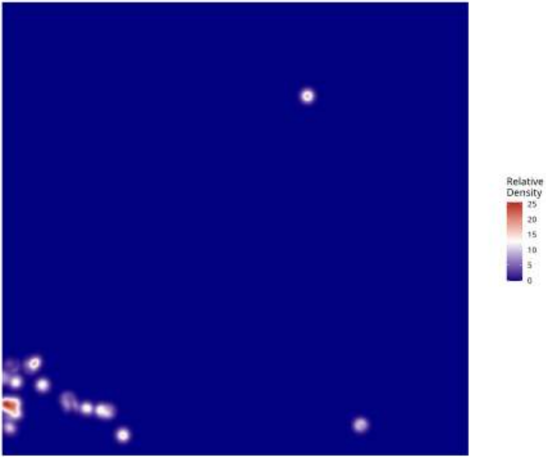

Relative Density - Patient: um12 | Target: MTC | Modeled: PPR

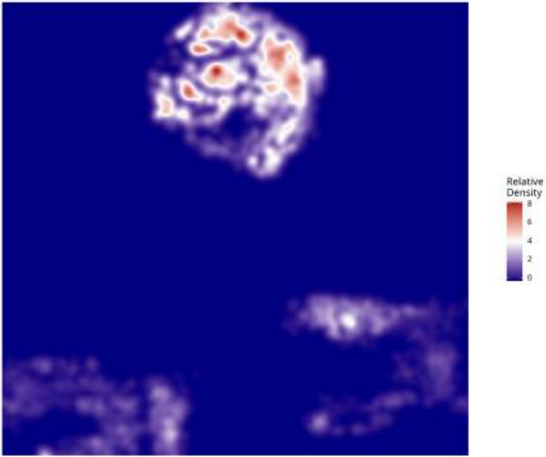

**Relative Density - Patient: um12 | Target: MTC | Modeled: Tcells**

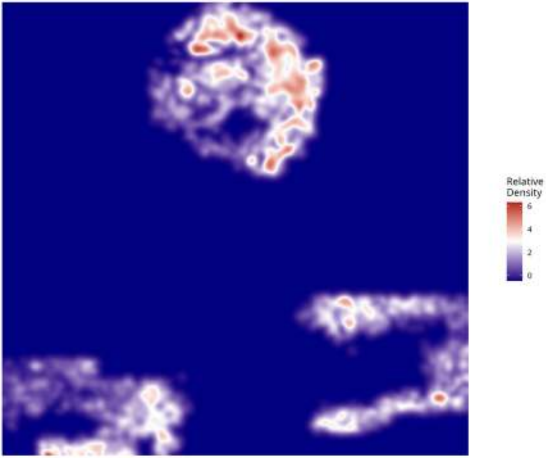

**Relative Density - Patient: um12 | Target: MTC | Modeled: vasculature**

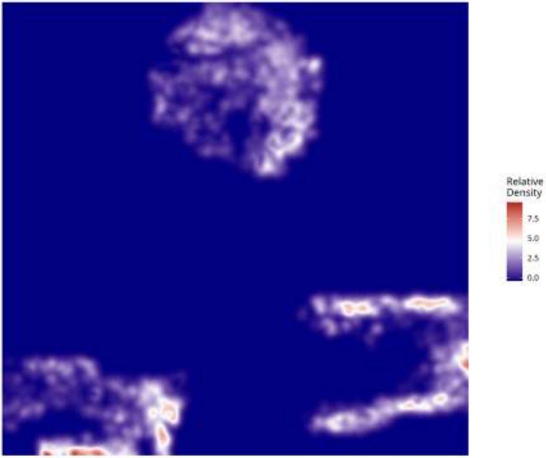

**Relative Density - Patient: um12 | Target: NEU | Modeled: astrocytes**

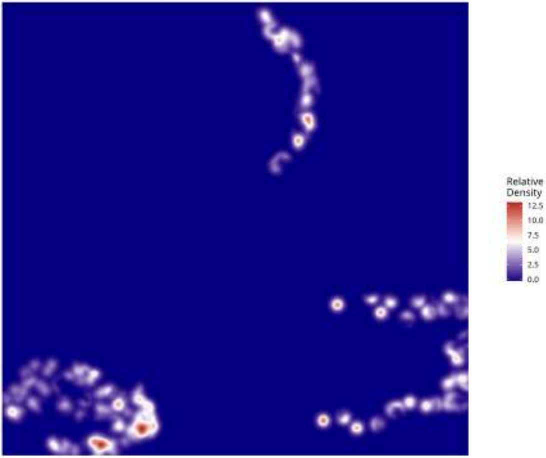

**Relative Density - Patient: um12 | Target: NEU | Modeled: cDC**

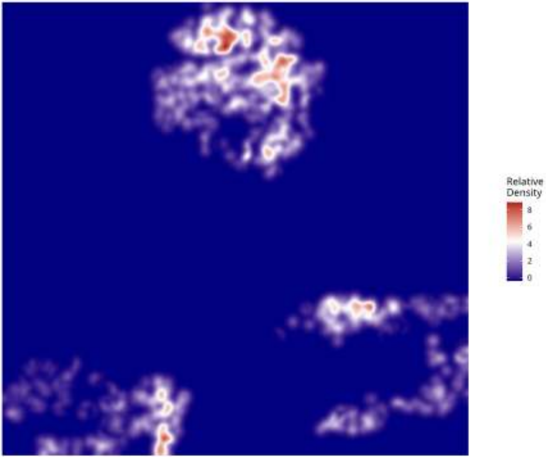

**Relative Density - Patient: um12 | Target: NEU | Modeled: GPM**

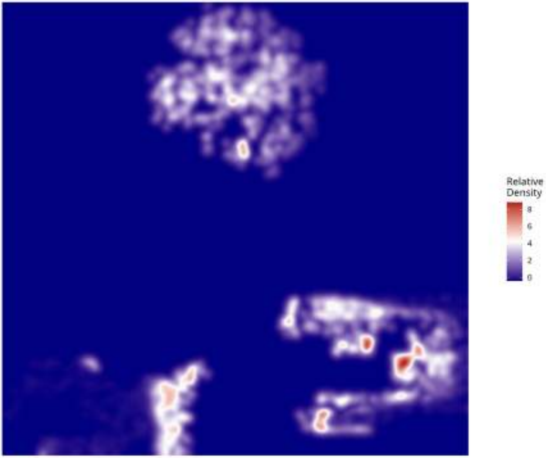

**Relative Density - Patient: um12 | Target: NEU | Modeled: Macrophage**

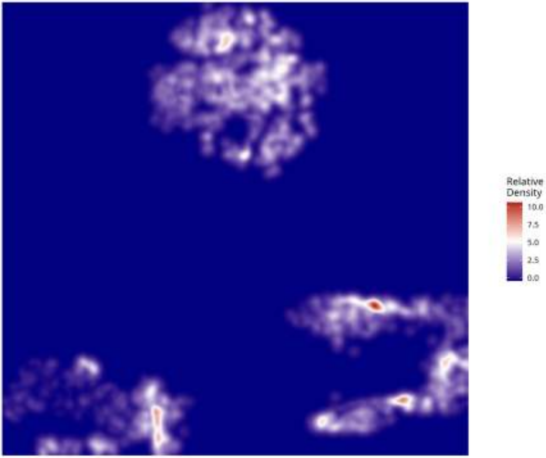

Relative Density - Patient: um12 | Target: NEU | Modeled: Microglia

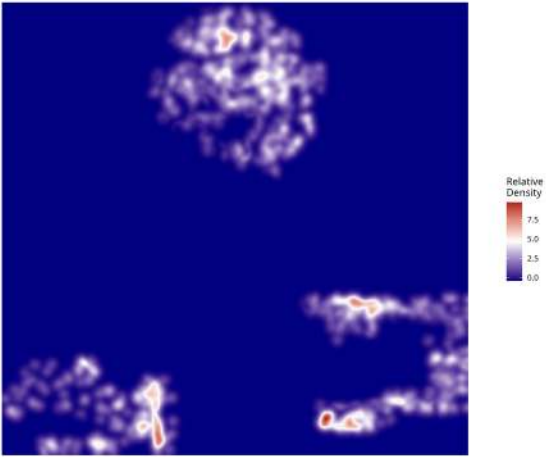

**Relative Density - Patient: um12 | Target: NEU | Modeled: Monocyte**

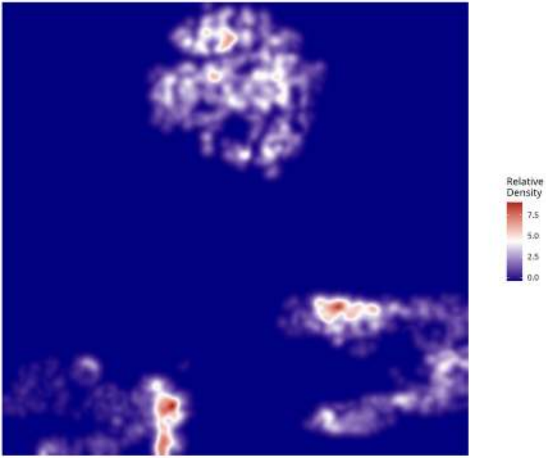

**Relative Density - Patient: um12 | Target: NEU | Modeled: MTC**

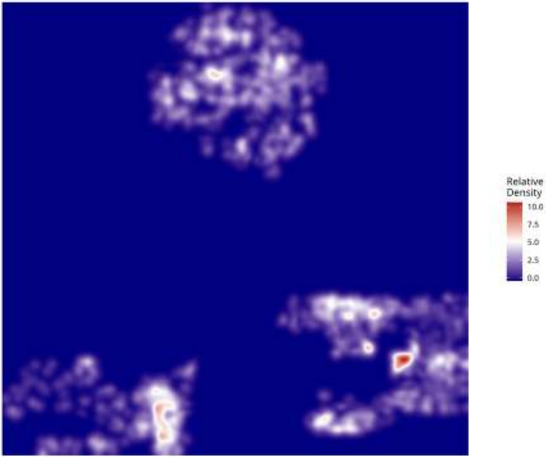

**Relative Density - Patient: um12 | Target: NEU | Modeled: NEU**

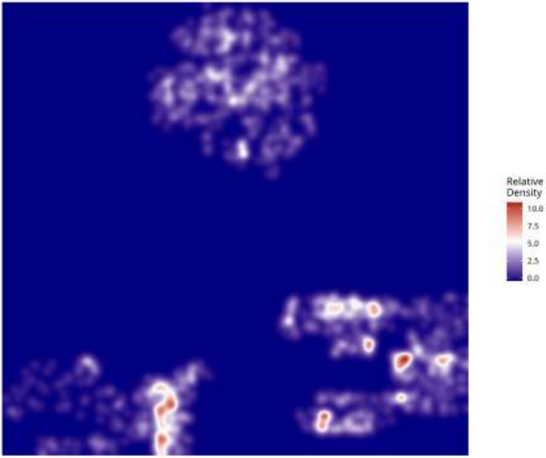

**Relative Density - Patient: um12 | Target: NEU | Modeled: Neutrophil**

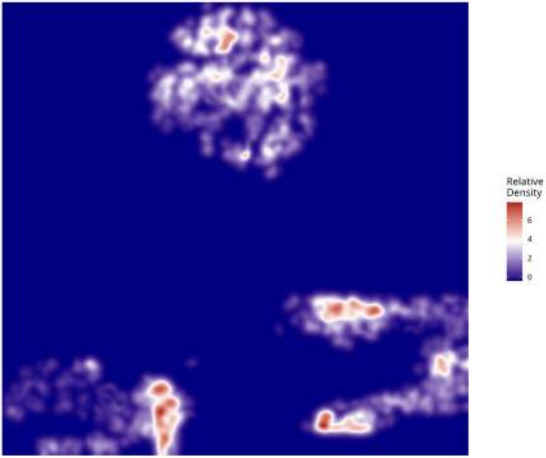

Relative Density - Patient: um12 | Target: NEU | Modeled: oligodendrocyte

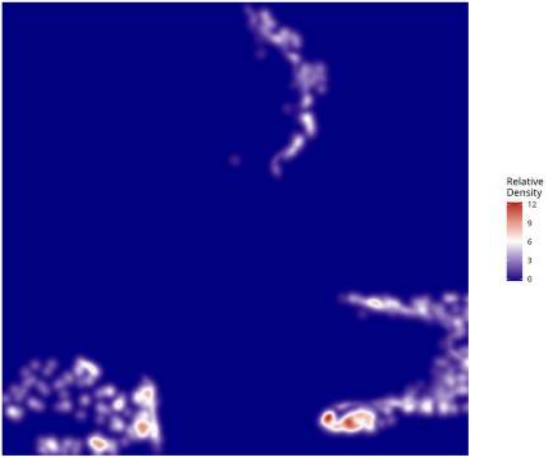

**Relative Density - Patient: um12 | Target: NEU | Modeled: OPC**

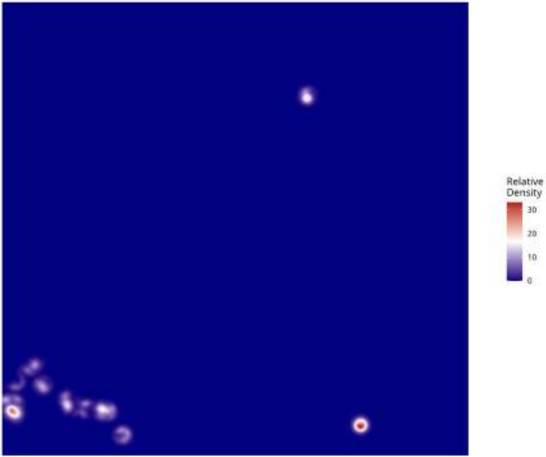

**Relative Density - Patient: um12 | Target: NEU | Modeled: PPR**

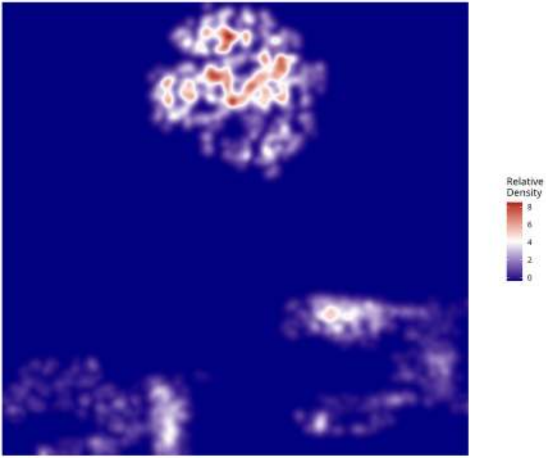

**Relative Density - Patient: um12 | Target: NEU | Modeled: Tcells**

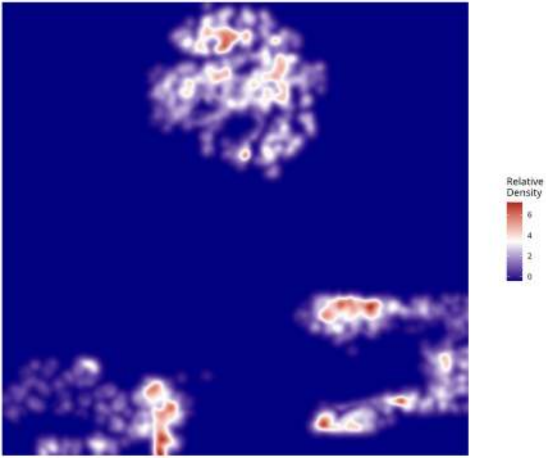

**Relative Density - Patient: um12 | Target: NEU | Modeled: vasculature**

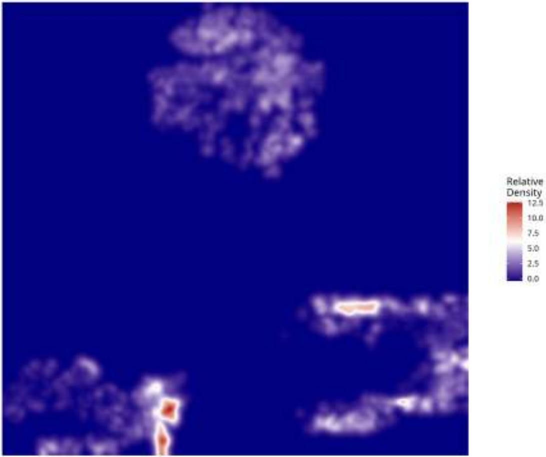

**Relative Density - Patient: um12 | Target: PPR | Modeled: astrocytes**

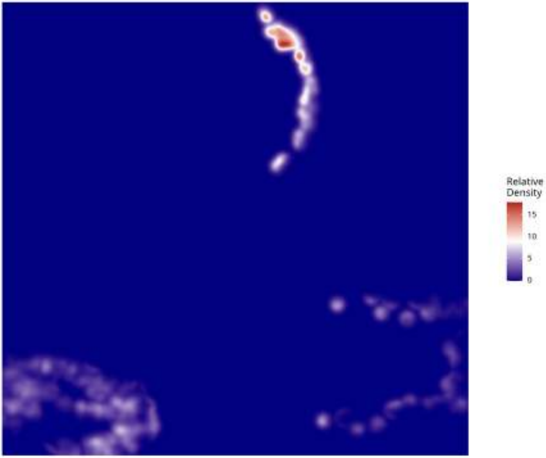

Relative Density - Patient: um12 | Target: PPR | Modeled: cDC

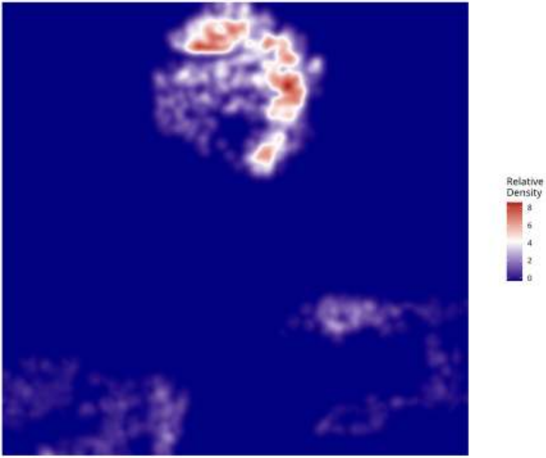

**Relative Density - Patient: um12 | Target: PPR | Modeled: GPM**

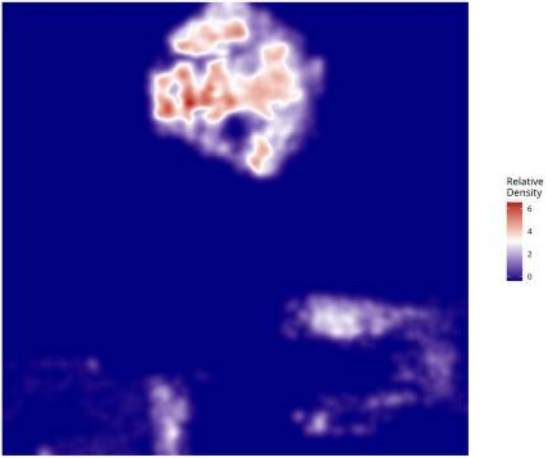

**Relative Density - Patient: um12 | Target: PPR | Modeled: Macrophage**

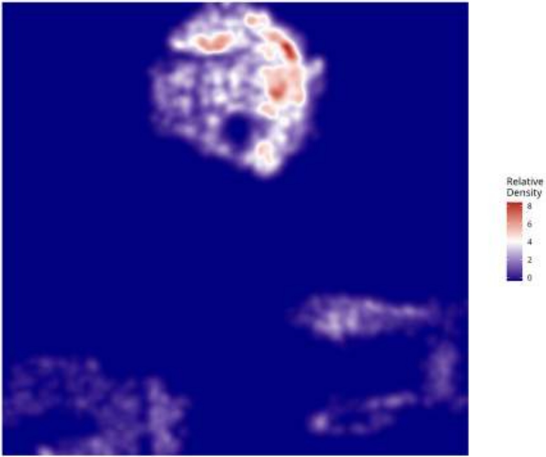

Relative Density - Patient: um12 | Target: PPR | Modeled: Microglia

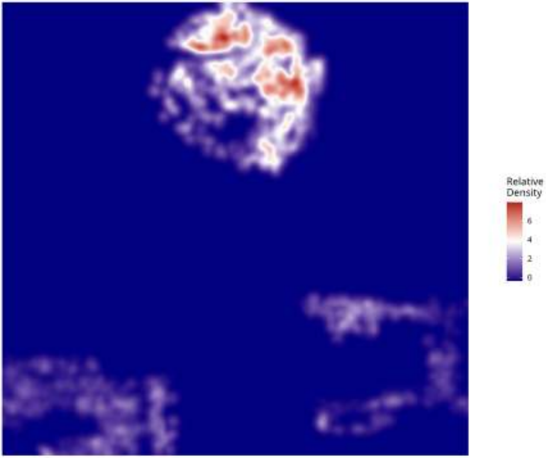

**Relative Density - Patient: um12 | Target: PPR | Modeled: Monocyte**

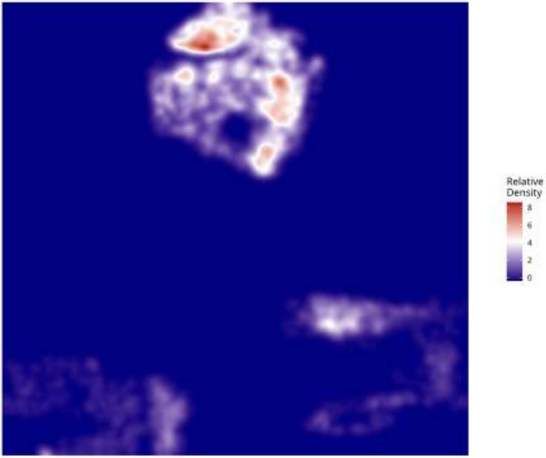

Relative Density - Patient: um12 | Target: PPR | Modeled: MTC

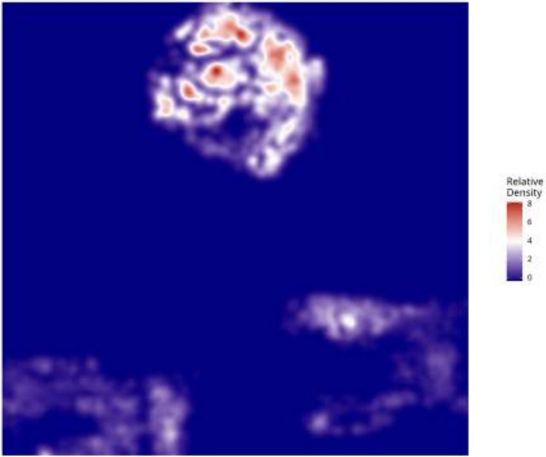

**Relative Density - Patient: um12 | Target: PPR | Modeled: NEU**

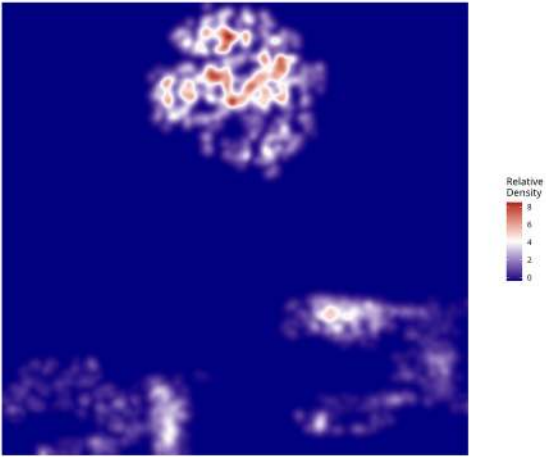

**Relative Density - Patient: um12 | Target: PPR | Modeled: Neutrophil**

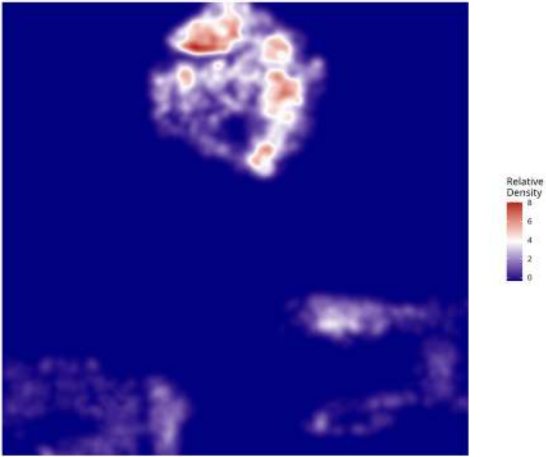

**Relative Density - Patient: um12 | Target: PPR | Modeled: oligodendrocyte**

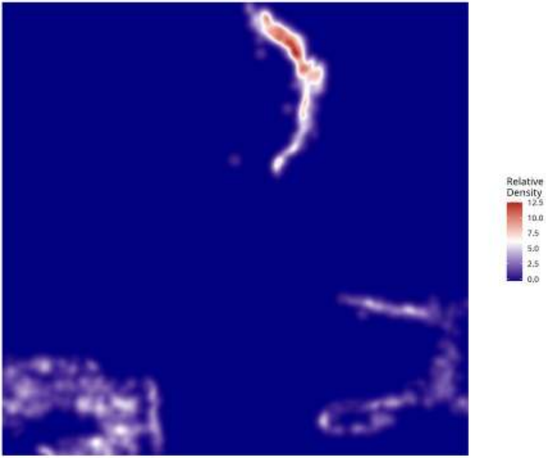

Relative Density - Patient: um12 | Target: PPR | Modeled: OPC

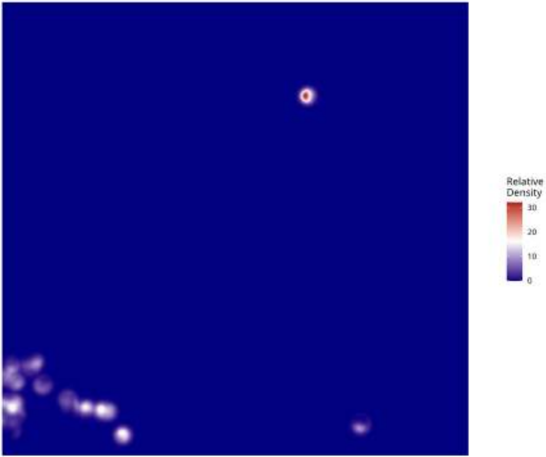

Relative Density - Patient: um12 | Target: PPR | Modeled: PPR

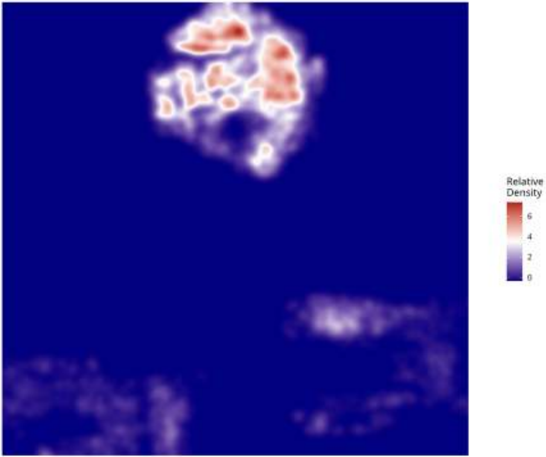

**Relative Density - Patient: um12 | Target: PPR | Modeled: Tcells**

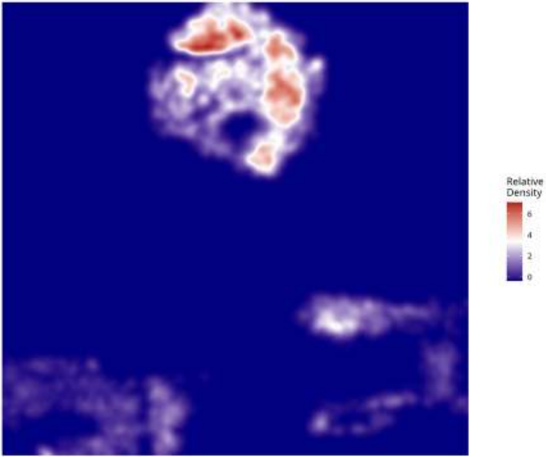

**Relative Density - Patient: um12 | Target: PPR | Modeled: vasculature**

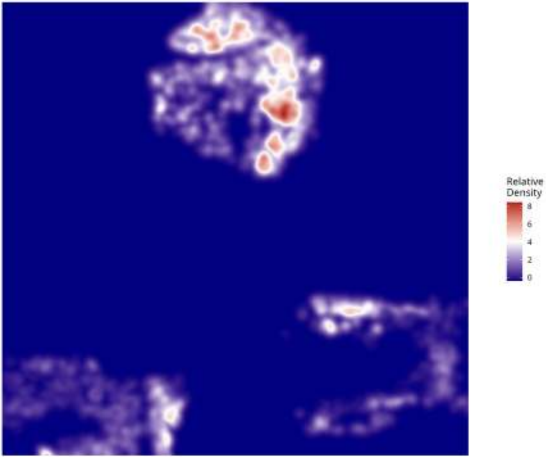

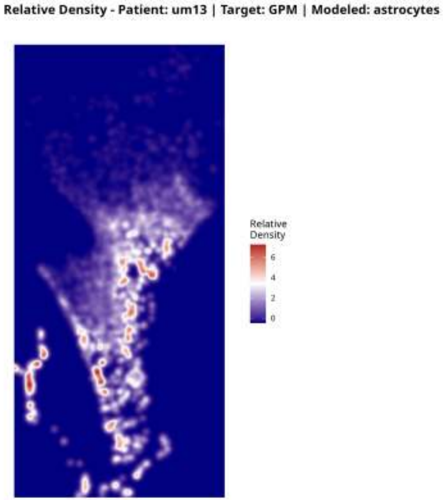
Relative Density - Patient: um13 | Target: GPM | Modeled: astrocytes
Relative
Density
6
4
2
0

**Relative Density - Patient: um13 | Target: GPM | Modeled: cDC**

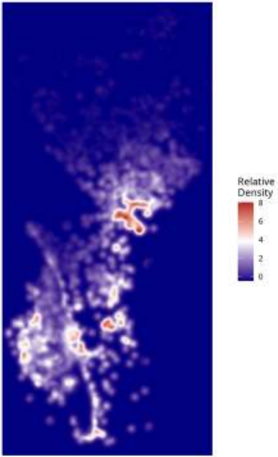

Relative Density - Patient: um13 | Target: GPM | Modeled: GPM

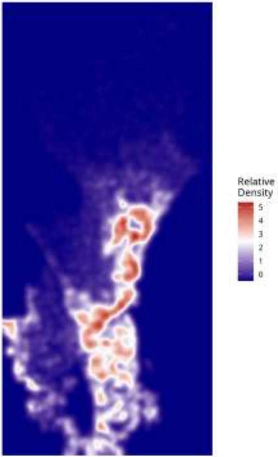

Relative Density - Patient: um13 | Target: GPM | Modeled: Macrophage

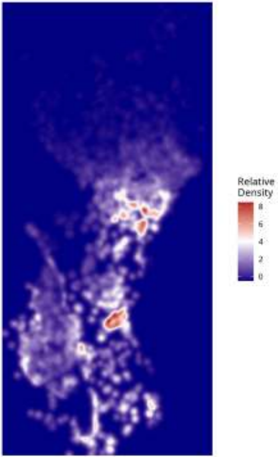

Relative Density - Patient: um13 | Target: GPM | Modeled: Microglia

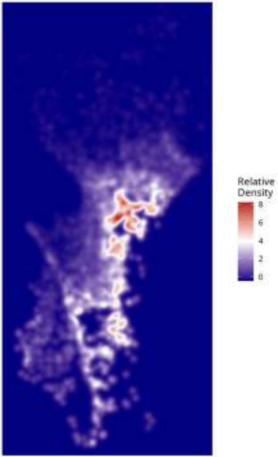

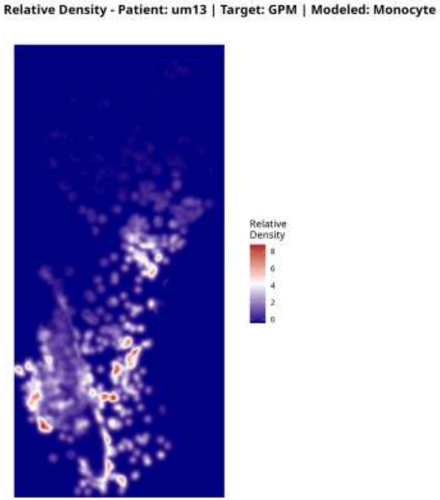
Relative Density - Patient: um13 | Target: GPM | Modeled: Monocyte
Relative Density
8
6
4
2
0

**Relative Density - Patient: um13 | Target: GPM | Modeled: MTC**

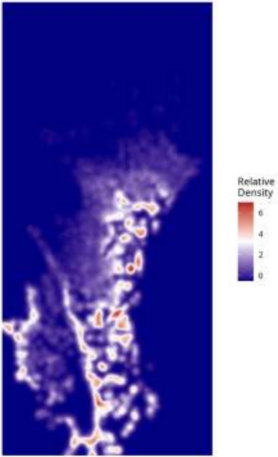

#### Relative Density - Patient: um13 | Target: GPM | Modeled: NEU

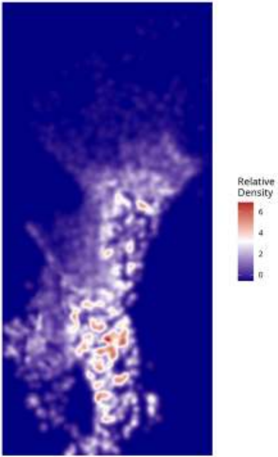

Relative Density - Patient: um13 | Target: GPM | Modeled: neurons

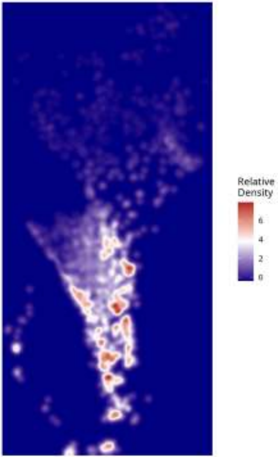

Relative Density - Patient: um13 | Target: GPM | Modeled: Neutrophil

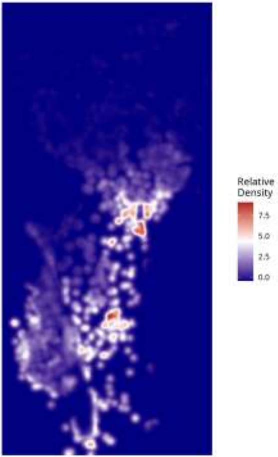

**Relative Density - Patient: um13 | Target: GPM | Modeled: oligodendrocyte**

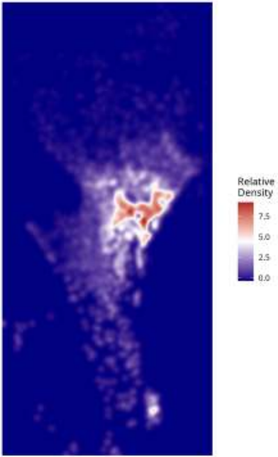

Relative Density - Patient: um13 | Target: GPM | Modeled: OPC

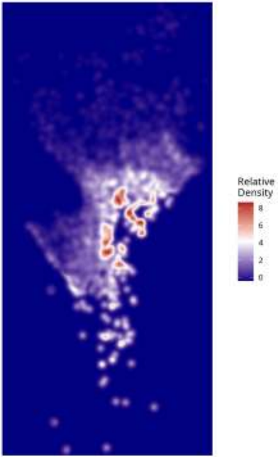

Relative Density - Patient: um13 | Target: GPM | Modeled: PPR

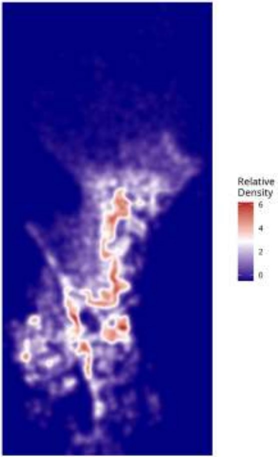

Relative Density - Patient: um13 | Target: GPM | Modeled: Tcells

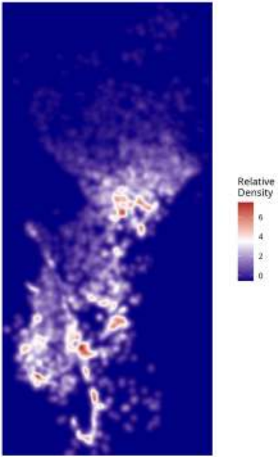

Relative Density - Patient: um13 | Target: GPM | Modeled: vasculature

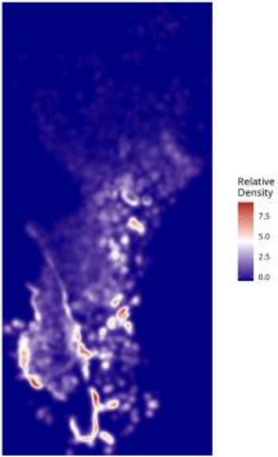

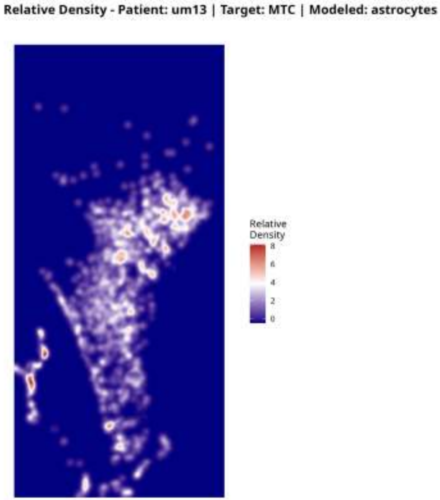
Relative Density - Patient: um13 | Target: MTC | Modeled: astrocytes
Relative Density
8
6
4
2
0

Relative Density - Patient: um13 | Target: MTC | Modeled: cDC

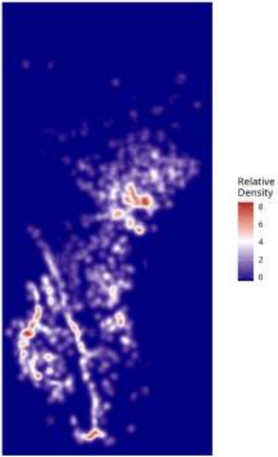

Relative Density - Patient: um13 | Target: MTC | Modeled: GPM

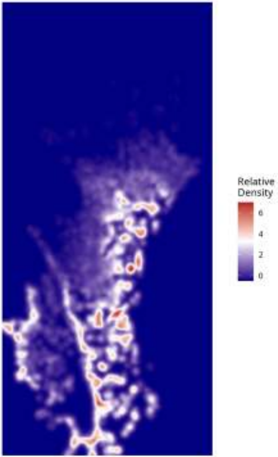

Relative Density - Patient: um13 | Target: MTC | Modeled: Macrophage

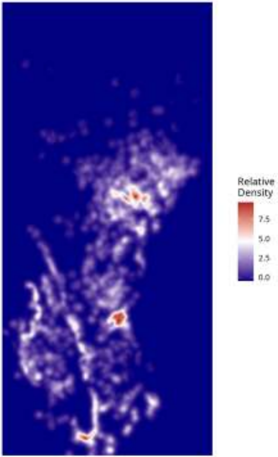

Relative Density - Patient: um13 | Target: MTC | Modeled: Microglia

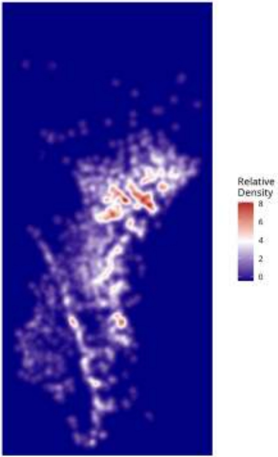

Relative Density - Patient: um13 | Target: MTC | Modeled: Monocyte

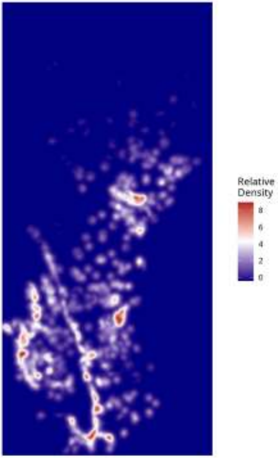

**Relative Density - Patient: um13 | Target: MTC | Modeled: MTC**

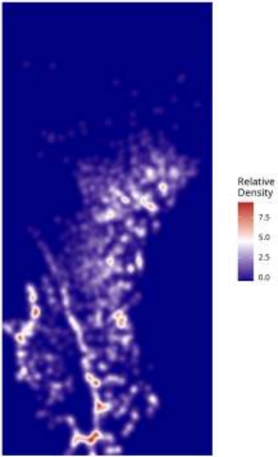

Relative Density - Patient: um13 | Target: MTC | Modeled: NEU

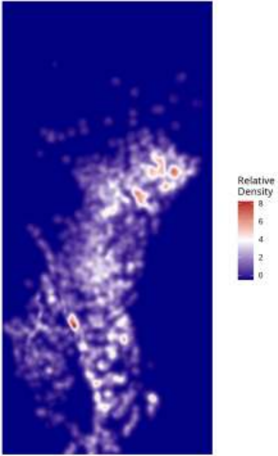

Relative Density - Patient: um13 | Target: MTC | Modeled: neurons

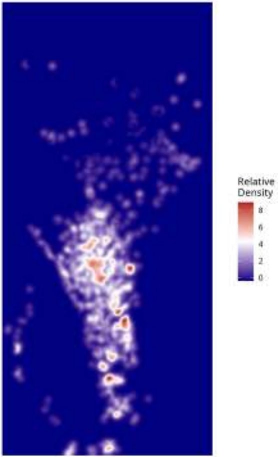

Relative Density - Patient: um13 | Target: MTC | Modeled: Neutrophil

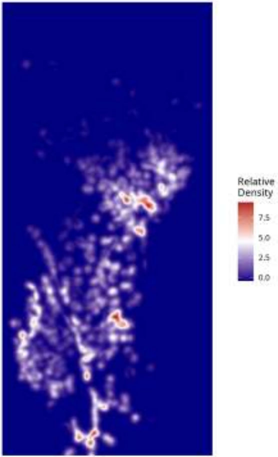

**Relative Density - Patient: um13 | Target: MTC | Modeled: oligodendrocyte**

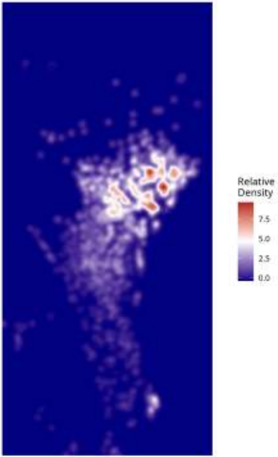

Relative Density - Patient: um13 | Target: MTC | Modeled: OPC

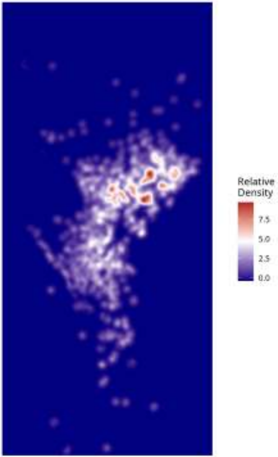

**Relative Density - Patient: um13 | Target: MTC | Modeled: PPR**

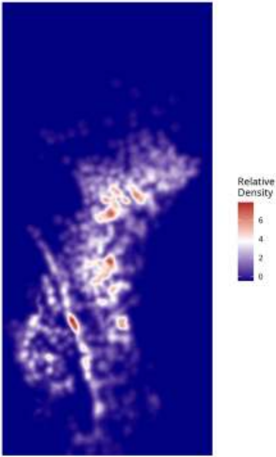

**Relative Density - Patient: um13 | Target: MTC | Modeled: Tcells**

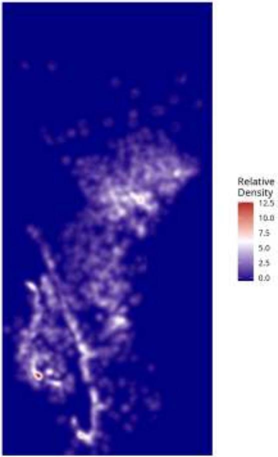

**Relative Density - Patient: um13 | Target: MTC | Modeled: vasculature**

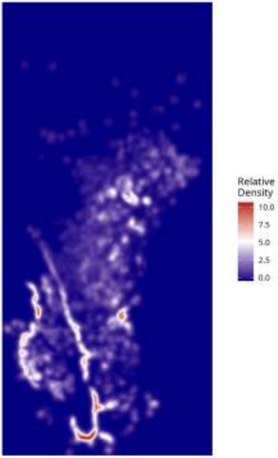

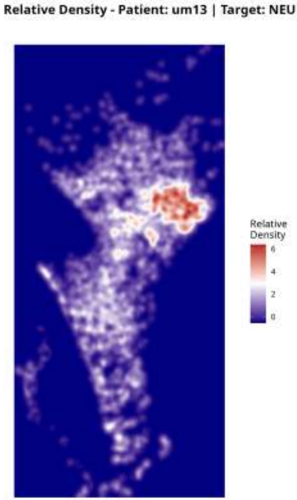

Relative Density - Patient: um13 | Target: NEU | Modeled: astrocytes
Relative Density
6
4
2
0

Relative Density - Patient: um13 | Target: NEU | Modeled: cDC

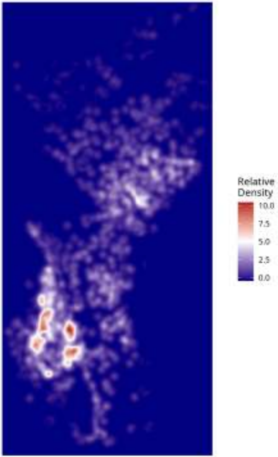

Relative Density - Patient: um13 | Target: NEU | Modeled: GPM

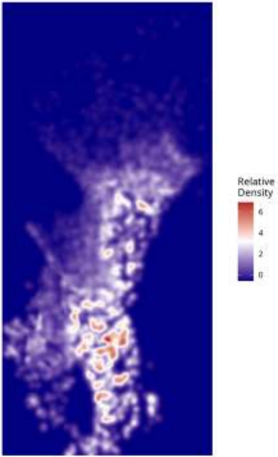

#### Relative Density - Patient: um13 | Target: NEU | Modeled: Macrophage

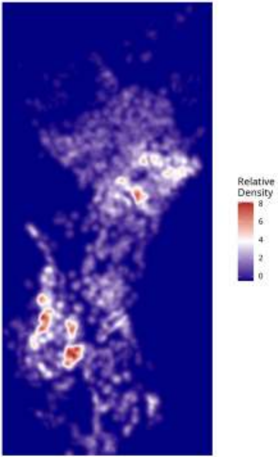

Relative Density - Patient: um13 | Target: NEU | Modeled: Microglia

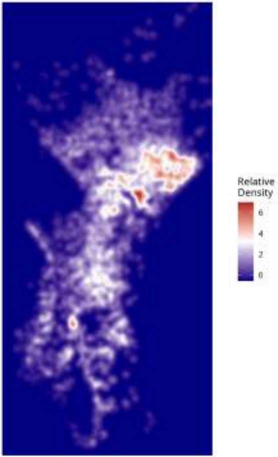

Relative Density - Patient: um13 | Target: NEU | Modeled: Monocyte

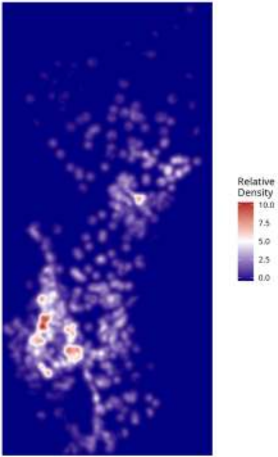

Relative Density - Patient: um13 | Target: NEU | Modeled: MTC

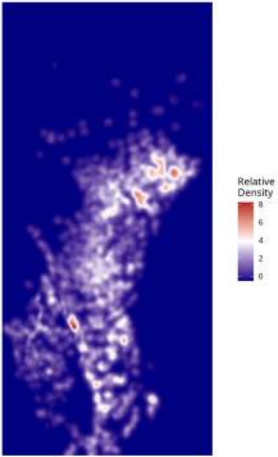

**Relative Density - Patient: um13 | Target: NEU | Modeled: NEU**

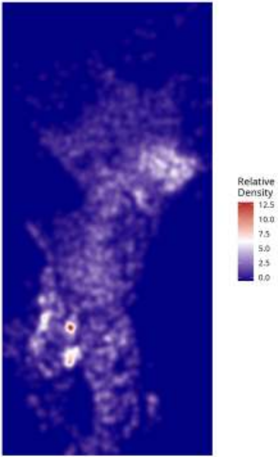

Relative Density - Patient: um13 | Target: NEU | Modeled: neurons

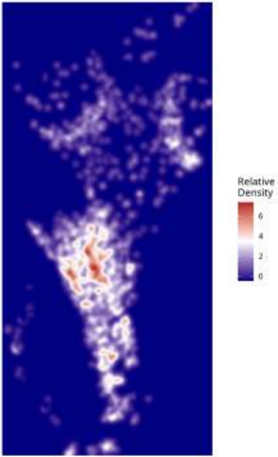

Relative Density - Patient: um13 | Target: NEU | Modeled: Neutrophil

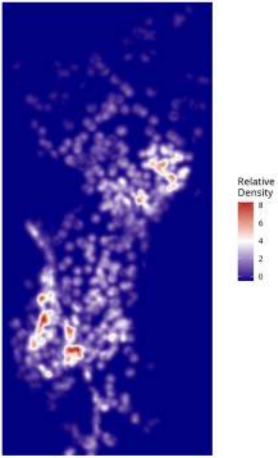

Relative Density - Patient: um13 | Target: NEU | Modeled: oligodendrocyte

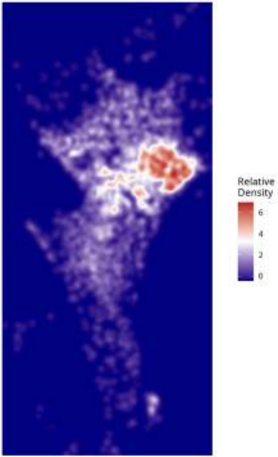

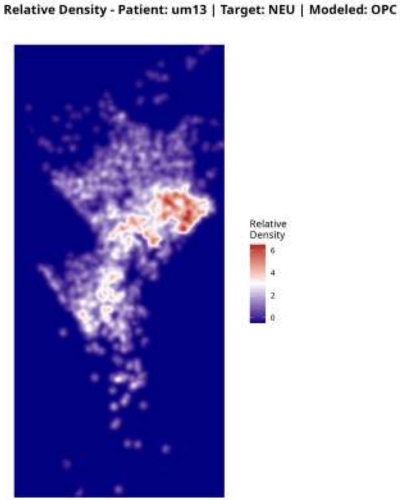
Relative Density - Patient: um13 | Target: NEU | Modeled: OPC
Relative
Density
6
4
2
0

**Relative Density - Patient: um13 | Target: NEU | Modeled: PPR**

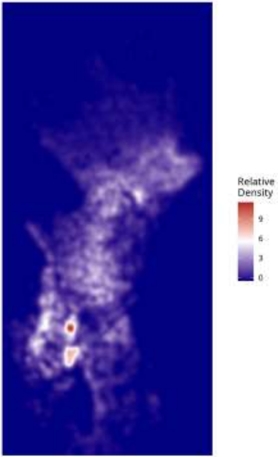

Relative Density - Patient: um13 | Target: NEU | Modeled: Tcells

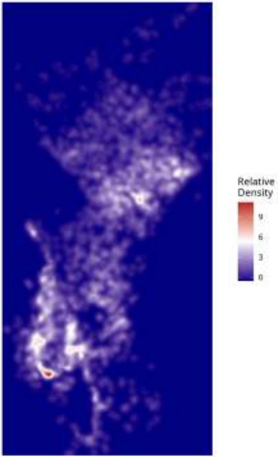

Relative Density - Patient: um13 | Target: NEU | Modeled: vasculature

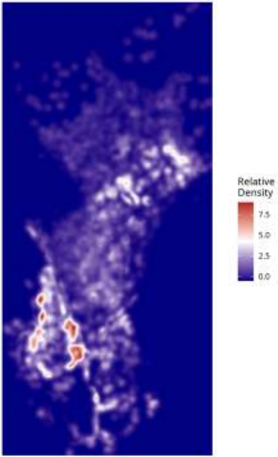

Relative Density - Patient: um13 | Target: PPR | Modeled: astrocytes

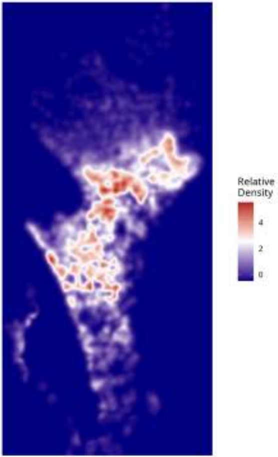

**Relative Density - Patient: um13 | Target: PPR | Modeled: cDC**

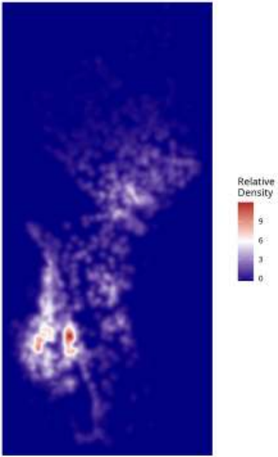

**Relative Density - Patient: um13 | Target: PPR | Modeled: GPM**

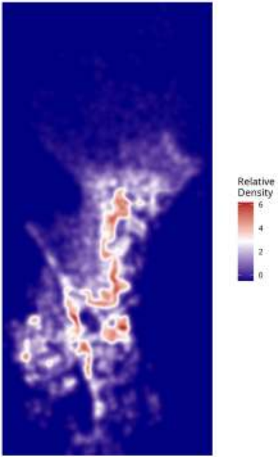

Relative Density - Patient: um13 | Target: PPR | Modeled: Macrophage

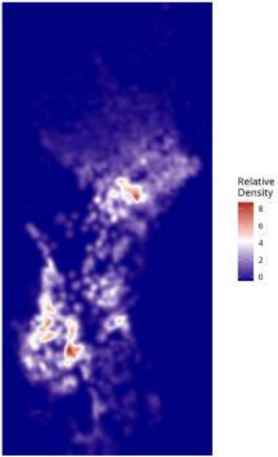

**Relative Density - Patient: um13 | Target: PPR | Modeled: Microglia**

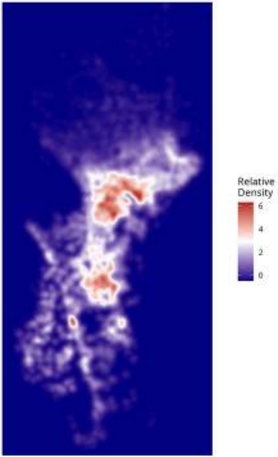

**Relative Density - Patient: um13 | Target: PPR | Modeled: Monocyte**

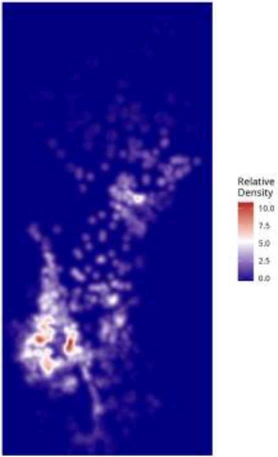

**Relative Density - Patient: um13 | Target: PPR | Modeled: MTC**

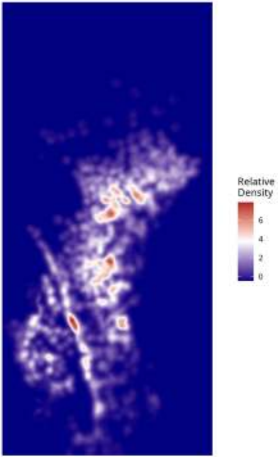

Relative Density - Patient: um13 | Target: PPR | Modeled: NEU

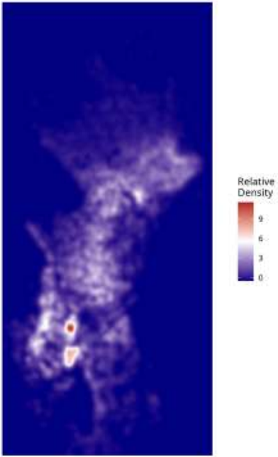

**Relative Density - Patient: um13 | Target: PPR | Modeled: neurons**

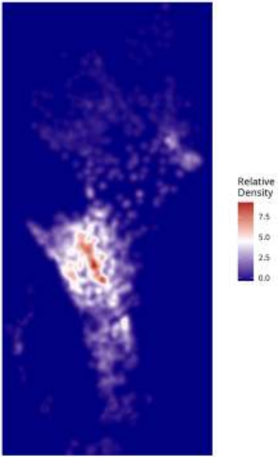

Relative Density - Patient: um13 | Target: PPR | Modeled: Neutrophil

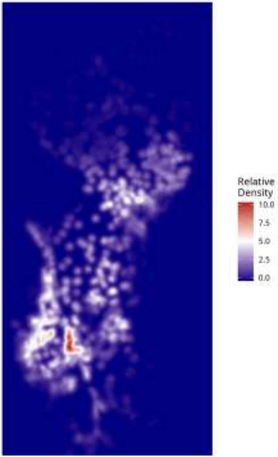

Relative Density - Patient: um13 | Target: PPR | Modeled: oligodendrocyte

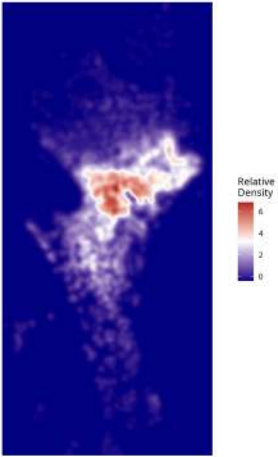

Relative Density - Patient: um13 | Target: PPR | Modeled: OPC

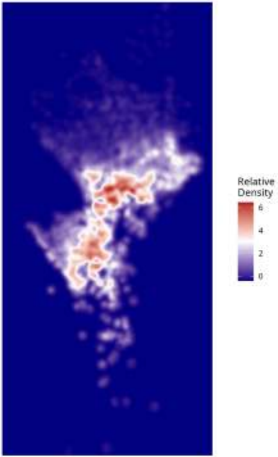

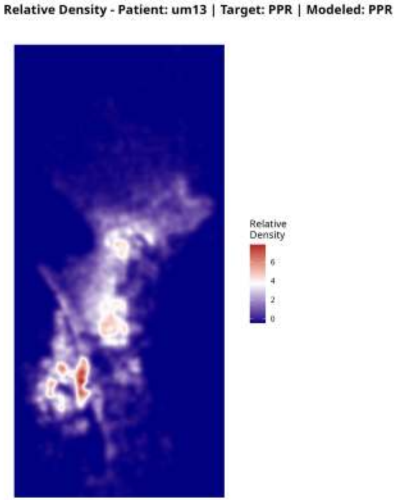
Relative Density - Patient: um13 | Target: PPR | Modeled: PPR
Relative Density
6
4
2
0

#### Relative Density - Patient: um13 | Target: PPR | Modeled: Tcells

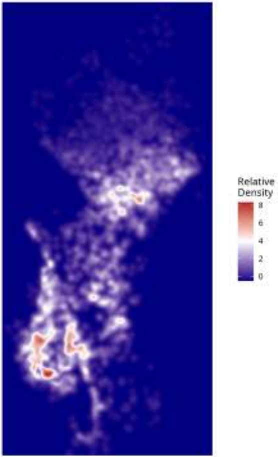

Relative Density - Patient: um13 | Target: PPR | Modeled: vasculature

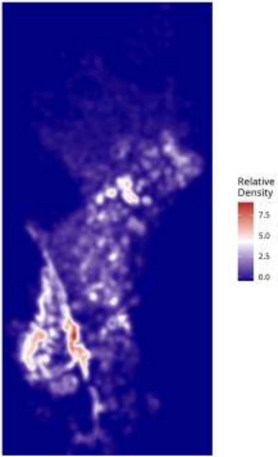

Relative Density - Patient: um14 | Target: GPM | Modeled: astrocytes

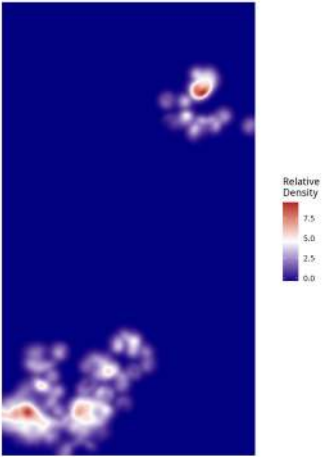

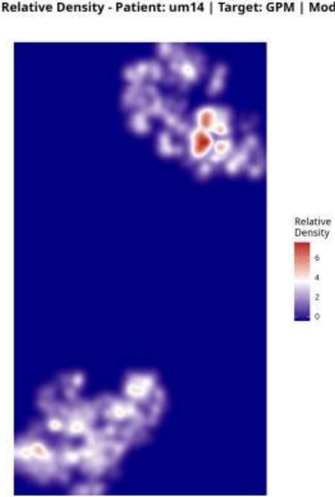
Relative Density - Patient: um14 | Target: GPM | Modeled: cDC
Relative
Density
6
4
2
0

**Relative Density - Patient: um14 | Target: GPM | Modeled: GPM**

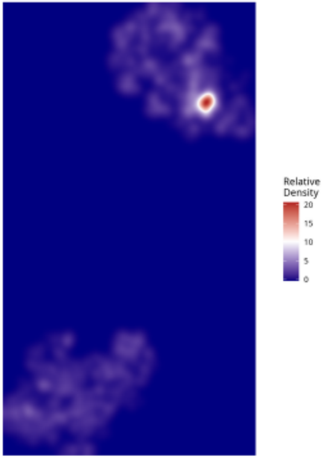

Relative Density - Patient: um14 | Target: GPM | Modeled: Macrophage

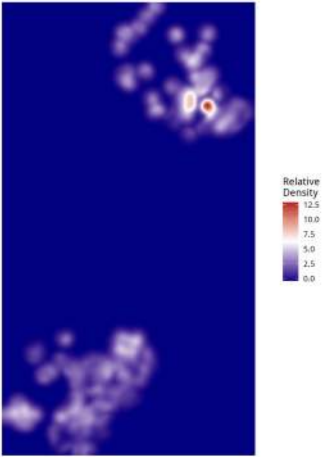

Relative Density - Patient: um14 | Target: GPM | Modeled: Microglia

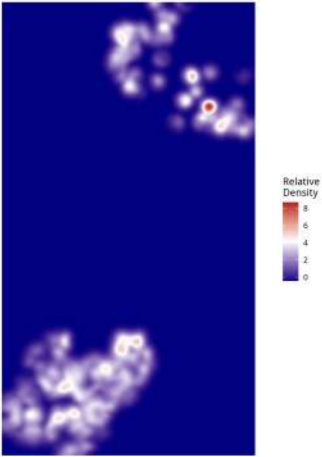

Relative Density - Patient: um14 | Target: GPM | Modeled: Monocyte

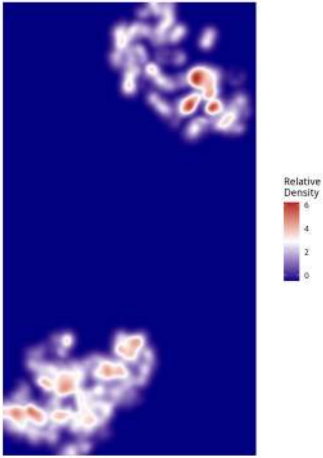

**Relative Density - Patient: um14 | Target: GPM | Modeled: MTC**

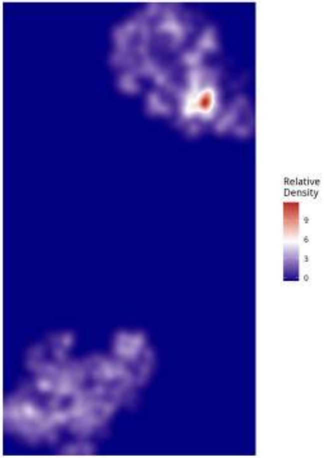

Relative Density - Patient: um14 | Target: GPM | Modeled: NEU

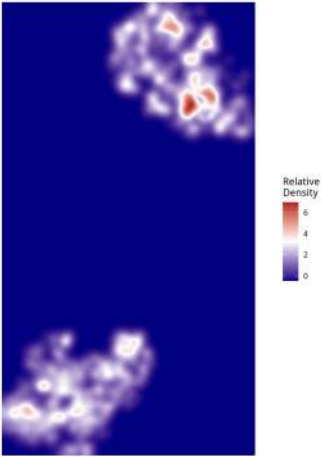

Relative Density - Patient: um14 | Target: GPM | Modeled: neurons

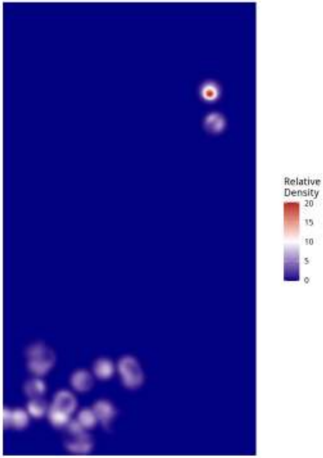

Relative Density - Patient: um14 | Target: GPM | Modeled: Neutrophil

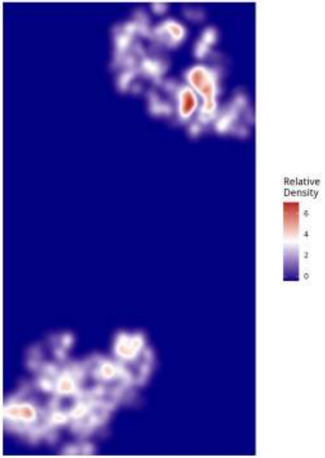

**Relative Density - Patient: um14 | Target: GPM | Modeled: oligodendrocyte**

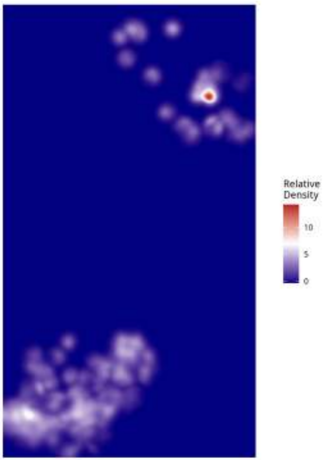

Relative Density - Patient: um14 | Target: GPM | Modeled: OPC

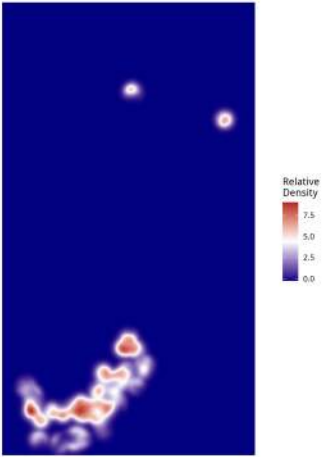

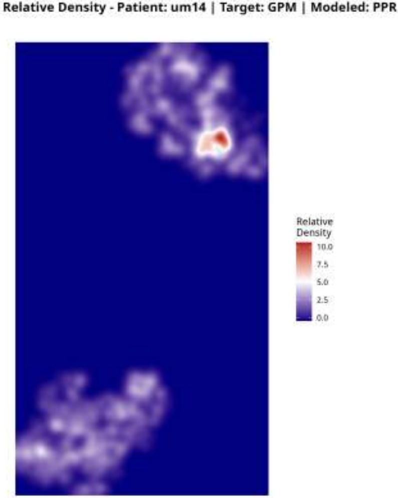
Relative Density - Patient: um14 | Target: GPM | Modeled: PPR
Relative
Density
10.0
7.5
5.0
2.5
0.0

#### Relative Density - Patient: um14 | Target: GPM | Modeled: Tcells

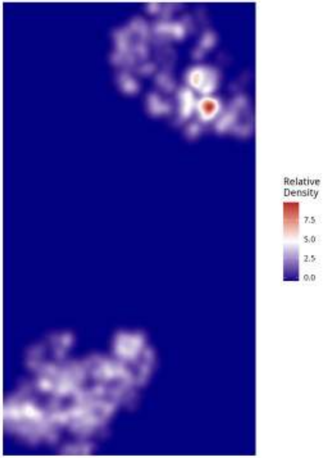

Relative Density - Patient: um14 | Target: GPM | Modeled: vasculature

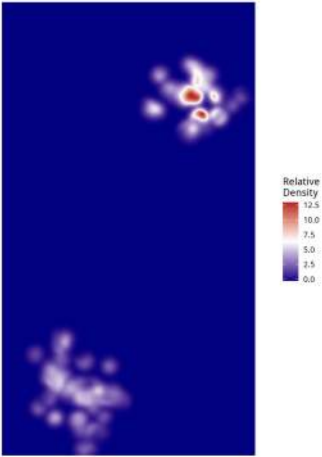

**Relative Density - Patient: um14 | Target: MTC | Modeled: astrocytes**

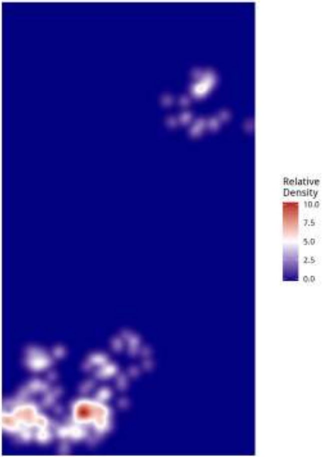

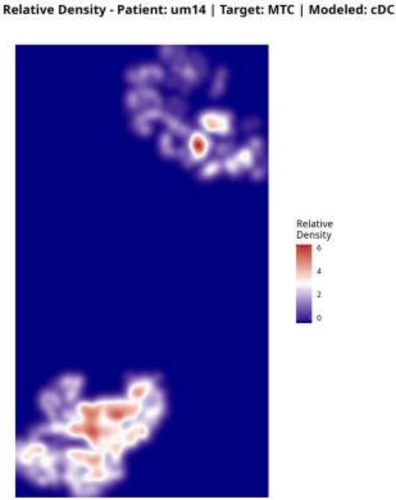
Relative Density - Patient: um14 | Target: MTC | Modeled: cDC
Relative Density
6
4
2
0

Relative Density - Patient: um14 | Target: MTC | Modeled: GPM

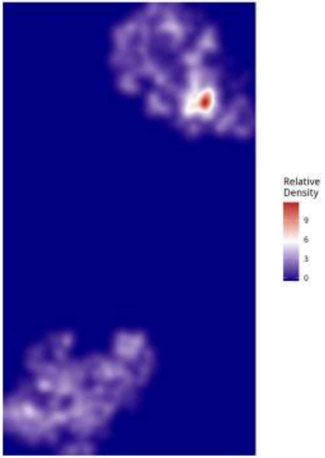

Relative Density - Patient: um14 | Target: MTC | Modeled: Macrophage

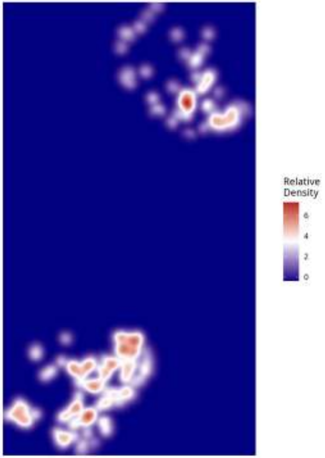

**Relative Density - Patient: um14 | Target: MTC | Modeled: Microglia**

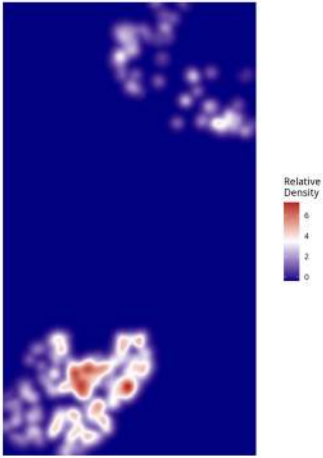

**Relative Density - Patient: um14 | Target: MTC | Modeled: Monocyte**

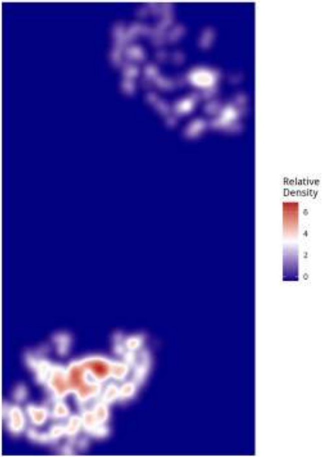

Relative Density - Patient: um14 | Target: MTC | Modeled: MTC

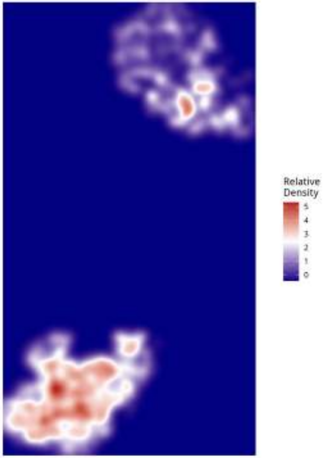

**Relative Density - Patient: um14 | Target: MTC | Modeled: NEU**

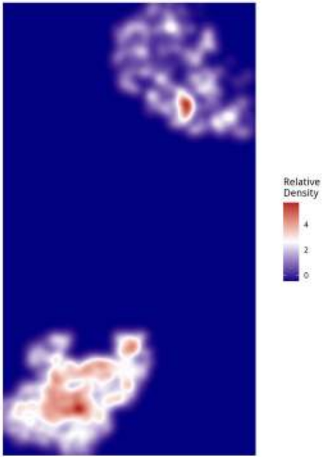

Relative Density - Patient: um14 | Target: MTC | Modeled: neurons

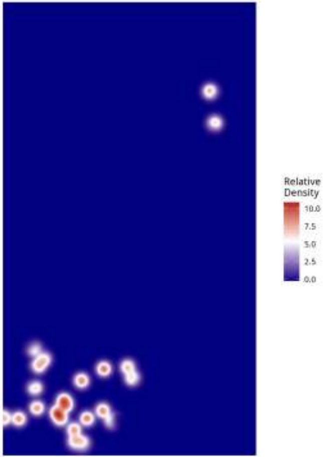

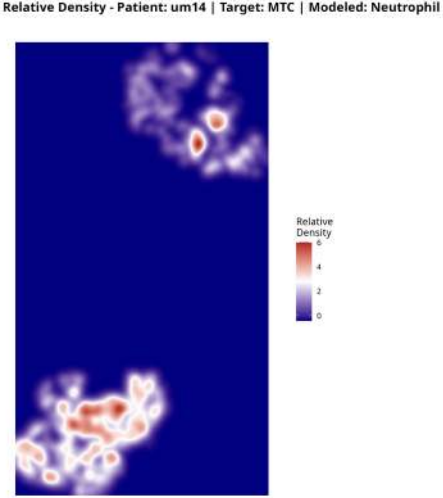
Relative Density - Patient: um14 | Target: MTC | Modeled: Neutrophil
Relative
Density
6
4
2
0

Relative Density - Patient: um14 | Target: MTC | Modeled: oligodendrocyte

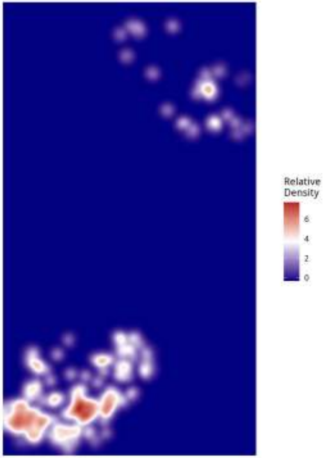

**Relative Density - Patient: um14 | Target: MTC | Modeled: OPC**

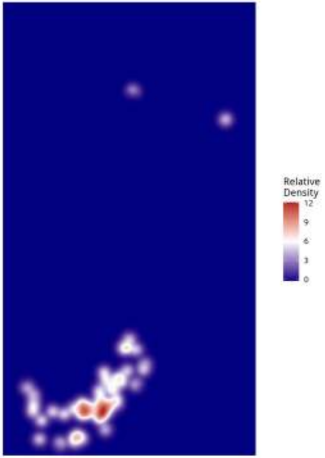

Relative Density - Patient: um14 | Target: MTC | Modeled: PPR

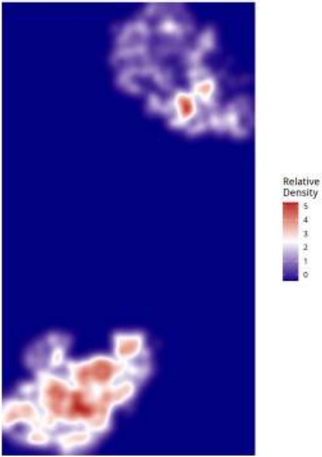

**Relative Density - Patient: um14 | Target: MTC | Modeled: Tcells**

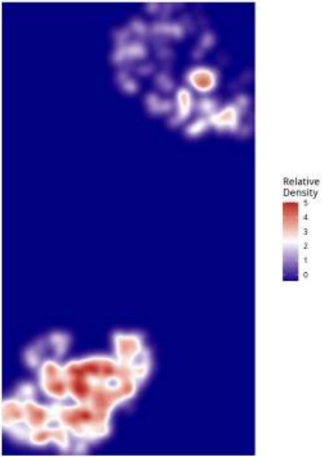

Relative Density - Patient: um14 | Target: MTC | Modeled: vasculature

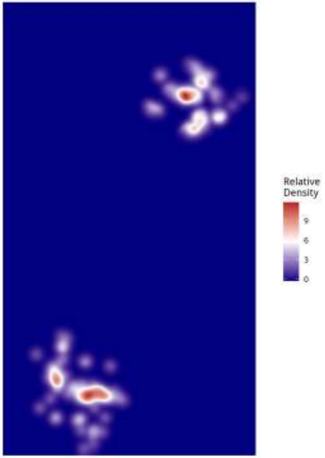

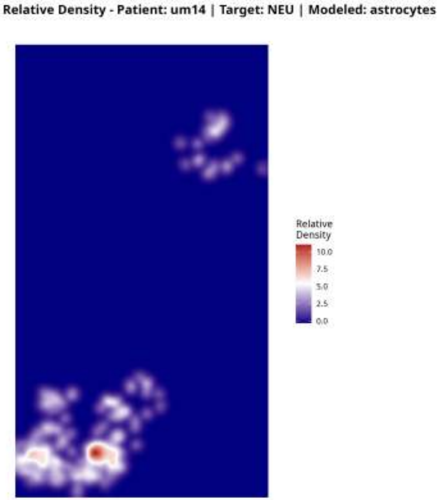
Relative Density - Patient: um14 | Target: NEU | Modeled: astrocytes
Relative
Density
10.0
7.5
5.0
2.5
0.0

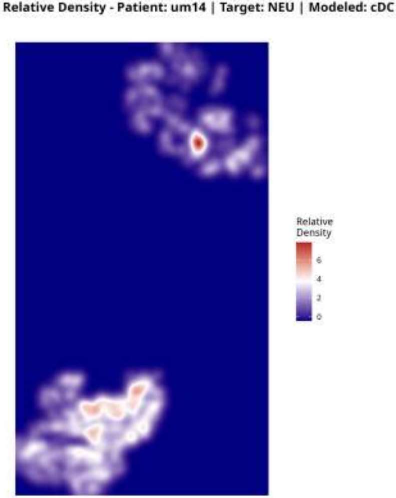

Relative Density - Patient: um14 | Target: NEU | Modeled: cDC
Relative
Density
6
4
2
0

**Relative Density - Patient: um14 | Target: NEU | Modeled: GPM**

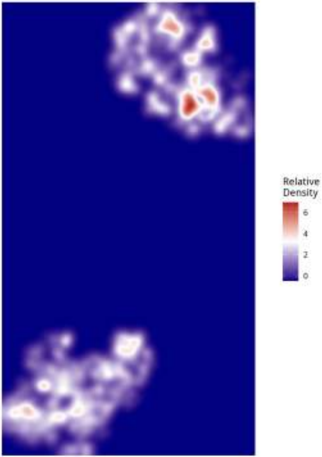

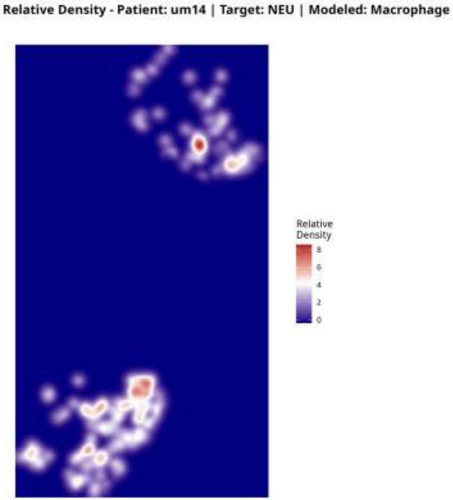

Relative Density - Patient: um14 | Target: NEU | Modeled: Macrophage
Relative Density
8
6
4
2
0

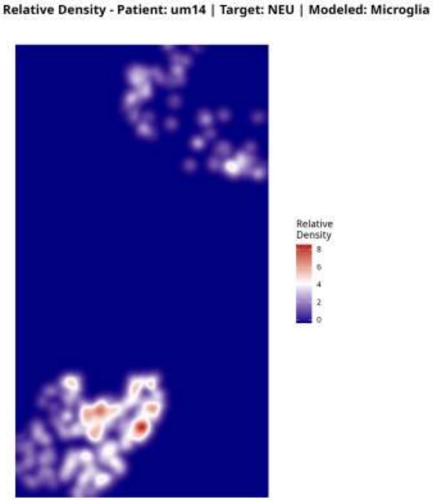
Relative Density - Patient: um14 | Target: NEU | Modeled: Microglia
Relative Density
8
6
4
2
0

Relative Density - Patient: um14 | Target: NEU | Modeled: Monocyte

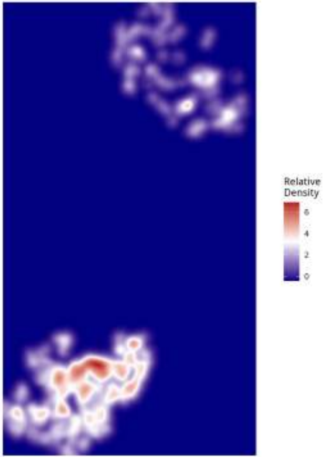

Relative Density - Patient: um14 | Target: NEU | Modeled: MTC

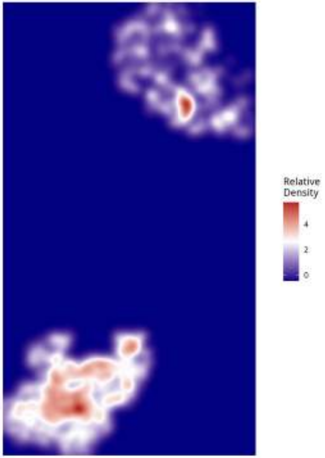

**Relative Density - Patient: um14 | Target: NEU | Modeled: NEU**

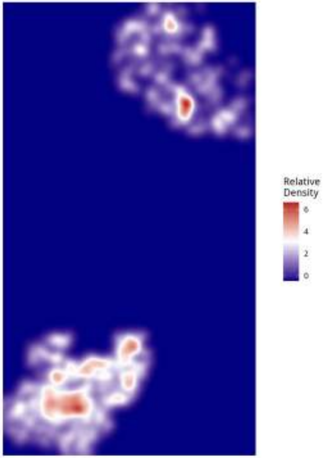

Relative Density - Patient: um14 | Target: NEU | Modeled: neurons

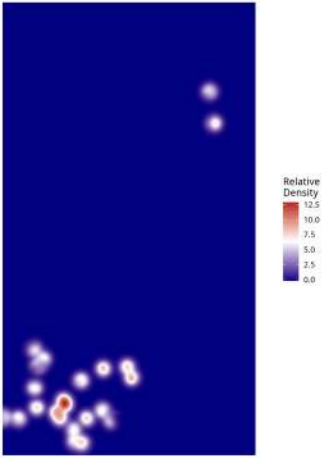

Relative Density - Patient: um14 | Target: NEU | Modeled: Neutrophil

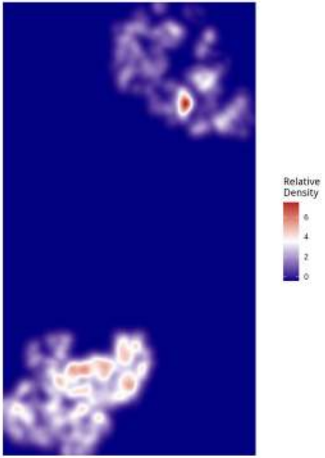

Relative Density - Patient: um14 | Target: NEU | Modeled: oligodendrocyte

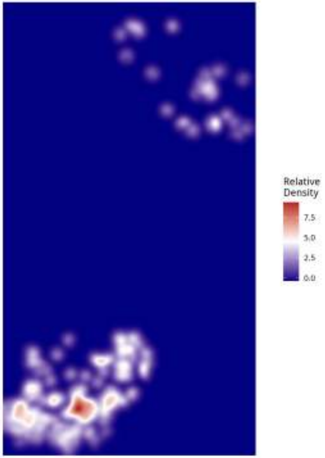

**Relative Density - Patient: um14 | Target: NEU | Modeled: OPC**

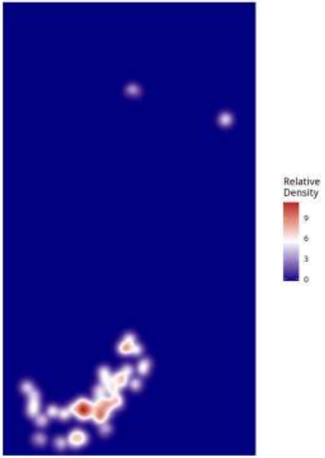

Relative Density - Patient: um14 | Target: NEU | Modeled: PPR

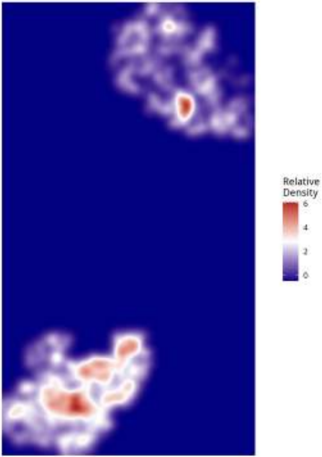

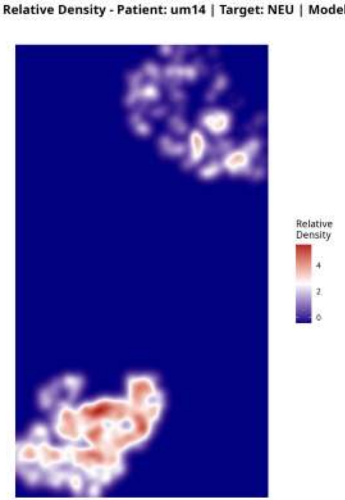
Relative Density - Patient: um14 | Target: NEU | Modeled: Tcells
Relative
Density
4
2
0

Relative Density - Patient: um14 | Target: NEU | Modeled: vasculature

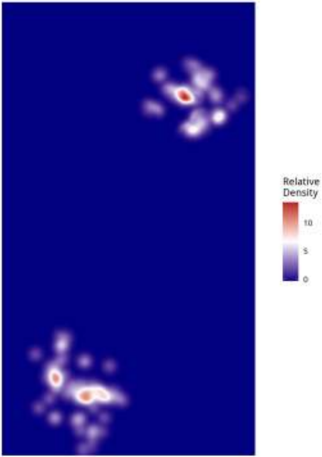

**Relative Density - Patient: um14 | Target: PPR | Modeled: astrocytes**

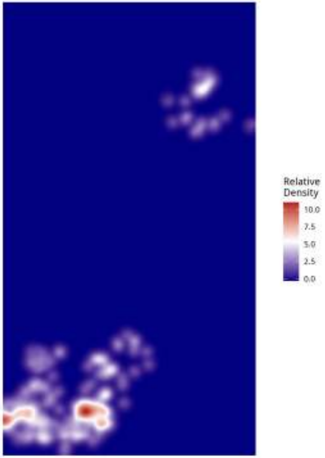

Relative Density - Patient: um14 | Target: PPR | Modeled: cDC

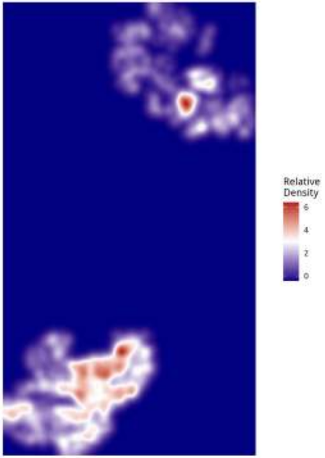

**Relative Density - Patient: um14 | Target: PPR | Modeled: GPM**

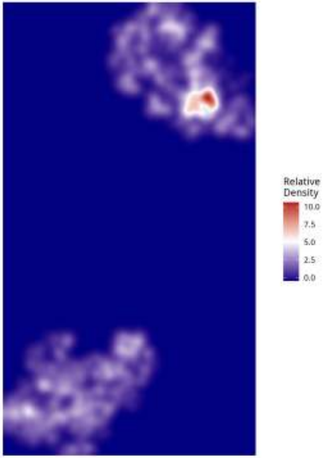

Relative Density - Patient: um14 | Target: PPR | Modeled: Macrophage

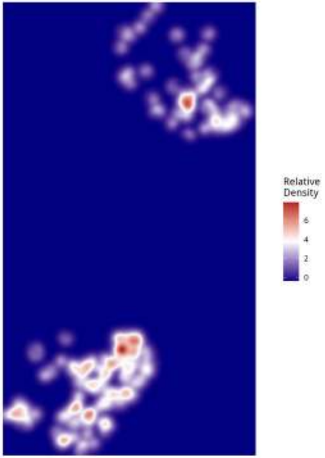

Relative Density - Patient: um14 | Target: PPR | Modeled: Microglia

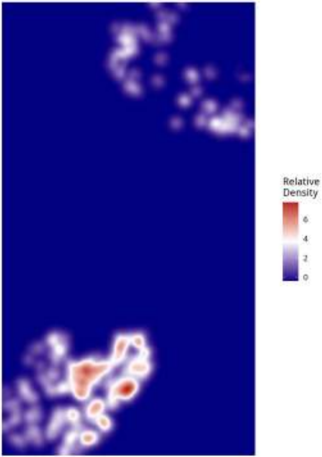

Relative Density - Patient: um14 | Target: PPR | Modeled: Monocyte

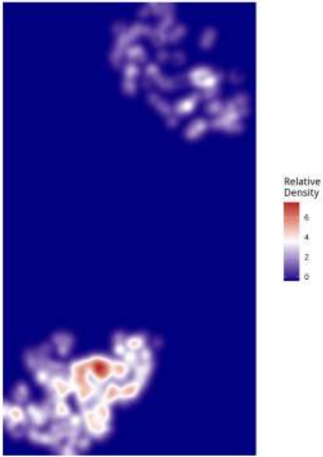

Relative Density - Patient: um14 | Target: PPR | Modeled: MTC

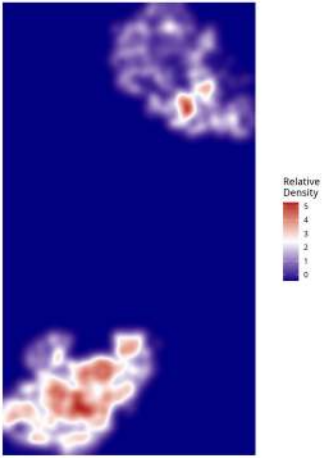

Relative Density - Patient: um14 | Target: PPR | Modeled: NEU

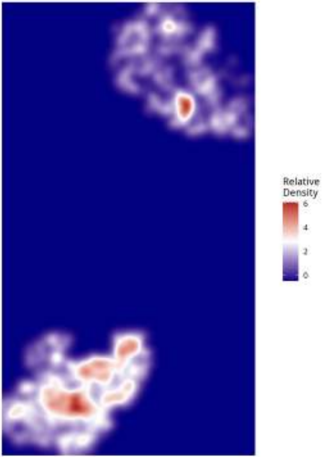

Relative Density - Patient: um14 | Target: PPR | Modeled: neurons

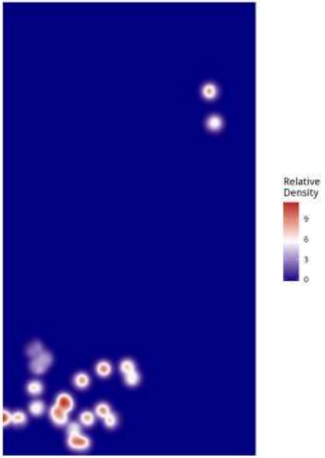

**Relative Density - Patient: um14 | Target: PPR | Modeled: Neutrophil**

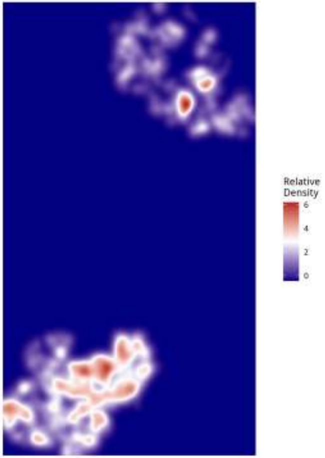

**Relative Density - Patient: um14 | Target: PPR | Modeled: oligodendrocyte**

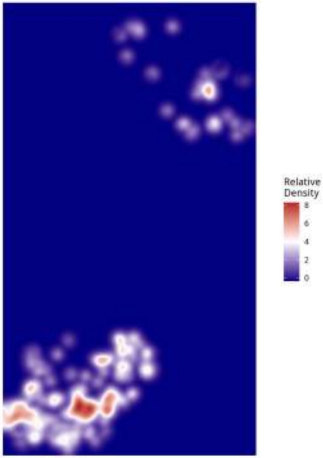

Relative Density - Patient: um14 | Target: PPR | Modeled: OPC

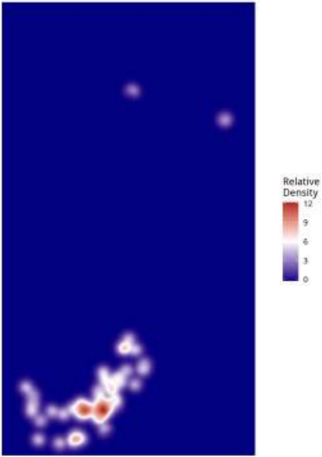

**Relative Density - Patient: um14 | Target: PPR | Modeled: PPR**

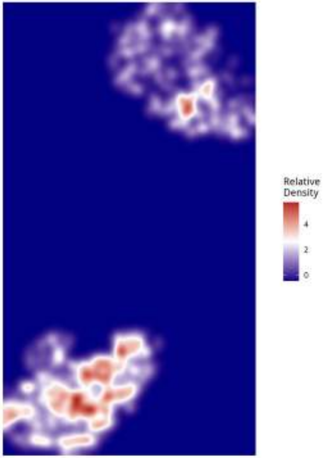

Relative Density - Patient: um14 | Target: PPR | Modeled: Tcells

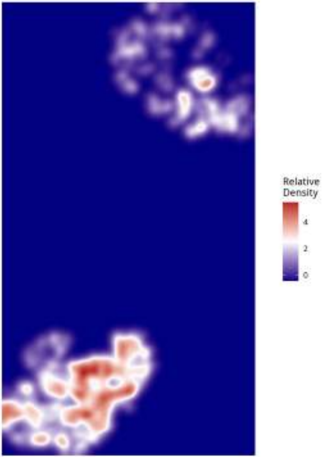

Relative Density - Patient: um14 | Target: PPR | Modeled: vasculature

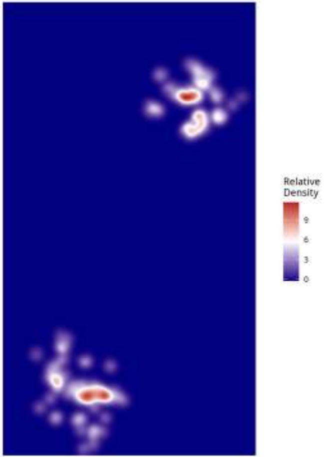

Relative Density - Patient: um15 | Target: GPM | Modeled: astrocytes

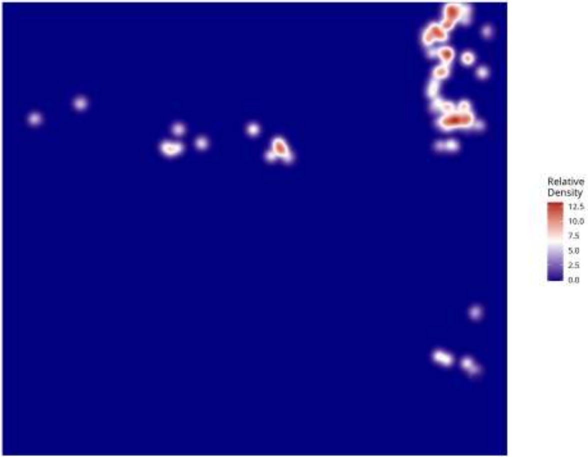

**Relative Density - Patient: um15 | Target: GPM | Modeled: cDC**

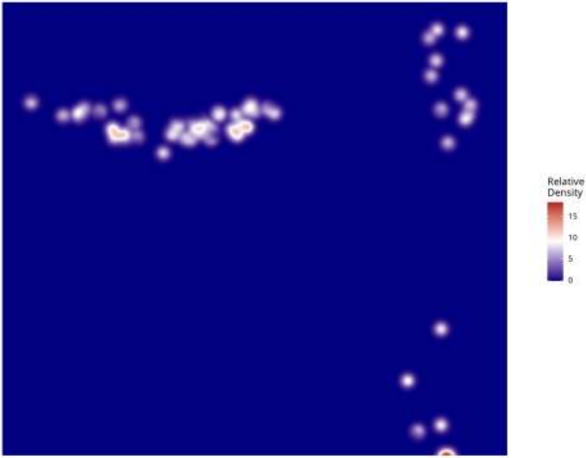

Relative Density - Patient: um15 | Target: GPM | Modeled: GPM

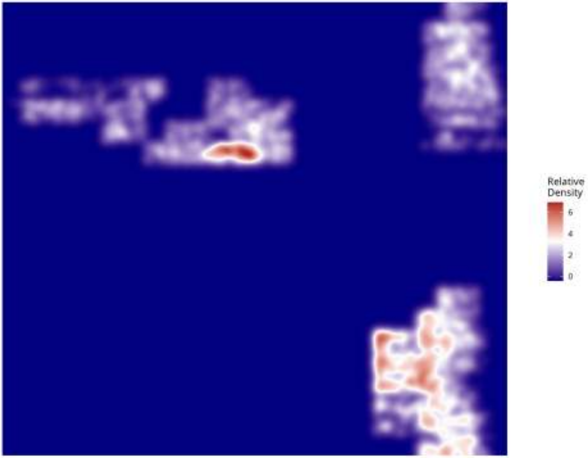

**Relative Density - Patient: um15 | Target: GPM | Modeled: Macrophage**

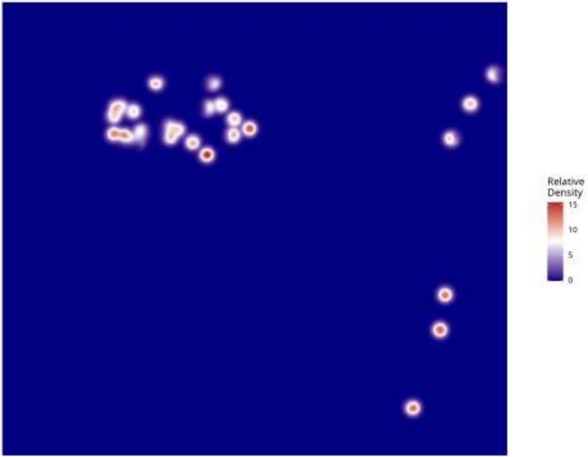

**Relative Density - Patient: um15 | Target: GPM | Modeled: Monocyte**

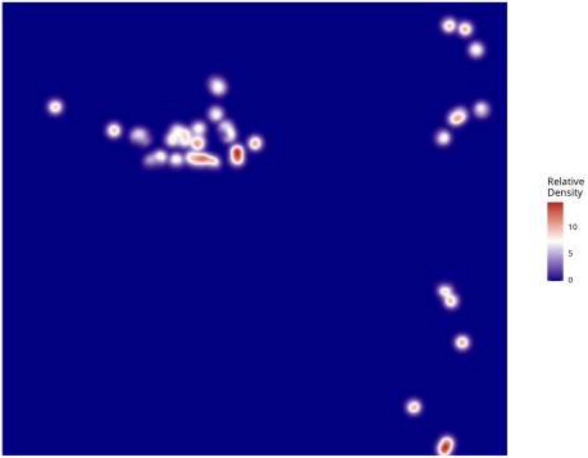

**Relative Density - Patient: um15 | Target: GPM | Modeled: MTC**

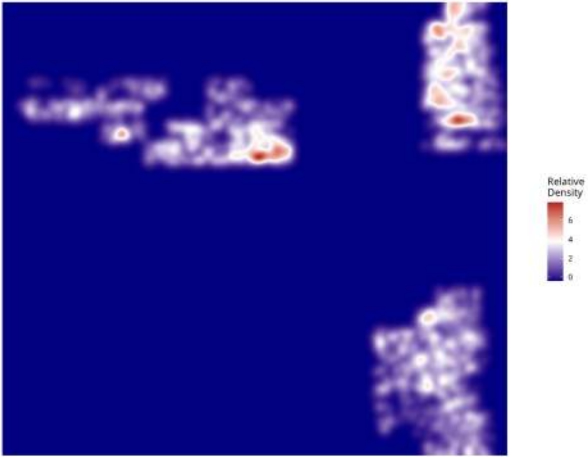

Relative Density - Patient: um15 | Target: GPM | Modeled: NEU

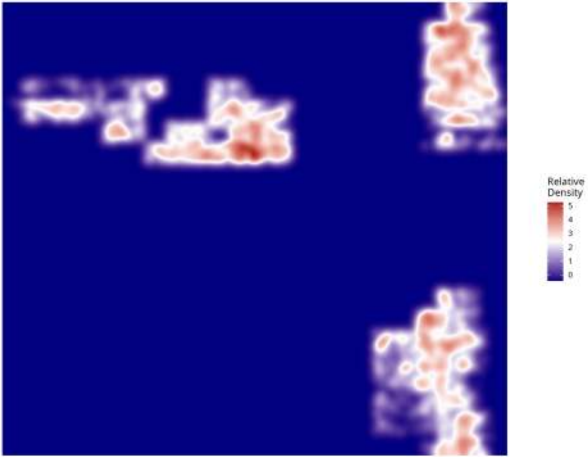

Relative Density - Patient: um15 | Target: GPM | Modeled: neurons

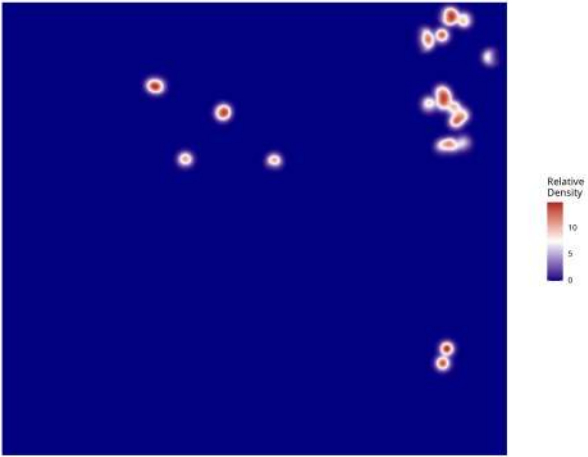

Relative Density - Patient: um15 | Target: GPM | Modeled: Neutrophil

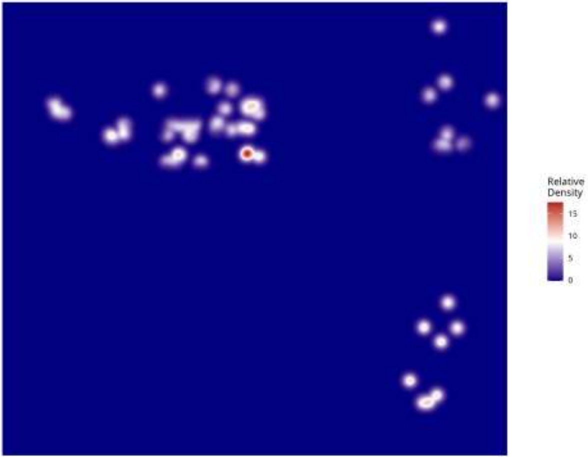

Relative Density - Patient: um15 | Target: GPM | Modeled: oligodendrocyte

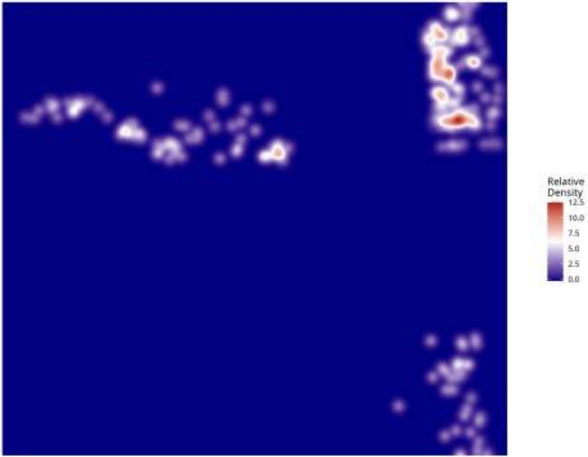

**Relative Density - Patient: um15 | Target: GPM | Modeled: OPC**

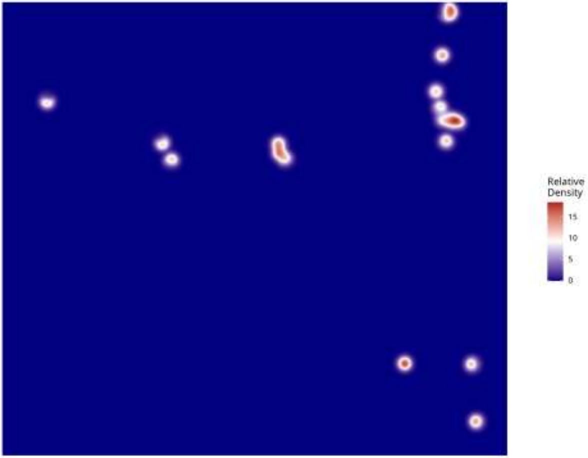

**Relative Density - Patient: um15 | Target: GPM | Modeled: PPR**

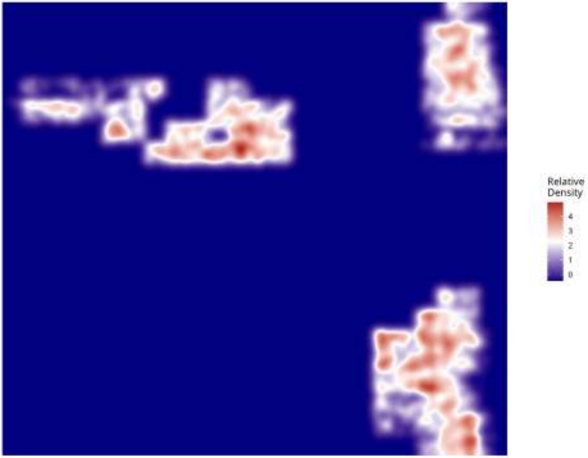

**Relative Density - Patient: um15 | Target: GPM | Modeled: Tcells**

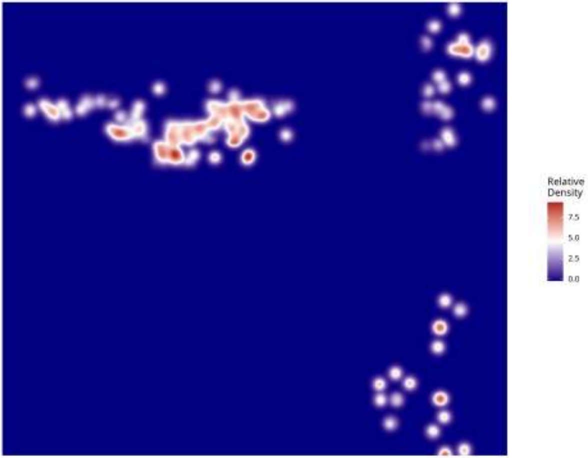

**Relative Density - Patient: um15 | Target: GPM | Modeled: vasculature**

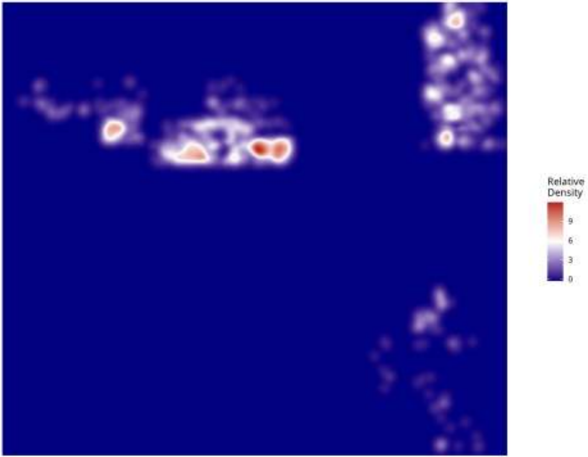

**Relative Density - Patient: um15 | Target: MTC | Modeled: astrocytes**

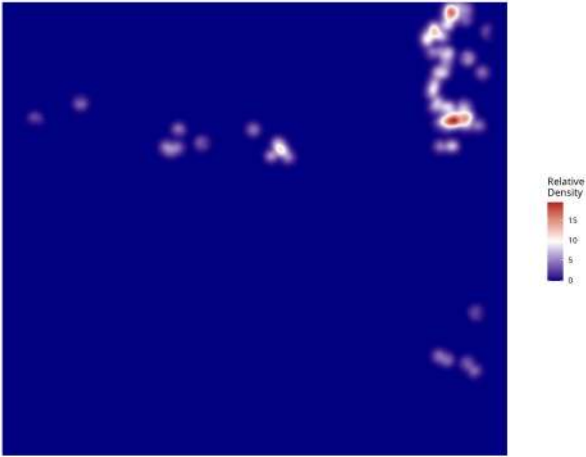

**Relative Density - Patient: um15 | Target: MTC | Modeled: cDC**

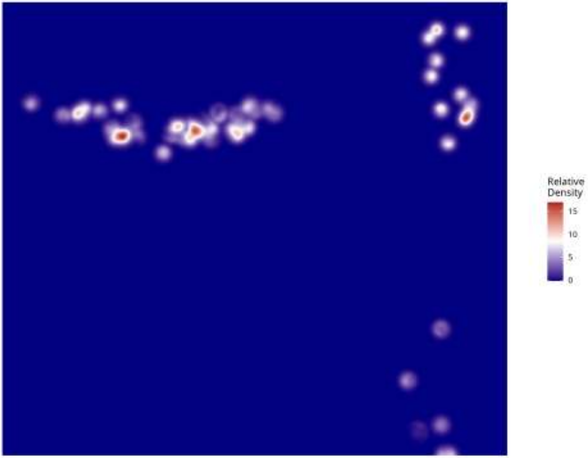

**Relative Density - Patient: um15 | Target: MTC | Modeled: GPM**

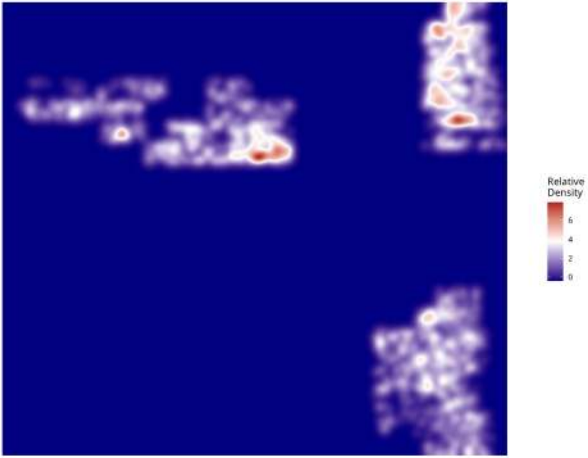

Relative Density - Patient: um15 | Target: MTC | Modeled: Macrophage

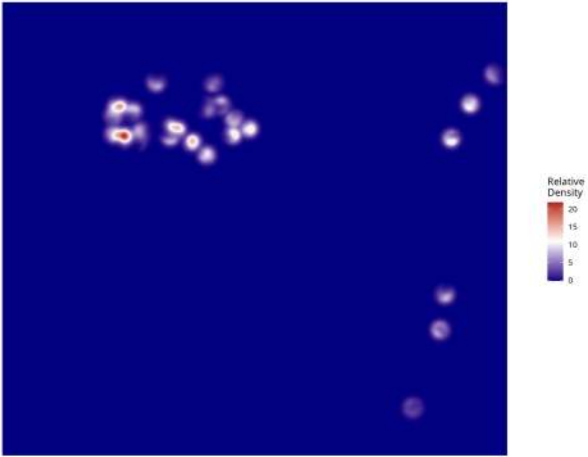

**Relative Density - Patient: um15 | Target: MTC | Modeled: Monocyte**

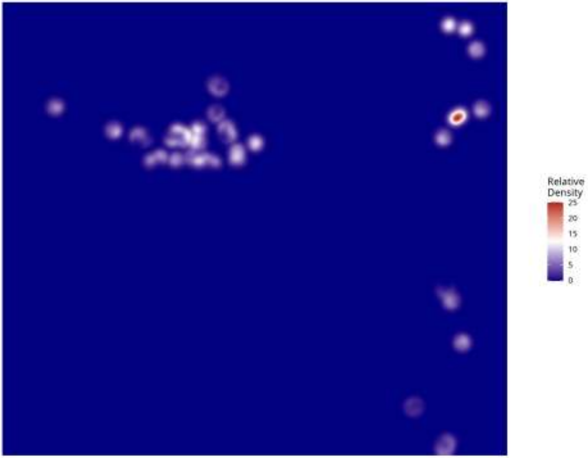

Relative Density - Patient: um15 | Target: MTC | Modeled: MTC

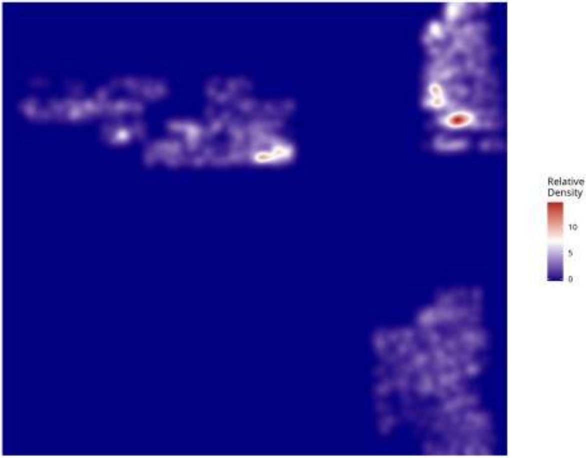

Relative Density - Patient: um15 | Target: MTC | Modeled: NEU

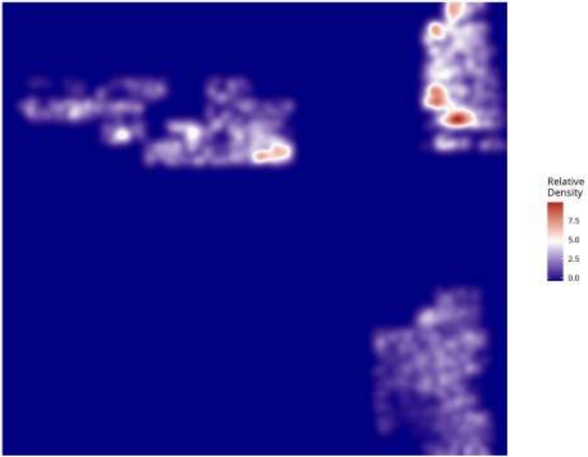

**Relative Density - Patient: um15 | Target: MTC | Modeled: neurons**

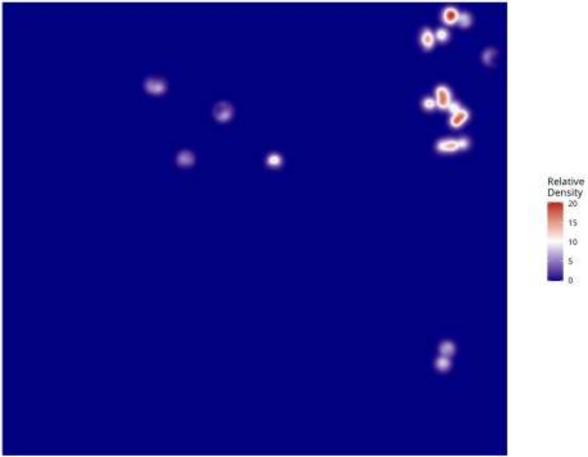

**Relative Density - Patient: um15 | Target: MTC | Modeled: Neutrophil**

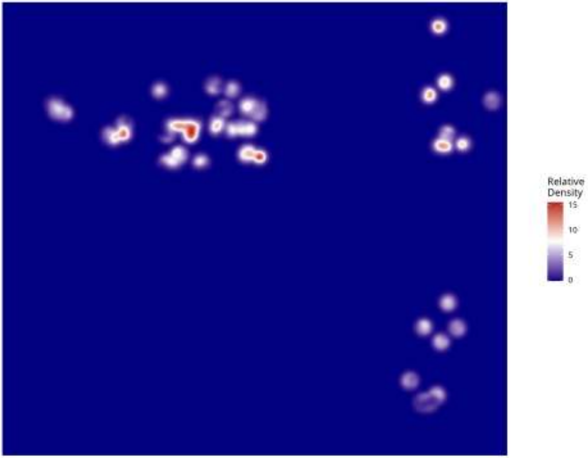

**Relative Density - Patient: um15 | Target: MTC | Modeled: oligodendrocyte**

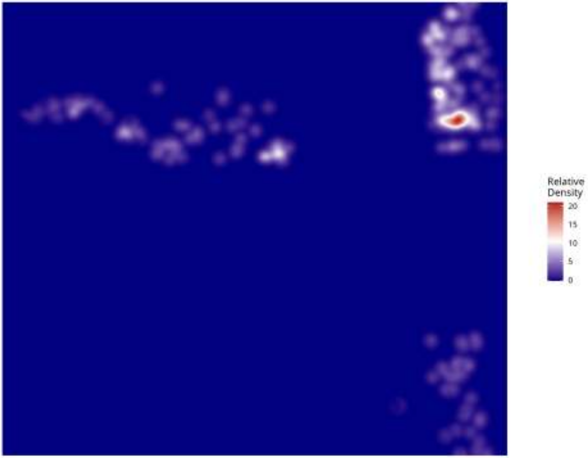

Relative Density - Patient: um15 | Target: MTC | Modeled: OPC

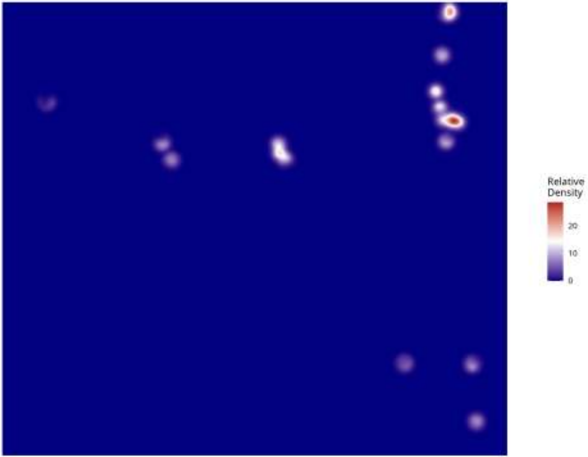

**Relative Density - Patient: um15 | Target: MTC | Modeled: PPR**

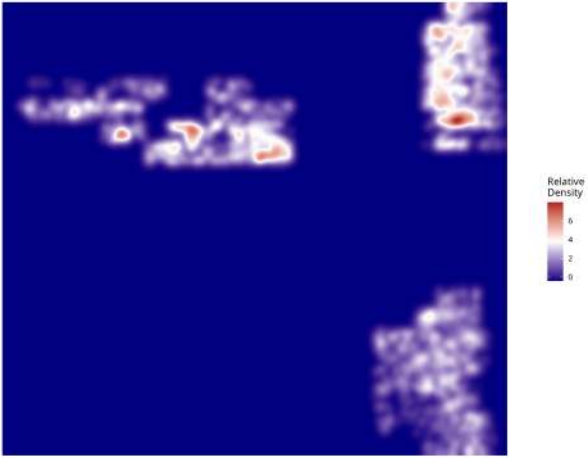

**Relative Density - Patient: um15 | Target: MTC | Modeled: Tcells**

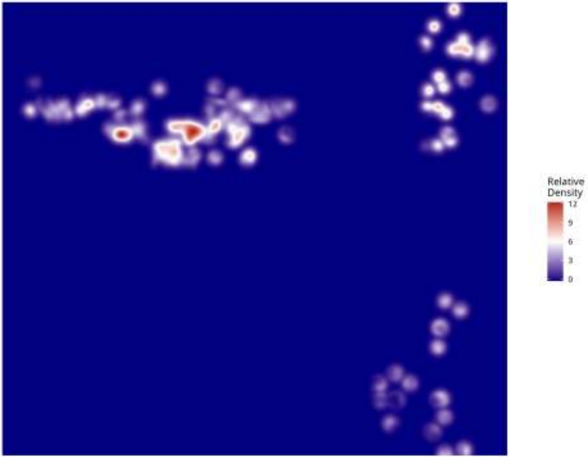

Relative Density - Patient: um15 | Target: MTC | Modeled: vasculature

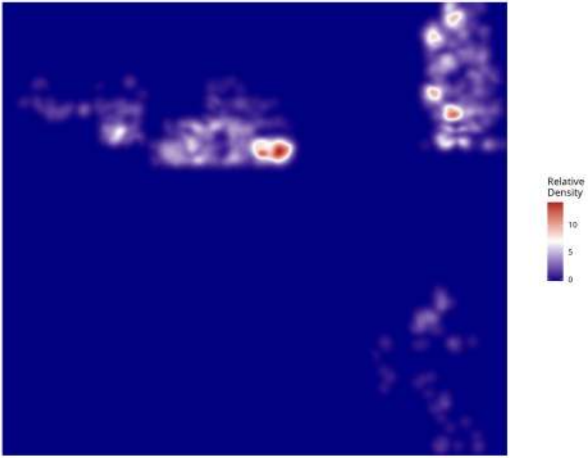

Relative Density - Patient: um15 | Target: NEU | Modeled: astrocytes

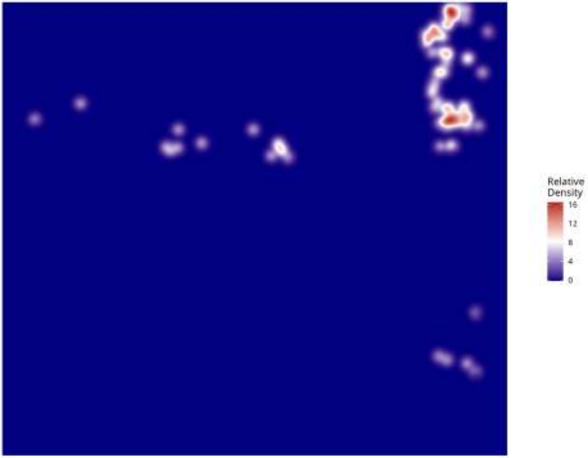

**Relative Density - Patient: um15 | Target: NEU | Modeled: cDC**

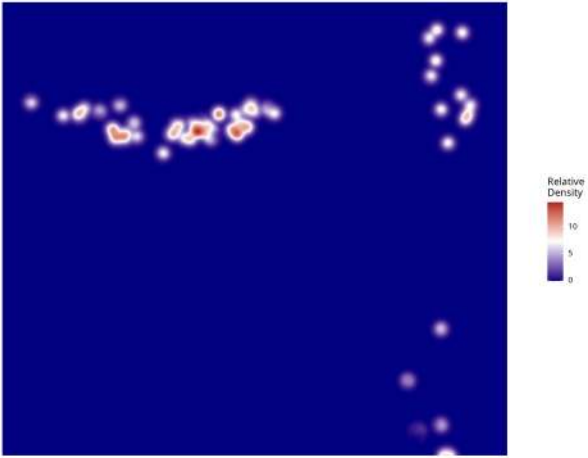

**Relative Density - Patient: um15 | Target: NEU | Modeled: GPM**

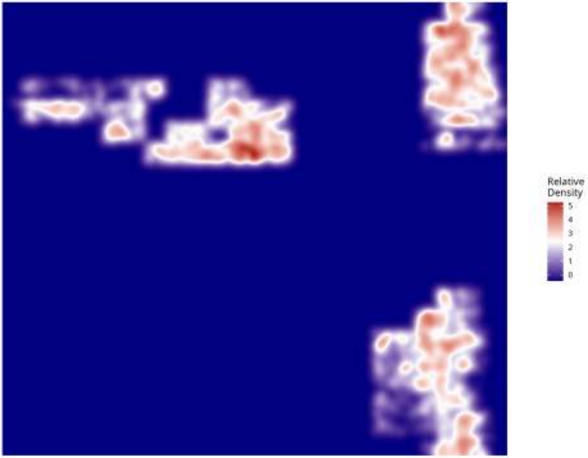

**Relative Density - Patient: um15 | Target: NEU | Modeled: Macrophage**

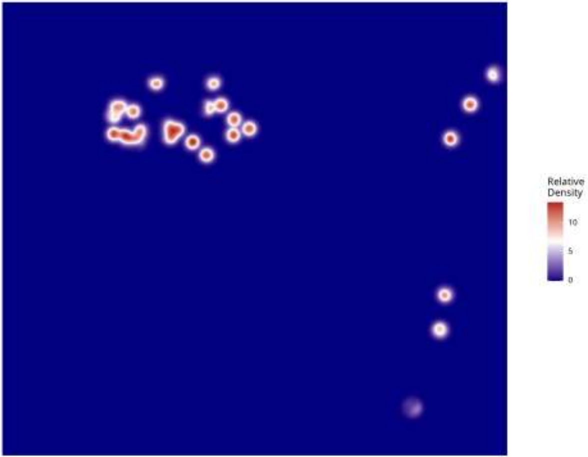

**Relative Density - Patient: um15 | Target: NEU | Modeled: Monocyte**

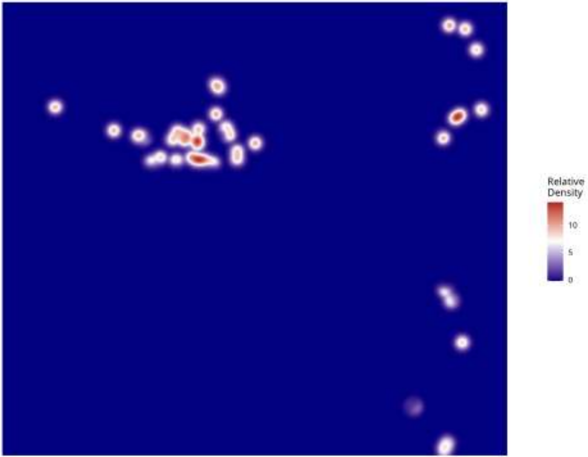

Relative Density - Patient: um15 | Target: NEU | Modeled: MTC

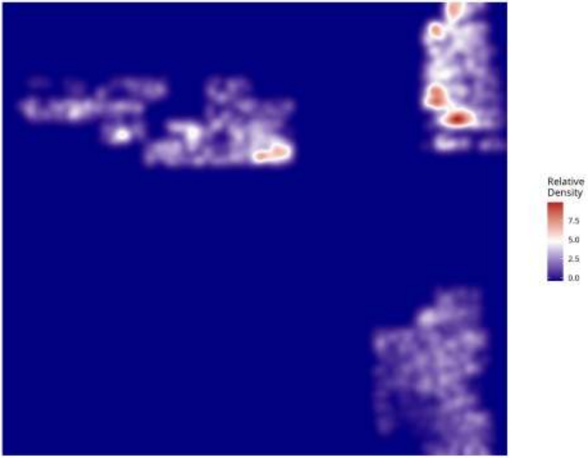

**Relative Density - Patient: um15 | Target: NEU | Modeled: NEU**

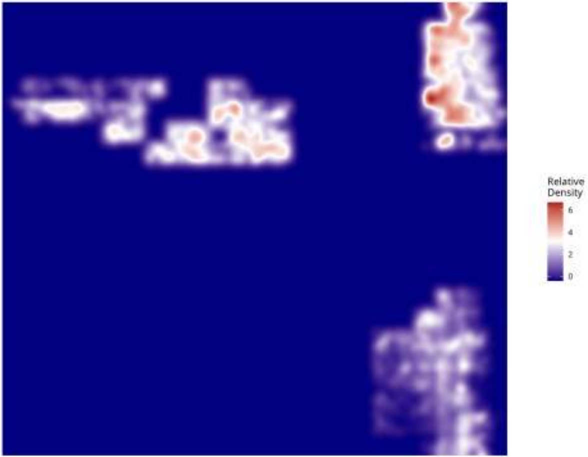

Relative Density - Patient: um15 | Target: NEU | Modeled: neurons

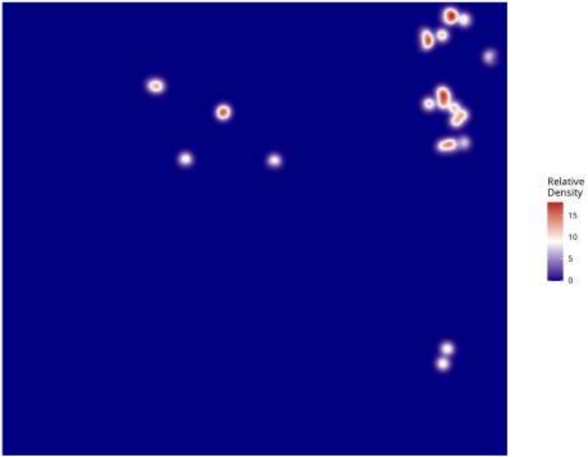

**Relative Density - Patient: um15 | Target: NEU | Modeled: Neutrophil**

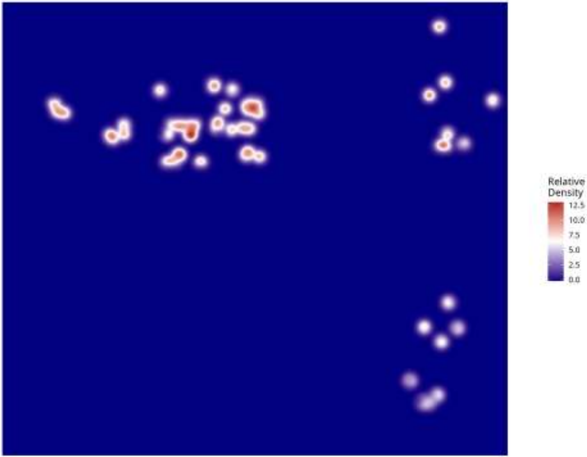

**Relative Density - Patient: um15 | Target: NEU | Modeled: oligodendrocyte**

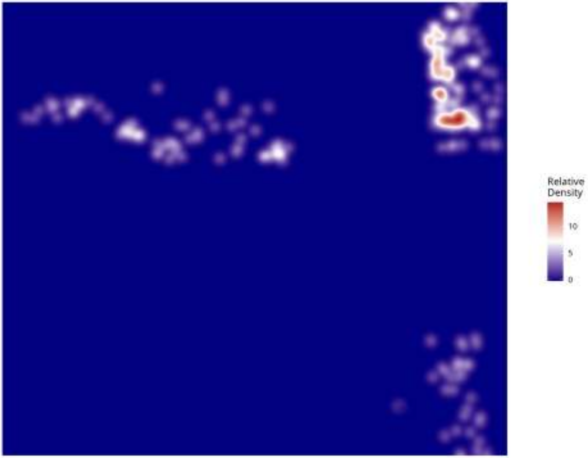

**Relative Density - Patient: um15 | Target: NEU | Modeled: OPC**

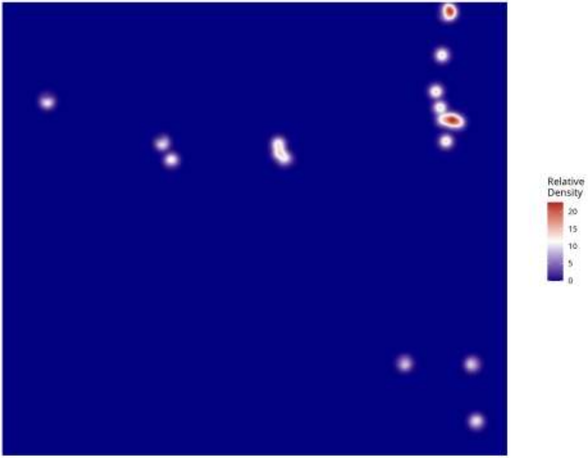

Relative Density - Patient: um15 | Target: NEU | Modeled: PPR

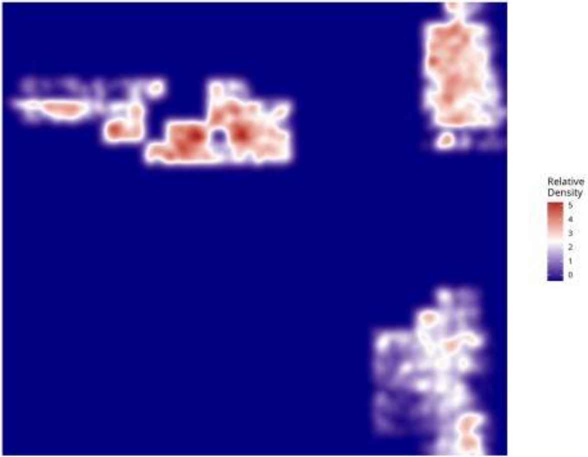

**Relative Density - Patient: um15 | Target: NEU | Modeled: Tcells**

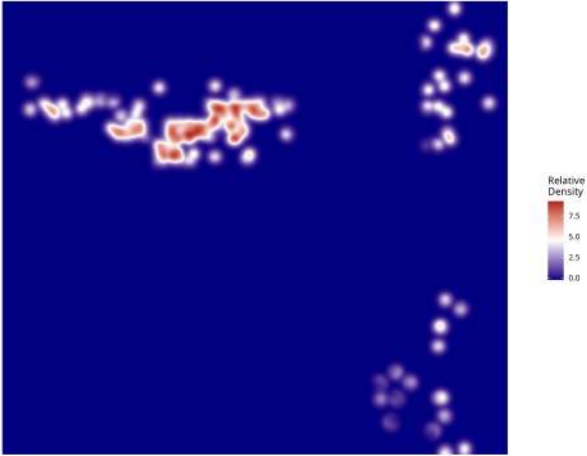

**Relative Density - Patient: um15 | Target: NEU | Modeled: vasculature**

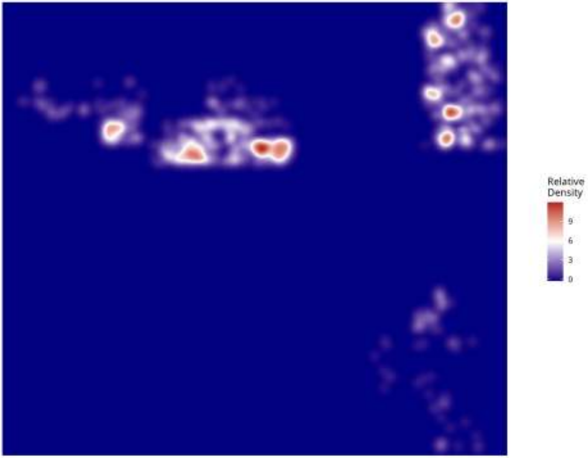

Relative Density - Patient: um15 | Target: PPR | Modeled: astrocytes

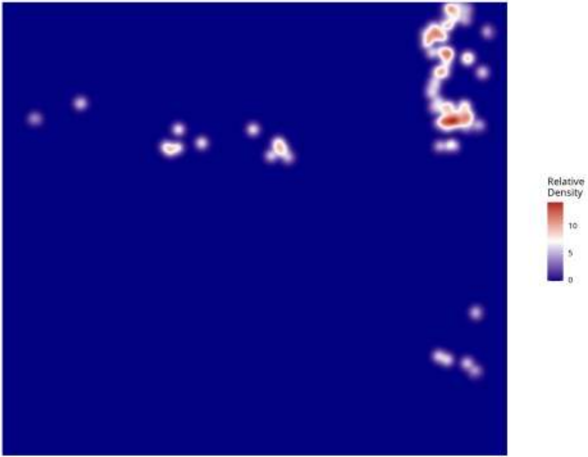

**Relative Density - Patient: um15 | Target: PPR | Modeled: cDC**

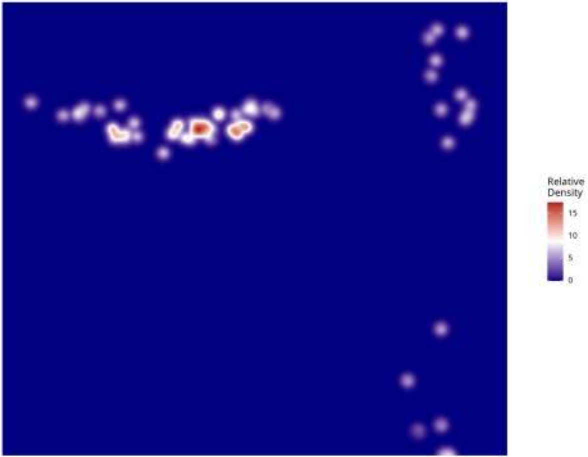

**Relative Density - Patient: um15 | Target: PPR | Modeled: GPM**

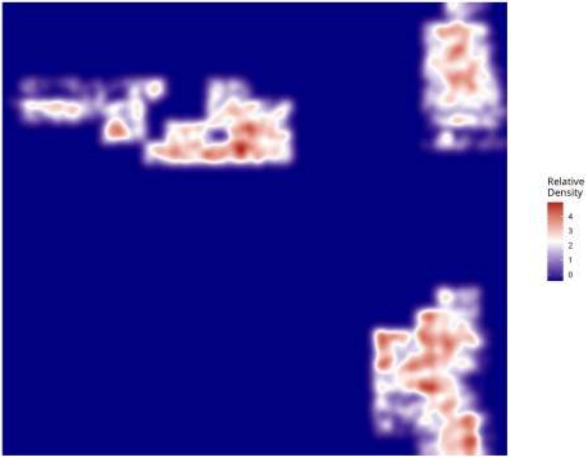

**Relative Density - Patient: um15 | Target: PPR | Modeled: Macrophage**

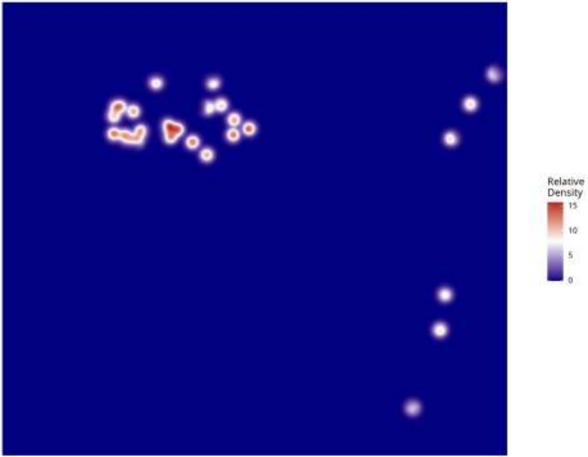

**Relative Density - Patient: um15 | Target: PPR | Modeled: Monocyte**

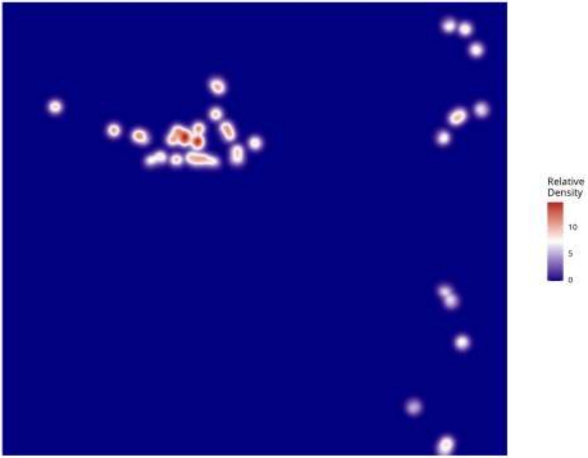

Relative Density - Patient: um15 | Target: PPR | Modeled: MTC

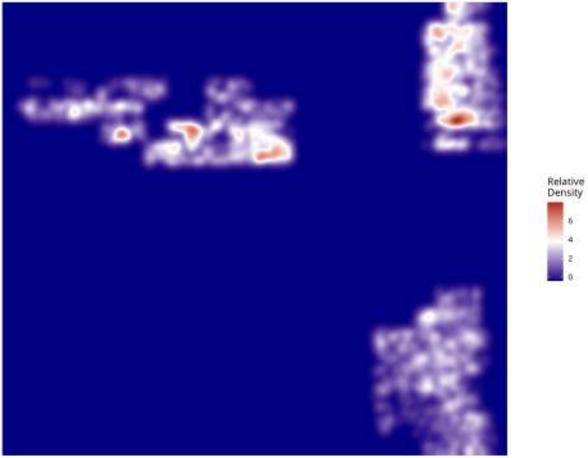

Relative Density - Patient: um15 | Target: PPR | Modeled: NEU

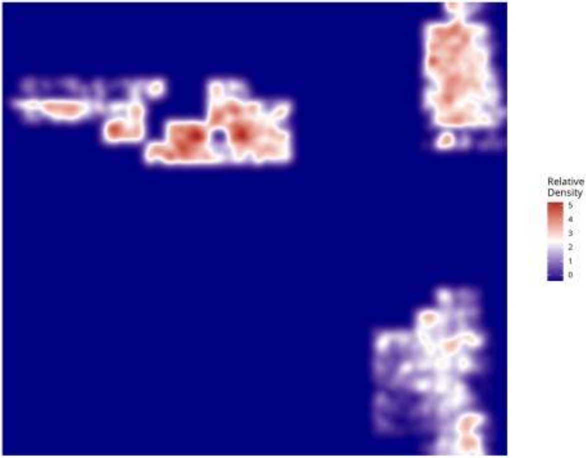

Relative Density - Patient: um15 | Target: PPR | Modeled: neurons

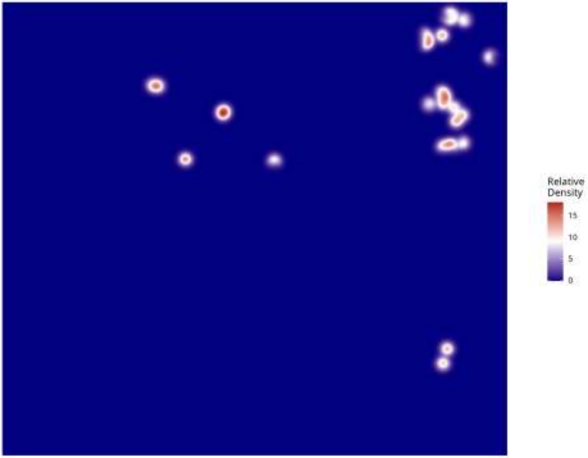

**Relative Density - Patient: um15 | Target: PPR | Modeled: Neutrophil**

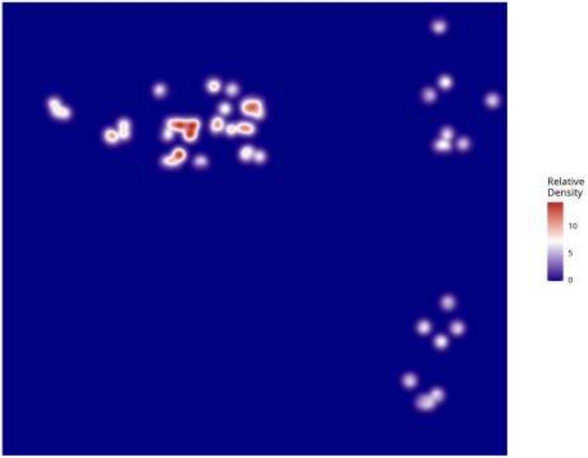

**Relative Density - Patient: um15 | Target: PPR | Modeled: oligodendrocyte**

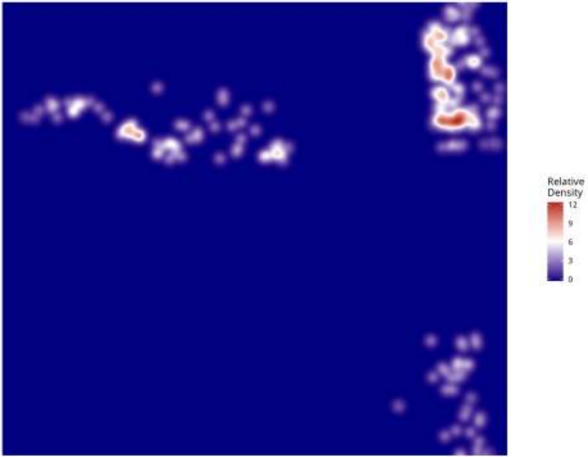

**Relative Density - Patient: um15 | Target: PPR | Modeled: OPC**

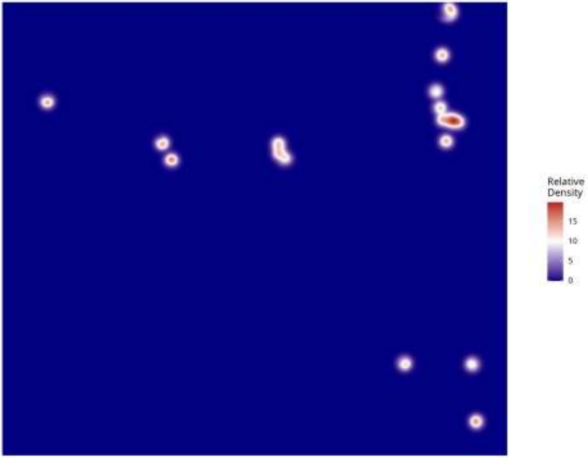

**Relative Density - Patient: um15 | Target: PPR | Modeled: PPR**

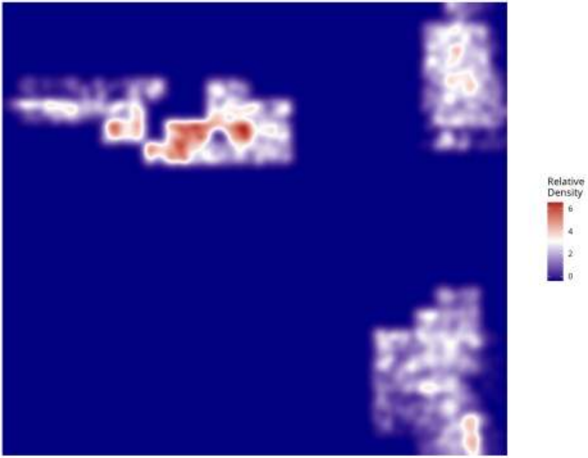

**Relative Density - Patient: um15 | Target: PPR | Modeled: Tcells**

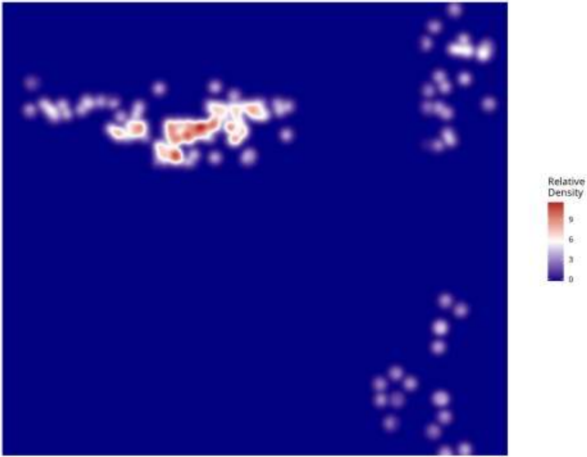

**Relative Density - Patient: um15 | Target: PPR | Modeled: vasculature**

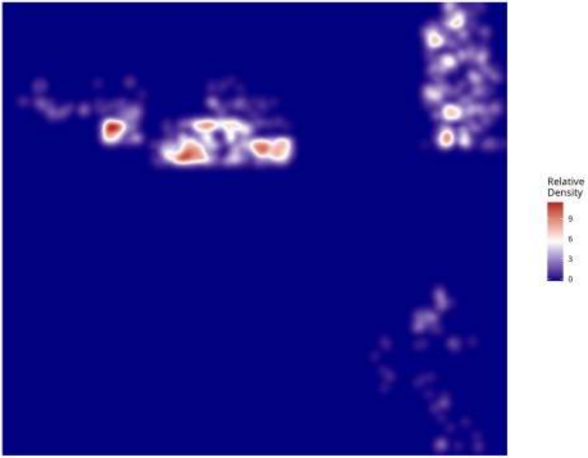
