## Supplementary Figures 5 for "Point-Process Modelling of Cell-Type Interaction Structure in CosMx Glioma Data for Hypothesis-Driven Tumor Inference from scRNA-seq"

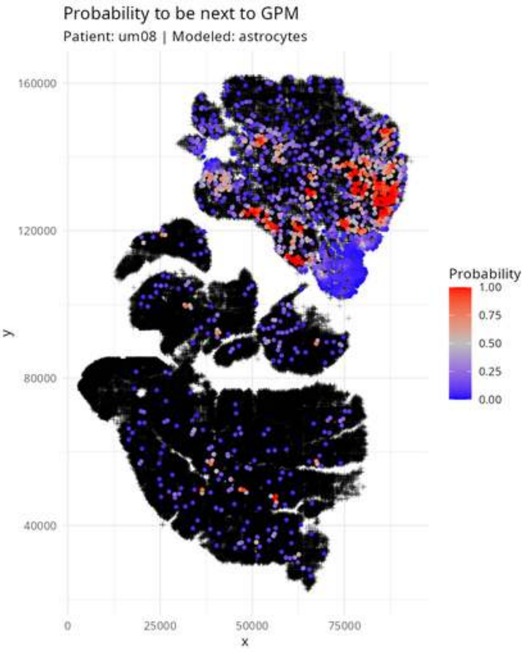
Probability to be next to GPM
Patient: um08 | Modeled: astrocytes
160000
120000
80000
40000
y
0
25000
50000
75000
x
Probability
1.00
0.75
0.50
0.25
0.00

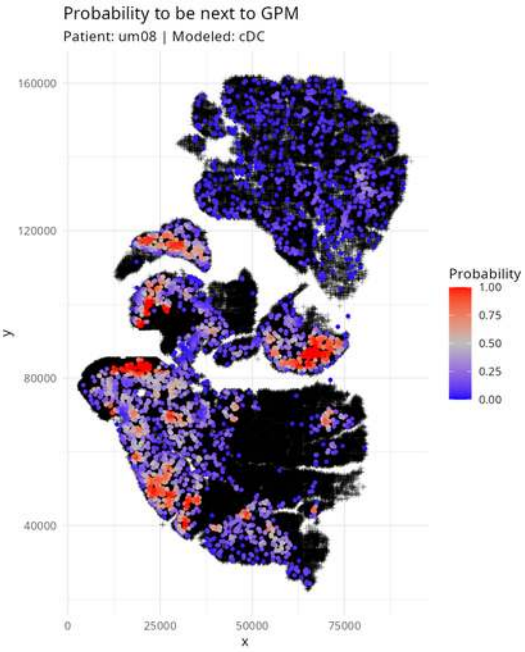
Probability to be next to GPM
Patient: um08 | Modeled: cDC
y
160000
120000
80000
40000
x
0
25000
50000
75000
Probability
1.00
0.75
0.50
0.25
0.00

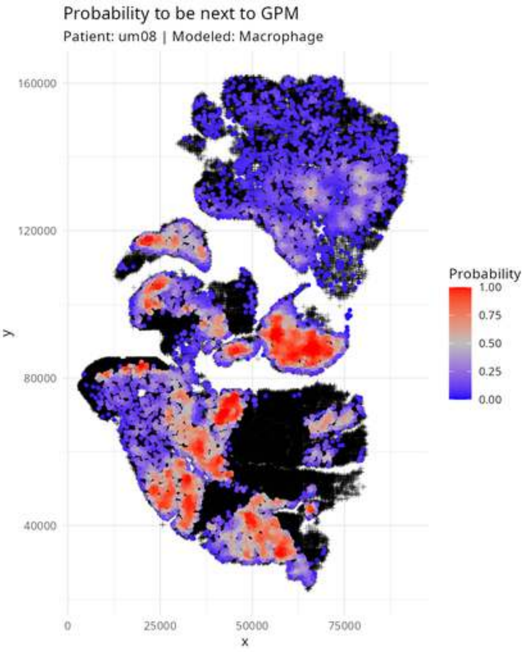

Probability to be next to GPM
Patient: um08 | Modeled: Macrophage
160000
120000
80000
40000
y
0
25000
50000
75000
x
Probability
1.00
0.75
0.50
0.25
0.00

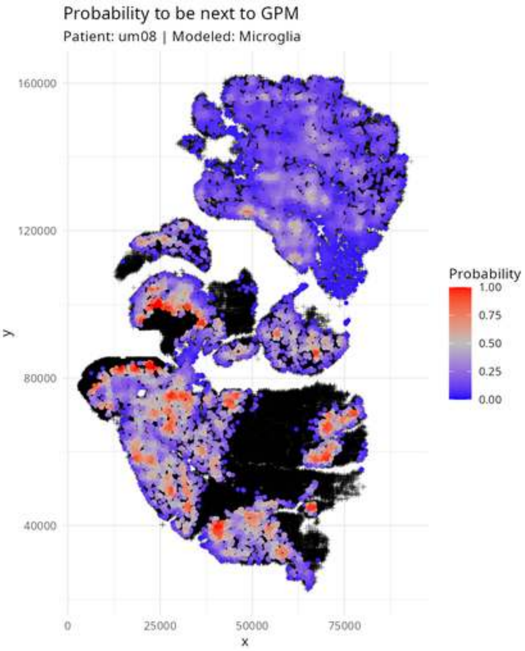
Probability to be next to GPM
Patient: um08 | Modeled: Microglia
160000
120000
80000
40000
y
0
25000
50000
75000
x
Probability
1.00
0.75
0.50
0.25
0.00

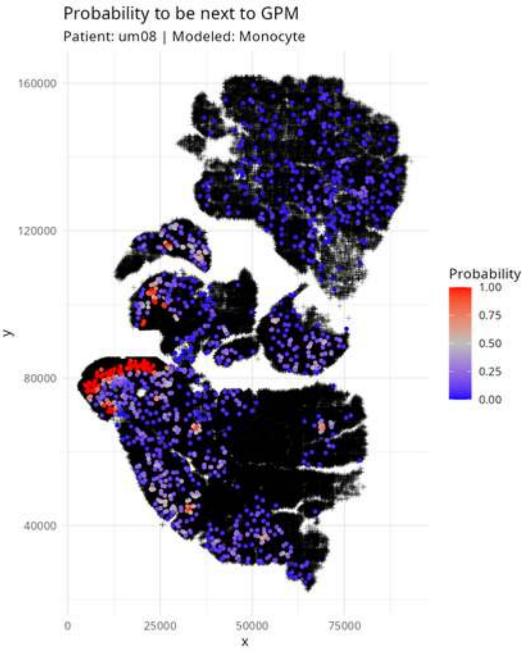
Probability to be next to GPM
Patient: um08 | Modeled: Monocyte
160000
120000
80000
40000
y
0
25000
50000
75000
x
Probability
1.00
0.75
0.50
0.25
0.00

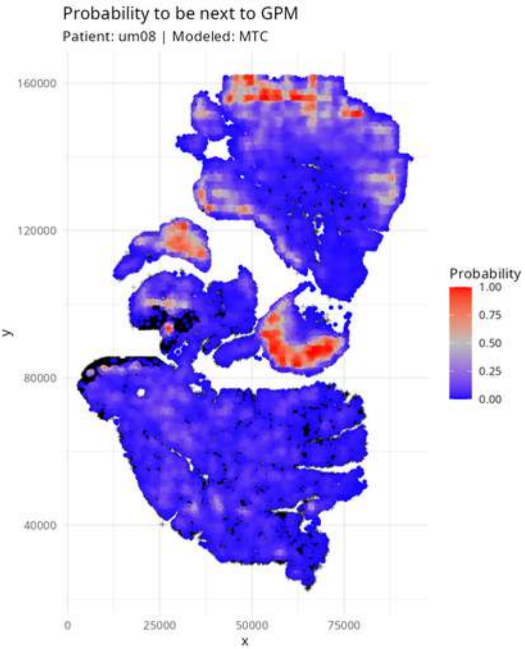
Probability to be next to GPM
Patient: um08 | Modeled: MTC
160000
120000
80000
40000
y
0
25000
50000
75000
x
Probability
1.00
0.75
0.50
0.25
0.00

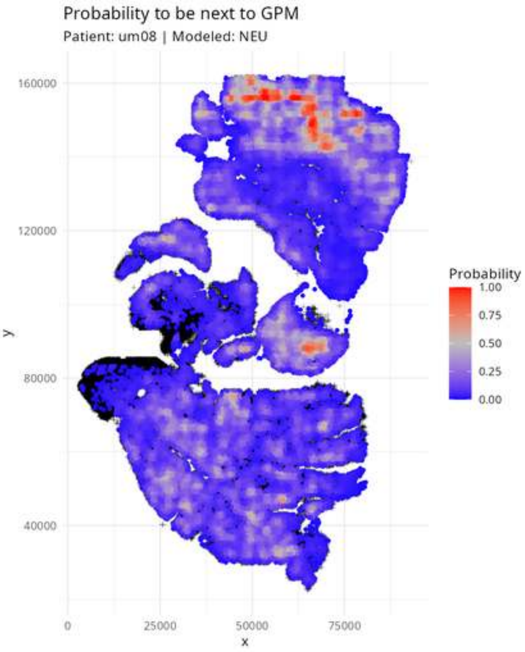
Probability to be next to GPM
Patient: um08 | Modeled: NEU
160000
120000
80000
40000
y
0
25000
50000
75000
x
Probability
1.00
0.75
0.50
0.25
0.00

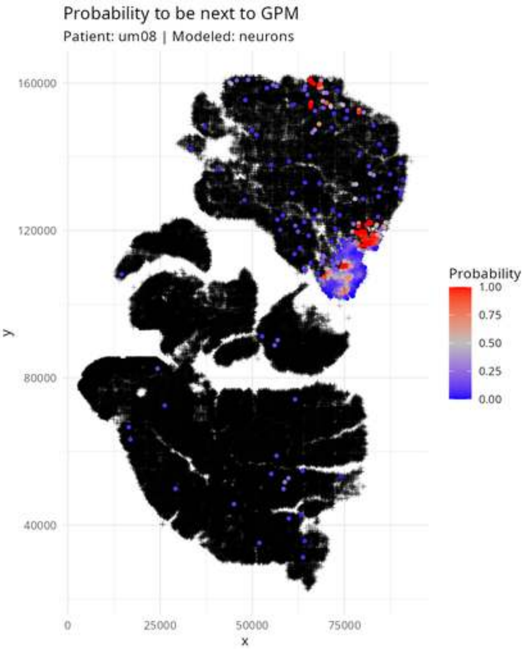

Probability to be next to GPM
Patient: um08 | Modeled: neurons
160000
120000
80000
40000
y
0
25000
50000
75000
x
Probability
1.00
0.75
0.50
0.25
0.00

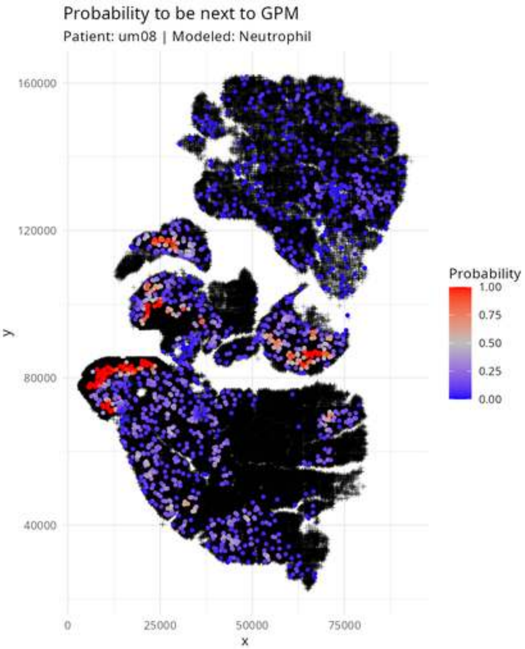

Probability to be next to GPM
Patient: um08 | Modeled: Neutrophil
160000
120000
80000
40000
y
0
25000
50000
75000
x
Probability
1.00
0.75
0.50
0.25
0.00

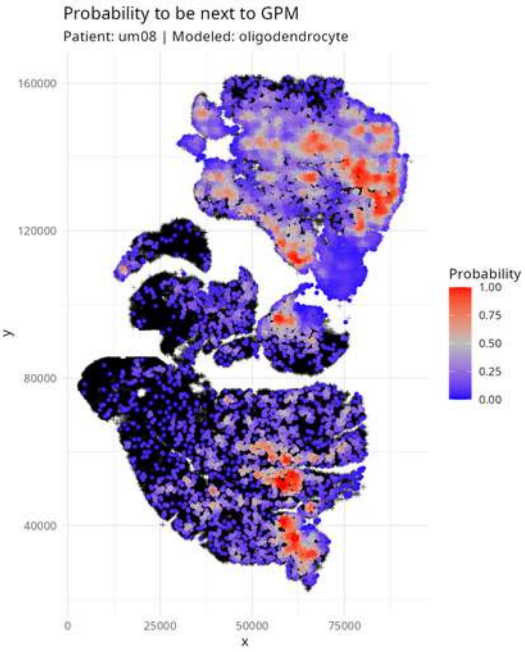

Probability to be next to GPM
Patient: um08 | Modeled: oligodendrocyte
160000
120000
80000
40000
y
0
25000
50000
75000
x
Probability
1.00
0.75
0.50
0.25
0.00

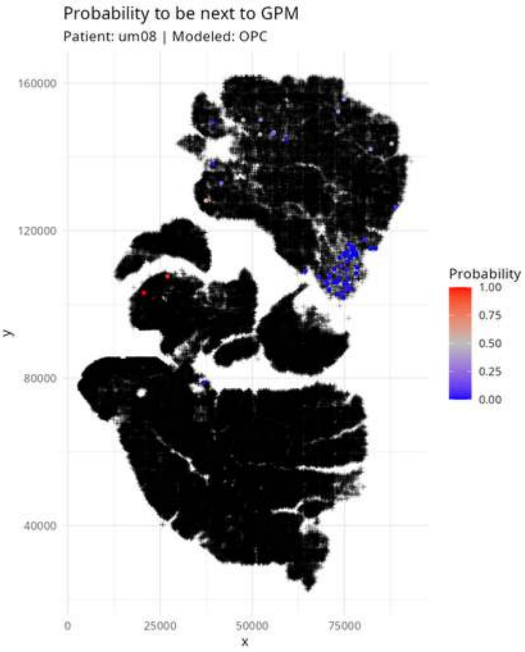

Probability to be next to GPM
Patient: um08 | Modeled: OPC
160000
120000
80000
40000
y
0
25000
50000
75000
x
Probability
1.00
0.75
0.50
0.25
0.00

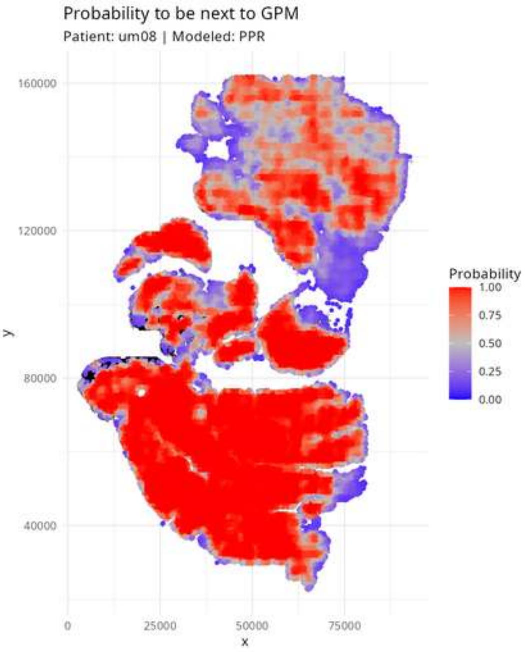
Probability to be next to GPM
Patient: um08 | Modeled: PPR
160000
120000
80000
40000
y
0
25000
50000
75000
x
Probability
1.00
0.75
0.50
0.25
0.00

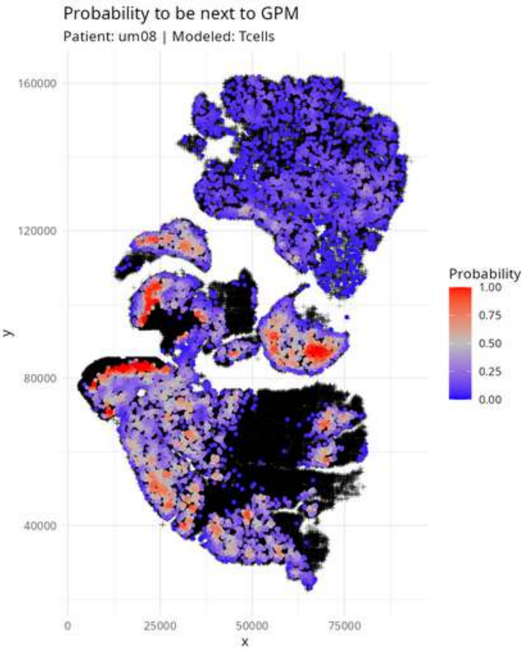
Probability to be next to GPM
Patient: um08 | Modeled: Tcells
y
160000
120000
80000
40000
x
0
25000
50000
75000
Probability
1.00
0.75
0.50
0.25
0.00

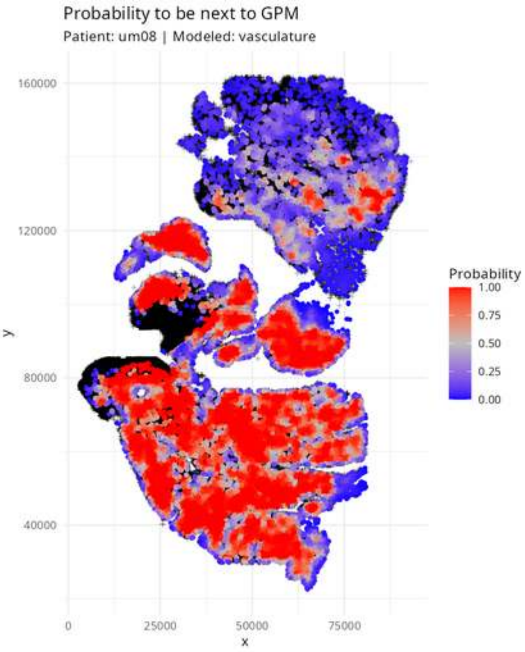

Probability to be next to GPM
Patient: um08 | Modeled: vasculature
160000
120000
80000
40000
y
0
25000
50000
75000
x
Probability
1.00
0.75
0.50
0.25
0.00

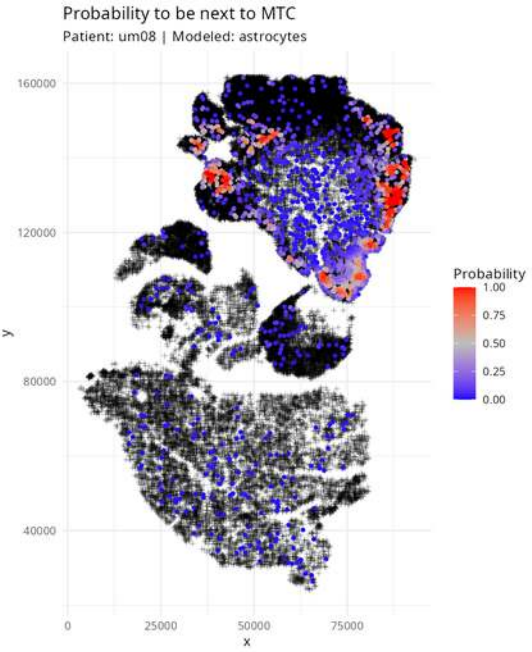
Probability to be next to MTC
Patient: um08 | Modeled: astrocytes
160000
120000
80000
40000
y
0
25000
50000
75000
x
Probability
1.00
0.75
0.50
0.25
0.00

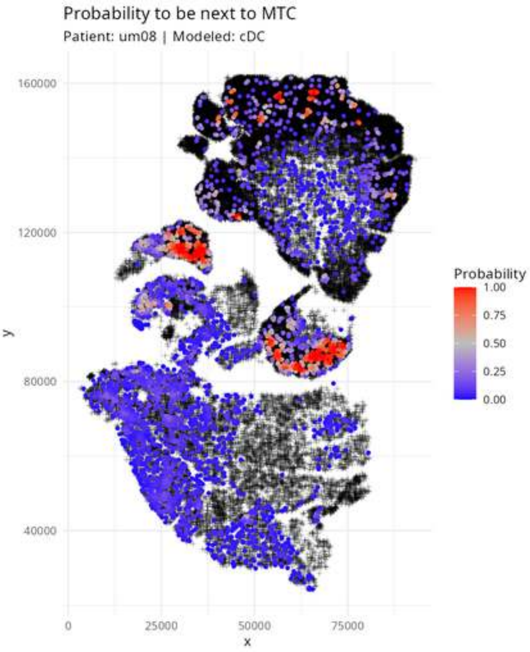

Probability to be next to MTC
Patient: um08 | Modeled: cDC
160000
120000
80000
40000
y
0
25000
50000
75000
x
Probability
1.00
0.75
0.50
0.25
0.00

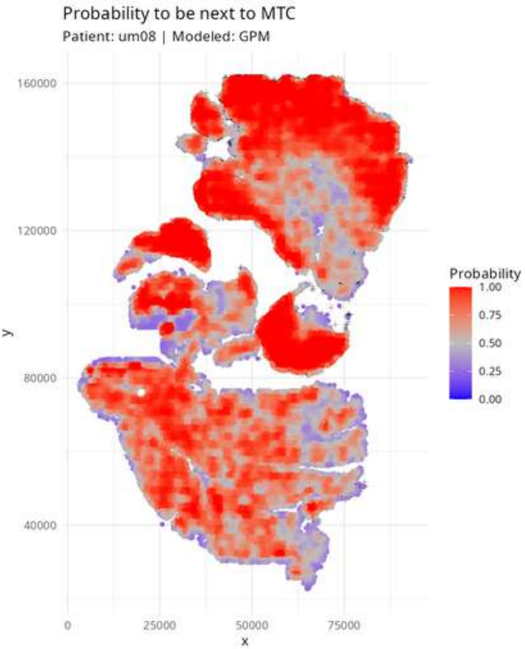

Probability to be next to MTC
Patient: um08 | Modeled: GPM
160000
120000
80000
40000
y
0
25000
50000
75000
x
Probability
1.00
0.75
0.50
0.25
0.00

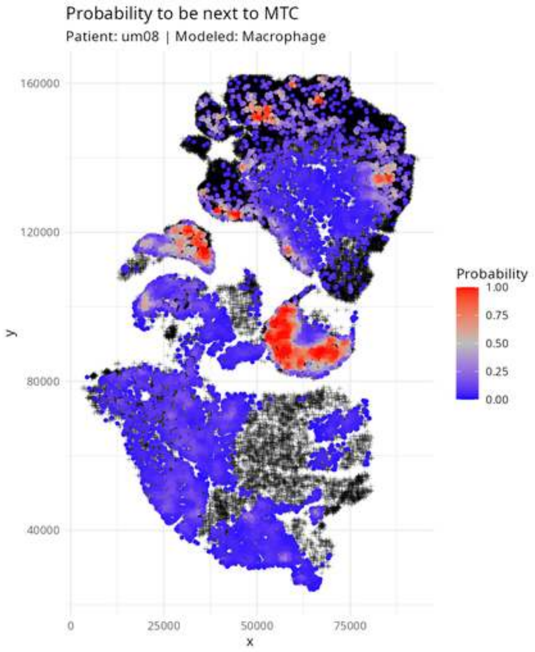

Probability to be next to MTC
Patient: um08 | Modeled: Macrophage
160000
120000
80000
40000
y
0
25000
50000
75000
x
Probability
1.00
0.75
0.50
0.25
0.00

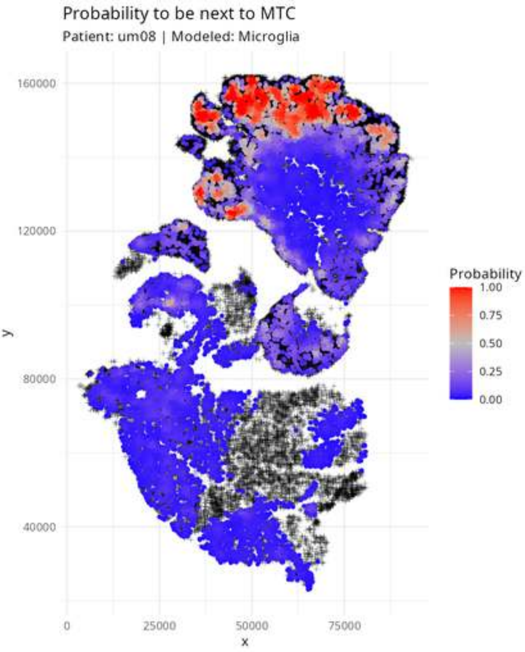
Probability to be next to MTC
Patient: um08 | Modeled: Microglia
160000
120000
80000
40000
y
0
25000
50000
75000
x
Probability
1.00
0.75
0.50
0.25
0.00

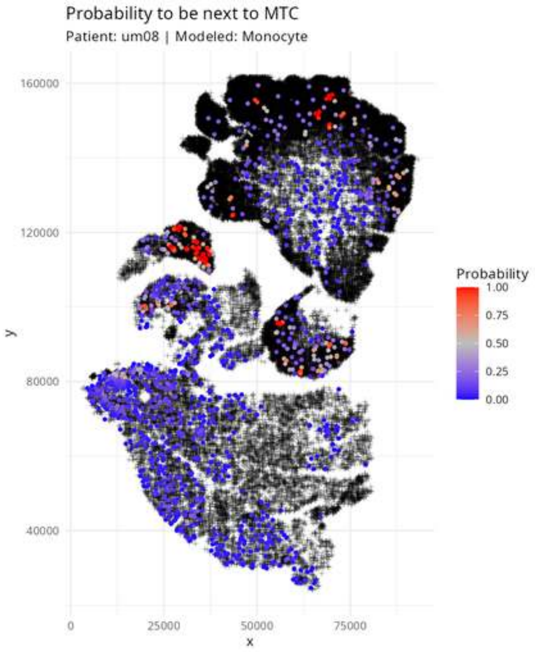

Probability to be next to MTC
Patient: um08 | Modeled: Monocyte
160000
120000
80000
40000
y
0
25000
50000
75000
x
Probability
1.00
0.75
0.50
0.25
0.00

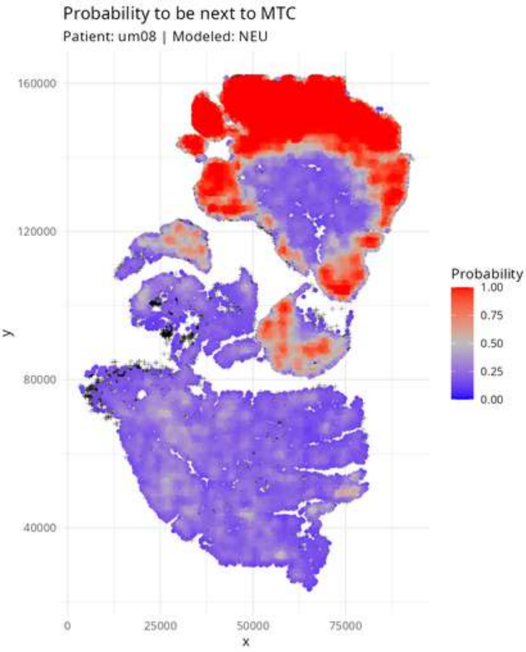

Probability to be next to MTC
Patient: um08 | Modeled: NEU
160000
120000
80000
40000
y
0
25000
50000
75000
x
Probability
1.00
0.75
0.50
0.25
0.00

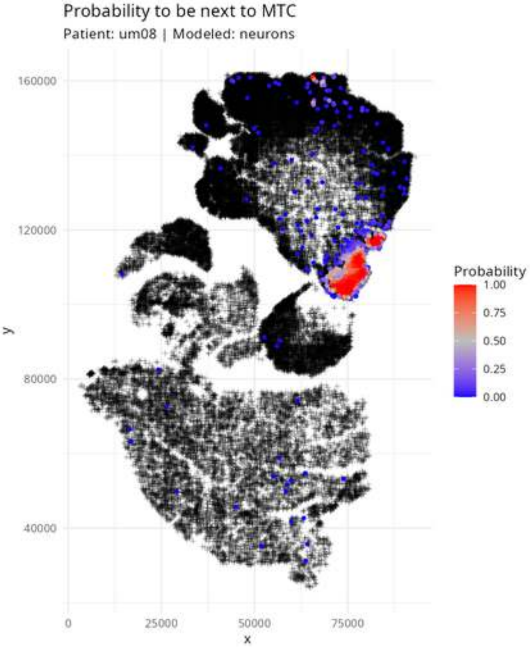
Probability to be next to MTC
Patient: um08 | Modeled: neurons
160000
120000
80000
40000
y
0
25000
50000
75000
x
Probability
1.00
0.75
0.50
0.25
0.00

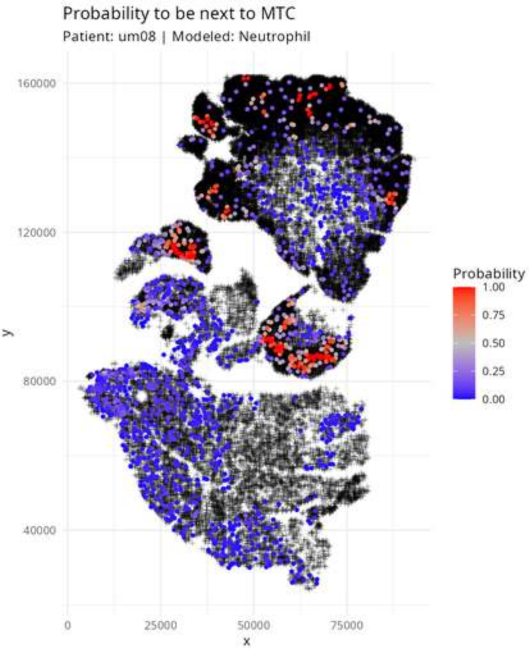

Probability to be next to MTC
Patient: um08 | Modeled: Neutrophil
160000
120000
80000
40000
y
0
25000
50000
75000
x
Probability
1.00
0.75
0.50
0.25
0.00

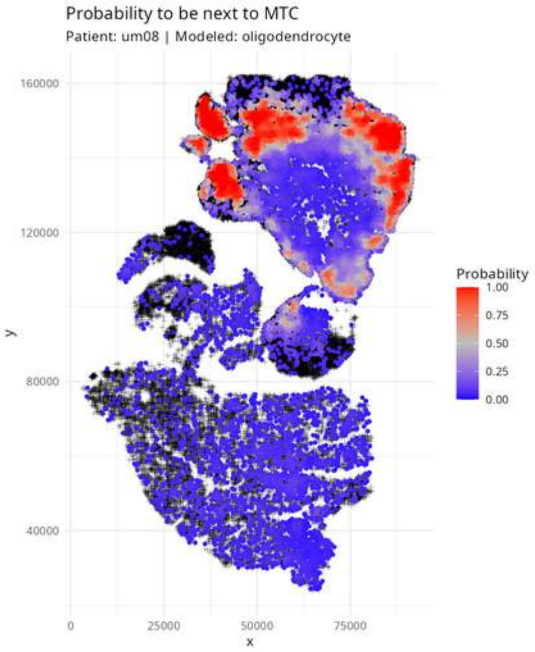
Probability to be next to MTC
Patient: um08 | Modeled: oligodendrocyte
160000
120000
80000
40000
y
0
25000
50000
75000
x
Probability
1.00
0.75
0.50
0.25
0.00

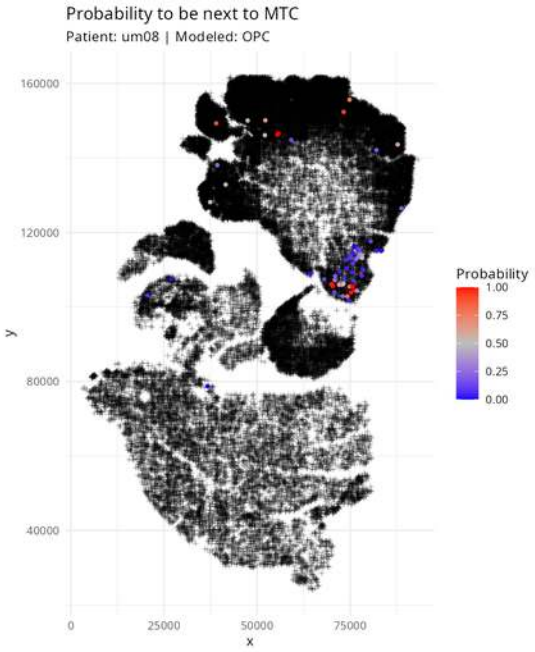
Probability to be next to MTC
Patient: um08 | Modeled: OPC
160000
120000
80000
40000
y
0
25000
50000
75000
x
Probability
1.00
0.75
0.50
0.25
0.00

Probability to be next to MTC

Patient: um08 | Modeled: PPR

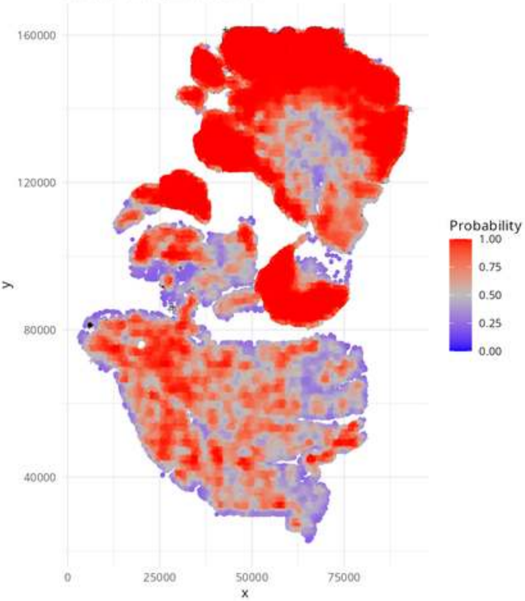

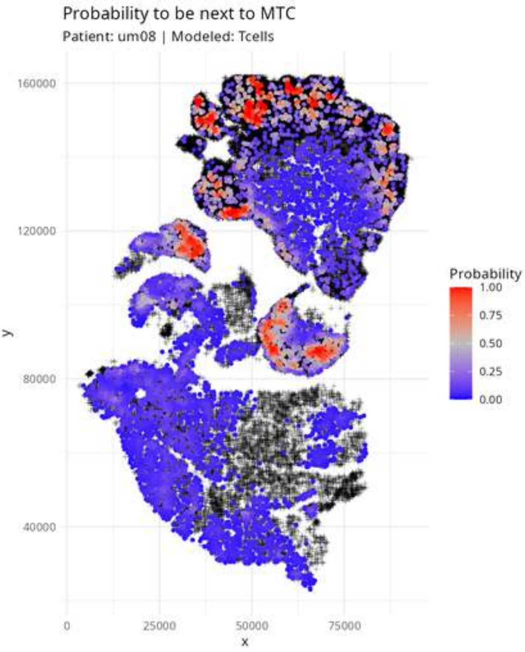

Probability to be next to MTC
Patient: um08 | Modeled: Tcells
160000
120000
80000
40000
y
0
25000
50000
75000
x
Probability
1.00
0.75
0.50
0.25
0.00

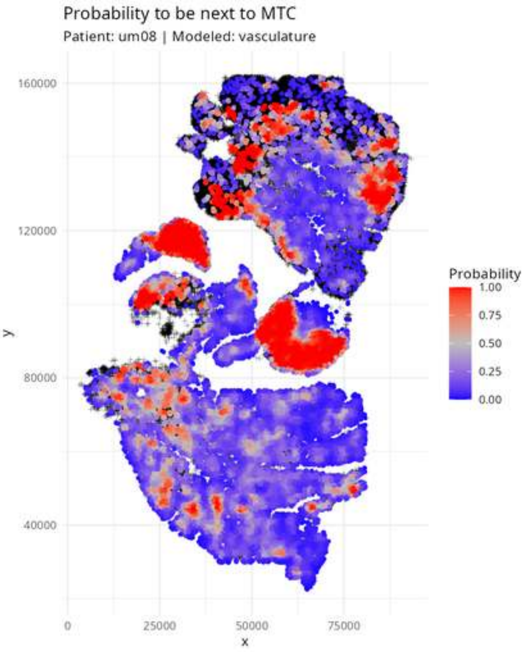

Probability to be next to MTC
Patient: um08 | Modeled: vasculature
y
x
160000
120000
80000
40000
0
25000
50000
75000
Probability
1.00
0.75
0.50
0.25
0.00

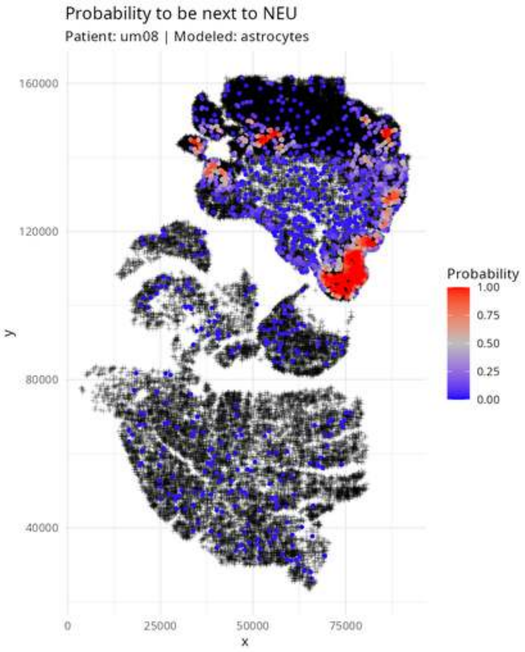

Probability to be next to NEU
Patient: um08 | Modeled: astrocytes
160000
120000
80000
40000
y
0
25000
50000
75000
x
Probability
1.00
0.75
0.50
0.25
0.00

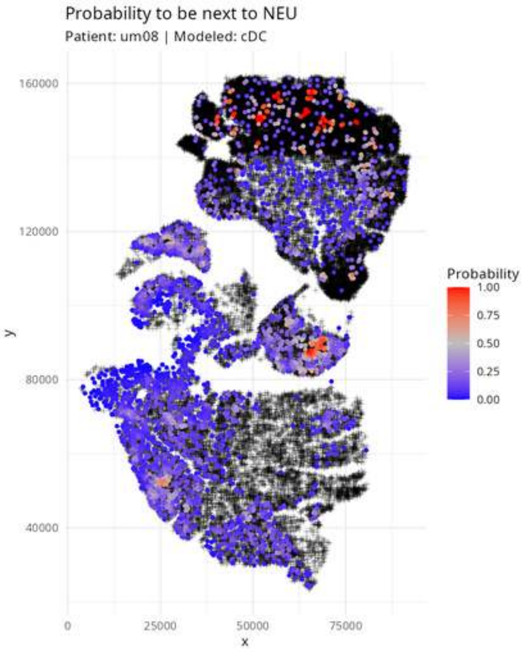

Probability to be next to NEU
Patient: um08 | Modeled: cDC
160000
120000
80000
40000
y
0
25000
50000
75000
x
Probability
1.00
0.75
0.50
0.25
0.00

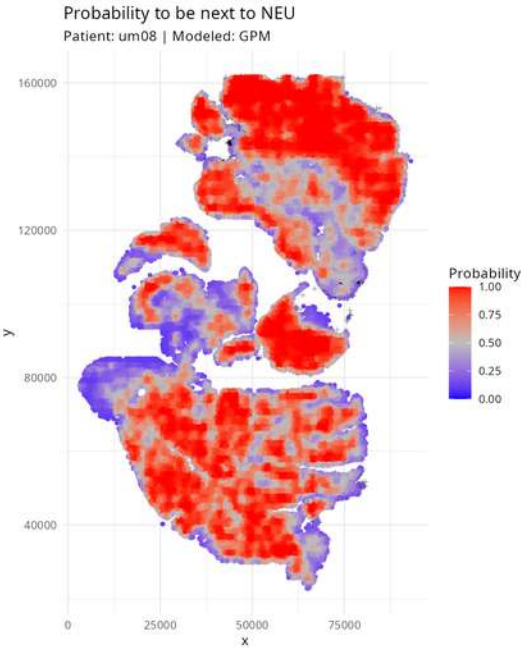
Probability to be next to NEU
Patient: um08 | Modeled: GPM
160000
120000
80000
40000
y
0
25000
50000
75000
x
Probability
1.00
0.75
0.50
0.25
0.00

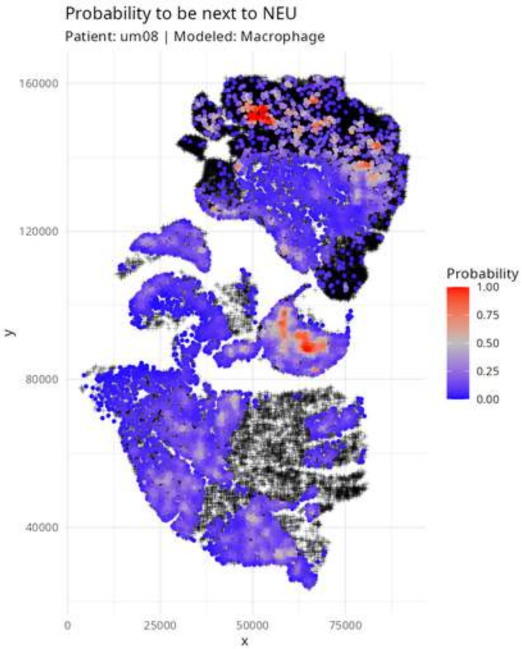
Probability to be next to NEU
Patient: um08 | Modeled: Macrophage
160000
120000
80000
40000
y
0
25000
50000
75000
x
Probability
1.00
0.75
0.50
0.25
0.00

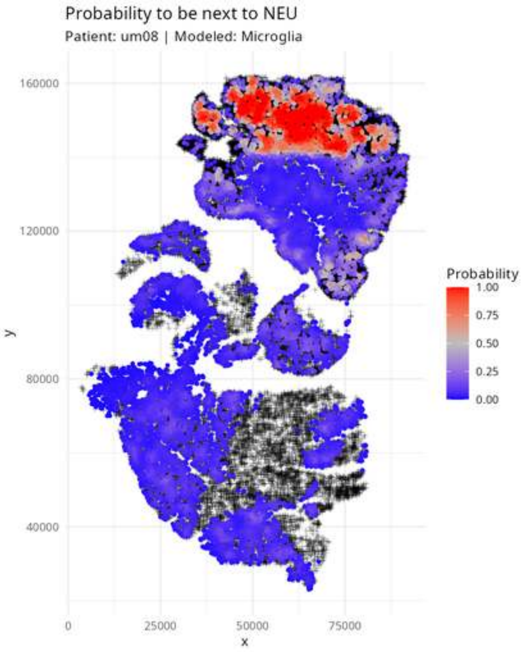
Probability to be next to NEU
Patient: um08 | Modeled: Microglia
160000
120000
80000
40000
y
0
25000
50000
75000
x
Probability
1.00
0.75
0.50
0.25
0.00

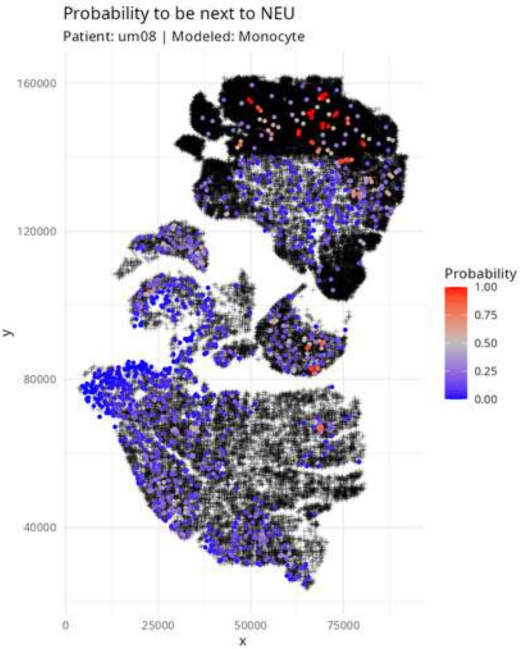

Probability to be next to NEU
Patient: um08 | Modeled: Monocyte
160000
120000
80000
40000
y
0
25000
50000
75000
x
Probability
1.00
0.75
0.50
0.25
0.00

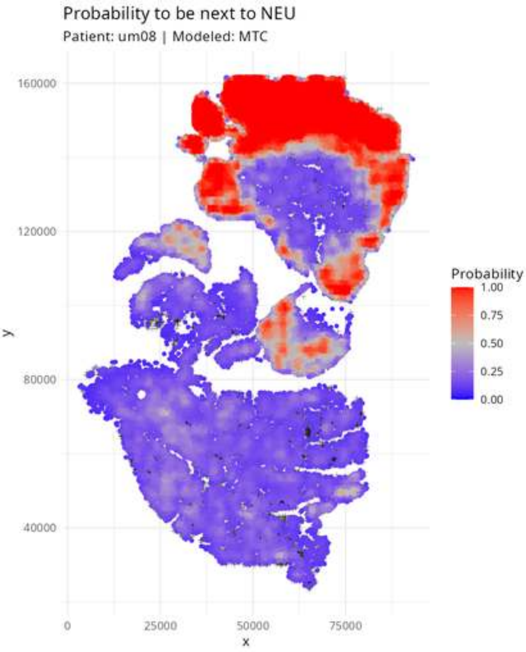

Probability to be next to NEU
Patient: um08 | Modeled: MTC
160000
120000
80000
40000
y
0
25000
50000
75000
x
Probability
1.00
0.75
0.50
0.25
0.00

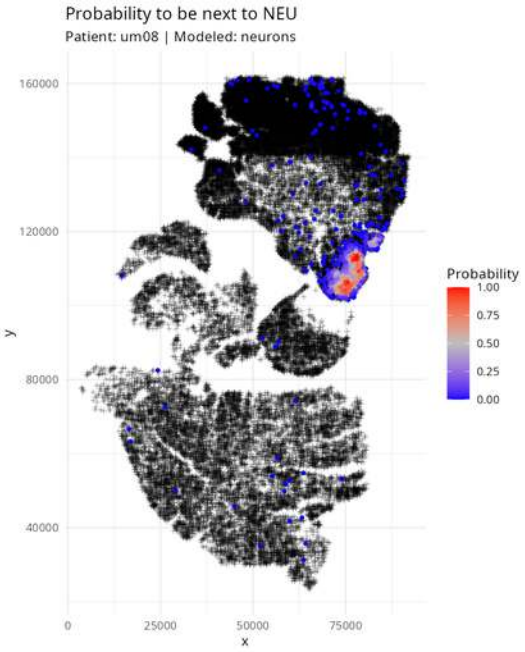

Probability to be next to NEU
Patient: um08 | Modeled: neurons
160000
120000
80000
40000
y
0
25000
50000
75000
x
Probability
1.00
0.75
0.50
0.25
0.00

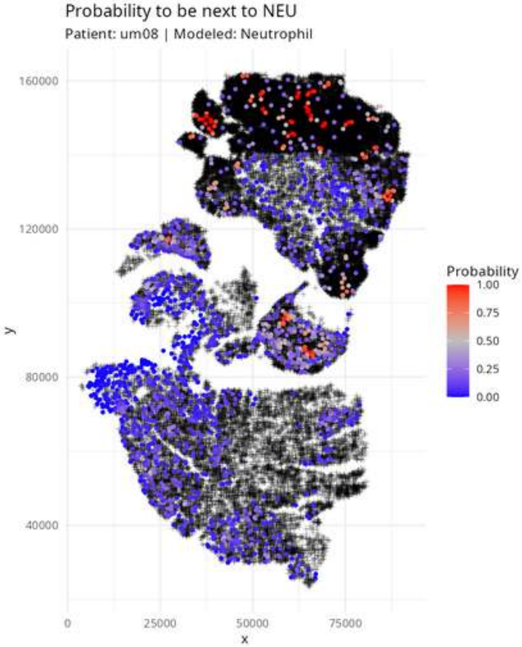
Probability to be next to NEU
Patient: um08 | Modeled: Neutrophil
160000
120000
80000
40000
y
0
25000
50000
75000
x
Probability
1.00
0.75
0.50
0.25
0.00

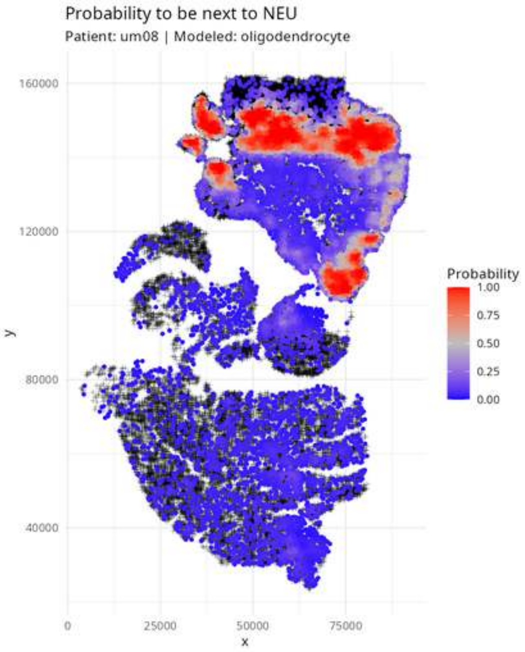

Probability to be next to NEU
Patient: um08 | Modeled: oligodendrocyte
160000
120000
80000
40000
y
0
25000
50000
75000
x
Probability
1.00
0.75
0.50
0.25
0.00

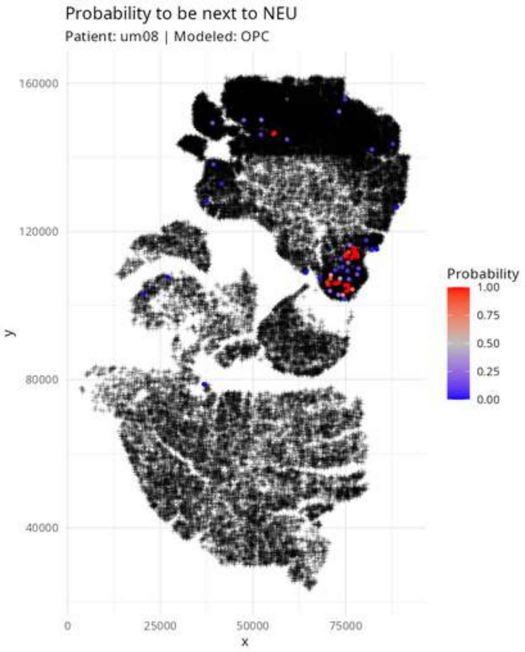

Probability to be next to NEU
Patient: um08 | Modeled: OPC
160000
120000
80000
40000
y
0
25000
50000
75000
x
Probability
1.00
0.75
0.50
0.25
0.00

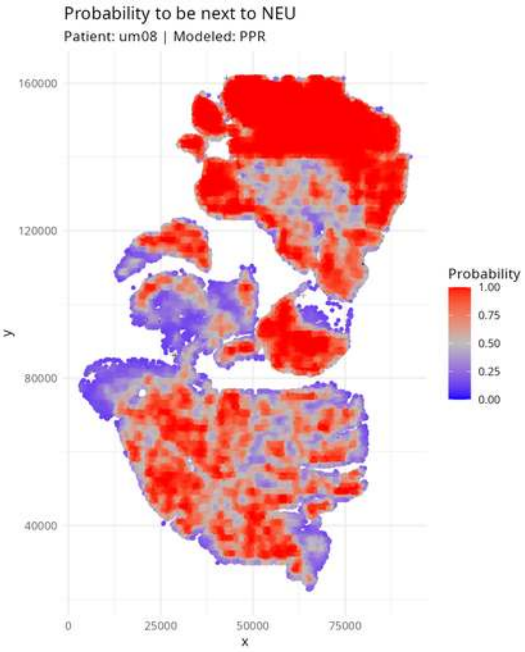

Probability to be next to NEU
Patient: um08 | Modeled: PPR
160000
120000
80000
40000
y
0
25000
50000
75000
x
Probability
1.00
0.75
0.50
0.25
0.00

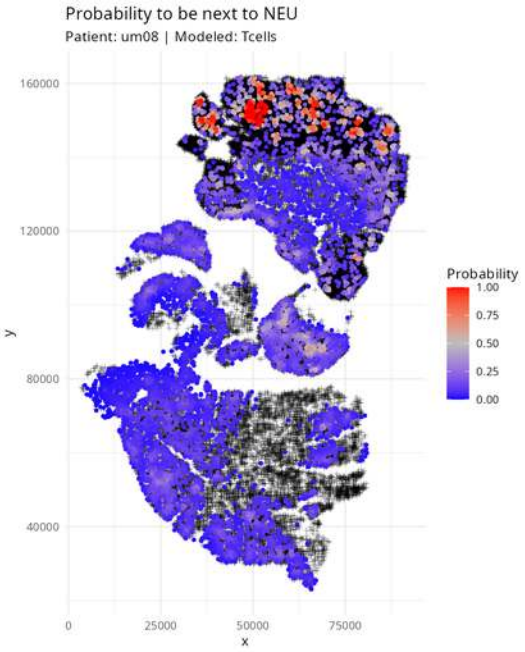
Probability to be next to NEU
Patient: um08 | Modeled: Tcells
160000
120000
80000
40000
y
0
25000
50000
75000
x
Probability
1.00
0.75
0.50
0.25
0.00

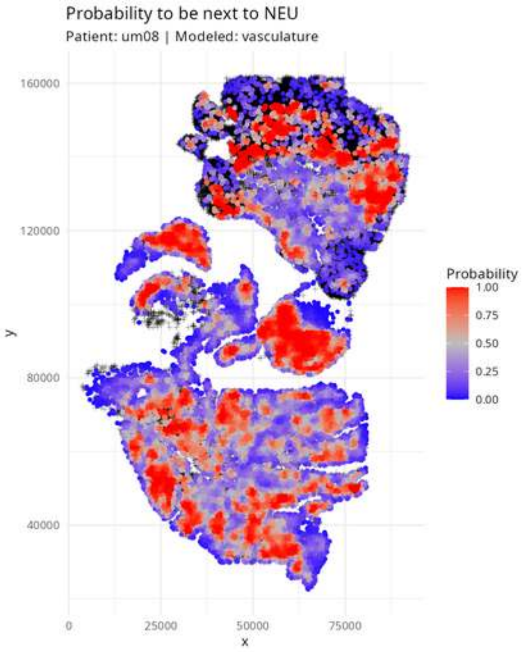

Probability to be next to NEU
Patient: um08 | Modeled: vasculature
160000
120000
80000
40000
y
0
25000
50000
75000
x
Probability
1.00
0.75
0.50
0.25
0.00

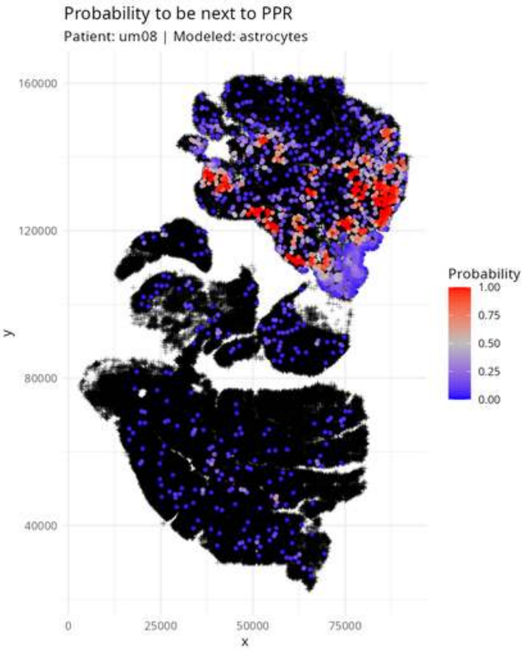

Probability to be next to PPR
Patient: um08 | Modeled: astrocytes
160000
120000
80000
40000
y
0
25000
50000
75000
x
Probability
1.00
0.75
0.50
0.25
0.00

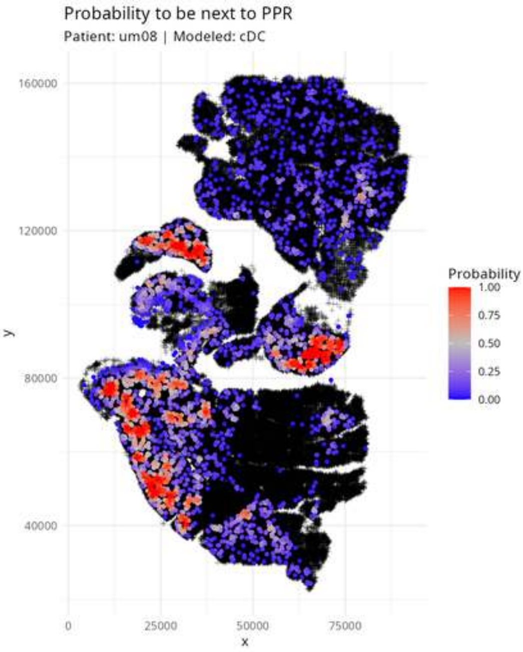
Probability to be next to PPR
Patient: um08 | Modeled: cDC
160000
120000
80000
40000
y
0
25000
50000
75000
x
Probability
1.00
0.75
0.50
0.25
0.00

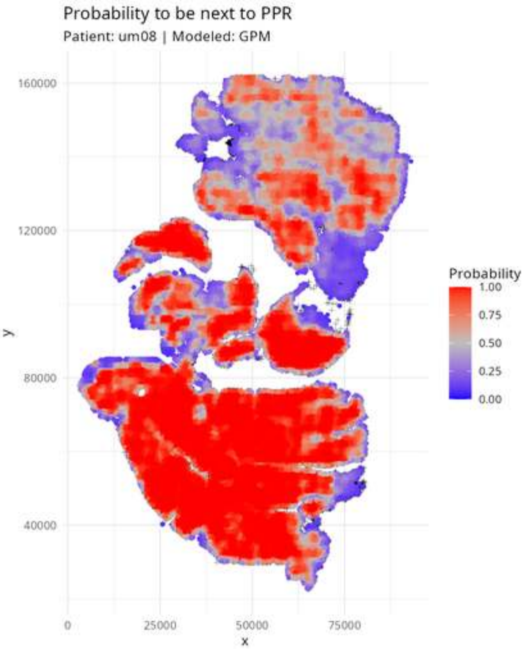

Probability to be next to PPR
Patient: um08 | Modeled: GPM
160000
120000
80000
40000
y
0
25000
50000
75000
x
Probability
1.00
0.75
0.50
0.25
0.00

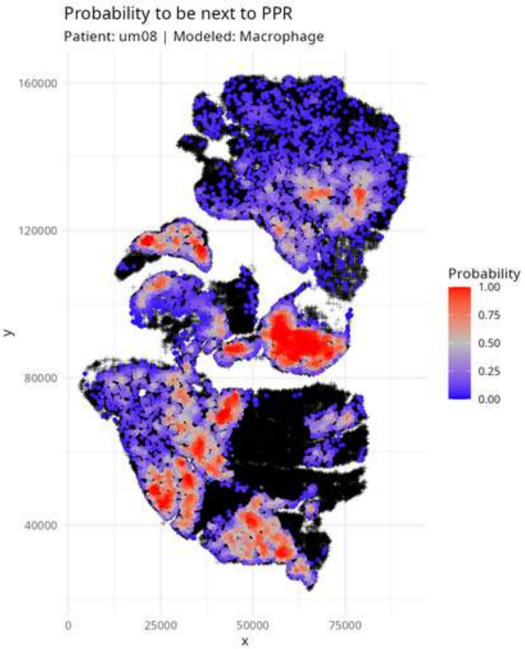

Probability to be next to PPR
Patient: um08 | Modeled: Macrophage
160000
120000
80000
40000
y
0
25000
50000
75000
x
Probability
1.00
0.75
0.50
0.25
0.00

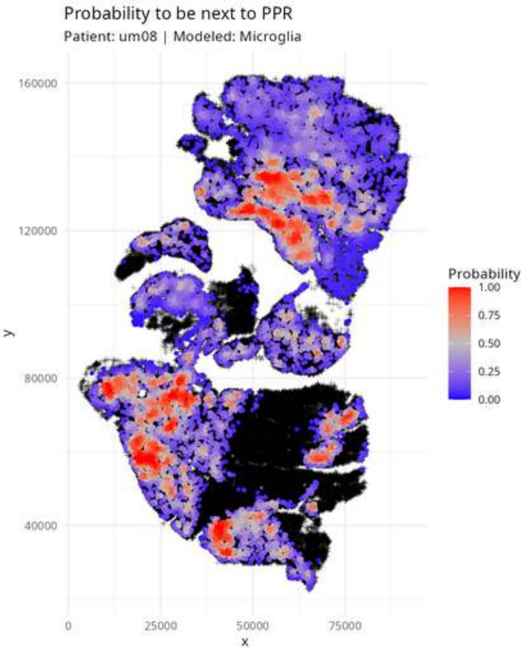

Probability to be next to PPR
Patient: um08 | Modeled: Microglia
160000
120000
80000
40000
y
0
25000
50000
75000
x
Probability
1.00
0.75
0.50
0.25
0.00

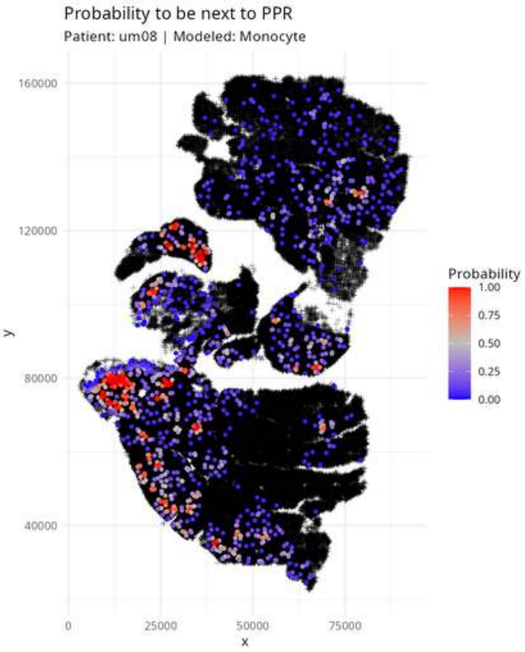

Probability to be next to PPR
Patient: um08 | Modeled: Monocyte
160000
120000
80000
40000
y
0
25000
50000
75000
x
Probability
1.00
0.75
0.50
0.25
0.00

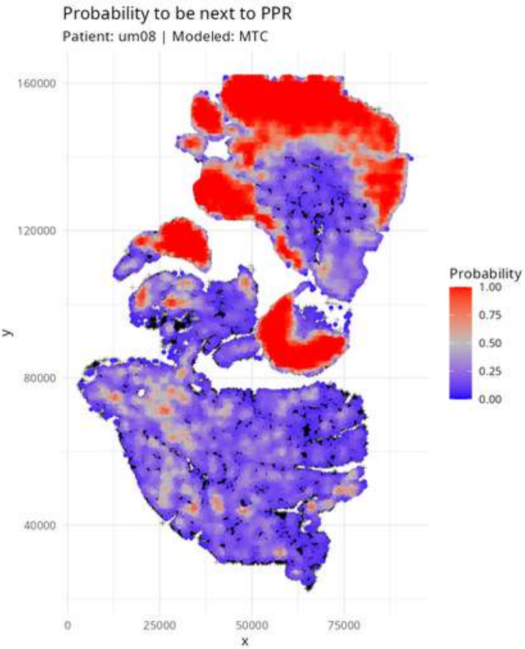
Probability to be next to PPR
Patient: um08 | Modeled: MTC
160000
120000
80000
40000
y
0
25000
50000
75000
x
Probability
1.00
0.75
0.50
0.25
0.00

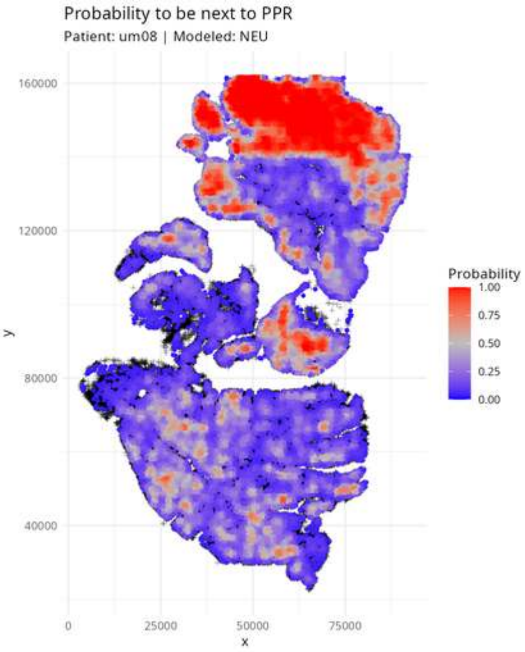

Probability to be next to PPR
Patient: um08 | Modeled: NEU
160000
120000
80000
40000
y
0
25000
50000
75000
x
Probability
1.00
0.75
0.50
0.25
0.00

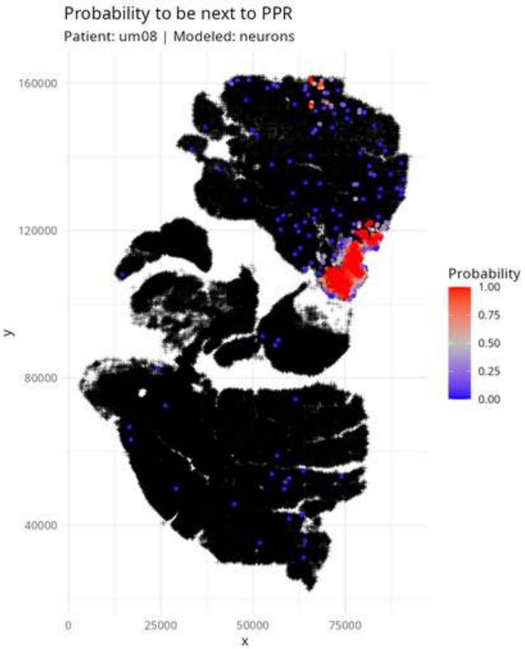

Probability to be next to PPR
Patient: um08 | Modeled: neurons
160000
120000
80000
40000
y
0
25000
50000
75000
x
Probability
1.00
0.75
0.50
0.25
0.00

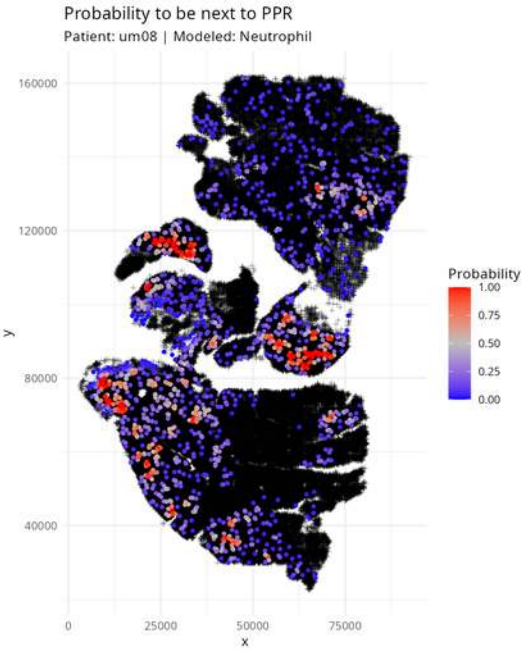
Probability to be next to PPR
Patient: um08 | Modeled: Neutrophil
160000
120000
80000
40000
y
0
25000
50000
75000
x
Probability
1.00
0.75
0.50
0.25
0.00

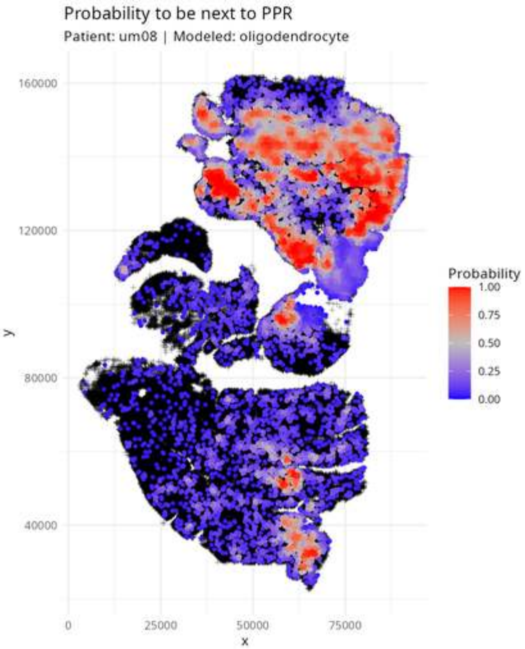
Probability to be next to PPR
Patient: um08 | Modeled: oligodendrocyte
160000
120000
80000
40000
y
0
25000
50000
75000
x
Probability
1.00
0.75
0.50
0.25
0.00

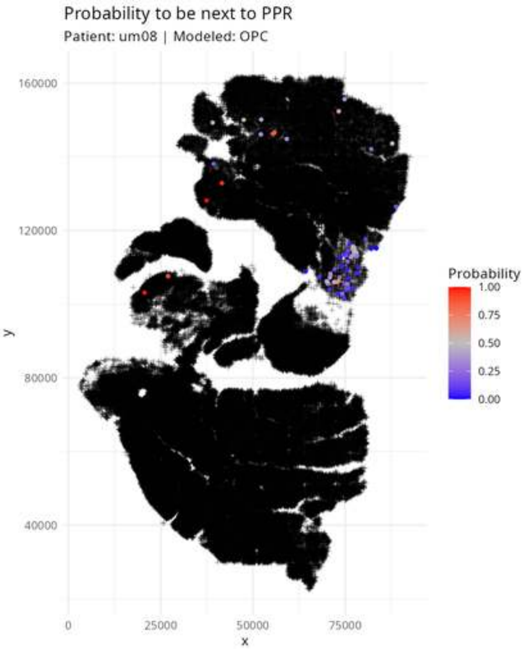

Probability to be next to PPR
Patient: um08 | Modeled: OPC
160000
120000
80000
40000
y
0
25000
50000
75000
x
Probability
1.00
0.75
0.50
0.25
0.00

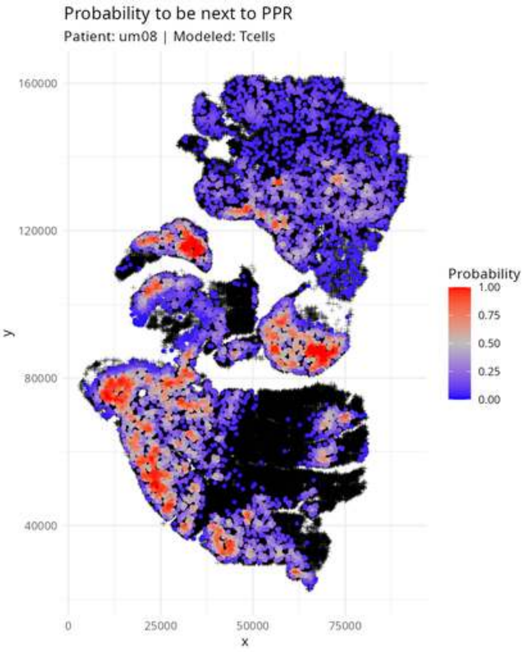
Probability to be next to PPR
Patient: um08 | Modeled: Tcells
160000
120000
80000
40000
y
0
25000
50000
75000
x
Probability
1.00
0.75
0.50
0.25
0.00

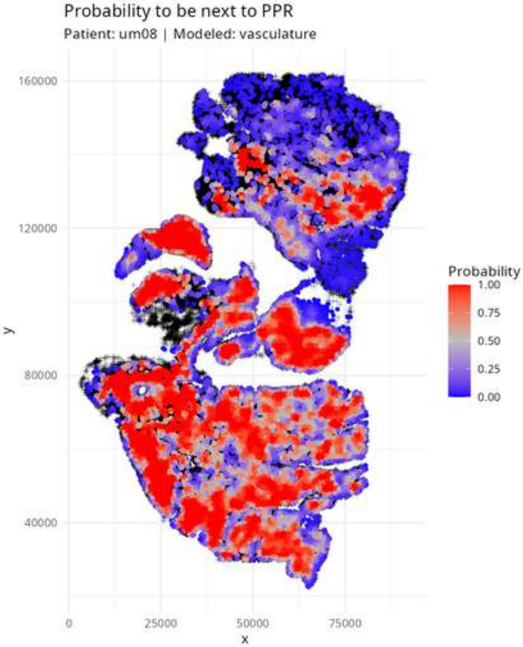
Probability to be next to PPR
Patient: um08 | Modeled: vasculature
160000
120000
80000
40000
y
0
25000
50000
75000
x
Probability
1.00
0.75
0.50
0.25
0.00

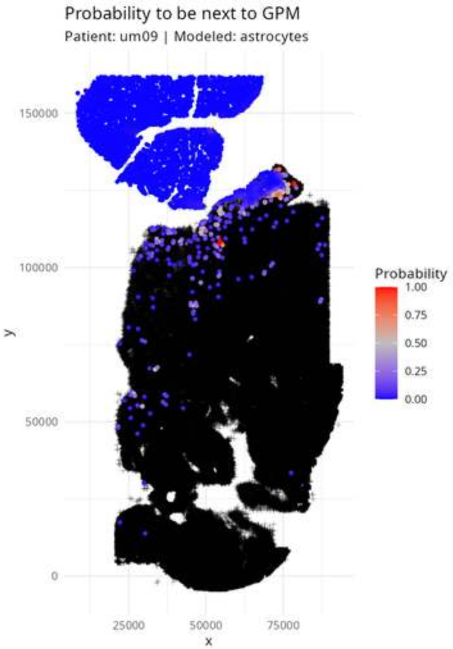
Probability to be next to GPM
Patient: um09 | Modeled: astrocytes
150000
100000
50000
0
y
25000
50000
75000
x
Probability
1.00
0.75
0.50
0.25
0.00

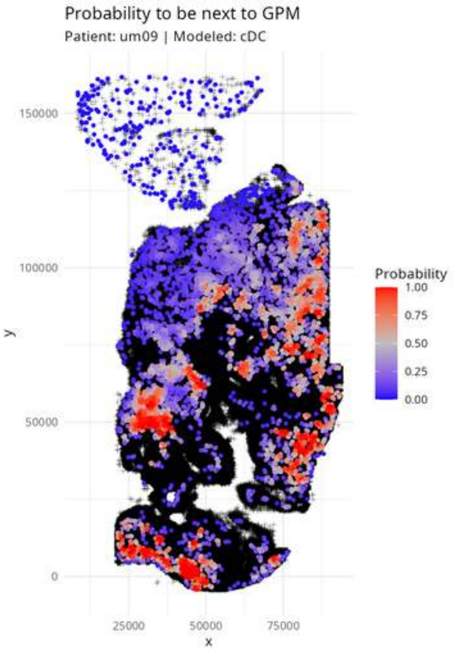
Probability to be next to GPM
Patient: um09 | Modeled: cDC
150000
100000
50000
0
y
25000
50000
75000
x
Probability
1.00
0.75
0.50
0.25
0.00

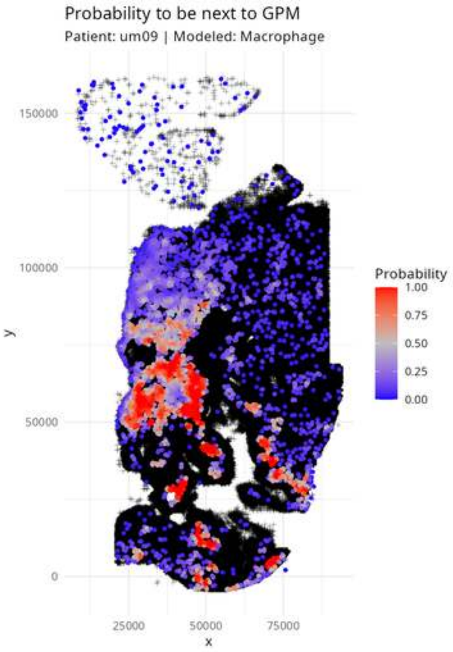

Probability to be next to GPM
Patient: um09 | Modeled: Macrophage
150000
100000
50000
0
y
25000
50000
75000
x
Probability
1.00
0.75
0.50
0.25
0.00

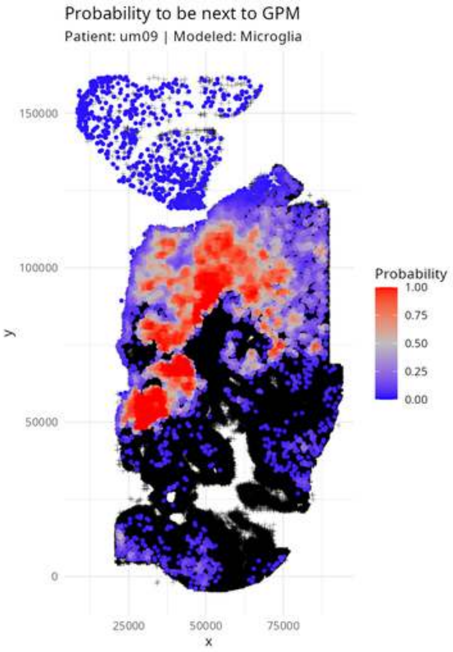

Probability to be next to GPM
Patient: um09 | Modeled: Microglia
150000
100000
50000
0
y
25000
50000
75000
x
Probability
1.00
0.75
0.50
0.25
0.00

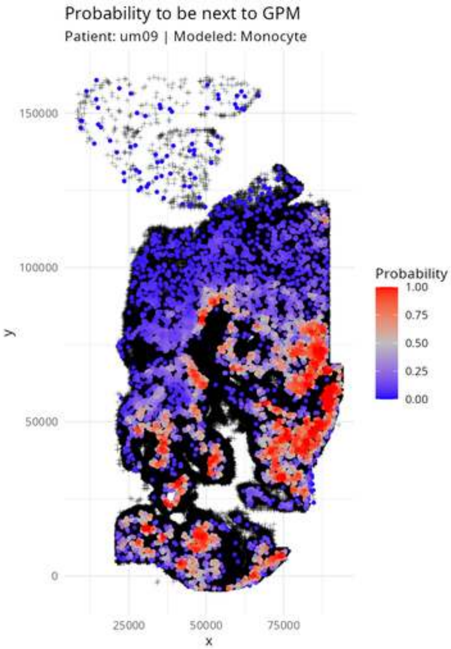

Probability to be next to GPM
Patient: um09 | Modeled: Monocyte
150000
100000
50000
0
y
25000
50000
75000
x
Probability
1.00
0.75
0.50
0.25
0.00

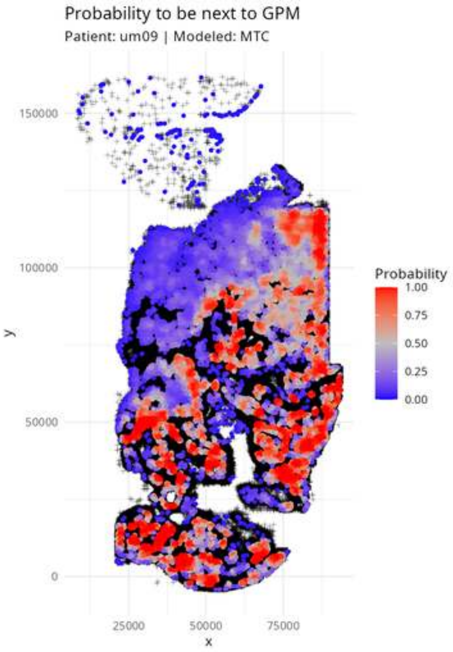
Probability to be next to GPM
Patient: um09 | Modeled: MTC
150000
100000
50000
0
y
25000
50000
75000
x
Probability
1.00
0.75
0.50
0.25
0.00

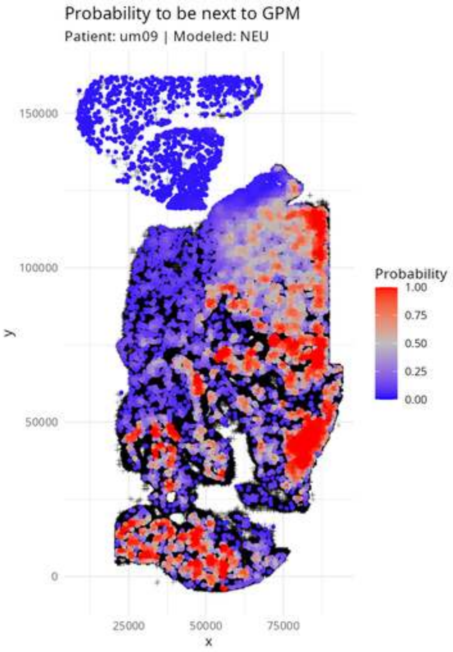

Probability to be next to GPM
Patient: um09 | Modeled: NEU
150000
100000
50000
0
y
25000
50000
75000
x
Probability
1.00
0.75
0.50
0.25
0.00

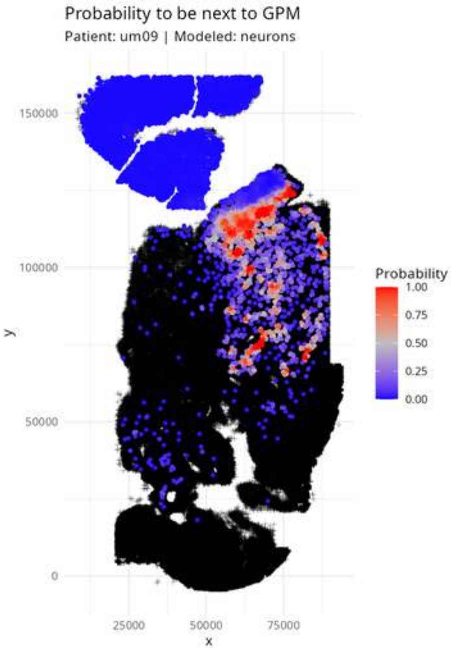
Probability to be next to GPM
Patient: um09 | Modeled: neurons
150000
100000
50000
0
y
25000
50000
75000
x
Probability
1.00
0.75
0.50
0.25
0.00

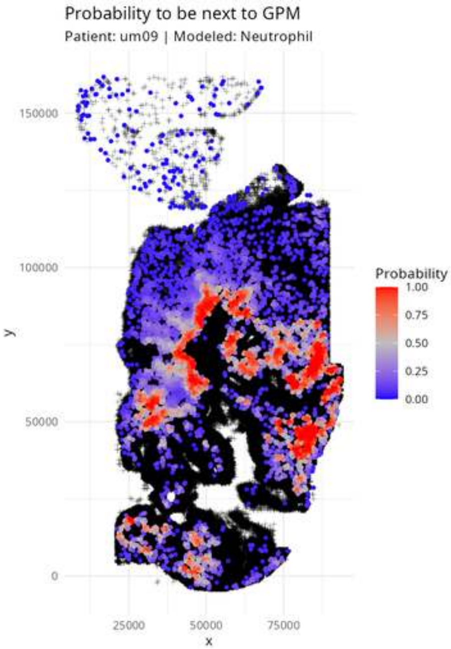

Probability to be next to GPM
Patient: um09 | Modeled: Neutrophil
150000
100000
50000
0
y
25000
50000
75000
x
Probability
1.00
0.75
0.50
0.25
0.00

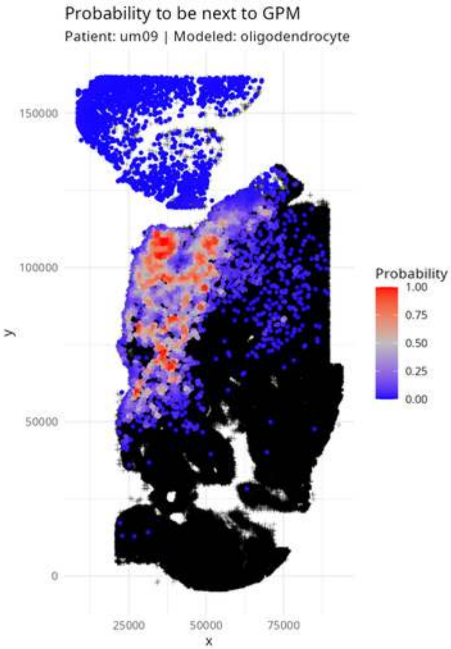

Probability to be next to GPM
Patient: um09 | Modeled: oligodendrocyte
150000
100000
50000
0
y
25000
50000
75000
x
Probability
1.00
0.75
0.50
0.25
0.00

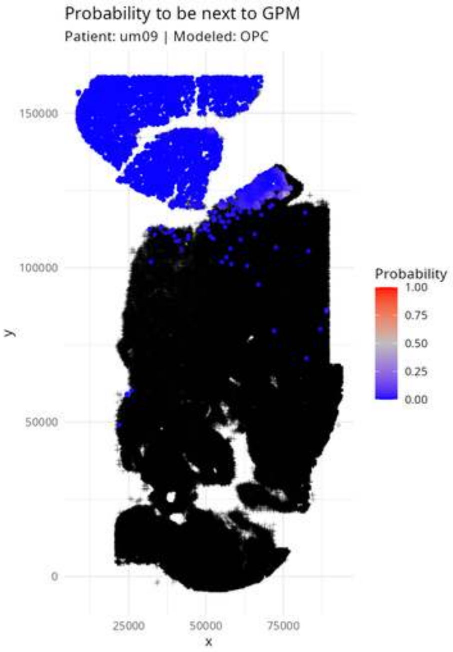
Probability to be next to GPM
Patient: um09 | Modeled: OPC
150000
100000
50000
0
y
25000
50000
75000
x
Probability
1.00
0.75
0.50
0.25
0.00

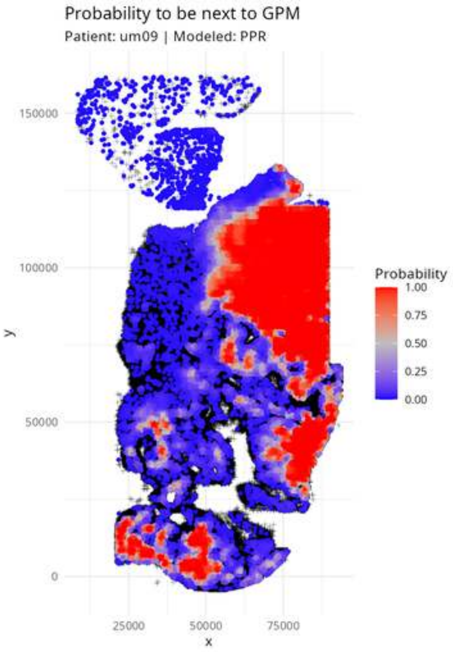

Probability to be next to GPM
Patient: um09 | Modeled: PPR
150000
100000
50000
0
y
25000
50000
75000
x
Probability
1.00
0.75
0.50
0.25
0.00

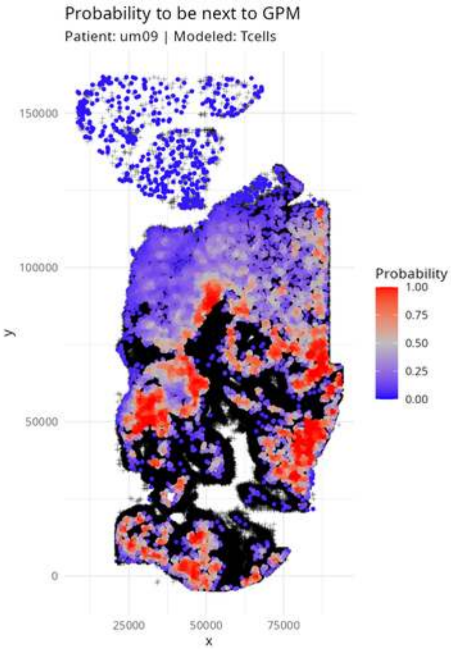
Probability to be next to GPM
Patient: um09 | Modeled: Tcells
150000
100000
50000
0
25000
50000
75000
x
y
Probability
1.00
0.75
0.50
0.25
0.00

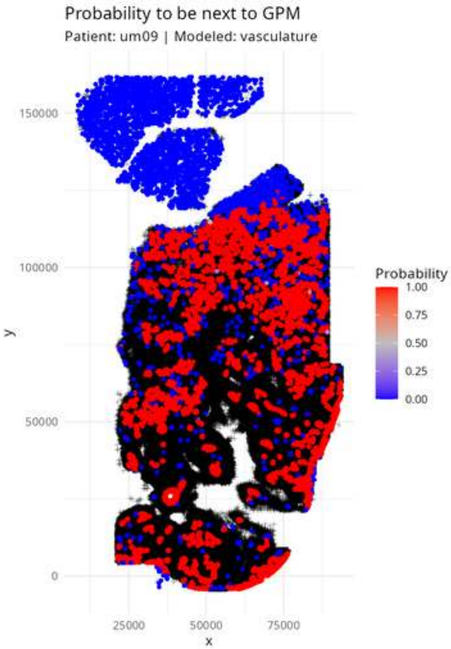

Probability to be next to GPM
Patient: um09 | Modeled: vasculature
150000
100000
50000
0
y
25000
50000
75000
x
Probability
1.00
0.75
0.50
0.25
0.00

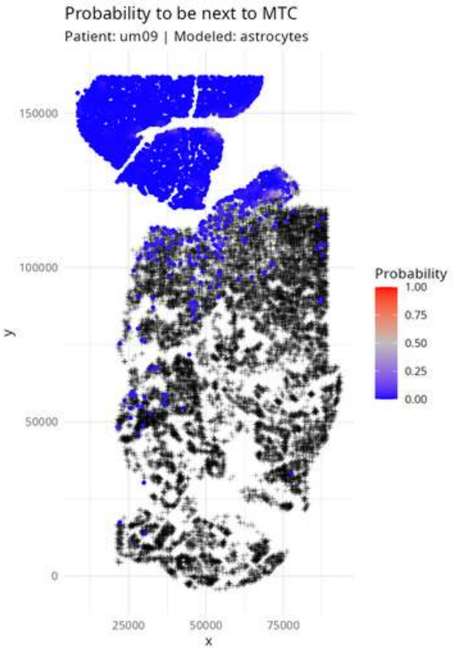

Probability to be next to MTC
Patient: um09 | Modeled: astrocytes
150000
100000
50000
0
y
25000
50000
75000
x
Probability
1.00
0.75
0.50
0.25
0.00

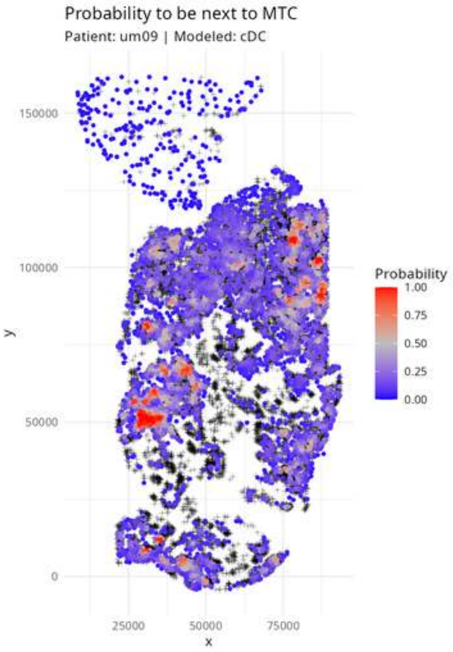
Probability to be next to MTC
Patient: um09 | Modeled: cDC
150000
100000
50000
0
y
25000
50000
75000
x
Probability
1.00
0.75
0.50
0.25
0.00

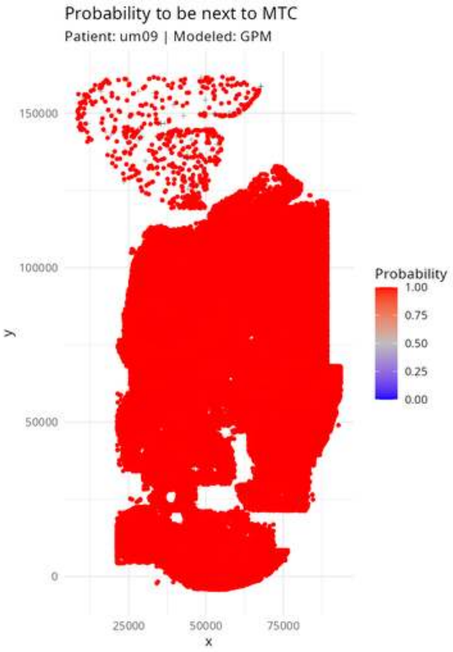
Probability to be next to MTC
Patient: um09 | Modeled: GPM
Probability
1.00
0.75
0.50
0.25
0.00
y
150000
100000
50000
0
x
25000
50000
75000

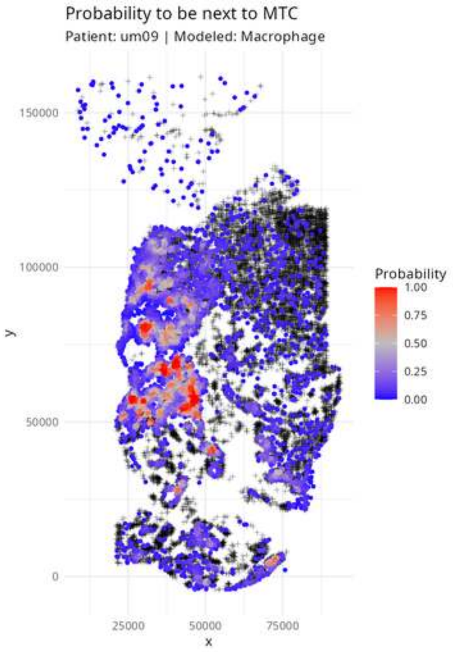
Probability to be next to MTC
Patient: um09 | Modeled: Macrophage
Probability
1.00
0.75
0.50
0.25
0.00
150000
100000
50000
0
25000
50000
75000
x
y

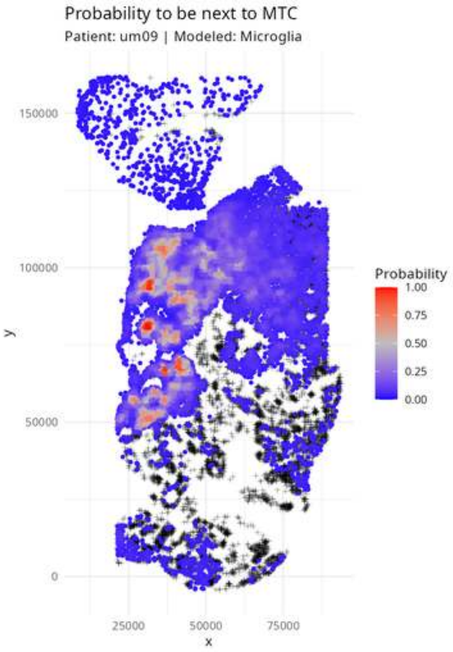

Probability to be next to MTC
Patient: um09 | Modeled: Microglia
150000
100000
50000
0
y
25000
50000
75000
x
Probability
1.00
0.75
0.50
0.25
0.00

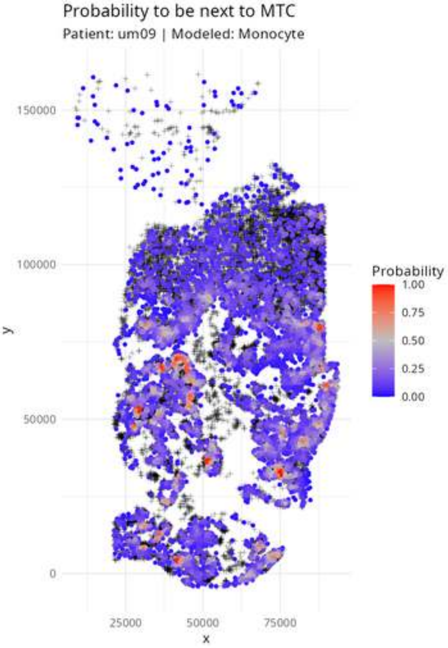
Probability to be next to MTC
Patient: um09 | Modeled: Monocyte
150000
100000
50000
0
y
25000
50000
75000
x
Probability
1.00
0.75
0.50
0.25
0.00

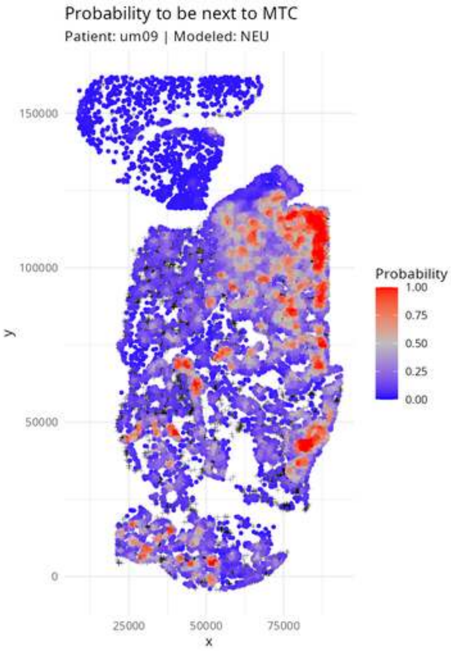
Probability to be next to MTC
Patient: um09 | Modeled: NEU
150000
100000
50000
0
y
25000
50000
75000
x
Probability
1.00
0.75
0.50
0.25
0.00

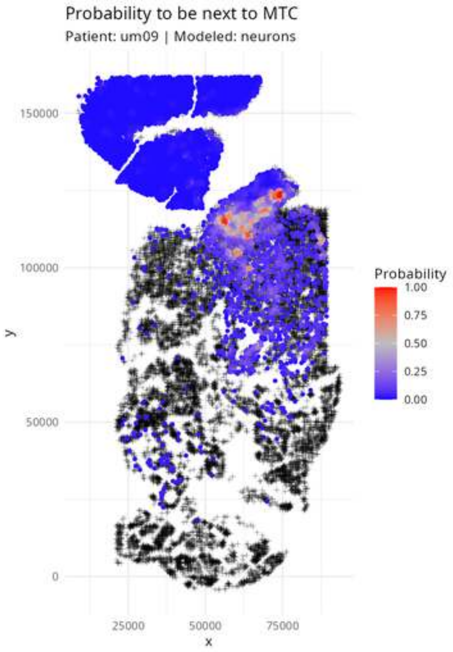
Probability to be next to MTC
Patient: um09 | Modeled: neurons
150000
100000
50000
0
25000
50000
75000
x
y
Probability
1.00
0.75
0.50
0.25
0.00

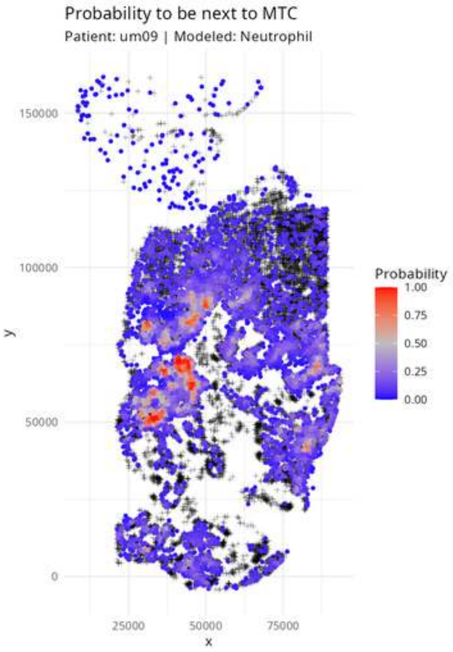

Probability to be next to MTC
Patient: um09 | Modeled: Neutrophil
y
150000
100000
50000
0
x
25000
50000
75000
Probability
1.00
0.75
0.50
0.25
0.00

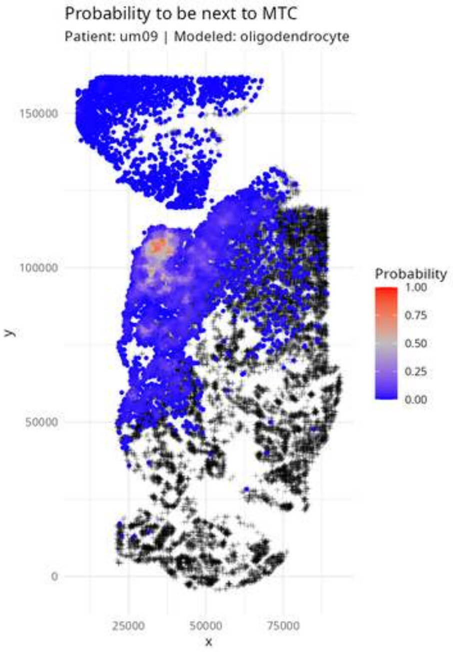

Probability to be next to MTC
Patient: um09 | Modeled: oligodendrocyte
150000
100000
50000
0
y
25000
50000
75000
x
Probability
1.00
0.75
0.50
0.25
0.00

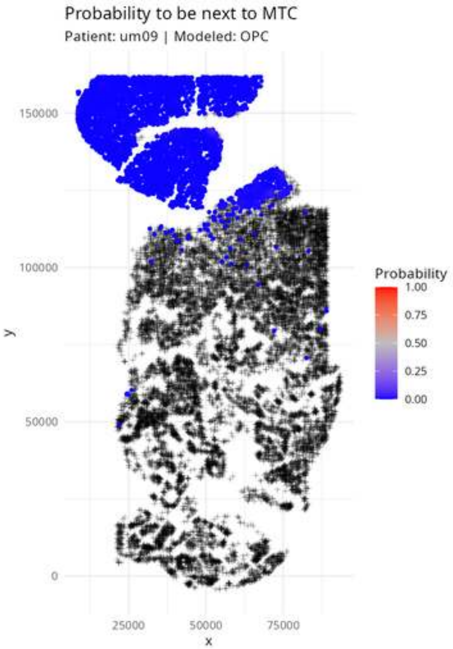
Probability to be next to MTC
Patient: um09 | Modeled: OPC
y
150000
100000
50000
0
x
25000
50000
75000
Probability
1.00
0.75
0.50
0.25
0.00

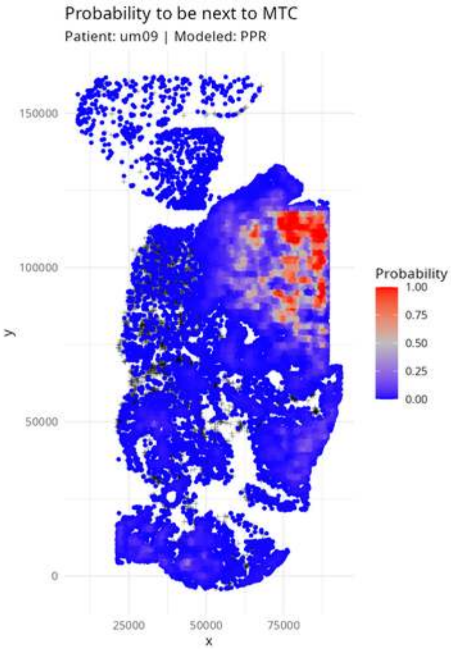

Probability to be next to MTC
Patient: um09 | Modeled: PPR
150000
100000
50000
0
y
25000
50000
75000
x
Probability
1.00
0.75
0.50
0.25
0.00

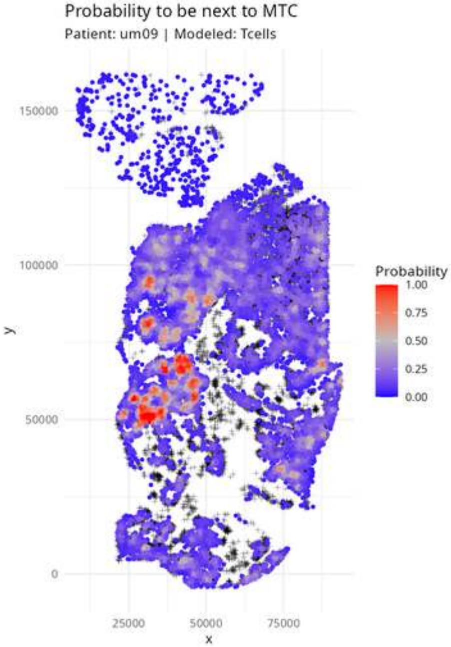
Probability to be next to MTC
Patient: um09 | Modeled: Tcells
150000
100000
50000
0
25000
50000
75000
x
y
Probability
1.00
0.75
0.50
0.25
0.00

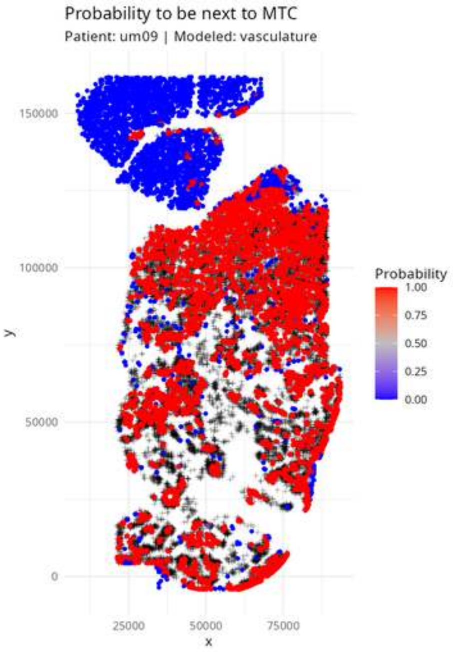

Probability to be next to MTC
Patient: um09 | Modeled: vasculature
150000
100000
50000
0
y
25000
50000
75000
x
Probability
1.00
0.75
0.50
0.25
0.00

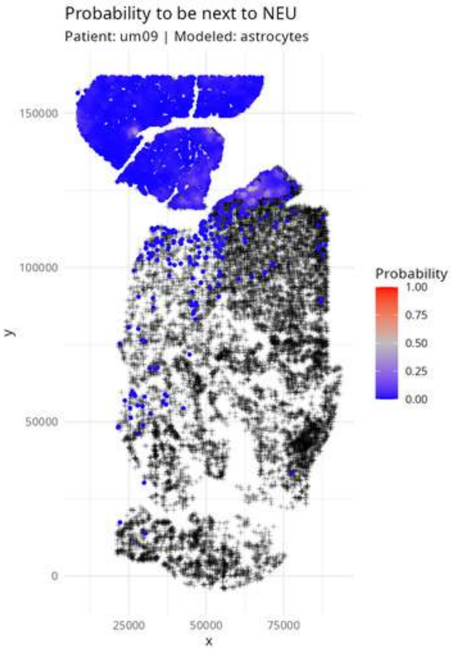

Probability to be next to NEU
Patient: um09 | Modeled: astrocytes
150000
100000
50000
0
y
25000
50000
75000
x
Probability
1.00
0.75
0.50
0.25
0.00

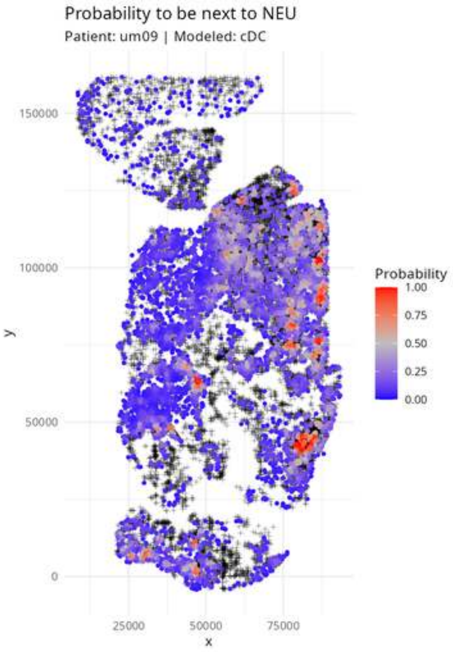
Probability to be next to NEU
Patient: um09 | Modeled: cDC
150000
100000
50000
0
y
25000
50000
75000
x
Probability
1.00
0.75
0.50
0.25
0.00

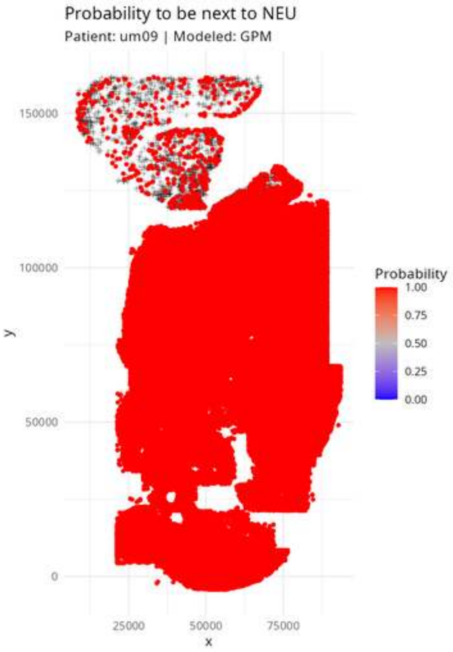

Probability to be next to NEU
Patient: um09 | Modeled: GPM
150000
100000
50000
0
y
25000
50000
75000
x
Probability
1.00
0.75
0.50
0.25
0.00

Probability to be next to NEU

Patient: um09 | Modeled: Macrophage

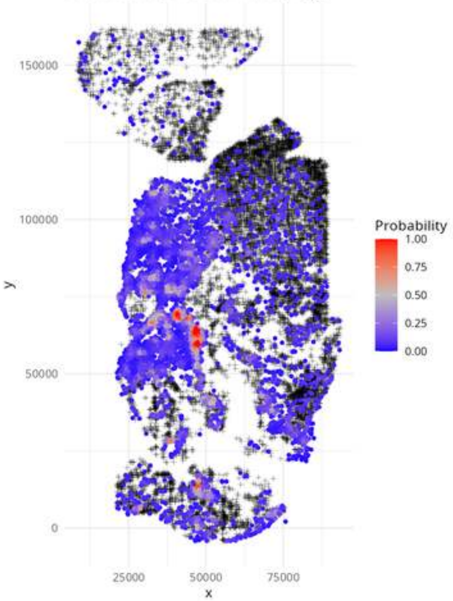

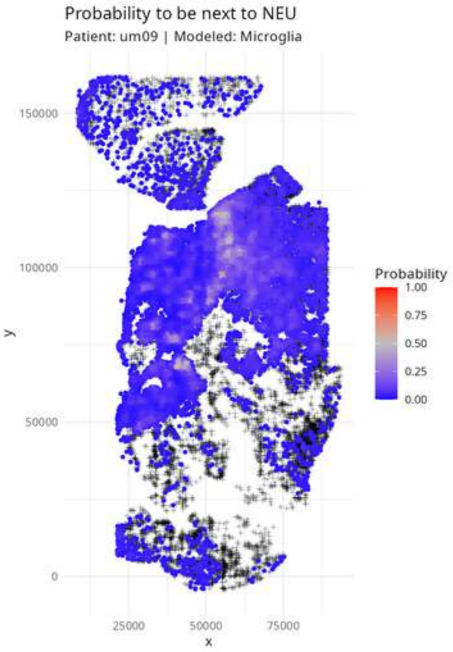
Probability to be next to NEU
Patient: um09 | Modeled: Microglia
Probability
1.00
0.75
0.50
0.25
0.00
150000
100000
50000
0
y
25000
50000
75000
x

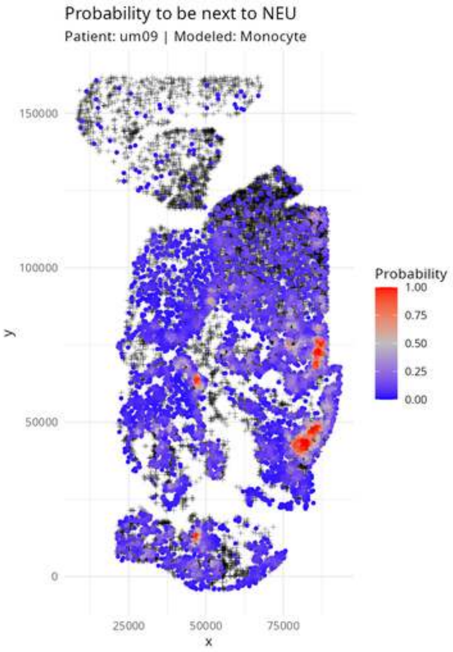

Probability to be next to NEU
Patient: um09 | Modeled: Monocyte
150000
100000
50000
0
y
25000
50000
75000
x
Probability
1.00
0.75
0.50
0.25
0.00

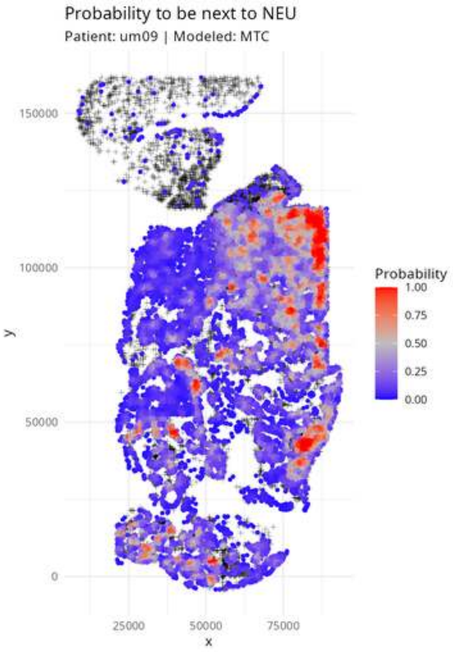

Probability to be next to NEU
Patient: um09 | Modeled: MTC
y
x
150000
100000
50000
0
25000
50000
75000
Probability
1.00
0.75
0.50
0.25
0.00

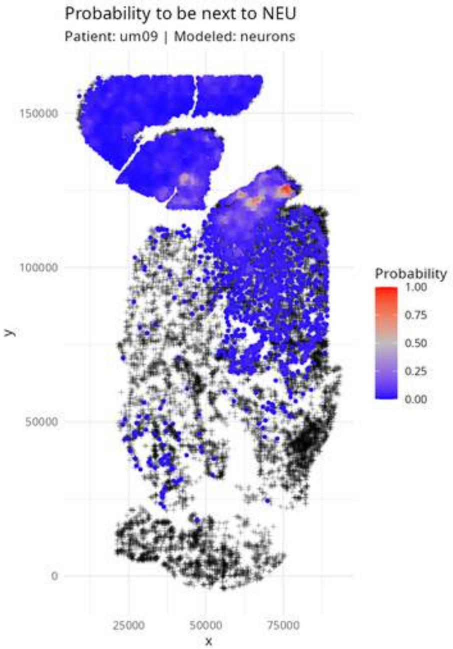
Probability to be next to NEU
Patient: um09 | Modeled: neurons
150000
100000
50000
0
25000
50000
75000
x
y
Probability
1.00
0.75
0.50
0.25
0.00

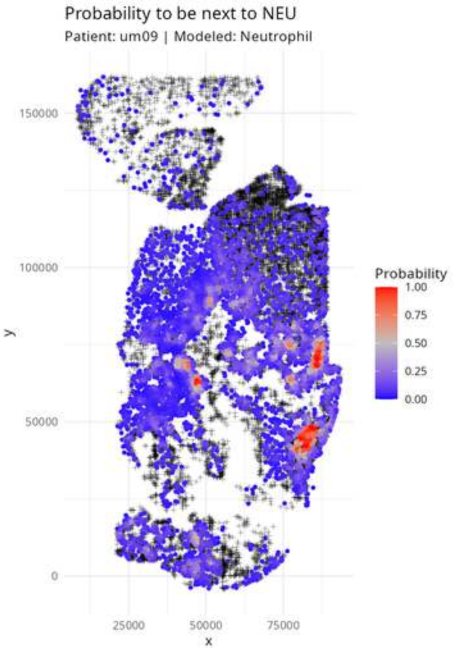
Probability to be next to NEU
Patient: um09 | Modeled: Neutrophil
150000
100000
50000
0
y
25000
50000
75000
x
Probability
1.00
0.75
0.50
0.25
0.00

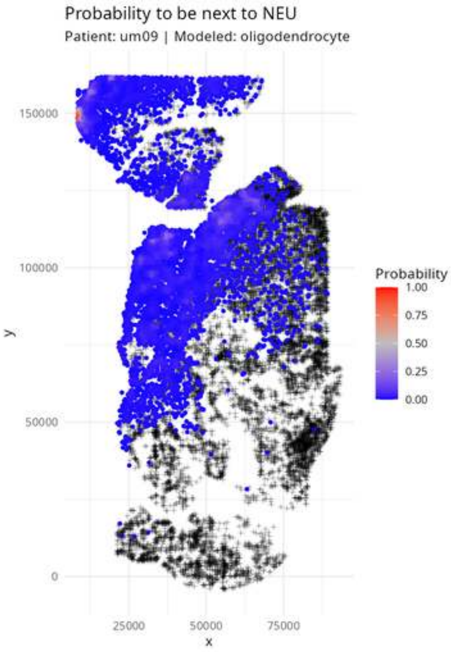

Probability to be next to NEU
Patient: um09 | Modeled: oligodendrocyte
150000
100000
50000
0
y
25000
50000
75000
x
Probability
1.00
0.75
0.50
0.25
0.00

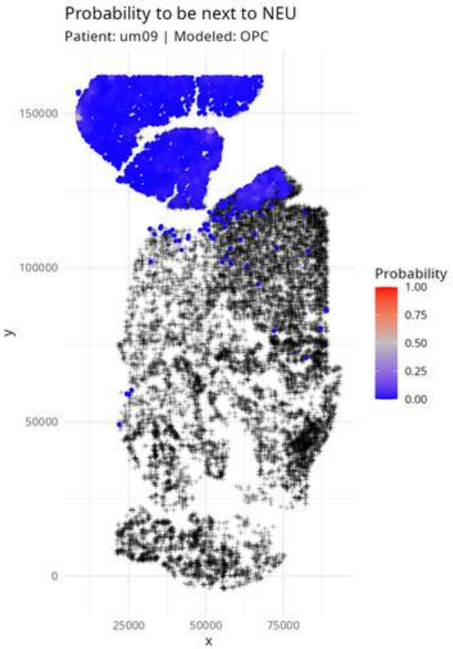

Probability to be next to NEU
Patient: um09 | Modeled: OPC
150000
100000
50000
0
y
25000
50000
75000
x
Probability
1.00
0.75
0.50
0.25
0.00

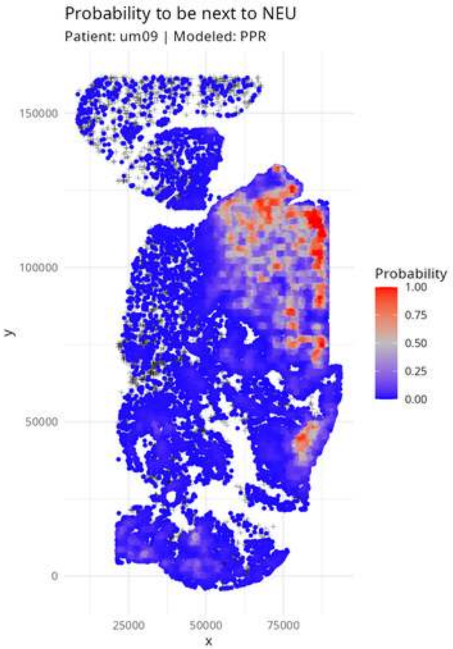
Probability to be next to NEU
Patient: um09 | Modeled: PPR
150000
100000
50000
0
y
25000
50000
75000
x
Probability
1.00
0.75
0.50
0.25
0.00

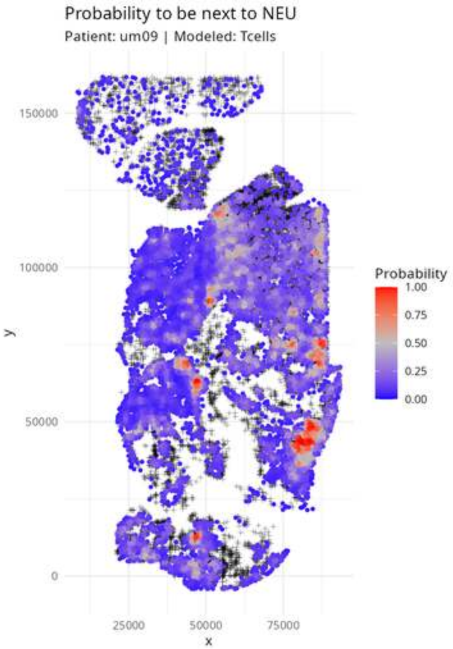

Probability to be next to NEU
Patient: um09 | Modeled: Tcells
150000
100000
50000
0
y
25000
50000
75000
x
Probability
1.00
0.75
0.50
0.25
0.00

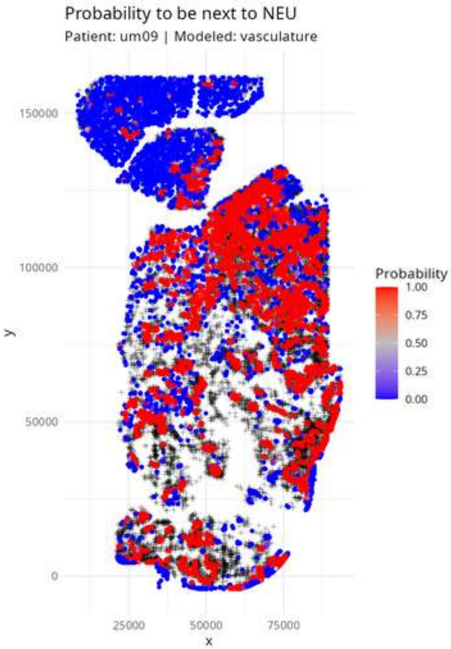

Probability to be next to NEU
Patient: um09 | Modeled: vasculature
y
150000
100000
50000
0
x
25000
50000
75000
Probability
1.00
0.75
0.50
0.25
0.00

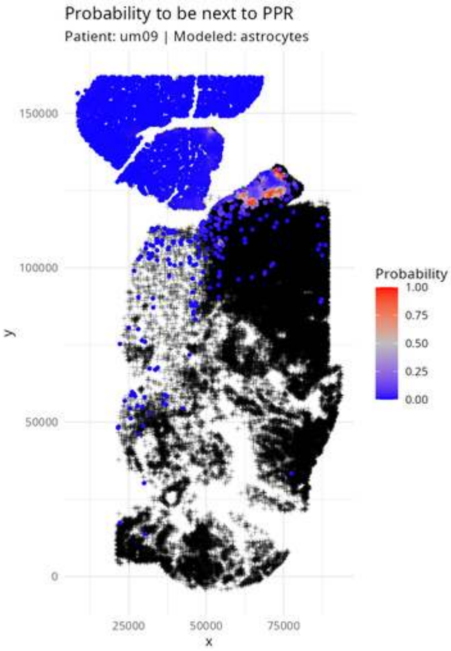

Probability to be next to PPR
Patient: um09 | Modeled: astrocytes
150000
100000
50000
0
y
25000
50000
75000
x
Probability
1.00
0.75
0.50
0.25
0.00

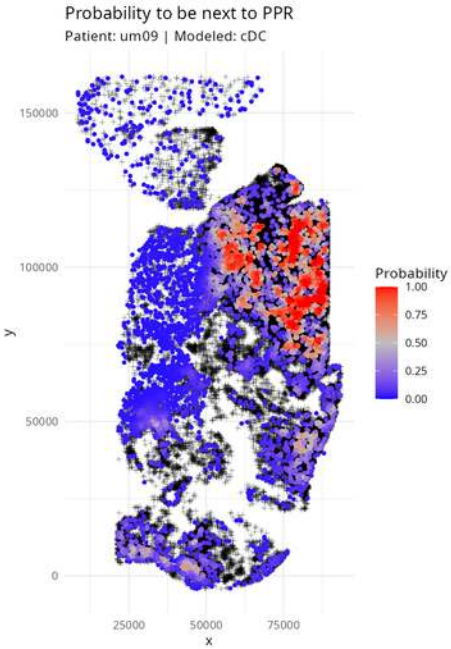
Probability to be next to PPR
Patient: um09 | Modeled: cDC
150000
100000
50000
0
y
25000
50000
75000
x
Probability
1.00
0.75
0.50
0.25
0.00

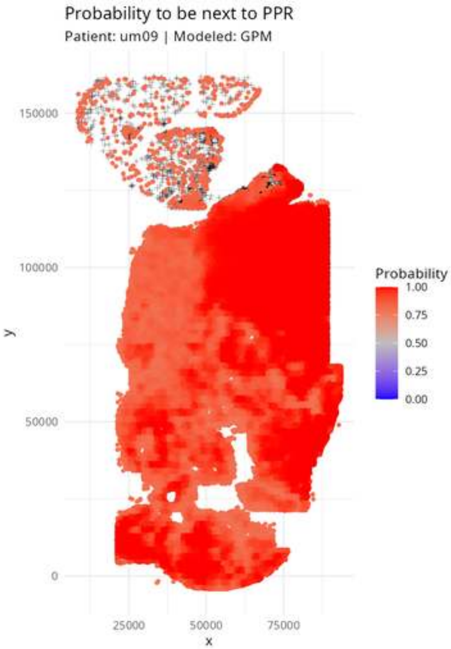

Probability to be next to PPR
Patient: um09 | Modeled: GPM
150000
100000
50000
0
y
25000
50000
75000
x
Probability
1.00
0.75
0.50
0.25
0.00

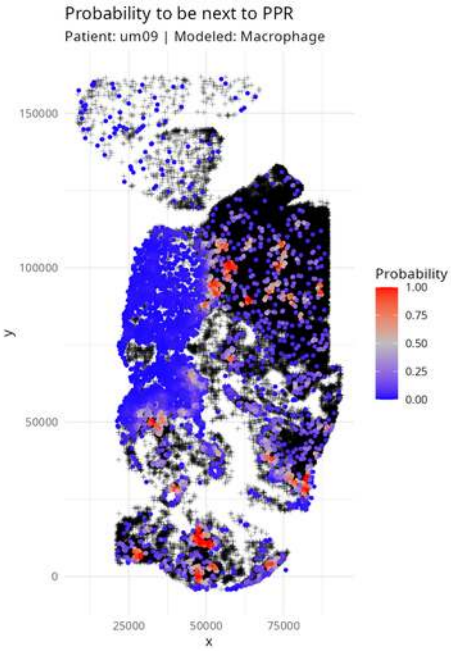
Probability to be next to PPR
Patient: um09 | Modeled: Macrophage
y
x
150000
100000
50000
0
25000
50000
75000
Probability
1.00
0.75
0.50
0.25
0.00

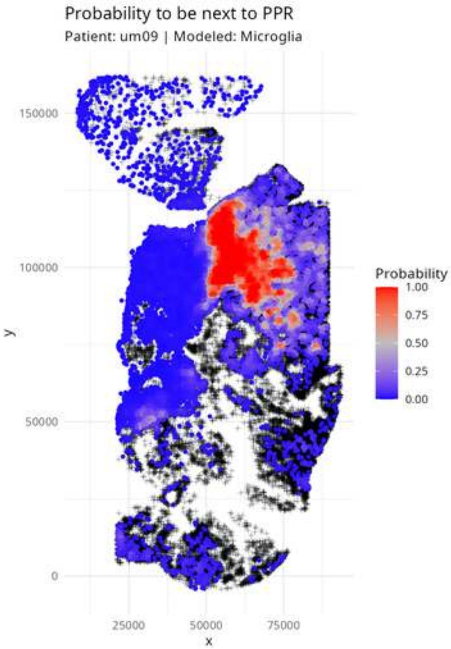

Probability to be next to PPR
Patient: um09 | Modeled: Microglia
150000
100000
50000
0
y
25000
50000
75000
x
Probability
1.00
0.75
0.50
0.25
0.00

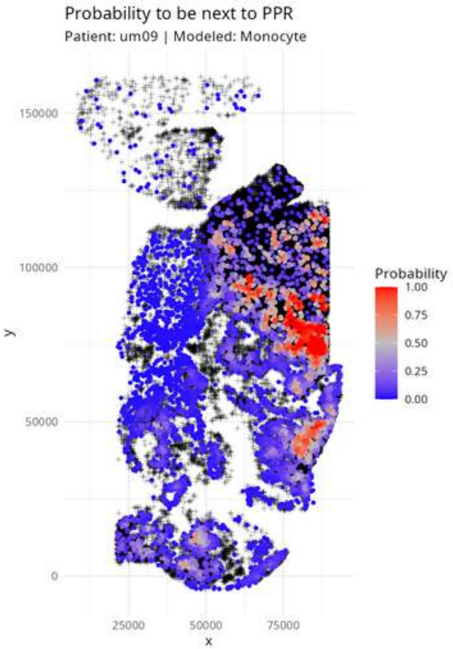

Probability to be next to PPR
Patient: um09 | Modeled: Monocyte
150000
100000
50000
0
y
25000
50000
75000
x
Probability
1.00
0.75
0.50
0.25
0.00

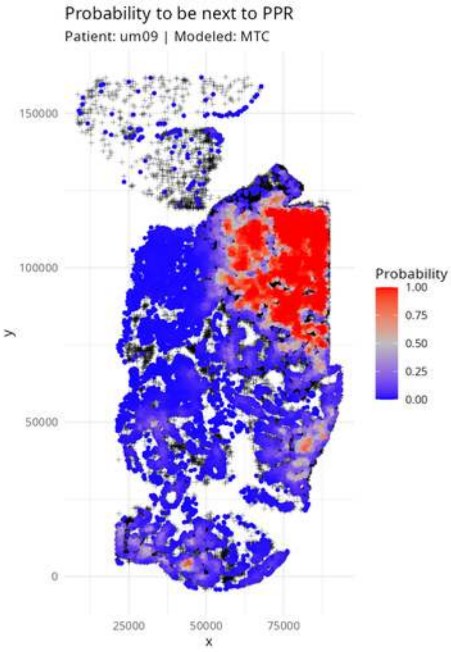
Probability to be next to PPR
Patient: um09 | Modeled: MTC
Probability
1.00
0.75
0.50
0.25
0.00
150000
100000
50000
0
y
25000
50000
75000
x

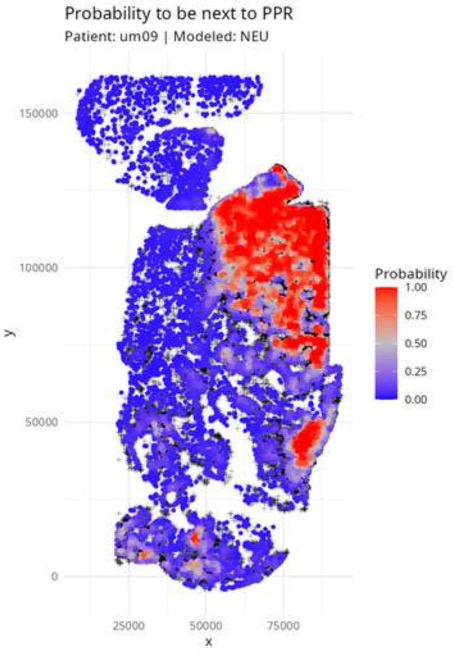

Probability to be next to PPR
Patient: um09 | Modeled: NEU
150000
100000
50000
0
y
25000
50000
75000
x
Probability
1.00
0.75
0.50
0.25
0.00

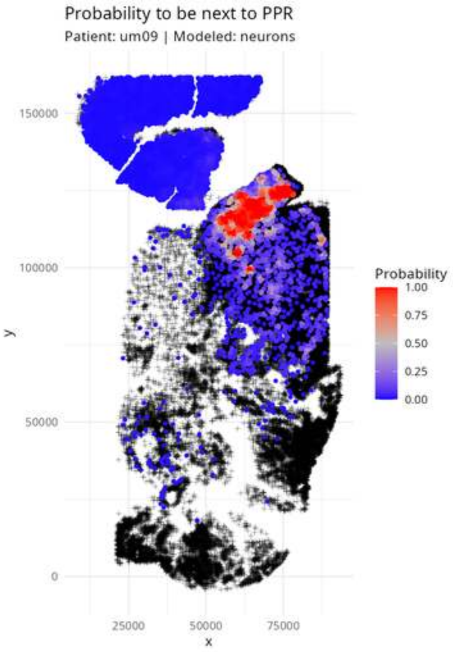

Probability to be next to PPR
Patient: um09 | Modeled: neurons
150000
100000
50000
0
y
25000
50000
75000
x
Probability
1.00
0.75
0.50
0.25
0.00

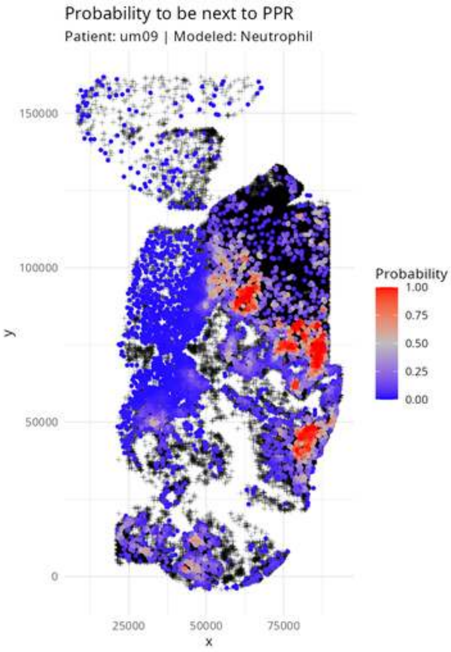

Probability to be next to PPR
Patient: um09 | Modeled: Neutrophil
150000
100000
50000
0
y
25000
50000
75000
x
Probability
1.00
0.75
0.50
0.25
0.00

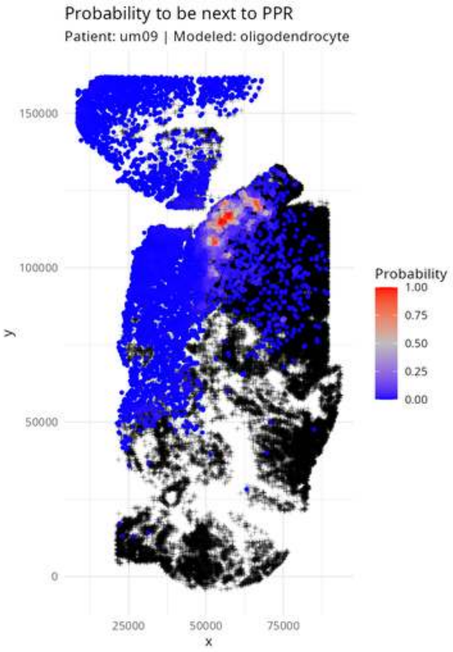
Probability to be next to PPR
Patient: um09 | Modeled: oligodendrocyte
150000
100000
50000
0
y
25000
50000
75000
x
Probability
1.00
0.75
0.50
0.25
0.00

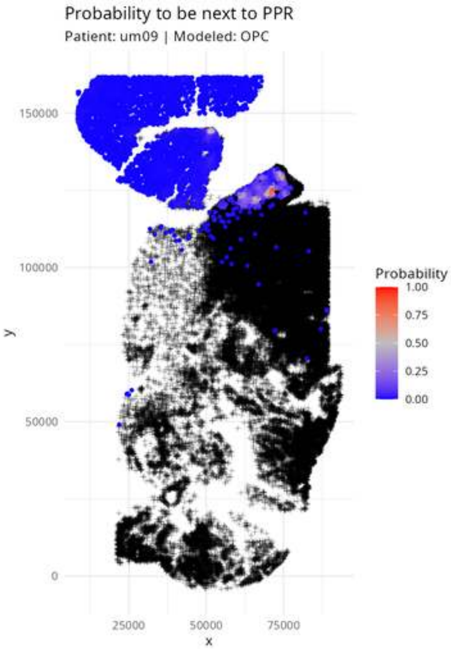

Probability to be next to PPR
Patient: um09 | Modeled: OPC
150000
100000
50000
0
y
25000
50000
75000
x
Probability
1.00
0.75
0.50
0.25
0.00

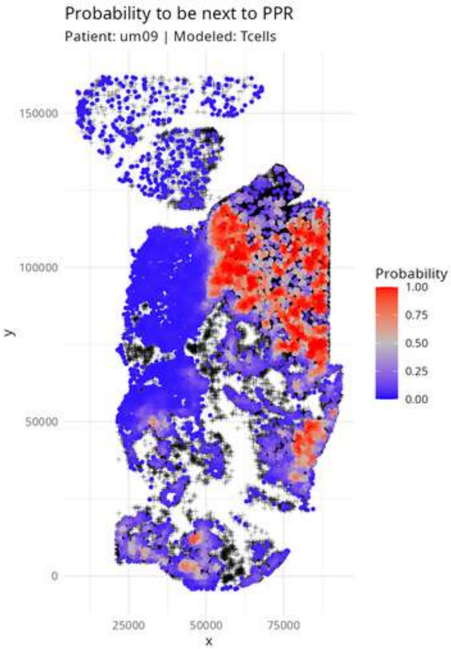
Probability to be next to PPR
Patient: um09 | Modeled: Tcells
150000
100000
50000
0
25000
50000
75000
x
y
Probability
1.00
0.75
0.50
0.25
0.00

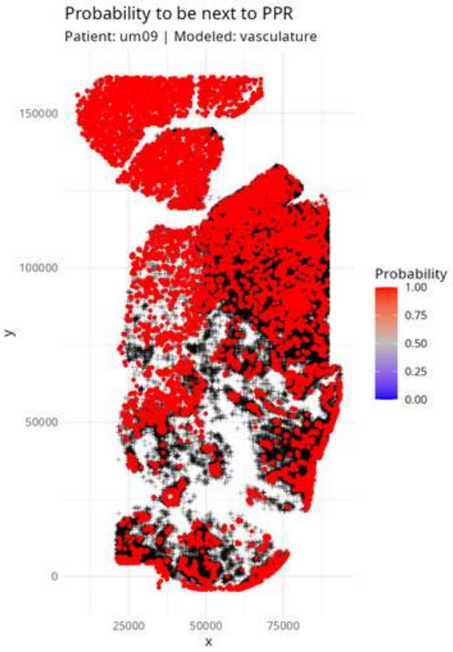

Probability to be next to PPR
Patient: um09 | Modeled: vasculature
150000
100000
50000
0
y
25000
50000
75000
x
Probability
1.00
0.75
0.50
0.25
0.00

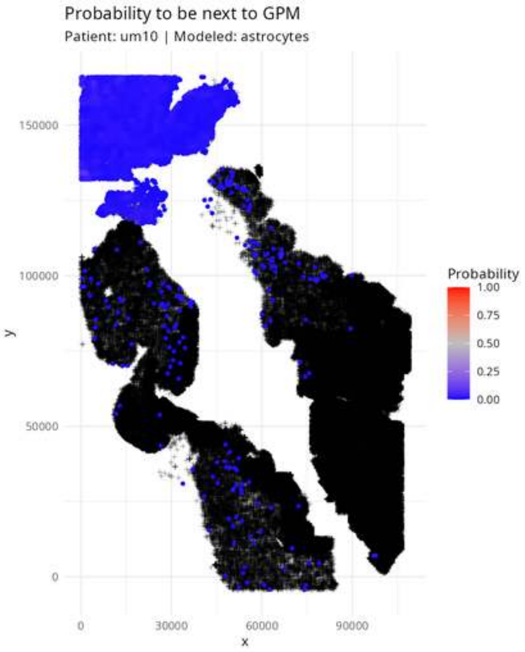

Probability to be next to GPM
Patient: um10 | Modeled: astrocytes
150000
100000
50000
0
y
0
30000
60000
90000
x
Probability
1.00
0.75
0.50
0.25
0.00

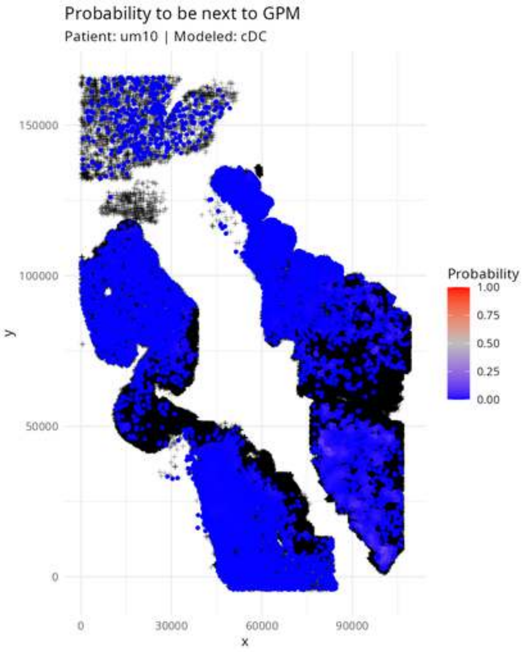

Probability to be next to GPM
Patient: um10 | Modeled: cDC
y
150000
100000
50000
0
0
30000
60000
90000
x
Probability
1.00
0.75
0.50
0.25
0.00

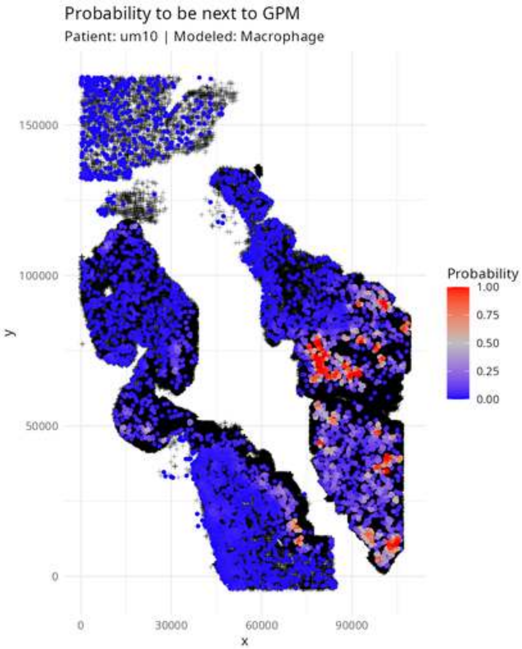

Probability to be next to GPM
Patient: um10 | Modeled: Macrophage
Probability
1.00
0.75
0.50
0.25
0.00
150000
100000
50000
0
y
0
30000
60000
90000
x

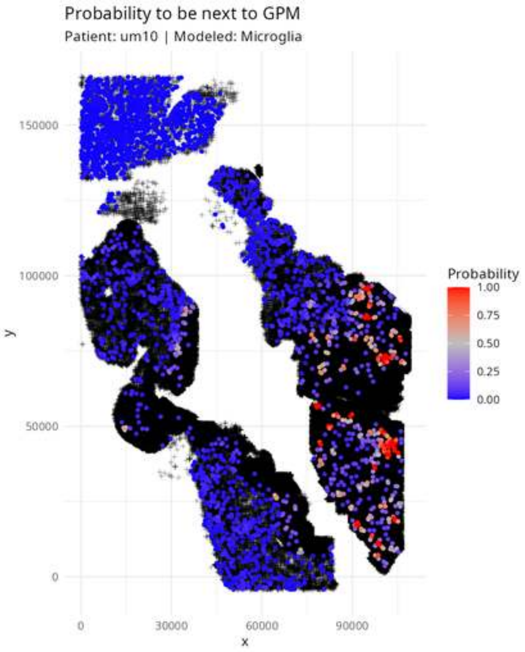
Probability to be next to GPM
Patient: um10 | Modeled: Microglia
150000
100000
50000
0
y
0
30000
60000
90000
x
Probability
1.00
0.75
0.50
0.25
0.00

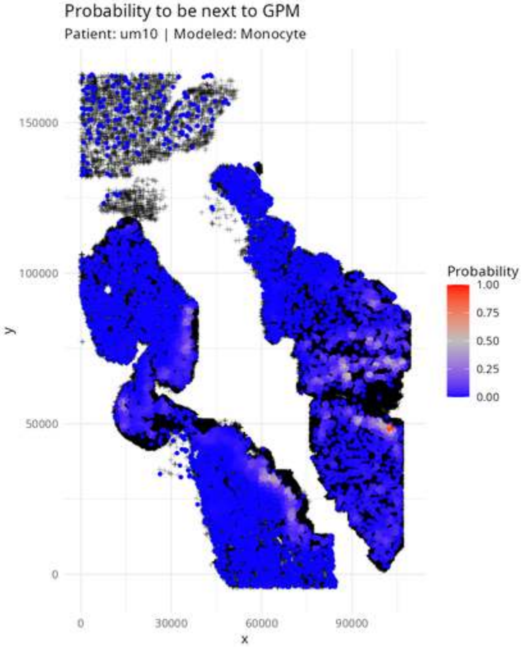

Probability to be next to GPM
Patient: um10 | Modeled: Monocyte
y
150000
100000
50000
0
x
0
30000
60000
90000
Probability
1.00
0.75
0.50
0.25
0.00

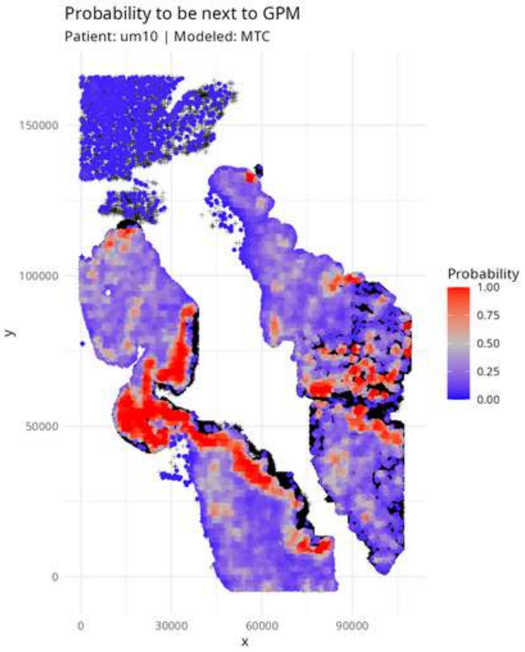
Probability to be next to GPM
Patient: um10 | Modeled: MTC
y
150000
100000
50000
0
x
0
30000
60000
90000
Probability
1.00
0.75
0.50
0.25
0.00

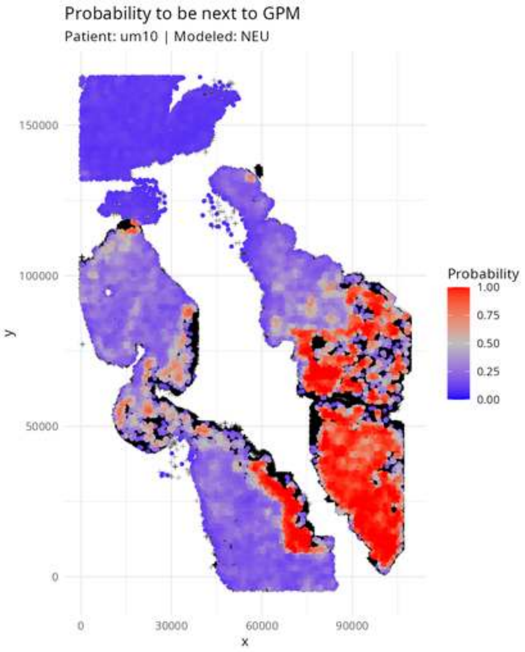

Probability to be next to GPM
Patient: um10 | Modeled: NEU
150000
100000
50000
0
y
0
30000
60000
90000
x
Probability
1.00
0.75
0.50
0.25
0.00

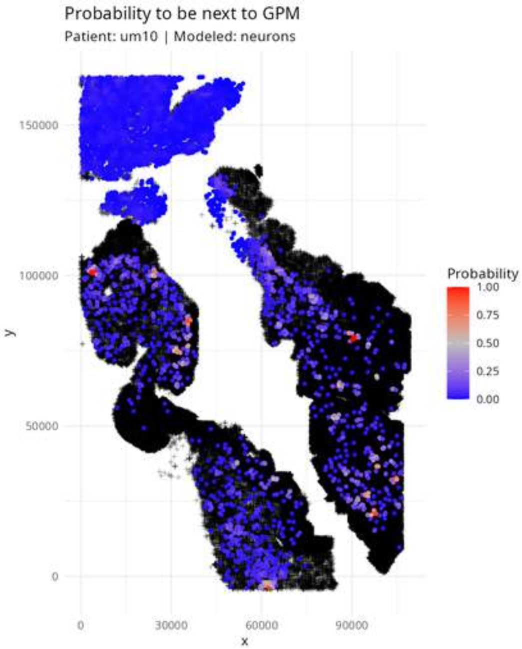

Probability to be next to GPM
Patient: um10 | Modeled: neurons
150000
100000
50000
0
y
0
30000
60000
90000
x
Probability
1.00
0.75
0.50
0.25
0.00

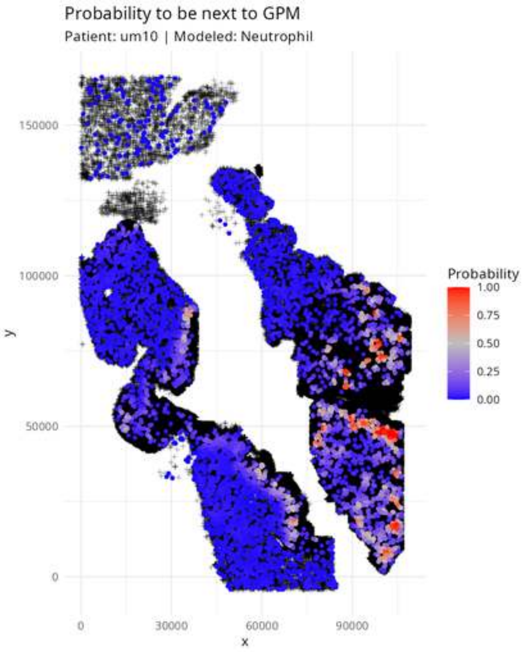

Probability to be next to GPM
Patient: um10 | Modeled: Neutrophil
Probability
1.00
0.75
0.50
0.25
0.00
150000
100000
50000
0
y
0
30000
60000
90000
x

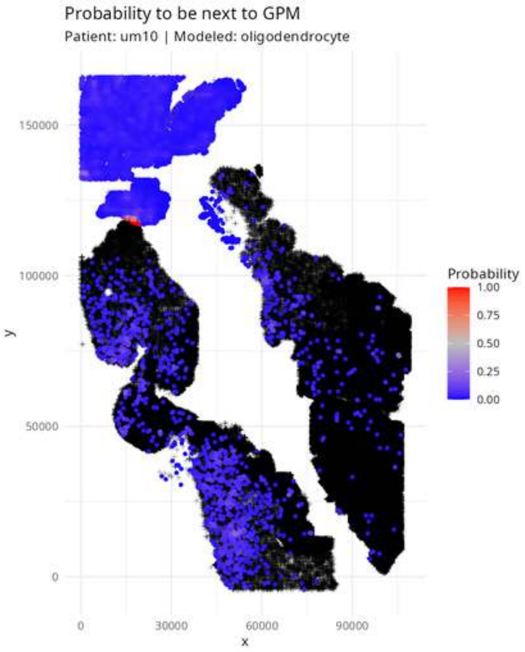

Probability to be next to GPM
Patient: um10 | Modeled: oligodendrocyte
y
150000
100000
50000
0
x
0
30000
60000
90000
Probability
1.00
0.75
0.50
0.25
0.00

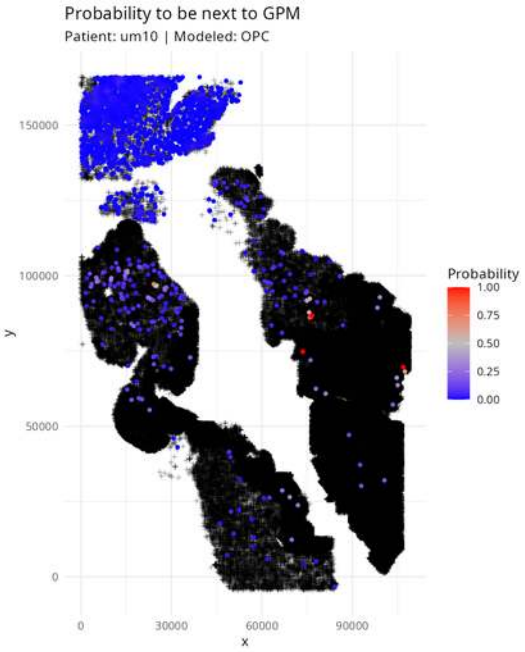

Probability to be next to GPM
Patient: um10 | Modeled: OPC
150000
100000
50000
0
y
0
30000
60000
90000
x
Probability
1.00
0.75
0.50
0.25
0.00

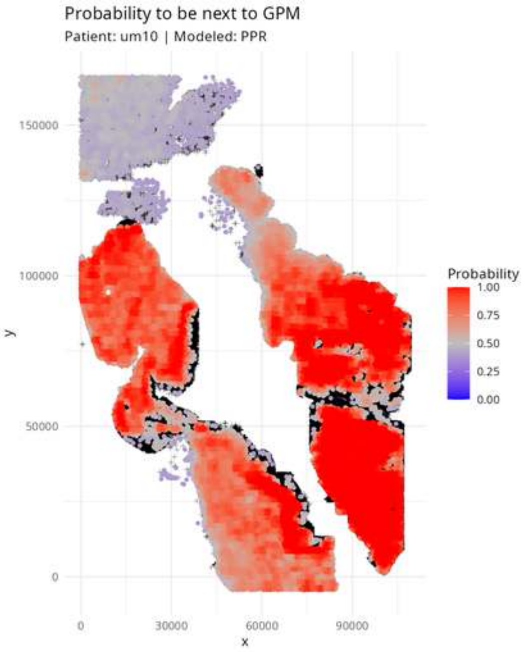

Probability to be next to GPM
Patient: um10 | Modeled: PPR
150000
100000
50000
0
y
0
30000
60000
90000
x
Probability
1.00
0.75
0.50
0.25
0.00

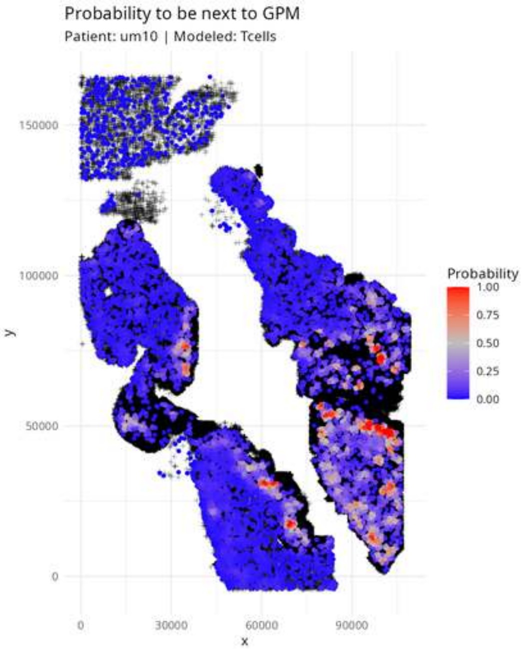

Probability to be next to GPM
Patient: um10 | Modeled: Tcells
y
150000
100000
50000
0
x
0
30000
60000
90000
Probability
1.00
0.75
0.50
0.25
0.00

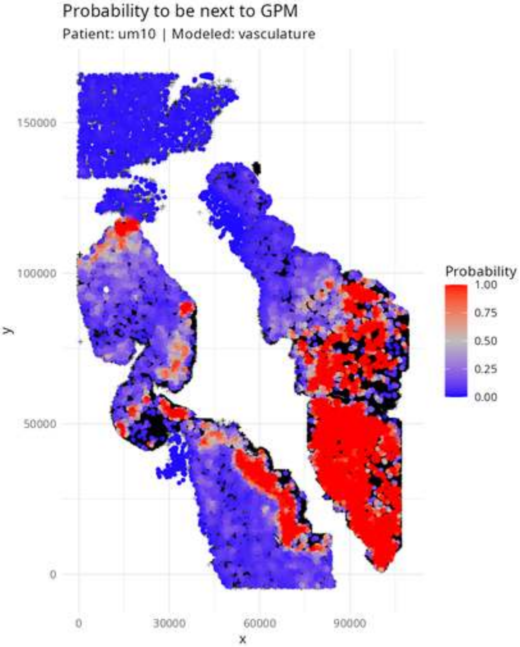
Probability to be next to GPM
Patient: um10 | Modeled: vasculature
Probability
1.00
0.75
0.50
0.25
0.00
150000
100000
50000
0
y
0
30000
60000
90000
x

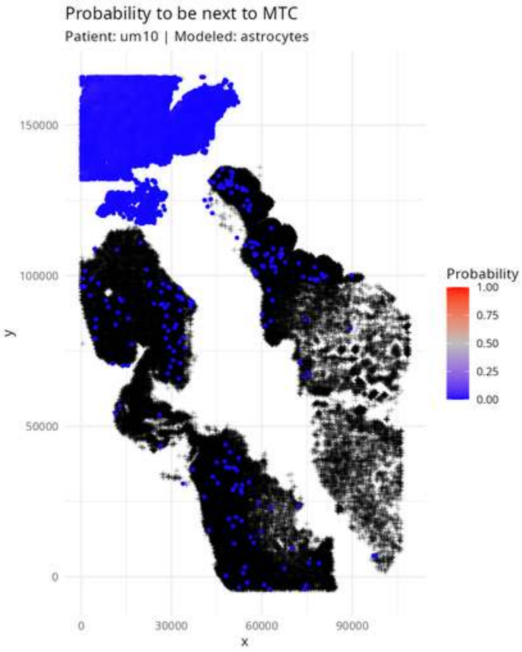
Probability to be next to MTC
Patient: um10 | Modeled: astrocytes
150000
100000
50000
0
y
0
30000
60000
90000
x
Probability
1.00
0.75
0.50
0.25
0.00

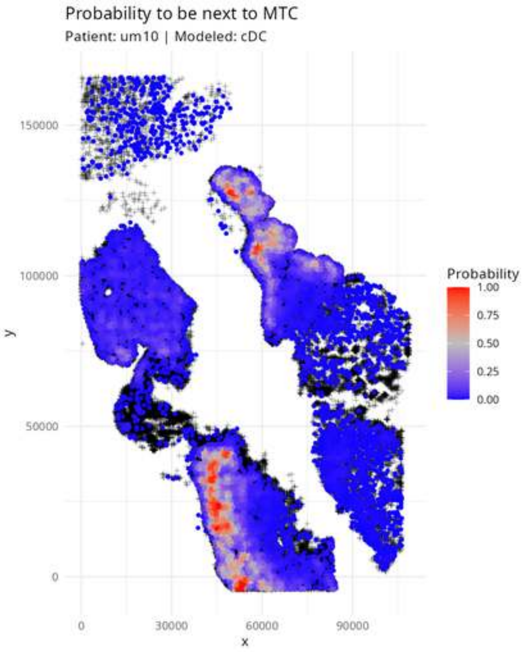
Probability to be next to MTC
Patient: um10 | Modeled: cDC
150000
100000
50000
0
y
0
30000
60000
90000
x
Probability
1.00
0.75
0.50
0.25
0.00

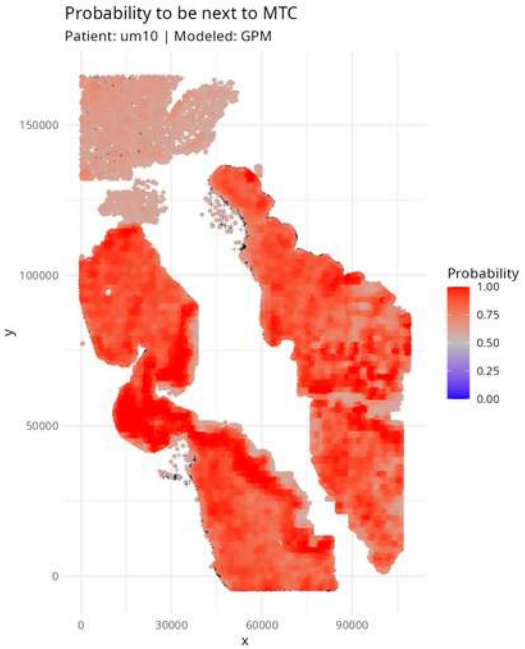

Probability to be next to MTC
Patient: um10 | Modeled: GPM
Probability
1.00
0.75
0.50
0.25
0.00
150000
100000
50000
0
y
0
30000
60000
90000
x

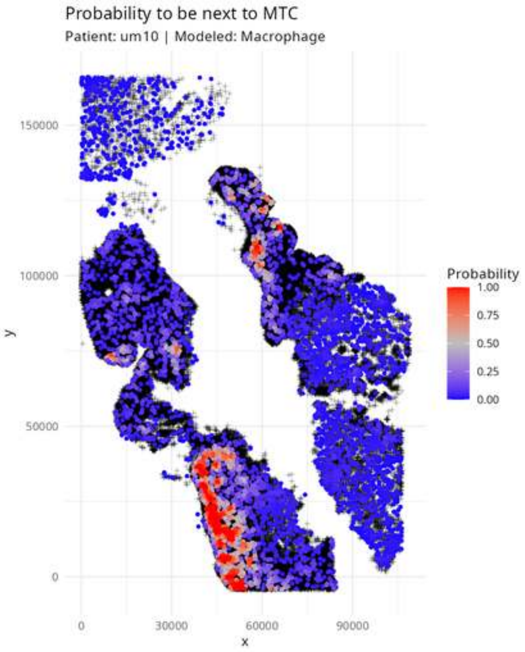
Probability to be next to MTC
Patient: um10 | Modeled: Macrophage
150000
100000
50000
0
y
0
30000
60000
90000
x
Probability
1.00
0.75
0.50
0.25
0.00

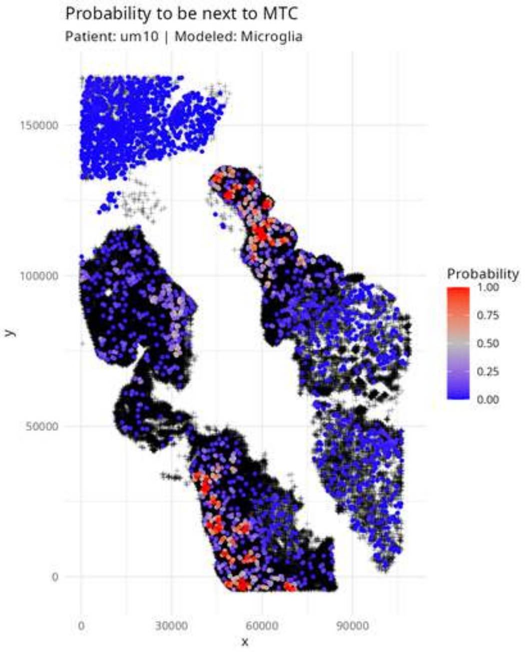

Probability to be next to MTC
Patient: um10 | Modeled: Microglia
Probability
1.00
0.75
0.50
0.25
0.00
150000
100000
50000
0
y
0
30000
60000
90000
x

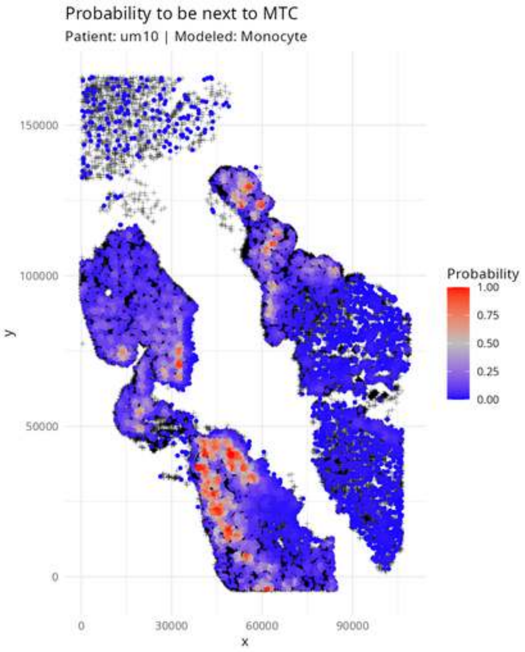

Probability to be next to MTC
Patient: um10 | Modeled: Monocyte
Probability
1.00
0.75
0.50
0.25
0.00
150000
100000
50000
0
y
0
30000
60000
90000
x

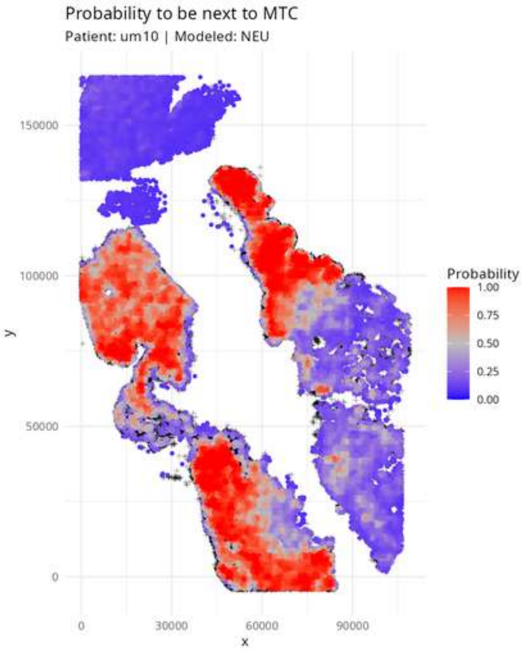

Probability to be next to MTC
Patient: um10 | Modeled: NEU
y
x
150000
100000
50000
0
0
30000
60000
90000
Probability
1.00
0.75
0.50
0.25
0.00

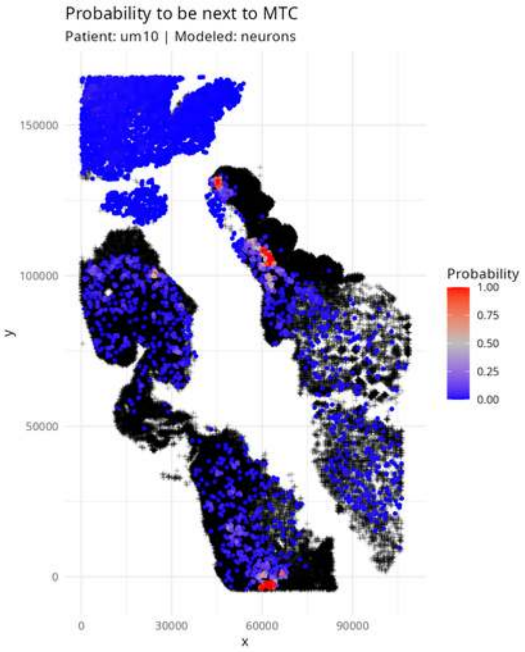
Probability to be next to MTC
Patient: um10 | Modeled: neurons
150000
100000
50000
0
y
0
30000
60000
90000
x
Probability
1.00
0.75
0.50
0.25
0.00

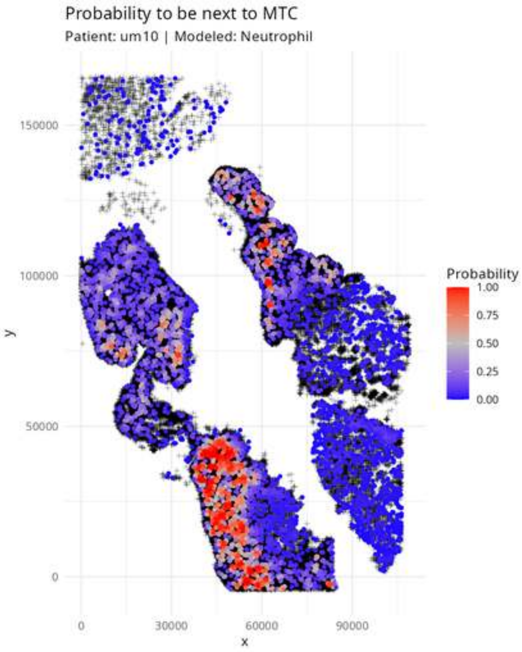

Probability to be next to MTC
Patient: um10 | Modeled: Neutrophil
y
150000
100000
50000
0
x
0
30000
60000
90000
Probability
1.00
0.75
0.50
0.25
0.00

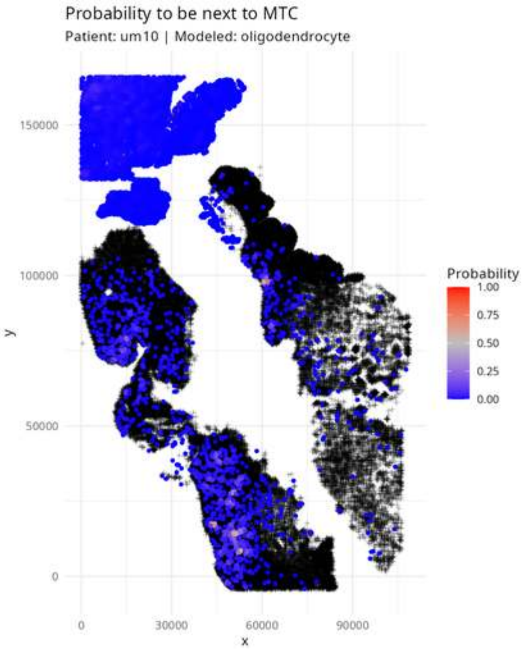

Probability to be next to MTC
Patient: um10 | Modeled: oligodendrocyte
y
150000
100000
50000
0
x
0
30000
60000
90000
Probability
1.00
0.75
0.50
0.25
0.00

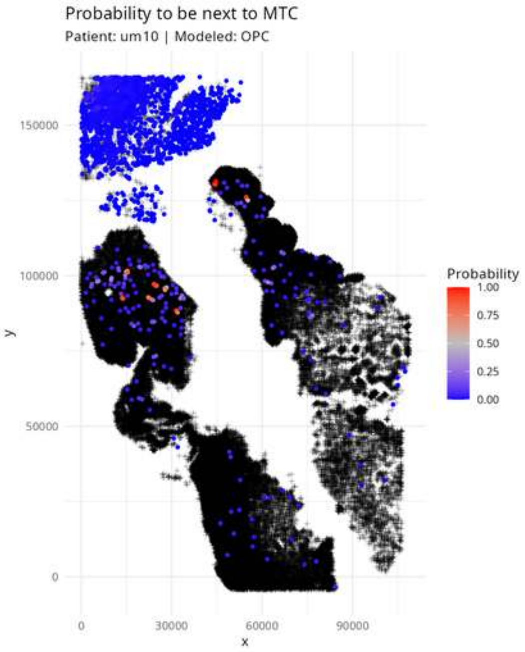

Probability to be next to MTC
Patient: um10 | Modeled: OPC
y
x
150000
100000
50000
0
0
30000
60000
90000
Probability
1.00
0.75
0.50
0.25
0.00

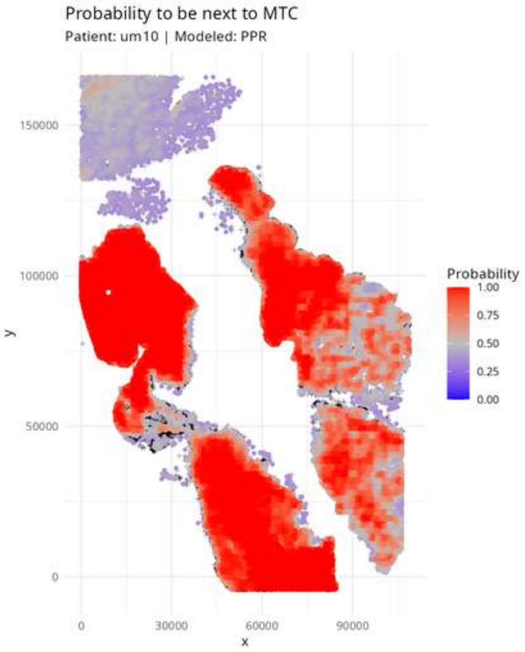

Probability to be next to MTC
Patient: um10 | Modeled: PPR
150000
100000
50000
0
y
0
30000
60000
90000
x
Probability
1.00
0.75
0.50
0.25
0.00

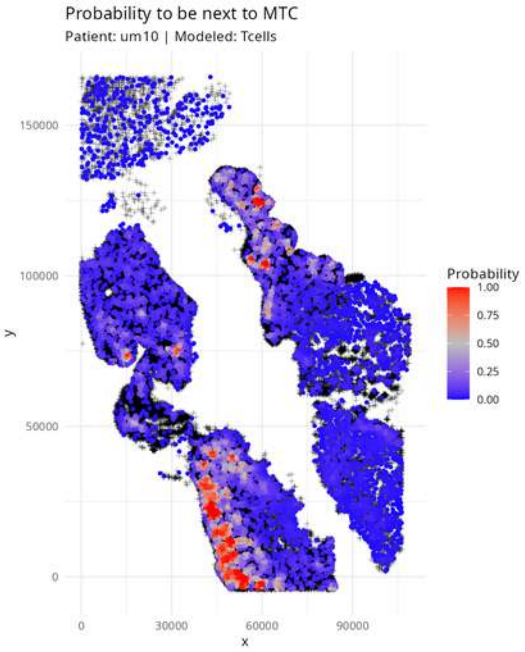

Probability to be next to MTC
Patient: um10 | Modeled: Tcells
150000
100000
50000
0
y
0
30000
60000
90000
x
Probability
1.00
0.75
0.50
0.25
0.00

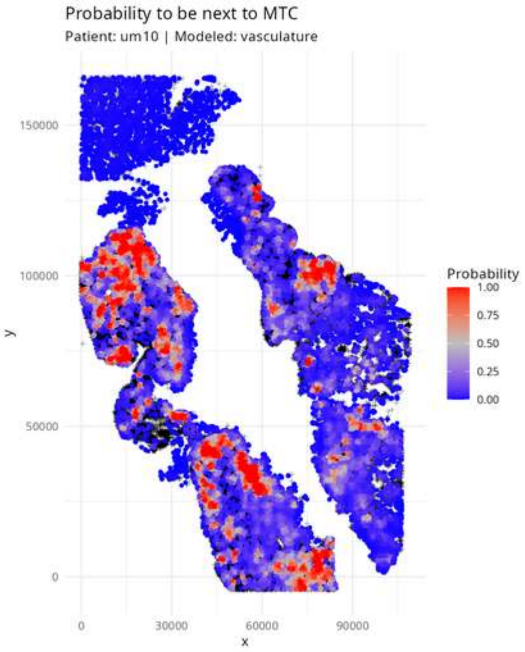

Probability to be next to MTC
Patient: um10 | Modeled: vasculature
Probability
1.00
0.75
0.50
0.25
0.00
150000
100000
50000
0
y
0
30000
60000
90000
x

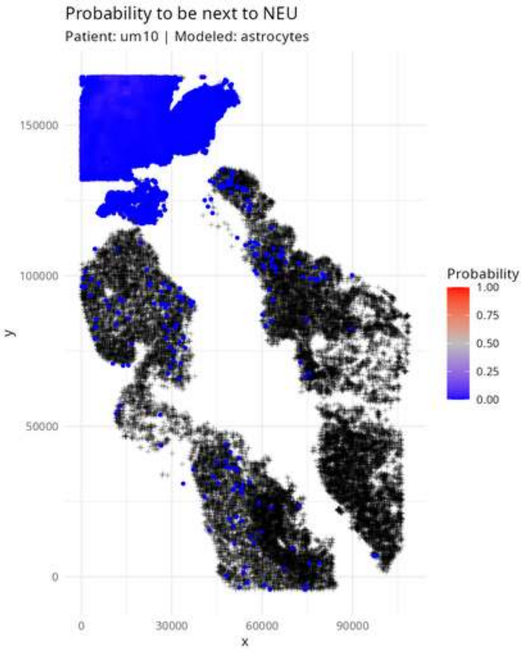

Probability to be next to NEU
Patient: um10 | Modeled: astrocytes
150000
100000
50000
0
y
0
30000
60000
90000
x
Probability
1.00
0.75
0.50
0.25
0.00

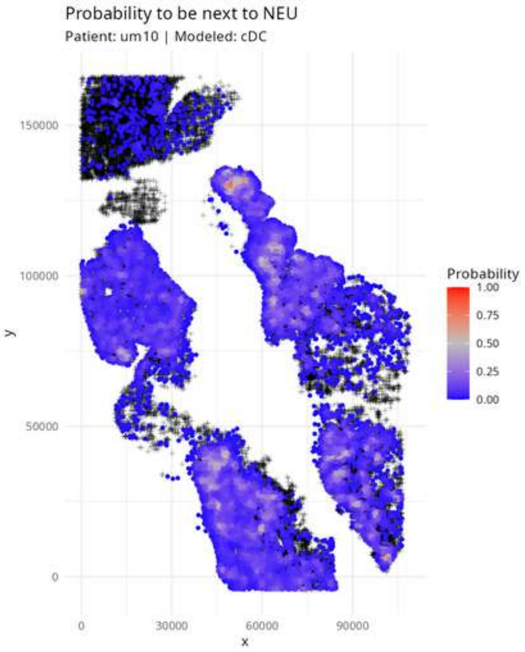

Probability to be next to NEU
Patient: um10 | Modeled: cDC
y
150000
100000
50000
0
x
0
30000
60000
90000
Probability
1.00
0.75
0.50
0.25
0.00

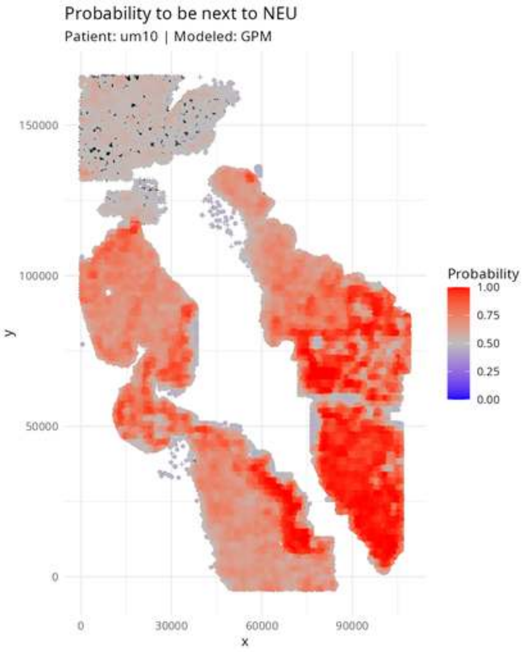
Probability to be next to NEU
Patient: um10 | Modeled: GPM
150000
100000
50000
0
y
0
30000
60000
90000
x
Probability
1.00
0.75
0.50
0.25
0.00

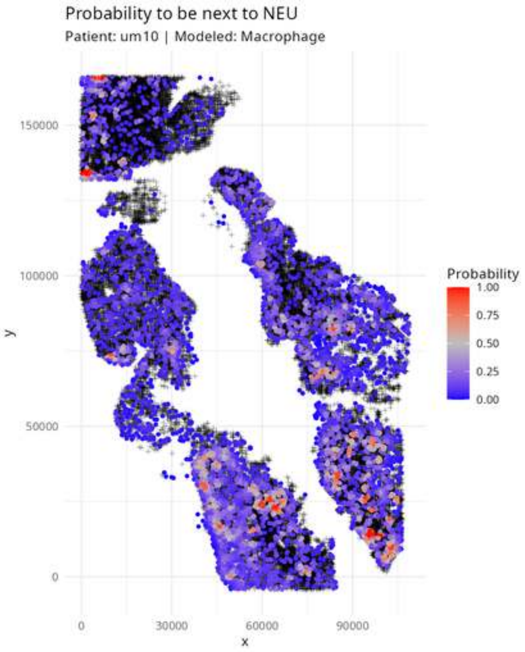

Probability to be next to NEU
Patient: um10 | Modeled: Macrophage
150000
100000
50000
0
y
0
30000
60000
90000
x
Probability
1.00
0.75
0.50
0.25
0.00

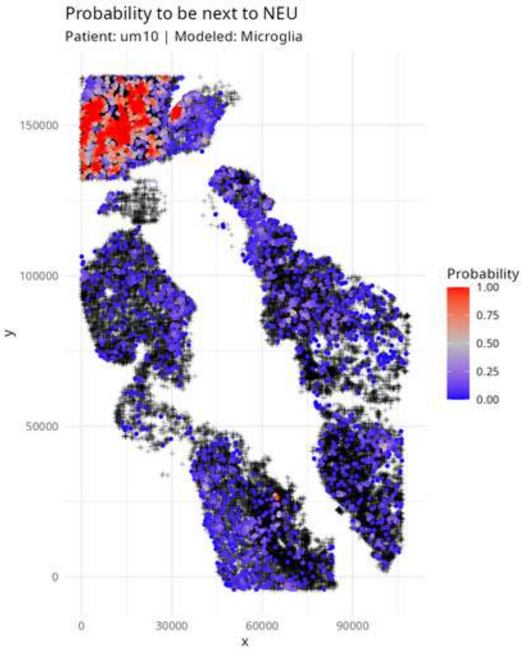

Probability to be next to NEU
Patient: um10 | Modeled: Microglia
150000
100000
50000
0
y
0
30000
60000
90000
x
Probability
1.00
0.75
0.50
0.25
0.00

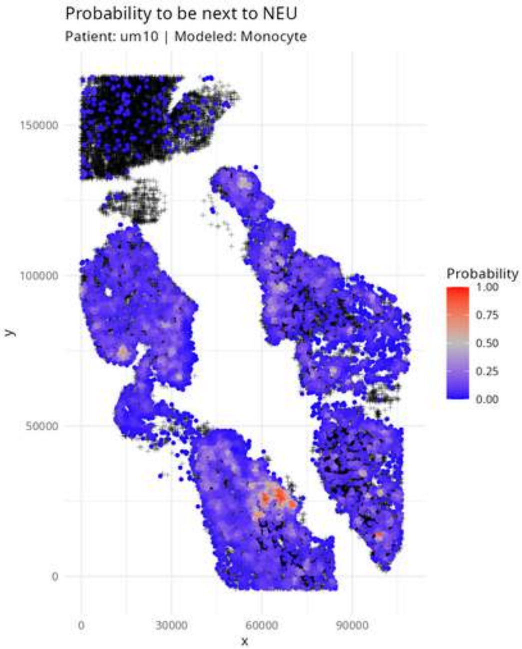
Probability to be next to NEU
Patient: um10 | Modeled: Monocyte
150000
100000
50000
0
y
0
30000
60000
90000
x
Probability
1.00
0.75
0.50
0.25
0.00

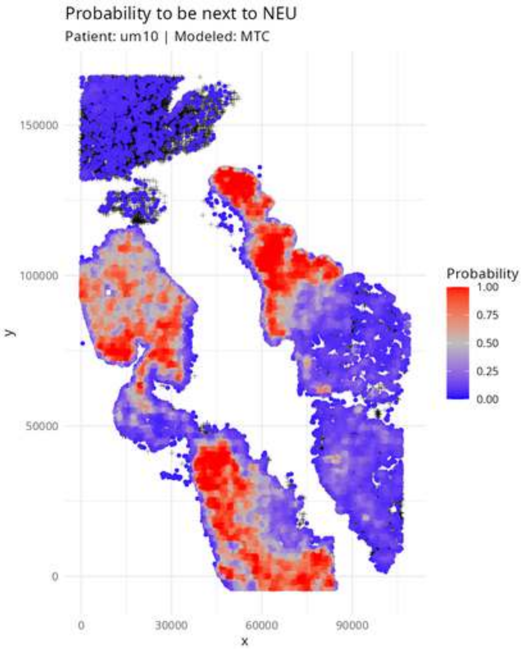

Probability to be next to NEU
Patient: um10 | Modeled: MTC
Probability
1.00
0.75
0.50
0.25
0.00
150000
100000
50000
0
y
0
30000
60000
90000
x

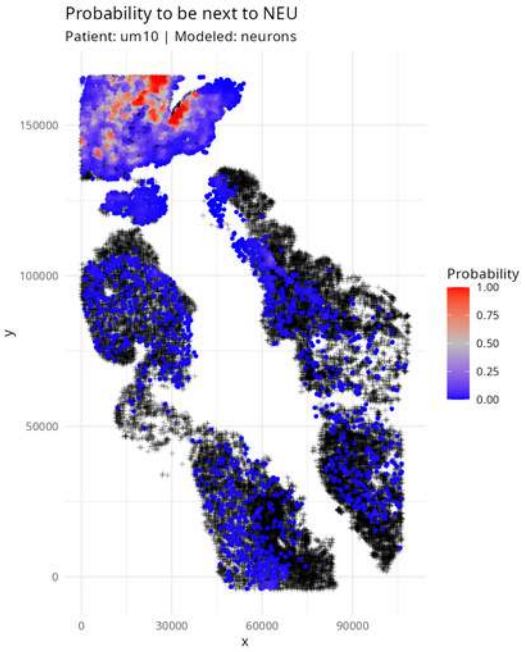

Probability to be next to NEU
Patient: um10 | Modeled: neurons
Probability
1.00
0.75
0.50
0.25
0.00
y
150000
100000
50000
0
x
0
30000
60000
90000

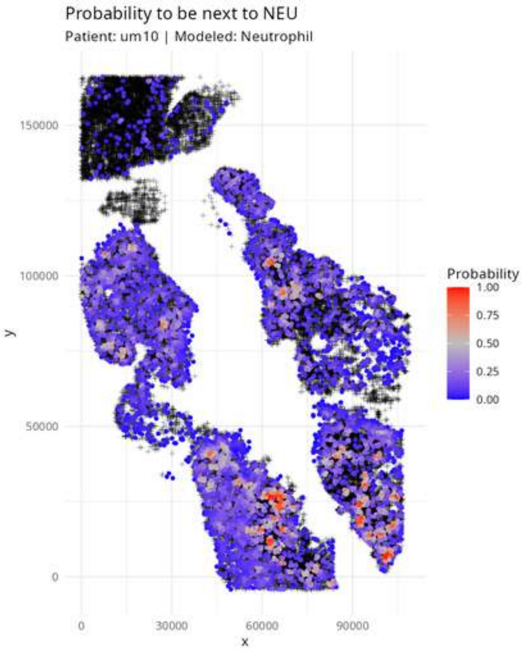

Probability to be next to NEU
Patient: um10 | Modeled: Neutrophil
150000
100000
50000
0
y
0
30000
60000
90000
x
Probability
1.00
0.75
0.50
0.25
0.00

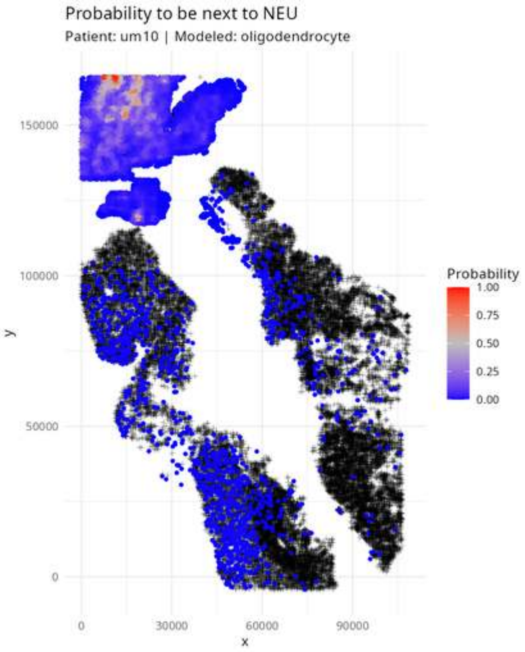

Probability to be next to NEU
Patient: um10 | Modeled: oligodendrocyte
150000
100000
50000
0
y
0
30000
60000
90000
x
Probability
1.00
0.75
0.50
0.25
0.00

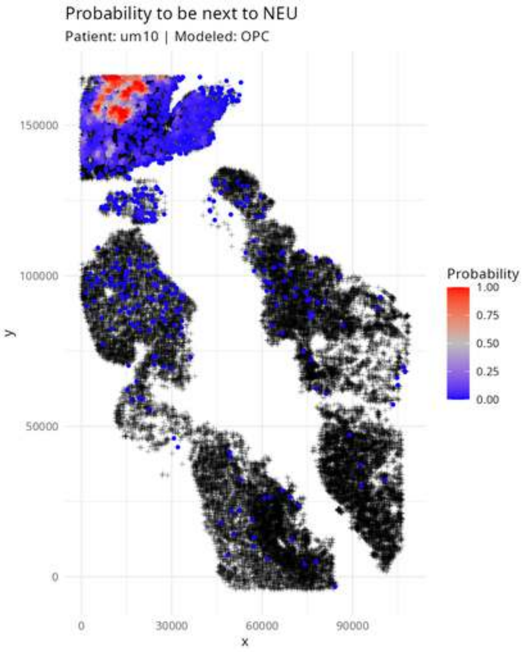
Probability to be next to NEU
Patient: um10 | Modeled: OPC
150000
100000
50000
0
0
30000
60000
90000
x
y
Probability
1.00
0.75
0.50
0.25
0.00

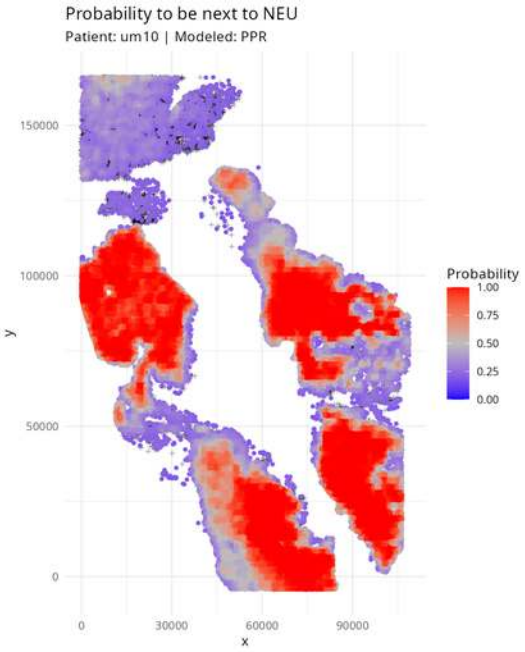

Probability to be next to NEU
Patient: um10 | Modeled: PPR
Probability
1.00
0.75
0.50
0.25
0.00
150000
100000
50000
0
y
0
30000
60000
90000
x

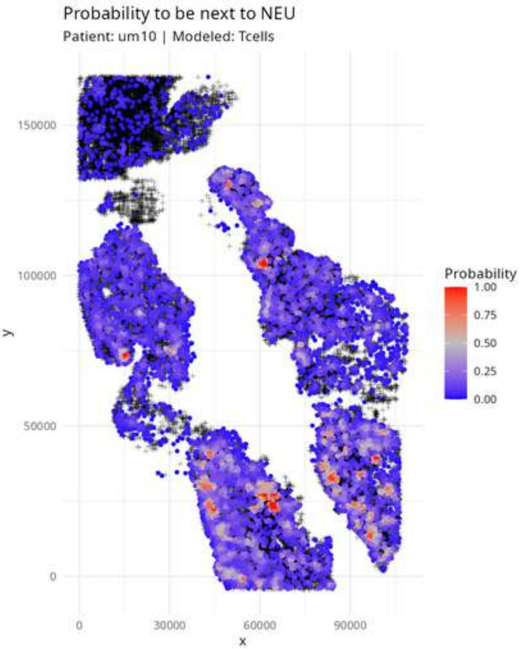
Probability to be next to NEU
Patient: um10 | Modeled: Tcells
150000
100000
50000
0
0
30000
60000
90000
x
y
Probability
1.00
0.75
0.50
0.25
0.00

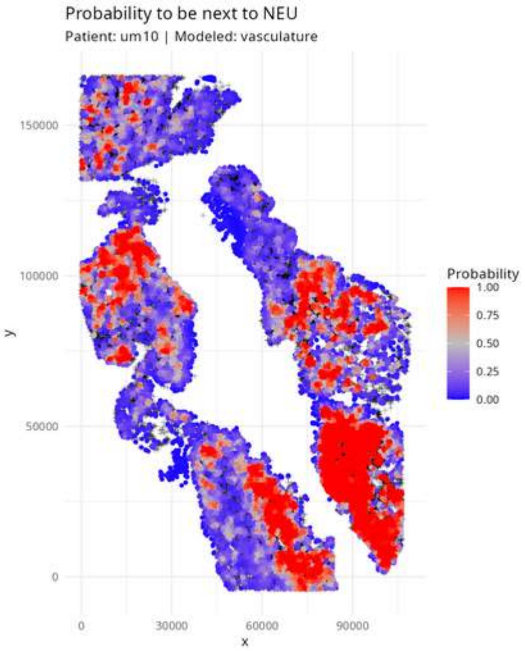
Probability to be next to NEU
Patient: um10 | Modeled: vasculature
y
150000
100000
50000
0
x
0
30000
60000
90000
Probability
1.00
0.75
0.50
0.25
0.00

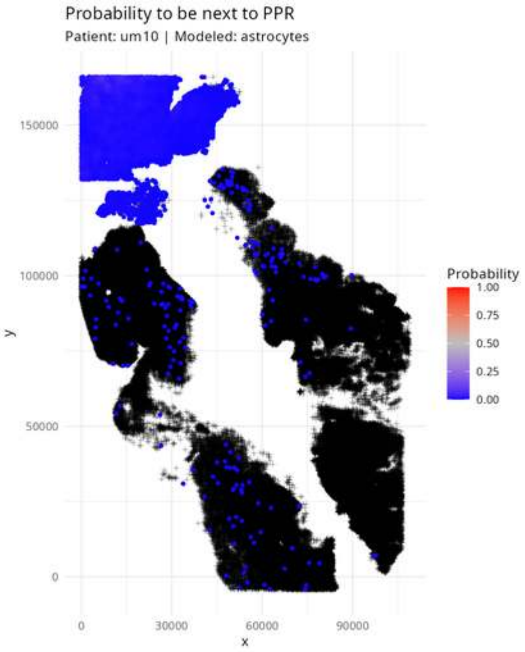
Probability to be next to PPR
Patient: um10 | Modeled: astrocytes
150000
100000
50000
0
y
0
30000
60000
90000
x
Probability
1.00
0.75
0.50
0.25
0.00

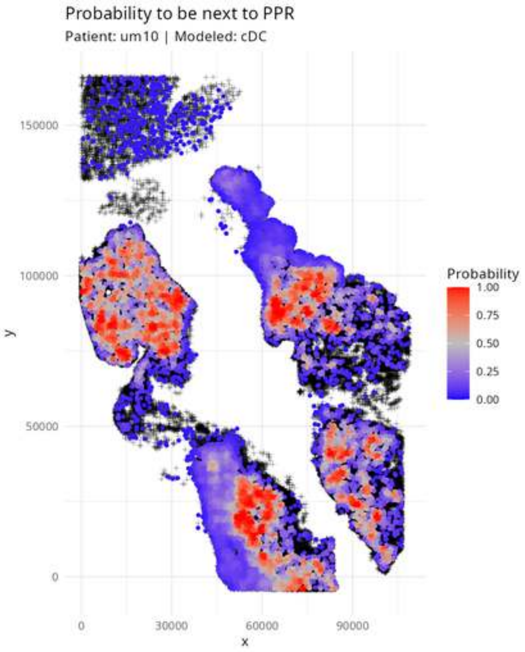

Probability to be next to PPR
Patient: um10 | Modeled: cDC
Probability
1.00
0.75
0.50
0.25
0.00
y
150000
100000
50000
0
x
0
30000
60000
90000

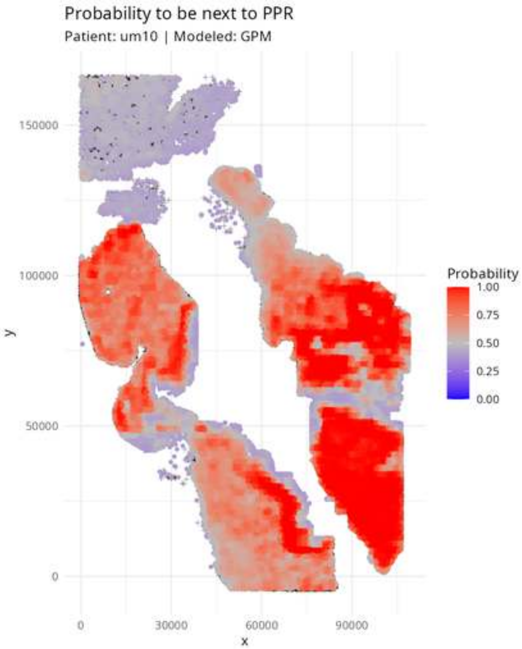

Probability to be next to PPR
Patient: um10 | Modeled: GPM
150000
100000
50000
0
y
0
30000
60000
90000
x
Probability
1.00
0.75
0.50
0.25
0.00

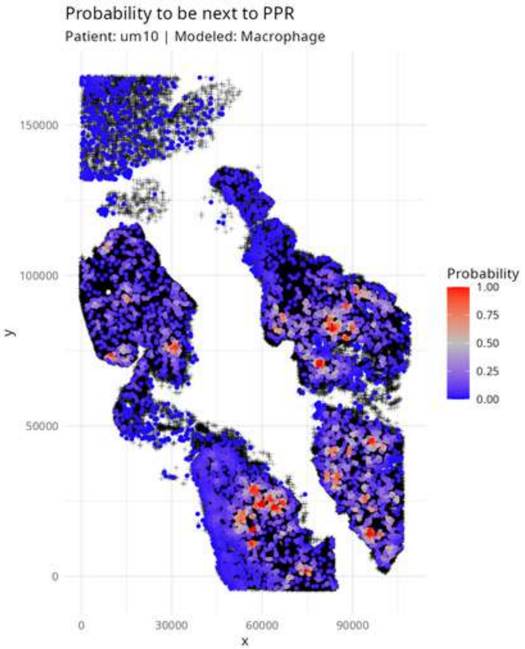
Probability to be next to PPR
Patient: um10 | Modeled: Macrophage
150000
100000
50000
0
y
0
30000
60000
90000
x
Probability
1.00
0.75
0.50
0.25
0.00

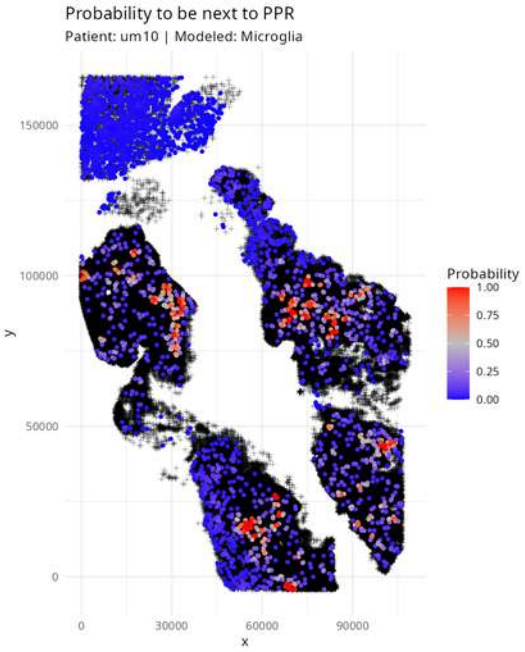

Probability to be next to PPR
Patient: um10 | Modeled: Microglia
Probability
1.00
0.75
0.50
0.25
0.00
y
150000
100000
50000
0
x
0
30000
60000
90000

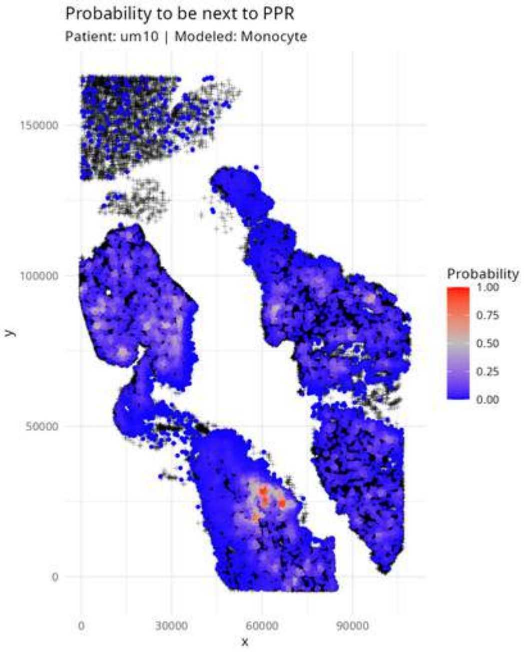

Probability to be next to PPR
Patient: um10 | Modeled: Monocyte
y
150000
100000
50000
0
x
0
30000
60000
90000
Probability
1.00
0.75
0.50
0.25
0.00

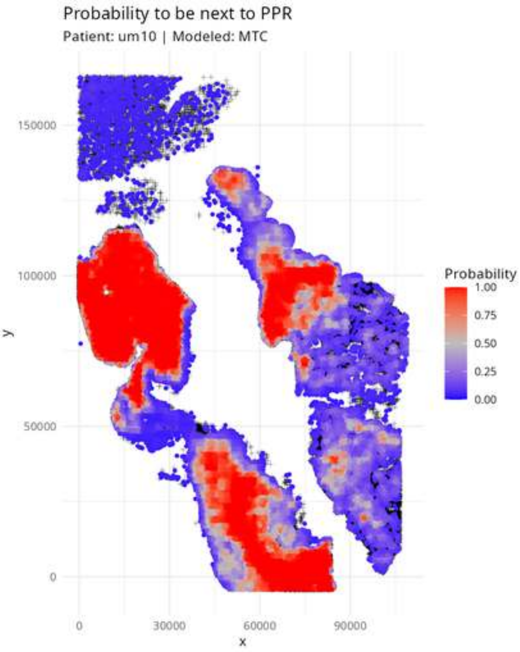
Probability to be next to PPR
Patient: um10 | Modeled: MTC
150000
100000
50000
0
y
0
30000
60000
90000
x
Probability
1.00
0.75
0.50
0.25
0.00

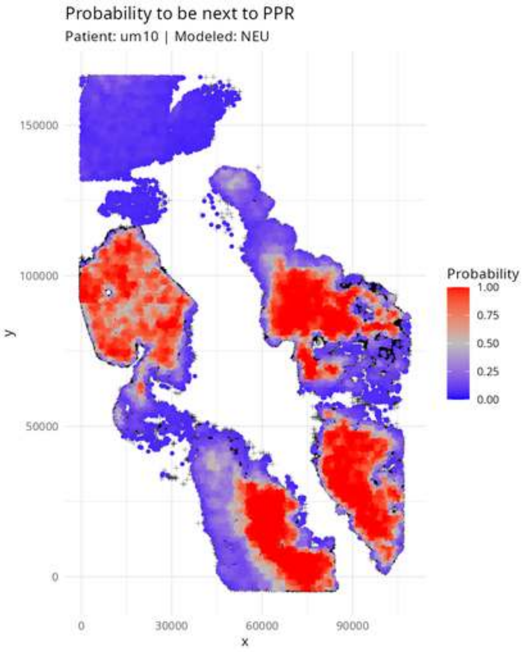
Probability to be next to PPR
Patient: um10 | Modeled: NEU
150000
100000
50000
0
y
0
30000
60000
90000
x
Probability
1.00
0.75
0.50
0.25
0.00

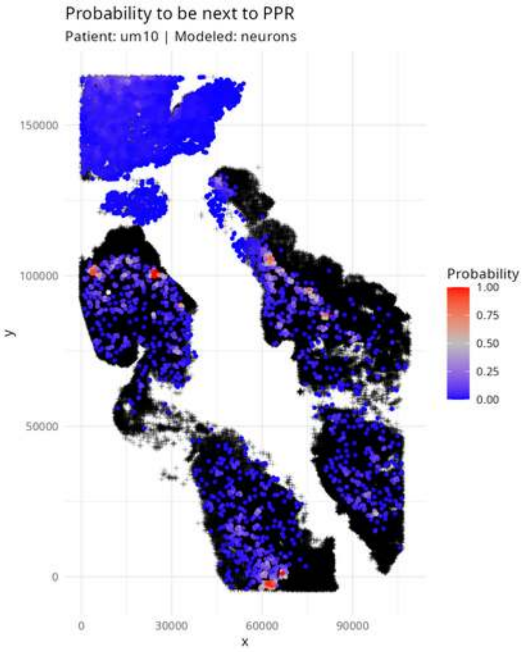

Probability to be next to PPR
Patient: um10 | Modeled: neurons
150000
100000
50000
0
y
0
30000
60000
90000
x
Probability
1.00
0.75
0.50
0.25
0.00

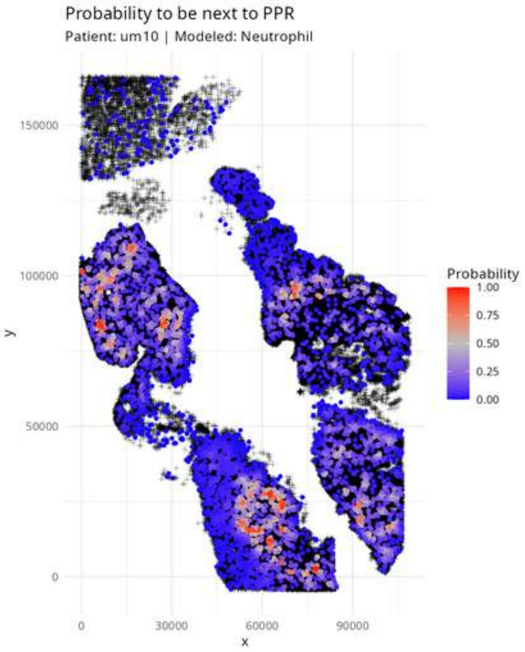

Probability to be next to PPR
Patient: um10 | Modeled: Neutrophil
150000
100000
50000
0
y
0
30000
60000
90000
x
Probability
1.00
0.75
0.50
0.25
0.00

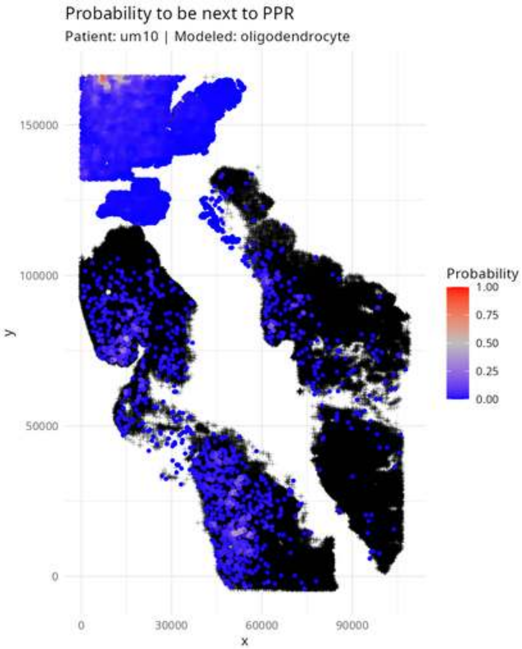

Probability to be next to PPR
Patient: um10 | Modeled: oligodendrocyte
y
150000
100000
50000
0
x
0
30000
60000
90000
Probability
1.00
0.75
0.50
0.25
0.00

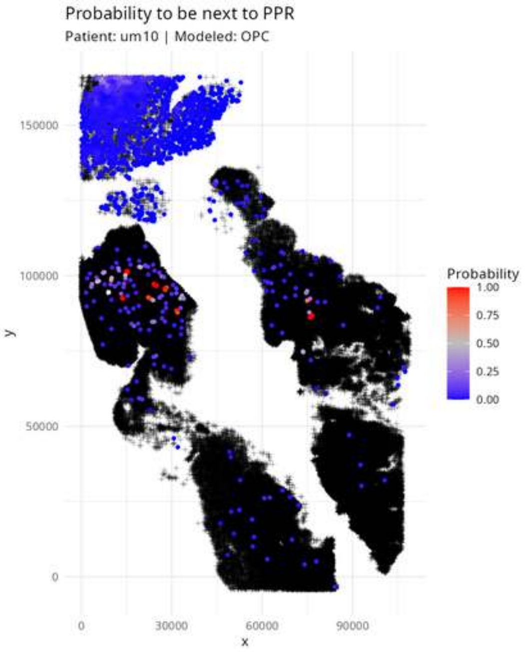

Probability to be next to PPR
Patient: um10 | Modeled: OPC
150000
100000
50000
0
y
0
30000
60000
90000
x
Probability
1.00
0.75
0.50
0.25
0.00

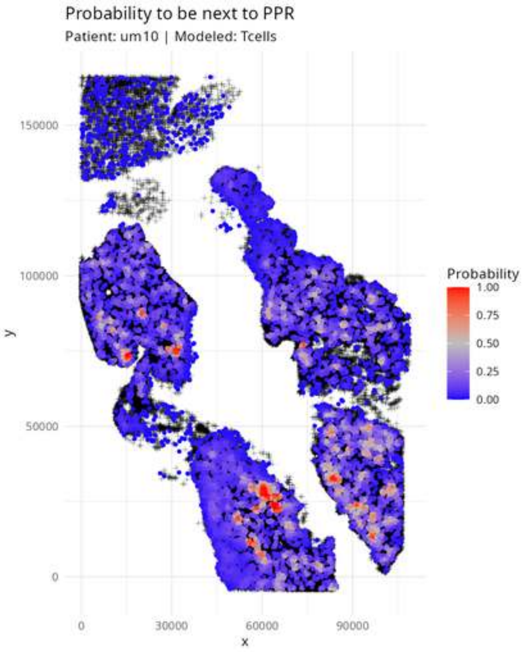

Probability to be next to PPR
Patient: um10 | Modeled: Tcells
y
x
0
50000
100000
150000
0
30000
60000
90000
Probability
1.00
0.75
0.50
0.25
0.00

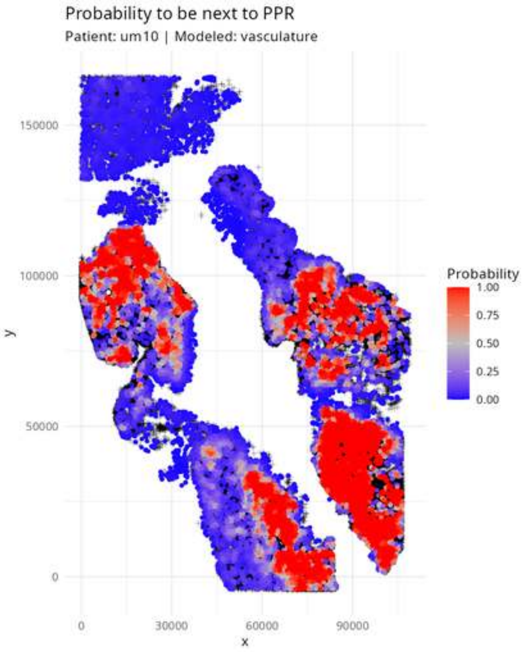

Probability to be next to PPR
Patient: um10 | Modeled: vasculature
Probability
1.00
0.75
0.50
0.25
0.00
150000
100000
50000
0
y
0
30000
60000
90000
x

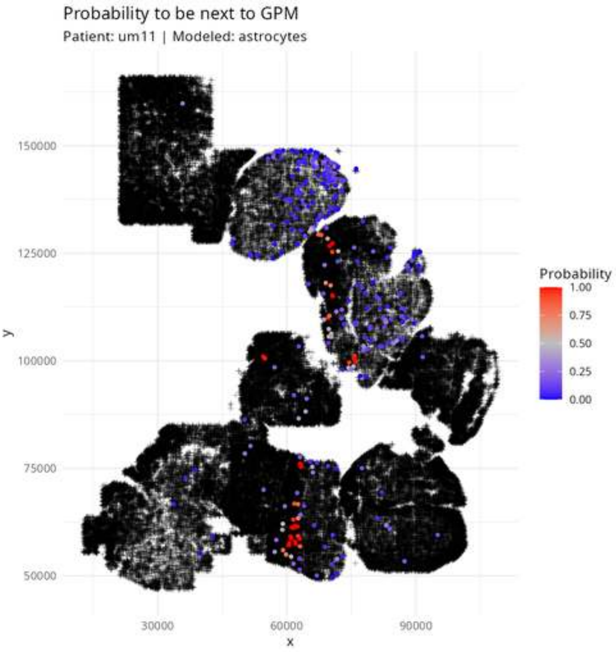

Probability to be next to GPM
Patient: um11 | Modeled: astrocytes
150000
125000
100000
75000
50000
y
30000
60000
90000
x
Probability
1.00
0.75
0.50
0.25
0.00

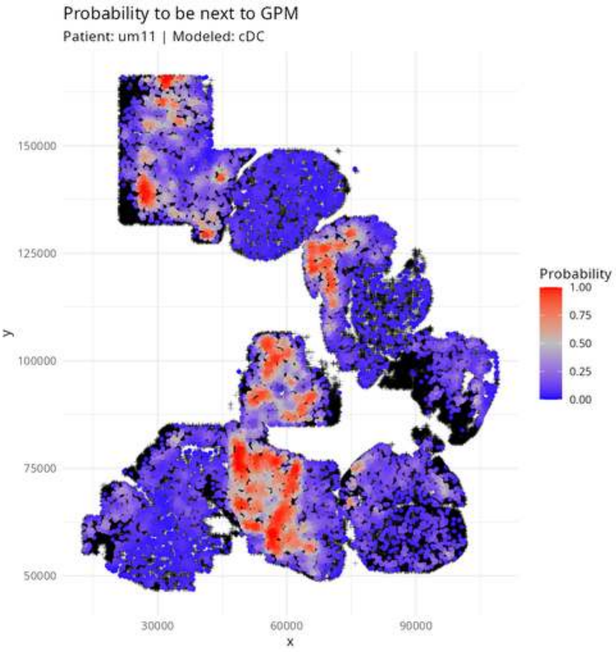
Probability to be next to GPM
Patient: um11 | Modeled: cDC
150000
125000
100000
75000
50000
y
30000
60000
90000
x
Probability
1.00
0.75
0.50
0.25
0.00

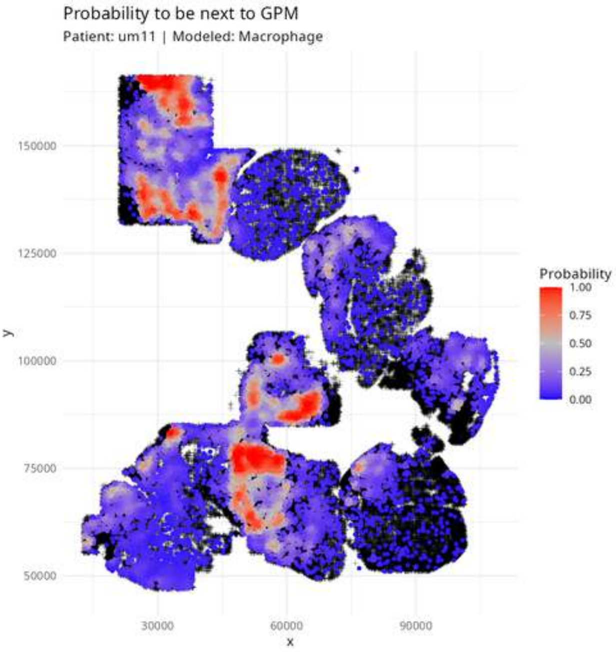
Probability to be next to GPM
Patient: um11 | Modeled: Macrophage
Probability
1.00
0.75
0.50
0.25
0.00
y
150000
125000
100000
75000
50000
30000
60000
90000
x

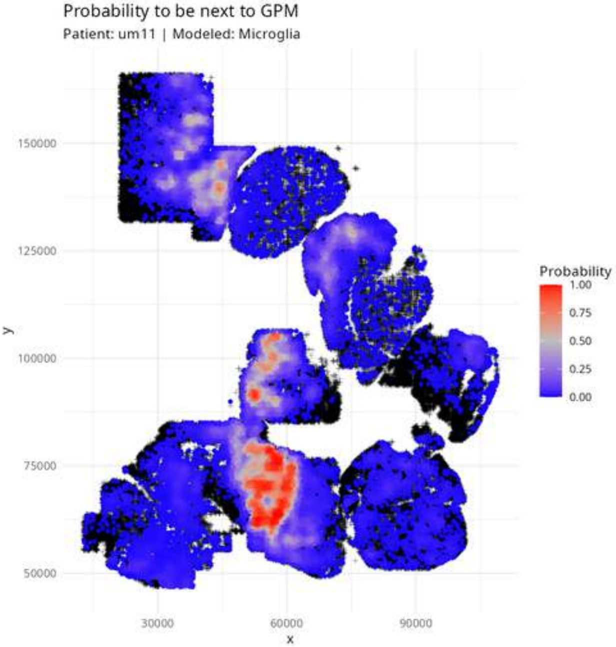

Probability to be next to GPM
Patient: um11 | Modeled: Microglia
Probability
1.00
0.75
0.50
0.25
0.00
150000
125000
100000
75000
50000
y
30000
60000
90000
x

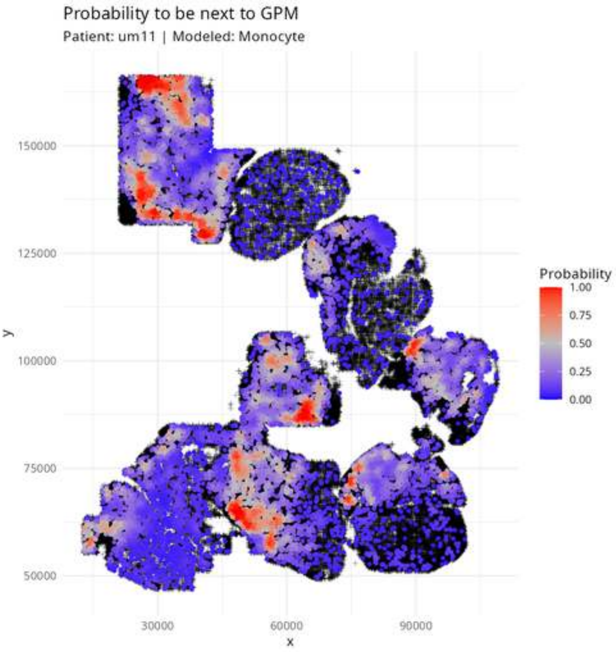
Probability to be next to GPM
Patient: um11 | Modeled: Monocyte
150000
125000
100000
75000
50000
y
30000
60000
90000
x
Probability
1.00
0.75
0.50
0.25
0.00

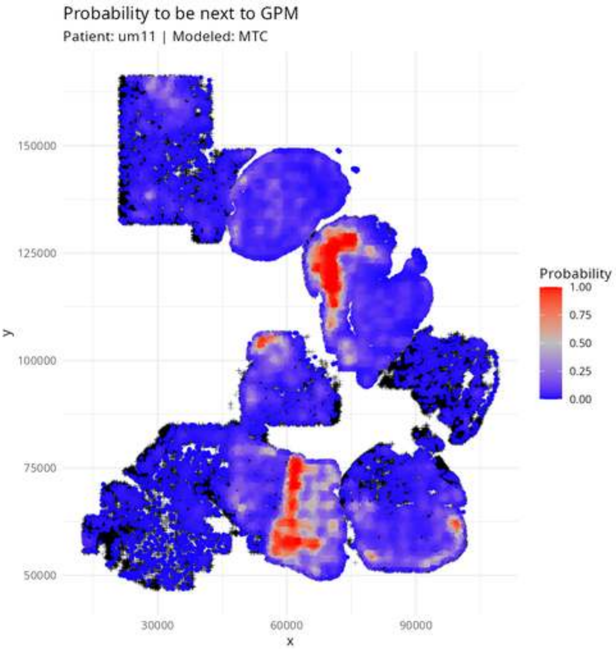

Probability to be next to GPM
Patient: um11 | Modeled: MTC
y
150000
125000
100000
75000
50000
30000
60000
90000
x
Probability
1.00
0.75
0.50
0.25
0.00

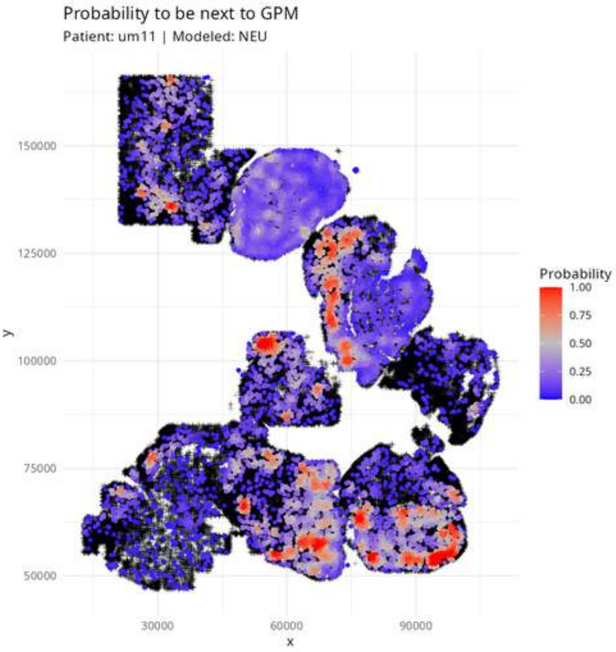

Probability to be next to GPM
Patient: um11 | Modeled: NEU
150000
125000
100000
75000
50000
y
30000
60000
90000
x
Probability
1.00
0.75
0.50
0.25
0.00

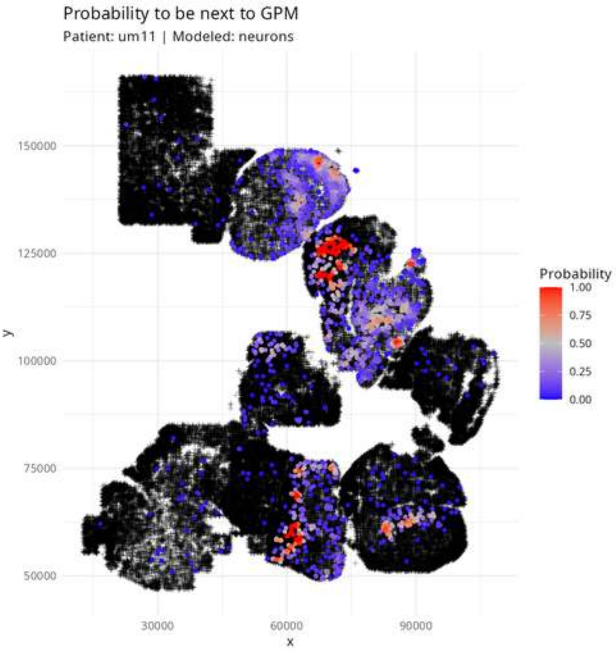

Probability to be next to GPM
Patient: um11 | Modeled: neurons
y
150000
125000
100000
75000
50000
x
30000
60000
90000
Probability
1.00
0.75
0.50
0.25
0.00

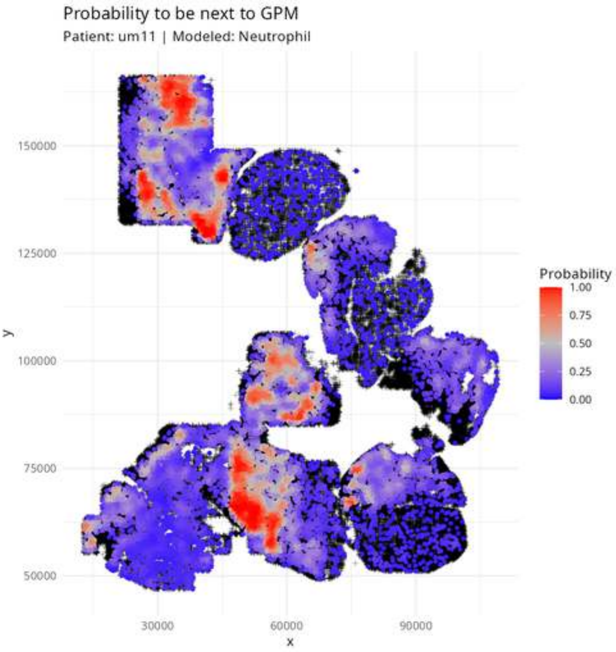

Probability to be next to GPM
Patient: um11 | Modeled: Neutrophil
Probability
1.00
0.75
0.50
0.25
0.00
150000
125000
100000
75000
50000
y
30000
60000
90000
x

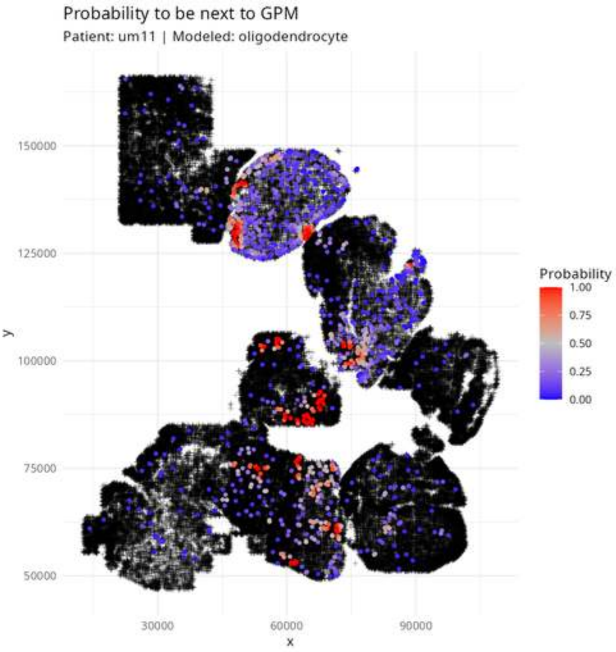

Probability to be next to GPM
Patient: um11 | Modeled: oligodendrocyte
y
150000
125000
100000
75000
50000
x
30000
60000
90000
Probability
1.00
0.75
0.50
0.25
0.00

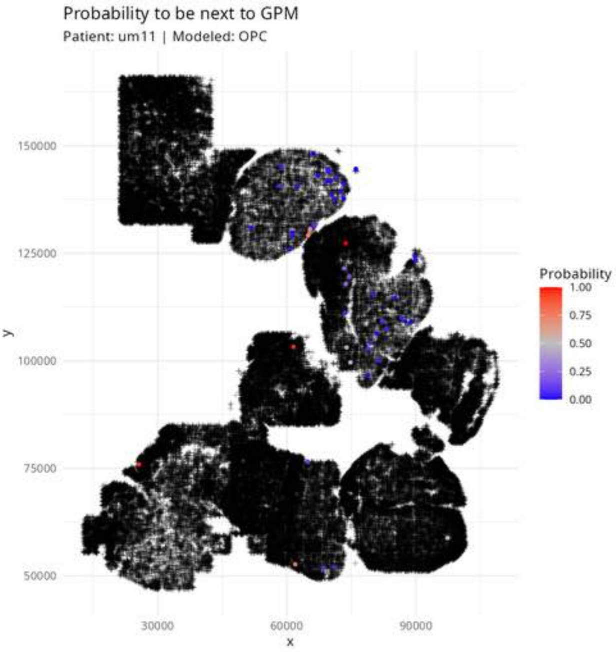

Probability to be next to GPM
Patient: um11 | Modeled: OPC
y
150000
125000
100000
75000
50000
x
30000
60000
90000
Probability
1.00
0.75
0.50
0.25
0.00

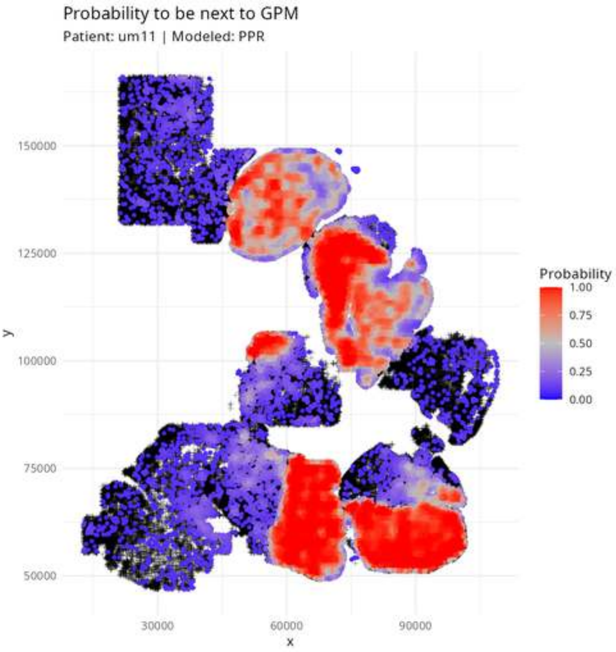

Probability to be next to GPM
Patient: um11 | Modeled: PPR
150000
125000
100000
75000
50000
y
30000
60000
90000
x
Probability
1.00
0.75
0.50
0.25
0.00

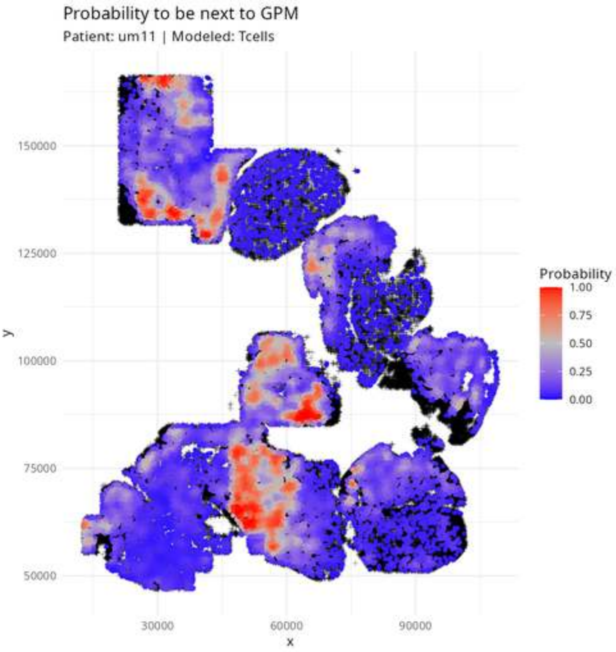

Probability to be next to GPM
Patient: um11 | Modeled: Tcells
150000
125000
100000
75000
50000
y
30000
60000
90000
x
Probability
1.00
0.75
0.50
0.25
0.00

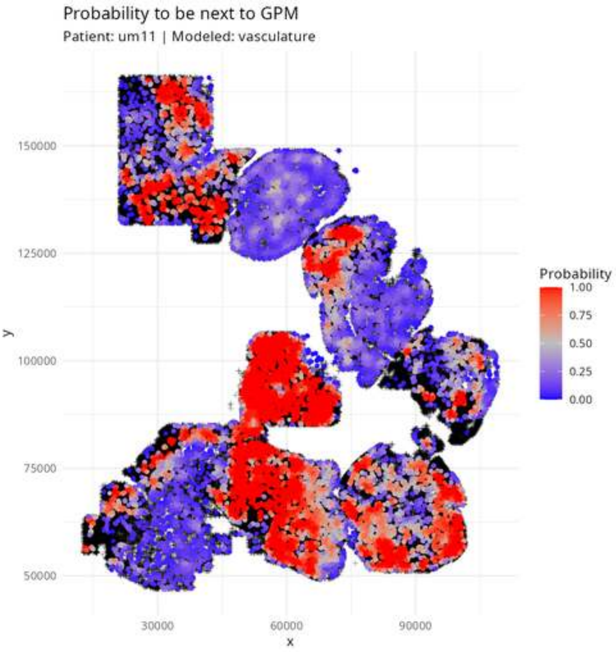
Probability to be next to GPM
Patient: um11 | Modeled: vasculature
150000
125000
100000
75000
50000
y
30000
60000
90000
x
Probability
1.00
0.75
0.50
0.25
0.00

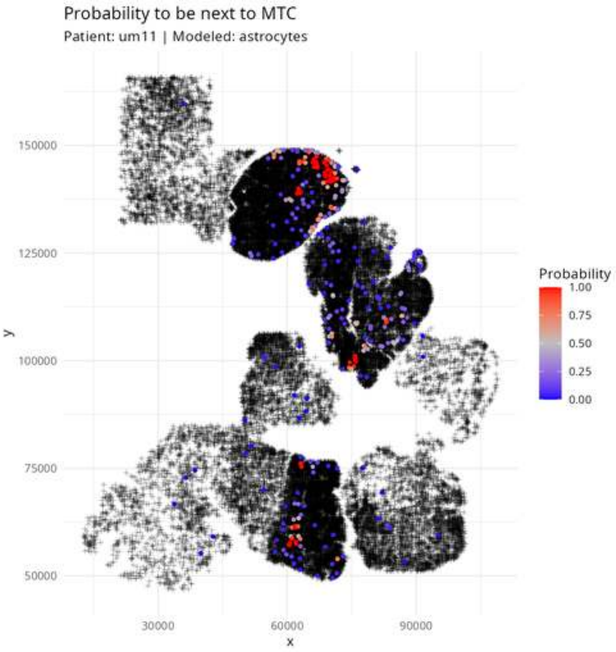

Probability to be next to MTC
Patient: um11 | Modeled: astrocytes
150000
125000
100000
75000
50000
y
30000
60000
90000
x
Probability
1.00
0.75
0.50
0.25
0.00

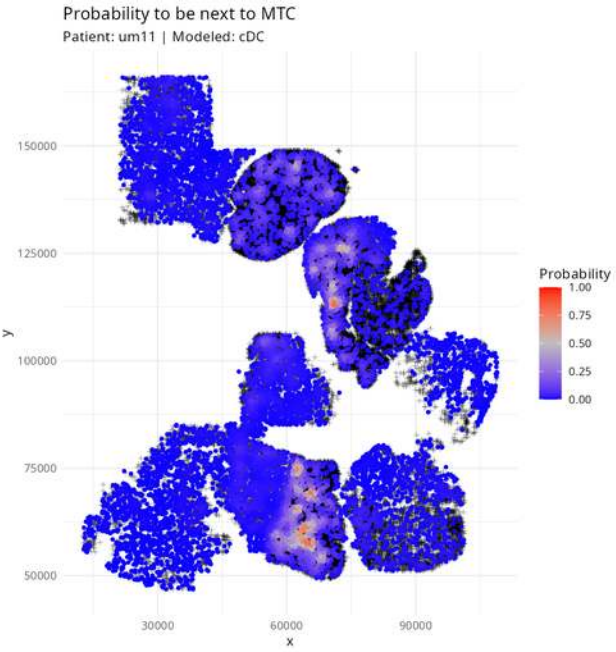

Probability to be next to MTC
Patient: um11 | Modeled: cDC
150000
125000
100000
75000
50000
y
30000
60000
90000
x
Probability
1.00
0.75
0.50
0.25
0.00

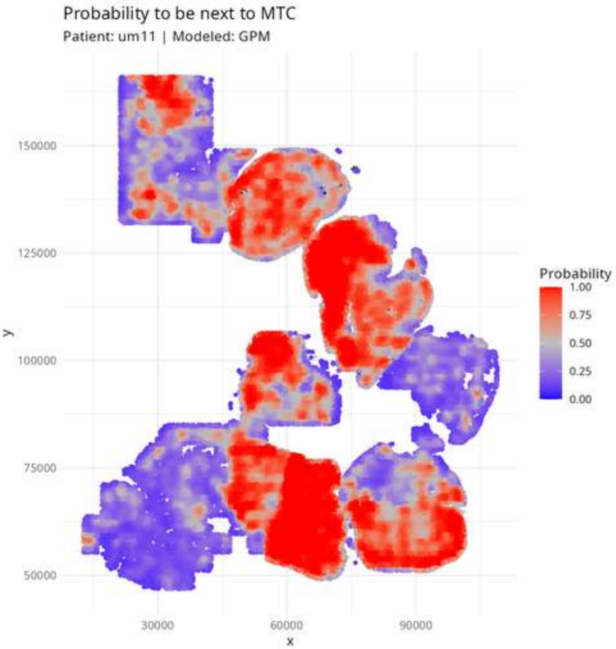

Probability to be next to MTC
Patient: um11 | Modeled: GPM
y
150000
125000
100000
75000
50000
x
30000
60000
90000
Probability
1.00
0.75
0.50
0.25
0.00

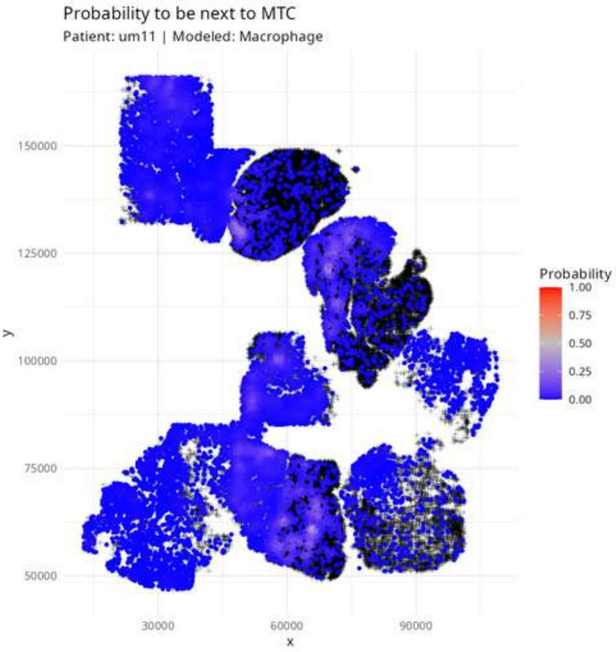

Probability to be next to MTC
Patient: um11 | Modeled: Macrophage
150000
125000
100000
75000
50000
y
30000
60000
90000
x
Probability
1.00
0.75
0.50
0.25
0.00

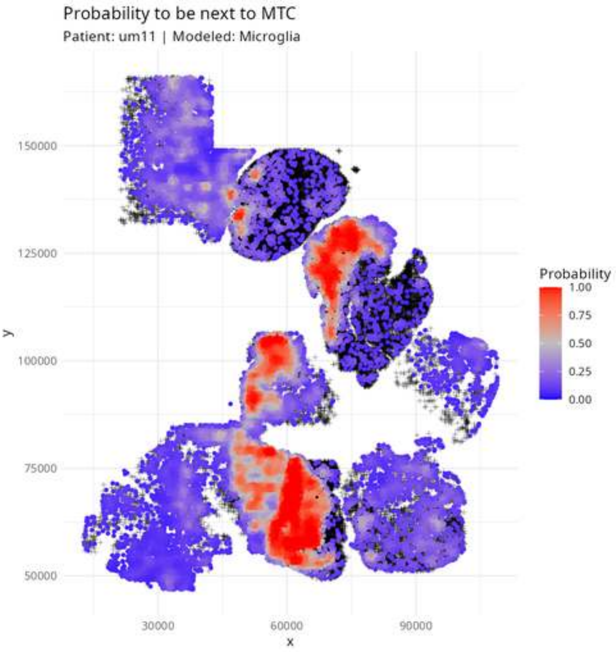
Probability to be next to MTC
Patient: um11 | Modeled: Microglia
150000
125000
100000
75000
50000
y
30000
60000
90000
x
Probability
1.00
0.75
0.50
0.25
0.00

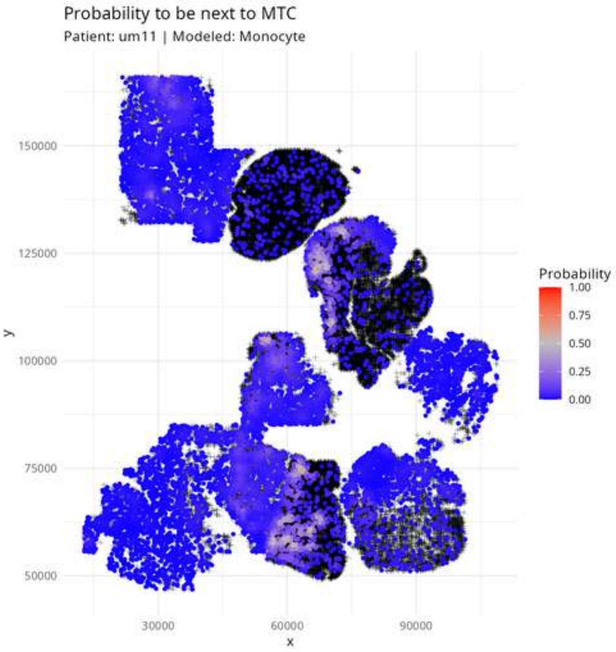

Probability to be next to MTC
Patient: um11 | Modeled: Monocyte
150000
125000
100000
75000
50000
y
30000
60000
90000
x
Probability
1.00
0.75
0.50
0.25
0.00

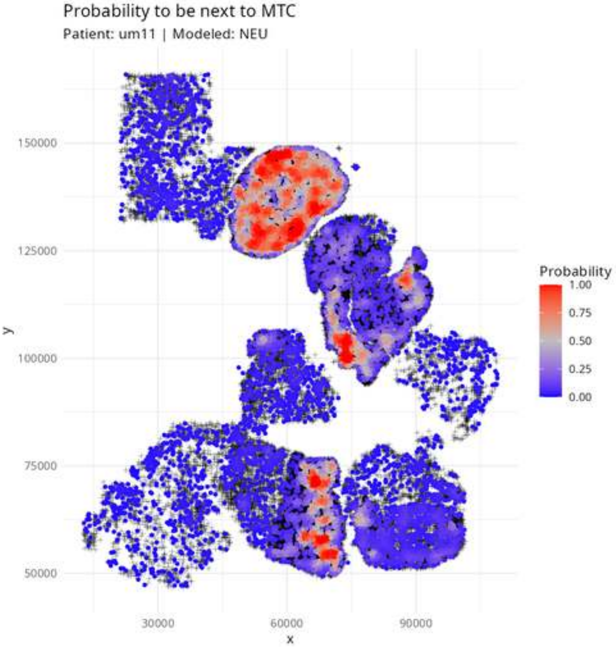

Probability to be next to MTC
Patient: um11 | Modeled: NEU
150000
125000
100000
75000
50000
y
30000
60000
90000
x
Probability
1.00
0.75
0.50
0.25
0.00

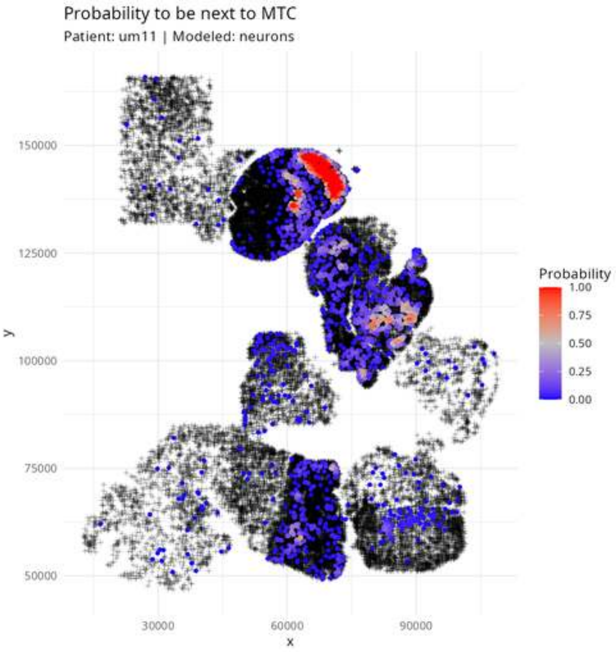

Probability to be next to MTC
Patient: um11 | Modeled: neurons
y
150000
125000
100000
75000
50000
x
30000
60000
90000
Probability
1.00
0.75
0.50
0.25
0.00

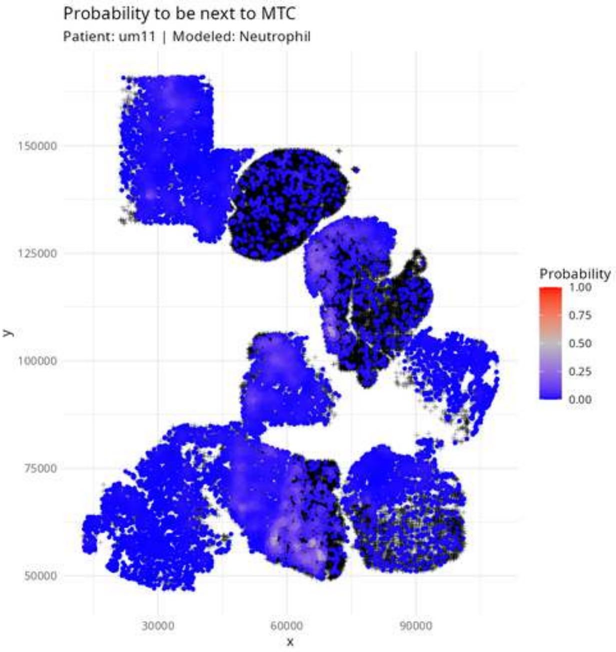

Probability to be next to MTC
Patient: um11 | Modeled: Neutrophil
150000
125000
100000
75000
50000
y
30000
60000
90000
x
Probability
1.00
0.75
0.50
0.25
0.00

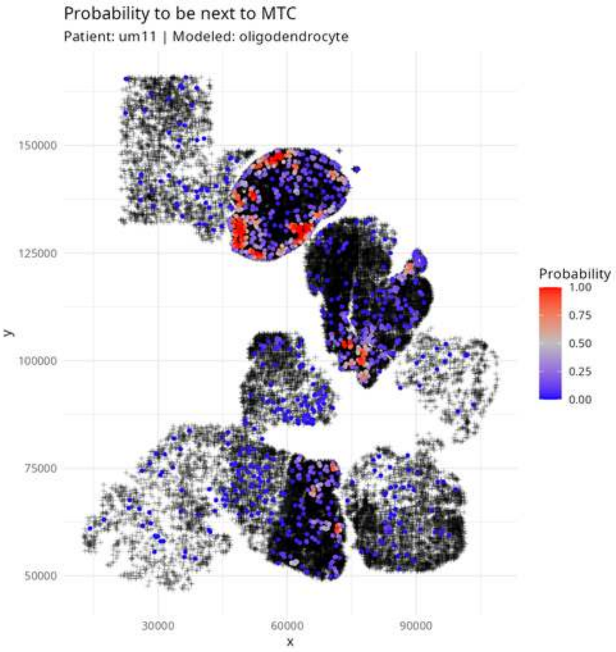

Probability to be next to MTC
Patient: um11 | Modeled: oligodendrocyte
150000
125000
100000
75000
50000
y
30000
60000
90000
x
Probability
1.00
0.75
0.50
0.25
0.00

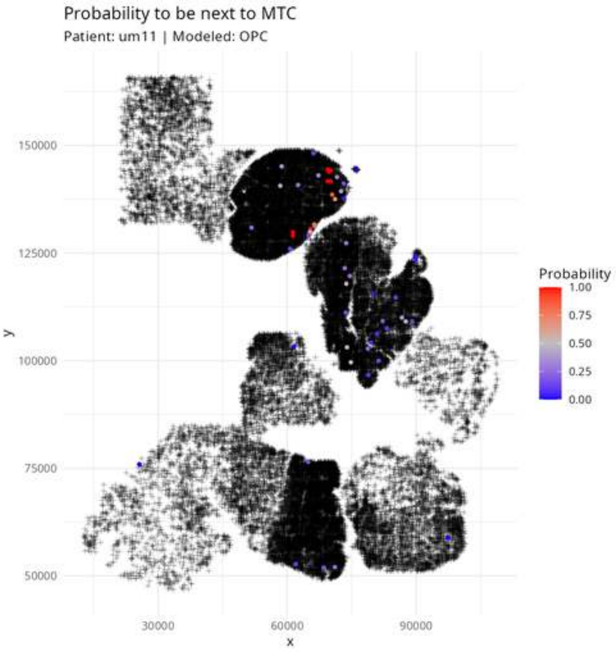
Probability to be next to MTC
Patient: um11 | Modeled: OPC
150000
125000
100000
75000
50000
y
30000
60000
90000
x
Probability
1.00
0.75
0.50
0.25
0.00

Probability to be next to MTC

Patient: um11 | Modeled: PPR

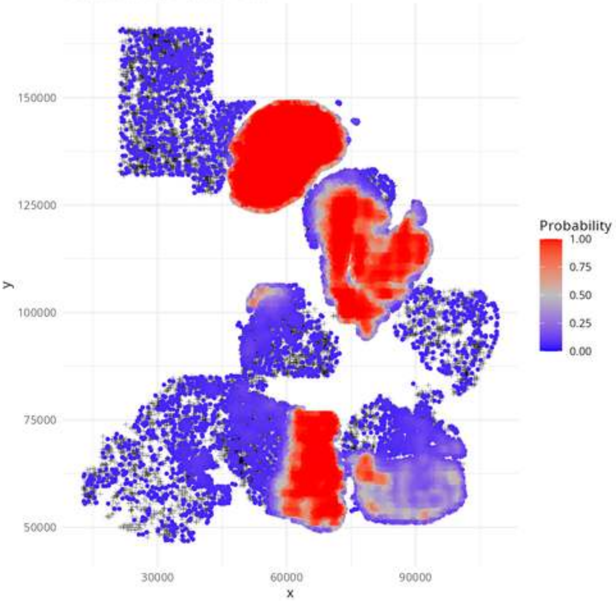

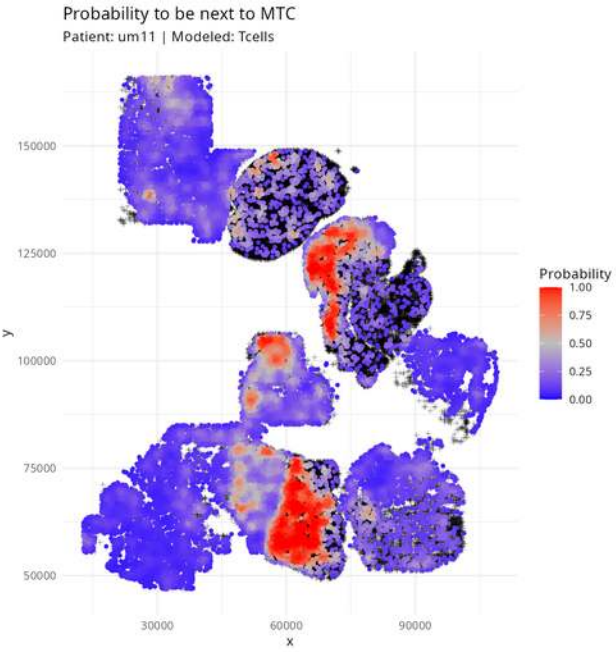
Probability to be next to MTC
Patient: um11 | Modeled: Tcells
150000
125000
100000
75000
50000
y
30000
60000
90000
x
Probability
1.00
0.75
0.50
0.25
0.00

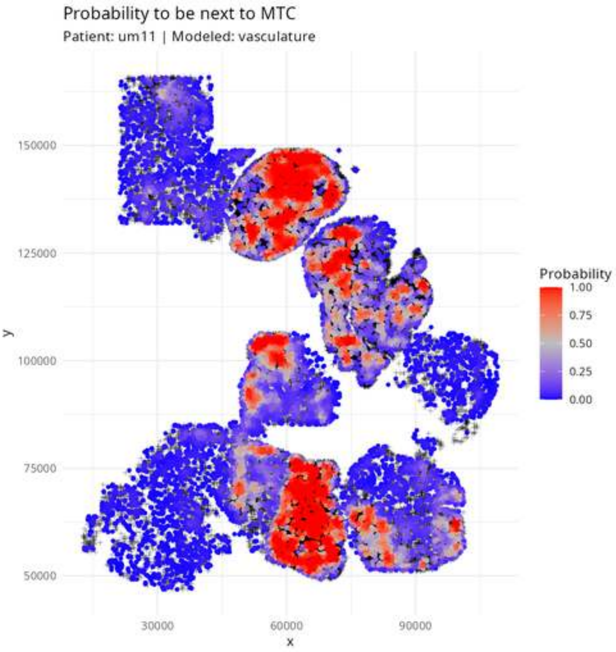
Probability to be next to MTC
Patient: um11 | Modeled: vasculature
150000
125000
100000
75000
50000
y
30000
60000
90000
x
Probability
1.00
0.75
0.50
0.25
0.00

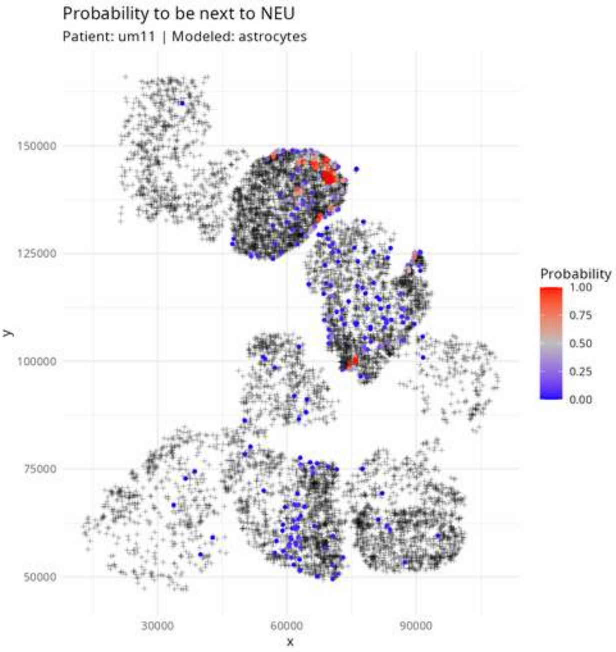

Probability to be next to NEU
Patient: um11 | Modeled: astrocytes
150000
125000
100000
75000
50000
y
30000
60000
90000
x
Probability
1.00
0.75
0.50
0.25
0.00

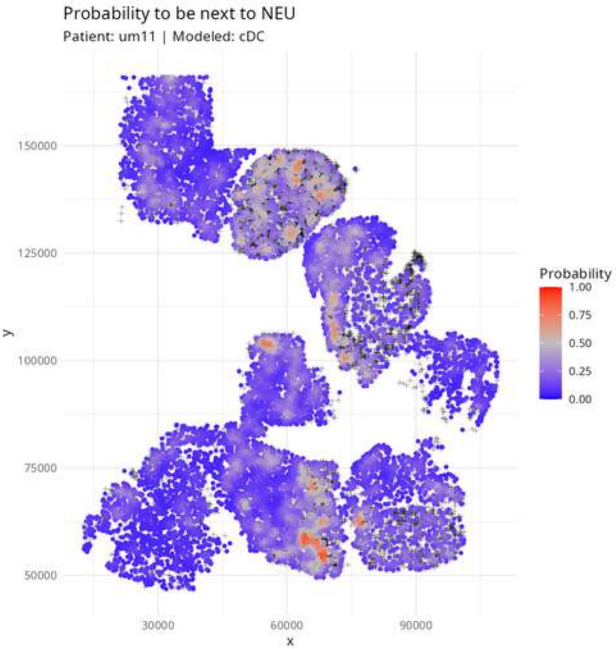

Probability to be next to NEU
Patient: um11 | Modeled: cDC
150000
125000
100000
75000
50000
y
30000
60000
90000
x
Probability
1.00
0.75
0.50
0.25
0.00

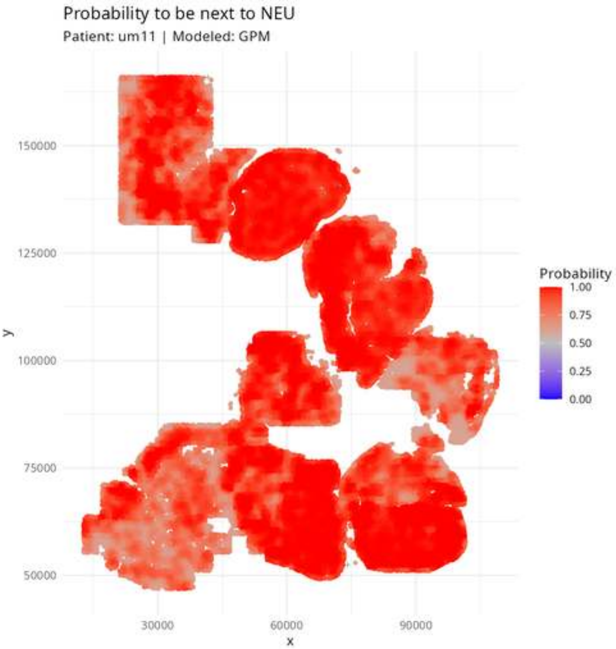

Probability to be next to NEU
Patient: um11 | Modeled: GPM
150000
125000
100000
75000
50000
y
30000
60000
90000
x
Probability
1.00
0.75
0.50
0.25
0.00

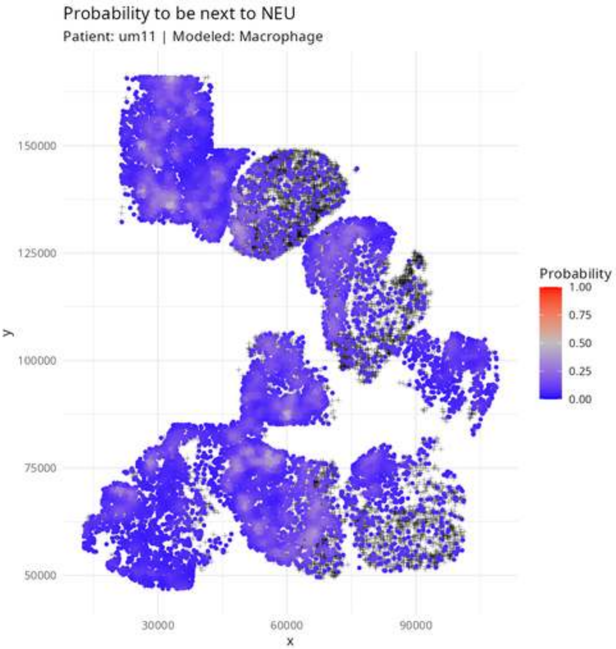

Probability to be next to NEU
Patient: um11 | Modeled: Macrophage
150000
125000
100000
75000
50000
y
30000
60000
90000
x
Probability
1.00
0.75
0.50
0.25
0.00

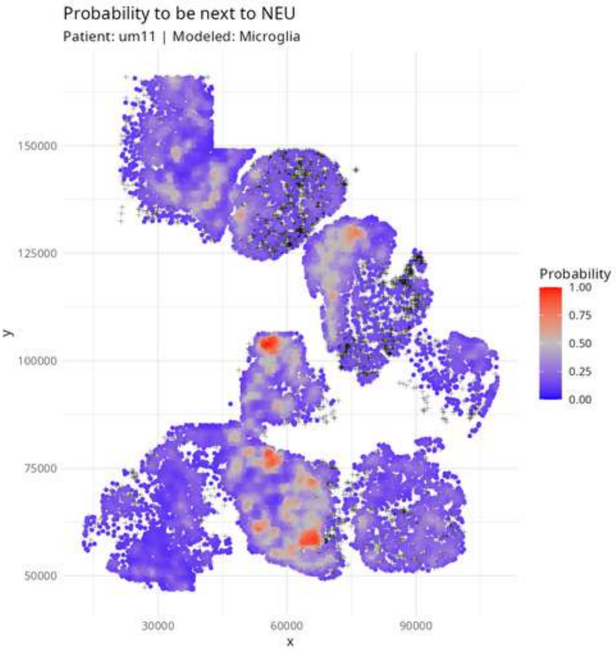

Probability to be next to NEU
Patient: um11 | Modeled: Microglia
Probability
1.00
0.75
0.50
0.25
0.00
y
150000
125000
100000
75000
50000
x
30000
60000
90000

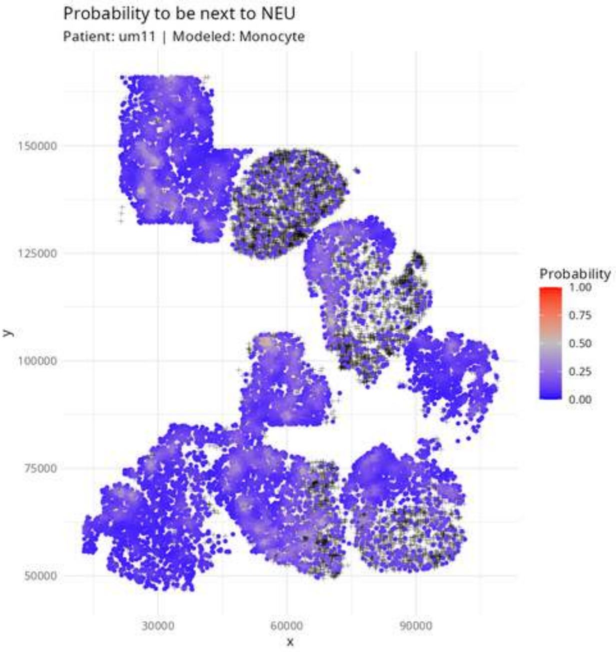

Probability to be next to NEU
Patient: um11 | Modeled: Monocyte
150000
125000
100000
75000
50000
y
30000
60000
90000
x
Probability
1.00
0.75
0.50
0.25
0.00

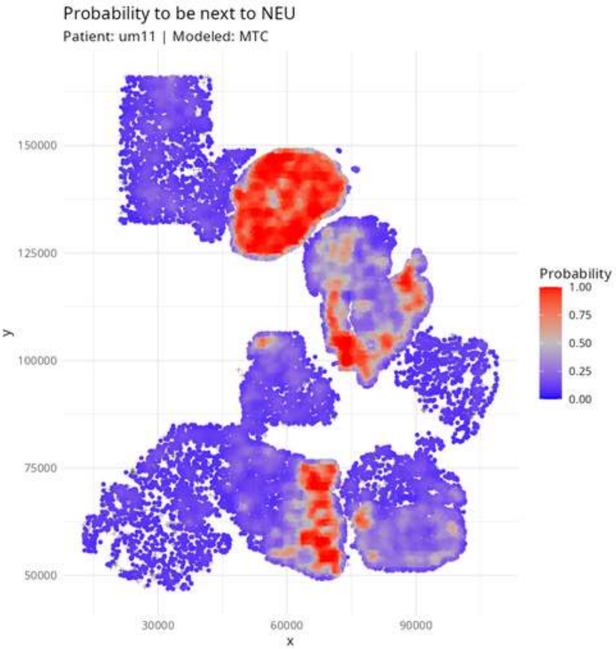
Probability to be next to NEU
Patient: um11 | Modeled: MTC
150000
125000
100000
75000
50000
y
30000
60000
90000
x
Probability
1.00
0.75
0.50
0.25
0.00

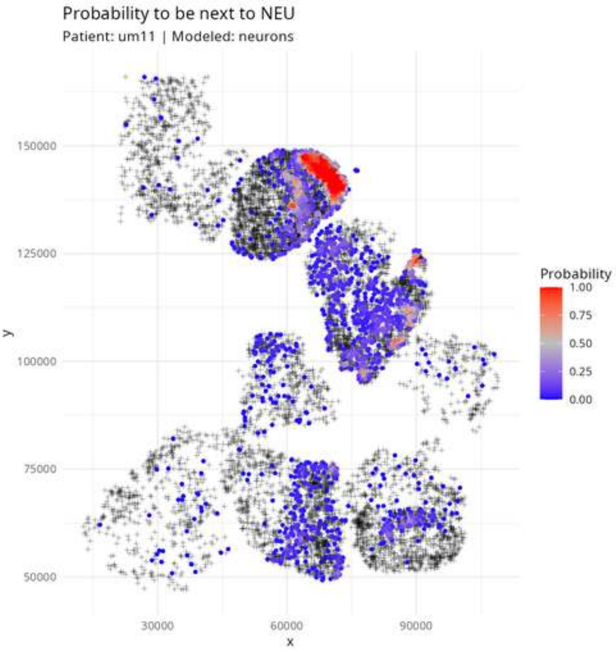

Probability to be next to NEU
Patient: um11 | Modeled: neurons
150000
125000
100000
75000
50000
y
30000
60000
90000
x
Probability
1.00
0.75
0.50
0.25
0.00

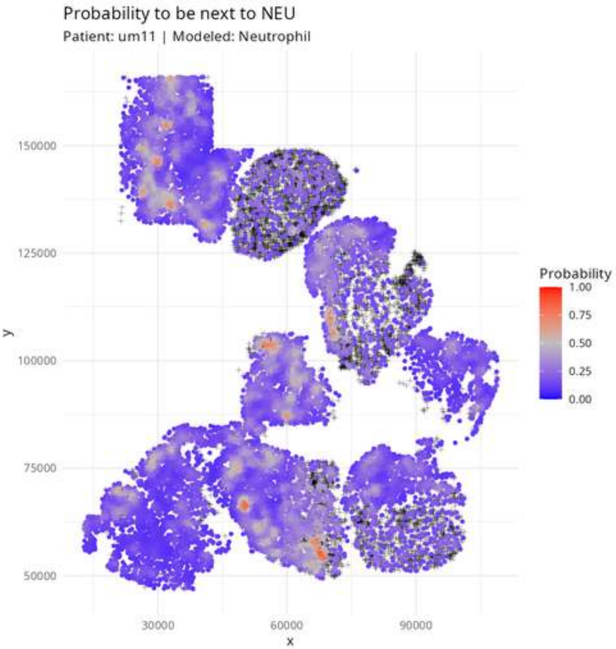
Probability to be next to NEU
Patient: um11 | Modeled: Neutrophil
150000
125000
100000
75000
50000
y
30000
60000
90000
x
Probability
1.00
0.75
0.50
0.25
0.00

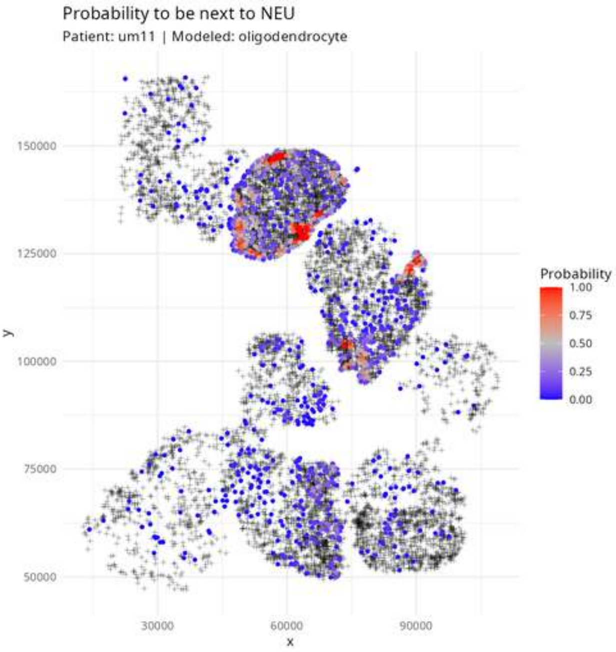

Probability to be next to NEU
Patient: um11 | Modeled: oligodendrocyte
y
150000
125000
100000
75000
50000
x
30000
60000
90000
Probability
1.00
0.75
0.50
0.25
0.00

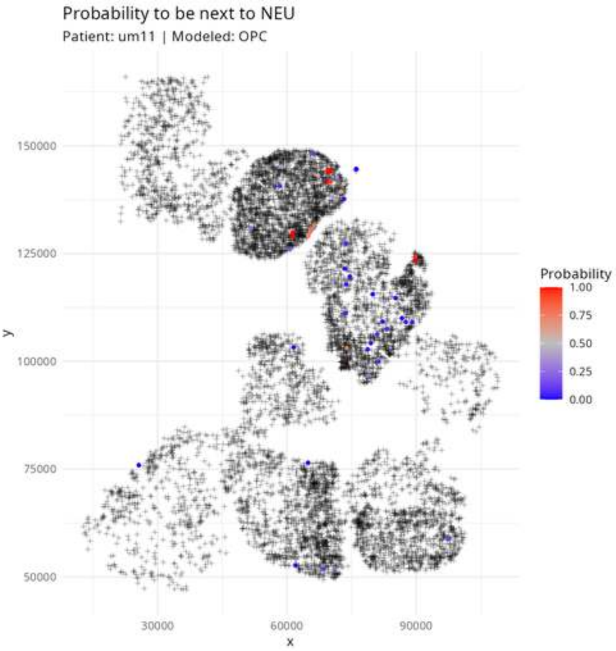
Probability to be next to NEU
Patient: um11 | Modeled: OPC
150000
125000
100000
75000
50000
y
30000
60000
90000
x
Probability
1.00
0.75
0.50
0.25
0.00

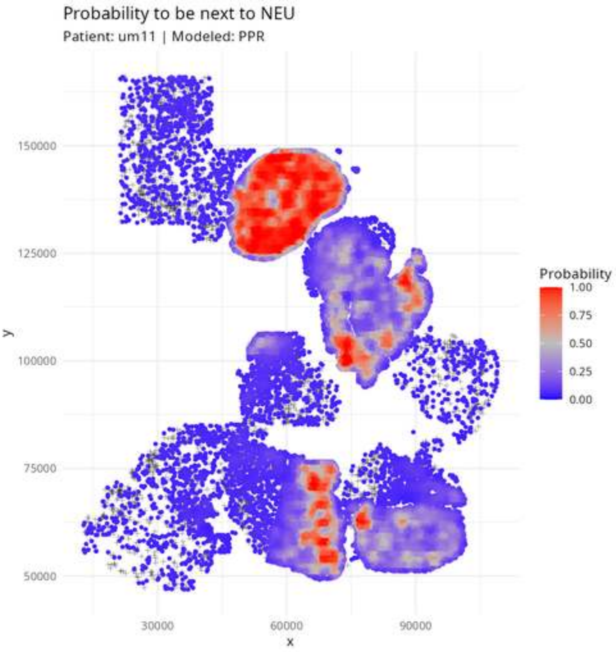

Probability to be next to NEU
Patient: um11 | Modeled: PPR
150000
125000
100000
75000
50000
y
30000
60000
90000
x
Probability
1.00
0.75
0.50
0.25
0.00

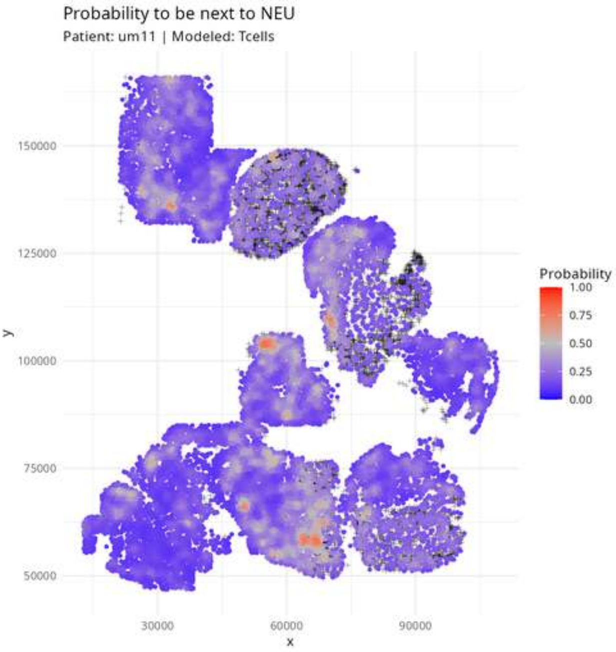

Probability to be next to NEU
Patient: um11 | Modeled: Tcells
y
150000
125000
100000
75000
50000
x
30000
60000
90000
Probability
1.00
0.75
0.50
0.25
0.00

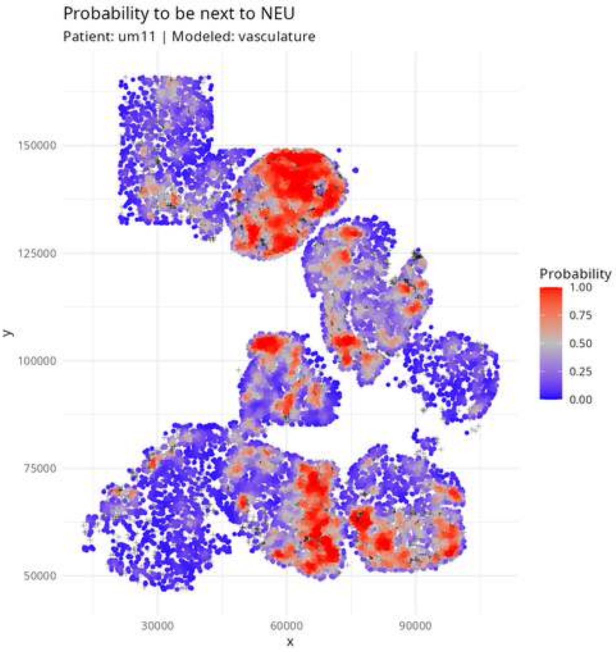
Probability to be next to NEU
Patient: um11 | Modeled: vasculature
150000
125000
100000
75000
50000
y
30000
60000
90000
x
Probability
1.00
0.75
0.50
0.25
0.00

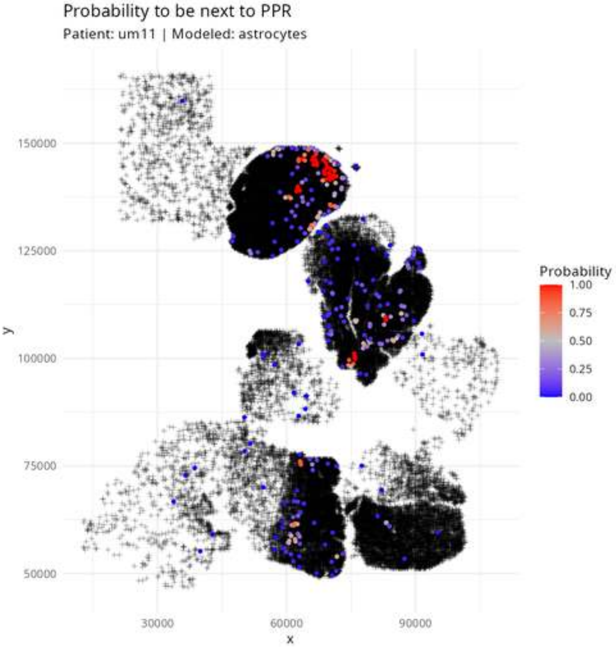
Probability to be next to PPR
Patient: um11 | Modeled: astrocytes
Probability
1.00
0.75
0.50
0.25
0.00
150000
125000
100000
75000
50000
y
30000
60000
90000
x

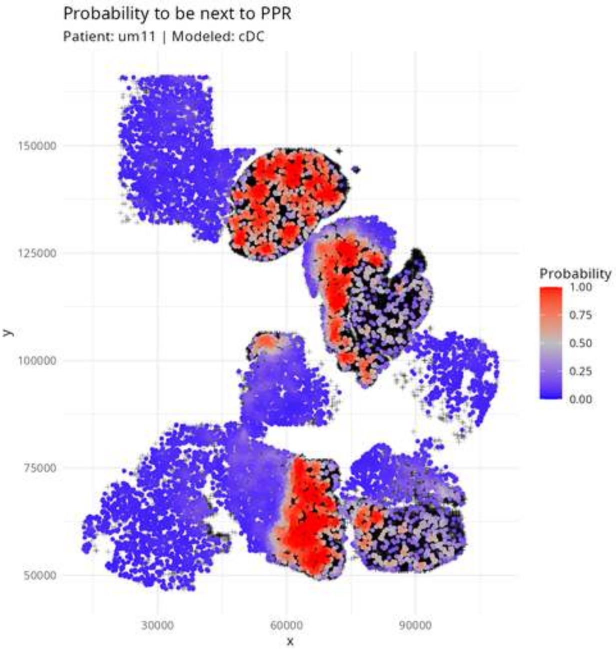
Probability to be next to PPR
Patient: um11 | Modeled: cDC
150000
125000
100000
75000
50000
y
30000
60000
90000
x
Probability
1.00
0.75
0.50
0.25
0.00

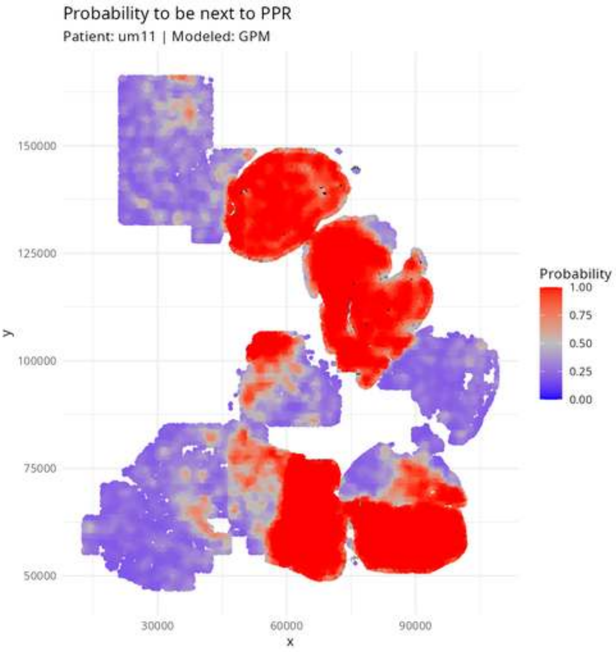
Probability to be next to PPR
Patient: um11 | Modeled: GPM
150000
125000
100000
75000
50000
y
30000
60000
90000
x
Probability
1.00
0.75
0.50
0.25
0.00

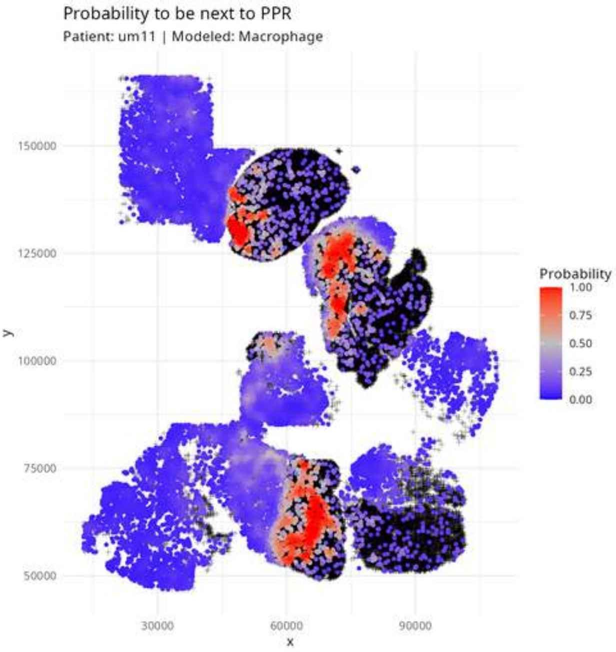
Probability to be next to PPR
Patient: um11 | Modeled: Macrophage
150000
125000
100000
75000
50000
y
30000
60000
90000
x
Probability
1.00
0.75
0.50
0.25
0.00

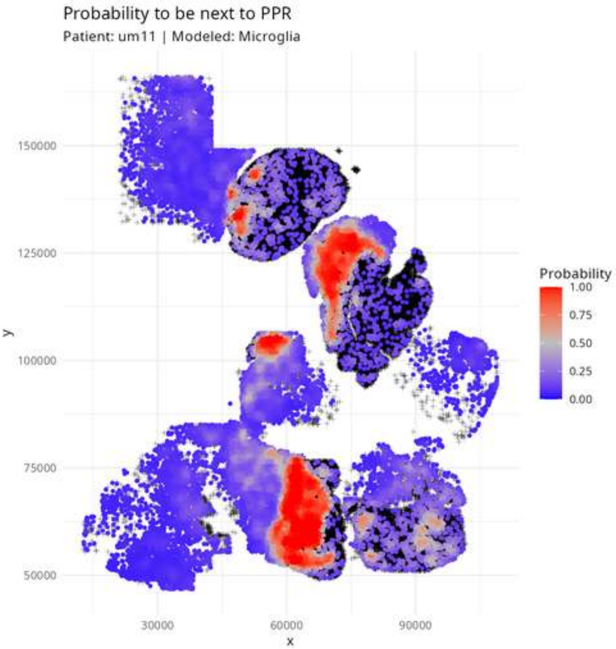
Probability to be next to PPR
Patient: um11 | Modeled: Microglia
150000
125000
100000
75000
50000
y
30000
60000
90000
x
Probability
1.00
0.75
0.50
0.25
0.00

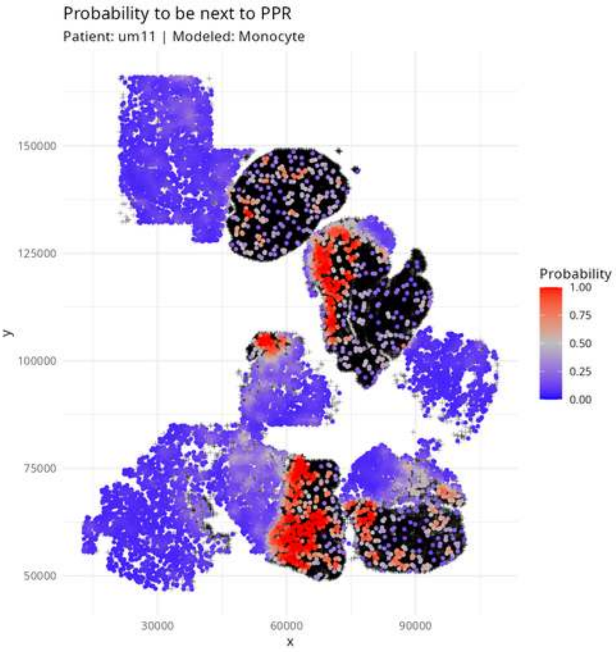
Probability to be next to PPR
Patient: um11 | Modeled: Monocyte
150000
125000
100000
75000
50000
y
30000
60000
90000
x
Probability
1.00
0.75
0.50
0.25
0.00

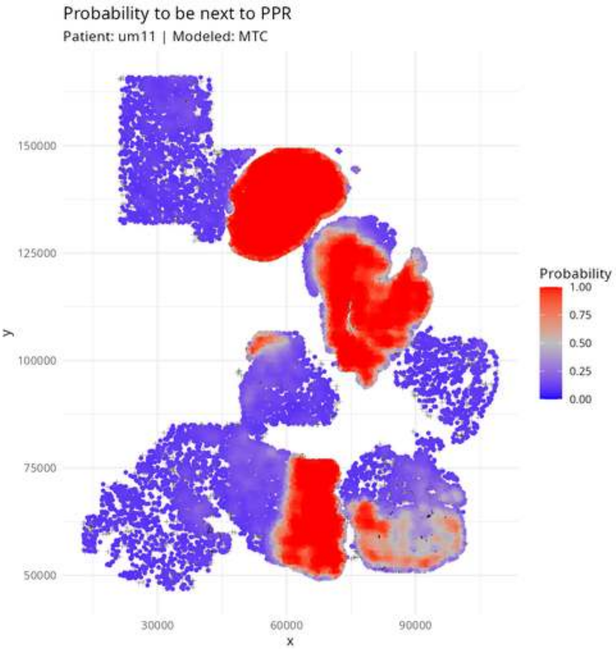

Probability to be next to PPR
Patient: um11 | Modeled: MTC
y
150000
125000
100000
75000
50000
30000
60000
90000
x
Probability
1.00
0.75
0.50
0.25
0.00

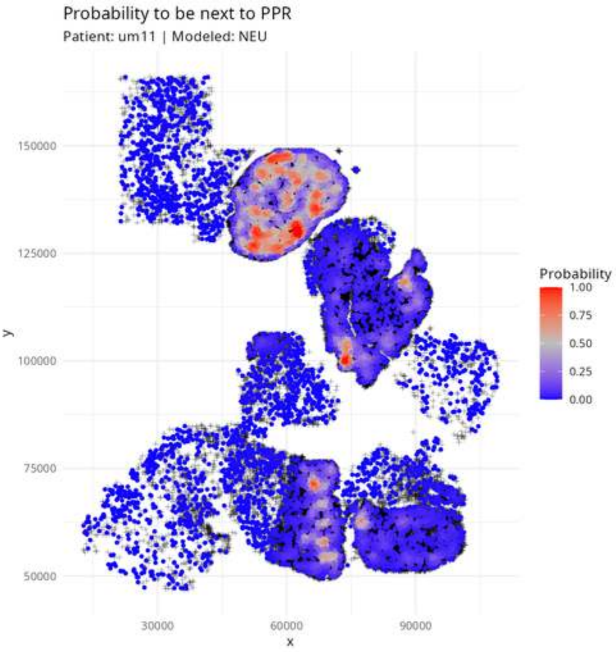
Probability to be next to PPR
Patient: um11 | Modeled: NEU
150000
125000
100000
75000
50000
y
30000
60000
90000
x
Probability
1.00
0.75
0.50
0.25
0.00

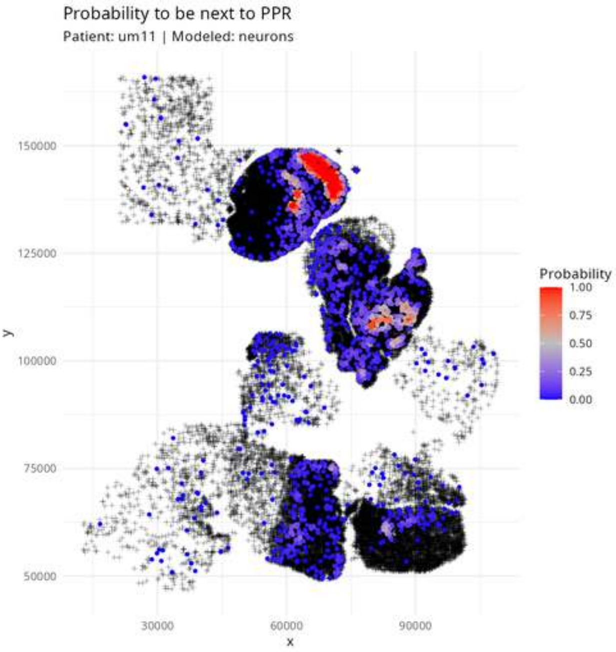

Probability to be next to PPR
Patient: um11 | Modeled: neurons
y
150000
125000
100000
75000
50000
x
30000
60000
90000
Probability
1.00
0.75
0.50
0.25
0.00

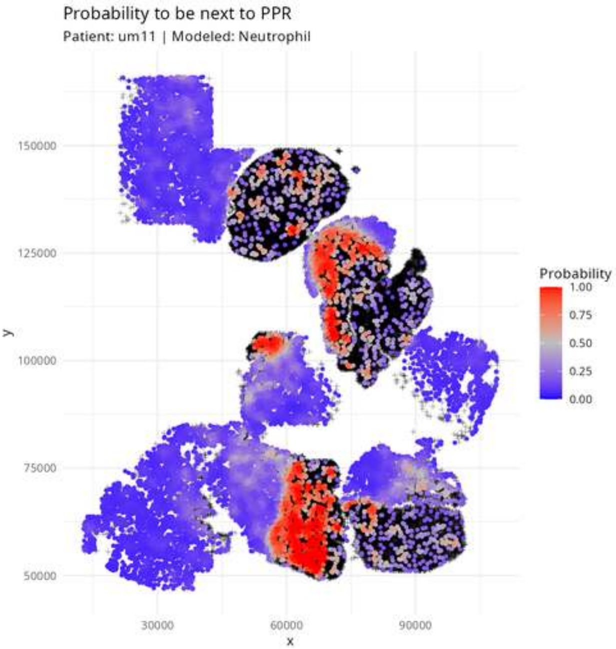
Probability to be next to PPR
Patient: um11 | Modeled: Neutrophil
150000
125000
100000
75000
50000
y
30000
60000
90000
x
Probability
1.00
0.75
0.50
0.25
0.00

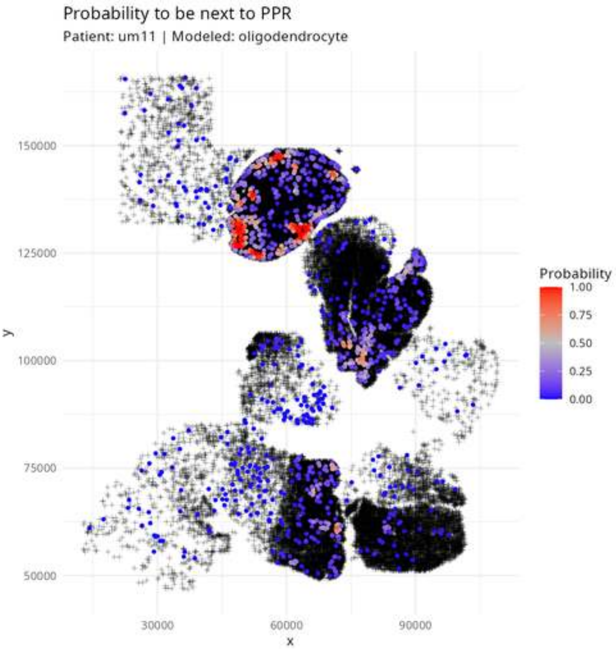

Probability to be next to PPR
Patient: um11 | Modeled: oligodendrocyte
150000
125000
100000
75000
50000
y
30000
60000
90000
x
Probability
1.00
0.75
0.50
0.25
0.00

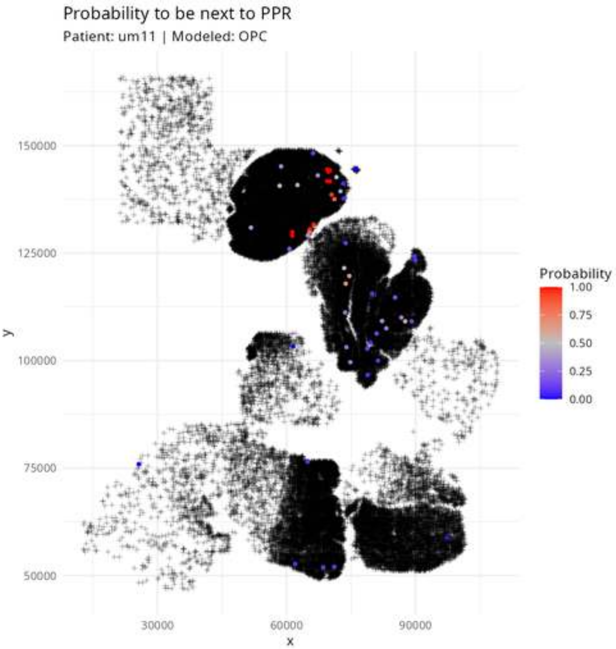
Probability to be next to PPR
Patient: um11 | Modeled: OPC
150000
125000
100000
75000
50000
y
30000
60000
90000
x
Probability
1.00
0.75
0.50
0.25
0.00

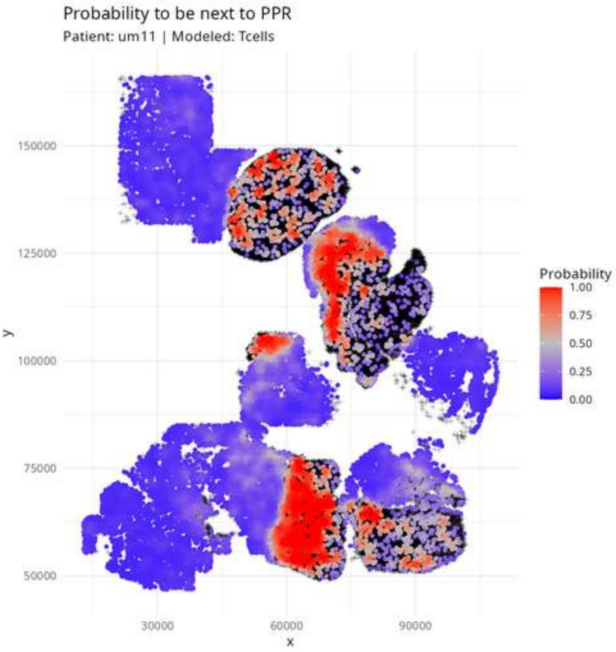

Probability to be next to PPR
Patient: um11 | Modeled: Tcells
y
150000
125000
100000
75000
50000
30000
60000
90000
x
Probability
1.00
0.75
0.50
0.25
0.00

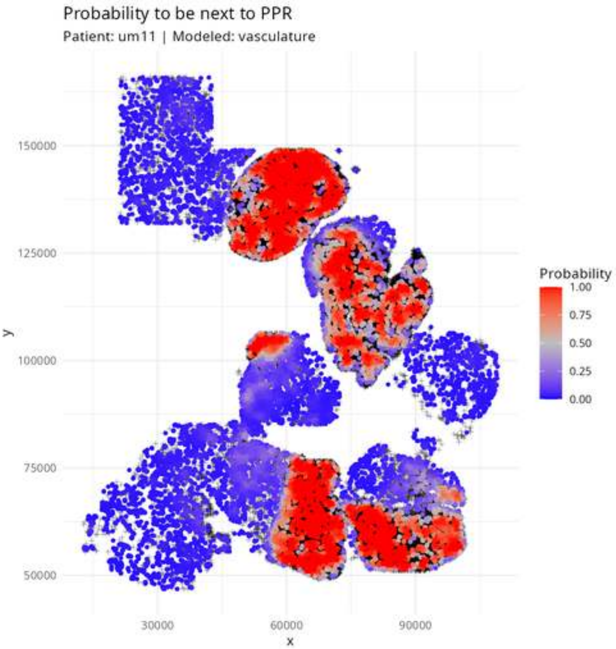
Probability to be next to PPR
Patient: um11 | Modeled: vasculature
150000
125000
100000
75000
50000
y
30000
60000
90000
x
Probability
1.00
0.75
0.50
0.25
0.00

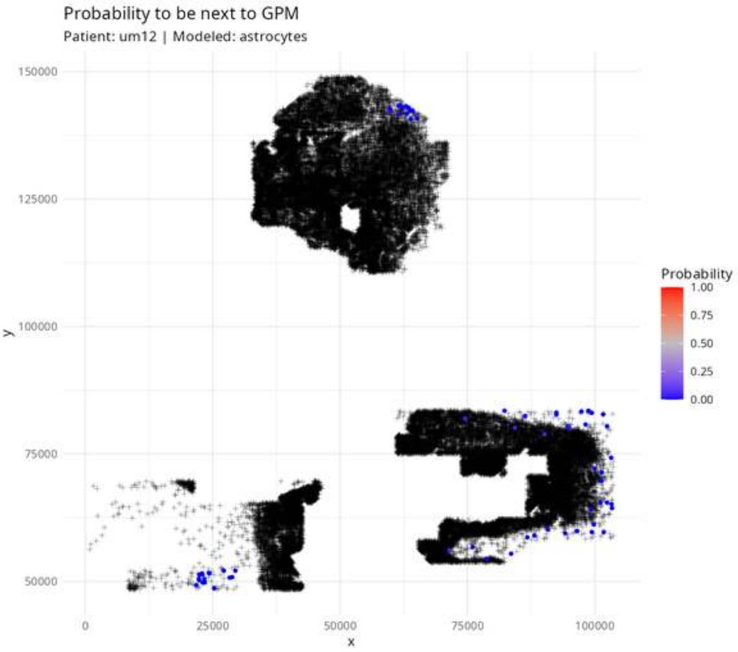

Probability to be next to GPM
Patient: um12 | Modeled: astrocytes
150000
125000
100000
75000
50000
y
0
25000
50000
75000
100000
x
Probability
1.00
0.75
0.50
0.25
0.00

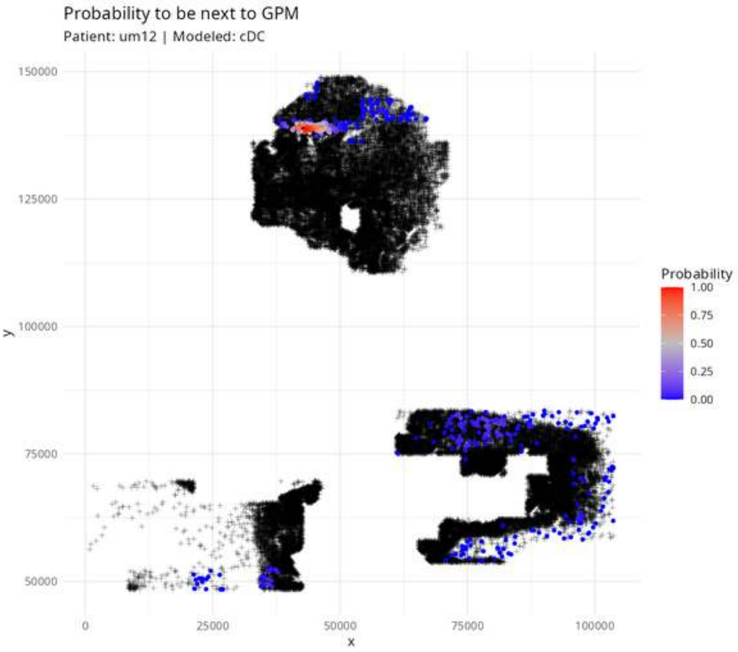
Probability to be next to GPM
Patient: um12 | Modeled: cDC
150000
125000
100000
75000
50000
y
0
25000
50000
75000
100000
x
Probability
1.00
0.75
0.50
0.25
0.00

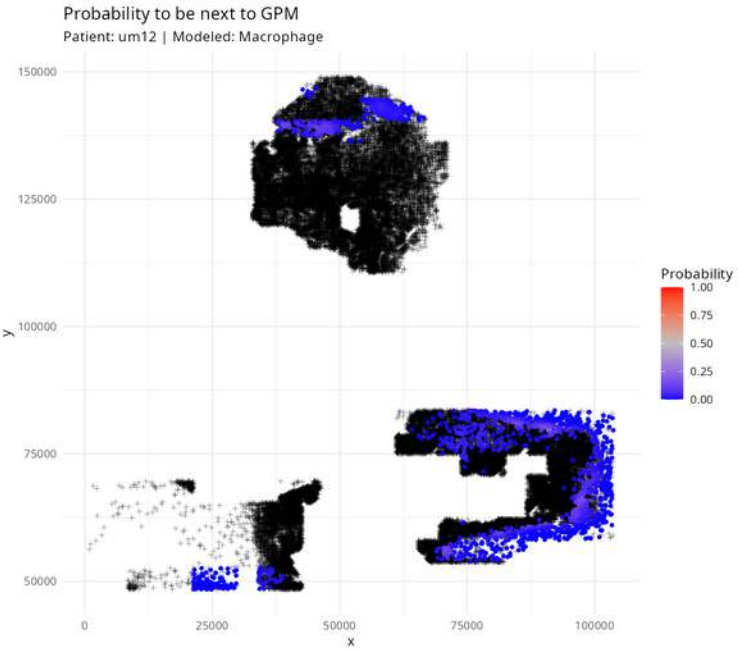

Probability to be next to GPM
Patient: um12 | Modeled: Macrophage
150000
125000
100000
75000
50000
y
0
25000
50000
75000
100000
x
Probability
1.00
0.75
0.50
0.25
0.00

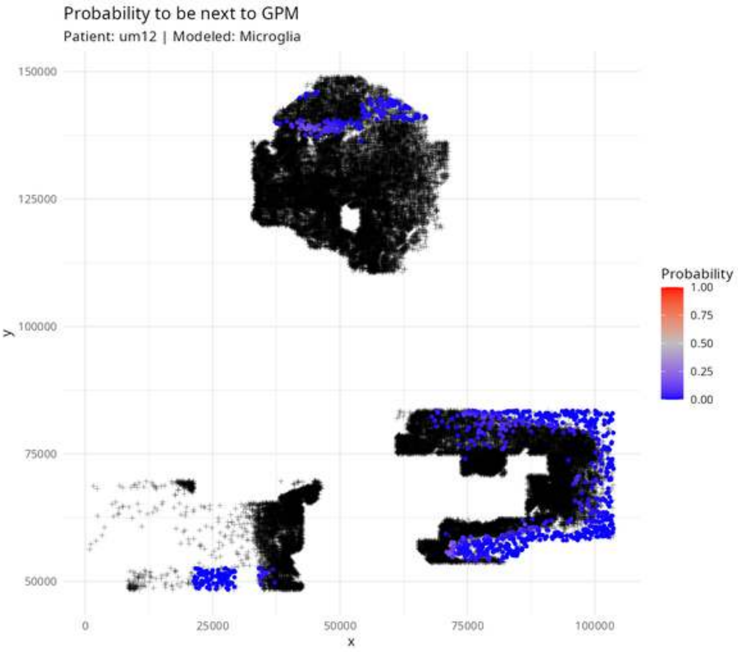

Probability to be next to GPM
Patient: um12 | Modeled: Microglia
150000
125000
100000
75000
50000
y
0
25000
50000
75000
100000
x
Probability
1.00
0.75
0.50
0.25
0.00

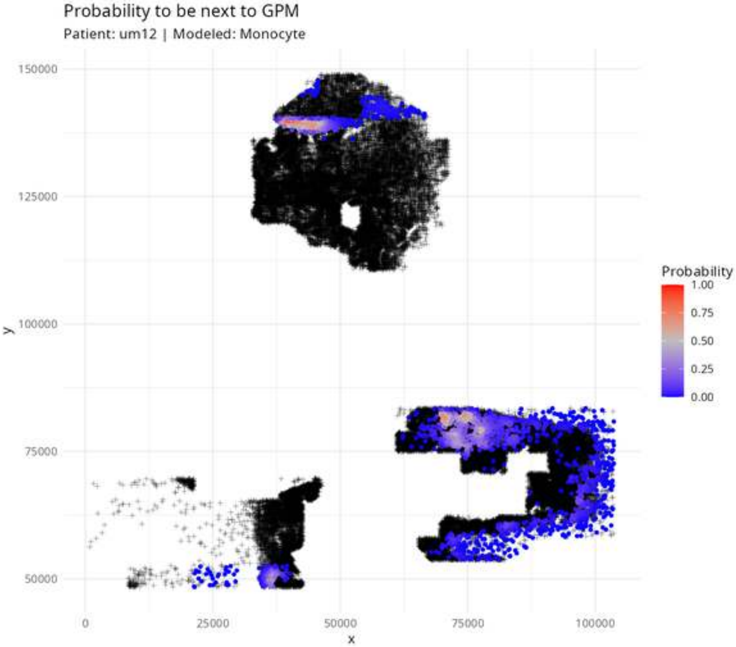
Probability to be next to GPM
Patient: um12 | Modeled: Monocyte
150000
125000
100000
75000
50000
y
0
25000
50000
75000
100000
x
Probability
1.00
0.75
0.50
0.25
0.00

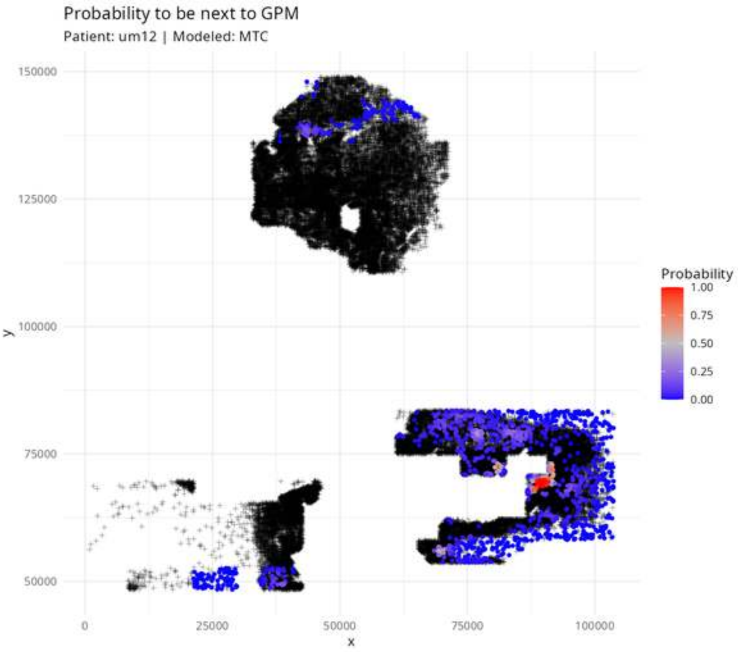
Probability to be next to GPM
Patient: um12 | Modeled: MTC
150000
125000
100000
75000
50000
y
0
25000
50000
75000
100000
x
Probability
1.00
0.75
0.50
0.25
0.00

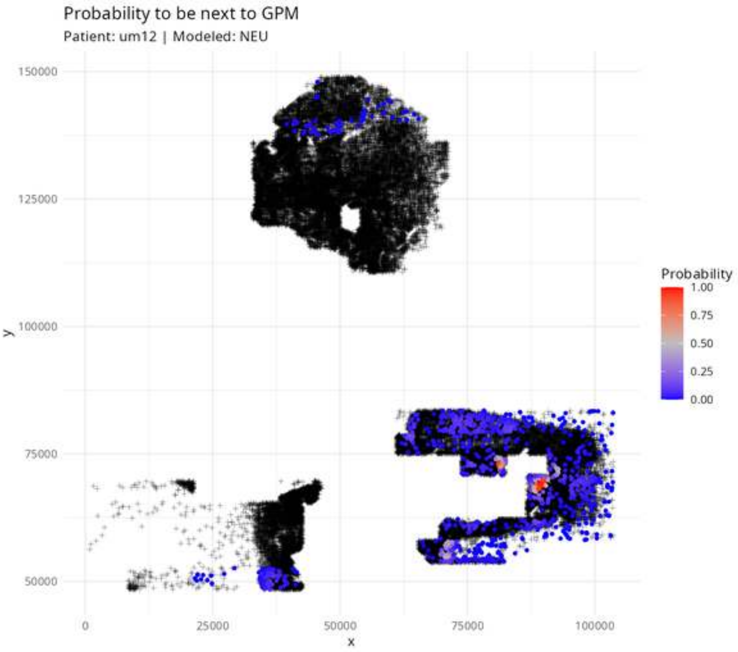
Probability to be next to GPM
Patient: um12 | Modeled: NEU
150000
125000
100000
75000
50000
y
0
25000
50000
75000
100000
x
Probability
1.00
0.75
0.50
0.25
0.00

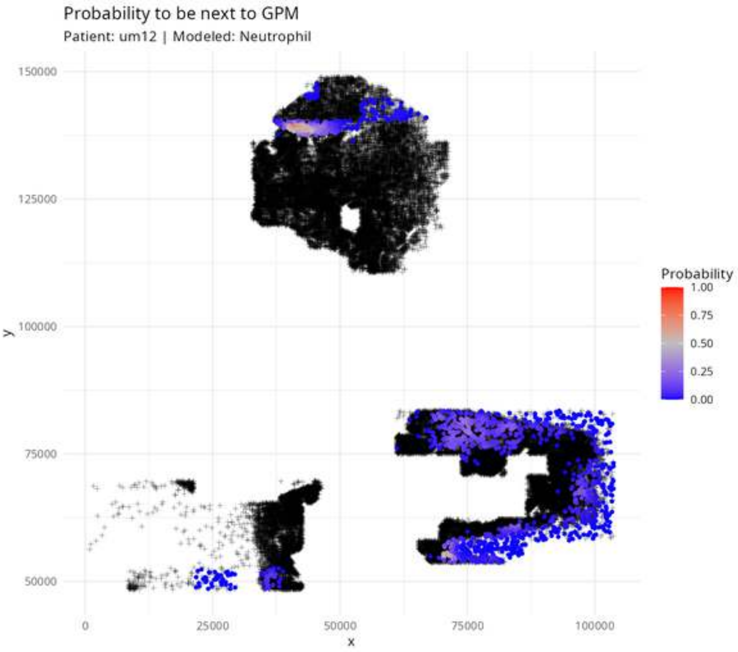

Probability to be next to GPM
Patient: um12 | Modeled: Neutrophil
150000
125000
100000
75000
50000
y
0
25000
50000
75000
100000
x
Probability
1.00
0.75
0.50
0.25
0.00

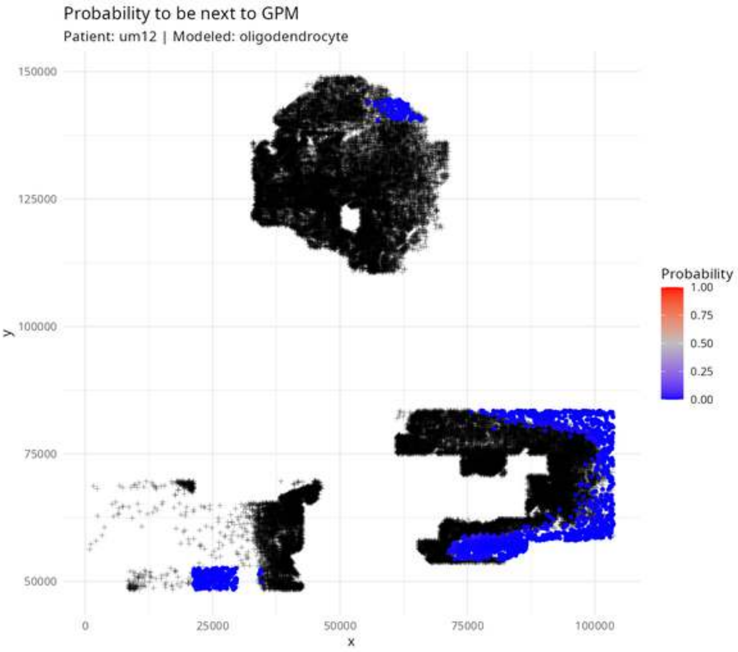

Probability to be next to GPM
Patient: um12 | Modeled: oligodendrocyte
150000
125000
100000
75000
50000
y
0
25000
50000
75000
100000
x
Probability
1.00
0.75
0.50
0.25
0.00

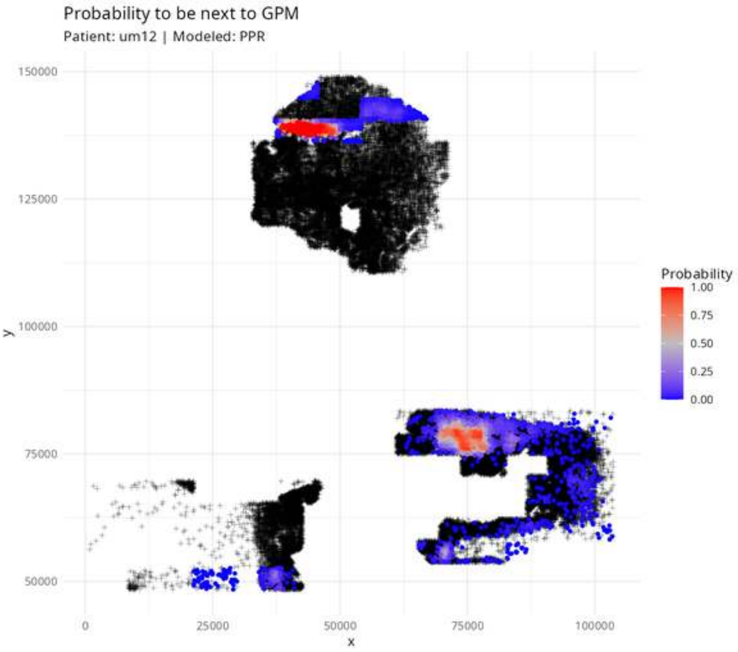

Probability to be next to GPM
Patient: um12 | Modeled: PPR
150000
125000
100000
75000
50000
y
0
25000
50000
75000
100000
x
Probability
1.00
0.75
0.50
0.25
0.00

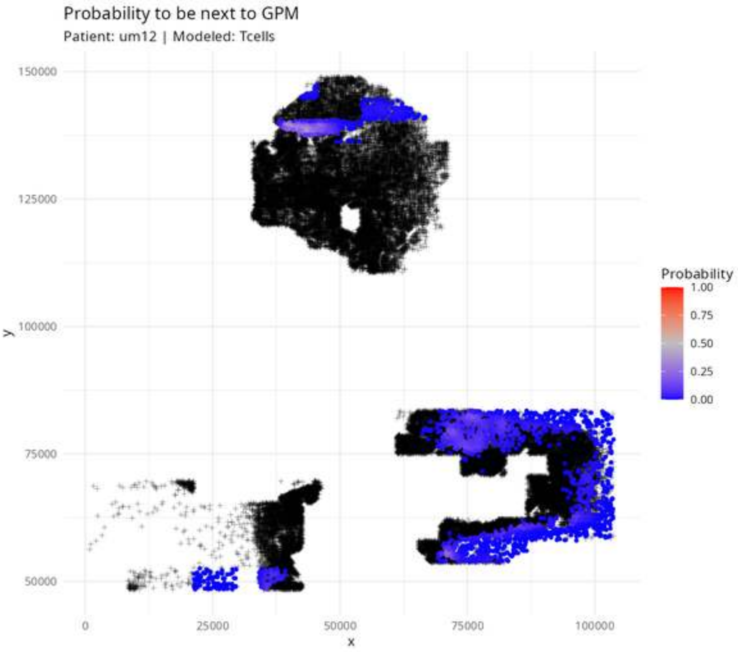
Probability to be next to GPM
Patient: um12 | Modeled: Tcells
150000
125000
100000
75000
50000
y
0
25000
50000
75000
100000
x
Probability
1.00
0.75
0.50
0.25
0.00

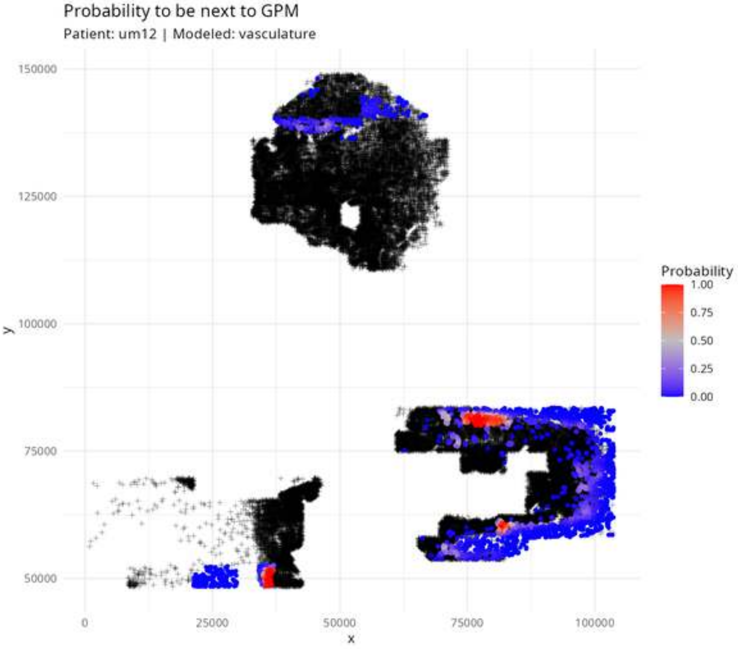

Probability to be next to GPM
Patient: um12 | Modeled: vasculature
150000
125000
100000
75000
50000
y
0
25000
50000
75000
100000
x
Probability
1.00
0.75
0.50
0.25
0.00

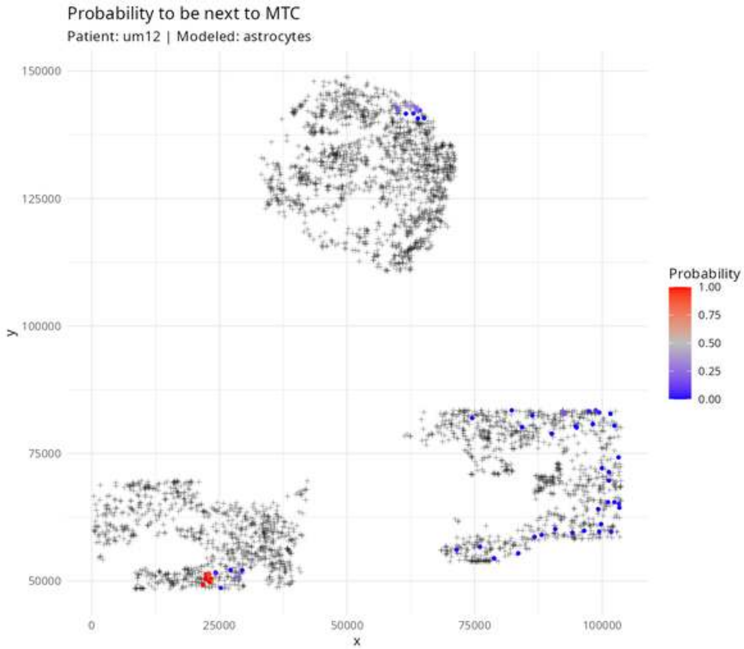

Probability to be next to MTC
Patient: um12 | Modeled: astrocytes
150000
125000
100000
75000
50000
y
0
25000
50000
75000
100000
x
Probability
1.00
0.75
0.50
0.25
0.00

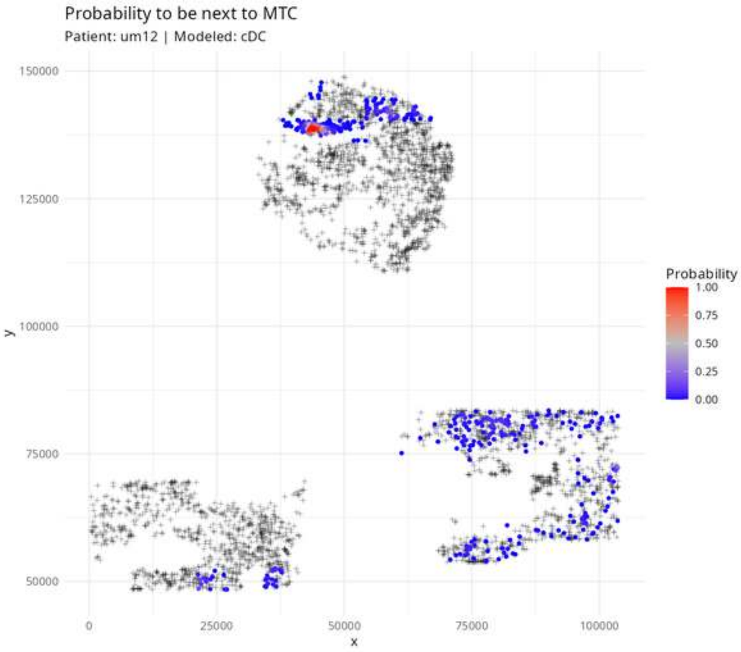
Probability to be next to MTC
Patient: um12 | Modeled: cDC
150000
125000
100000
75000
50000
y
0
25000
50000
75000
100000
x
Probability
1.00
0.75
0.50
0.25
0.00

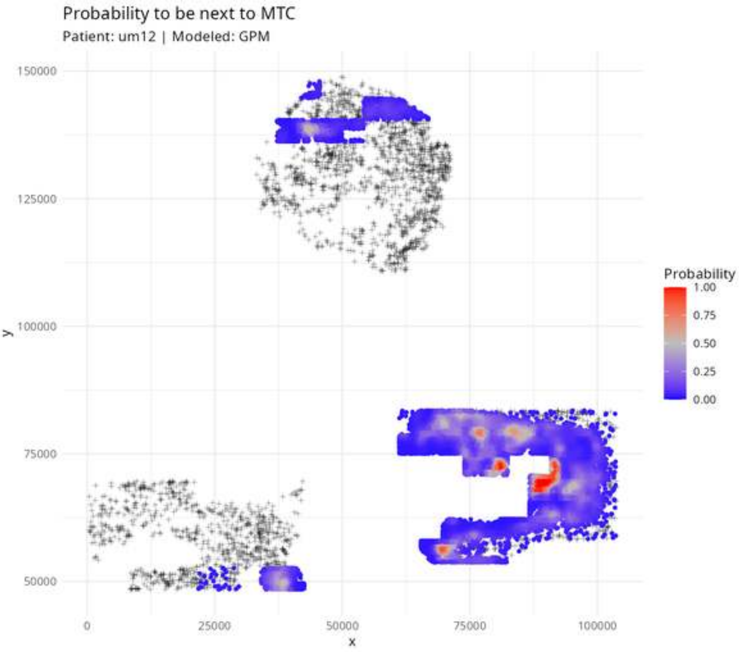

Probability to be next to MTC
Patient: um12 | Modeled: GPM
150000
125000
100000
75000
50000
y
0
25000
50000
75000
100000
x
Probability
1.00
0.75
0.50
0.25
0.00

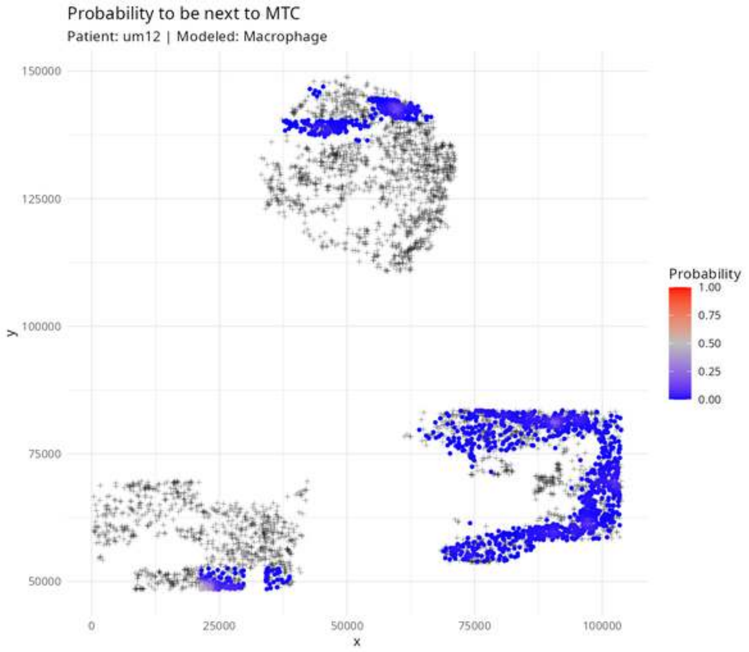
Probability to be next to MTC
Patient: um12 | Modeled: Macrophage
150000
125000
100000
75000
50000
y
0
25000
50000
75000
100000
x
Probability
1.00
0.75
0.50
0.25
0.00

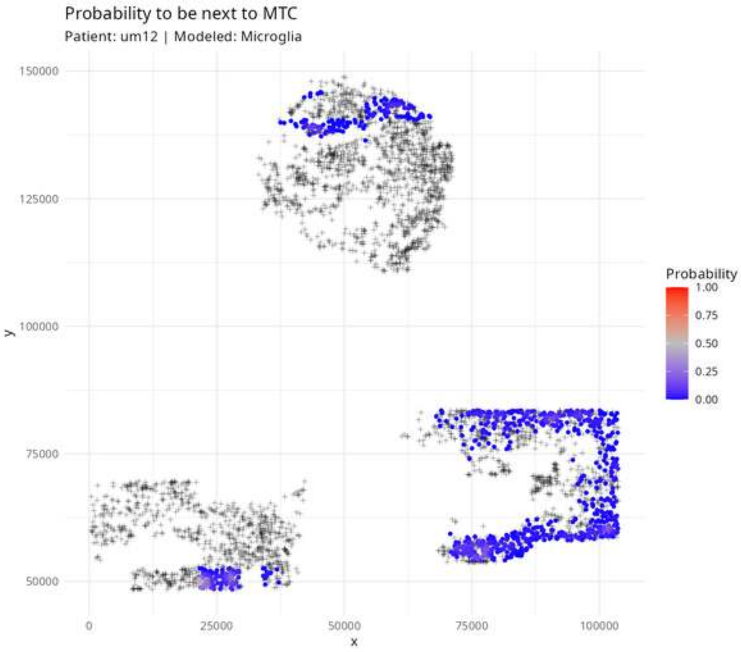
Probability to be next to MTC
Patient: um12 | Modeled: Microglia
150000
125000
100000
75000
50000
y
0
25000
50000
75000
100000
x
Probability
1.00
0.75
0.50
0.25
0.00

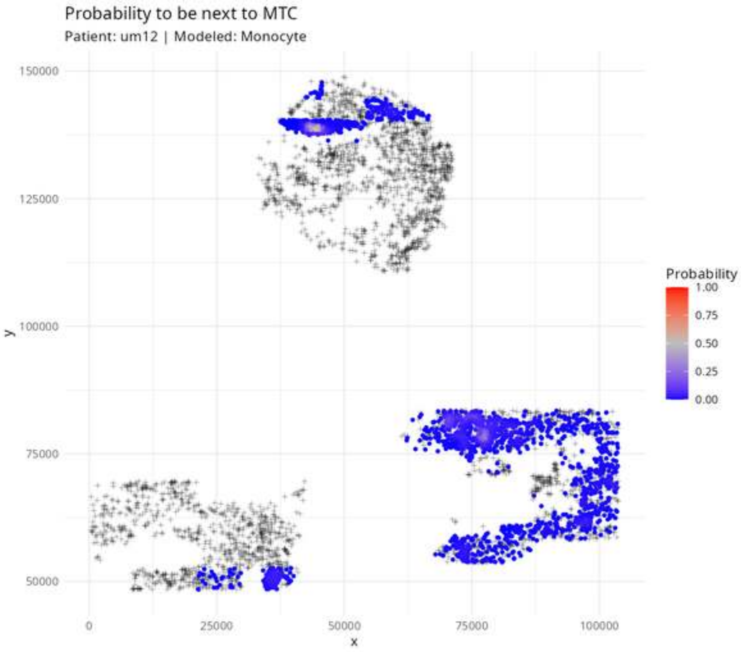

Probability to be next to MTC
Patient: um12 | Modeled: Monocyte
150000
125000
100000
75000
50000
y
0
25000
50000
75000
100000
x
Probability
1.00
0.75
0.50
0.25
0.00

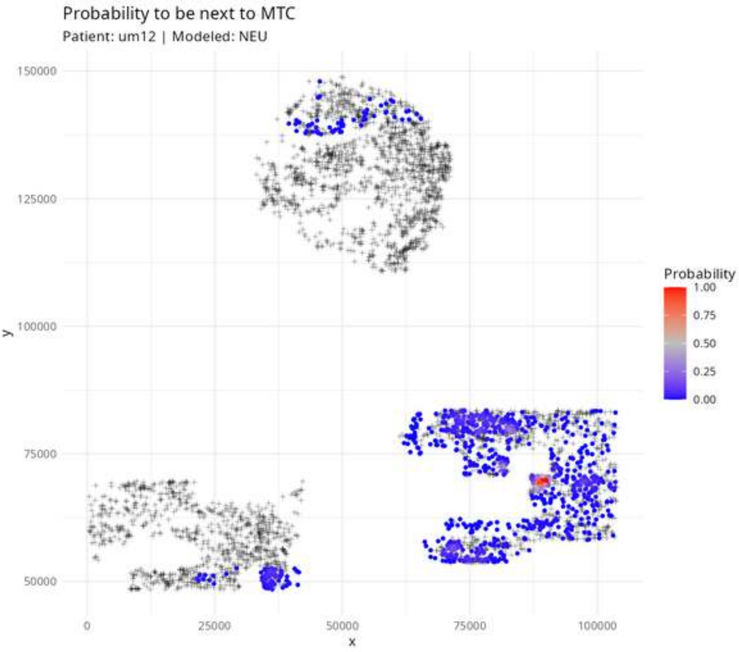
Probability to be next to MTC
Patient: um12 | Modeled: NEU
150000
125000
100000
75000
50000
y
0
25000
50000
75000
100000
x
Probability
1.00
0.75
0.50
0.25
0.00

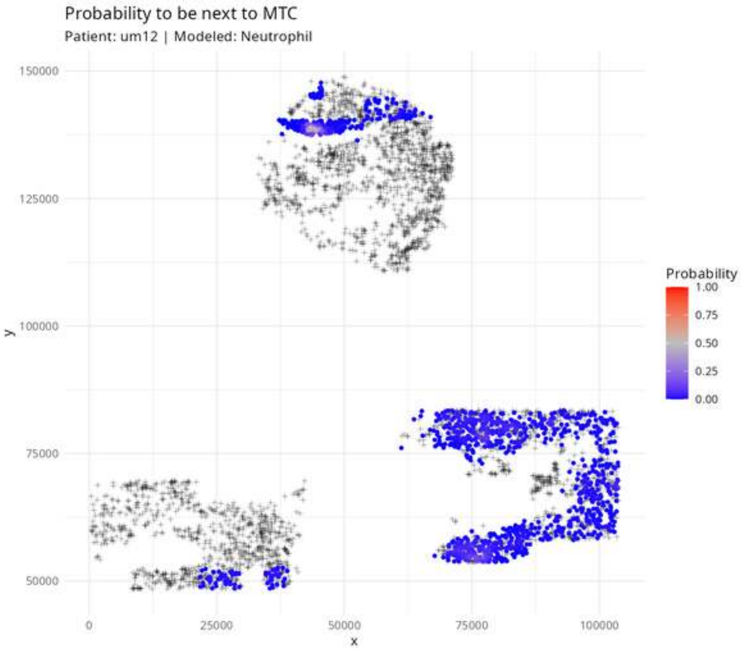

Probability to be next to MTC
Patient: um12 | Modeled: Neutrophil
y
150000
125000
100000
75000
50000
x
0
25000
50000
75000
100000
Probability
1.00
0.75
0.50
0.25
0.00

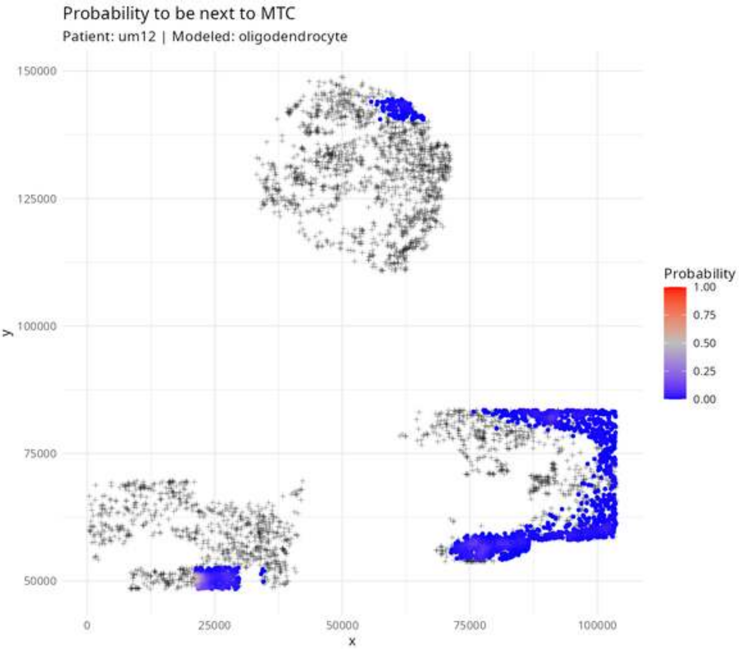
Probability to be next to MTC
Patient: um12 | Modeled: oligodendrocyte
150000
125000
100000
75000
50000
y
0
25000
50000
75000
100000
x
Probability
1.00
0.75
0.50
0.25
0.00

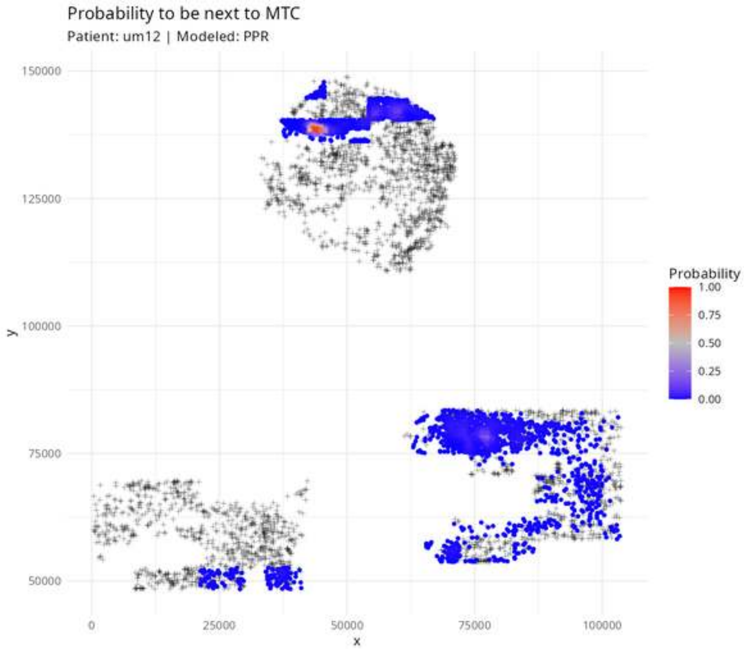
Probability to be next to MTC
Patient: um12 | Modeled: PPR
150000
125000
100000
75000
50000
y
0
25000
50000
75000
100000
x
Probability
1.00
0.75
0.50
0.25
0.00

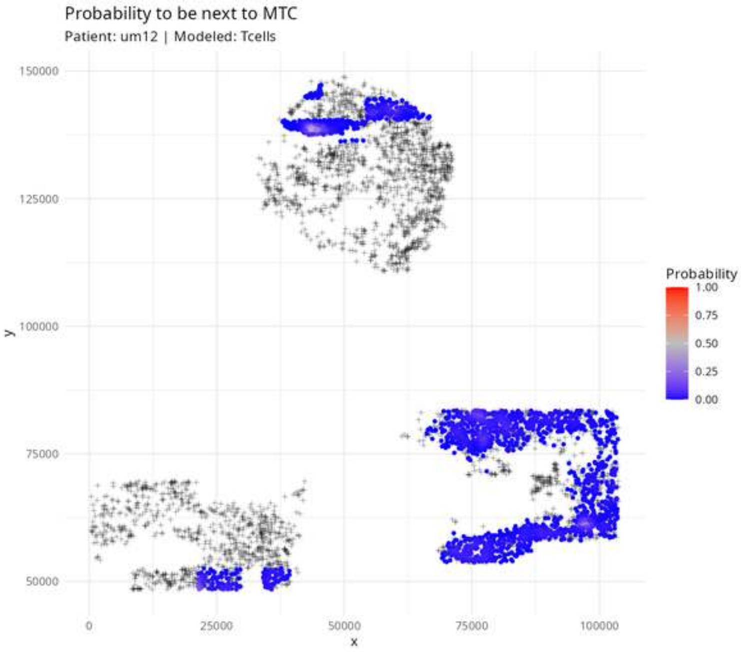
Probability to be next to MTC
Patient: um12 | Modeled: Tcells
150000
125000
100000
75000
50000
y
0
25000
50000
75000
100000
x
Probability
1.00
0.75
0.50
0.25
0.00

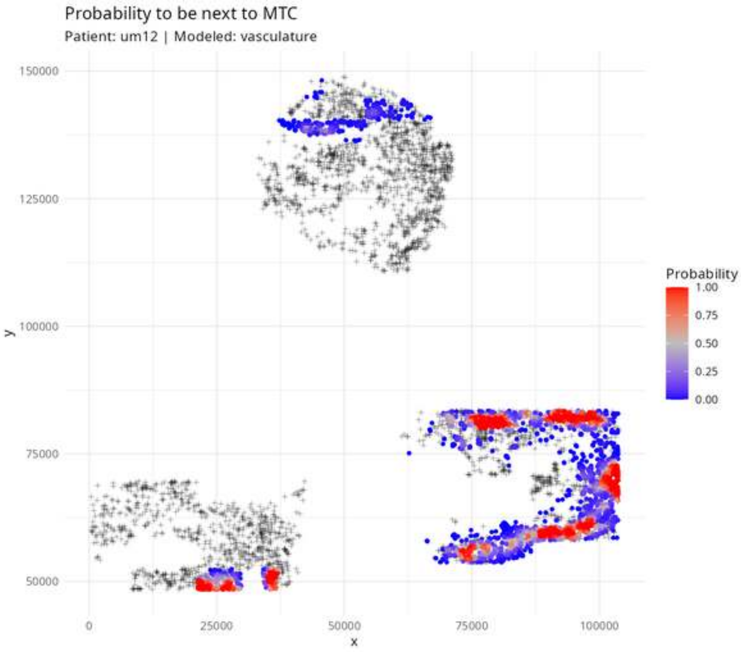
Probability to be next to MTC
Patient: um12 | Modeled: vasculature
150000
125000
100000
75000
50000
y
0
25000
50000
75000
100000
x
Probability
1.00
0.75
0.50
0.25
0.00

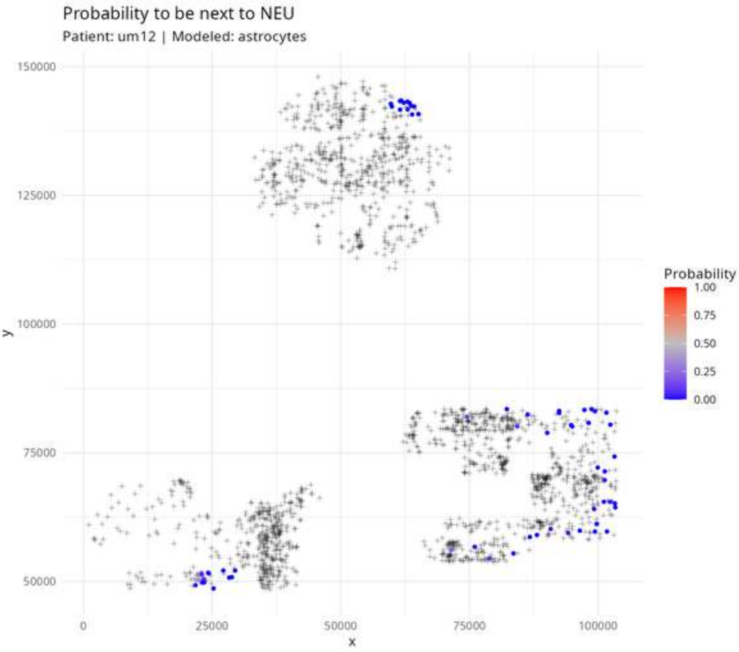
Probability to be next to NEU
Patient: um12 | Modeled: astrocytes
150000
125000
100000
75000
50000
y
0
25000
50000
75000
100000
x
Probability
1.00
0.75
0.50
0.25
0.00

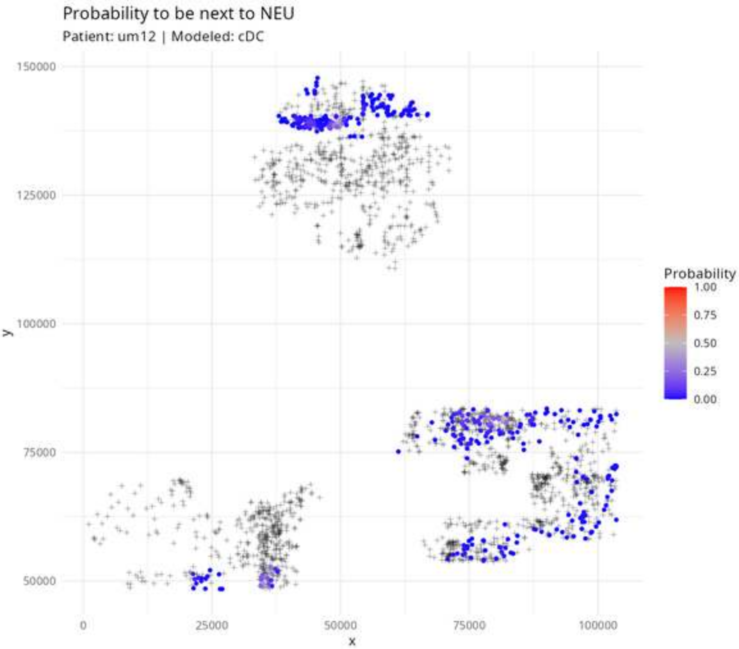
Probability to be next to NEU
Patient: um12 | Modeled: cDC
150000
125000
100000
75000
50000
y
0
25000
50000
75000
100000
x
Probability
1.00
0.75
0.50
0.25
0.00

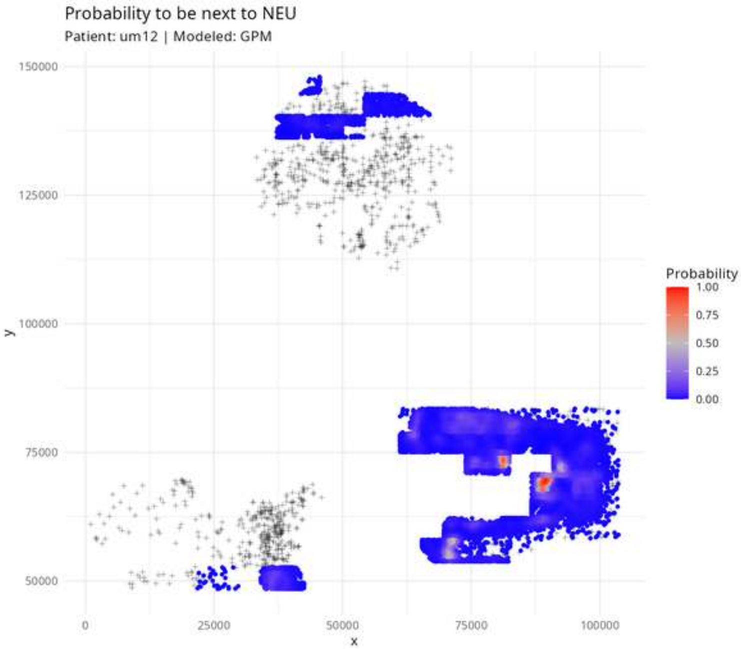

Probability to be next to NEU
Patient: um12 | Modeled: GPM
150000
125000
100000
75000
50000
y
0
25000
50000
75000
100000
x
Probability
1.00
0.75
0.50
0.25
0.00

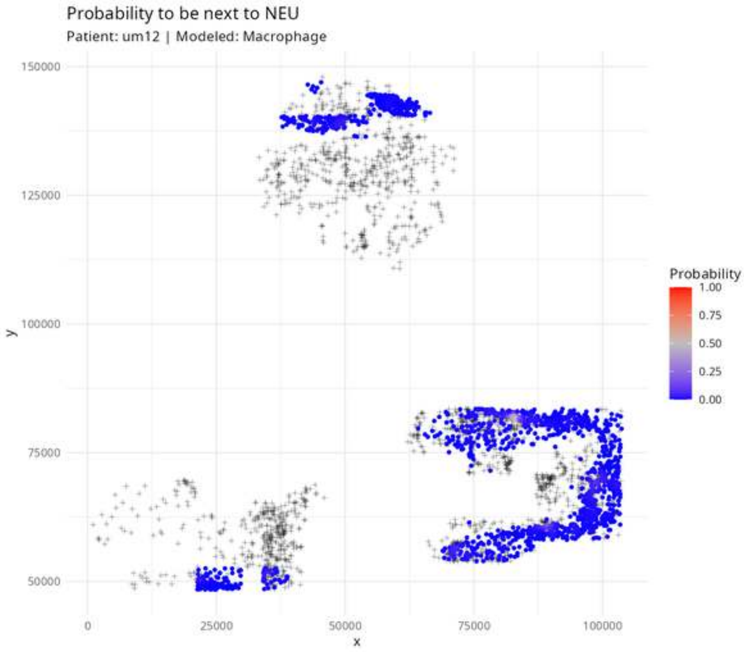

Probability to be next to NEU
Patient: um12 | Modeled: Macrophage
150000
125000
100000
75000
50000
y
0
25000
50000
75000
100000
x
Probability
1.00
0.75
0.50
0.25
0.00

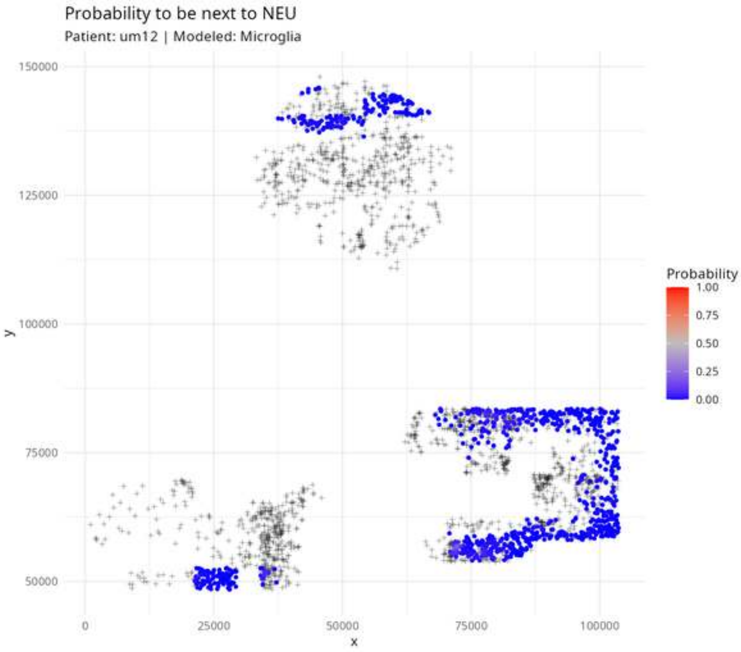
Probability to be next to NEU
Patient: um12 | Modeled: Microglia
150000
125000
100000
75000
50000
y
0
25000
50000
75000
100000
x
Probability
1.00
0.75
0.50
0.25
0.00

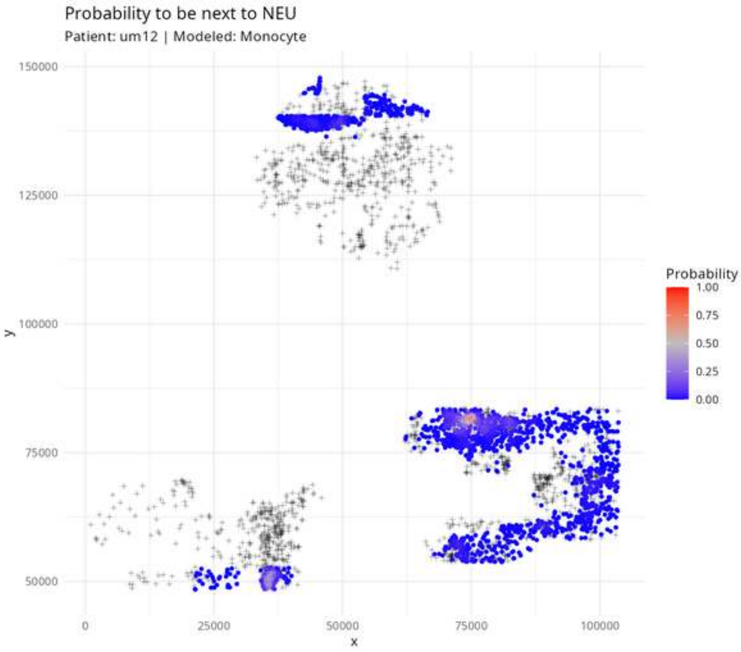

Probability to be next to NEU
Patient: um12 | Modeled: Monocyte
150000
125000
100000
75000
50000
y
0
25000
50000
75000
100000
x
Probability
1.00
0.75
0.50
0.25
0.00

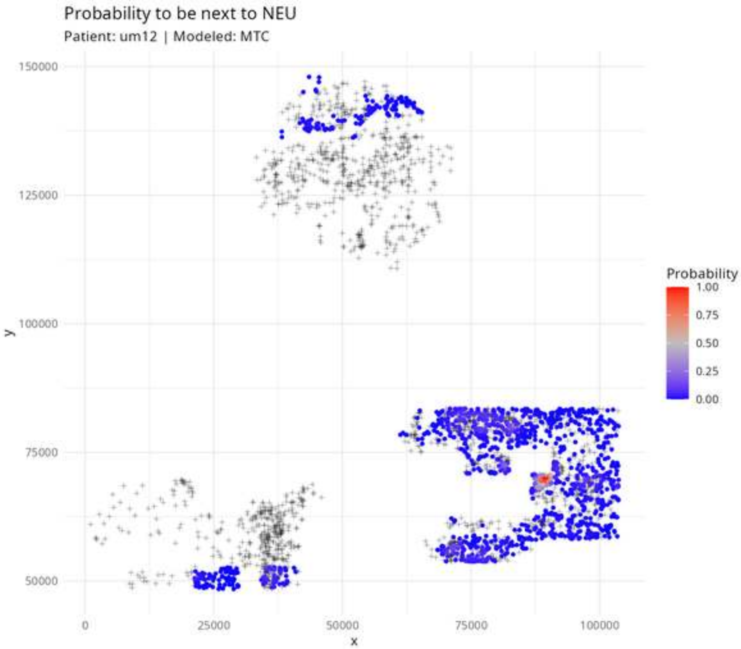
Probability to be next to NEU
Patient: um12 | Modeled: MTC
150000
125000
100000
75000
50000
y
0
25000
50000
75000
100000
x
Probability
1.00
0.75
0.50
0.25
0.00

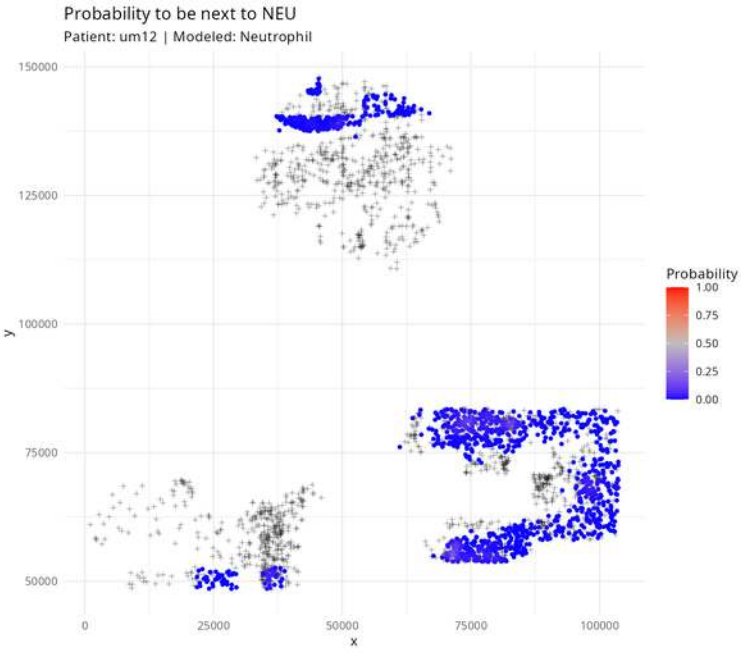
Probability to be next to NEU
Patient: um12 | Modeled: Neutrophil
150000
125000
100000
75000
50000
y
0
25000
50000
75000
100000
x
Probability
1.00
0.75
0.50
0.25
0.00

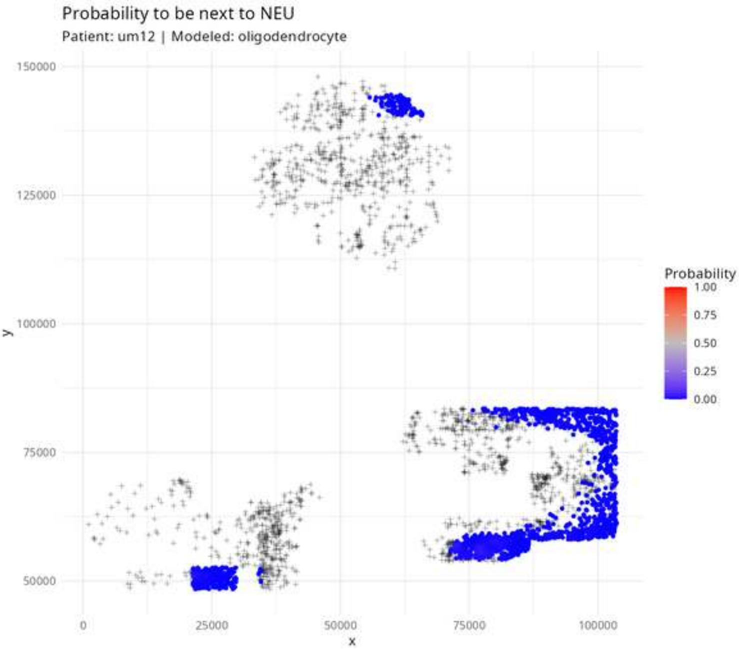
Probability to be next to NEU
Patient: um12 | Modeled: oligodendrocyte
150000
125000
100000
75000
50000
y
0
25000
50000
75000
100000
x
Probability
1.00
0.75
0.50
0.25
0.00

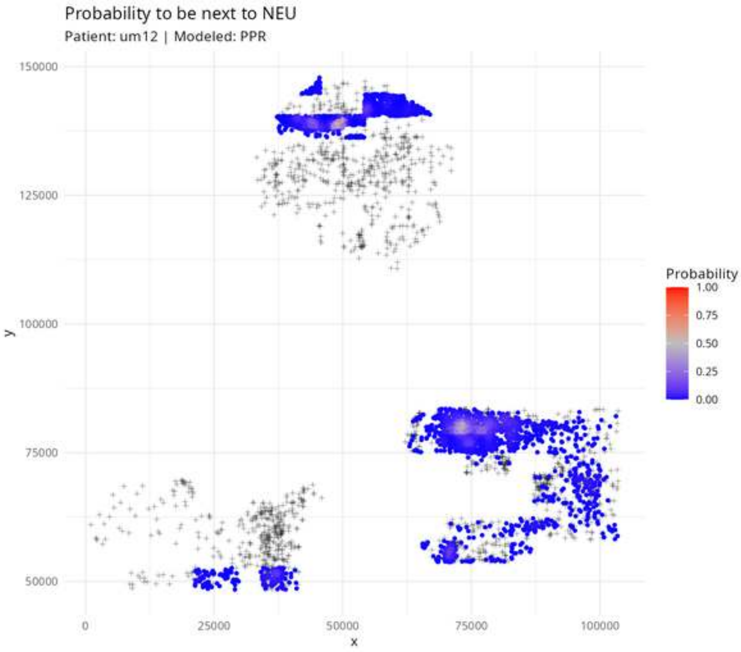
Probability to be next to NEU
Patient: um12 | Modeled: PPR
150000
125000
100000
75000
50000
y
0
25000
50000
75000
100000
x
Probability
1.00
0.75
0.50
0.25
0.00

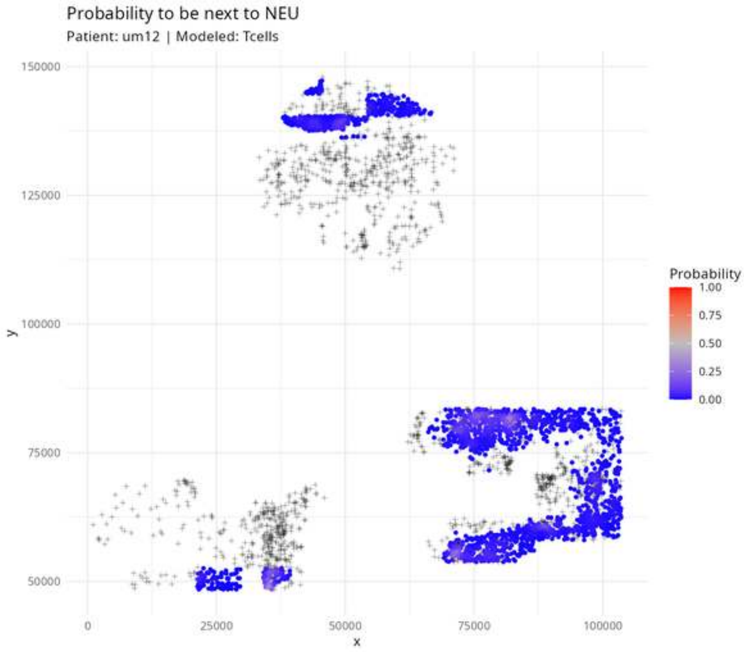

Probability to be next to NEU
Patient: um12 | Modeled: Tcells
150000
125000
100000
75000
50000
y
0
25000
50000
75000
100000
x
Probability
1.00
0.75
0.50
0.25
0.00

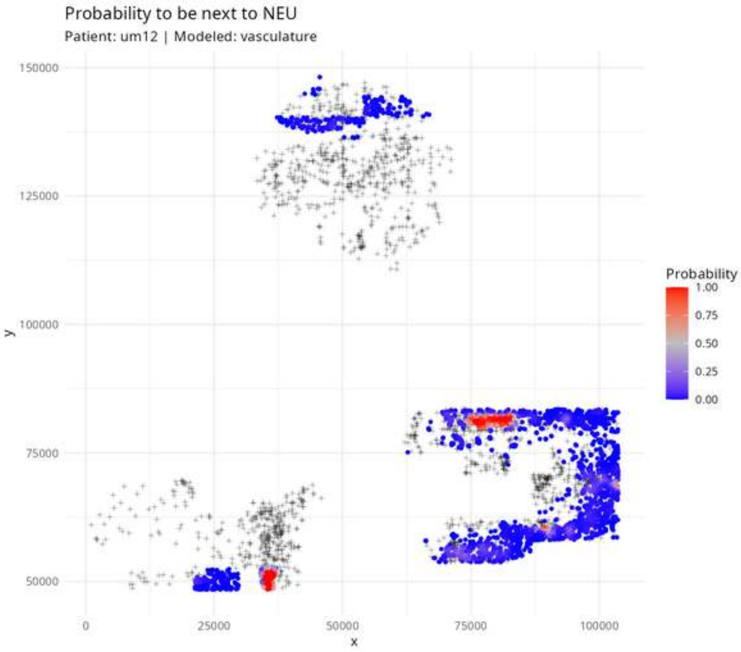

Probability to be next to NEU
Patient: um12 | Modeled: vasculature
150000
125000
100000
75000
50000
y
0
25000
50000
75000
100000
x
Probability
1.00
0.75
0.50
0.25
0.00

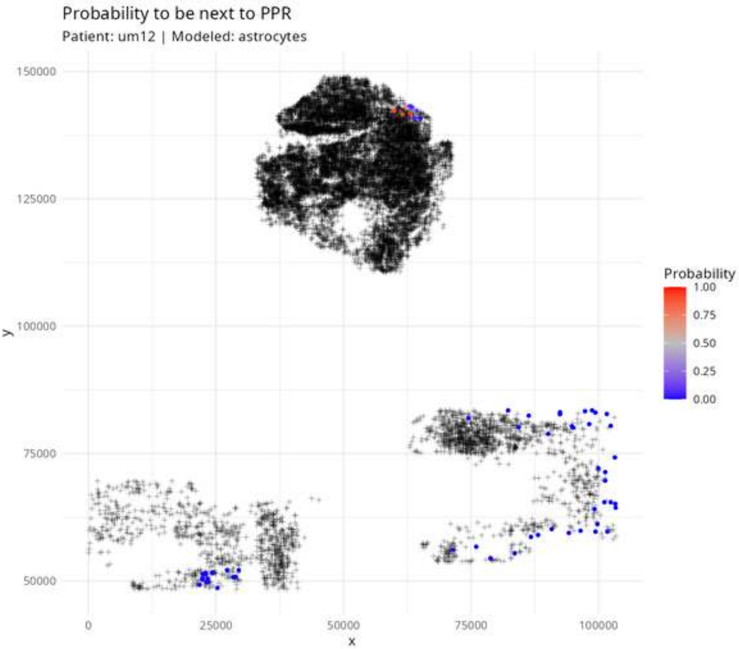

Probability to be next to PPR
Patient: um12 | Modeled: astrocytes
Probability
1.00
0.75
0.50
0.25
0.00
150000
125000
100000
75000
50000
y
0
25000
50000
75000
100000
x

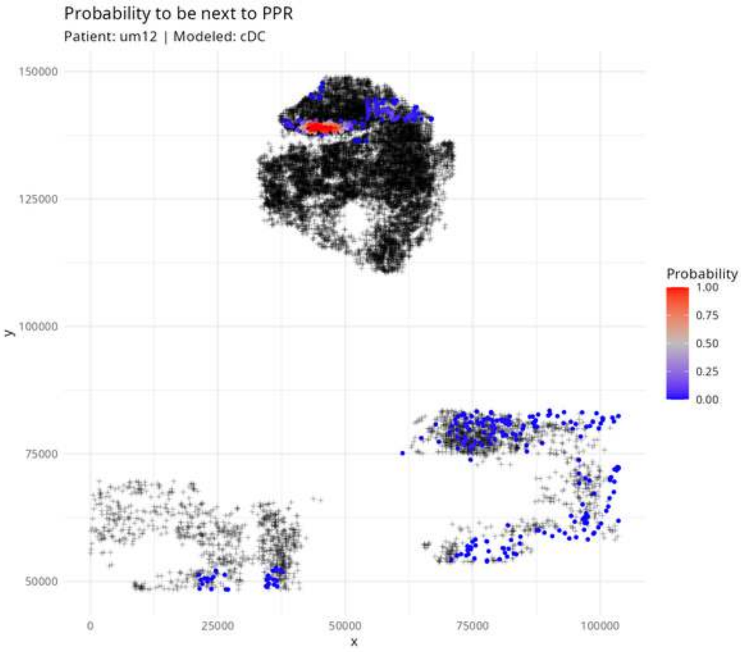

Probability to be next to PPR
Patient: um12 | Modeled: cDC
150000
125000
100000
75000
50000
y
0
25000
50000
75000
100000
x
Probability
1.00
0.75
0.50
0.25
0.00

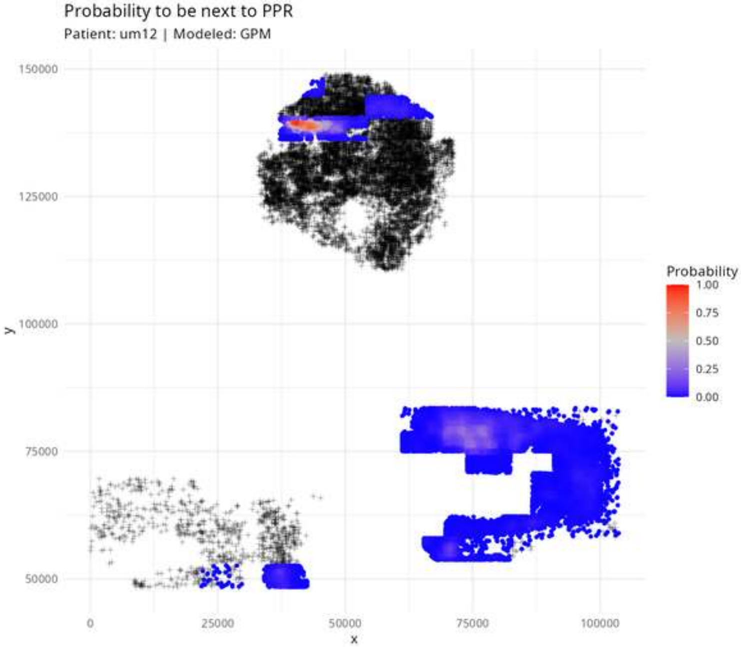

Probability to be next to PPR
Patient: um12 | Modeled: GPM
150000
125000
100000
75000
50000
y
0
25000
50000
75000
100000
x
Probability
1.00
0.75
0.50
0.25
0.00

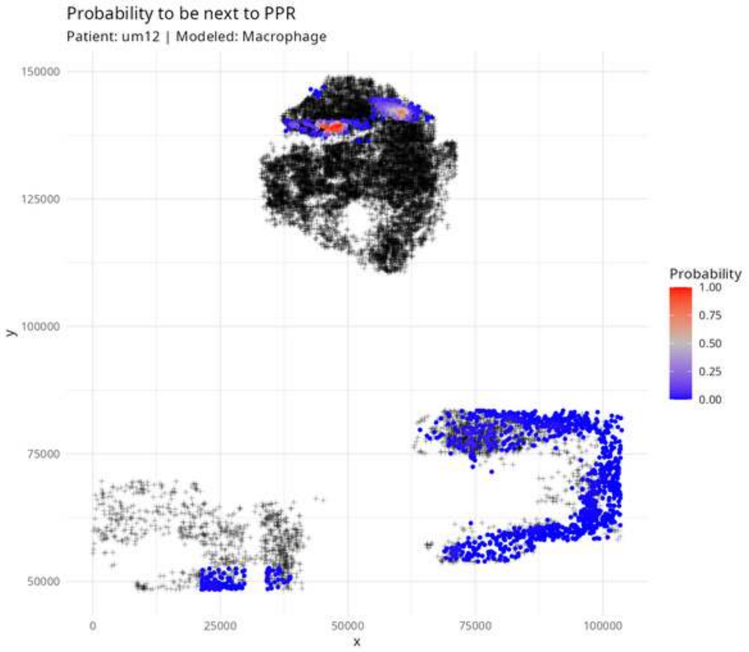

Probability to be next to PPR
Patient: um12 | Modeled: Macrophage
150000
125000
100000
75000
50000
y
0
25000
50000
75000
100000
x
Probability
1.00
0.75
0.50
0.25
0.00

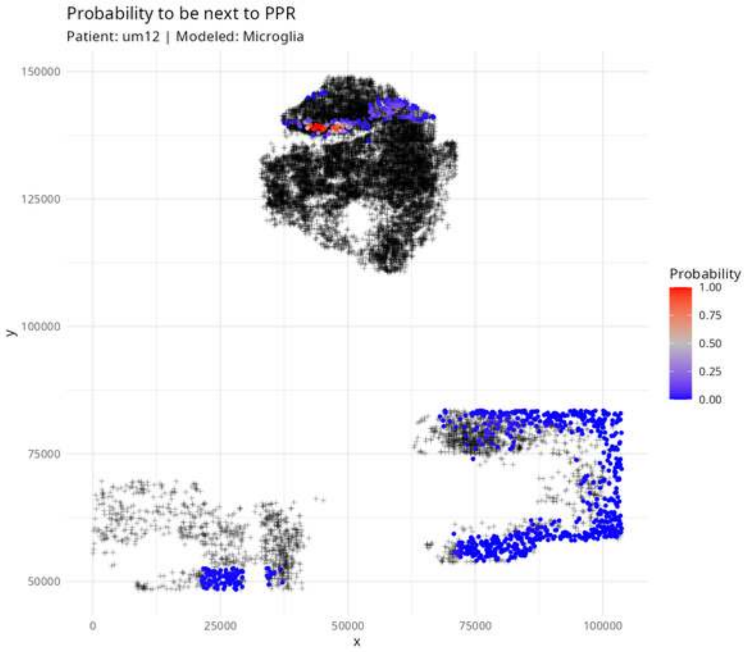
Probability to be next to PPR
Patient: um12 | Modeled: Microglia
150000
125000
100000
75000
50000
y
0
25000
50000
75000
100000
x
Probability
1.00
0.75
0.50
0.25
0.00

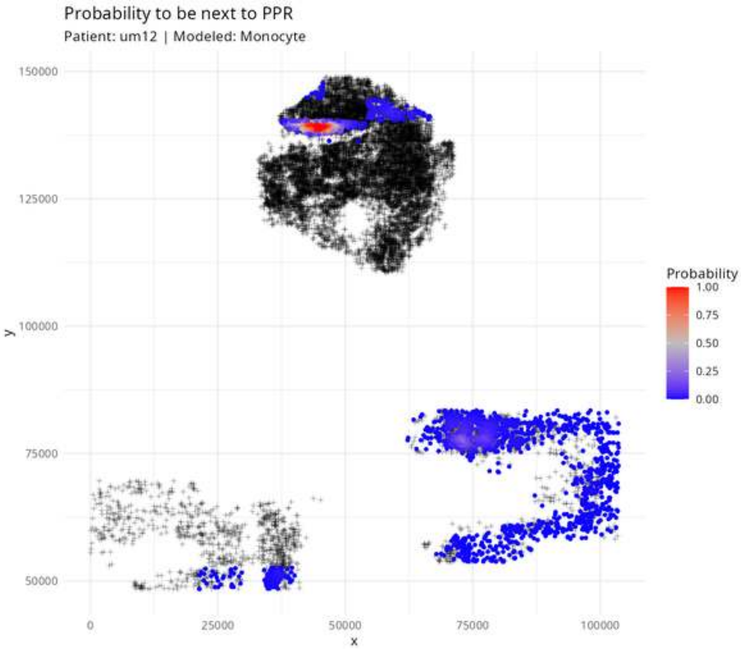

Probability to be next to PPR
Patient: um12 | Modeled: Monocyte
150000
125000
100000
75000
50000
y
0
25000
50000
75000
100000
x
Probability
1.00
0.75
0.50
0.25
0.00

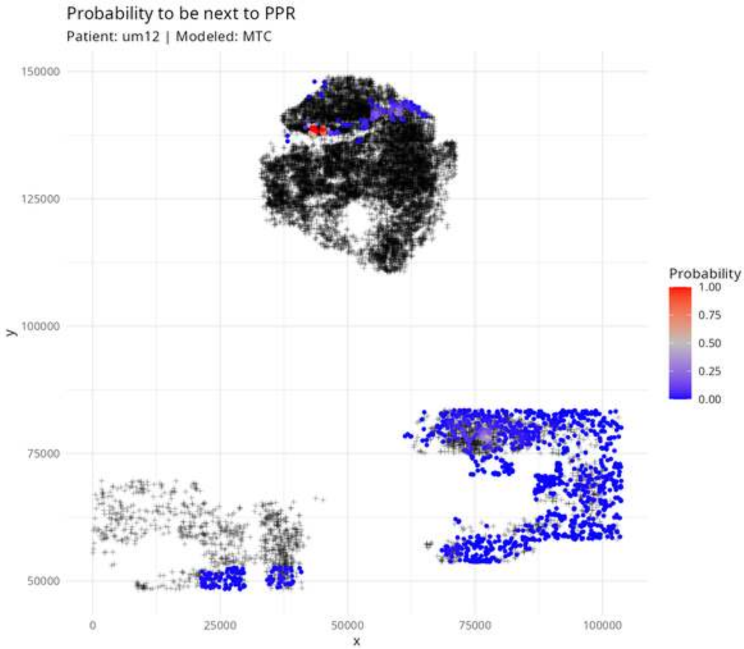

Probability to be next to PPR
Patient: um12 | Modeled: MTC
150000
125000
100000
75000
50000
y
0
25000
50000
75000
100000
x
Probability
1.00
0.75
0.50
0.25
0.00

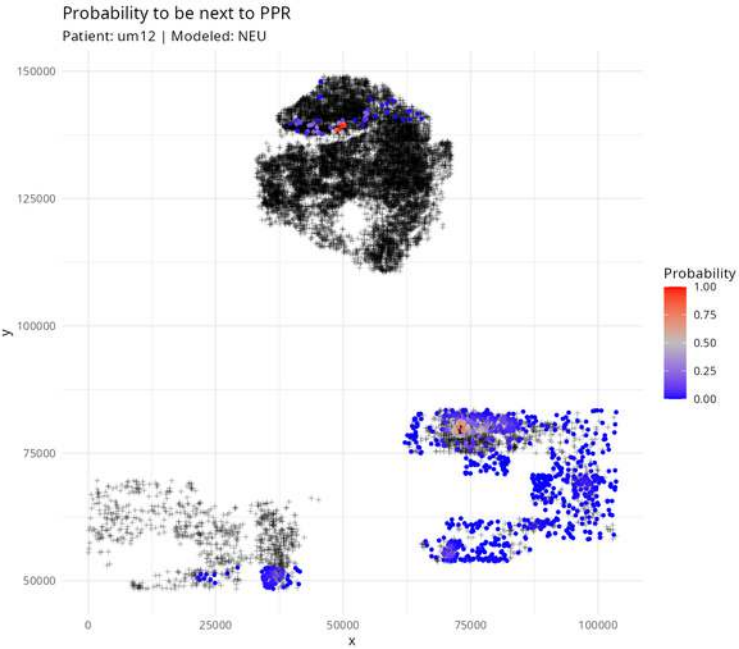

Probability to be next to PPR
Patient: um12 | Modeled: NEU
150000
125000
100000
75000
50000
y
0
25000
50000
75000
100000
x
Probability
1.00
0.75
0.50
0.25
0.00

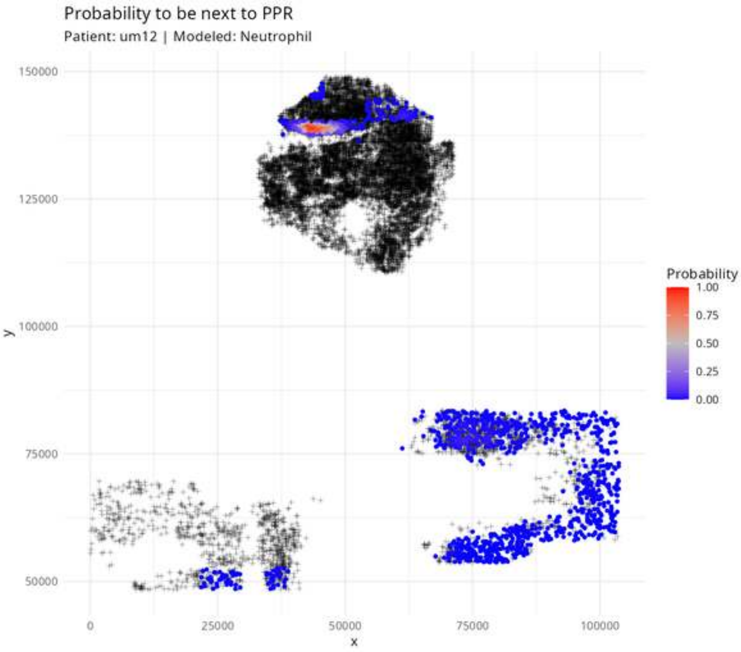
Probability to be next to PPR
Patient: um12 | Modeled: Neutrophil
150000
125000
100000
75000
50000
y
0
25000
50000
75000
100000
x
Probability
1.00
0.75
0.50
0.25
0.00

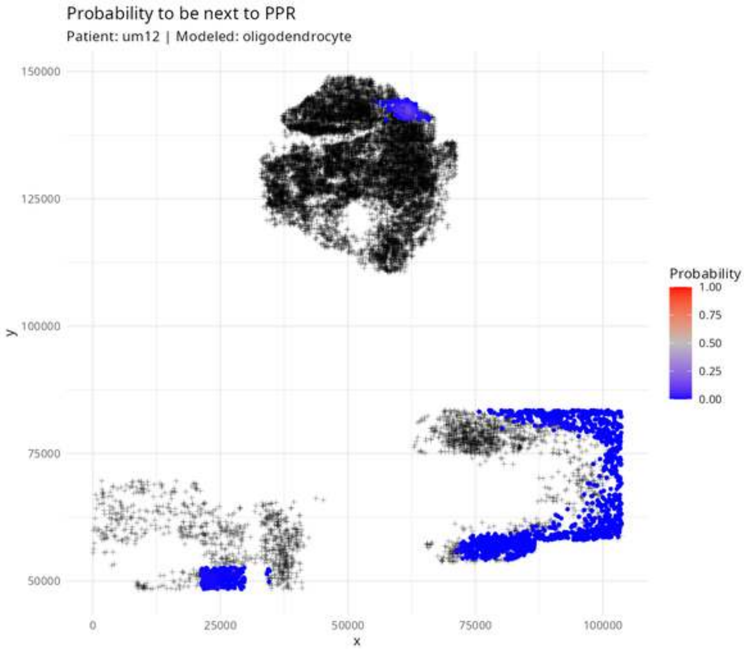

Probability to be next to PPR
Patient: um12 | Modeled: oligodendrocyte
150000
125000
100000
75000
50000
y
0
25000
50000
75000
100000
x
Probability
1.00
0.75
0.50
0.25
0.00

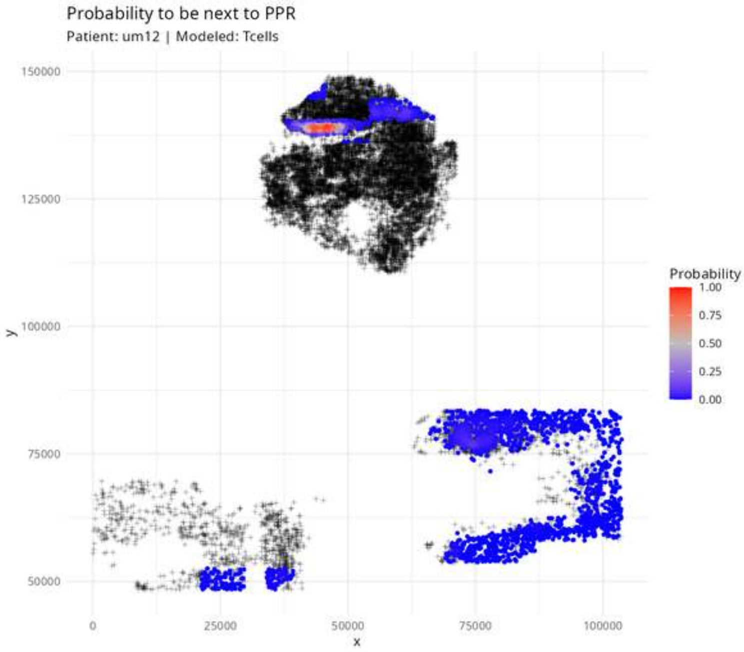
Probability to be next to PPR
Patient: um12 | Modeled: Tcells
150000
125000
100000
75000
50000
y
0
25000
50000
75000
100000
x
Probability
1.00
0.75
0.50
0.25
0.00

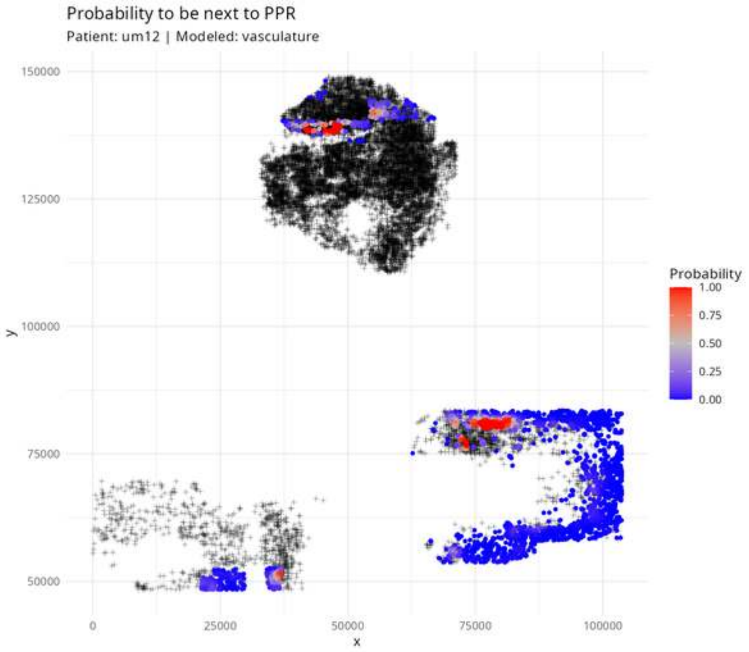

Probability to be next to PPR
Patient: um12 | Modeled: vasculature
150000
125000
100000
75000
50000
y
0
25000
50000
75000
100000
x
Probability
1.00
0.75
0.50
0.25
0.00

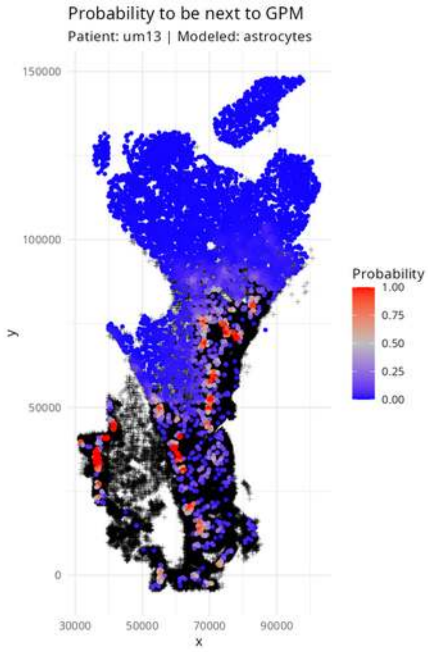
Probability to be next to GPM
Patient: um13 | Modeled: astrocytes
150000
100000
50000
0
y
30000
50000
70000
90000
x
Probability
1.00
0.75
0.50
0.25
0.00

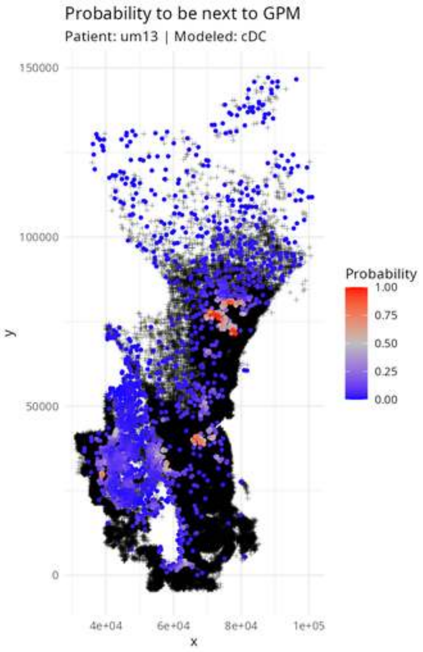

Probability to be next to GPM
Patient: um13 | Modeled: cDC
150000
100000
50000
0
y
4e+04
6e+04
8e+04
1e+05
x
Probability
1.00
0.75
0.50
0.25
0.00

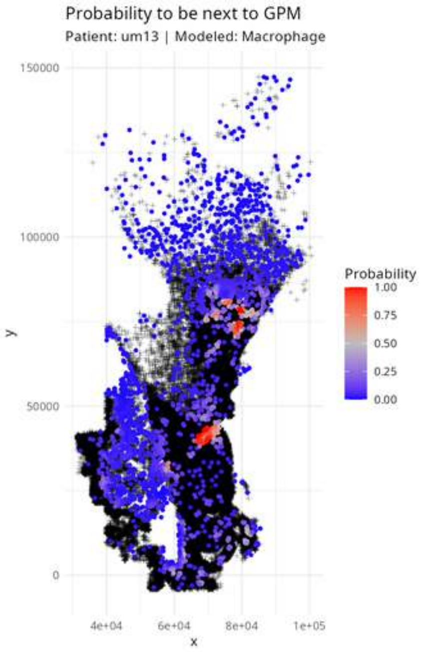

Probability to be next to GPM
Patient: um13 | Modeled: Macrophage
150000
100000
50000
0
y
4e+04
6e+04
8e+04
1e+05
x
Probability
1.00
0.75
0.50
0.25
0.00

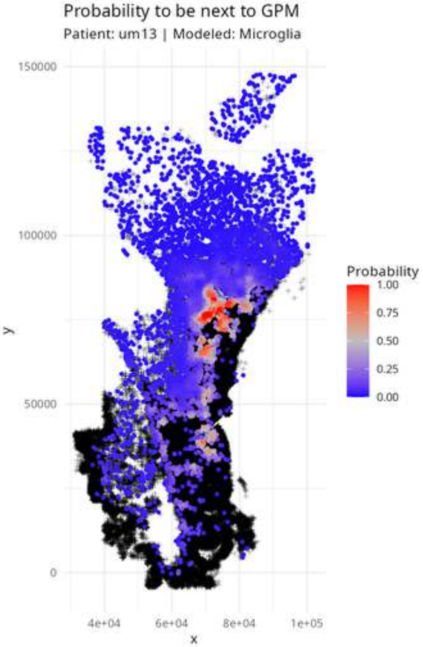

Probability to be next to GPM
Patient: um13 | Modeled: Microglia
150000
100000
50000
0
y
4e+04
6e+04
8e+04
1e+05
x
Probability
1.00
0.75
0.50
0.25
0.00

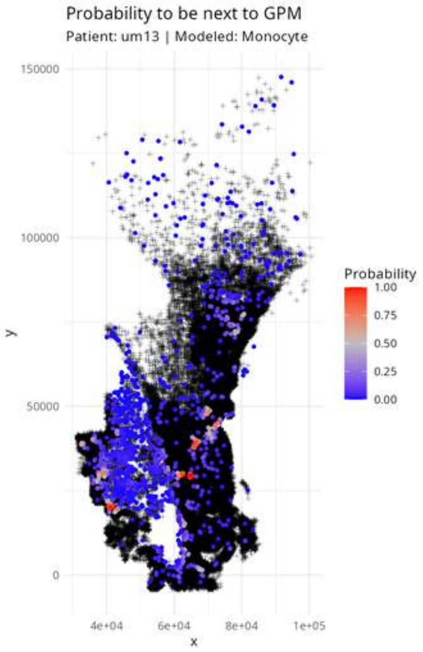
Probability to be next to GPM
Patient: um13 | Modeled: Monocyte
150000
100000
50000
0
4e+04
6e+04
8e+04
1e+05
x
y
Probability
1.00
0.75
0.50
0.25
0.00

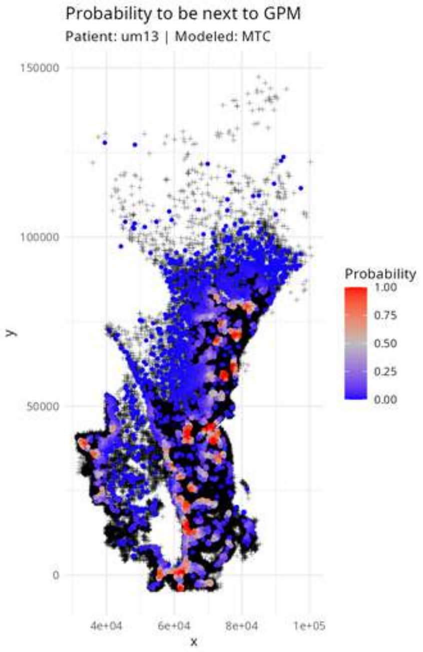
Probability to be next to GPM
Patient: um13 | Modeled: MTC
150000
100000
50000
0
y
4e+04
6e+04
8e+04
1e+05
x
Probability
1.00
0.75
0.50
0.25
0.00

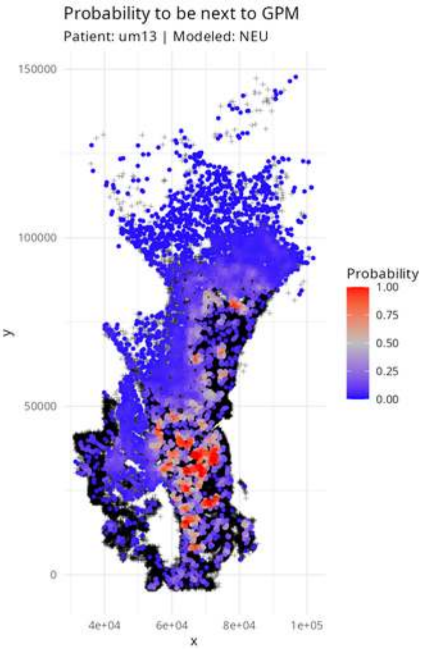

Probability to be next to GPM
Patient: um13 | Modeled: NEU
150000
100000
50000
0
y
4e+04
6e+04
8e+04
1e+05
x
Probability
1.00
0.75
0.50
0.25
0.00

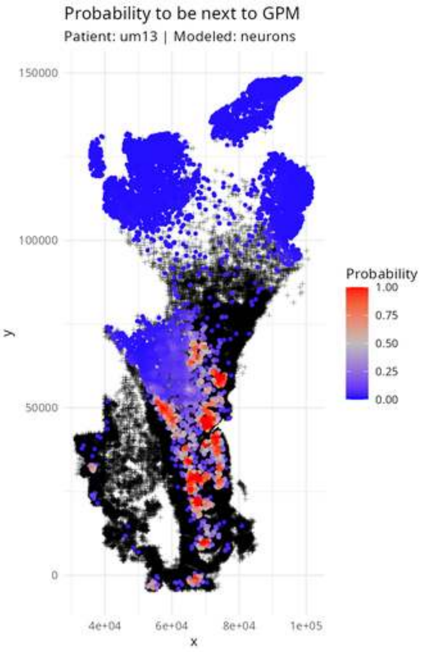

Probability to be next to GPM
Patient: um13 | Modeled: neurons
150000
100000
50000
0
y
4e+04
6e+04
8e+04
1e+05
x
Probability
1.00
0.75
0.50
0.25
0.00

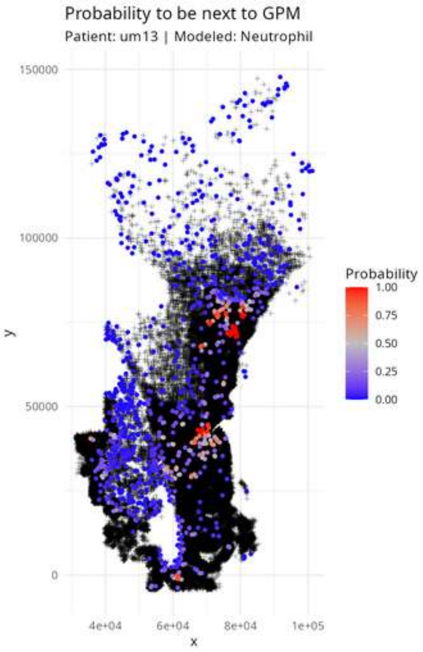

Probability to be next to GPM
Patient: um13 | Modeled: Neutrophil
150000
100000
50000
0
y
4e+04
6e+04
8e+04
1e+05
x
Probability
1.00
0.75
0.50
0.25
0.00

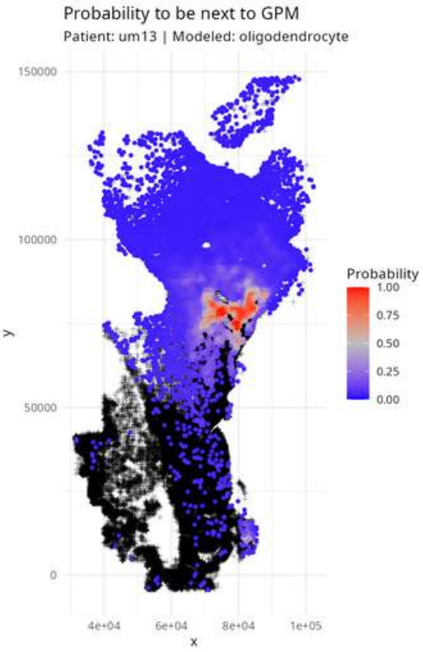

Probability to be next to GPM
Patient: um13 | Modeled: oligodendrocyte
150000
100000
50000
0
y
4e+04
6e+04
8e+04
1e+05
x
Probability
1.00
0.75
0.50
0.25
0.00

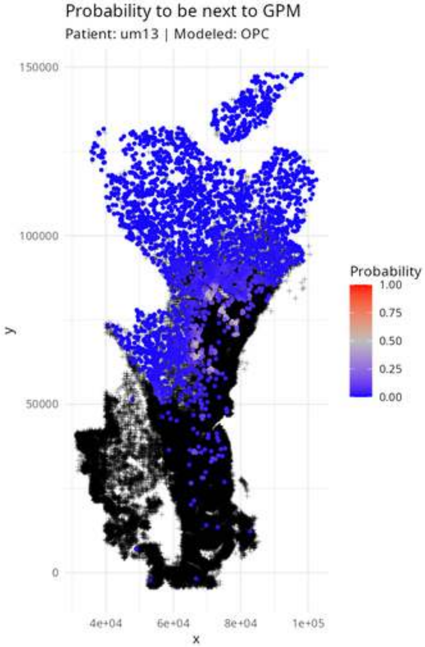

Probability to be next to GPM
Patient: um13 | Modeled: OPC
150000
100000
50000
0
y
4e+04
6e+04
8e+04
1e+05
x
Probability
1.00
0.75
0.50
0.25
0.00

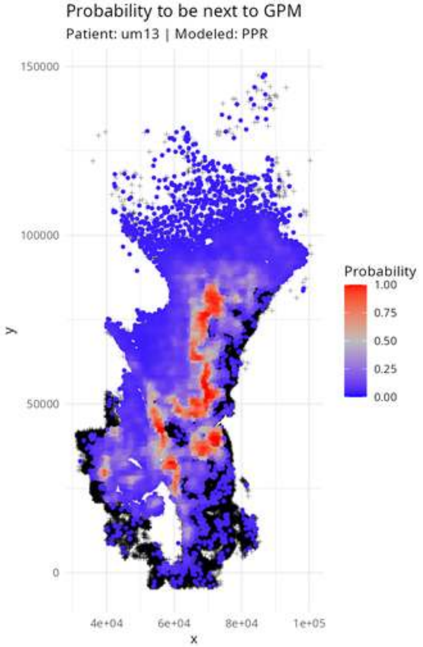

Probability to be next to GPM
Patient: um13 | Modeled: PPR
150000
100000
50000
0
y
4e+04
6e+04
8e+04
1e+05
x
Probability
1.00
0.75
0.50
0.25
0.00

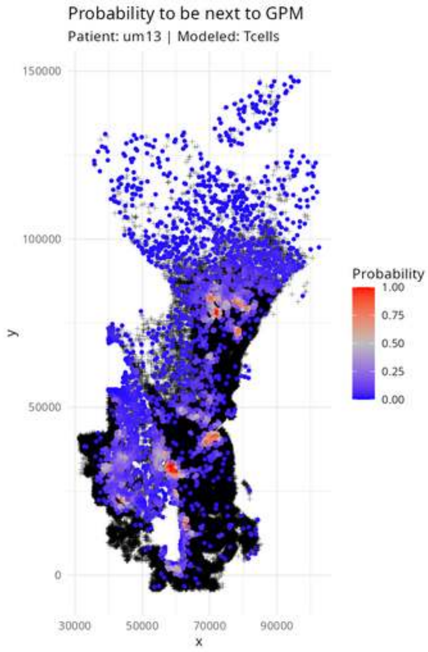
Probability to be next to GPM
Patient: um13 | Modeled: Tcells
150000
100000
50000
0
30000
50000
70000
90000
x
y
Probability
1.00
0.75
0.50
0.25
0.00

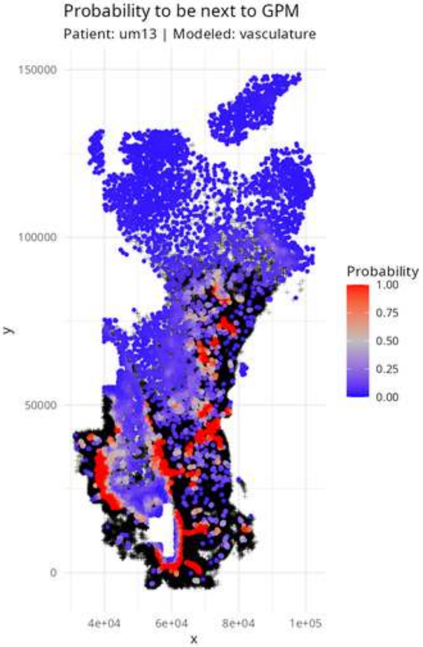

Probability to be next to GPM
Patient: um13 | Modeled: vasculature
150000
100000
50000
0
y
4e+04
6e+04
8e+04
1e+05
x
Probability
1.00
0.75
0.50
0.25
0.00

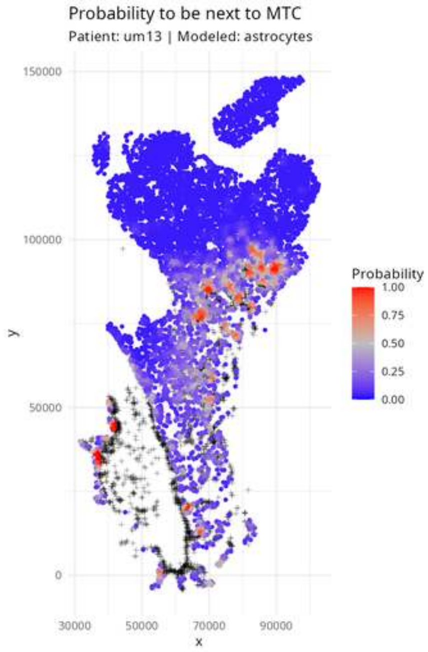

Probability to be next to MTC
Patient: um13 | Modeled: astrocytes
150000
100000
50000
0
y
30000
50000
70000
90000
x
Probability
1.00
0.75
0.50
0.25
0.00

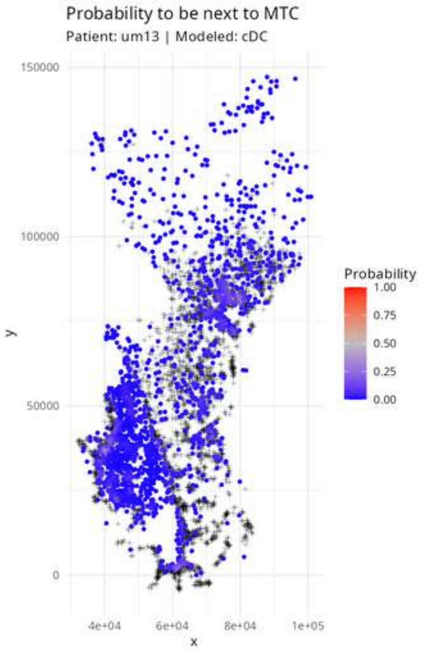

Probability to be next to MTC
Patient: um13 | Modeled: cDC
150000
100000
50000
0
y
4e+04
6e+04
8e+04
1e+05
x
Probability
1.00
0.75
0.50
0.25
0.00

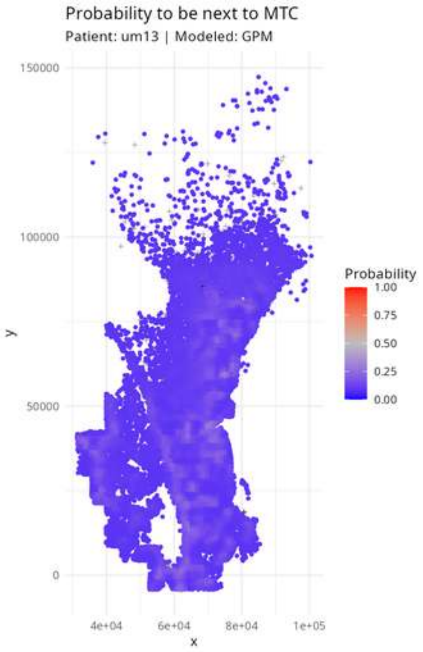

Probability to be next to MTC
Patient: um13 | Modeled: GPM
150000
100000
50000
0
y
4e+04
6e+04
8e+04
1e+05
x
Probability
1.00
0.75
0.50
0.25
0.00

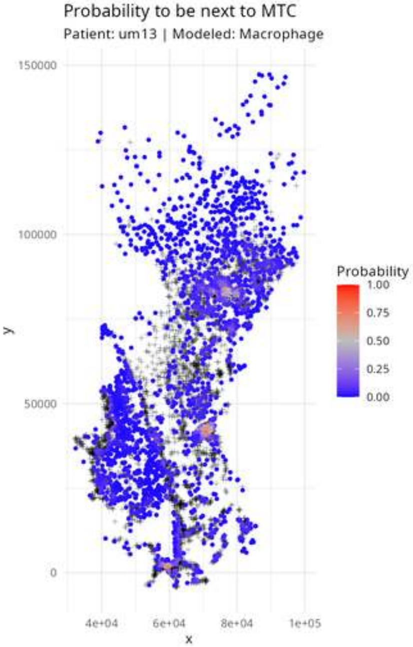
Probability to be next to MTC
Patient: um13 | Modeled: Macrophage
150000
100000
50000
0
y
4e+04
6e+04
8e+04
1e+05
x
Probability
1.00
0.75
0.50
0.25
0.00

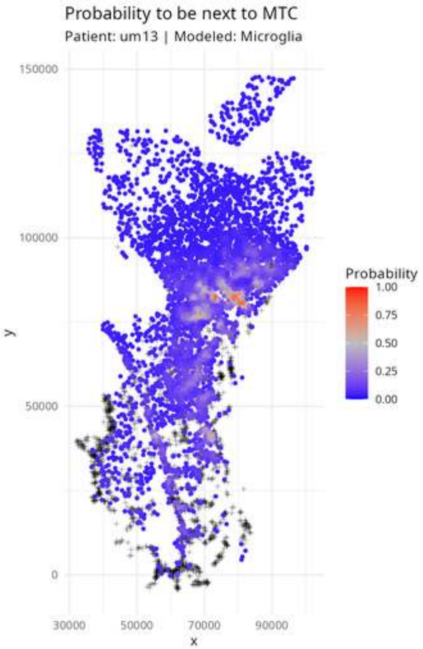

Probability to be next to MTC
Patient: um13 | Modeled: Microglia
150000
100000
50000
0
y
30000
50000
70000
90000
x
Probability
1.00
0.75
0.50
0.25
0.00

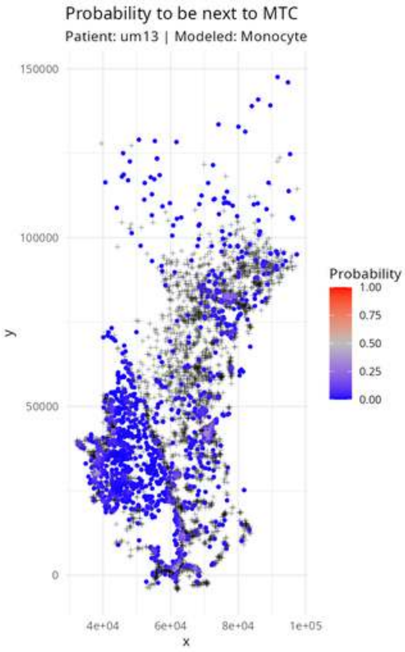

Probability to be next to MTC
Patient: um13 | Modeled: Monocyte
150000
100000
50000
0
y
4e+04
6e+04
8e+04
1e+05
x
Probability
1.00
0.75
0.50
0.25
0.00

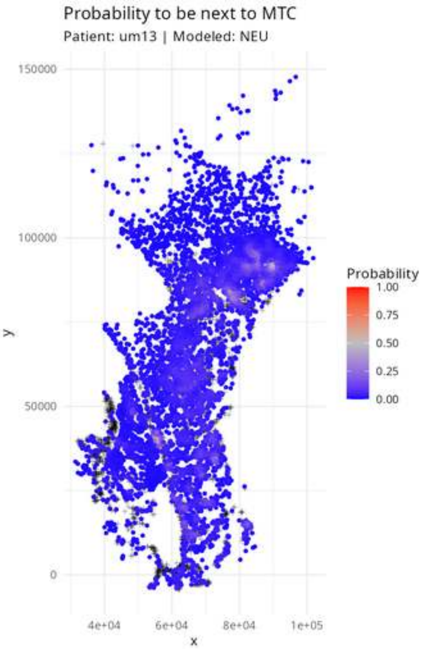

Probability to be next to MTC
Patient: um13 | Modeled: NEU
150000
100000
50000
0
y
4e+04
6e+04
8e+04
1e+05
x
Probability
1.00
0.75
0.50
0.25
0.00

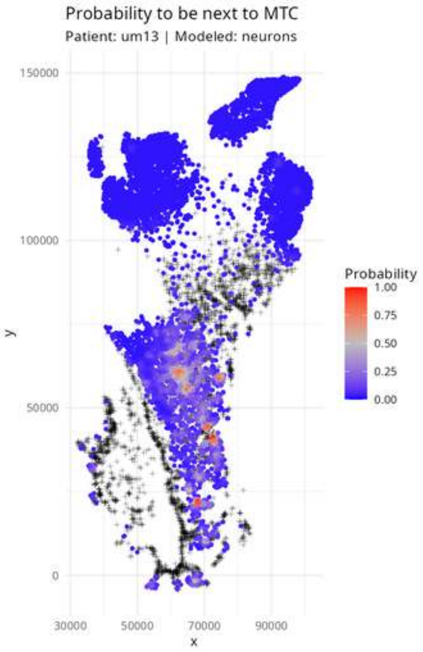

Probability to be next to MTC
Patient: um13 | Modeled: neurons
150000
100000
50000
0
y
30000
50000
70000
90000
x
Probability
1.00
0.75
0.50
0.25
0.00

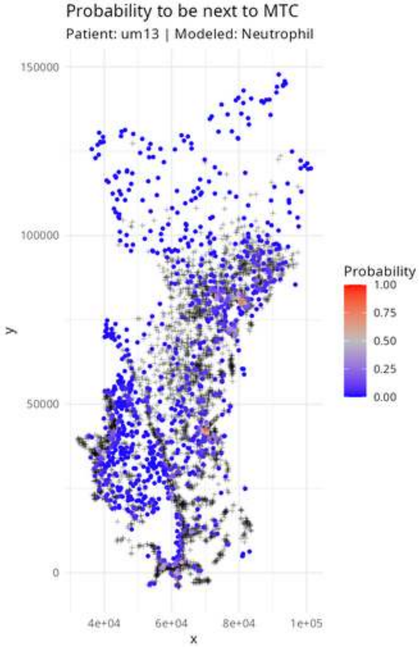
Probability to be next to MTC
Patient: um13 | Modeled: Neutrophil
150000
100000
50000
0
y
4e+04
6e+04
8e+04
1e+05
x
Probability
1.00
0.75
0.50
0.25
0.00

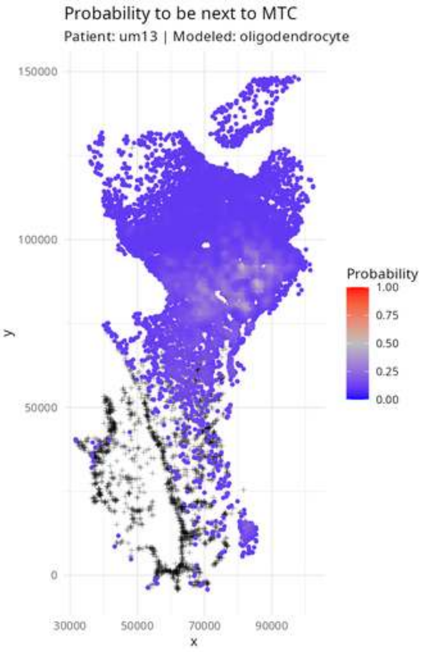

Probability to be next to MTC
Patient: um13 | Modeled: oligodendrocyte
150000
100000
50000
0
y
30000
50000
70000
90000
x
Probability
1.00
0.75
0.50
0.25
0.00

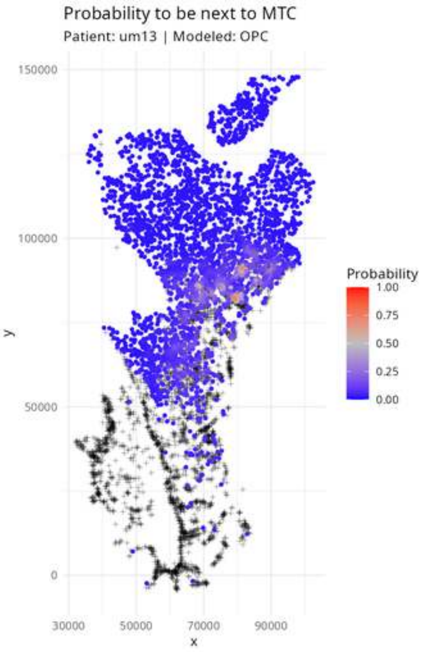

Probability to be next to MTC
Patient: um13 | Modeled: OPC
150000
100000
50000
0
30000
50000
70000
90000
x
y
Probability
1.00
0.75
0.50
0.25
0.00

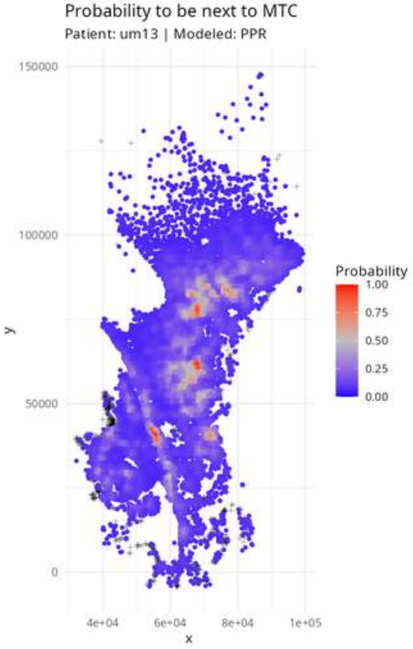

Probability to be next to MTC
Patient: um13 | Modeled: PPR
150000
100000
50000
0
y
4e+04
6e+04
8e+04
1e+05
x
Probability
1.00
0.75
0.50
0.25
0.00

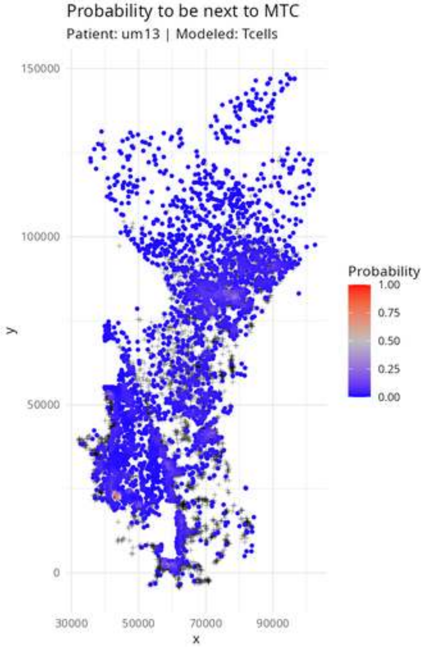

Probability to be next to MTC
Patient: um13 | Modeled: Tcells
150000
100000
50000
0
y
30000
50000
70000
90000
x
Probability
1.00
0.75
0.50
0.25
0.00

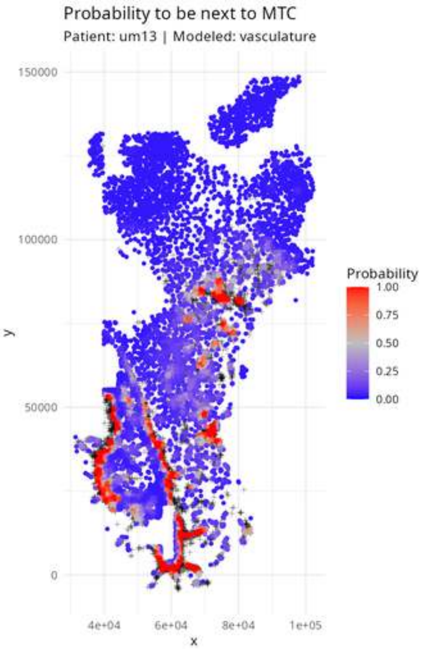

Probability to be next to MTC
Patient: um13 | Modeled: vasculature
150000
100000
50000
0
y
4e+04
6e+04
8e+04
1e+05
x
Probability
1.00
0.75
0.50
0.25
0.00

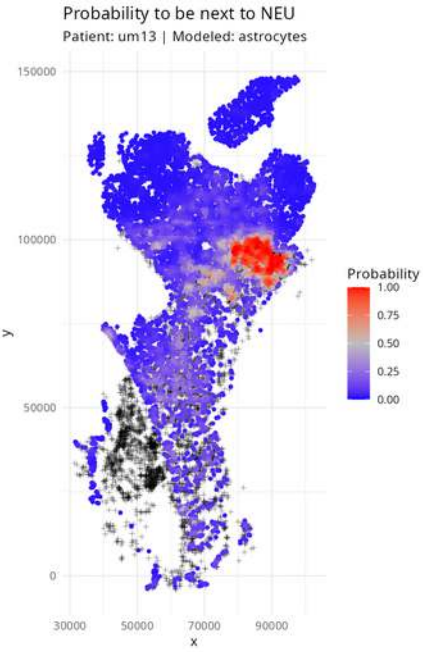

Probability to be next to NEU
Patient: um13 | Modeled: astrocytes
150000
100000
50000
0
y
30000
50000
70000
90000
x
Probability
1.00
0.75
0.50
0.25
0.00

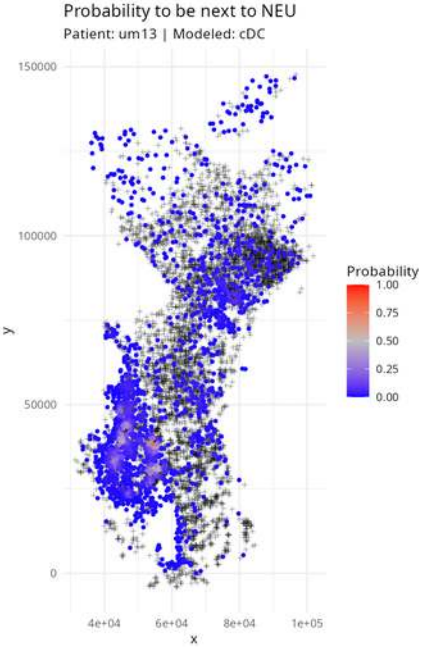
Probability to be next to NEU
Patient: um13 | Modeled: cDC
150000
100000
50000
0
4e+04
6e+04
8e+04
1e+05
x
y
Probability
1.00
0.75
0.50
0.25
0.00

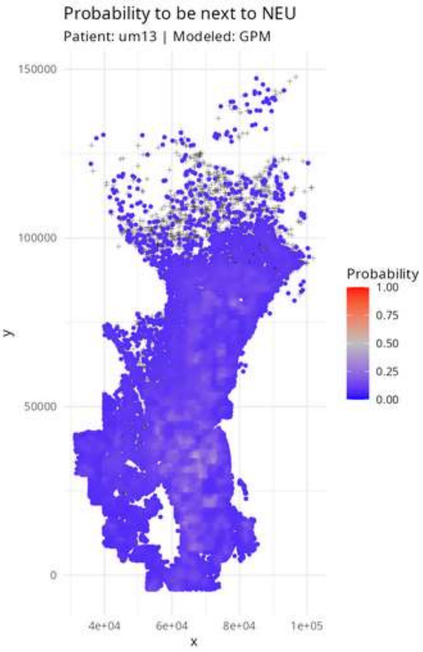

Probability to be next to NEU
Patient: um13 | Modeled: GPM
150000
100000
50000
0
y
4e+04
6e+04
8e+04
1e+05
x
Probability
1.00
0.75
0.50
0.25
0.00

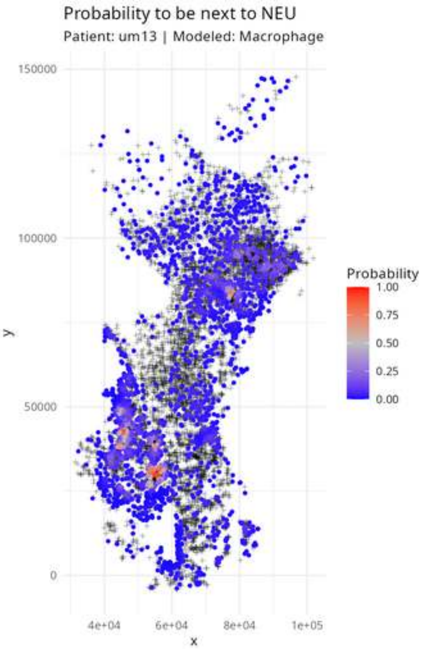

Probability to be next to NEU
Patient: um13 | Modeled: Macrophage
150000
100000
50000
0
y
4e+04
6e+04
8e+04
1e+05
x
Probability
1.00
0.75
0.50
0.25
0.00

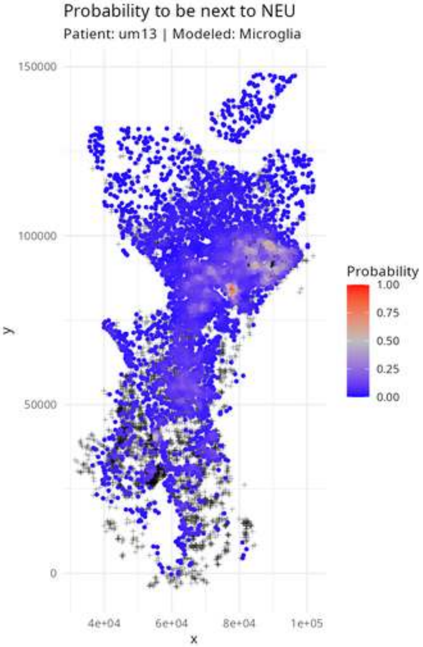

Probability to be next to NEU
Patient: um13 | Modeled: Microglia
150000
100000
50000
0
y
4e+04
6e+04
8e+04
1e+05
x
Probability
1.00
0.75
0.50
0.25
0.00

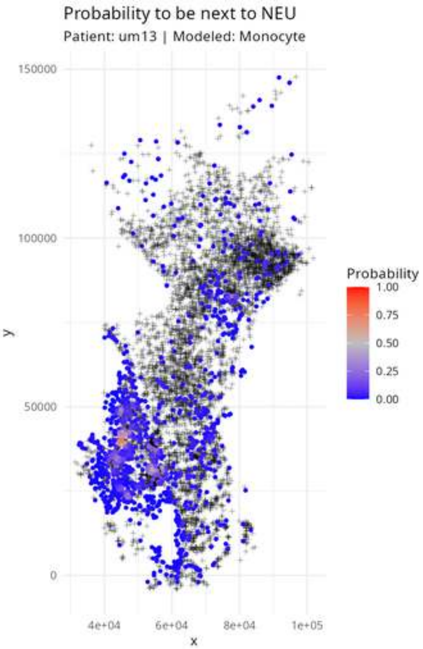
Probability to be next to NEU
Patient: um13 | Modeled: Monocyte
150000
100000
50000
0
y
4e+04
6e+04
8e+04
1e+05
x
Probability
1.00
0.75
0.50
0.25
0.00

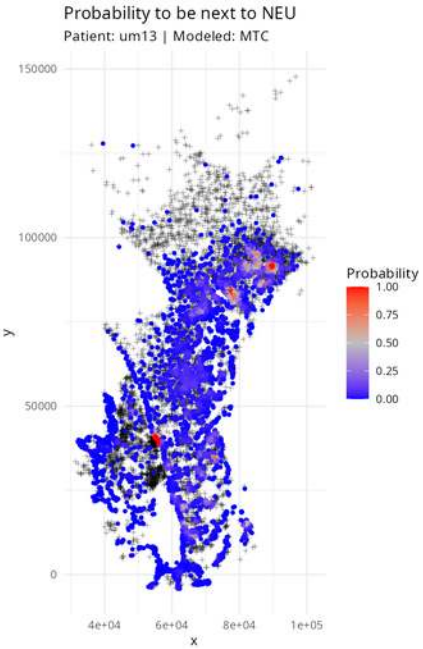
Probability to be next to NEU
Patient: um13 | Modeled: MTC
150000
100000
50000
0
y
4e+04
6e+04
8e+04
1e+05
x
Probability
1.00
0.75
0.50
0.25
0.00

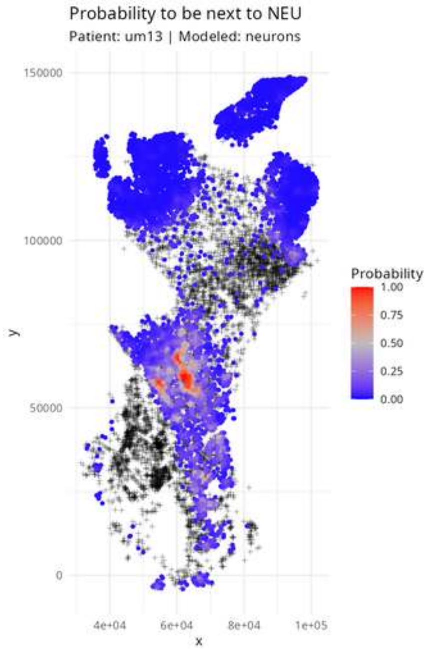
Probability to be next to NEU
Patient: um13 | Modeled: neurons
150000
100000
50000
0
y
4e+04
6e+04
8e+04
1e+05
x
Probability
1.00
0.75
0.50
0.25
0.00

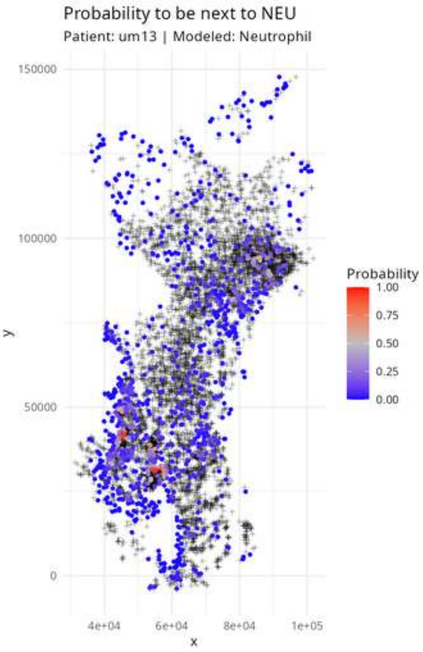
Probability to be next to NEU
Patient: um13 | Modeled: Neutrophil
150000
100000
50000
0
4e+04
6e+04
8e+04
1e+05
x
y
Probability
1.00
0.75
0.50
0.25
0.00

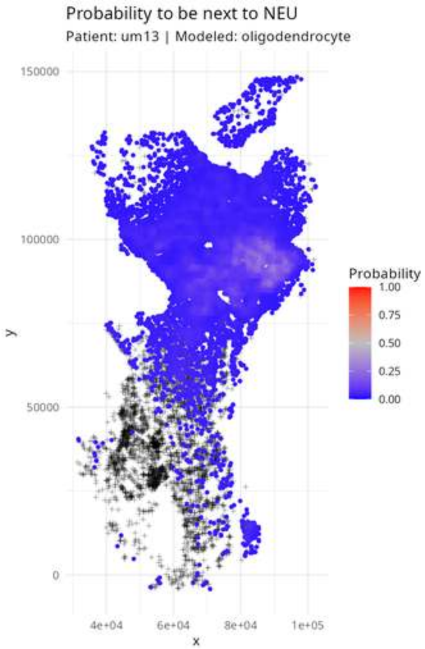

Probability to be next to NEU
Patient: um13 | Modeled: oligodendrocyte
150000
100000
50000
0
y
4e+04
6e+04
8e+04
1e+05
x
Probability
1.00
0.75
0.50
0.25
0.00

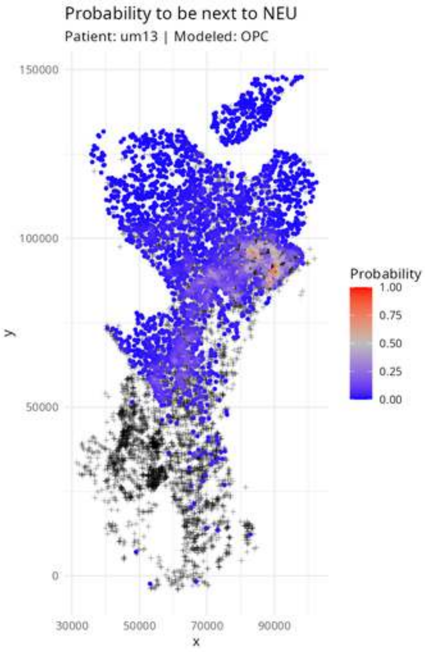
Probability to be next to NEU
Patient: um13 | Modeled: OPC
150000
100000
50000
0
y
30000
50000
70000
90000
x
Probability
1.00
0.75
0.50
0.25
0.00

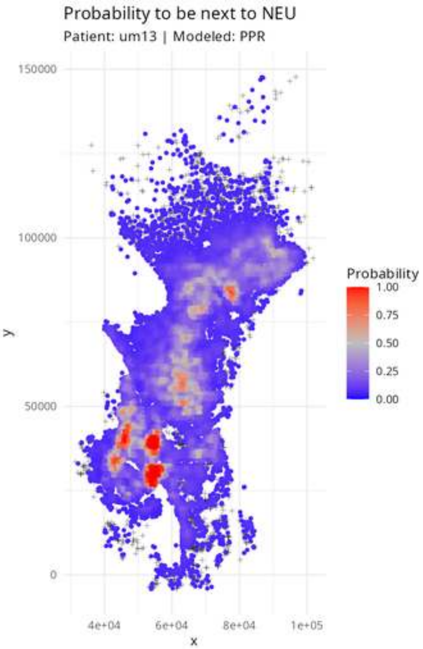

Probability to be next to NEU
Patient: um13 | Modeled: PPR
150000
100000
50000
0
y
4e+04
6e+04
8e+04
1e+05
x
Probability
1.00
0.75
0.50
0.25
0.00

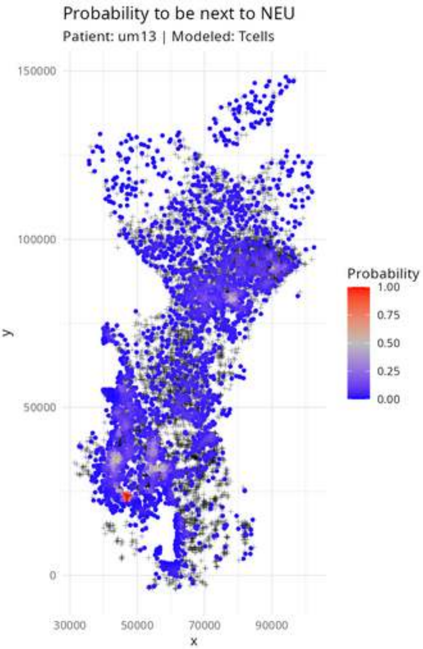
Probability to be next to NEU
Patient: um13 | Modeled: Tcells
150000
100000
50000
0
y
30000
50000
70000
90000
x
Probability
1.00
0.75
0.50
0.25
0.00

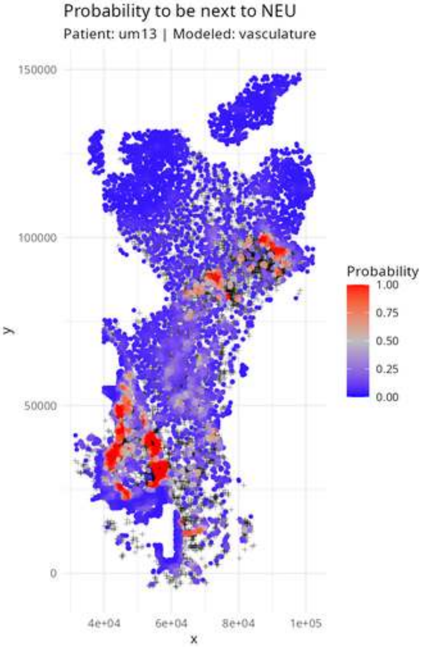
Probability to be next to NEU
Patient: um13 | Modeled: vasculature
150000
100000
50000
0
y
4e+04
6e+04
8e+04
1e+05
x
Probability
1.00
0.75
0.50
0.25
0.00

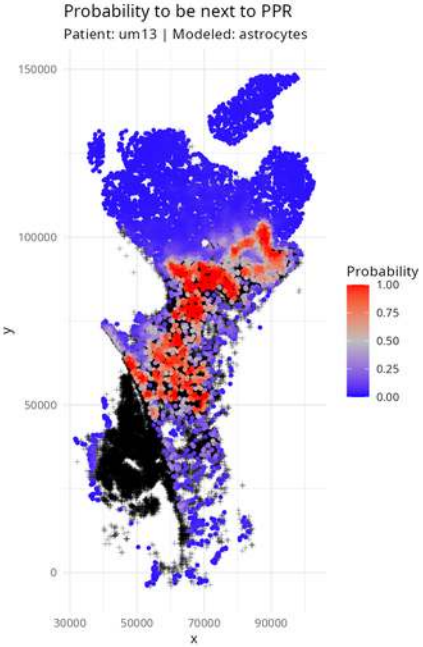

Probability to be next to PPR
Patient: um13 | Modeled: astrocytes
150000
100000
50000
0
y
30000
50000
70000
90000
x
Probability
1.00
0.75
0.50
0.25
0.00

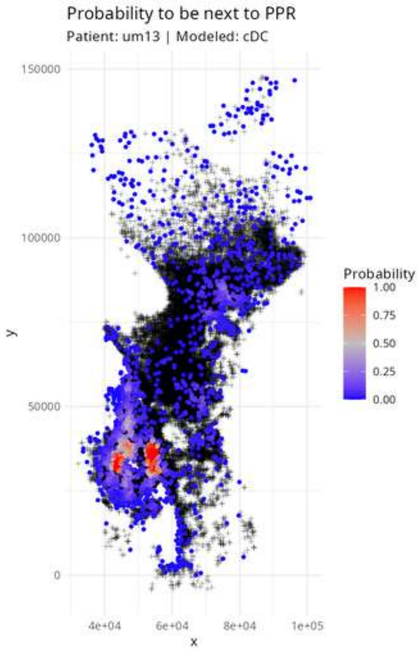

Probability to be next to PPR
Patient: um13 | Modeled: cDC
150000
100000
50000
0
y
4e+04
6e+04
8e+04
1e+05
x
Probability
1.00
0.75
0.50
0.25
0.00

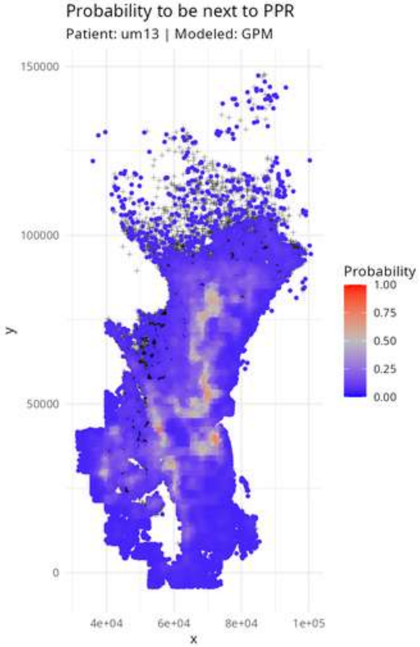

Probability to be next to PPR
Patient: um13 | Modeled: GPM
150000
100000
50000
0
y
4e+04
6e+04
8e+04
1e+05
x
Probability
1.00
0.75
0.50
0.25
0.00

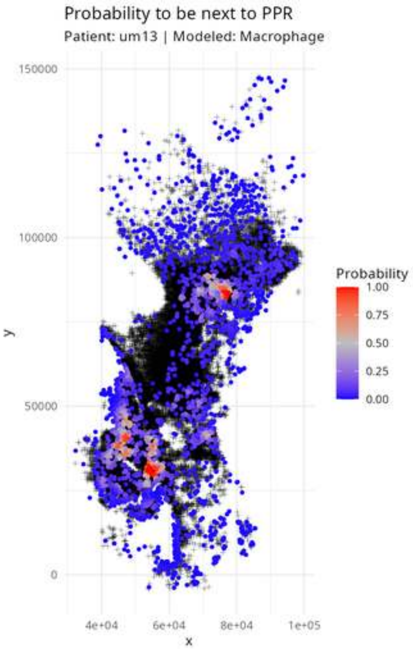

Probability to be next to PPR
Patient: um13 | Modeled: Macrophage
150000
100000
50000
0
y
4e+04
6e+04
8e+04
1e+05
x
Probability
1.00
0.75
0.50
0.25
0.00

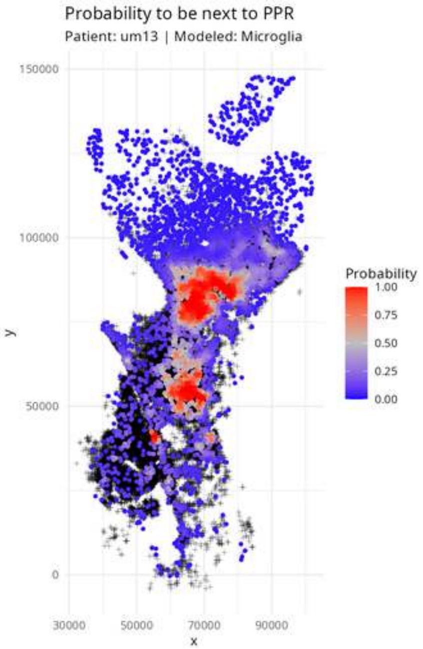

Probability to be next to PPR
Patient: um13 | Modeled: Microglia
150000
100000
50000
0
y
30000
50000
70000
90000
x
Probability
1.00
0.75
0.50
0.25
0.00

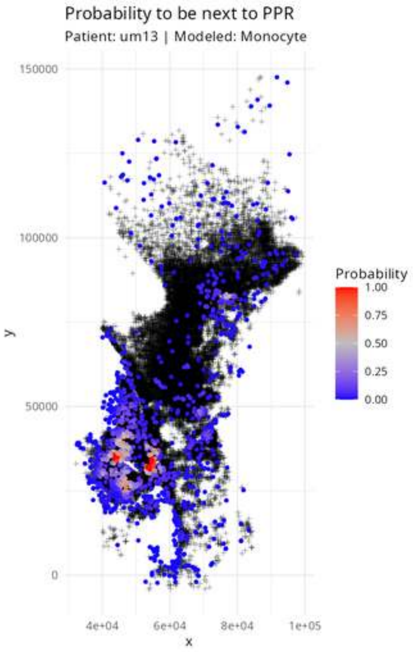

Probability to be next to PPR
Patient: um13 | Modeled: Monocyte
150000
100000
50000
0
y
4e+04
6e+04
8e+04
1e+05
x
Probability
1.00
0.75
0.50
0.25
0.00

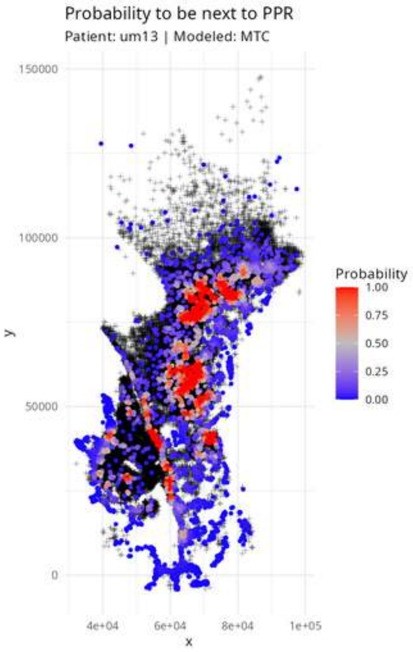
Probability to be next to PPR
Patient: um13 | Modeled: MTC
150000
100000
50000
0
4e+04
6e+04
8e+04
1e+05
x
y
Probability
1.00
0.75
0.50
0.25
0.00

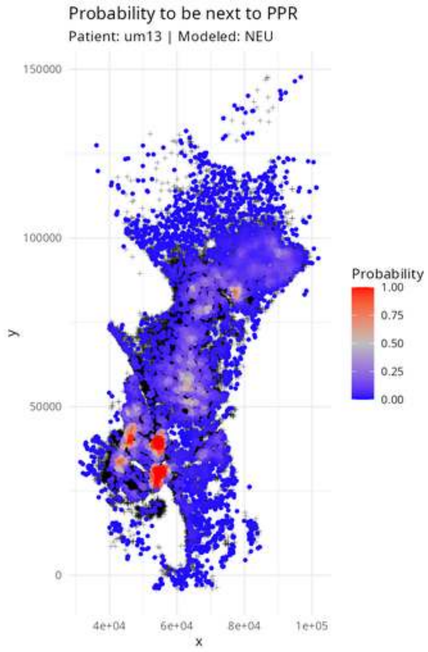

Probability to be next to PPR
Patient: um13 | Modeled: NEU
150000
100000
50000
0
y
4e+04
6e+04
8e+04
1e+05
x
Probability
1.00
0.75
0.50
0.25
0.00

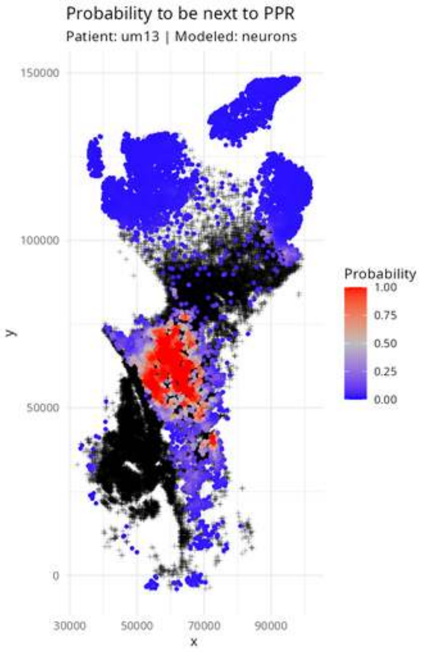

Probability to be next to PPR
Patient: um13 | Modeled: neurons
150000
100000
50000
0
y
30000
50000
70000
90000
x
Probability
1.00
0.75
0.50
0.25
0.00

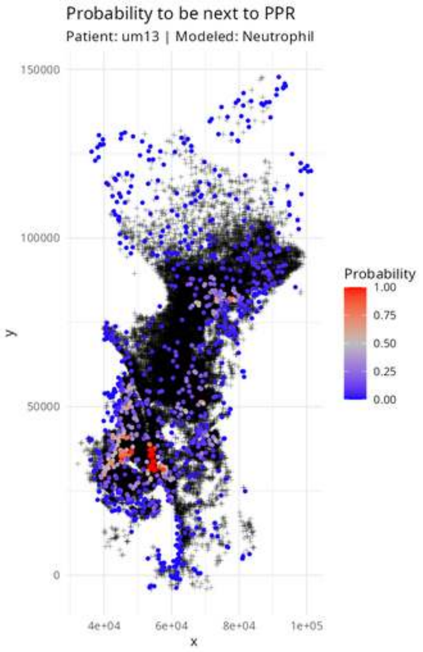

Probability to be next to PPR
Patient: um13 | Modeled: Neutrophil
150000
100000
50000
0
y
4e+04
6e+04
8e+04
1e+05
x
Probability
1.00
0.75
0.50
0.25
0.00

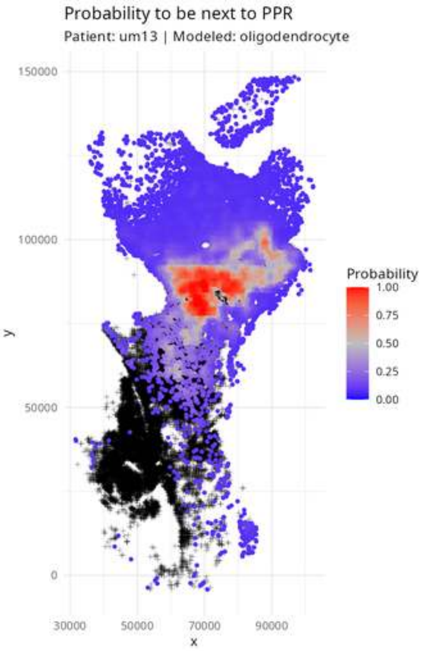

Probability to be next to PPR
Patient: um13 | Modeled: oligodendrocyte
150000
100000
50000
0
y
30000
50000
70000
90000
x
Probability
1.00
0.75
0.50
0.25
0.00

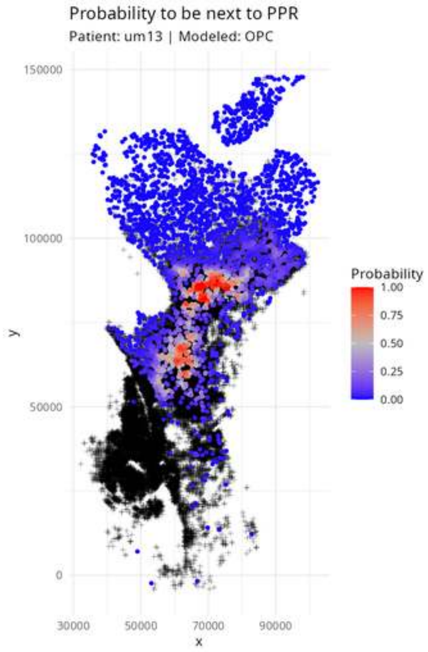

Probability to be next to PPR
Patient: um13 | Modeled: OPC
150000
100000
50000
0
30000
50000
70000
90000
x
y
Probability
1.00
0.75
0.50
0.25
0.00

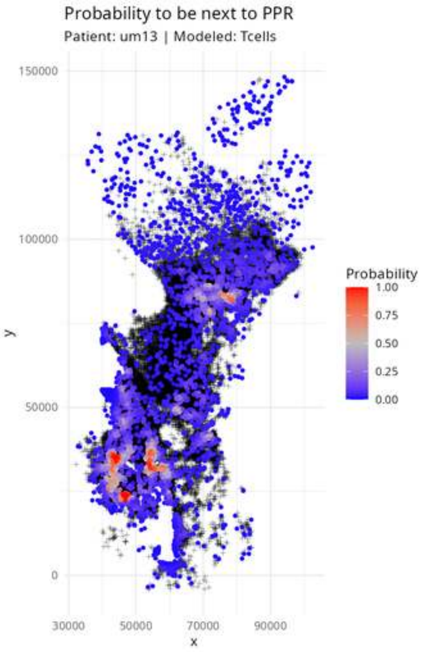

Probability to be next to PPR
Patient: um13 | Modeled: Tcells
150000
100000
50000
0
y
30000
50000
70000
90000
x
Probability
1.00
0.75
0.50
0.25
0.00

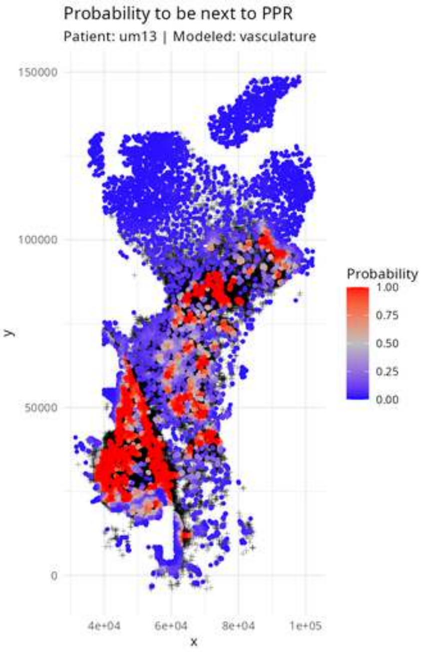
Probability to be next to PPR
Patient: um13 | Modeled: vasculature
150000
100000
50000
0
y
4e+04
6e+04
8e+04
1e+05
x
Probability
1.00
0.75
0.50
0.25
0.00

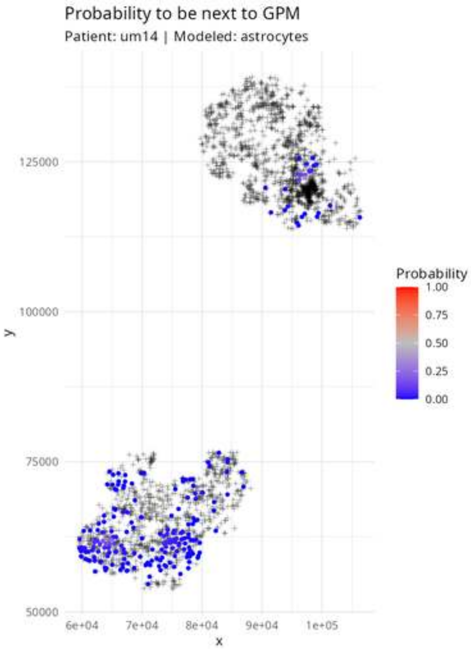

Probability to be next to GPM
Patient: um14 | Modeled: astrocytes
125000
100000
75000
50000
y
6e+04
7e+04
8e+04
9e+04
1e+05
x
Probability
1.00
0.75
0.50
0.25
0.00

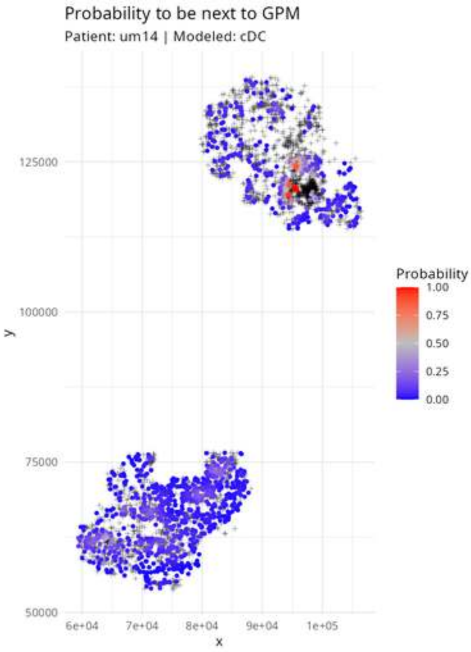

Probability to be next to GPM
Patient: um14 | Modeled: cDC
125000
100000
75000
50000
y
6e+04
7e+04
8e+04
9e+04
1e+05
x
Probability
1.00
0.75
0.50
0.25
0.00

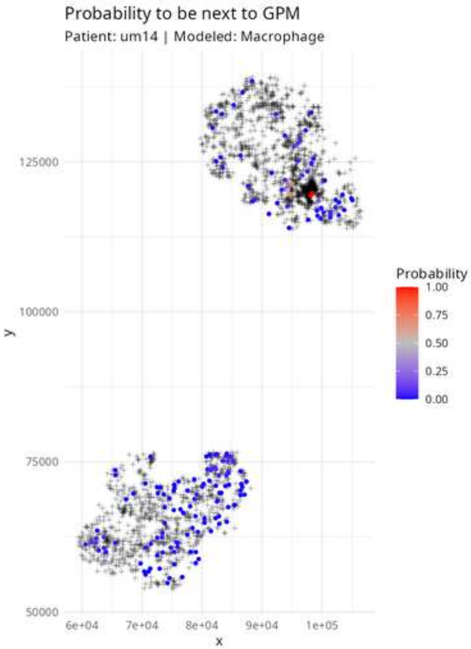

Probability to be next to GPM
Patient: um14 | Modeled: Macrophage
125000
100000
75000
50000
y
6e+04
7e+04
8e+04
9e+04
1e+05
x
Probability
1.00
0.75
0.50
0.25
0.00

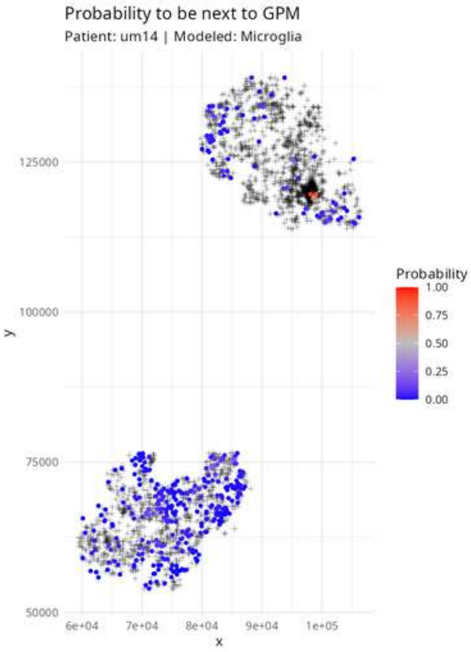

Probability to be next to GPM
Patient: um14 | Modeled: Microglia
Probability
1.00
0.75
0.50
0.25
0.00
125000
100000
75000
50000
y
6e+04
7e+04
8e+04
9e+04
1e+05
x

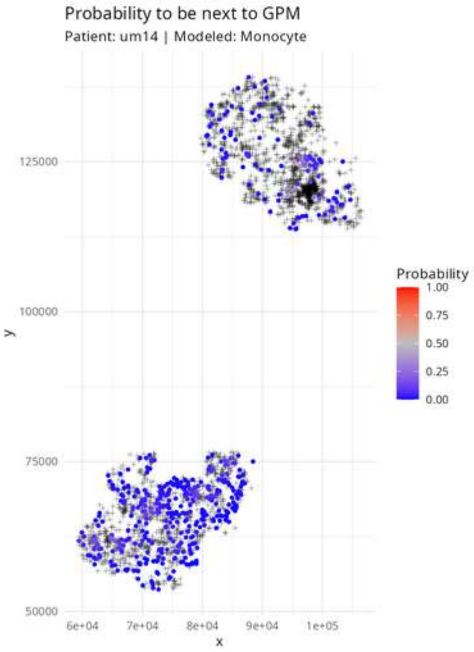

Probability to be next to GPM
Patient: um14 | Modeled: Monocyte
125000
100000
75000
50000
y
6e+04
7e+04
8e+04
9e+04
1e+05
x
Probability
1.00
0.75
0.50
0.25
0.00

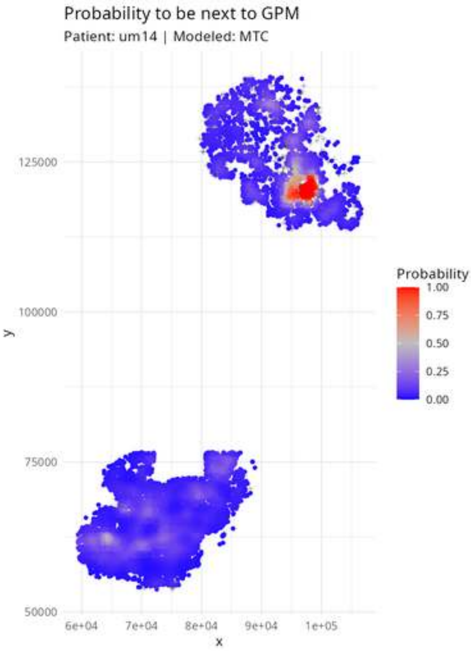
Probability to be next to GPM
Patient: um14 | Modeled: MTC
125000
100000
75000
50000
y
6e+04
7e+04
8e+04
9e+04
1e+05
x
Probability
1.00
0.75
0.50
0.25
0.00

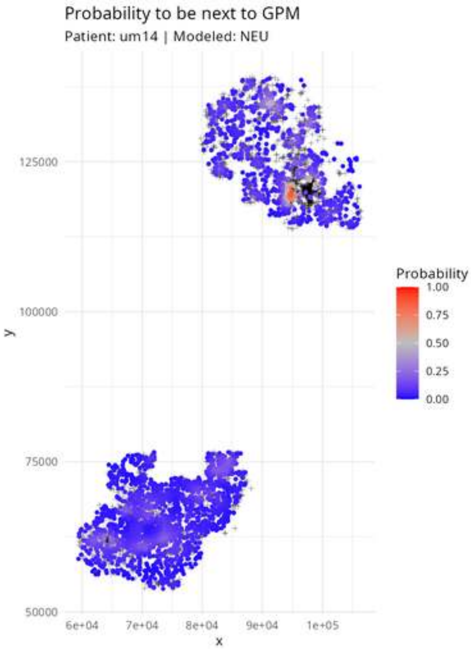

Probability to be next to GPM
Patient: um14 | Modeled: NEU
125000
100000
75000
50000
y
6e+04
7e+04
8e+04
9e+04
1e+05
x
Probability
1.00
0.75
0.50
0.25
0.00

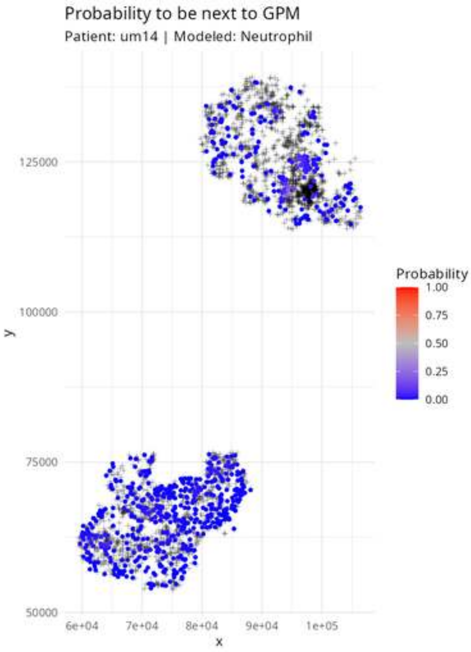

Probability to be next to GPM
Patient: um14 | Modeled: Neutrophil
125000
100000
75000
50000
y
6e+04
7e+04
8e+04
9e+04
1e+05
x
Probability
1.00
0.75
0.50
0.25
0.00

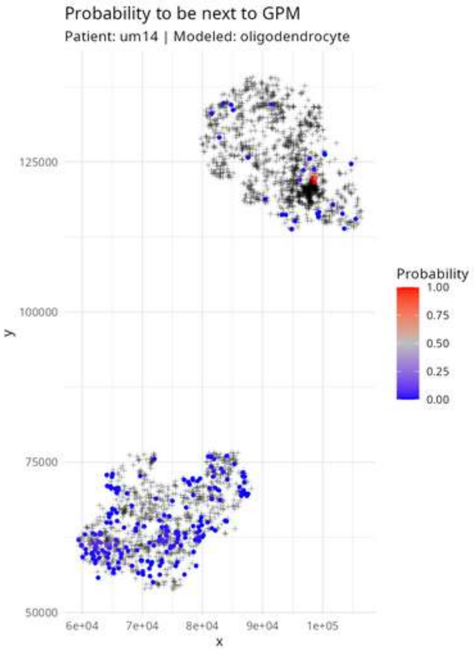

Probability to be next to GPM
Patient: um14 | Modeled: oligodendrocyte
125000
100000
75000
50000
y
6e+04
7e+04
8e+04
9e+04
1e+05
x
Probability
1.00
0.75
0.50
0.25
0.00

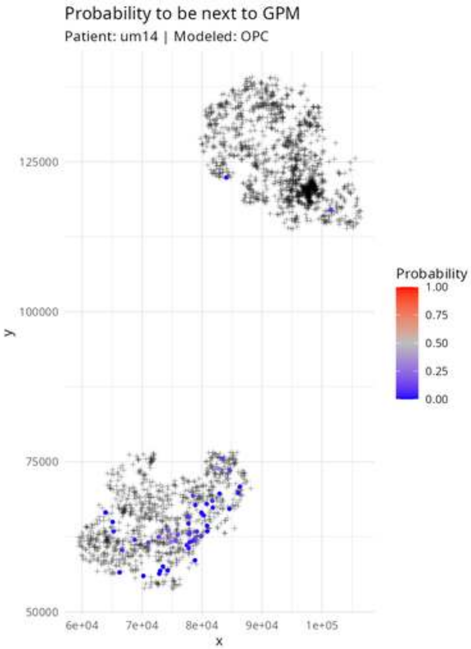
Probability to be next to GPM
Patient: um14 | Modeled: OPC
125000
100000
75000
50000
y
6e+04
7e+04
8e+04
9e+04
1e+05
x
Probability
1.00
0.75
0.50
0.25
0.00

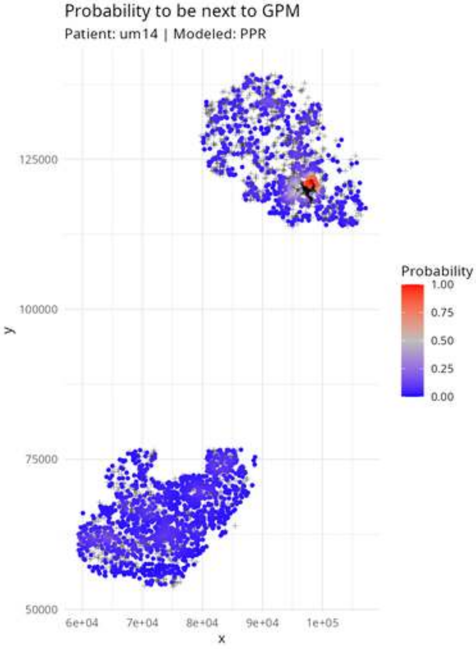
Probability to be next to GPM
Patient: um14 | Modeled: PPR
125000
100000
75000
50000
y
6e+04
7e+04
8e+04
9e+04
1e+05
x
Probability
1.00
0.75
0.50
0.25
0.00

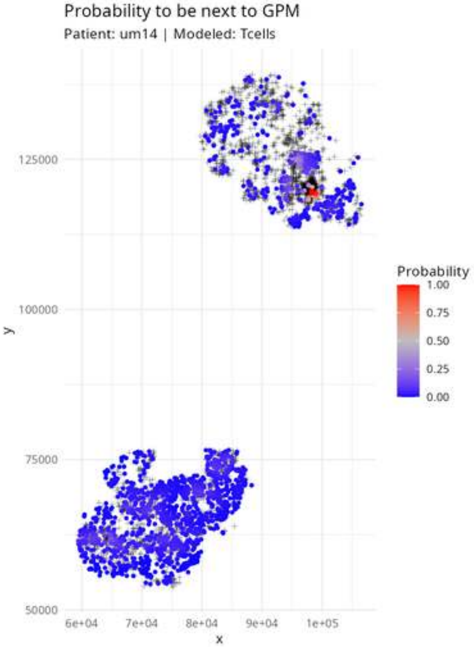

Probability to be next to GPM
Patient: um14 | Modeled: Tcells
125000
100000
75000
50000
y
6e+04
7e+04
8e+04
9e+04
1e+05
x
Probability
1.00
0.75
0.50
0.25
0.00

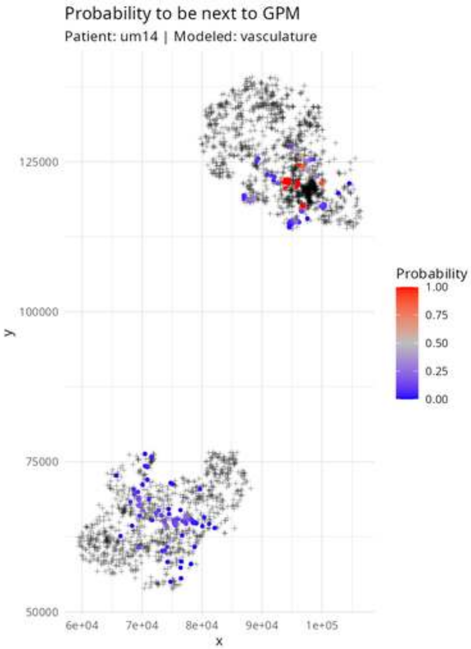
Probability to be next to GPM
Patient: um14 | Modeled: vasculature
125000
100000
75000
50000
y
6e+04
7e+04
8e+04
9e+04
1e+05
x
Probability
1.00
0.75
0.50
0.25
0.00

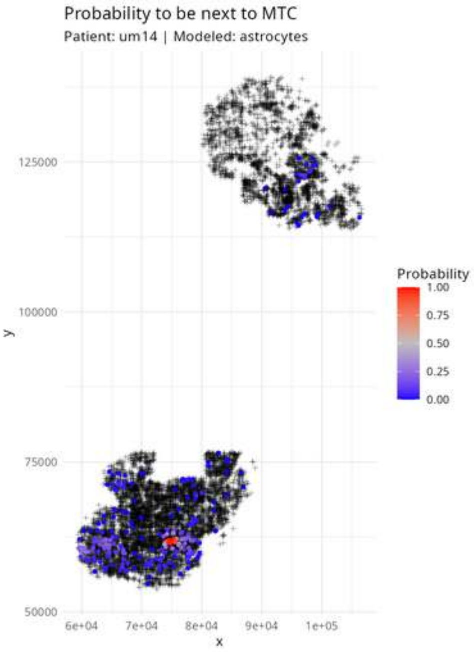

Probability to be next to MTC
Patient: um14 | Modeled: astrocytes
125000
100000
75000
50000
y
6e+04
7e+04
8e+04
9e+04
1e+05
x
Probability
1.00
0.75
0.50
0.25
0.00

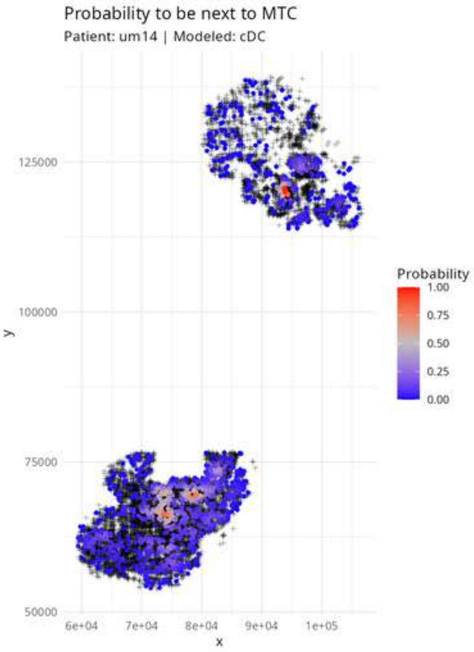

Probability to be next to MTC
Patient: um14 | Modeled: cDC
125000
100000
75000
50000
y
6e+04
7e+04
8e+04
9e+04
1e+05
x
Probability
1.00
0.75
0.50
0.25
0.00

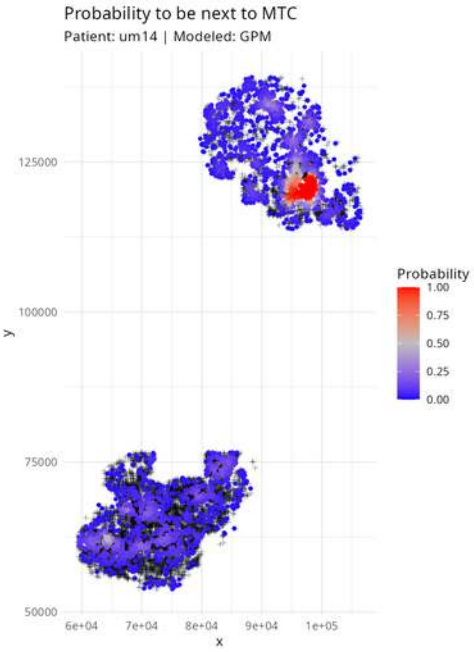
Probability to be next to MTC
Patient: um14 | Modeled: GPM
125000
100000
75000
50000
y
6e+04
7e+04
8e+04
9e+04
1e+05
x
Probability
1.00
0.75
0.50
0.25
0.00

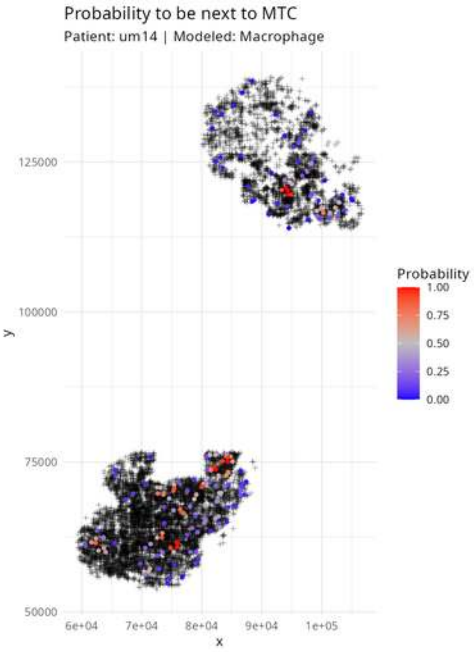
Probability to be next to MTC
Patient: um14 | Modeled: Macrophage
Probability
1.00
0.75
0.50
0.25
0.00
125000
100000
75000
50000
y
6e+04
7e+04
8e+04
9e+04
1e+05
x

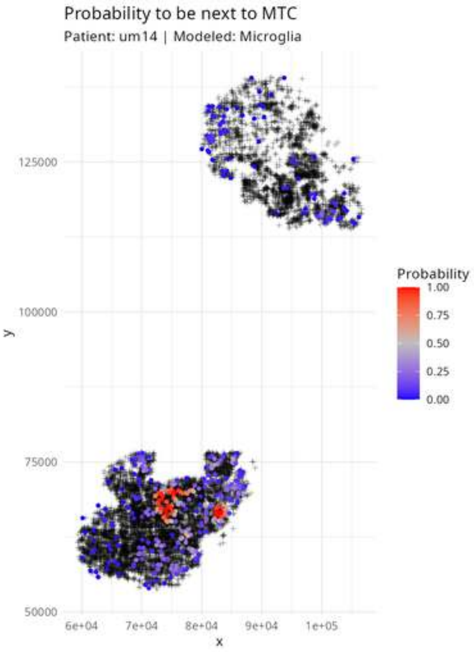

Probability to be next to MTC
Patient: um14 | Modeled: Microglia
125000
100000
75000
50000
y
6e+04
7e+04
8e+04
9e+04
1e+05
x
Probability
1.00
0.75
0.50
0.25
0.00

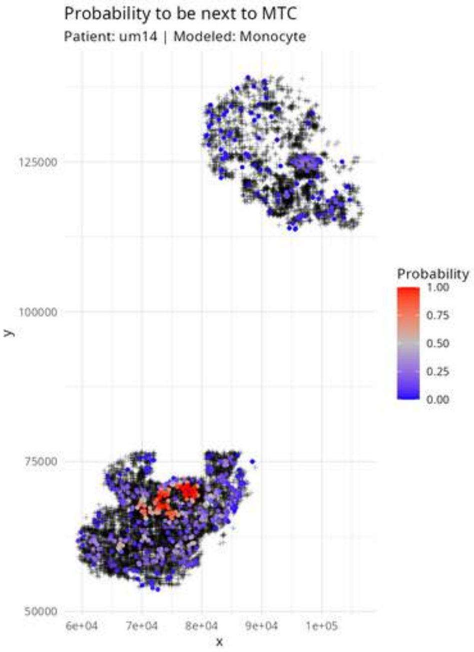
Probability to be next to MTC
Patient: um14 | Modeled: Monocyte
125000
100000
75000
50000
y
6e+04
7e+04
8e+04
9e+04
1e+05
x
Probability
1.00
0.75
0.50
0.25
0.00

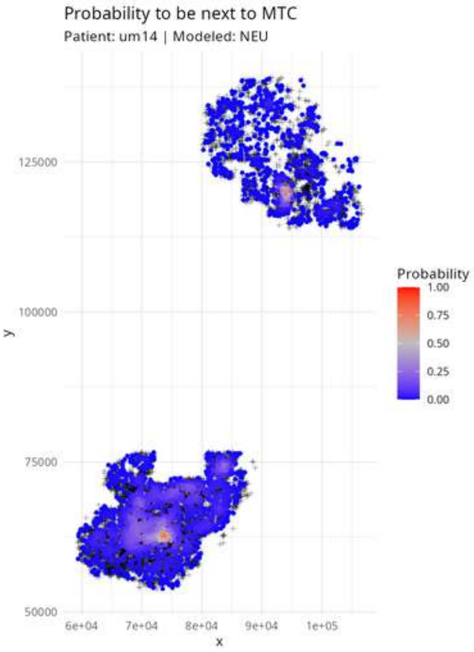

Probability to be next to MTC
Patient: um14 | Modeled: NEU
125000
100000
75000
50000
y
6e+04
7e+04
8e+04
9e+04
1e+05
x
Probability
1.00
0.75
0.50
0.25
0.00

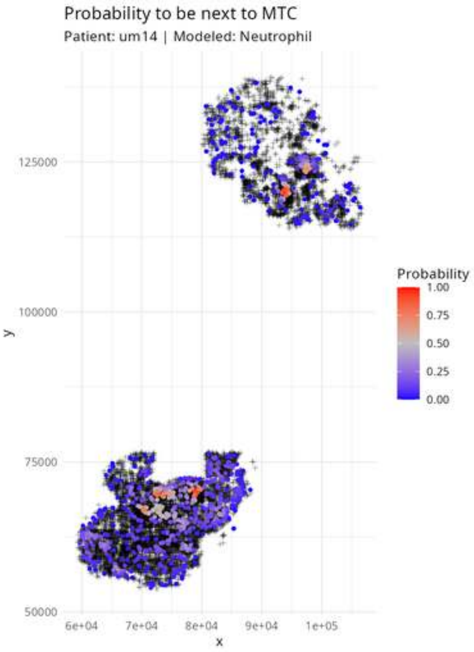
Probability to be next to MTC
Patient: um14 | Modeled: Neutrophil
125000
100000
75000
50000
y
6e+04
7e+04
8e+04
9e+04
1e+05
x
Probability
1.00
0.75
0.50
0.25
0.00

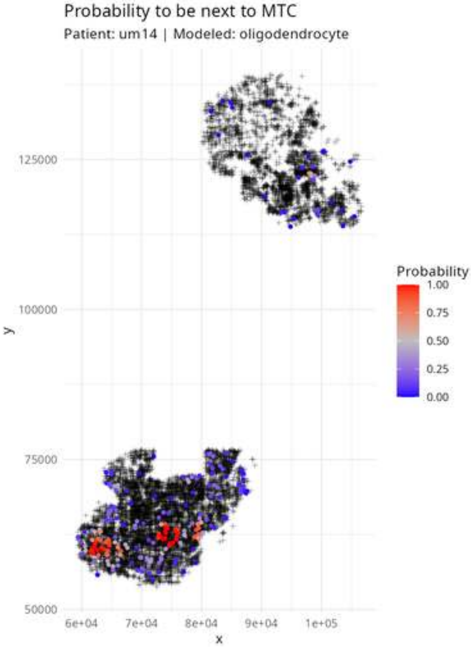

Probability to be next to MTC
Patient: um14 | Modeled: oligodendrocyte
Probability
1.00
0.75
0.50
0.25
0.00
125000
100000
75000
50000
y
6e+04
7e+04
8e+04
9e+04
1e+05
x

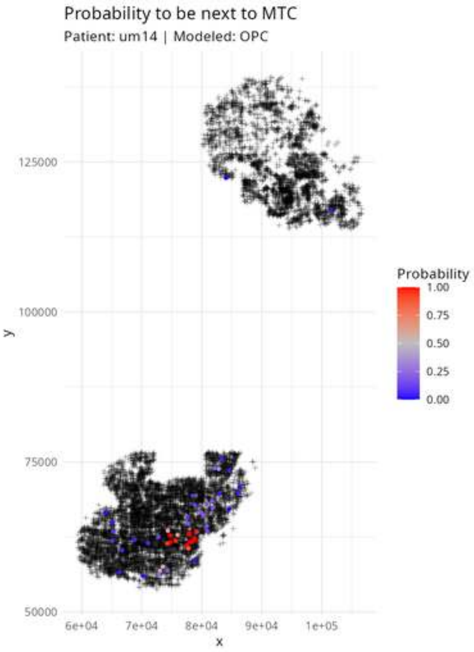
Probability to be next to MTC
Patient: um14 | Modeled: OPC
125000
100000
75000
50000
y
6e+04
7e+04
8e+04
9e+04
1e+05
x
Probability
1.00
0.75
0.50
0.25
0.00

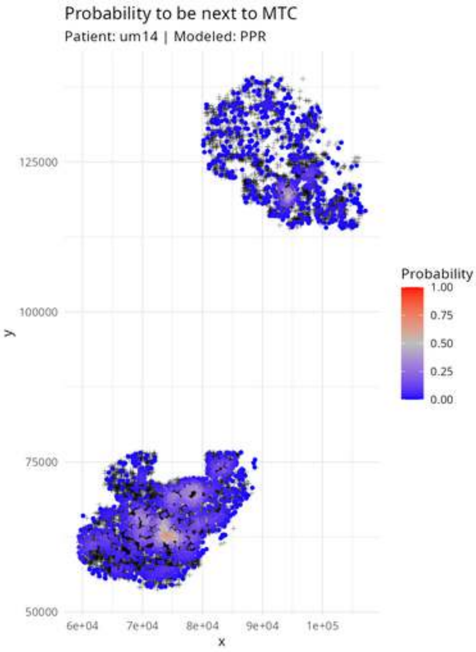
Probability to be next to MTC
Patient: um14 | Modeled: PPR
125000
100000
75000
50000
y
6e+04
7e+04
8e+04
9e+04
1e+05
x
Probability
1.00
0.75
0.50
0.25
0.00

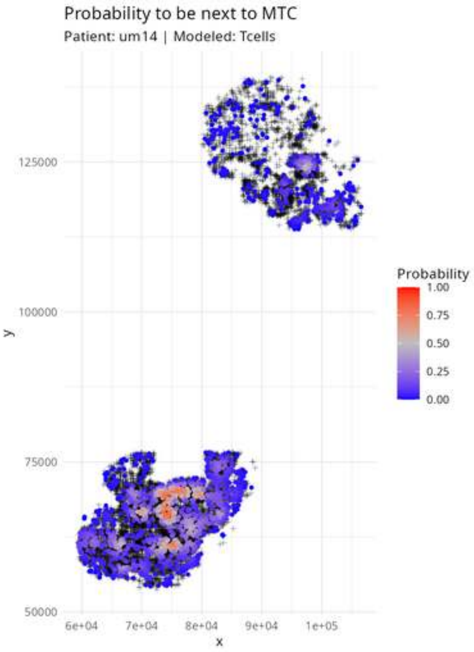
Probability to be next to MTC
Patient: um14 | Modeled: Tcells
125000
100000
75000
50000
y
6e+04
7e+04
8e+04
9e+04
1e+05
x
Probability
1.00
0.75
0.50
0.25
0.00

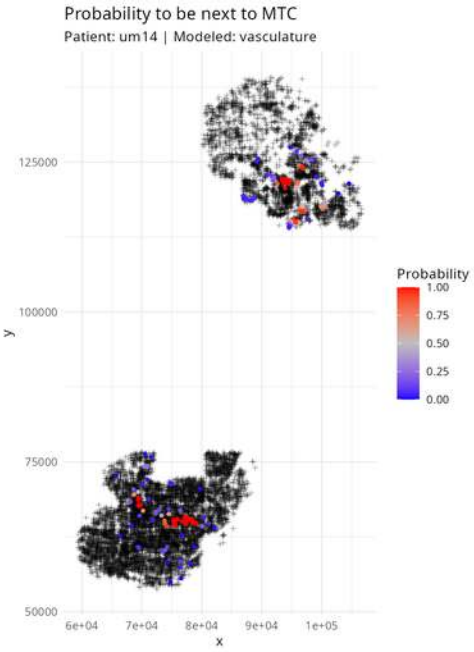
Probability to be next to MTC
Patient: um14 | Modeled: vasculature
y
125000
100000
75000
50000
x
6e+04
7e+04
8e+04
9e+04
1e+05
Probability
1.00
0.75
0.50
0.25
0.00

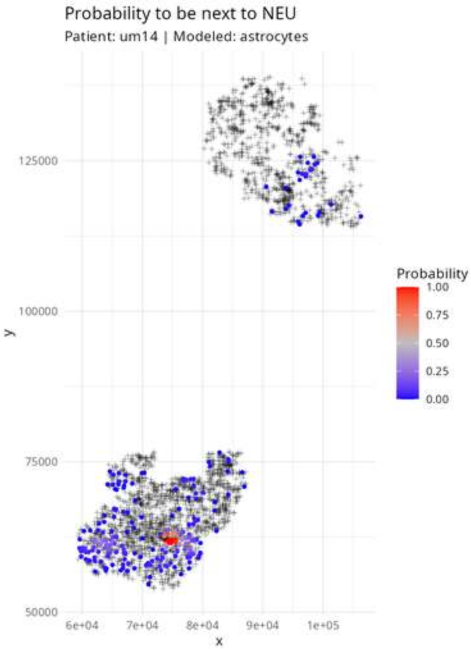

Probability to be next to NEU
Patient: um14 | Modeled: astrocytes
y
125000
100000
75000
50000
x
6e+04
7e+04
8e+04
9e+04
1e+05
Probability
1.00
0.75
0.50
0.25
0.00

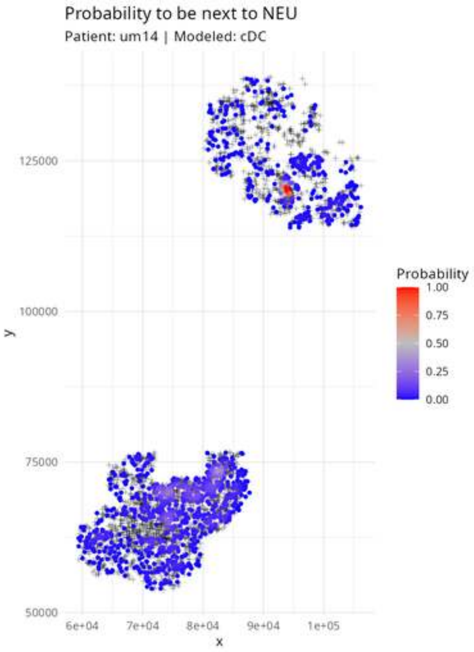

Probability to be next to NEU
Patient: um14 | Modeled: cDC
Probability
1.00
0.75
0.50
0.25
0.00
y
125000
100000
75000
50000
x
6e+04
7e+04
8e+04
9e+04
1e+05

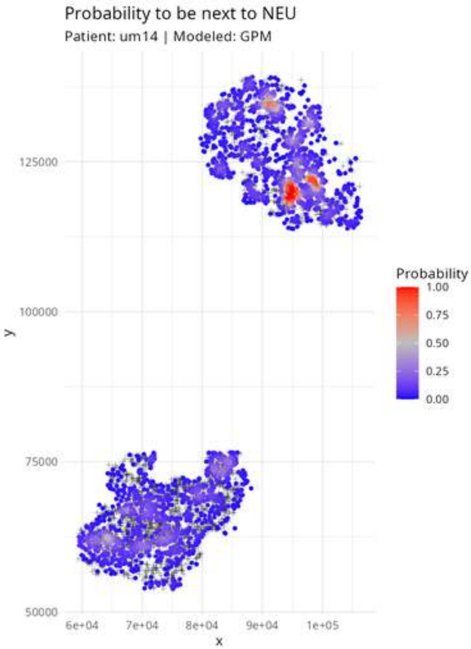

Probability to be next to NEU
Patient: um14 | Modeled: GPM
125000
100000
75000
50000
y
6e+04
7e+04
8e+04
9e+04
1e+05
x
Probability
1.00
0.75
0.50
0.25
0.00

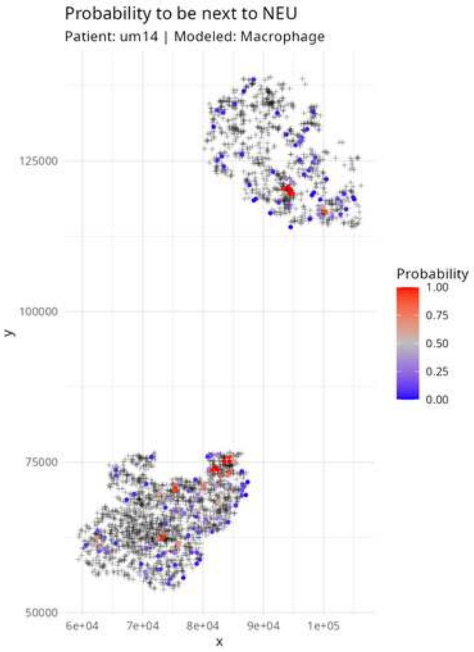

Probability to be next to NEU
Patient: um14 | Modeled: Macrophage
Probability
1.00
0.75
0.50
0.25
0.00
125000
100000
75000
50000
y
6e+04
7e+04
8e+04
9e+04
1e+05
x

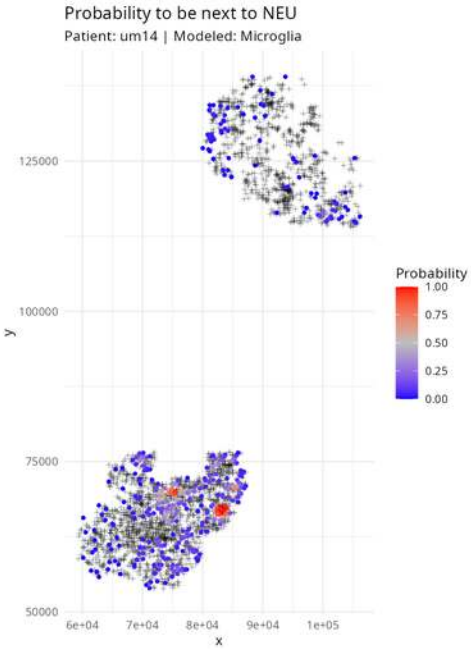

Probability to be next to NEU
Patient: um14 | Modeled: Microglia
125000
100000
75000
50000
y
6e+04
7e+04
8e+04
9e+04
1e+05
x
Probability
1.00
0.75
0.50
0.25
0.00

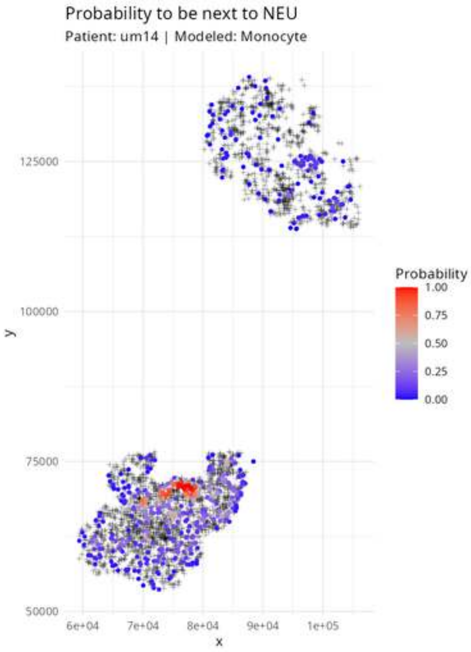
Probability to be next to NEU
Patient: um14 | Modeled: Monocyte
y
125000
100000
75000
50000
x
6e+04
7e+04
8e+04
9e+04
1e+05
Probability
1.00
0.75
0.50
0.25
0.00

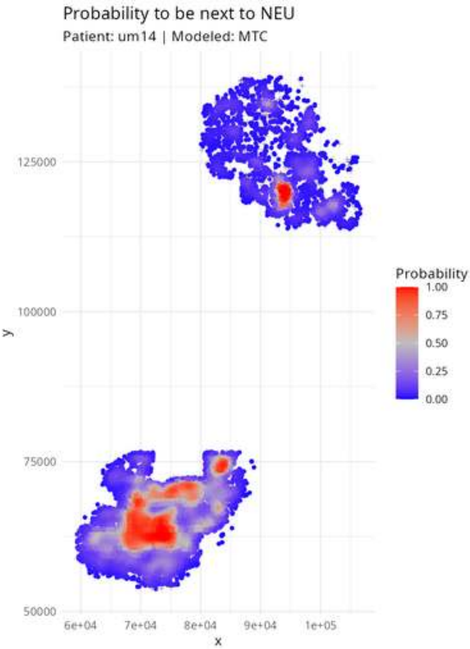

Probability to be next to NEU
Patient: um14 | Modeled: MTC
125000
100000
75000
50000
y
6e+04
7e+04
8e+04
9e+04
1e+05
x
Probability
1.00
0.75
0.50
0.25
0.00

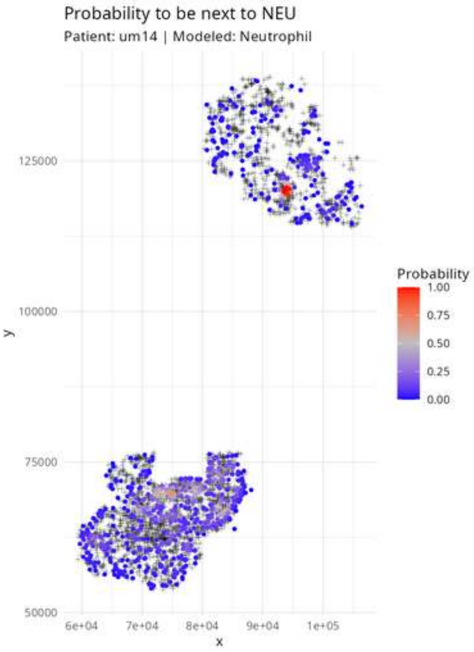

Probability to be next to NEU
Patient: um14 | Modeled: Neutrophil
125000
100000
75000
50000
y
6e+04
7e+04
8e+04
9e+04
1e+05
x
Probability
1.00
0.75
0.50
0.25
0.00

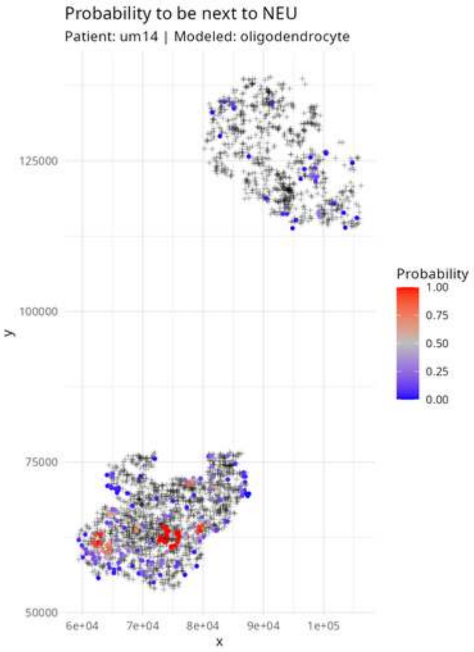

Probability to be next to NEU
Patient: um14 | Modeled: oligodendrocyte
Probability
1.00
0.75
0.50
0.25
0.00
125000
100000
75000
50000
y
6e+04
7e+04
8e+04
9e+04
1e+05
x

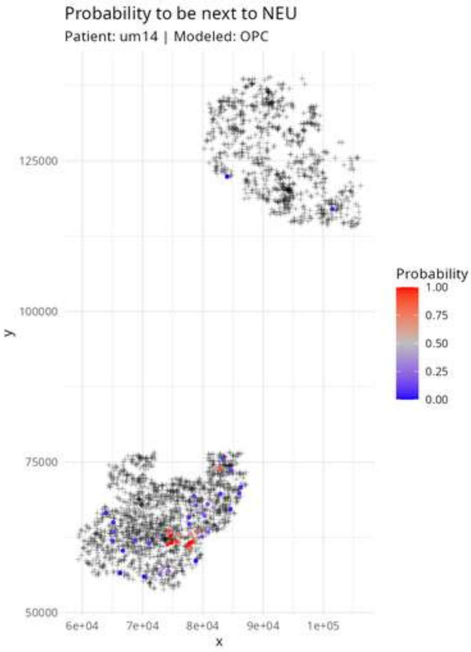
Probability to be next to NEU
Patient: um14 | Modeled: OPC
y
125000
100000
75000
50000
x
6e+04
7e+04
8e+04
9e+04
1e+05
Probability
1.00
0.75
0.50
0.25
0.00

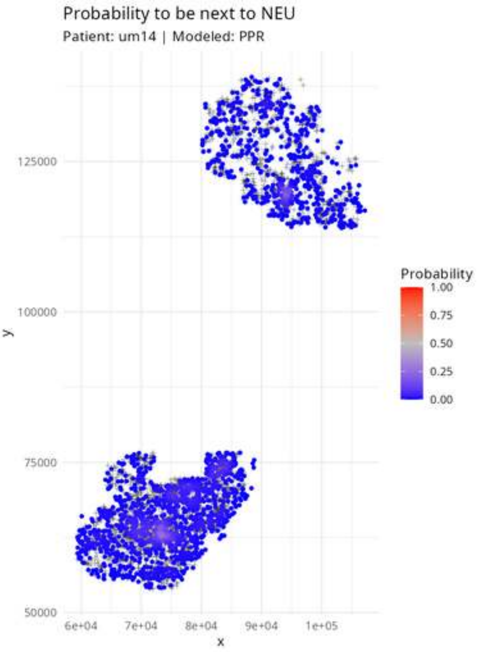
Probability to be next to NEU
Patient: um14 | Modeled: PPR
125000
100000
75000
50000
y
6e+04
7e+04
8e+04
9e+04
1e+05
x
Probability
1.00
0.75
0.50
0.25
0.00

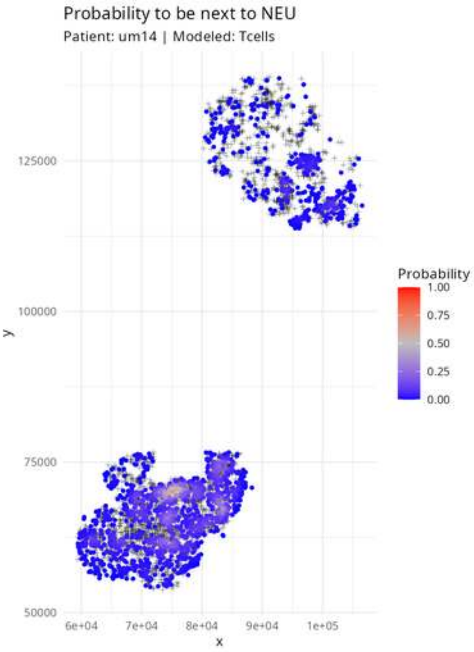

Probability to be next to NEU
Patient: um14 | Modeled: Tcells
125000
100000
75000
50000
y
6e+04
7e+04
8e+04
9e+04
1e+05
x
Probability
1.00
0.75
0.50
0.25
0.00

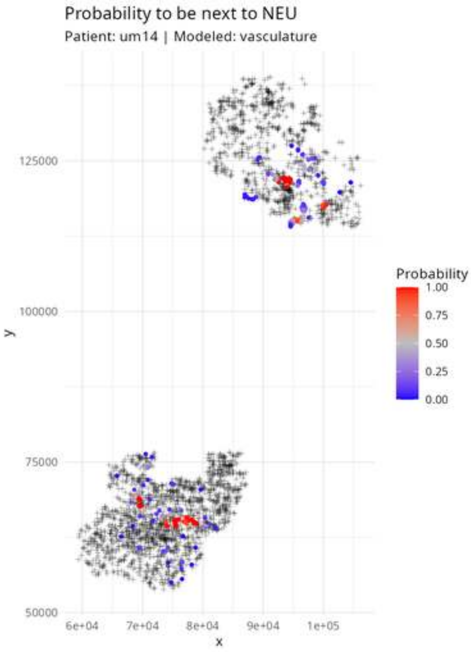
Probability to be next to NEU
Patient: um14 | Modeled: vasculature
125000
100000
75000
50000
y
6e+04
7e+04
8e+04
9e+04
1e+05
x
Probability
1.00
0.75
0.50
0.25
0.00

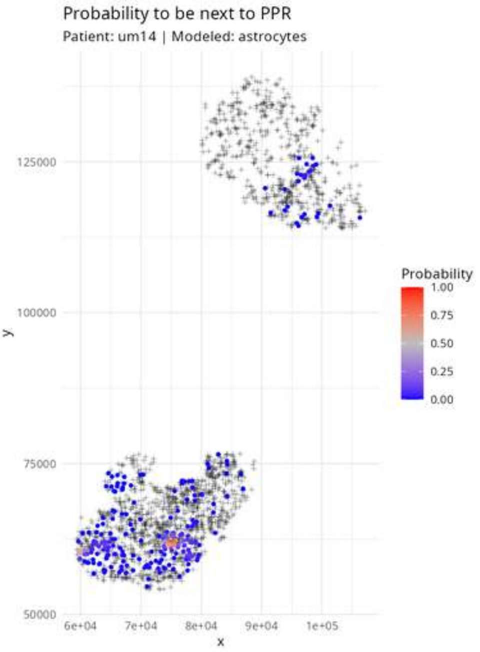

Probability to be next to PPR
Patient: um14 | Modeled: astrocytes
125000
100000
75000
50000
y
6e+04
7e+04
8e+04
9e+04
1e+05
x
Probability
1.00
0.75
0.50
0.25
0.00

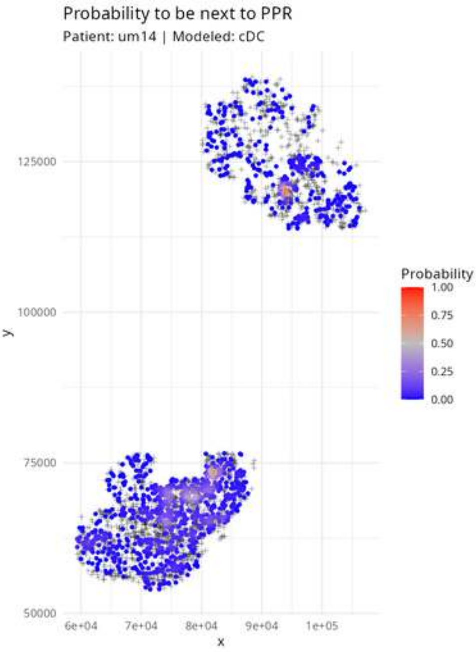
Probability to be next to PPR
Patient: um14 | Modeled: cDC
125000
100000
75000
50000
y
6e+04
7e+04
8e+04
9e+04
1e+05
x
Probability
1.00
0.75
0.50
0.25
0.00

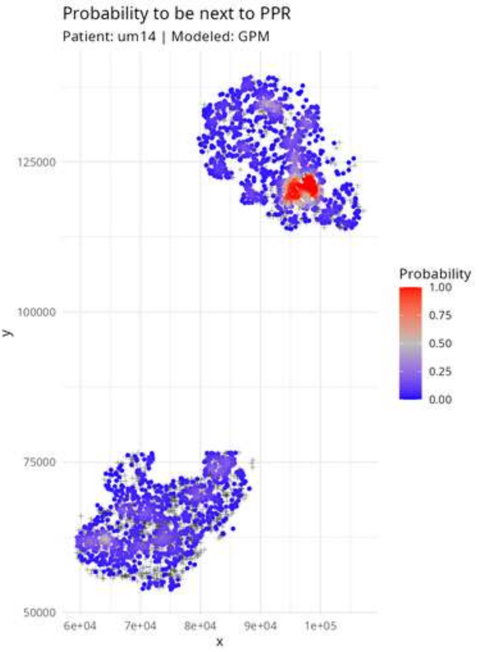
Probability to be next to PPR
Patient: um14 | Modeled: GPM
125000
100000
75000
50000
y
6e+04
7e+04
8e+04
9e+04
1e+05
x
Probability
1.00
0.75
0.50
0.25
0.00

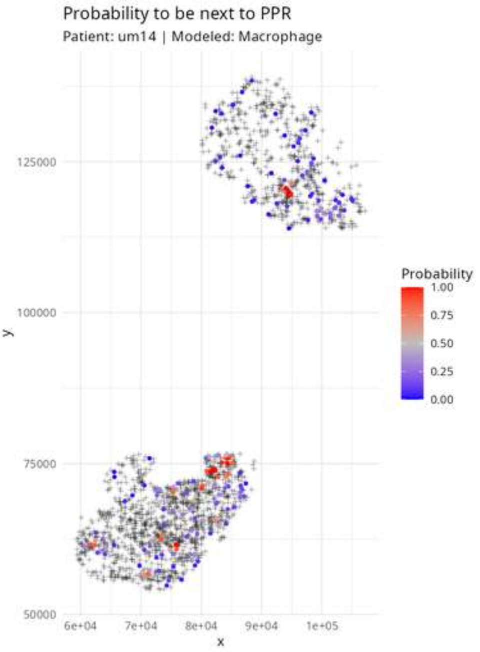

Probability to be next to PPR
Patient: um14 | Modeled: Macrophage
Probability
1.00
0.75
0.50
0.25
0.00
125000
100000
75000
50000
y
6e+04
7e+04
8e+04
9e+04
1e+05
x

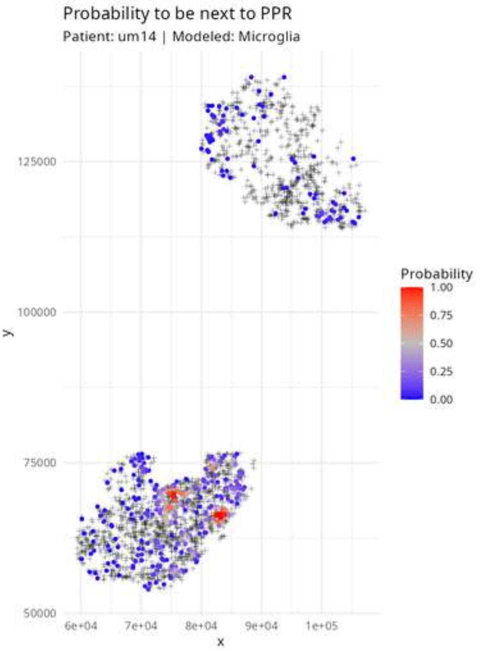

Probability to be next to PPR
Patient: um14 | Modeled: Microglia
Probability
1.00
0.75
0.50
0.25
0.00
125000
100000
75000
50000
y
6e+04
7e+04
8e+04
9e+04
1e+05
x

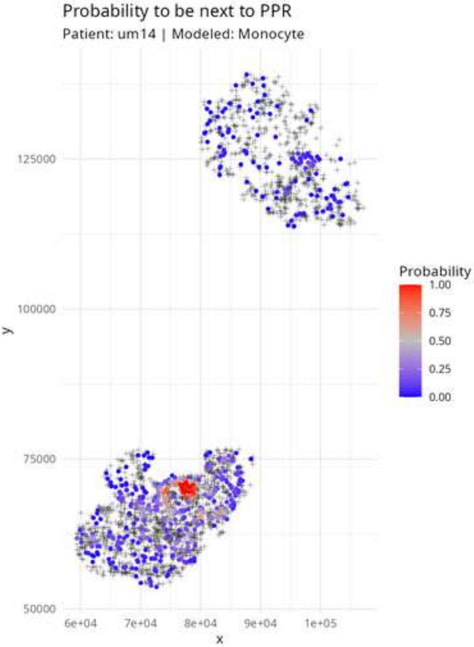

Probability to be next to PPR
Patient: um14 | Modeled: Monocyte
y
125000
100000
75000
50000
6e+04
7e+04
8e+04
9e+04
1e+05
x
Probability
1.00
0.75
0.50
0.25
0.00

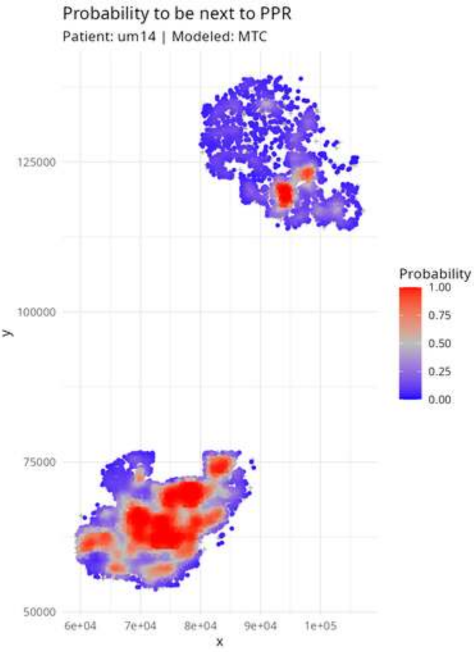

Probability to be next to PPR
Patient: um14 | Modeled: MTC
125000
100000
75000
50000
y
6e+04
7e+04
8e+04
9e+04
1e+05
x
Probability
1.00
0.75
0.50
0.25
0.00

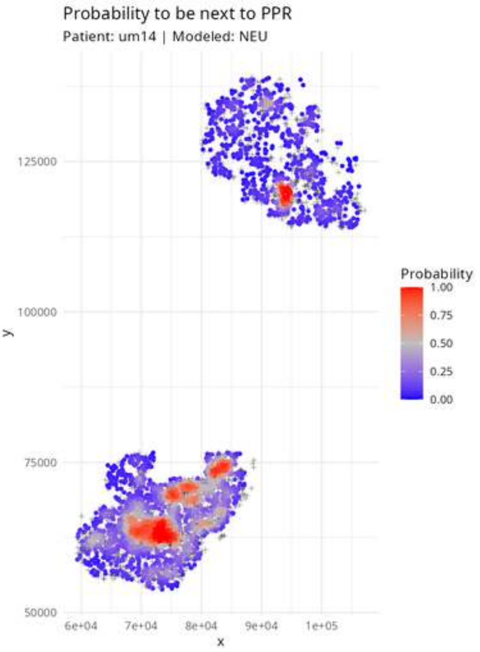

Probability to be next to PPR
Patient: um14 | Modeled: NEU
125000
100000
75000
50000
y
6e+04
7e+04
8e+04
9e+04
1e+05
x
Probability
1.00
0.75
0.50
0.25
0.00

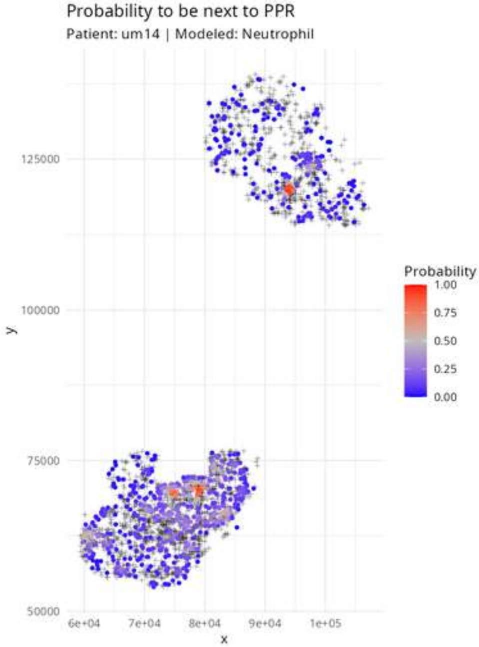
Probability to be next to PPR
Patient: um14 | Modeled: Neutrophil
125000
100000
75000
50000
y
6e+04
7e+04
8e+04
9e+04
1e+05
x
Probability
1.00
0.75
0.50
0.25
0.00

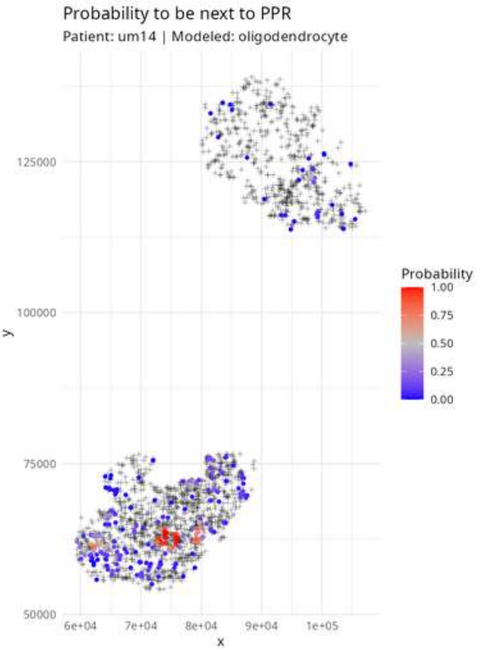

Probability to be next to PPR
Patient: um14 | Modeled: oligodendrocyte
Probability
1.00
0.75
0.50
0.25
0.00
125000
100000
75000
50000
y
6e+04
7e+04
8e+04
9e+04
1e+05
x

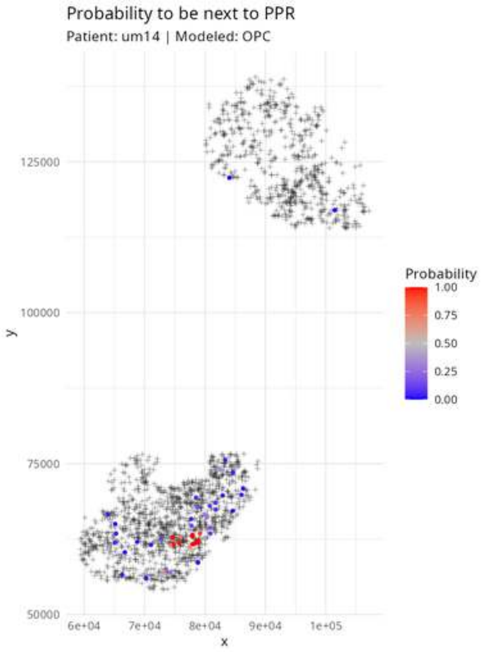

Probability to be next to PPR
Patient: um14 | Modeled: OPC
125000
100000
75000
50000
y
6e+04
7e+04
8e+04
9e+04
1e+05
x
Probability
1.00
0.75
0.50
0.25
0.00

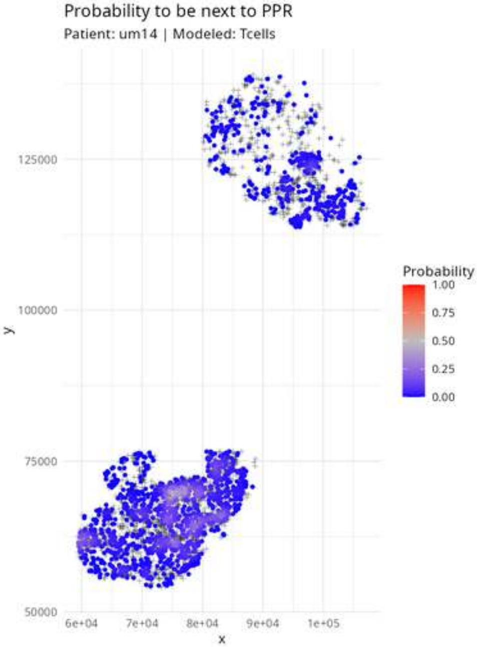
Probability to be next to PPR
Patient: um14 | Modeled: Tcells
125000
100000
75000
50000
y
6e+04
7e+04
8e+04
9e+04
1e+05
x
Probability
1.00
0.75
0.50
0.25
0.00

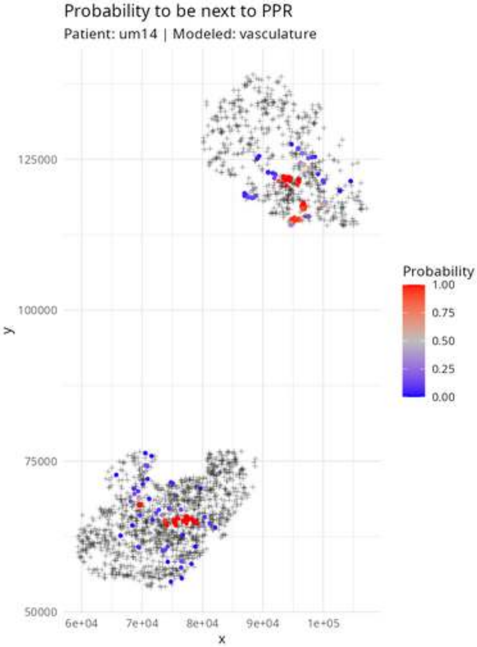
Probability to be next to PPR
Patient: um14 | Modeled: vasculature
125000
100000
75000
50000
y
6e+04
7e+04
8e+04
9e+04
1e+05
x
Probability
1.00
0.75
0.50
0.25
0.00

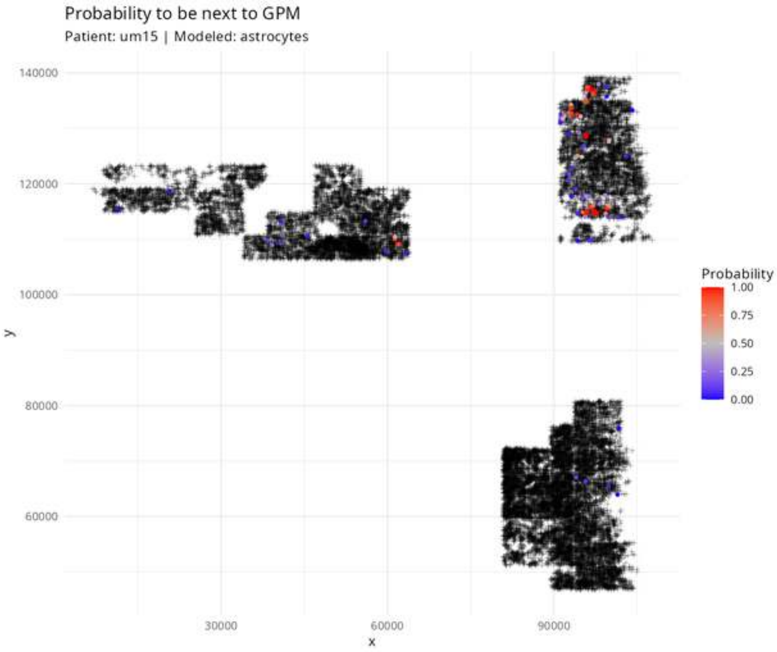

Probability to be next to GPM
Patient: um15 | Modeled: astrocytes
140000
120000
100000
80000
60000
y
30000
60000
90000
x
Probability
1.00
0.75
0.50
0.25
0.00

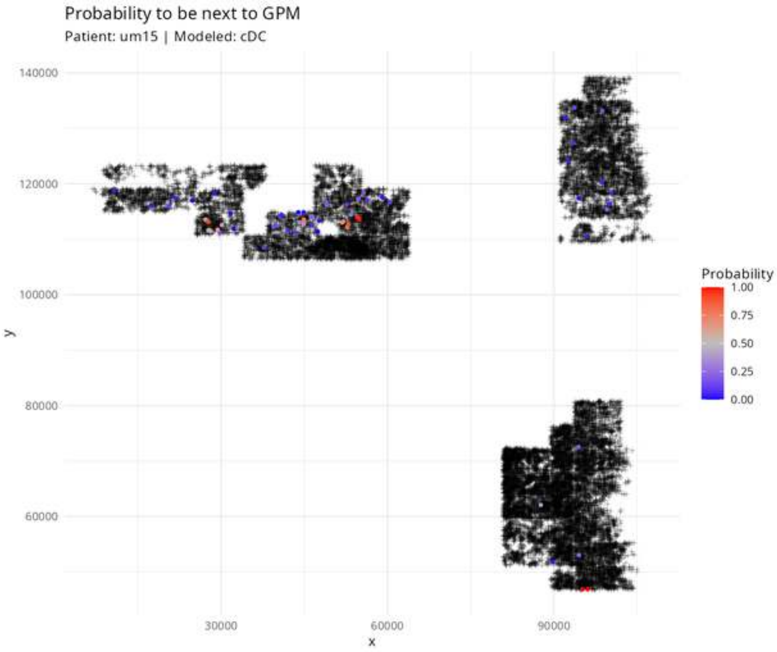

Probability to be next to GPM
Patient: um15 | Modeled: cDC
140000
120000
100000
80000
60000
y
30000
60000
90000
x
Probability
1.00
0.75
0.50
0.25
0.00

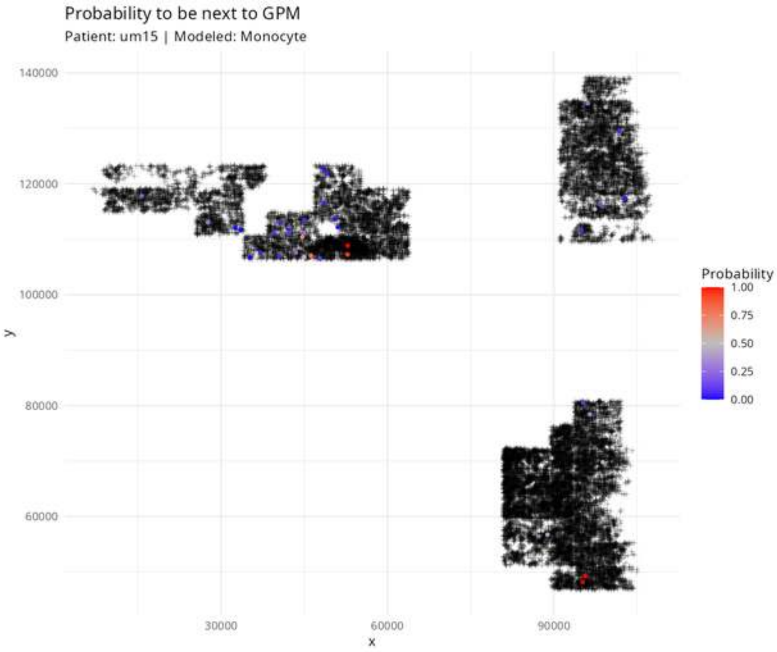
Probability to be next to GPM
Patient: um15 | Modeled: Monocyte
140000
120000
100000
80000
60000
y
30000
60000
90000
x
Probability
1.00
0.75
0.50
0.25
0.00

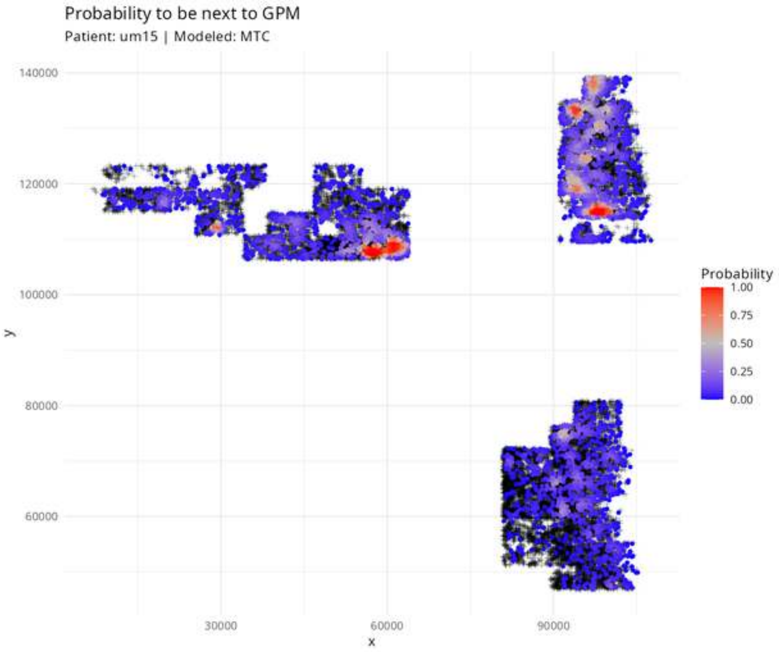
Probability to be next to GPM
Patient: um15 | Modeled: MTC
140000
120000
100000
80000
60000
y
30000
60000
90000
x
Probability
1.00
0.75
0.50
0.25
0.00

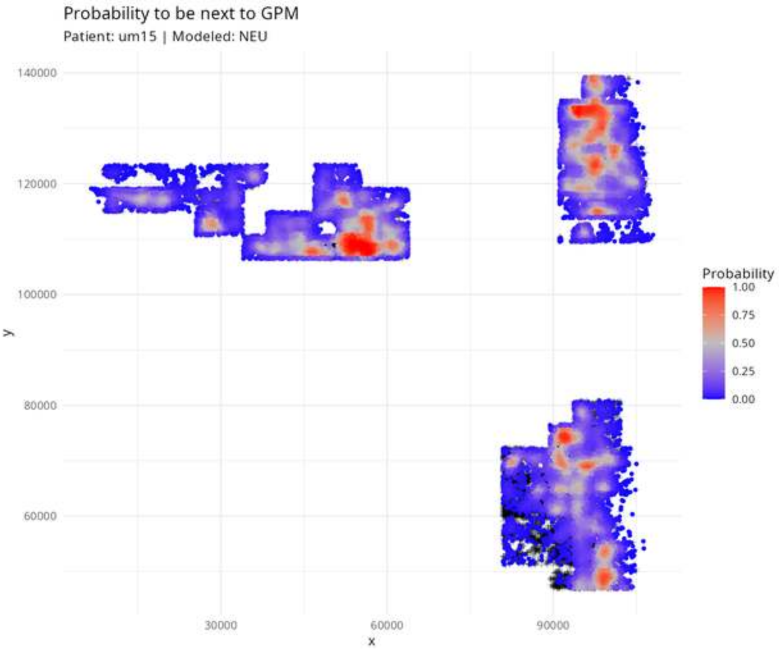

Probability to be next to GPM
Patient: um15 | Modeled: NEU
140000
120000
100000
80000
60000
y
30000
60000
90000
x
Probability
1.00
0.75
0.50
0.25
0.00

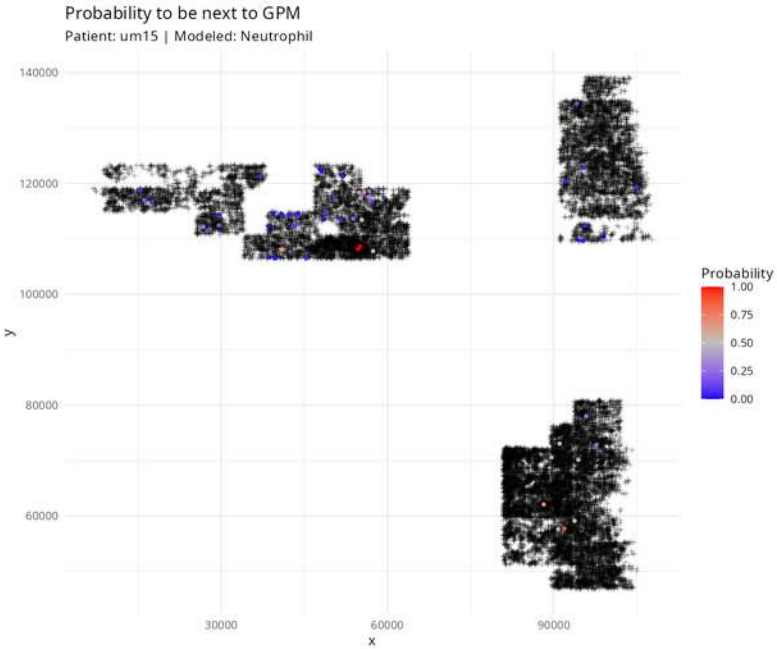
Probability to be next to GPM
Patient: um15 | Modeled: Neutrophil
140000
120000
100000
80000
60000
y
30000
60000
90000
x
Probability
1.00
0.75
0.50
0.25
0.00

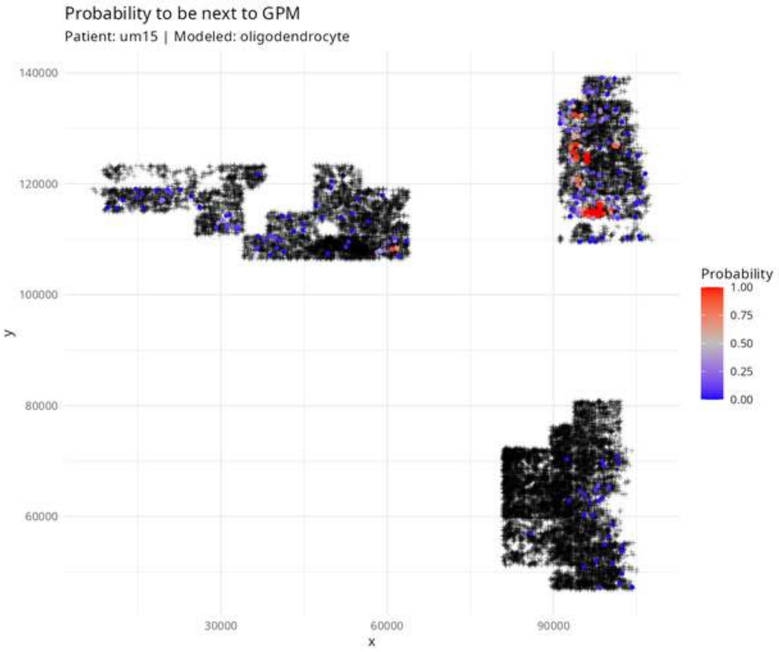

Probability to be next to GPM
Patient: um15 | Modeled: oligodendrocyte
140000
120000
100000
80000
60000
y
30000
60000
90000
x
Probability
1.00
0.75
0.50
0.25
0.00

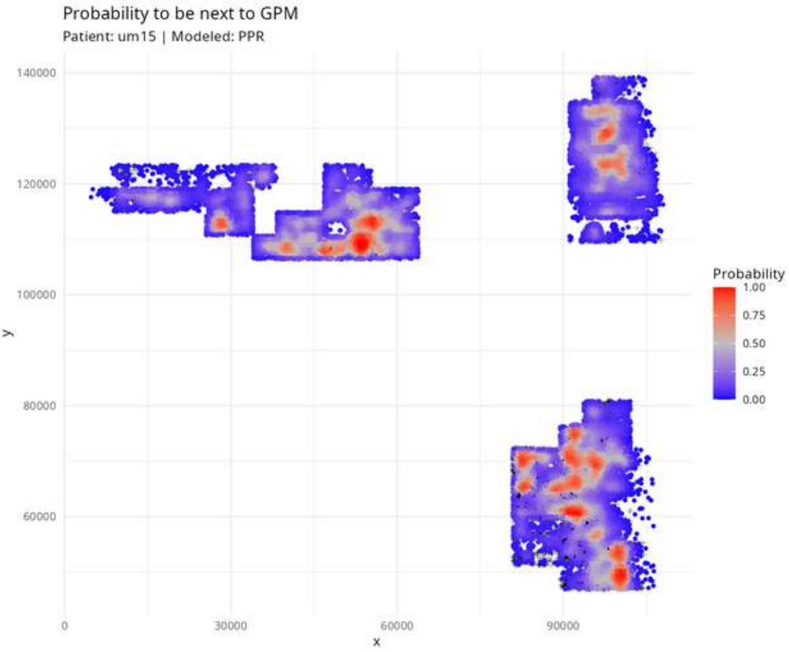
Probability to be next to GPM
Patient: um15 | Modeled: PPR
140000
120000
100000
80000
60000
y
0
30000
60000
90000
x
Probability
1.00
0.75
0.50
0.25
0.00

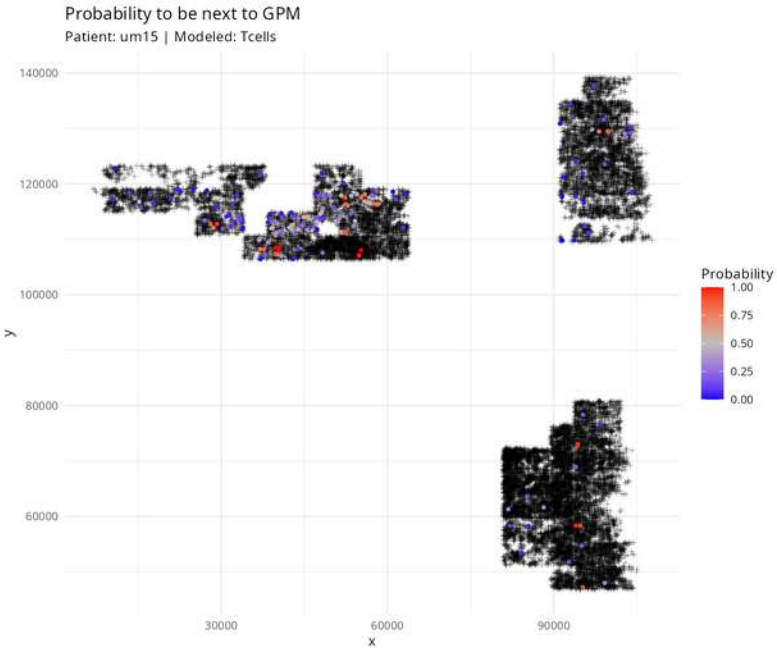
Probability to be next to GPM
Patient: um15 | Modeled: Tcells
140000
120000
100000
80000
60000
y
30000
60000
90000
x
Probability
1.00
0.75
0.50
0.25
0.00

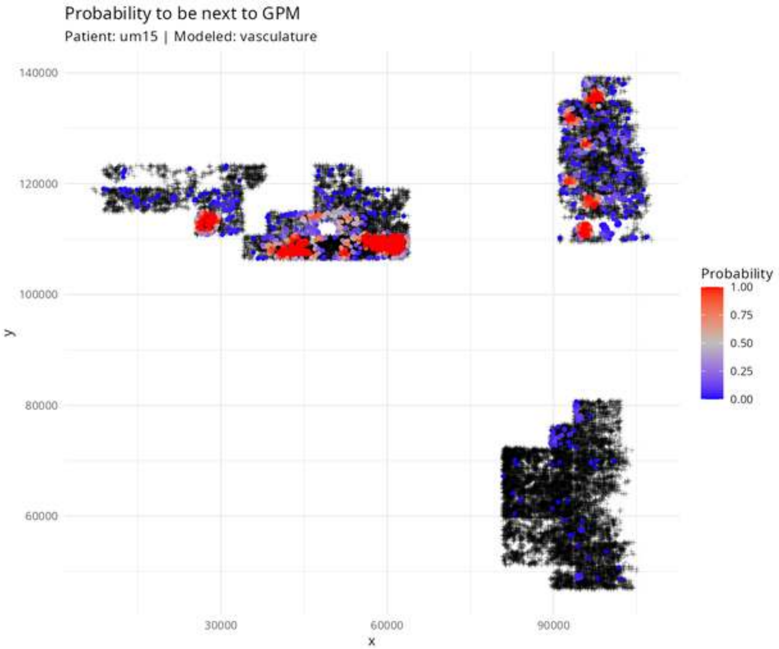
Probability to be next to GPM
Patient: um15 | Modeled: vasculature
140000
120000
100000
80000
60000
y
30000
60000
90000
x
Probability
1.00
0.75
0.50
0.25
0.00

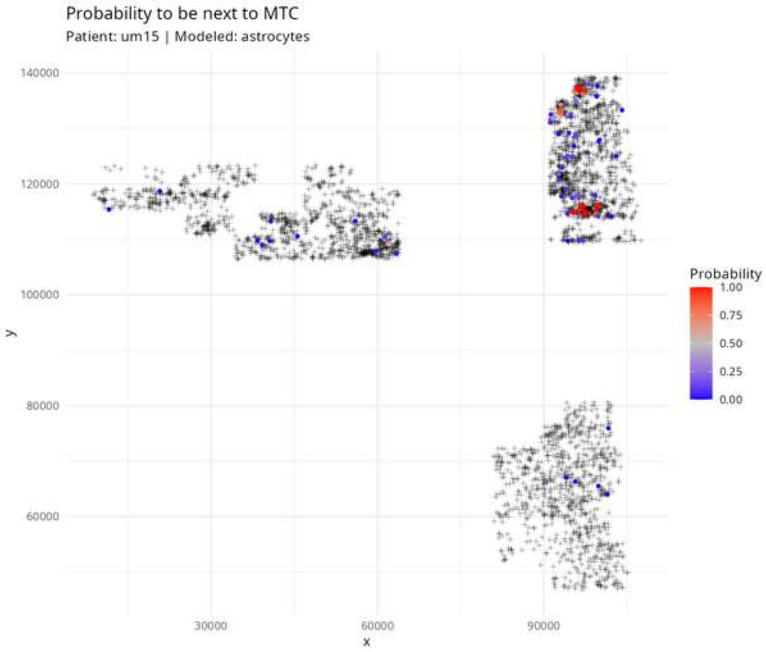

Probability to be next to MTC
Patient: um15 | Modeled: astrocytes
140000
120000
100000
80000
60000
y
30000
60000
90000
x
Probability
1.00
0.75
0.50
0.25
0.00

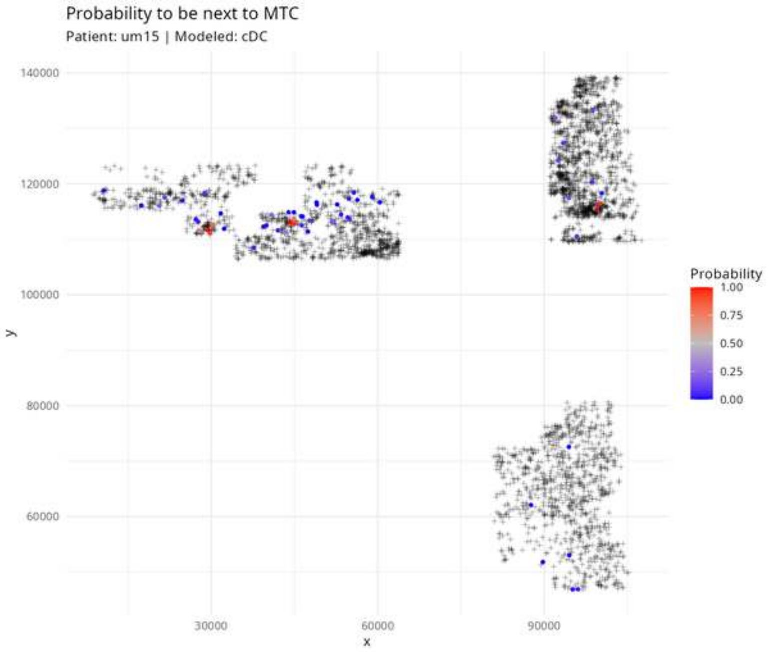
Probability to be next to MTC
Patient: um15 | Modeled: cDC
140000
120000
100000
80000
60000
y
30000
60000
90000
x
Probability
1.00
0.75
0.50
0.25
0.00

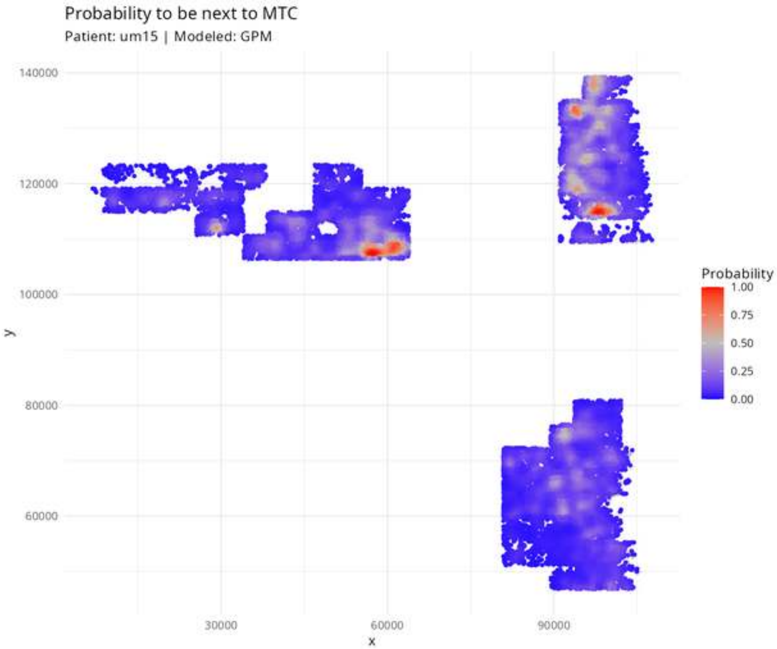

Probability to be next to MTC
Patient: um15 | Modeled: GPM
140000
120000
100000
80000
60000
y
30000
60000
90000
x
Probability
1.00
0.75
0.50
0.25
0.00

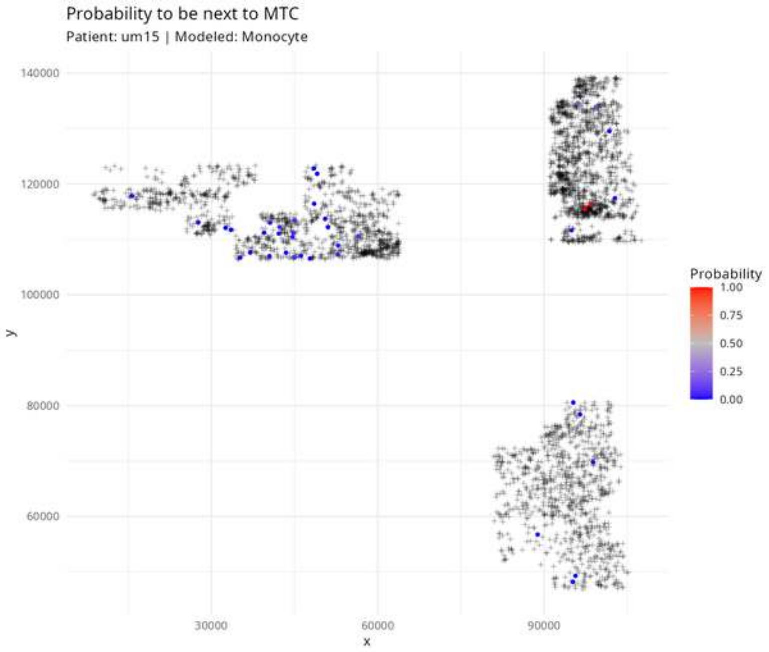
Probability to be next to MTC
Patient: um15 | Modeled: Monocyte
140000
120000
100000
80000
60000
y
30000
60000
90000
x
Probability
1.00
0.75
0.50
0.25
0.00

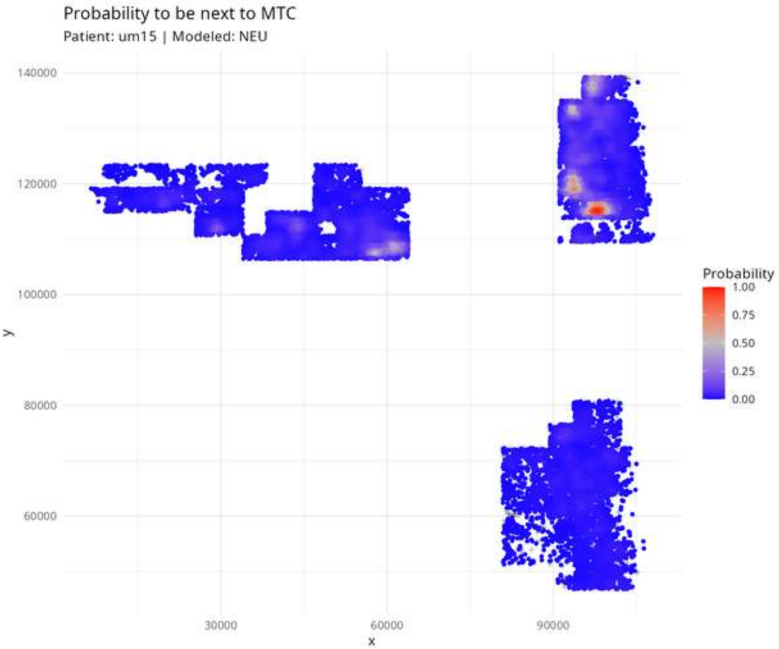

Probability to be next to MTC
Patient: um15 | Modeled: NEU
140000
120000
100000
80000
60000
y
30000
60000
90000
x
Probability
1.00
0.75
0.50
0.25
0.00

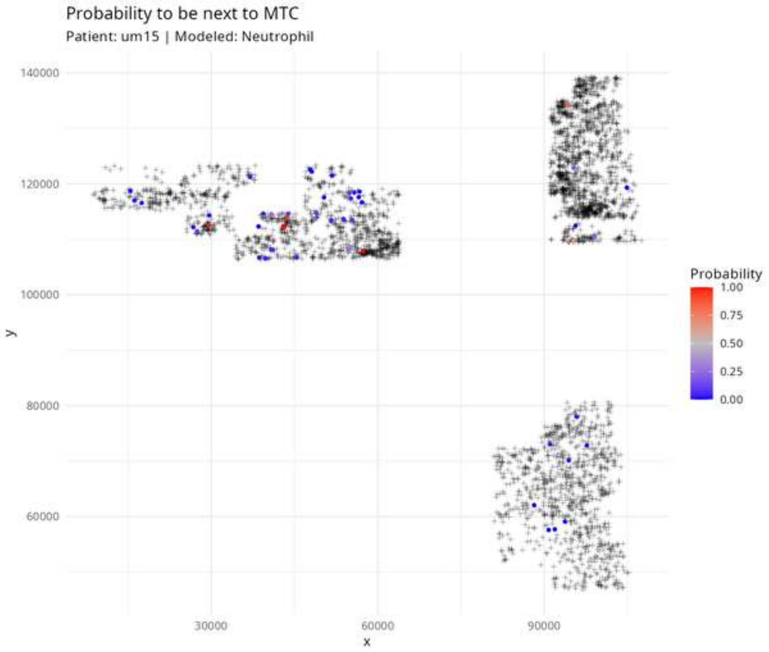

Probability to be next to MTC
Patient: um15 | Modeled: Neutrophil
140000
120000
100000
80000
60000
y
30000
60000
90000
x
Probability
1.00
0.75
0.50
0.25
0.00

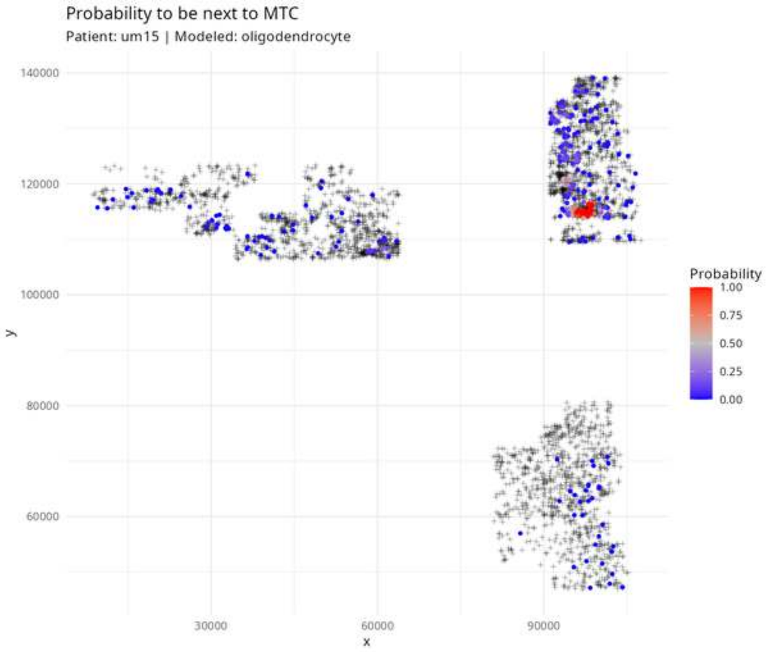
Probability to be next to MTC
Patient: um15 | Modeled: oligodendrocyte
140000
120000
100000
80000
60000
y
30000
60000
90000
x
Probability
1.00
0.75
0.50
0.25
0.00

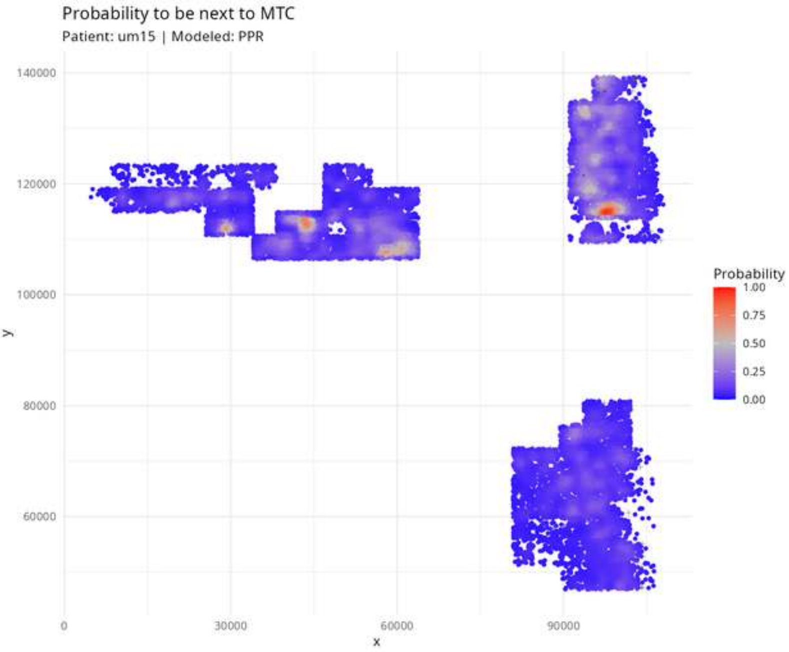
Probability to be next to MTC
Patient: um15 | Modeled: PPR
140000
120000
100000
80000
60000
y
0
30000
60000
90000
x
Probability
1.00
0.75
0.50
0.25
0.00

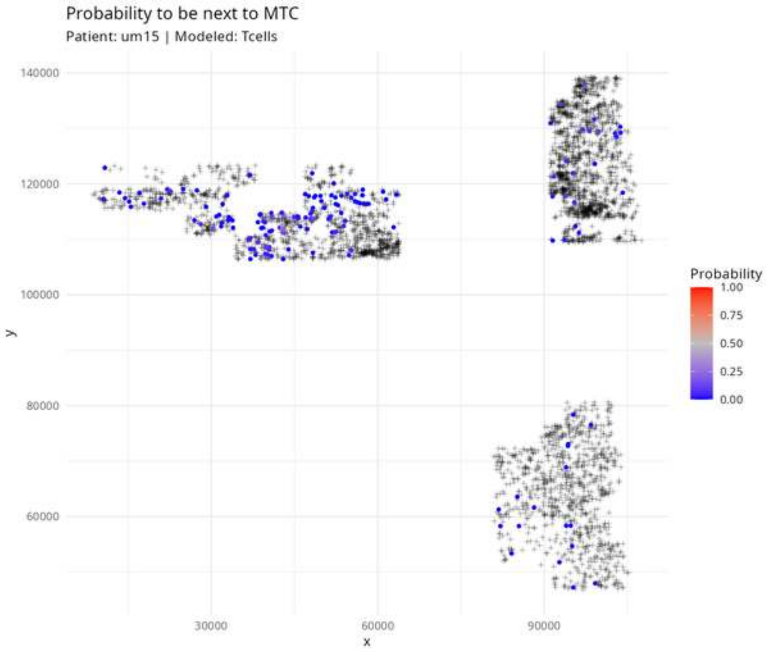

Probability to be next to MTC
Patient: um15 | Modeled: Tcells
140000
120000
100000
80000
60000
y
30000
60000
90000
x
Probability
1.00
0.75
0.50
0.25
0.00

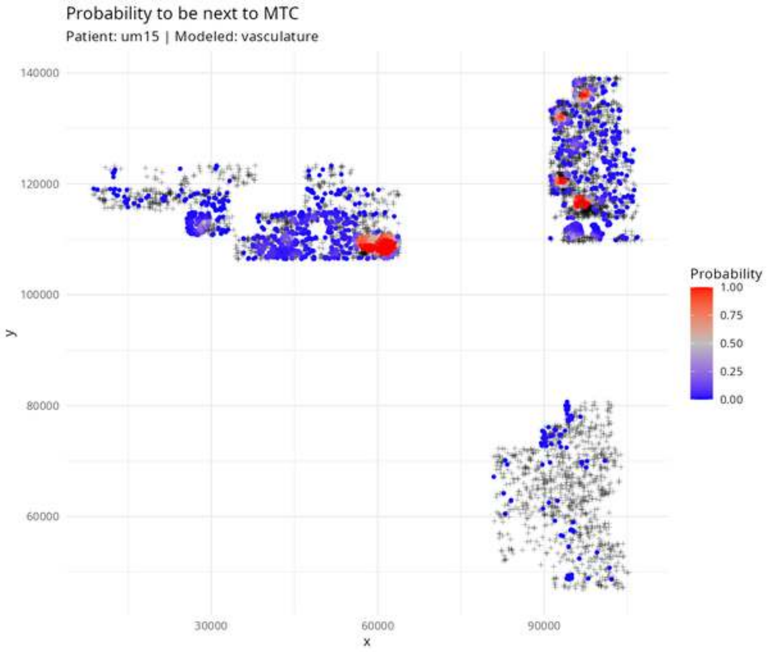
Probability to be next to MTC
Patient: um15 | Modeled: vasculature
140000
120000
100000
80000
60000
y
30000
60000
90000
x
Probability
1.00
0.75
0.50
0.25
0.00

Probability to be next to NEU

Patient: um15 | Modeled: astrocytes

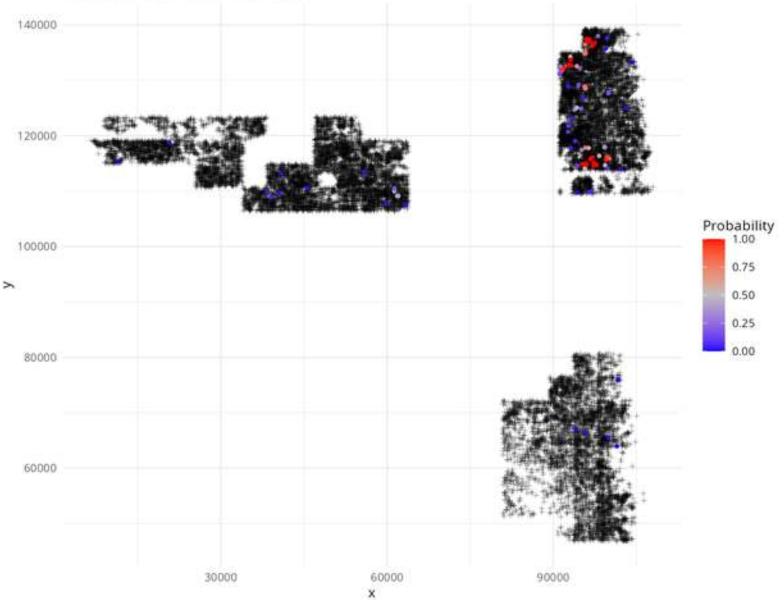

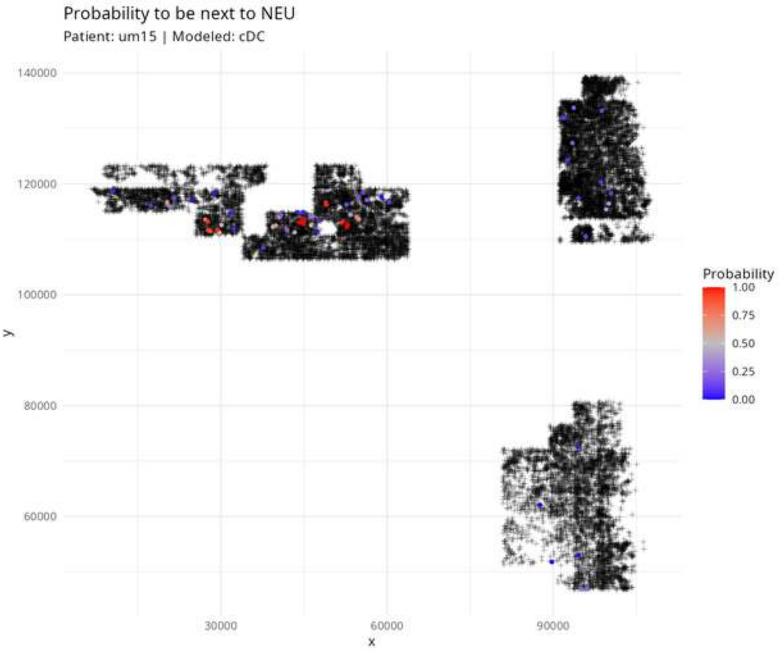

Probability to be next to NEU
Patient: um15 | Modeled: cDC
140000
120000
100000
80000
60000
y
30000
60000
90000
x
Probability
1.00
0.75
0.50
0.25
0.00

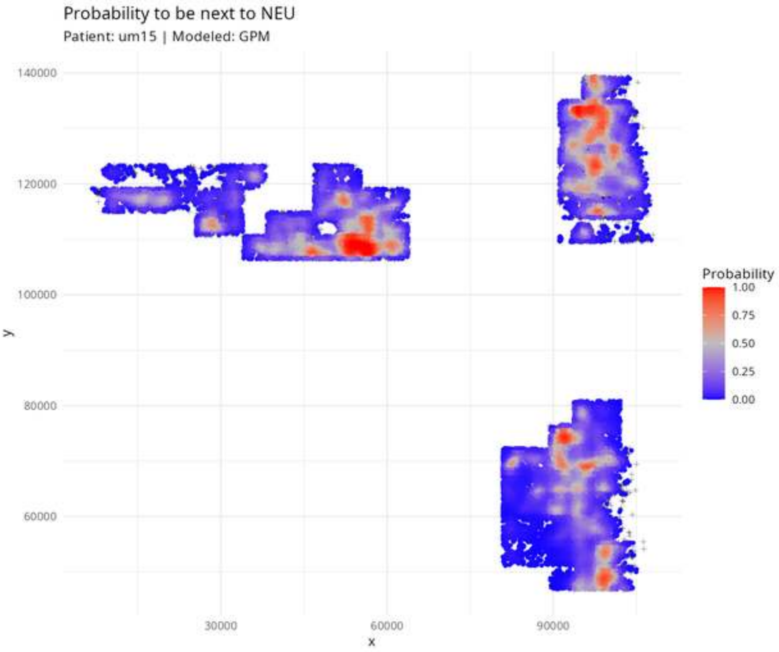
Probability to be next to NEU
Patient: um15 | Modeled: GPM
140000
120000
100000
80000
60000
y
30000
60000
90000
x
Probability
1.00
0.75
0.50
0.25
0.00

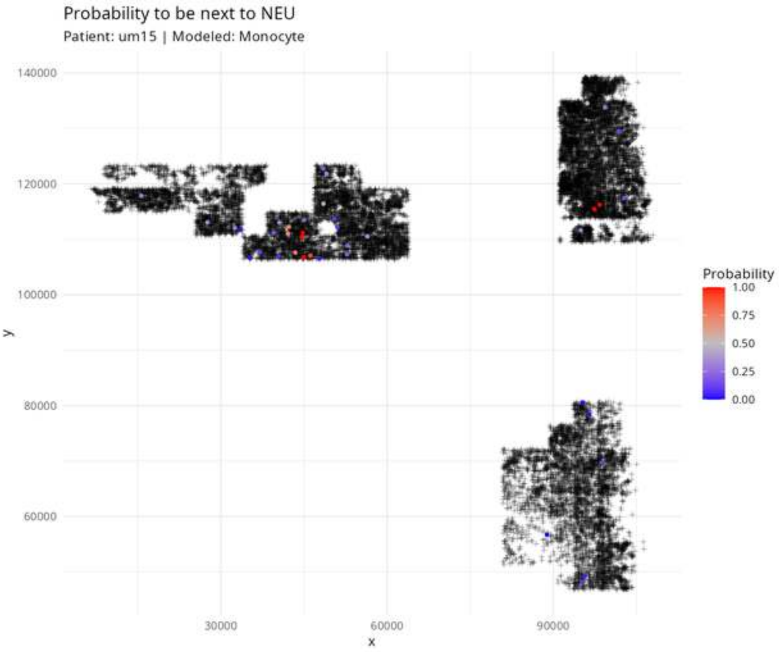
Probability to be next to NEU
Patient: um15 | Modeled: Monocyte
140000
120000
100000
80000
60000
y
30000
60000
90000
x
Probability
1.00
0.75
0.50
0.25
0.00

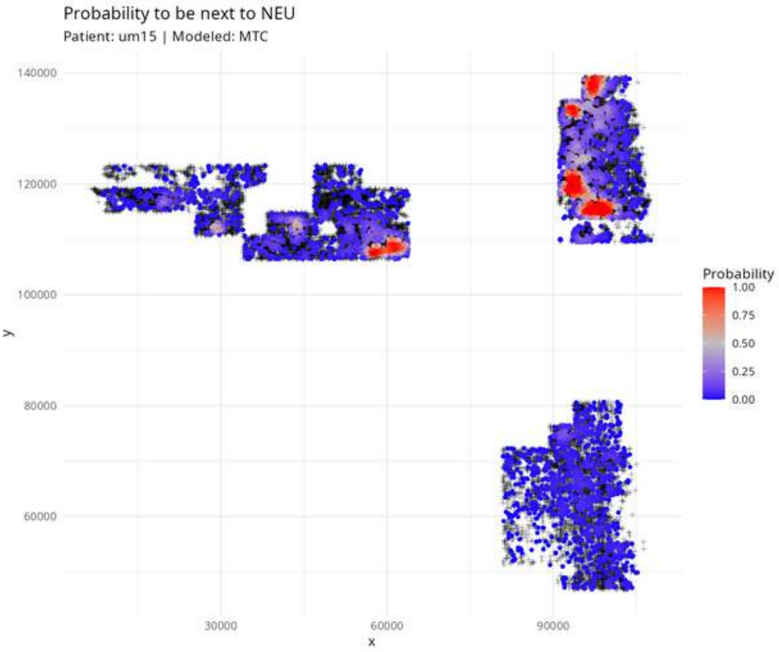

Probability to be next to NEU
Patient: um15 | Modeled: MTC
140000
120000
100000
80000
60000
y
30000
60000
90000
x
Probability
1.00
0.75
0.50
0.25
0.00

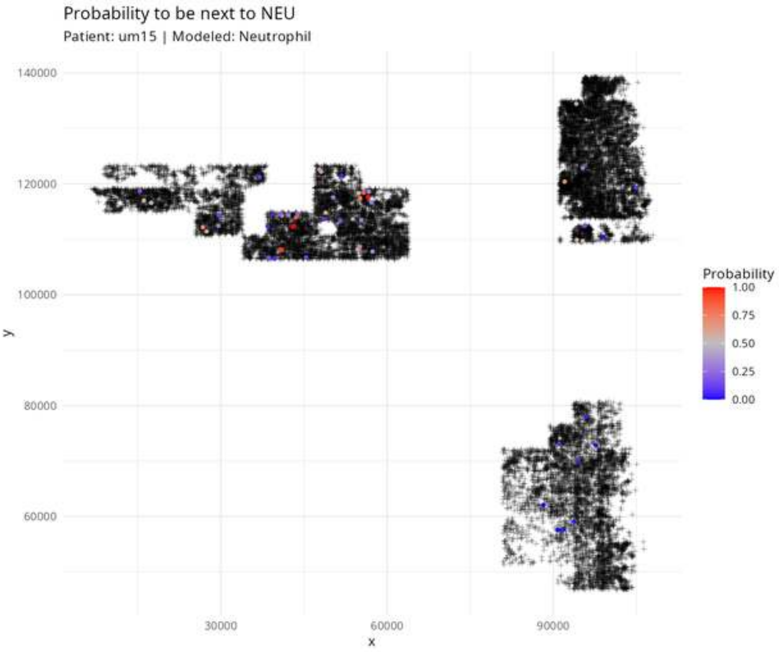

Probability to be next to NEU
Patient: um15 | Modeled: Neutrophil
140000
120000
100000
80000
60000
y
30000
60000
90000
x
Probability
1.00
0.75
0.50
0.25
0.00

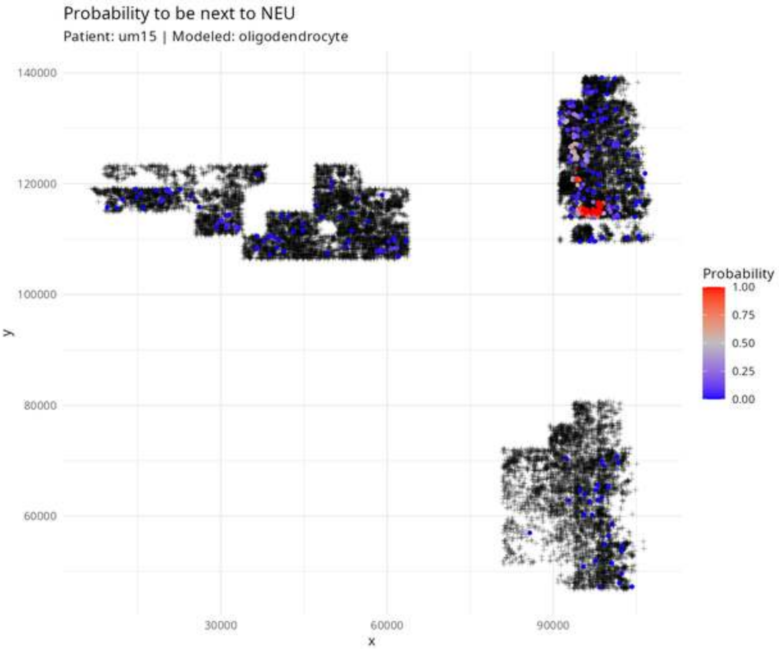

Probability to be next to NEU
Patient: um15 | Modeled: oligodendrocyte
140000
120000
100000
80000
60000
y
30000
60000
90000
x
Probability
1.00
0.75
0.50
0.25
0.00

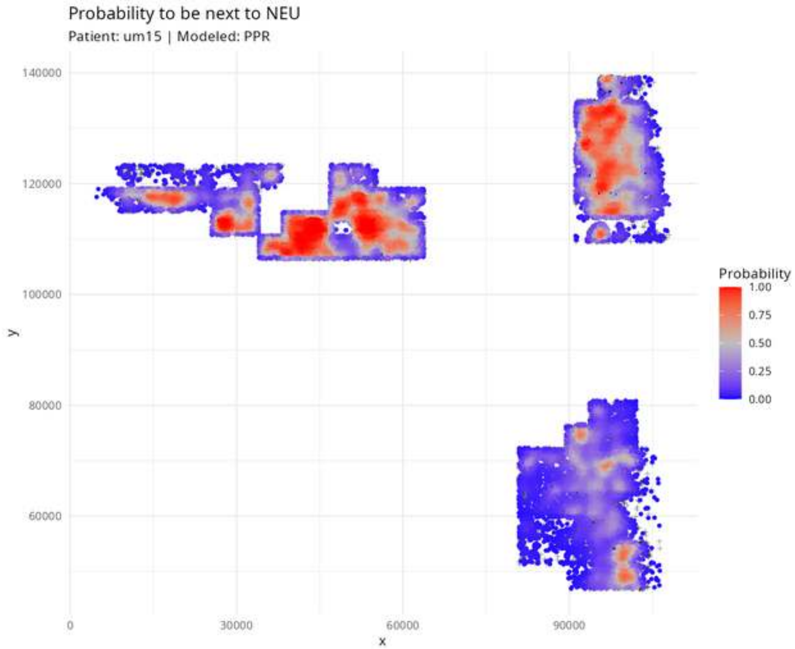
Probability to be next to NEU
Patient: um15 | Modeled: PPR
140000
120000
100000
80000
60000
y
0
30000
60000
90000
x
Probability
1.00
0.75
0.50
0.25
0.00

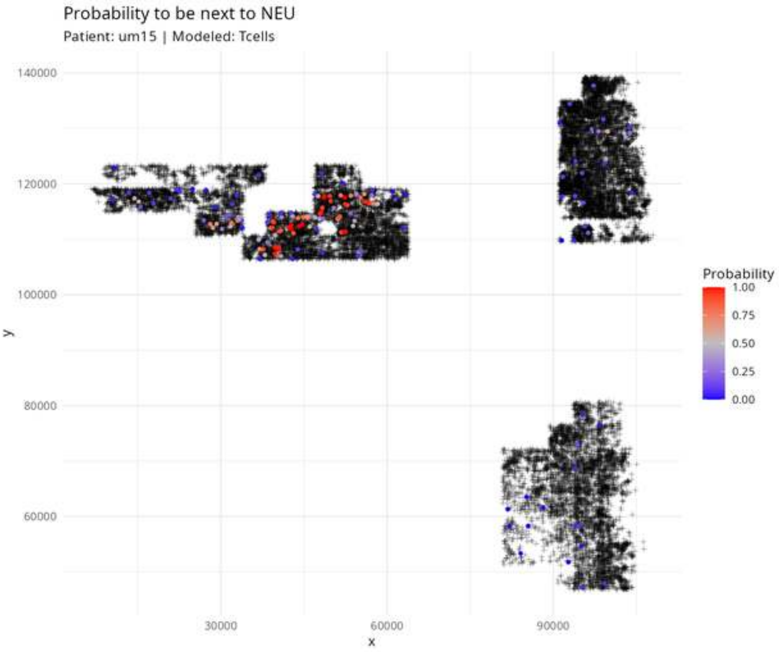
Probability to be next to NEU
Patient: um15 | Modeled: Tcells
140000
120000
100000
80000
60000
y
30000
60000
90000
x
Probability
1.00
0.75
0.50
0.25
0.00

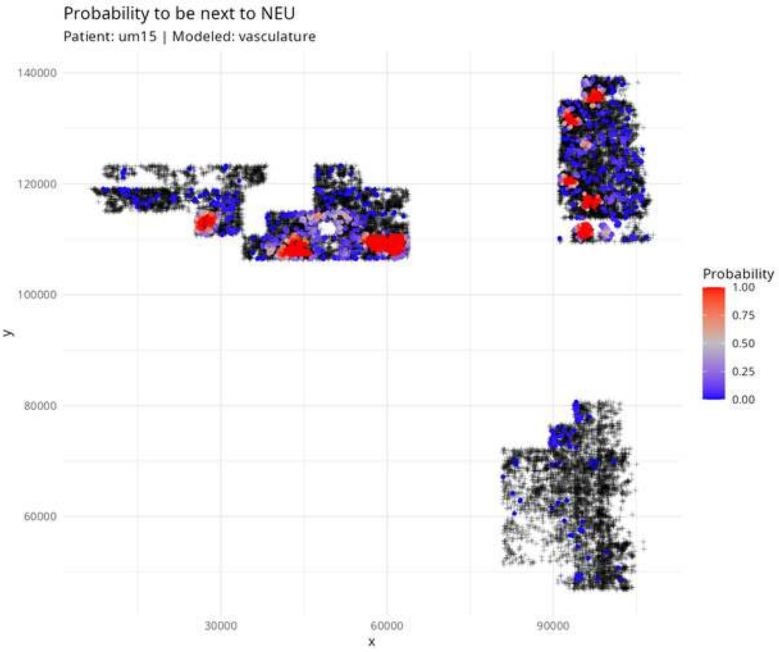

Probability to be next to NEU
Patient: um15 | Modeled: vasculature
140000
120000
100000
80000
60000
y
30000
60000
90000
x
Probability
1.00
0.75
0.50
0.25
0.00

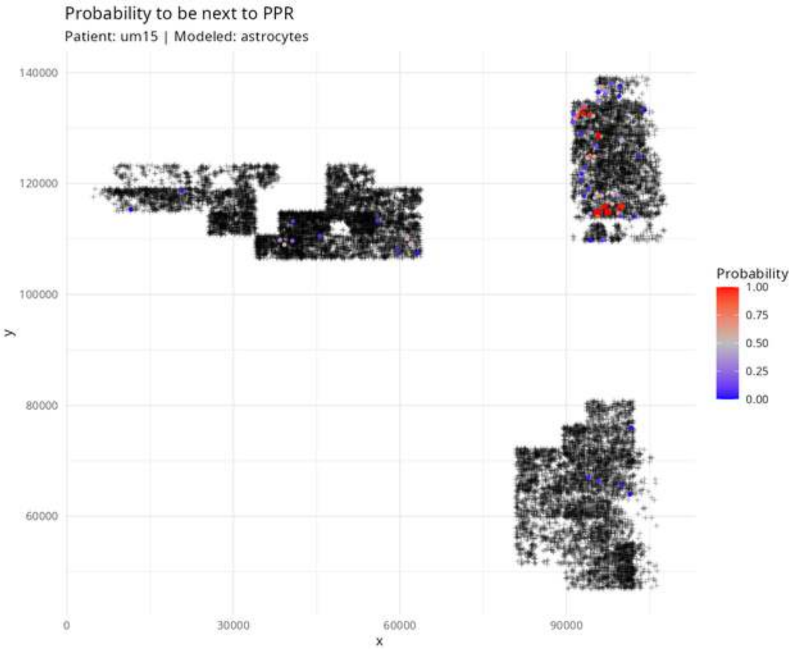
Probability to be next to PPR
Patient: um15 | Modeled: astrocytes
140000
120000
100000
80000
60000
y
0
30000
60000
90000
x
Probability
1.00
0.75
0.50
0.25
0.00

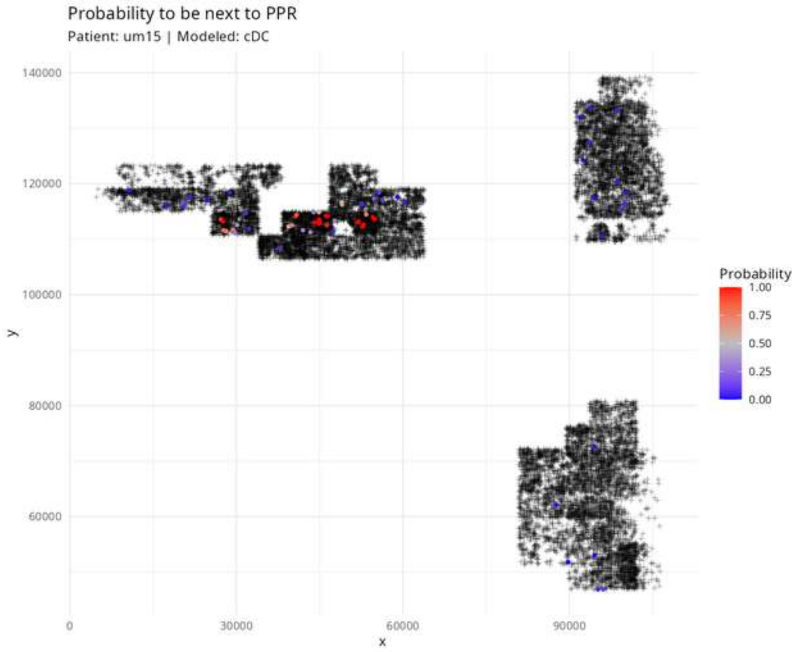

Probability to be next to PPR
Patient: um15 | Modeled: cDC
y
x
140000
120000
100000
80000
60000
0
30000
60000
90000
Probability
1.00
0.75
0.50
0.25
0.00

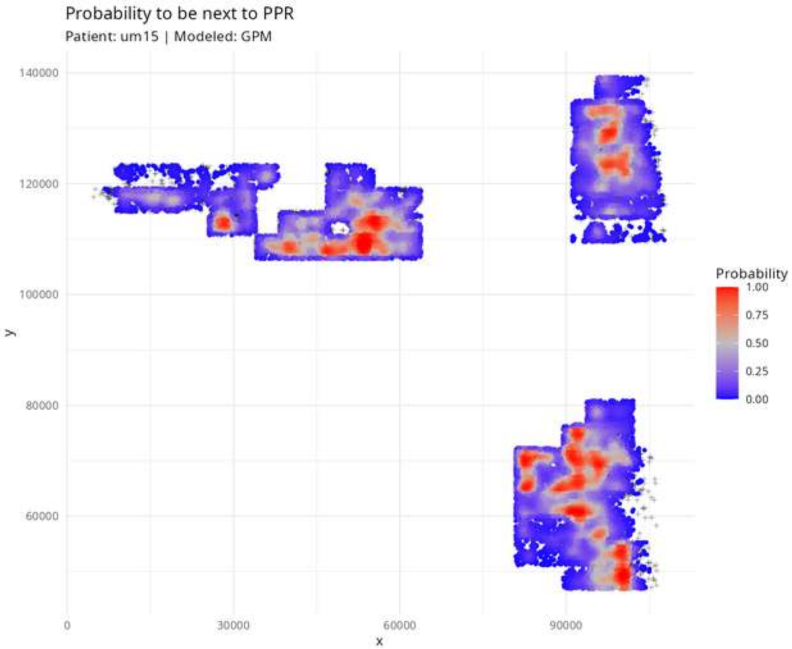
Probability to be next to PPR
Patient: um15 | Modeled: GPM
140000
120000
100000
80000
60000
y
0
30000
60000
90000
x
Probability
1.00
0.75
0.50
0.25
0.00

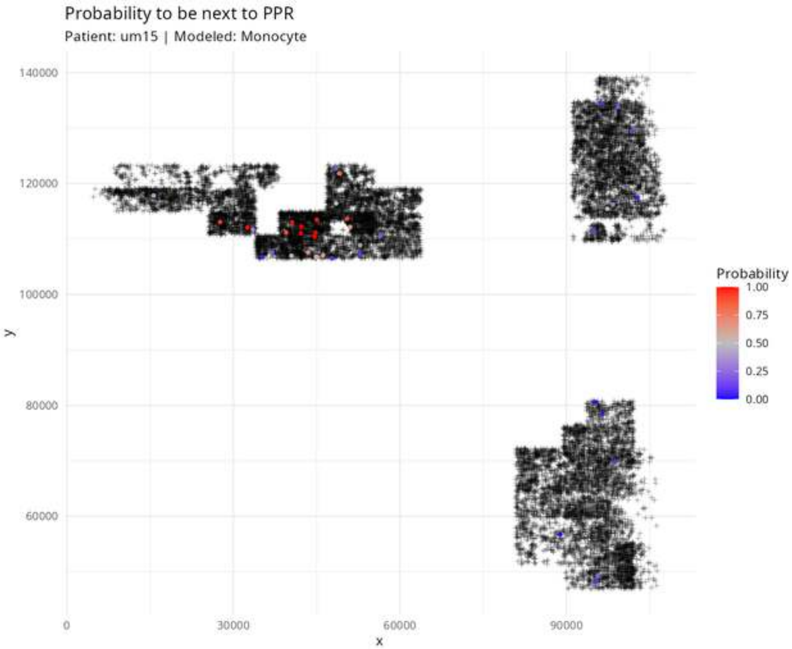

Probability to be next to PPR
Patient: um15 | Modeled: Monocyte
140000
120000
100000
80000
60000
y
0
30000
60000
90000
x
Probability
1.00
0.75
0.50
0.25
0.00

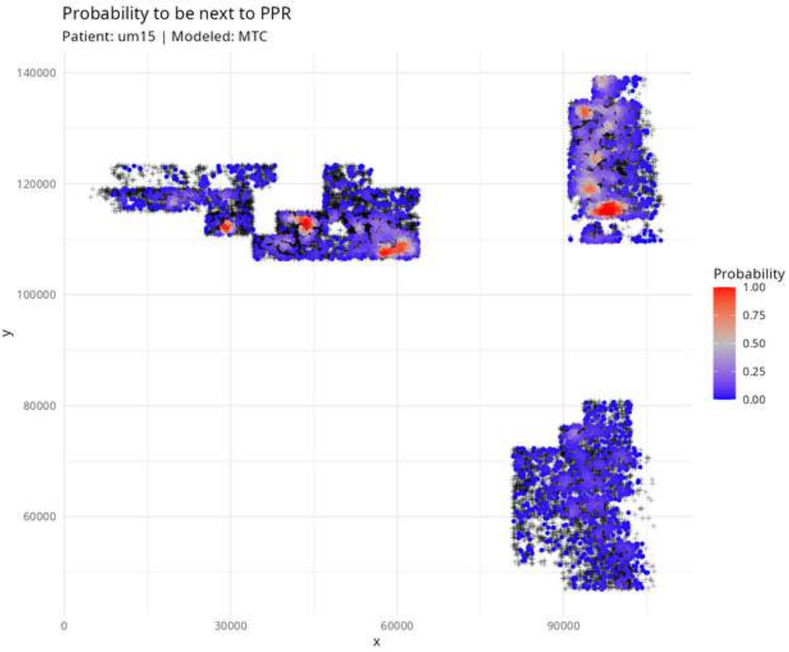

Probability to be next to PPR
Patient: um15 | Modeled: MTC
140000
120000
100000
80000
60000
y
0
30000
60000
90000
x
Probability
1.00
0.75
0.50
0.25
0.00

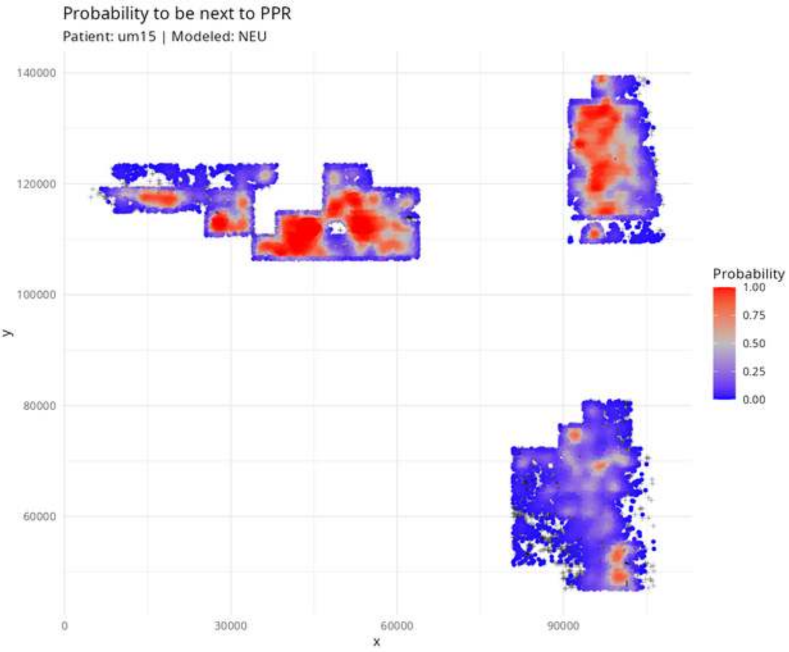

Probability to be next to PPR
Patient: um15 | Modeled: NEU
140000
120000
100000
80000
60000
y
0
30000
60000
90000
x
Probability
1.00
0.75
0.50
0.25
0.00

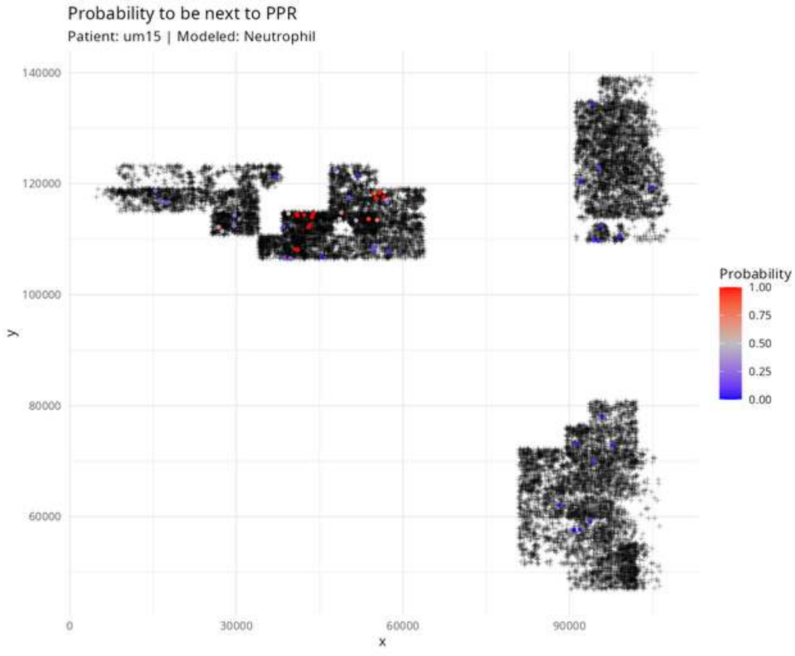

Probability to be next to PPR
Patient: um15 | Modeled: Neutrophil
140000
120000
100000
80000
60000
y
0
30000
60000
90000
x
Probability
1.00
0.75
0.50
0.25
0.00

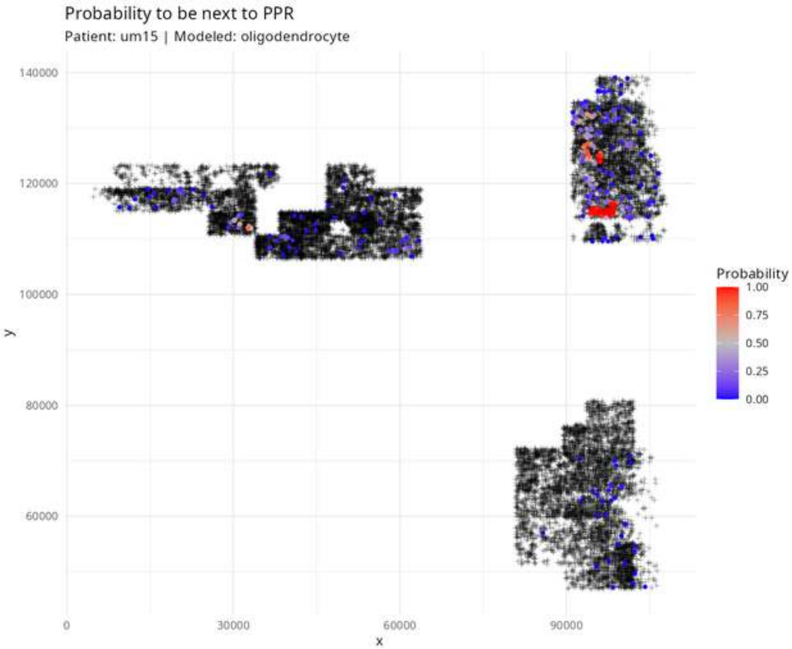
Probability to be next to PPR
Patient: um15 | Modeled: oligodendrocyte
140000
120000
100000
80000
60000
y
0
30000
60000
90000
x
Probability
1.00
0.75
0.50
0.25
0.00

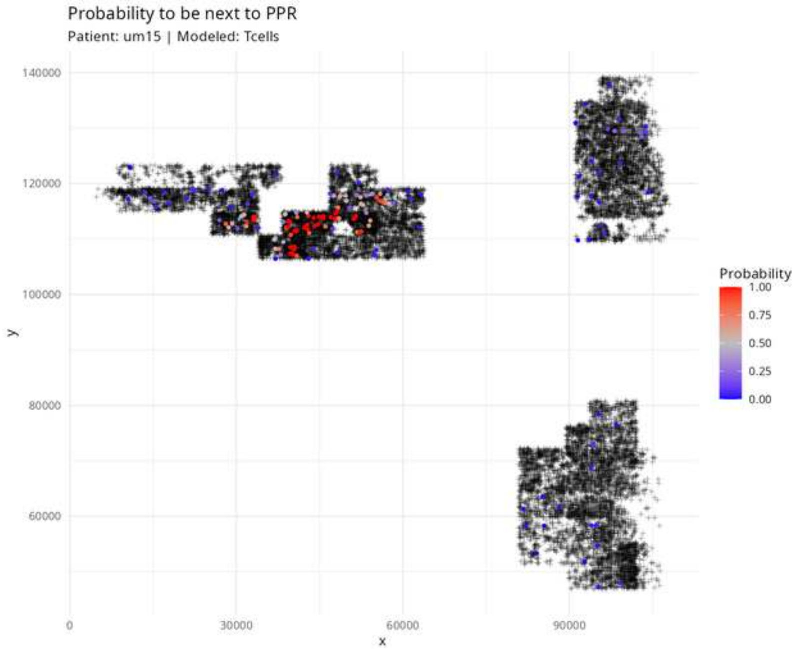
Probability to be next to PPR
Patient: um15 | Modeled: Tcells
140000
120000
100000
80000
60000
y
0
30000
60000
90000
x
Probability
1.00
0.75
0.50
0.25
0.00

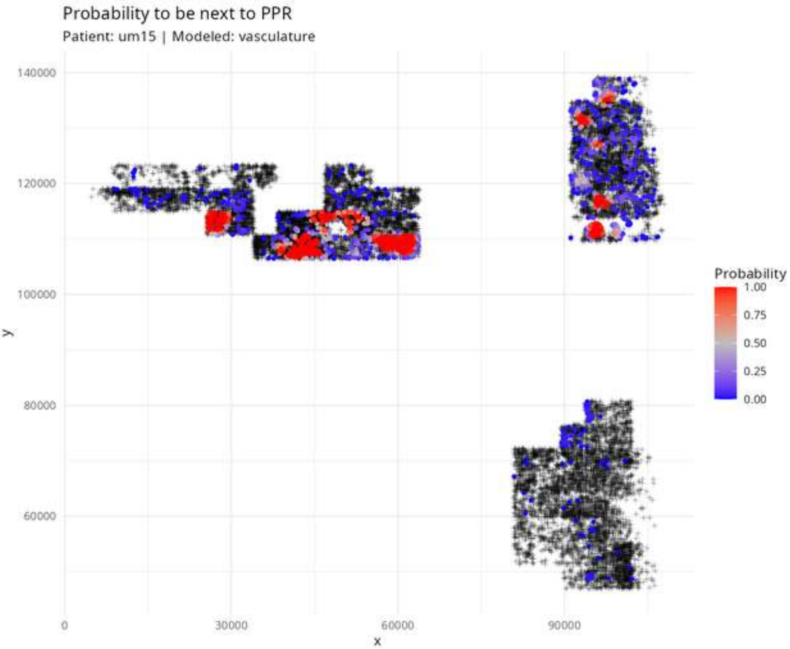

Probability to be next to PPR
Patient: um15 | Modeled: vasculature
140000
120000
100000
80000
60000
y
0
30000
60000
90000
x
Probability
1.00
0.75
0.50
0.25
0.00
