## Supplementary Figures 6 for "Point-Process Modelling of Cell-Type Interaction Structure in CosMx Glioma Data for Hypothesis-Driven Tumor Inference from scRNA-seq"

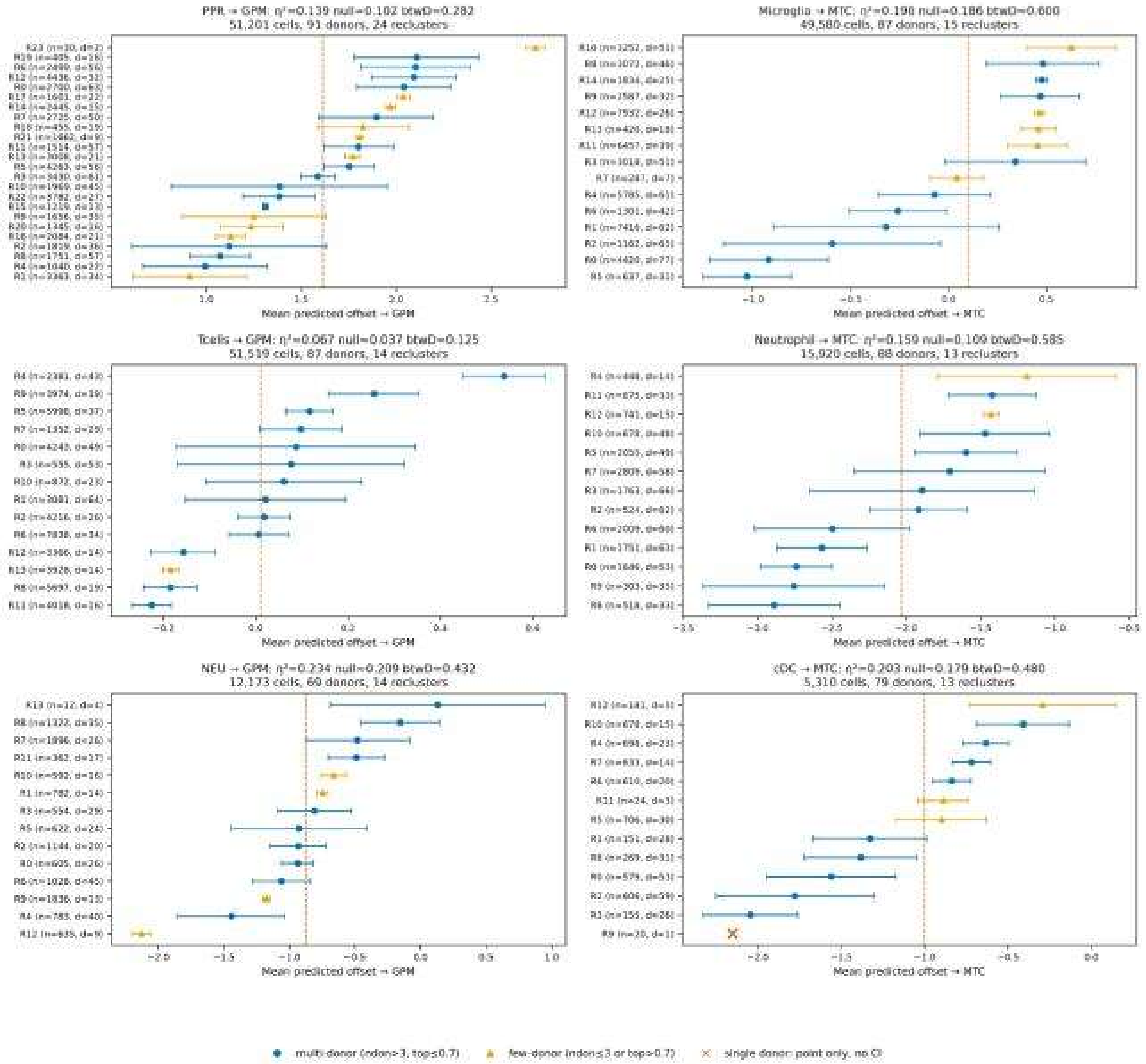
PPR → GPM: η²=0.139 null=0.102 btwD=0.282
51,201 cells, 91 donors, 24 reclusters
Mean predicted offset → GPM
Microglia → MTC: η²=0.198 null=0.186 btwD=0.600
49,580 cells, 87 donors, 15 reclusters
Mean predicted offset → MTC
Tcells → GPM: η²=0.067 null=0.037 btwD=0.125
51,519 cells, 87 donors, 14 reclusters
Mean predicted offset → GPM
Neutrophil → MTC: η²=0.159 null=0.109 btwD=0.585
15,920 cells, 88 donors, 13 reclusters
Mean predicted offset → MTC
NEU → GPM: η²=0.234 null=0.209 btwD=0.432
12,173 cells, 69 donors, 14 reclusters
Mean predicted offset → GPM
cDC → MTC: η²=0.203 null=0.179 btwD=0.480
5,310 cells, 79 donors, 13 reclusters
Mean predicted offset → MTC
multi-donor (ndon>3, top≤0.7)
few-donor (ndon≤3 or top>0.7)
single donor: point only, no CI

Figure S1 — Subcluster means (supplement): types with no state signal beyond donor structure
Points: per-recluster mean; bars: ±1.96×donor-clustered SE; dashed red: type mean. UMAP overlays retained in E035 for QC only and carry no inferential weight.

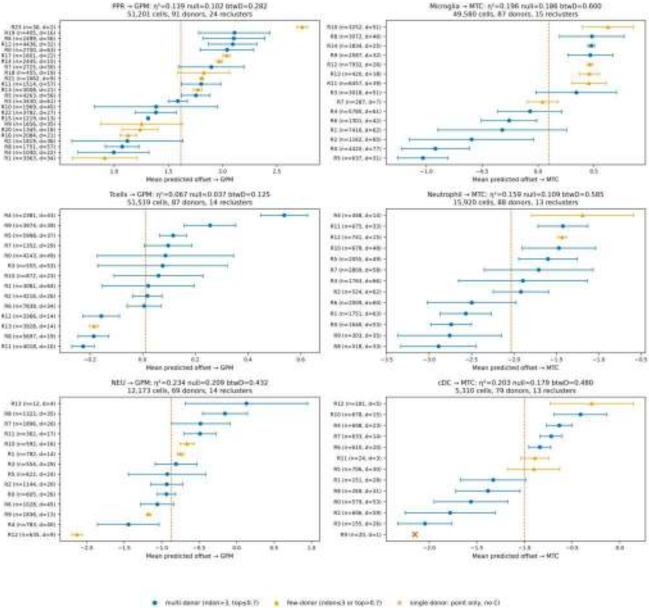
